# Calling small variants with universality and Bayesian-frequentist hybridism

**DOI:** 10.1101/2020.08.23.263749

**Authors:** Xiaofei Zhao, Allison Hu, Sizhen Wang, Xiaoyue Wang

## Abstract

The accuracy of variant calling is crucially important in clinical settings, as the misdiagnosis of a genetic disease such as cancer can compromise patient survival. Although many variant callers were developed, variant-calling accuracy is still insufficient for clinical applications.

Here we describe UVC, a method for calling small variants of germline or somatic origin. By combining contrary assumptions with sublation, we found two principles to improve variant calling. First, we discovered the following power-law universality: allele fraction is inversely proportional to the cubic root of variant-calling error rate. Second, we found that zero inflation can combine Bayesian and frequentist models of sequencing bias.

We evaluated UVC with other state-of-the-art variant callers by considering a variety of calling modes (germline, somatic, tumor-only, and cell-free DNA with unique molecular identifiers (UMIs)), sequencing platforms (Illumina, BGI, and IonTorrent), sequencing types (whole-genome, whole-exome, and PCR-amplicon), human reference genomes (hg19, hs37d5, and GRCh38), aligners (BWA and NovoAlign), and representative sequencing depths and purities for both tumor and normal. UVC generally outperformed other germline variant callers on the GIAB germline truth sets. UVC strongly outperformed other somatic variant callers on 192 scenarios of *in silico* mixtures simulating 192 combinations of tumor/normal sequencing depths and tumor/normal purities. UVC strongly outperformed other somatic variant callers on the GIAB somatic truth sets derived from physical mixture and on the SEQC2 somatic reference sets derived from the breast-cancer cell-line HCC1395. UVC achieved 100% concordance with the manual review conducted by multiple independent researchers on a Qiagen 71-gene-panel dataset derived from 16 patients with colon adenoma. Additionally, UVC outperformed Mageri and smCounter2, the state-of-the-art UMI-aware variant callers, on the tumor-only datasets used for publishing these two variant callers. Performance is measured by using sensitivity-specificity trade off for all called variants. The improved variant calls generated by UVC from previously published UMI-based sequencing data are able to provide additional biological insight about DNA damage repair.

UVC enables highly accurate calling of small variants from a variety of sequencing data, which can directly benefit patients in clinical settings. UVC is open-sourced under the BSD 3-Clause license at https://github.com/genetronhealth/uvc and quay.io/genetronhealth/gcc-6-3-0-uvc-0-6-0-441a694.

## 1 Background

Variant calling is a fundamental problem. Accurate detection of germline variants is crucial to the evaluation of predisposition to many diseases, to the study of biological pathways, etc. Although a plethora of algorithms and software packages were designed and developed for calling germline variants, there are still opportunities to improve. Accurate detection of somatic variants is crucial to the diagnosis, prognosis, and treatment monitoring of cancer. Various next-generation sequencing (NGS)-based methods have been routinely used to discover variants at different allele fractions in different types of tumor samples. However, the accuracy of software that calls these variants is compromised by various biases and errors in NGS experiments. In a research setting, we are interested in drawing conclusions about populations of patients, so the consequences of technical issues with variant callers that affect occasional samples will be mitigated by sample size ^27^. In a clinical setting, we are interested in the mutations in each specific patient, so these issues become substantially more important ^27^. Variant refinement has the potential to filter out false positive variant calls but is unable to rescue the false negative variants that were not called ^13^. Moreover, manual variant refinement is both labor-intensive and time-consuming ^13^.

Moreover, some enhanced library-preparation techniques for NGS, such as molecular barcoding with unique molecular identifiers (UMIs) and duplex sequencing with duplex UMIs, were applied in clinical settings for detecting mutations from cell-free DNA (cfDNA). Currently, UMI-aware variant callers are strongly coupled with their associated experimental protocols, so the academic and clinical communities both need a generic UMI-aware variant caller that performs well on multiple independently generated UMI datasets.

Here we described and evaluated UVC, a versatile variant caller, which uses universality and Bayesian-frequentist hybrid zero-inflated modeling of biases. Universality is the observation that properties of a large class of systems are independent of the dynamic detail of the system. Here, universality refers to the following observation that we discovered: if the coverage depth of a variant is sufficiently high, then allele fraction is inversely proportional to the cubic root of variant-calling error probability regardless of variant type and error type. Bayesian and frequentist statistics were respectively developed by two schools of thought about the nature of statistics. And zero-inflation is often used to adjust a model by introducing a special mechanism for generating the statistical outcome represented by zero. Here, zero denotes the lack of a bias assumed by the frequentist null hypothesis, and nonzero denotes the effect size of the bias estimated by Bayesian inference.

UVC is able to call somatic single-nucleotide variants (SNVs) and insertions-deletions (InDels) without using any training data. The variant quality generated by UVC can reliably rank variants by confidence, where variant quality refers to the QUAL column in the VCF file format specification ^5^. Thus, we can simply apply different cutoffs on variant quality to filter variants for different applications, respectively. Furthermore, the variant statistics produced by UVC can be used as input features for both filtering variants (e.g., hard filter in VarScan2^14^ and MuTect ^3^) and recalling variants (e.g., ensemble mode in NeuSomatic ^20^). UVC is designed to be aware of both UMI (including duplex UMI) and the tumor-matched normal control. Nevertheless, if UMI and/or the tumor-matched normal control is not provided, UVC can still accurately call variants. Thus, UVC uses as much information as possible from sequencing data but does not require any specific information to be present in such data. Finally, UVC runs fast, making its usage possible in clinical settings.

## 2 Methods

This section, along with Fig. 1, summarizes our main method by presenting our ideas to improve the calling of small variants. More details can be found in the supplementary methods. We utilized the idea of combining, through sublation, a pair of contrary assumptions which negate each other to the maximum degree. For example, in the NGS settings, the statements “sequencing depth is low” and “sequencing depth is high” constitute a pair of contrary assumptions because “low” and “high” are the antonyms of each other. By applying this idea to NGS bioinformatics, we made the following two discoveries. First, we discovered that the probability of a variant candidate being false positive at high depth of coverage is inversely proportional to the cube (i.e., third power) of its allele fraction. Second, we discovered that a Bayesian model with an inflated degree of belief for the frequentist null hypothesis can model various biases in NGS.

**Figure 1:**
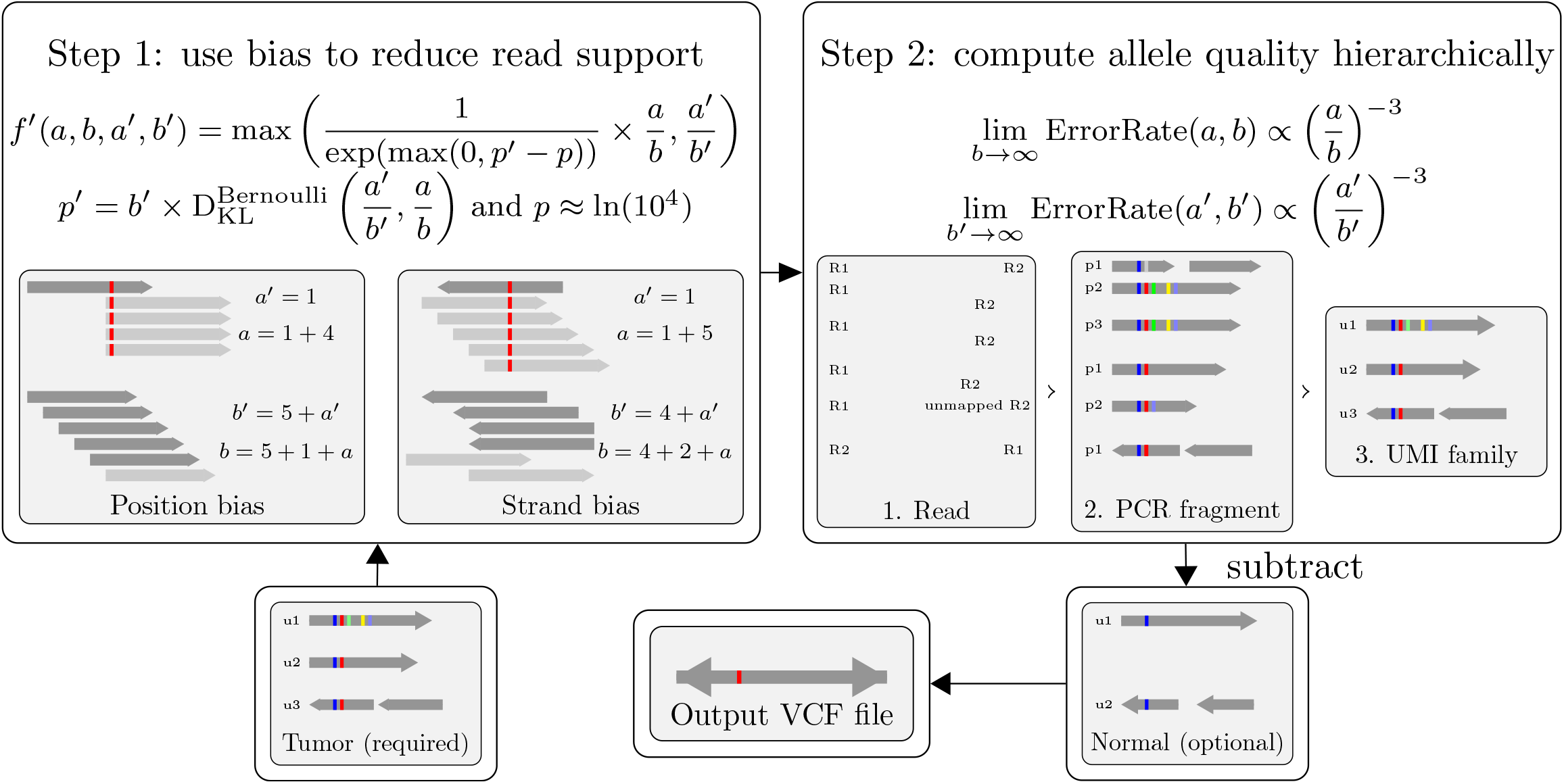
Overview of the variant-calling method in UVC.

First, we discovered the following universal power-law relationship (referred to as the NGS power law) between allele fraction and NGS error: given the expected and observed allele fractions *f* and *g* of a variant candidate, the probability that the candidate is false positive is approximately proportional to 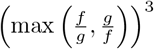 at high sequencing depth. Thus, the NGS power law translates the allele fraction of each variant candidate into an upper bound for the probability that the candidate is false positive. In addition, the NGS power law is inherently aware of the unique molecular identifiers (UMIs, which are also known as molecular barcodes) that are often used in the sequencing of cell-free DNA (cfDNA) in blood plasma. Thus, the NGS power law is universally applied by UVC for calling germline SNP, germline InDel, somatic SNV, and/or somatic InDel.

Second, we discovered that zero inflation can combine Bayesian and frequentist statistics to model NGS biases, where zero denotes the absence of a bias. Similar to traditional frequentist statistics, our model considers the following two hypotheses.

1. Null hypothesis: there is no bias in the variant candidate.
2. Alternative hypothesis: there is some bias in the variant candidate.

Our model computes a likelihood ratio of the alternative hypothesis to the null hypothesis. The likelihood ratio is similar to a *P* value. If the likelihood ratio exceeds a predefined threshold, then the null hypothesis is rejected, but the variant itself is not rejected. Instead, the probabilty *p*_1_ that the variant candidate is false positive increases in a way that is directly proportional to such likelihood. At the same time, our model uses only the reads not characterized by any bias to compute the probability *p*_2_ that the variant candidate is false positive. By default, the non-informative Jeffrey’s prior is applied to the computation of all allele fractions used for estimating the probability that the variant candidate is false positive. Then, the maximum of *p*_1_ (the frequentist probability) and *p*_2_ (the Bayesian probability) is used as the probability that the variant candidate is false positive. In brief, our inference formalizes the following two rules of thumb with the manual review of variant candidates:

1. The frequentist rule: if the sequencing depth is low, then we estimate the likelihood that the variant candidate has some bias. If the likelihood exceeds some threshold, then we increase the probability that the variant candidate is false positive accordingly.
2. The Bayesian rule: if the sequencing depth is sufficiently high, then we assume that there is some bias and compute the effect size of the bias.

The frequentist and Bayesian inference rules are incorporated into UVC to achieve high sensitivity at low depth of coverage and robustness to any systematic bias at high depth of coverage, respectively. These two discoveries are summarized in Fig. 1 which also provides an overview of our variant-calling method. In sum, for each variant candidate, UVC sequentially performs the following tasks.

1. UVC uses bias, such as position bias and strand bias, to reduce the number of reads effectively supporting the variant candidate.
2. UVC groups reads into fragments and then groups fragments into unique molecular identifier (UMI) families.
3. UVC uses the bias-reduced reads, fragments, and UMI families to compute their respective variant allele fractions. Then, UVC uses these allele fractions to compute the probability that the variant candidate is false positive.
4. UVC compares the tumor with its matched normal to refine the variant candidate if the matched normal is provided as input.

## 3 Results

We evaluated the performance of UVC using the metrics F-score and PrAUC. F-score denotes the maximum harmonic mean between precision and recall. PrAUC, which is equivalent to average precision, denotes the area under the curve (AUC) of precision-versus-recall.

### 3.1 Evaluation results on whole-genome-sequencing (WGS) and amplicon-sequencing data to validate the power-law universality

First, we validated the NGS power law with two datasets: a whole-genome-sequencing (WGS) dataset of the HG001 (or equivalently NA12878) cell-line sequenced at 300x average depth by Illumina HiSeq, and an amplicon-sequencing dataset of a cell-line mixture consisting of 1% HG001 and 99% HG002 (or equivalently NA24385) sequenced by Illumina NextSeq. Indeed, the probability of a variant being false positive is inversely proportional to the cubic power of the allele fraction of the variant if the range of allele fractions is between 0.1% and 100%, and this range spans more than the two orders of magnitude mentioned by Stumpf and Porter ^25^ (Figures S1 and S2). Moreover, this probability is always close to 1 at the allele fraction of 0.1% computed using raw sequenced segments regardless of sequencing technology and assay type, which is consistent with an NGS nucleotide substitution error rate of approximately 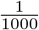 per sequenced base ^23^ (Figures S1 and S2). Thus, the range of allele fractions between 0.1% and 100% is exhaustive if such fractions are computed using raw sequenced segments.

### 3.2 Evaluation results on WGS datasets for calling germline variants

We evaluated UVC with the germline variant callers HaplotypeCaller ^18^, Strelka2^22^, FreeBayes ^11^, and bcftools ^5;16^ with a combination of two performance metrics (F-score and PrAUC), two types of small variants (SNV and InDel), two sequencing platforms (Illumina HiSeq with NovoAlign and BGI MGISEQ with BWA MEM ^17^), three Genome-in-a-Bottle (GIAB) reference samples (HG001, HG002, and HG005) ^30^, and two average sequencing depths (30x and 60x). Our evaluation shows that UVC is the best for calling germline SNPs and the second best for calling germline InDels, whereas HaplotypeCaller is the best for calling germline InDels (Figures S3-S8). We manually reviewed the germline InDel calls and found that UVC sometimes cannot determine if an InDel is homyzygous or heterozygous, whereas HaplotypeCaller can, presumably because of the strategy of localized assembly employed by HaplotypeCaller.

### 3.3 Evaluation results on WGS datasets for calling variants in tumor-only mode, with 64 *in silico* mixtures considering both Illumina and BGI platforms

We evaluated UVC with the variant callers Mutect2^1^ and LoFreq ^28^, all in tumor-only mode, with a combination of two performance metrics (F-score and PrAUC), two sequencing platforms and aligners (Illumina HiSeq with NovoAlign and BGI MGISEQ with BWA MEM ^17^), four tumor sequencing depths (HiSeq: 240x and 120x. MGISEQ: 48x and 24x), eight tumor purities (1.0, 0.75, 0.5, 0.25, 0.125, 0.0625, 0.03125, and 0.015625), and two simulated sources of tumor and normal cells (HG001/HG002 as tumor/normal and HG002/HG001 as tumor/normal). The *in silico* mixture was prepared using the technique proposed by Cibulskis et al. ^3^. In our evaluation, the germline-versus-somatic origin is not determined by any caller. Thus, the groundtruth germline variants in the normal were first subtracted from the called variants, then the remaining called variants were assessed for sensitivity-specificity trade offs. Our evaluation shows that UVC is always the best for calling somatic SNVs and InDels in tumor-only mode without any exception (Figures S9-S40).

### 3.4 Evaluation results on WGS datasets for calling somatic variants from tumor-normal pairs, with 128 *in silico* mixtures considering both Illumina and BGI platforms

We evaluated UVC with the somatic variant callers Mutect2^1^, Strelka2^22^, VarScan2^14^, LoFreq ^28^, SomaticSniper ^15^, and LoLoPicker ^2^ with a combination of two performance metrics (F-score and PrAUC), two types of small variants (SNV and InDel), two sequencing platforms and aligners (Illumina HiSeq with NovoAlign and BGI MGISEQ with BWA MEM ^17^), four tumor sequencing depths (HiSeq: 240x and 120x. MGISEQ: 48x and 24x), four normal sequencing depths (HiSeq: 120x and 60x. MGISEQ: 24x and 12x), eight tumor purities (1.0, 0.75, 0.5, 0.25, 0.125, 0.0625, 0.03125, and 0.015625), two tumor-in-normal contamination (TiN) rates (0.0 and 0.046875), and two simulated sources of tumor and normal cells (HG001/HG002 as tumor/normal and HG002/HG001 as tumor/normal). Taylor-Weiner et al. ^26^ showed that, even for liquid tumor which is characterized by heavy TiN contamination, the TiN contamination rate is still usually below 0.04. Therefore, We used a TiN contamination rate of 0.046875 to generate the corresponding normal purity from a given tumor purity. Our evaluation shows that UVC is always the best for calling somatic SNVs and InDels from tumor-normal pairs mode except for the following three cases (Figures S41-S104).

1. Case 1 consists of the combination of Streka2, SNV, F-score, MGISEQ with BWA MEM, 24x tumor depth, 24x normal depth, 0.03125 tumor purity, 1.0 normal purity, and HG001/HG002 as tumor/normal.
2. Case 2 consists of the combination of Streka2, SNV, F-score, MGISEQ with BWA MEM, 24x tumor depth, 24x normal depth, 0.015625 tumor purity, 1.0 normal purity, and HG001/HG002 as tumor/normal.
3. Case 3 consists of the combination of Streka2, SNV, PrAUC, MGISEQ with BWA MEM, 48x tumor depth, 24x normal depth, 0.25 tumor purity, 1.0 normal purity, and HG002/HG001 as tumor/normal.

These three cases can be explained by stochastic effects because

1. the difference between the best performance and the UVC performance is negligible and
2. UVC achieves the best performance if the sources of tumor and normal cells (i.e., HG001 and HG002) are swapped in simulation.

A representative portion of the results mentioned in this subsection is provided in Tables 1 and 2.

**Table 1:** Summary F-scores for calling SNVs from the *in silico* mixtures of tumor/normal (T/N) reads sequenced by Illumina HiSeq. More detail can be found in the supplementary results corresponding to Section 3.4.

| T/N depths | T/N purities | UVC-version-441a694 | Mutect2-all | Mutect2-filter | Strelka2 | Varscan2-all | Varscan2-filter | LoFreq* | LoLoPicker |
| --- | --- | --- | --- | --- | --- | --- | --- | --- | --- |
| 240.0/120.0 | 1.0000/1.0000 | **0.9992** | 0.9979 | 0.9352 | 0.9962 | 0.9980 | 0.9978 | 0.9813 | 0.5611 |
| 240.0/120.0 | 0.7500/1.0000 | **0.9992** | 0.9981 | 0.9364 | 0.9966 | 0.9978 | 0.9976 | 0.9788 | N/A |
| 240.0/120.0 | 0.5000/1.0000 | **0.9992** | 0.9979 | 0.9370 | 0.9960 | 0.9376 | 0.9375 | 0.9690 | 0.5593 |
| 240.0/120.0 | 0.2500/1.0000 | **0.9987** | 0.9948 | 0.9364 | 0.9947 | 0.1759 | 0.1759 | 0.9201 | 0.5561 |
| 240.0/120.0 | 0.1250/1.0000 | **0.9917** | 0.9771 | 0.9188 | 0.9852 | 0.0003 | 0.0003 | 0.6883 | 0.5595 |
| 240.0/120.0 | 0.0625/1.0000 | **0.9375** | 0.9009 | 0.8018 | 0.9076 | 0.0000 | N/A | 0.2162 | 0.5395 |
| 240.0/120.0 | 0.0312/1.0000 | **0.7367** | 0.6135 | 0.4266 | 0.5929 | 0.0000 | N/A | 0.0253 | N/A |
| 240.0/120.0 | 0.0156/1.0000 | **0.4436** | 0.2369 | 0.1075 | 0.2144 | 0.0000 | N/A | 0.0015 | N/A |
| 120.0/120.0 | 1.0000/1.0000 | **0.9991** | 0.9981 | 0.9389 | 0.9964 | 0.9980 | 0.9977 | 0.9782 | 0.8050 |
| 120.0/120.0 | 0.7500/1.0000 | **0.9992** | 0.9980 | 0.9396 | 0.9965 | 0.9976 | 0.9973 | 0.9734 | 0.8063 |
| 120.0/120.0 | 0.5000/1.0000 | **0.9988** | 0.9972 | 0.9379 | 0.9967 | 0.9011 | 0.9007 | 0.9574 | 0.8079 |
| 120.0/120.0 | 0.2500/1.0000 | **0.9957** | 0.9890 | 0.9287 | 0.9922 | 0.1770 | 0.1771 | 0.8511 | 0.8092 |
| 120.0/120.0 | 0.1250/1.0000 | **0.9534** | 0.9424 | 0.8612 | 0.9425 | 0.0025 | 0.0025 | 0.4804 | 0.7813 |
| 120.0/120.0 | 0.0625/1.0000 | **0.7970** | 0.7288 | 0.6161 | 0.7344 | 0.0000 | 0.0000 | 0.1166 | 0.6190 |
| 120.0/120.0 | 0.0312/1.0000 | **0.5341** | 0.3552 | 0.2572 | 0.3747 | 0.0000 | 0.0000 | 0.0142 | 0.3340 |
| 120.0/120.0 | 0.0156/1.0000 | **0.2620** | 0.0999 | 0.0601 | 0.1117 | 0.0000 | 0.0000 | 0.0011 | 0.1203 |
| 240.0/60.0 | 1.0000/1.0000 | **0.9990** | 0.9980 | 0.9356 | 0.9957 | 0.9978 | 0.9977 | 0.9818 | 0.5518 |
| 240.0/60.0 | 0.7500/1.0000 | **0.9991** | 0.9983 | 0.9372 | 0.9959 | 0.9975 | 0.9974 | 0.9794 | N/A |
| 240.0/60.0 | 0.5000/1.0000 | **0.9991** | 0.9981 | 0.9378 | 0.9956 | 0.9372 | 0.9374 | 0.9696 | 0.5500 |
| 240.0/60.0 | 0.2500/1.0000 | **0.9986** | 0.9947 | 0.9367 | 0.9943 | 0.1758 | 0.1758 | 0.9212 | 0.5467 |
| 240.0/60.0 | 0.1250/1.0000 | **0.9885** | 0.9773 | 0.9231 | 0.9823 | 0.0003 | 0.0003 | 0.6884 | 0.5459 |
| 240.0/60.0 | 0.0625/1.0000 | **0.9252** | 0.9114 | 0.8300 | 0.8874 | 0.0000 | 0.0000 | 0.2166 | 0.5242 |
| 240.0/60.0 | 0.0312/1.0000 | **0.7057** | 0.6539 | 0.5052 | 0.5719 | 0.0000 | 0.0000 | 0.0252 | N/A |
| 240.0/60.0 | 0.0156/1.0000 | **0.4038** | 0.2764 | 0.1455 | 0.2042 | 0.0000 | 0.0000 | 0.0015 | N/A |
| 240.0/120.0 | 1.0000/0.9375 | **0.9981** | 0.9570 | 0.8399 | 0.9962 | 0.9977 | 0.9975 | 0.2336 | 0.2715 |
| 240.0/120.0 | 0.7500/0.9531 | **0.9985** | 0.9831 | 0.8370 | 0.9963 | 0.9975 | 0.9973 | 0.3646 | N/A |
| 240.0/120.0 | 0.5000/0.9688 | **0.9989** | 0.9951 | 0.8121 | 0.9921 | 0.9375 | 0.9374 | 0.5379 | 0.4582 |
| 240.0/120.0 | 0.2500/0.9844 | **0.9962** | 0.9947 | 0.7897 | 0.9771 | 0.1759 | 0.1759 | 0.7106 | 0.5317 |
| 240.0/120.0 | 0.1250/0.9922 | **0.9814** | 0.9771 | 0.7702 | 0.9759 | 0.0003 | 0.0003 | 0.5894 | 0.5547 |
| 240.0/120.0 | 0.0625/0.9961 | **0.9264** | 0.9035 | 0.7255 | 0.9028 | 0.0000 | N/A | 0.1893 | 0.5386 |
| 240.0/120.0 | 0.0312/0.9980 | **0.7245** | 0.6210 | 0.4000 | 0.5900 | 0.0000 | N/A | 0.0241 | N/A |
| 240.0/120.0 | 0.0156/0.9990 | **0.4370** | 0.2422 | 0.1019 | 0.2138 | 0.0000 | N/A | 0.0015 | N/A |

**Table 2:** Summary F-scores for calling InDels from the *in silico* mixtures of tumor/normal (T/N) reads sequenced by Illumina HiSeq. More detail can be found in the supplementary results corresponding to Section 3.4.

| T/N depths | T/N purities | UVC-version-441a694 | Mutect2-all | Mutect2-filter | Strelka2 | Varscan2-all | Varscan2-filter | LoFreq* | LoLoPicker |
| --- | --- | --- | --- | --- | --- | --- | --- | --- | --- |
| 240.0/120.0 | 1.0000/1.0000 | **0.9896** | 0.9537 | 0.8978 | 0.9727 | 0.9639 | 0.9636 | 0.9083 | N/A |
| 240.0/120.0 | 0.7500/1.0000 | **0.9885** | 0.9677 | 0.9050 | 0.9760 | 0.9540 | 0.9534 | 0.9088 | N/A |
| 240.0/120.0 | 0.5000/1.0000 | **0.9883** | 0.9664 | 0.8265 | 0.9748 | 0.7470 | 0.7470 | 0.9059 | N/A |
| 240.0/120.0 | 0.2500/1.0000 | **0.9869** | 0.9564 | 0.6462 | 0.9428 | 0.1208 | 0.1208 | 0.8860 | N/A |
| 240.0/120.0 | 0.1250/1.0000 | **0.9598** | 0.9130 | 0.6131 | 0.8948 | 0.0000 | 0.0000 | 0.7877 | N/A |
| 240.0/120.0 | 0.0625/1.0000 | **0.8377** | 0.7656 | 0.5204 | 0.7159 | 0.0000 | 0.0000 | 0.5559 | N/A |
| 240.0/120.0 | 0.0312/1.0000 | **0.6138** | 0.4435 | 0.2637 | 0.3693 | 0.0000 | 0.0000 | 0.2553 | N/A |
| 240.0/120.0 | 0.0156/1.0000 | **0.3406** | 0.1755 | 0.0740 | 0.1238 | 0.0000 | 0.0000 | 0.0749 | N/A |
| 120.0/120.0 | 1.0000/1.0000 | **0.9888** | 0.9577 | 0.9016 | 0.9747 | 0.9640 | 0.9631 | 0.9086 | N/A |
| 120.0/120.0 | 0.7500/1.0000 | **0.9888** | 0.9704 | 0.8938 | 0.9770 | 0.9476 | 0.9467 | 0.9067 | N/A |
| 120.0/120.0 | 0.5000/1.0000 | **0.9888** | 0.9691 | 0.7291 | 0.9784 | 0.7348 | 0.7335 | 0.8988 | N/A |
| 120.0/120.0 | 0.2500/1.0000 | **0.9773** | 0.9413 | 0.6219 | 0.9473 | 0.1233 | 0.1233 | 0.8239 | N/A |
| 120.0/120.0 | 0.1250/1.0000 | **0.8845** | 0.8478 | 0.5513 | 0.7919 | 0.0000 | 0.0000 | 0.6304 | N/A |
| 120.0/120.0 | 0.0625/1.0000 | **0.6712** | 0.5731 | 0.3308 | 0.4364 | 0.0000 | 0.0000 | 0.3506 | N/A |
| 120.0/120.0 | 0.0312/1.0000 | **0.4278** | 0.2497 | 0.1152 | 0.1486 | 0.0000 | 0.0000 | 0.1267 | N/A |
| 120.0/120.0 | 0.0156/1.0000 | **0.2044** | 0.0798 | 0.0257 | 0.0351 | 0.0000 | 0.0000 | 0.0277 | N/A |
| 240.0/60.0 | 1.0000/1.0000 | **0.9891** | 0.9549 | 0.8951 | 0.9561 | 0.9607 | 0.9604 | 0.9265 | N/A |
| 240.0/60.0 | 0.7500/1.0000 | **0.9883** | 0.9670 | 0.9030 | 0.9749 | 0.9523 | 0.9517 | 0.9260 | N/A |
| 240.0/60.0 | 0.5000/1.0000 | **0.9883** | 0.9660 | 0.8310 | 0.9743 | 0.7443 | 0.7443 | 0.9224 | N/A |
| 240.0/60.0 | 0.2500/1.0000 | **0.9825** | 0.9560 | 0.6475 | 0.9434 | 0.1204 | 0.1204 | 0.8997 | N/A |
| 240.0/60.0 | 0.1250/1.0000 | **0.9477** | 0.9135 | 0.6132 | 0.8909 | 0.0000 | 0.0000 | 0.7985 | N/A |
| 240.0/60.0 | 0.0625/1.0000 | **0.8146** | 0.7759 | 0.5248 | 0.6968 | 0.0000 | 0.0000 | 0.5653 | N/A |
| 240.0/60.0 | 0.0312/1.0000 | **0.5810** | 0.4633 | 0.2750 | 0.3665 | 0.0000 | 0.0000 | 0.2587 | N/A |
| 240.0/60.0 | 0.0156/1.0000 | **0.3144** | 0.1880 | 0.0859 | 0.1405 | 0.0000 | 0.0000 | 0.0771 | N/A |
| 240.0/120.0 | 1.0000/0.9375 | **0.9877** | 0.9315 | 0.7954 | 0.9732 | 0.9634 | 0.9631 | 0.3380 | N/A |
| 240.0/120.0 | 0.7500/0.9531 | **0.9872** | 0.9569 | 0.7749 | 0.9760 | 0.9539 | 0.9533 | 0.4219 | N/A |
| 240.0/120.0 | 0.5000/0.9688 | **0.9874** | 0.9626 | 0.6944 | 0.9710 | 0.7468 | 0.7468 | 0.5102 | N/A |
| 240.0/120.0 | 0.2500/0.9844 | **0.9825** | 0.9553 | 0.5389 | 0.9075 | 0.1208 | 0.1208 | 0.6453 | N/A |
| 240.0/120.0 | 0.1250/0.9922 | **0.9476** | 0.9125 | 0.5216 | 0.8640 | 0.0000 | 0.0000 | 0.6453 | N/A |
| 240.0/120.0 | 0.0625/0.9961 | **0.8255** | 0.7674 | 0.4785 | 0.6943 | 0.0000 | 0.0000 | 0.4826 | N/A |
| 240.0/120.0 | 0.0312/0.9980 | **0.6024** | 0.4496 | 0.2423 | 0.3631 | 0.0000 | 0.0000 | 0.2404 | N/A |
| 240.0/120.0 | 0.0156/0.9990 | **0.3364** | 0.1748 | 0.0700 | 0.1223 | 0.0000 | 0.0000 | 0.0709 | N/A |

### 3.5 Evaluation results on WGS datasets for calling somatic variants from tumor-normal pairs, with physical mixtures

We evaluated UVC with the somatic variant callers Mutect2^1^, Strelka2^22^, VarScan2^14^, LoFreq ^28^, SomaticSniper ^15^, and LoLoPicker ^2^ with a combination of two performance metrics (F-score and PrAUC) and two types of small variants (SNV and InDel) on the GIAB somatic truth sets following the guideline at https://github.com/hbc/projects/tree/master/giab_somatic/. Our evaluation shows that UVC is always the best for calling somatic SNVs and InDels for the GIAB somatic truth sets without any exception (Figure S105).

### 3.6 Evaluation results on whole-exome-sequencing and amplicon-sequencing datasets for calling somatic variants from tumor-normal pairs, with the breast-cancer cell-line HCC1395

We evaluated UVC with the somatic variant callers Mutect2^1^, Strelka2^22^, VarScan2^14^, LoFreq ^28^, and SomaticSniper ^15^ with a combination of two performance metrics (F-score and PrAUC) and two types of small variants (SNV and InDel) on the FDA-led Sequence-Quality-Control Consortium (SEQC2) somatic reference sets (ftp://ftp-trace.ncbi.nlm.nih.gov/ReferenceSamples/seqc/) ^9^. We evaluated with two strategies for generating truth variants: keeping both medium and high confidence calls, and keeping only high confidence calls. We also evaluated with the quality control (QC) criteria of the following minimum tumor/normal depths: 100x/50x, 200x/50x, 100x/100x, and 200x/100x. Variant loci that fail the QC criteria are not assessed in the QC-criteria-associated evaluation, which is usually done as part the standard operating procedure (SOP) for NGS in clinical settings. We used the seven non-WGS tumor-normal-paired sequencing runs that are publicly available. In sum, our evaluation incorporated an additional combination of two confidence levels for truth variants, four QC criteria, and seven NGS experiments.

Our evaluation shows that UVC is always the best for calling somatic SNV and InDel for the SEQC2 somatic reference sets without any exception (Figures S106-S133). A representative portion of the results mentioned in this subsection is provided in Table 3.

**Table 3:** Summary F-scores for calling SNVs (top) and InDels (bottom) from the SEQC2 datasets obtained by selecting amplicon-sequencing or whole-exome-sequencing (WXS) runs with tumor-normal pairs, evaluating with high-confidence calls in the reference-variant set, and by using 100x tumor and normal minimum sequencing depths as the quality-control (QC) requirements to generate callable regions. More detail can be found in the supplementary results corresponding to Section 3.6.

| Tumor/normal accessions | assay-type | UVC-version-441a694 | Mutect2-all | Mutect2-filter | Strelka2 | Varscan2-all | Varscan2-filter | LoFreq* |
| --- | --- | --- | --- | --- | --- | --- | --- | --- |
| SRR7890876/SRR7890881 | WXS | **0.9622** | 0.8078 | 0.9521 | 0.8928 | 0.7131 | 0.7017 | 0.9492 |
| SRR7890879/SRR7890880 | WXS | **0.9632** | 0.8140 | 0.9490 | 0.8758 | 0.7019 | 0.7279 | 0.9333 |
| SRR7890918/SRR7890919 | WXS | **0.9623** | 0.8349 | 0.9592 | 0.9061 | 0.7678 | 0.7736 | 0.9428 |
| SRR8955955/SRR8955956 | AMPLICON | **0.9616** | 0.7689 | 0.7578 | 0.9091 | 0.8248 | 0.7718 | 0.7978 |
| SRR8955957/SRR8955958 | AMPLICON | **0.9638** | 0.7739 | 0.8075 | 0.9091 | 0.8351 | 0.7934 | 0.8213 |
| SRR8955959/SRR8955960 | AMPLICON | **0.9608** | 0.7929 | 0.7865 | 0.9078 | 0.8307 | 0.7890 | 0.8229 |
| SRR8955982/SRR8955981 | WXS | **0.9527** | 0.8588 | 0.4959 | 0.9130 | 0.7531 | 0.7554 | 0.8571 |
| SRR7890876/SRR7890881 | WXS | **0.8571** | 0.6716 | 0.7965 | N/A | 0.2667 | 0.2667 | 0.7500 |
| SRR7890879/SRR7890880 | WXS | **1.0000** | 0.6667 | 0.7778 | N/A | 0.2500 | 0.2500 | **1.0000** |
| SRR7890918/SRR7890919 | WXS | **0.8679** | 0.7101 | 0.8175 | N/A | 0.3333 | 0.3429 | 0.7925 |
| SRR8955955/SRR8955956 | AMPLICON | **0.8966** | 0.7507 | 0.7595 | N/A | 0.8462 | 0.8333 | 0.8889 |
| SRR8955957/SRR8955958 | AMPLICON | **0.8889** | 0.7814 | 0.7792 | N/A | 0.8000 | 0.7407 | 0.8000 |
| SRR8955959/SRR8955960 | AMPLICON | **0.9164** | 0.8211 | 0.8462 | N/A | 0.8889 | 0.8333 | 0.8462 |
| SRR8955982/SRR8955981 | WXS | **0.8828** | 0.5833 | 0.3636 | 0.0000 | 0.4286 | 0.4286 | 0.4286 |

### 3.7 Evaluation results on amplicon-sequencing datasets for calling somatic variants from tumor-normal pairs, with samples from colon-cancer patients

We compared the performance of UVC with appreci8, which uses machine learning to combine the results of eight variant callers, on the colorectal cancer dataset used to evaluate appreci8^21^. To generate this dataset, human colon organoids and the adjcacent normal tissue were profiled for known colorectal-cancer-associated mutations using the Qiagen Qiaseq Colorectal Cancer Panel, which provides targeted sequencing information for 71 genes. More detail about this dataset is provided by Sandmann et al.; Dame et al. ^21;4^.

This dataset has been manually reviewed by Sandmann et al.; Dame et al. ^21;4^. Thus, Sandmann et al. ^21^ used concordance with their manual review as the variant-calling performance metrics. Sandmann et al. ^21^ already showed that appreci8 outperformed each of the eight individual callers, so we only compared UVC with appreci. The meta variant-caller appreci generated zero false positive calls and seven false negative calls, but UVC generated zero false positive calls and zero false negative calls to achieve 100% concordance with their manual review (Fig. 2).

**Figure 2:**
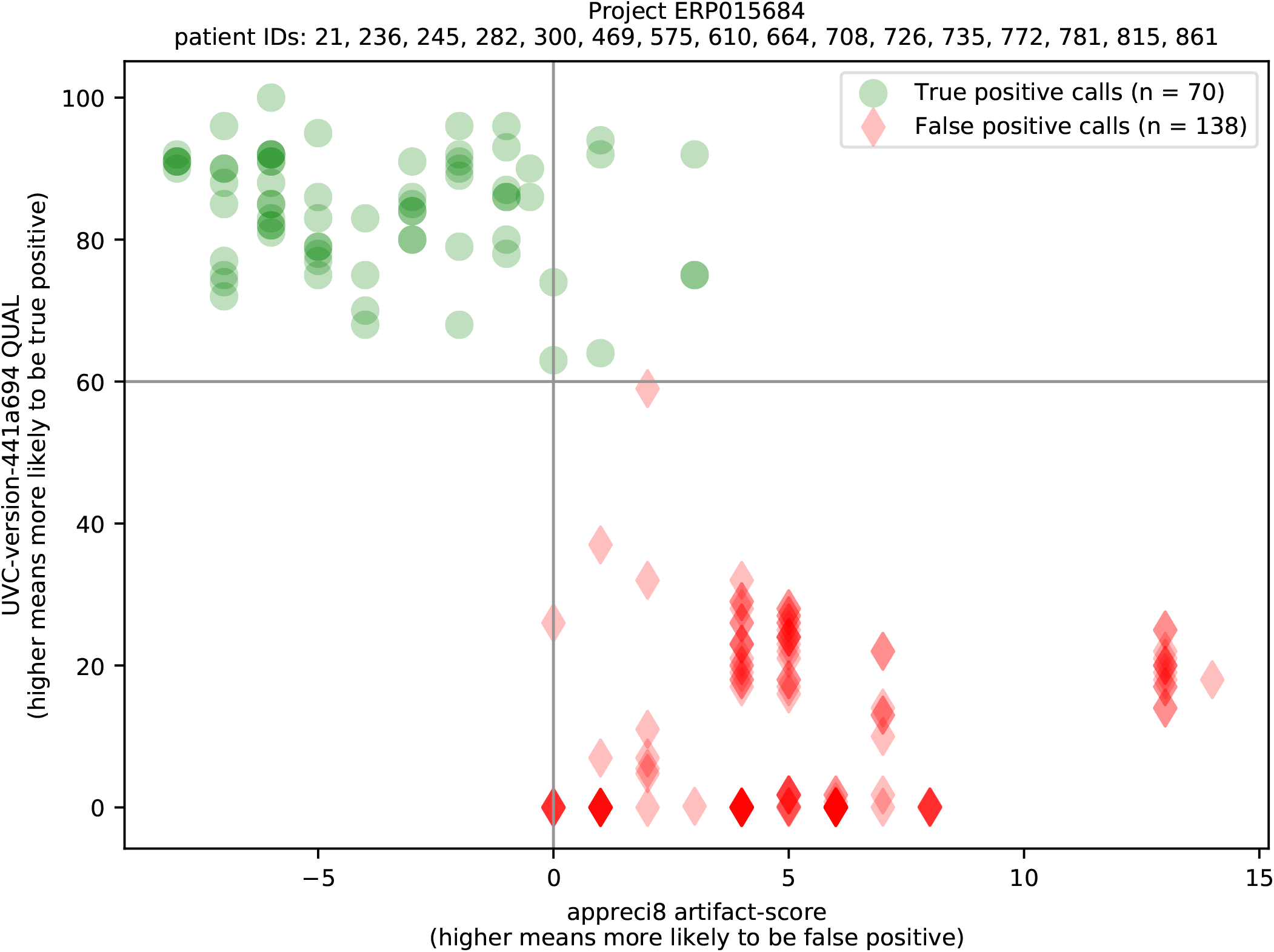
Distribution of variants labeled with the manual review done by Dame et al.; Sandmann et al. ^4;21^. The artifact score was generated by appreci8 by using a machine-learning approach to combine the results of eight variant callers ^21^.

According to UVC, there is an outlier false positive that seems to be clustered with true positives (Fig. 2). Hence, we manually reviewed this outlier variant and found that this outlier variant seems to be real (although still not as real as the true positive calls in general) (Supplementary File 1). Therefore, the probabilistic ranking of variants provided by UVC is extremely accurate.

### 3.8 Evaluation results on amplicon-sequencing datasets with UMI for calling variants in tumor-only mode

To evaluate the performance of UVC on UMI data, we used the dataset of the physical mixtures (mixed at 1:9 ratio) of the reference standard HD734 and the blood of healthy donors to simulate variants which are mostly expected to have an allele fraction of 0.1%. This dataset was originally published by Mageri ^24^. Mageri first trained its model on the dataset of the eight normal samples from the healthy donors, and this dataset is exactly the ground truth for all the true negative calls. Without any training, UVC outperformed Mageri by achieving higher sensitivity (Table S1).

Another UMI-based dataset used in our evaluation is the physical mixture (mixed at 1:99 ratio) of the reference materials HG001 and HG002 which are both well-characterized for germline polymorphisms ^29^. Xu et al. ^29^ have shown that smCounter2 outperformed all other applicable methods on this dataset. Therefore, we compared UVC with only smCounter2. The UMI-specific variant caller smCounter2 already trained its machine-learning model on multiple independent datasets generated by the same assay on the same sample ^29^. Thus, smCounter2 already knows *a priori* the sample-specific, assay-specific, and technology-specific error profiles. Meanwhile, UVC was not aware of any such error profiles. UVC outperformed smCounter2 for calling SNVs and InDels, even though smCounter2 was enhanced with UMI-specific repetitive-region filters ^29^ (Figure S134).

### 3.9 Reanalysis results on an amplicon-sequencing dataset with UMI for calling variants in tumor-normal pairs, which provides additional in-sight about DNA damage repair

UVC is the only variant caller that is aware of the UMIs in both tumor and normal samples, so we applied it to a ultraviolet (UV)-treated subclonal mutation dataset ^8^. In this dataset, SiMSen-seq ^10^, an UMI-based ultra-sensitive amplicon sequencing technique, was used to detect subclonal mutations at selected promoter regions in cells with deficiency in DNA repair. Using the four samples not treated with UV as the “control” and the corresponding four samples treated with UV as “tumor”, UVC identified the previously reported subclonal mutations at the *RPL13A* -116 bp `TTCCG` promoter hotspot site with an allele fraction between 0.05% and 0.5% in all four samples on default settings (Table S2). These mutations were all `C>T` transversions with a flanking `TTCCG` sequence, consistent with previously identified mutational signatures caused by UV radiation in cancer ^10;6^. On the other hand, if the untreated samples were used as “tumor” and the treated samples were used as “normal”, no variant was called by UVC, suggesting a low false positive rate of the variants called on this dataset. In the regions that the original paper did not include, UVC discovered a `C>T` variant before the sequence `TTCCG` 8 bp upstream of the transcription start site of the *DPH3* gene (position 3:16306504 on the GRCh37 genomic coordinates). In the raw data, the mutation is supported by a lot of UMI families in each UV-treated sample but is not supported by any UMI families in each control sample (Figure S135 and Table S2). The newly discovered variant further supports the conclusion made by Elliott et al. ^8^: intrinsic properties of UV radiation, instead of locally inhibited DNA repair, is the core mechanism resulting in the high prevalence of `C>T` base substitution before the sequence `TTCCG`.

### 3.10 Computational resources

The wall-clock running time, CPU running time, and random-access memory consumed by UVC are reasonable and should not be at the bottleneck of a typical variant-calling pipeline (Table S3). Hence, UVC can be incorporated into any variant-calling pipeline in practice.

## 4 Discussion

Here we demonstrated that UVC, a universality-based variant caller that uses a zero-inflated model to combine Bayesian and frequentist inferences of NGS biases, is able to call SNVs and InDels with high accuracy. UVC is both UMI-aware and able to compare the signal in the tumor sample against the signal in the matched normal sample. UVC strongly outperformed other state-of-the-art variant callers on a variety of NGS datasets in terms of sensitivity-specificity trade off. In addition, the improved variant calling of UVC leads to more comprehensive insight about DNA damage repair from a well analyzed existing dataset.

As mentioned in Section 3.2, UVC performs sub-optimally for determining the zygosity of long germline InDels because of strong reference bias. In the future, we may incorporate InDel realignment ^7^, localized assembly ^19^, and/or variation graph ^12^ into UVC to alleviate reference bias.

We hypothesize that the techniques used in UVC can also be used for calling structural variants (SVs) with high accuracy. The reason is that, fundamentally, SV is the same as SNV and InDel: they are all characterized by a reference allele and a non-reference allele, and the reference allele is usually supported by many times more reads than the non-reference allele. Then, for calling SVs, the only additional remaining work is to estimate the probability that a chimeric read is artifactual. In the future, we might implement the feature of calling SVs in UVC.

## 5 Conclusions

UVC significantly improved the state of the art in variant calling from NGS data, which in turn improves the diagnosis, prognosis, and treatment monitoring of cancer in clinical settings. Our work has important clinical applications, such as the development of more accurate cancer screening test and better recommendation system for cancer treatment.

## 6 Availability and requirements

Project name: UVC

Project home page: https://github.com/genetronhealth/uvc

Operating systems: UNIX/Linux

Programming language: C++ complying with the C++14 standard Other requirements: BASH 4.0 or higher

License: e.g. BSD 3-Clause

Any restrictions to use by non-academics: none

## Supporting information

Supplemental results

