## Supplemental methods for "Calling small variants with universality and Bayesian-frequentist hybridism"

### 1 Supplemental methods

Our core variant-calling algorithm is developed by heuristically applying Hegelian dialectics to next-generation sequencing (NGS). Hegelian dialectics is the fundamental cornerstone of modern philosophy as it influenced the development of Nationalism, Marxism-Leninism, etc. Hegelian dialectics consists of a triad of the following.

1. Thesis: a formal statement illustrating a point (e.g., sequencing depth is low). In this paper, our thesis is treated as an assumption in NGS.

2. Antithesis: the contrary, or equivalently the complete negation, of the thesis (e.g., sequencing depth is high). In this paper, our antithesis is treated as a counter-assumption in NGS.

3. Synthesis: the resolution of the conflict between the thesis and its corresponding antithesis (e.g., sequencing depth can be at any level). In this paper, the assumption and its corresponding counter-assumption are resolved (i.e., integrated together) by using a maximum-likelihood interpolation to either perform model selection or generate a mixture model.

By using such triads, we rediscovered many principles found in the literature of NGS bioinformatics, and most importantly, we discovered the following two novel principles.

1. The counter-assumption (antithesis) of infinite sequencing depth reveals a power-law relationship with the exponent of 3 between deviation from theoretical allele fraction and probability of false-positive variant call for both germline and somatic variants for a wide range of allele fractions. Such power-law is described in more detail in Section 1.3.

2. The interpolation (synthesis) between low NGS bias and high NGS bias reveals a Bayesian-frequentist hybrid model with zero inflation, where zero denotes the frequentist null hypothesis of having no bias. Such model is described in more detail in Section 1.4.

In this section, we will use the following slight abuse of notation to achieve more conciseness without losing clarity.

1. The domain of a mathematical function is omitted in equations if the range and meaning of the domain are clear.
2. A logical quantifier is omitted if the quantifier can be inferred from context.

For example, with such abuse of notation, the equation $\left(\tan = \frac{\sin}{\cos}\right)$ is equivalent the expression $\left(\forall_{x\in\mathbb{R}}\left(\tan(x) = \frac{\sin(x)}{\cos(x)}\right)\right)$.

#### 1.1 Interpolation between pairs of contrary assumptions: a thought experiment involving extreme cases

Variant calling is a complex problem. To tackle this problem, we begin by making the following assumptions.

1. The depth of coverage at a variant locus is very low.
2. The variant signal has no bias.
3. The reads covering a variant site are all not labeled with unique molecular identifiers (UMIs).
4. The variant origin, which can be either germline or somatic, cannot be determined.
5. The variant is a single nucleotide variant (SNV).
6. The variant signal is not characterized by any systematic error.
7. The sequencing platform is Illumina.

Then, we iteratively relax each assumption. Section 1.2 describes the variant-calling model if no assumption is relaxed. When an assumption is relaxed, it is first relaxed to the greatest degree with strong negation, which means that the key adjective or adverb in the original assumption is replaced by its antonym to make a new contrary assumption. The corresponding new contrary assumptions are as follows.

1. The depth of coverage at a variant locus is very high.
2. The variant signal has the highest possible bias.
3. The reads covering a variant site are all labeled with UMIs.
4. The true variant origin has already been determined with 100% accuracy. For example, we already know that a variant can only be of germline origin.
5. The variant is not an SNV.
6. The variant signal is strongly characterized by all sources of systematic error.
7. The sequencing platform is not Illumina.

Then, we interpolate between each original assumption and its corresponding contrary assumption, where the two paired assumptions are the strong negations of each other. Our interpolation is from extreme cases. Hence, our interpolation results in a variant-calling model that is both theoretically sound and broadly applicable to all NGS scenarios. Such interpolations result in the synthesis of formulae that either perform model selection or generate new mixture models using maximum likelihood. Sections 1.3 to 1.9 describe the progressive updates to our variant-calling model by incorporating the contrary assumptions 1, 2, 3, 4, 5, 6, and 7, respectively.

#### 1.2 Idealistic variant calls: at low sequencing depth, without any bias, without using molecular barcodes, without determination of germline-versus-somatic origin, ignoring insertion-deletions (InDels) to only call single-nucleotide variants (SNVs), without any systematic error, and considering only the Illumina sequencing platform

Illumina sequencers usually sequence each fragment twice, one time from each end. Hence, the middle region of a fragment, or even an entire fragment, can be sequenced twice. The twice-sequenced region should be counted only once instead of twice. At the same time, the quality of each base in the twice-sequenced region needs to be adjusted. There are various methods to adjust such base qualities [10;11;23]. Most methods assume that errors coming from the two read ends are independent of each other. Such assumption is quite strong as a lot of sources of errors, such as PCR errors, can affect both ends. Instead, we assume that there is up to 100% correlation between the errors on both ends. If the two bases from both ends agree at a position, then the quality of the merged base is simply the maximum of the qualities of the two bases. If the two bases from both ends disagree at a position, then the most likely non-erroneous base is the one with higher base quality, and the difference of the two qualities of the two disagreeing bases is approximately the Phred-scaled probability that the most likely non-erroneous base is erroneous. The above reasoning leads to the following formula for computing the combined fragment quality in the twice-sequenced region where the R1 and R2 ends overlap with each other.

$$\text{R1R2overlapQ}(q_1, q_2) = \begin{cases} \max(q_1, q_2) & \text{if } b_1 = b_2 \\ \max(q_1, q_2) - \min(q_1, q_2) & \text{if } b_1 \neq b_2 \end{cases} \quad (1)$$

where $q_1$ and $q_2$ are the qualities of the bases $b_1$ and $b_2$ from the paired ends R1 and R2, respectively, such that $b_1$ and $b_2$ originated from the same fragment. If the data of interest is produced by single-end sequencing or the region of interest is only covered by either the R1 end or the R2 end of a fragment, then Eq. (1) is not applicable and is thus not used. In fact, Eq. (1) is highly concordant with the empirical relationship used by NGmerge [6], so our merging procedure is not only theoretically sound but also empirically valid.

After merging the R1 and R2 ends, the next step is to compute the quality of the variant of interest. By our assumption, the depth of coverage is sufficiently low, so the statistical noise caused by the stochasticity in basecalling is the most significant source of errors. Therefore, the signals on different reads can be considered to be statistically independent. Then, we can statistically test whether the observed basecalls can be explained by the errors associated with their base qualities. By assuming that basecalling is the only source of errors, we obtain the following formula to compute variant quality.

$$\text{LowDepthQ}(a, B) = \max_{b_1 \in B} \text{LowDepthQ2}(a, \{x | x \in B \wedge x \geq b_1\}) \quad (2)$$

where

$$\text{LowDepthQ2}(a, B) = |B| \times \left( \min(B) + 10 \times \log_{10}\left(\frac{|B|}{a}\right)\right) \quad (3)$$

and where $a$ is the total fragment depth of all alleles at a locus and $B$ is the set of all base qualities of the variant allele of interest. In other words, Eq. (2) tries each threshold of base quality, filters out all reads below the threshold, and then calculates the signal-to-noise ratio (SNR) of the variant by multiplying the SNR of each read support.

#### 1.3 Variant calls: generalization from low sequencing depth to high sequencing depth

It has been well known that variant candidates with low allele fractions are likely to be false positive. No universal standard has been established for the allele fraction threshold for sequencing analysis, and thresholds of 10%, 5%, and 3% have been reported in the literature[9]. However, the relationship between allele fraction and false positive rate has not been explicitly modeled using a wide range of allele fractions. Mathematical models that fit to a wide range of values are typically characterized by the property of scale invariance which is a key characteristic of power law. Hence, we used a power-law model of false positive rate using allele fraction.

In our power-law model, we assume that allele depth is infinitely high. Although our assumption is invalid, it leads to a much simplified version of the complex problem of variant calling. Then, under this assumption, we hypothesize that the observed and expected allele fractions $f$ and $g$ are related to each other as follows.

$$\text{HighDepthR}(f, g) = \left(\max\left(\frac{f}{g}, \frac{g}{f}\right)\right)^{-3} \tag{4}$$

which is equivalent to the following formula in Phred-scale.

$$\text{HighDepthQ}(f, g) = 3 \times 10 \times \log_{10} \min\left(\frac{f}{g}, \frac{g}{f}\right) \tag{5}$$

where $\text{HighDepthR}(f, g)$ and $\text{HighDepthQ}(f, g)$ denote the raw and Phred-scaled likelihoods that the observed allele with a fraction of $f$ is a true biological variant that is expected to have a fraction of $g$. In fact, Eq. (5) can be verbally expressed as follows.

**Empirical Law 1.** *If the observed allele fraction is doubled or halved and deviates further from its expected allele fraction, then the likelihood that the observed allele is false positive increases by approximately eight.*

Equation (5) is a power-law distribution with the exponent of 3. The exponent of 3 is at the bifurcation point above/below which the variance of the power-law distribution is finite/infinite, which conforms to our intuition that the allele fractions of false positive variants have extremely large but finite variance. In addition, a power-law distribution with the exponent of 3 can be theoretically generated by the well-known process of linear preferential attachment[1]. Furthermore, a duplex consensus sequence (DCS) which requires two single-strand consensus sequences (SSCSs) has an error rate that is approximately $\frac{1}{10}$ of the error rate of an SSCS[16], and $\frac{1}{10}$ is close to the $\frac{1}{2^3}$ predicted by the power law with the exponent of 3. Most importantly, the first subsection in the supplementary results shows that, indeed, the relationship between false positive rate and allele fraction empirically follows a power law with the exponent of 3. Furthermore, Muyas et al.[14] showed that Eq. (5) also approximately holds for germline variants if allele fraction is replaced by allele odds ratio.

Then, Eq. (5) implies that the allele fraction $f$ is heuristically related to $\text{HighDepthR}(f)$, which is its odds of being false positive, as follows.

$$\text{HighDepthR}(f) = 10^{-9.5} \times f^{-3} \tag{6}$$

which is equivalent to the following formula in Phred-scale.

$$\text{HighDepthQ}(f) = 95 + 3 \times 10 \times \log_{10}(f) \tag{7}$$

where $\text{HighDepthR}(f)$ and $\text{HighDepthQ}(f)$ are the raw and Phred-scaled odds that a variant at a base position is false positive. Interestingly, Eq. (7) strongly conforms to our intuition as demonstrated by the following facts.

1. HighDepthR(1) $\approx 10^{-9.5}$. The constant $10^{-9.5}$ is approximately the rate of germline mosaicism[22], which is approximately the probability that a *de novo* germline mutation, which cannot be distinguished from a somatic mutation, occurs *in vivo* at a base position.

2. For Illumina sequencing data, HighDepthR($10^{-3}$) $\approx 1$ according to the common practice of discarding bases with basecall qualities of less than 30.

3. For any type of sequencing data, HighDepthR(0) $= \infty$. If a variant candidate is not supported by any read at all at extremely high depth of coverage, then this variant candidate is almost surely a true negative.

Most importantly, the first subsection in the supplementary results shows that Eq. (7) is empirically valid.

Then, we interpolate between Eq. (7) and Eq. (2), which were defined in Section 1.2, to obtain a formula that computes the variant quality at any depth of coverage. More specifically, we select either Eq. (7) or Eq. (2) such that the selected formula generates lower variant quality because such selected formula provides a more likely explanation for the observed signal. Thus, the interpolated formula is as follows.

$$\text{AnyDepthQ}(a, B) = \min\left(\text{HighDepthQ}\left(\frac{|B|}{a}\right), \text{LowDepthQ}(a, B)\right) \tag{8}$$

where $a$ and $B$ were previously defined in Eq. (3). The formula AnyDepthQ describes the interplay between allele fraction and the combination of sequencing depth and base quality. Most importantly, AnyDepthQ implies that the variant quality AnyDepthQ($a, B$) is capped by either HighDepthQ, which corresponds to an allele-fraction cutoff, or LowDepthQ, which corresponds to a cutoff derived from a combination of base quality and sequencing depth. Hence, AnyDepthQ calculates the optimal cutoff based on allele fraction given a combination of cutoffs for base quality and sequencing depth, and vice versa. As a result, AnyDepthQ provides theoretically optimal cutoffs for allele fraction, base quality, and sequencing depth as a by-product of computing variant qualities. Please note that, in Eq. (8), HighDepthQ provides an alternative model to LowDepthQ instead of providing prior information to LowDepthQ.

#### 1.4 Variant calls: incorporation of bias

Bias plays a significant role in NGS. Bias in NGS is similar to bias in other phenomena such as coin toss. In fact, all biases have three important properties.

1. There is always exactly one ideal unbiased situation with the assumption of uniform distribution. For example, in an ideal unbiased coin toss, the coin has 50%/50% chances of landing in heads/tails, respectively.

2. Biases cannot be distinguished from statistical noise if the number of observations is low. For example, the effect size, or equivalently extent, of bias cannot be determined from one single coin toss.

3. If the number of observations is sufficiently high, then biases can always be distinguished from statistical noise regardless of how small the effect size of the bias is. For example, if the tail of a coin is heavier than its head, then the probability of the coin landing in its tail is slightly higher than the probability of landing in its head, so the probability of the coin landing in its head will converge to such probability that is less than 50% as the number of observed outcomes of coin toss approaches infinity. Importantly, we did not know *a priori* that the tail is heavier than the head, but we did know *a priori* that one side is heavier than the other side even though the weight difference can be extremely small.

These three simple observations result in three simple heuristics.

1. If the number of observations is low, then we should assume that there is no bias in the underlying phenomena.
2. If the number of observations is sufficiently high, then the effect size of the bias can be estimated with sufficient accuracy. For example, if a coin lands on its head 1000 times and on its tail 9000 times, then the probability of this coin landing in its head is estimated to be 10%.
3. If the number of observations is neither low nor sufficiently high, then we should interpolate between the two cases: the case of no bias and the case of bias with the effect size inferred from observations.

Now, let us apply our heuristics to NGS bioinformatics. First, given a type of bias, a sequenced segment can be either biased or unbiased. For example, if we consider the reads on the reverse strand to be biased, then the reads on the forward strand are unbiased; if we consider the reads close to their read ends to be biased, then the reads far away from their ends are unbiased; etc. Then, let us define the following variables.

1. Let $a$ denote the number of all reads supporting the variant allele.
2. Let $b$ denote the number of all reads supporting all alleles including the reference allele.
3. Let $a'$ denote the number of unbiased reads supporting the variant allele.
4. Let $b'$ denote the number of unbiased reads supporting all alleles including the reference allele.
5. Let $p$ denote a threshold of information gain which is similar to a $P$ value.

The bias-adjusted allele fraction $f'$ is then defined as follows.

$$f'(a,b,a',b',p) = \max\left(\frac{1}{\exp(\max(0,p'-p))} \times \frac{a}{b}, \frac{a'}{b'}\right) \tag{9}$$

$$p' = b' \times \mathrm{D}_{\mathrm{KL}}^{\mathrm{Bernoulli}}\left(\frac{a'}{b'}, \frac{a}{b}\right) \tag{10}$$

where $\mathrm{D}_{\mathrm{KL}}^{\mathrm{Bernoulli}}(x,y)$ denotes the Kullback–Leibler (KL) divergence, or equivalently relative entropy, from an expected Bernoulli distribution with the rate parameter $y$ to another observed Bernoulli distribution with the rate parameter $x$. More specifically,

$$\mathrm{D}_{\mathrm{KL}}^{\mathrm{Bernoulli}}(x,y) = x \times \log\left(\frac{x}{y}\right) + (1-x) \times \log\left(\frac{1-x}{1-y}\right) \tag{11}$$

where $x$ and $y$ denote two probabilities of observing the variant allele, respectively. Equation (9) provides an information-theoretic interpolation of our simple heuristics. If the allele fraction $\frac{a'}{b'}$ of reads without bias deviates away from the allele fraction $\frac{a}{b}$ of all reads, then $\frac{a'}{b'}$ provides a higher level of surprise, so we gain more information about the bias, and vice versa. Similarly, if the depth of coverage $b'$ increases, then the variance of the bias decreases, so we gain more information about the bias, and vice versa. If the information gain is below the threshold $p$, then the null hypothesis that no bias is present is accepted, resulting in no adjustment of allele fraction. Otherwise, the null hypothesis is rejected, but the variant is not rejected. In this case, the allele fraction of all reads is reduced by $\exp(p'-p)$ fold, where $(p'-p)$ is exactly equal to the amount of information gain measured in nats. At the same time, the allele fraction cannot be reduced below $\frac{a'}{b'}$ which is computed using only reads without bias.

Interestingly, the value of $f'$ as a function of $p'$ has a sigmoid shape. The minimum and maximum of the sigmoid function represent allele fractions with very sufficient and very insufficient information gain due to bias, respectively. This sigmoid function is a combination of two rectified linear units in logarithmic scale so that Eq. (9) is similar to a perceptron, the fundamental unit of neural networks. Hence, Eq. (9) can be interpreted in terms of neural networks.

More interestingly, the variable $p$ can be interpreted as a Phred-scaled $P$ value, whereas the variable $\frac{a'}{b'}$ provides a Bayesian estimate of the allele fraction that is not affected by bias. Thus, if $p' < p$, then Eq. (9) deactivates Bayesian inference to use only frequentist inference. Otherwise, Bayesian inference is gradually mixed with frequentist inference as the number of observations increases. In the other extreme case, if $\frac{1}{\exp(\max(0, p'-p))} \times \frac{a}{b} < \frac{a'}{b'}$, then Eq. (9) deactivates frequentist inference to consider only Bayesian inference. Hence, Eq. (9) can be interpreted in terms of inferential statistics with model selection.

In this work, we considered four main types of biases: position bias, insert bias, strand bias, and read-orientation bias. A sequenced read segment is considered to be position-biased for a variant to the left/right segment end if the variant is within $5 + x$ number of bases to the left/right segment end, respectively, where $x$ is the root-mean-square number of bases inserted or deleted at this position. For position bias, $p$ varies depending on the sequence and read context of the variant as follows.

1. $p = \ln(10) \times \frac{40}{10}$ by default because $10^{-4}$ is approximately the probability that a genomic position is characterized by a true positive InDel variant that may induce false positive SNV variants nearby.

2. $p = \ln(10) \times \frac{20}{10}$ if the number of reads covering the locus of the ALT allele and supporting any non-ALT InDel at any position is higher than the number of reads supporting the ALT allele at this locus, because $10^{-2}$ is approximately the probability that a genomic position is characterized by a true positive InDel variant, given that a strong candidate for such InDel variant exists within approximately 50 bases.

Insert bias is similar to position bias. For insert bias, we let $p = \ln(10) \times \frac{q-15}{10}$, where $q$ is the root-mean-square value of the mapping qualities. We empirically observed that $10^{-15/10}$ is approximately the background fraction of reads that are not properly paired in a typical NGS run, so the formula for $p$ computes the likelihood ratio of random background noise to systematic mapping error. A sequenced read segment is considered to be insert-biased for a variant to the left/right insert end if the variant is not within $\frac{1}{2} \times x$ to $2 \times x$ number of bases to the left/right insert end, respectively, where $x$ is the average number of bases to the left/right insert end, respectively.

For strand bias, we let $p = \ln(10) \times \frac{q+10}{10}$, where $q$ is the root-mean-square value of the base qualities. The formula for $p$ captures the heuristic that strand bias tends to co-occur with low base qualities. For position bias, a read is biased to be closer to its read end, whereas for strand bias, a read is biased to be on either the forward strand or the reverse strand. Hence, we can obtain two allele fractions for strand biases on the forward and reverse stands, respectively, and the minimum allele fraction is considered to be the strand-bias allele fraction.

Read-orientation (abbreviated as orientation unless stated otherwise) bias is similar to strand bias. The forward and reverse strands are similar to the R1R2 and R2R1 orientations. For orientation bias, we let $p = \ln(10) \times (\frac{45}{10} + \log_{10}(f))$, where $f$ is the allele fraction of the variant of interest, because the probability of observing sequencing artifact in formalin-fixed tumors is approximately inversely proportional to allele fraction[21], and because the expected allele fraction caused by sequencing artifact is approximately $10^{-4.5}$ prior to PCR amplification[19].

In the end, each bias has a corresponding bias-reduced allele fraction by applying Eq. (9). Finally, the lowest bias-reduced allele fraction is treated as the effective allele fraction. Unless stated otherwise, the effective allele fraction is used for all purposes in situations involving allele fractions, such as in Eq. (2) and Eq. (5).

##### 1.4.1 Duplication bias

Duplication bias is special because we have to first define the criteria for considering two reads as being the duplicates of each other. Typically, reads mapped to the same chromosome, insert start position, and insert end position are assumed to be duplicates of each other. Unfortunately, this assumption may not hold as Sena [17] observed the following paradox: the PCR products presumably derived from the same molecule according to their unique molecular identifiers (UMIs) may map to slightly different genomic locations. This paradox can be partially explained by PCR stutter [17]. Therefore, to better cluster reads into duplicate families, we developed a density-based deduplication algorithm.

First, at each genomic position, the number of reads that start and/or end at this position is computed. Then, each position, referred to as the centroid, attracts other nearly positions, referred to as the satellites. The attraction strength is directly proportional to the number $x$ of reads that start and/or end at the centroid, is inversely proportional to the number $y$ of reads that start and/or end at the satellite, and decays exponentially as a function of the number $z$ of positions between the centroid and the satellite. More specifically, the attraction strength $s$ is defined as follows.

$$s = \frac{x}{y} \times 5^{-z} \tag{12}$$

where the constant 5 is approximately estimated from the most extreme mapping shifts shown by Sena [17] Figure 6. Positions that are strongly attracted to a second position (i.e., $s > 1$) are considered to be in the same duplicate family as the second position and are therefore clustered with the second position. And the second position is the centroid of the cluster if the second position is not attracted to any other position. Our clustering procedure is similar to the one presented by Edgar and Flyvbjerg [5] Figure 2, except that correction is performed for start and end positions instead of sequencing errors. This clustering procedure is supposed to especially improve the deduplication of PCR-amplified and UMI-labeled reads.

Afterwards, duplication bias is computed by setting $p = -\infty$, effectively turning Eq. (9) into the following.

$$f'(a, b, a', b', p) = \max\left(\frac{a'}{b'}\right) \tag{13}$$

where $a$ is the number of non-deduplicated (with duplicates kept) reads supporting the ALT allele, $b$ is the number of non-deduplicated reads supporting any allele. $a'$ is the number of deduplicated (with duplicates removed) reads supporting the ALT allele, and $b'$ is the number of deduplicated reads supporting any allele. Simply put, the allele fraction adjusted by duplication bias is the one computed using only deduplicated reads.

#### 1.5 Variant calls: incorporation of unique molecular identifiers (UMIs)

The introduction of unique molecular identifiers (UMIs), which are also known as molecular barcodes, greatly improves the specificity and sensitivity of somatic variant detection, in the situation that multiple PCR copies of the fragment of each original DNA molecule are often sequenced.

First, all read supports with base qualities (BQs) of less than 25 are filtered out. If a UMI family contains a sufficient number of base-calls covering a locus, then a UMI-derived BQ can be estimated for this locus. Intuitively, if the consensus allele of a UMI family of reads is different than a second allele supported by one single read in the UMI family, then the second allele is highly likely to be caused by PCR or sequencing error, so the comparison between the allele supported by the consensus of each UMI family and the allele supported by each read in terms of read count can provide an estimation for the NGS error rate after UMI attachment. More formally, the UMI-derived BQ of an allele $a$ supported by a multi-set $\mathbb{B}$ of UMI families, where each family is a multi-set of basecalls supporting the locus of $a$, is defined as follows.

$$\mathrm{BQ}^{\mathrm{UMI}}(a, \mathbb{B}) = \frac{\sum_{B\in\mathbb{B}'}(|\{b \in B | b = a\}|)}{\sum_{B\in\mathbb{B}'}(|B|)} \tag{14}$$

where

$$\mathbb{B}' = \{B \in \mathbb{B} | (|B| > 4 \wedge \mathrm{cons}(B) \neq a \wedge \frac{|\{b \in B | b = \mathrm{cons}(B)\}|}{|B|} \geq \frac{2}{3})\} \tag{15}$$

Here, $B$ denotes a multi-set of basecalls supporting the locus of interest in a UMI family, $b$ denotes one single basecall in $B$, $\mathrm{cons}(B)$ denotes the consensus base (i.e., the one with the most frequently occurring basecall) in the family $B$, and the symbol "|" means "such that". The UMI-derived BQ is especially useful to call InDels for Illumina-like sequencing data and to call SNVs for IonTorrent sequencing data because Illumina sequencers do not generate any BQs for InDels and IonTorrent sequencers do not generate any BQs for SNVs (Sections 1.7 and 1.9).

Then, to incorporate UMIs into our computation, we modify our parameters in Section 1.2 and Section 1.3.

Equation (2) can be adapted from reads to families. For each family of reads sharing the same UMI at each position, the consensus base of the family is defined as the most frequently occurring base. Then, the sum of the UMI-derived base qualities of the consensus base, subtracted by the sum of the base qualities of the non-consensus bases, is the consensus quality of the consensus base for this family. Then, we apply Eq. (2) to families instead of reads, where $a$ is the total family depth of all alleles at a locus and $B$ is the set of all consensus qualities of the variant allele of interest.

Equation (5) can also be adapted from reads to families. A family of reads sharing the same UMI is of good quality at a position if the family is supported by at least 2 reads and 80% of the bases in the family agree with the consensus base. Then, the allele fraction of the variant of interest computed using good-quality families instead of reads is used in Eq. (5). Due to the nature of consensus, families are less prone to error than reads. Hence, for families,

$$\mathrm{HighDepthQ}(f) = 95 + 3 \times 10 \times \log_{10}(f) + 41 \tag{16}$$

where $f$ is the allele fraction computed using families of good quality. The constant 41 is derived from the fact that the empirical error probabilities of Q30 bases and high-quality families are $q_1 = 2.7 \times 10^{-3}$–$3.5\times10^{-5}$ and $q_2 = 1.5\times10^{-4}$–$3.5\times10^{-5}$, respectively[16], where $41 \approx 3\times10/\log(10)\times\log(q_1/q_2)$.

The bias in reads should also be present in families which are formed by grouping reads. Hence, Eq. (9) can be easily adapted from reads to families: the effective allele fraction is modified by multiplying by the allele fraction computed using families and then dividing by the allele fraction computed using reads. Unless stated otherwise, the modified effective allele fraction is used for all purposes in situations involving allele fractions, such as in Eq. (2) and Eq. (5).

UMI-families, which represent the original DNA molecules, can be modeled with statistically independent generation by drawing with replacement. However, if a bigger number of sequenced segments (i.e., FASTQ records, each consisting of four consecutive lines) are derived from a smaller number of molecules (which is the case for the over-sequenced cfDNA molecules labeled with UMIs), then the sequenced segments are neither statistically independent of each other nor drawn with replacement. Fortunately, in Eq. (9), the effects of generation without statistical independence and of drawing without replacement practically annihilate each other: statistical tests have smaller-than-expected $P$ values due to statistical dependence, but the smaller $P$ values also have less impact in Eq. (9) because almost all original DNA molecules are observed in the sequencing data. Hence, although all biases are computed using only sequenced segments, these biases are applied to both sequenced segments and UMI-families.

Finally, the variant qualities computed without considering any UMIs and the variant qualities computed with all UMIs considered are combined. In Section 1.3, different error-generation processes are combined into one single model. Thus, the error-generation process resulting in the lowest variant quality is selected because such process provides the most likely explanation for the observed signal. On the contrary, in this section, the same error-generation process is fit with two models: with UMI and without UMI. Thus, the model resulting in the highest variant quality is selected because such model provides the most comprehensive explanation for the observed signal.

#### 1.6 Variant calls: determination of germline-versus-somatic origin

A real biological variant can be of either germline or somatic origin. The genome of a diploid living organism such as human has the following four possible combinations of alleles at each genetic locus: homozygous reference (denoted as HomRef or `0/0`), heterozygous alternated (denoted as Hetero or `0/1`), homozygous alternated (denoted as HomAlt or `1/1`), and heterozygous triallelic (denoted as Het3al or `1/2`). According to our empirical observation, the SNP genotypes `0/0`, `0/1`, `1/1`, and `1/2` have the Phred-scaled prior probabilities of 0, 31, 33, and 58, respectively (single-nucleotide variants (SNVs) of germline origin are referred to as single-nucleotide variants (SNVs)). To get the posterior probabilities of each genotype, we can apply the binomial and power-law models presented in Section 1.2 and Section 1.3. The binomial model assumes that alleles supported by different reads at the same locus are independent and identically distributed. Thus, given the REF, first ALT, and second ALT alleles with allele fractions $f_0$, $f_1$, and $f_2$, we obtain the following formulae to compute the binomial likelihood of each genotype.

$$\mathrm{GL}_{\mathtt{0/0}}^{\mathrm{Binomial}}(f_0, f_1) = \frac{10}{\log(10)} \times d \times \mathrm{D}_{\mathrm{KL}}^{\mathrm{Bernoulli}}\left(\frac{f_1}{f_0 + f_1}, \epsilon\right) \tag{17}$$

$$\mathrm{GL}_{\mathtt{0/1}}^{\mathrm{Binomial}}(f_0, f_1) = \frac{10}{\log(10)} \times d \times \mathrm{D}_{\mathrm{KL}}^{\mathrm{Bernoulli}}\left(\frac{f_1}{f_0 + f_1}, \frac{1}{2}\right) \tag{18}$$

$$\mathrm{GL}_{\mathtt{1/1}}^{\mathrm{Binomial}}(f_0, f_1) = \frac{10}{\log(10)} \times d \times \mathrm{D}_{\mathrm{KL}}^{\mathrm{Bernoulli}}\left(\frac{f_0}{f_0 + f_1}, \epsilon\right) \tag{19}$$

$$\mathrm{GL}_{\mathtt{1/2}}^{\mathrm{Binomial}}(f_0, f_1) = \frac{10}{\log(10)} \times d \times \mathrm{D}_{\mathrm{KL}}^{\mathrm{Bernoulli}}\left(\frac{f_1}{f_1 + f_2}, \frac{1}{2}\right) \tag{20}$$

where $d$ is the total number of reads that cover the locus. The function $\mathrm{D}_{\mathrm{KL}}^{\mathrm{Bernoulli}}(x, y)$, which is defined in Eq. (11), denotes the Kullback–Leibler divergence from an expected Bernoulli distribution with the rate parameter $y$ to another observed Bernoulli distribution with the rate parameter $x$. Cibulskis et al.[2] observed that usually less than 1.5% of read depth is caused by contamination from other samples. Thus, we let $\epsilon = 0.02$ by default to account for any contamination from other samples and other sources of errors. Then, by applying Empirical Law 1 and Eq. (5) to germline variants, we can derive the following formulae to compute the power-law likelihood of each genotype.

$$\mathrm{GL}_{\mathtt{0/0}}^{\mathrm{PowerLaw}}(f_0, f_1) = 3 \times \min\left(0, 10 \times \log_{10}\left(\frac{\epsilon \times f_0}{f_1}\right)\right) \tag{21}$$

$$\mathrm{GL}_{\mathtt{0/1}}^{\mathrm{PowerLaw}}(f_0, f_1) = 3 \times \min\left(0, 10 \times \log_{10}\left(\min\left(\frac{f_1}{f_0}, \frac{f_0}{f_1}\right)\right)\right) \tag{22}$$

$$\mathrm{GL}_{\mathtt{1/1}}^{\mathrm{PowerLaw}}(f_0, f_1) = 3 \times \min\left(0, 10 \times \log_{10}\left(\frac{\epsilon \times f_1}{f_0}\right)\right) \tag{23}$$

$$\mathrm{GL}_{\mathtt{1/2}}^{\mathrm{PowerLaw}}(f_1, f_2) = 3 \times \min\left(0, 10 \times \log_{10}\left(\min\left(\frac{f_1}{f_2}, \frac{f_2}{f_1}\right)\right)\right) \tag{24}$$

To let the computation of each genotype likelihood incorporate both binomial and power-law models, we combine Eqs. (17) to (20) and Eqs. (21) to (24) as follows.

$$\mathrm{GL}_{\mathcal{G}} = \mathrm{GL}_{\mathcal{G}}^{\mathrm{Prior}} + \max(\mathrm{GL}_{\mathcal{G}}^{\mathrm{Binomial}}, \mathrm{GL}_{\mathcal{G}}^{\mathrm{PowerLaw}}) \tag{25}$$

where $\mathcal{G}$ is the genotype for which the likelihood is computed and $\mathrm{GL}_{\mathcal{G}}^{\mathrm{Prior}}$ is the Phred-scaled prior probability for $\mathcal{G}$. We empirically observed that, for SNPs,

$$\mathrm{GL}_{\texttt{0/0}}^{\mathrm{Prior}} = 0 \qquad \mathrm{GL}_{\texttt{0/1}}^{\mathrm{Prior}} = -31 \qquad \mathrm{GL}_{\texttt{1/1}}^{\mathrm{Prior}} = -33 \qquad \mathrm{GL}_{\texttt{1/2}}^{\mathrm{Prior}} = -58 \tag{26}$$

In Eq. (25), the max function performs model selection: the model that best describes the data is used. As the sequencing depth becomes sufficiently high, the power-law model becomes increasingly more likely to be selected, and vice versa, which conforms to our intuition that allele fraction becomes increasingly more important than sequencing depth for calling variants as sequencing depth increases.

Finally, without being aware of the tumor and given only the normal BAM file, the Phred-scaled probability that a variant candidate is not of germline origin is estimated with $\mathrm{HomRefQ}^{\mathrm{N}}$ which is defined as follows.

$$\mathrm{HomRefQ}^{\mathrm{N}} = \mathrm{GL}_{\texttt{0/0}} - \max(\mathrm{GL}_{\texttt{0/1}}, \mathrm{GL}_{\texttt{1/1}}, \mathrm{GL}_{\texttt{1/2}}) \tag{27}$$

Finally, the genotype GT and genotype quality GQ of each germline-variant candidate are computed according to the way that the VCF specification is interpreted by GATK HaplotypeCaller[4;12]. More specifically,

$$\mathrm{GT} = \left( \underset{\mathcal{G} \in \{\texttt{0/0,0/1,1/1,1/2}\}}{\arg\max} \mathrm{GL}_{\mathcal{G}} \right) \tag{28}$$

$$\mathrm{GQ} = \mathrm{GL}_{\mathrm{GT}} - \left( \max_{\mathcal{G} \in (\{\texttt{0/0,0/1,1/1,1/2}\} \setminus \{\mathrm{GT}\})} \mathrm{GL}_{\mathcal{G}} \right) \tag{29}$$

where the operator $\setminus$ denotes set minus.

##### 1.6.1 Variant calls: adjustment of tumor variant quality by the matched normal

Let AnyAltQ be the tumor-sample variant quality obtained by applying the techniques in Sections 1.4 and 1.5 to the quality AnyDepthQ computed by Eq. (8) in Section 1.3. Then, for somatic variant call, AnyAltQ is modified by adding $\mathrm{TN}_{\mathrm{plus}}$ and subtracting $\mathrm{TN}_{\mathrm{minus}}$. The function $\mathrm{TN}_{\mathrm{plus}}$ has a power-law component that models high depth of coverage and a binomial component that models low depth of coverage. The formula for $\mathrm{TN}_{\mathrm{plus}}^{\mathrm{PowerLaw}}$ is defined as follows.

$$\mathrm{TN}_{\mathrm{plus}}^{\mathrm{PowerLaw}}(t, n) = 3 \times 10 \times \log_{10}(\min(t/n, 2)) \tag{30}$$

where $t$ denotes the tumor variant allele fraction (VAF) and $n$ denotes the normal VAF. The number 2 in Eq. (30) implies that $\frac{t}{n}$ is effectively capped at $3 \times 10 \times \log_{10}(2)$. The number 2 can be heuristically justified with the following extension of Empirical Law 1: given a sufficient number of observations, if the number of observations is doubled and the additional observations provide additional evidence for the absence of any artifact, then the probability that all observations are generated by some artifact becomes one-eighth of such probability before the doubling. In fact, we observed that approximately 80% of high-quality false positive variant calls in HG001 are also found in HG002 (data not shown) in the Illumina HiSeq 300x dataset. However, HG001 and HG002 were derived from two different genomes of two different persons, and some NGS errors are genome-specific. In practice, the tumor and matched-normal samples are from one single genome of the same patient, so more than 80% false positive calls should be shared between the tumor sample and the matched normal sample. Thus, we estimate that the tumor variant quality increases by at most $\frac{10}{\log(10)} \times \log(2^3)$ given a matched normal sample to compare with, which results in the number 2 in Eq. (30).

The formula for $\mathrm{TN}_{\mathrm{plus}}^{\mathrm{Binomial}}$ is defined in terms of information gain as follows.

$$\mathrm{TN}_{\mathrm{plus}}^{\mathrm{Binomial}}(a_1, a_2, b_1, b_2) = \begin{cases} \frac{10}{\log(10)} \times \left( b_2 \times \mathrm{D}_{\mathrm{KL}}^{\mathrm{Bernoulli}} \left( \frac{b_1}{b_2}, \frac{a_1}{a_2} \right) \right) & \text{if } \frac{a_1}{a_2} > \frac{b_1}{b_2} \\ 0 & \text{otherwise} \end{cases} \tag{31}$$

where $a_1$, $a_2$, $b_1$, and $b_2$ are the tumor allele depth, tumor total depth, normal allele depth, and normal total depth, respectively. The function $\mathrm{TN}_{\mathrm{plus}}$ then outputs the minimum of $\mathrm{TN}_{\mathrm{plus}}^{\mathrm{PowerLaw}}$ and $\mathrm{TN}_{\mathrm{plus}}^{\mathrm{Binomial}}$ as the final reward to variant quality by comparing the tumor with its matched normal. The function $\mathrm{TN}_{\mathrm{minus}}$ basically subtracts the normal-sample variant quality $\mathrm{AnyAltQ}^{\mathrm{N}}$ from the tumor-sample variant quality $\mathrm{AnyAltQ}^{\mathrm{T}}$, and such subtraction is adjusted according to the properties of systematic error. The function $\mathrm{TN}_{\mathrm{minus}}$ is defined as follows.

$$\mathrm{TN}_{\mathrm{minus}} = \max\left(0, \mathrm{AnyAltQ}^{\mathrm{N}} - \rho \times \frac{a_1/a_2}{b_1/b_2}\right) \tag{32}$$

where $a_1$, $a_2$, $b_1$, and $b_2$ are the tumor allele depth, tumor total depth, normal allele depth, and normal total depth, respectively. Higher $\rho$ results in higher decrease in likelihood if the tumor allele fraction deviates from the normal allele fraction. By default, we let $\rho = 15$ which is estimated from the variability in the observation of copy numbers[15].

As a reminder, $\mathrm{AnyAltQ}^{\mathrm{T}}$ and $\mathrm{AnyAltQ}^{\mathrm{N}}$ are the Phred-scaled tumor and normal false positive odds ratios and are computed by using only the tumor and normal BAM files, respectively. Given the matched normal BAM file, $\mathrm{AnyAltQ}^{\mathrm{T}}$ is adjusted to generated the tumor log-odds TLOD as follows.

$$10 \times \mathrm{TLOD} = \mathrm{AnyAltQ}^{\mathrm{T}} + \min(\mathrm{TN}_{\mathrm{plus}}^{\mathrm{Binomial}}, \mathrm{TN}_{\mathrm{plus}}^{\mathrm{PowerLaw}}) - \mathrm{TN}_{\mathrm{minus}} \tag{33}$$

As a reminder, the Phred-scaled odds ratio that a variant candidate is homozygous reference in the normal BAM file is $\mathrm{HomRefQ}^{\mathrm{N}}$, which is defined as $(\mathrm{GL}_{0/0} - \max(\mathrm{GL}_{0/1}, \mathrm{GL}_{1/1}, \mathrm{GL}_{1/2}))$ in Eq. (27). Given the tumor BAM file matched with the normal BAM file, the normal log-odds NLOD is defined as follows.

$$10 \times \mathrm{NLOD} = \mathrm{HomRefQ}^{\mathrm{N}} + \min(\mathrm{TN}_{\mathrm{plus}}^{\mathrm{Binomial}}, \mathrm{TN}_{\mathrm{plus}}^{\mathrm{PowerLaw}}) \tag{34}$$

Our TLOD and NLOD have the same interpretations as the ones presented by Cibulskis et al. [3].

#### 1.7 Variant calls: generalization from single-nucleotide variants (SNVs) to insertions-deletions (InDels)

So far, our method seems to be applicable to only SNVs. Nevertheless, InDels can be treated in the same way as SNVs. This subsection describes the differences between calling SNVs and calling InDels.

First, each genomic position is split into two sub-positions. One sub-position has the statistics of each nucleotide class among {A,C,G,T}, and the other sub-position has the statistics of each class of InDel gap. InDel gaps are grouped into the following seven classes by the short tandem repeat (STR) pattern of the inserted or deleted sequence.

1. neither insertion nor deletion
2. insertion of one STR unit
3. insertion of two STR units
4. insertion of three or more STR units or insertion of a non-STR sequence
5. deletion of one STR unit
6. deletion of two STR units
7. deletion of three or more STR units or deletion of a non-STR sequence

The occurrence of each InDel observed in sequencing data is assigned to one of the seven classes mentioned above. Then, each class of InDel is simply treated as a class of SNV.

The Illumina sequencing platform generates, in each read, basecall quality scores for mismatches but not for InDels. In addition, unlike mismatches, the number of possible InDels in a read is theoretically infinite because any sequence can be inserted at any position. UVC estimates the basecall-like quality of an InDel in a read with the following formula.

$$\text{InDelQual}(x, y, a, b, c) = \min\left(x + z, y + z, \text{STRQual}(a, b)\right) + 10 \times \log_{10}(c) \tag{35}$$

where

$$\text{STRQual}(a, b) = 44 - 10 \times \log_{10}\left(1 + \frac{8 \times \text{softplus}\left(a \times b - 8\right)}{\theta \times a^2}\right) \tag{36}$$

$$\text{softplus}(v) = \log(1 + \exp(v)) \tag{37}$$

and where $x$ and $y$ are respectively the two qualities of the two bases that are immediately before and after the InDel, $a$ is the length of each repeating unit in the STR region, $b$ is the number of repeating units in the STR region, $c$ is the number of repeating units that are inserted or deleted, and $z = 10$. If the InDel is not in an STR region, then $a = b = 1$.

Equation (36) is formulated according to the profile of PCR errors estimated by Shinde et al. [18]. In Eq. (36), 8 is approximately the number of bases in the reactive site of a typical PCR polymerase such as *Taq* [18], and 44 is approximately the InDel error rate of the Illumina sequencing platform [7].

The value of $\theta$ in Eq. (36) is also assigned according to the statistics provided by Shinde et al. [18]: if the InDel is the deletion of one STR unit, then $\theta = \frac{1}{5}$; otherwise, $\theta = 1$.

The treatment of the twice-sequenced class of InDel in the overlap between the R1 and R2 ends is similar to that of SNV in Eq. (1). However, for InDels, $\text{R1R2overlapQ}(q_1, q_2) = \max(q_1, q_2)$ regardless of whether $b_1 = b_2$. The reason is that the read end with the lower quality of InDel class is often mis-aligned, and mis-alignment effectively results in a quality of zero for its corresponding InDel class.

If all InDels in an input BAM file are left-aligned, then the expansion/contraction of a repeating unit occurring anywhere in an STR track would always result in an insertion/deletion of the repeating unit at the beginning of the STR track, respectively. Hence, the error probability of an STR expansion/contraction is positively correlated with the length of the STR track. Moreover, longer InDels result in bigger sequence changes and therefore are less likely to be erroneously generated from their corresponding reference alleles. Hence, the error probability of an InDel is negatively correlated with the size of the InDel. To incorporate these two correlations, the following Bonferri-like correction in the probability space is applied to the method in Section 1.3 for an InDel of $n$ bases that is an expansion or contraction found within an STR track of $L$ bases.

$$p' = p \times \frac{L}{n} \quad \text{or equivalently} \quad q' = q + 10 \times \log_{10}\left(\frac{n}{L}\right) \tag{38}$$

where $p$ and $p'$ are respectively the new and old raw error probabilities, and where $q$ and $q'$ are respectively the new and old Phred-scaled error probabilities. If the InDel is not within an STR track, then $L = 1$.

For InDels, the filter threshold of 25 for base qualities (BQs) mentioned in Section 1.5 is not applied before computing the UMI-derived BQs. In addition, the Illumina sequencers do not generate any basecall qualities for InDels, so the UMI-derived BQs are especially useful for calling InDels.

For InDels, the empirical Phred-scaled prior probabilities for the genotypes `0/0`, `0/1`, `1/1`, and `1/2` are observed to be 0, -40, -42, and -49, respectively, which are different than such probabilities for SNVs listed in Eq. (26).

#### 1.8 Variant calls: incorporation of systematic error

Systematic error is known to exist in NGS. Each source of systematic error offers a plausible explanation for the observed signals along with a Phred-scaled error probability that the explanation is correct. Thus, the source of systematic error with the lowest Phred value of error probability is used to impose a maximum to the final variant quality. Here, we describe the incorporation of three sources of systematic errors: basecalling error, mapping error, and tumor-in-normal (TiN) contamination.

##### 1.8.1 Systematic basecalling error

Meacham et al.[13] mentioned that, for the Illumina sequencing platform, the probability that a genomic position is subject to systematic basecalling error is approximately $\frac{1}{1000}$ which is equivalently to 30 in Phred scale. If a genomic position is subject to systematic basecalling error, then the sequencing machine should generate basecalls with low qualities. To eliminate as many false positive variant calls as possible, we assume the following worst-case scenario: at a genomic position with systematic basecalling error, the sequencer generates a random base quality between 0 and 37, where 37 is the maximum that can ge generated by the NovaSeq™ 6000 System. In this worse-case scenario, the root-mean-square base quality (RMS-BQ) is 20 when rounded down. Hence, the RMS-BQ is first subtracted by 20 and then multiplied by the number of bases called, resulting in the following formula to compute the maximum variant quality imposed by systematic basecalling error.

$$\text{SysErrBQ} = b \times (\text{RMS-BQ} - 20) + 30 \tag{39}$$

where $b$ is the number of bases called.

##### 1.8.2 Systematic mapping error

An alignment record contains information specifying the genome location to which the read is mapped. If a read is mapped to a different genome location which is not the one specified in the alignment record, then the read is characterized by mapping error. Mapping error can be caused by incomplete reference genome, alternative reference genome, rare structural variation, etc. To let variant-call quality account for mapping error, UVC uses the following heuristic formula to compute the maximum variant quality imposed by systematic mapping error.

$$\text{SysErrMQ} = 60 - 20 + \frac{\text{MQ}^{\text{ALT}}}{3} + (\text{MQ}^{\text{ALT}} - \text{MQ}^{\text{REF}}) - \text{GL}^{\text{Prior}}_{1/1} + 10 \times \log_{10}(f) \tag{40}$$

In the above formula, $\text{MQ}^{\text{REF}}$ and $\text{MQ}^{\text{ALT}}$ are the root-mean-squared (RMS) mapping qualities (MQs) (RMS-MQs) of the REF and ALT alleles, respectively, and $f$ is the allele fraction of the variant of interest. The additional numbers used in Eq. (40) can be justified as follows.

1. The number 60 is equal to the maximum mapping quality at which $\text{MQ}^{\text{REF}}$ and $\text{MQ}^{\text{ALT}}$ are capped. The number 60 is equal to the highest mapping quality that can be generated by BWA[8].

2. The number 20 is approximately equal to the probability that an alignment has zero mapping quality. The number 20 is obtained according to our empirical observation.

3. $\text{GL}^{\text{Prior}}_{1/1}$, which was previously defined in Eq. (26), is the Phred-scaled probability that a homozygous ALT germline variant occurs at a position. As previously mentioned, $\text{GL}^{\text{Prior}}_{1/1}$ is set to 33 for SNVs, 42 for InDels, and 0 for all reference alleles.

##### 1.8.3 Systematic tumor-in-normal contamination

Taylor-Weiner et al.[20] showed that, even for liquid tumor which is known to be heavily affected by tumor-in-normal (TiN) contamination, about 75% of the tumor-matched normal samples are only affected by TiN contamination rates of less than 4%. Thus, UVC uses by default a TiN contamination rate ($\eta$) of 5%, which is slighly higher than 4%, to exclude background noise in the observed variant signal in the tumor-matched normal samples. If there is strong evidence for very heavy TiN contamination, then we can increase the value of $\eta$ accordingly.

By incorporating TiN contamination, the formula for LowDepthQ2 in Eq. (3) is modified by capping base quality for the normal. Then, for the normal, Eq. (3) is modified as follows to be aware of the contamination from the tumor.

$$\text{LowDepthQ2}(a, B) = |B| \times \left(\min(B, f^T \times \eta) + 10 \times \log_{10}\left(\frac{|B|}{a}\right)\right) \quad (41)$$

where $f^T$ is the allele fraction in the tumor, $\eta$ is the aforementioned TiN contamination rate, and where $\min(B, f^T \times \eta) = \min(\min(B), f^T \times \eta)$. In brief, the expression $\min(B)$ in Eq. (3) is transformed into $\min(B, f^T \times \eta)$ in Eq. (41) to account for TiN contamination.

##### 1.9 Variant calls: generalization from Illumina to other sequencing platforms

We empirically observed that the technology of sequencing by synthesis used by Illumina and the technology of DNA nanoball sequencing used by BGI share the same error profile. Thus, everything that is applicable to the Illumina sequencing platform is also applicable to the BGI sequencing platform, without any modification whatsoever. Nevertheless, the technology of semiconductor sequencing used by IonTorrent/Life Technologies/Thermo Fisher Scientific is quite different: basecall qualities (BQs) generated by the IonTorrent sequencing platform denote probabilities of generating erroneous InDels at the corresponding base positions. Thus, for IonTorrent sequencing data, the Illumina-like nucleotide-substitution BQ is estimated to be the IonTorrent raw BQ plus 8, and the Illumina-like BQ is used in Eq. (41). Moreover, raw IonTorrent BQs directly estimate InDel sequencing-error rates, so the constant $z$ in Eq. (35) is set to zero for the IonTorrent platform. In addition, we found that the Illumina systematic basecalling error is not applicable to IonTorrent, but we did not find a well tested model for the IonTorrent systematic basecalling error. Thus, Eq. (39) is simply not applied to the IonTorrent platform.

[17] Johnny Sena. Unique Molecular Identifiers reveal a novel sequencing artefact with implications for RNA-Seq based gene expression analysis. *Scientific Reports*, 8, 09 2018.
