## Supplemental results for "Calling small variants with universality and Bayesian-frequentist hybridism"

### 1 Supplementary results

We evaluated the performance of UVC using the metrics F-score and PrAUC F-score denotes the maximum harmonic mean between precision and recall. PrAUC, which is equivalent to average precision, denotes the area under the curve (AUC) of precision-versus-recall.

In our evaluation, we tried to run the latest release of each software, with an allocation of 32 logical CPU cores, on a Red Hat 4.8.5-28 server with 128 GB of RAM and 28/56 physical/logical CPU cores (Intel(R) Xeon(R) CPU E5-2680 v4 @ 2.40GHz). BWA MEM version 0.7.17 was used to align reads to the human genome in the binary alignment/map (BAM) format if the reads were in the FASTQ format. The software packages UVC version 0.6.0.441a694, GATK version 4.1.9.0, Strelka2 version 2.9.10, VarScan version 2.4.2, LoFreq version 2.1.5, LoLoPicker commit de725bd, freebayes version 1.3.2, and bcftools version 1.11 were used to preprocess the BAM files or call variants from the BAM files. For GATK, deduplication and base recalibration were performed, which is recommended by the best practice for using GATK. For LoFreq, InDel-quality estimation by Dindel was performed on the deduplicated and base-calibrated BAM files, which is recommended by the best practice for using LoFreq. The other variant callers all used the raw BAM files, which were directly generated by BWA MEM or NovoAlign, as their inputs.

Unless stated otherwise, we ran each variant caller with its default command-line parameters, and we used the vcfeval utility from the RTG tools version 3.11.1 to perform our evaluation (`https://github.com/RealTimeGenomics/rtg-tools`). The vcfeval utility considers representational difference in the VCF format: if the application of two or more variants on the same genome would result in the same remaining sequence, then these variants are considered to be equivalent even if these variants are represented in different forms. The same NCBI dbSNP reference SNP (rs) number is assigned to variants of the same type at the same position, and vice versa[5]. Hence, if the ground-truth VCF for assessing variant calls is not provided, then we used the command `bcftools isec -c both` to create such ground-truth VCF, so SNPs that are at the same position are considered to be identical, and InDels that are at the same position are considered to be identical. VCF files generated by FilterMutectCall (as part of the recommended workflow of GATK4 Mutect2) and Varscan2 may have hard filters applied in the FILTER column. Hence, we evaluated each of these two callers with two filtering strategies: without applying any hard filter and with all hard filters applied. Unless stated otherwise, the NISTv3.3.2 versions of high-confidence BED files and VCF files at `ftp.ncbi.nlm.nih.gov/giab/ftp/release/` were used as the germline ground truth when applicable.

Additionally, samtools version 1.11 and bedtools version v2.29.2 were used for some intermediate steps in data processing (e.g., compression of SAM files into BAM files, generation of the intersection between two BED files of high-confidence regions, etc).

#### 1.1 Supplemental evaluation results on a WGS dataset to validate the power-law universality

To validate the power-law universality in NGS, we examine two sequencing runs.

1. The HiSeq 300x BAM file at `https://ftp.ncbi.nlm.nih.gov/giab/ftp/data/NA12878/NIST_NA12878_HG001_HiSeq_300x/NHGRI_Illumina300X_novoalign_bams/` (generated by the whole-genome sequencing (WGS) of HG001) was directly downloaded and used. For this WGS BAM file, a position is used to assess the NGS universality if and only if this position is covered by a sequencing depth of at least 200x.

2. The NextSeq FASTQ files with accession SRR7526729 (generated by targeted amplicon sequencing (TAS)) were aligned with BWA MEM to the human reference genome hs37d5 to generate a bam file. For this TAS BAM file, a position is used to assess the NGS universality if and only if this position is covered by a sequencing depth of at least 2000x.

In addition of evaluating only within the high-confidence BED regions, we excluded genomic positions that are within 8 bases of each variant in the truth VCF file to prevent true positive calls from being labeled as false positive due to potentially different representations of the same variant. For the TAS BAM file, the intersection of HG001 and HG002 high-confidence BED regions and the union of HG001 and HG002 truth variants were used. We used the command `bcftools mpileup -a AD,ADF,ADR,DP,SP -i 1000000 --max-idepth 1000000 -f $hs37d5 $bam` to generate the sequencing depth at each genomic position covered by any read.

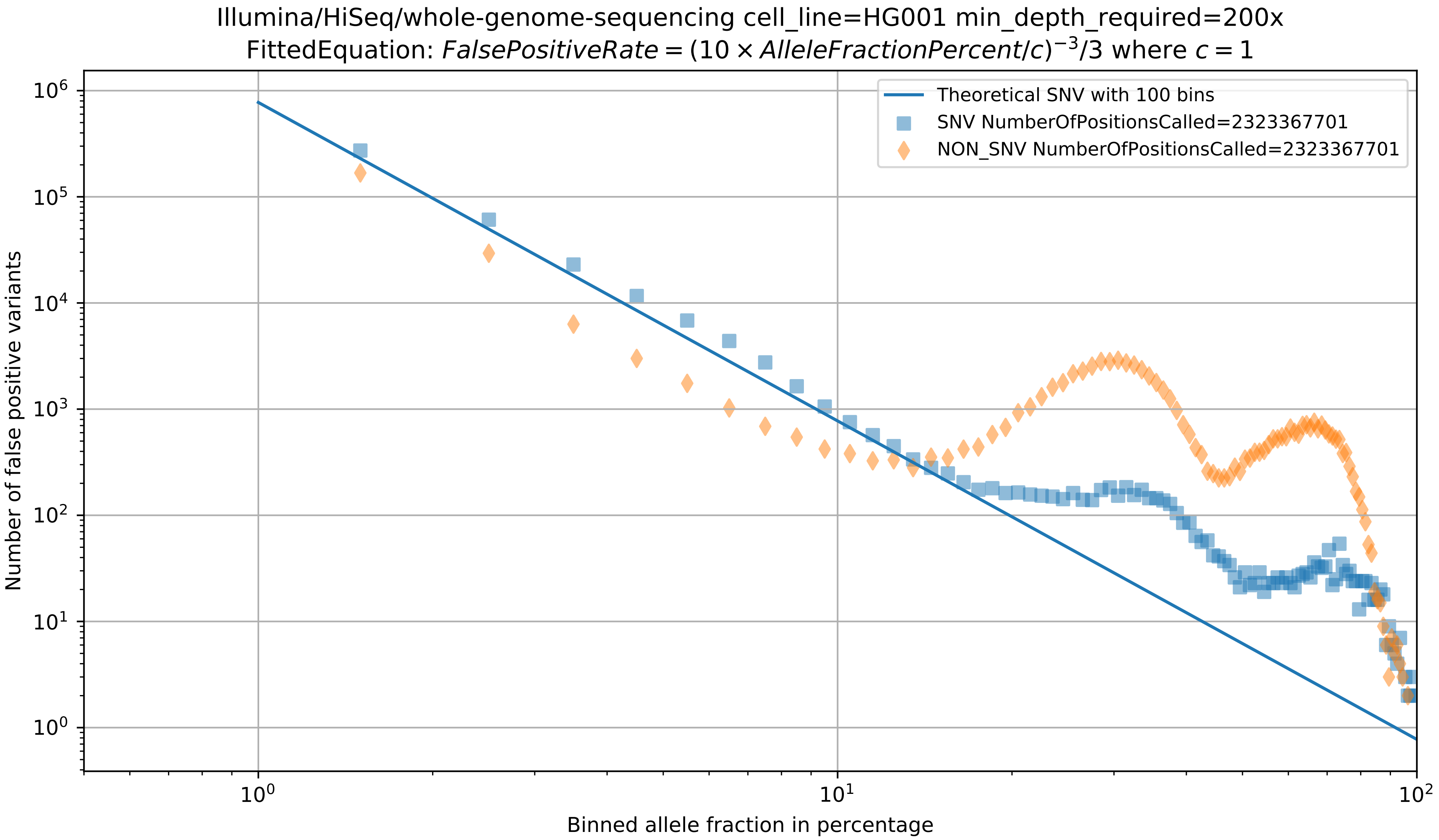

Supplementary Figure 1: Validation of the power-law relationship between the allele fraction of a variant and the probability that the variant is false positive. To generate this figure, the whole-genome-sequencing data generated by HiSeq (URL: `https://ftp.ncbi.nlm.nih.gov/giab/ftp/data/NA12878/NIST_NA12878_HG001_HiSeq_300x/NHGRI_Illumina300X_novoalign_bams/`, with an average sequencing depth of 300x) and pre-aligned by NovoAlign to the hs37d5 human reference genome were directly downloaded. The emergence of SNVs at the allele fractions of around 35% and 70% are presumably caused by mapping error, error in the human reference genome, and/or incomplete truth set of variants. The emergence of InDels at the allele fractions of around 30% and 65% are presumably caused by incomplete truth set of variants: some of these false positive InDels can be real germline InDels that are not known yet (in the truth set, the number of germline InDels increased to 505169 in v3.3.1 from 358753 in v3.3[8], so the current truth set v3.3.2 is still expected to exclude some real germline InDels).

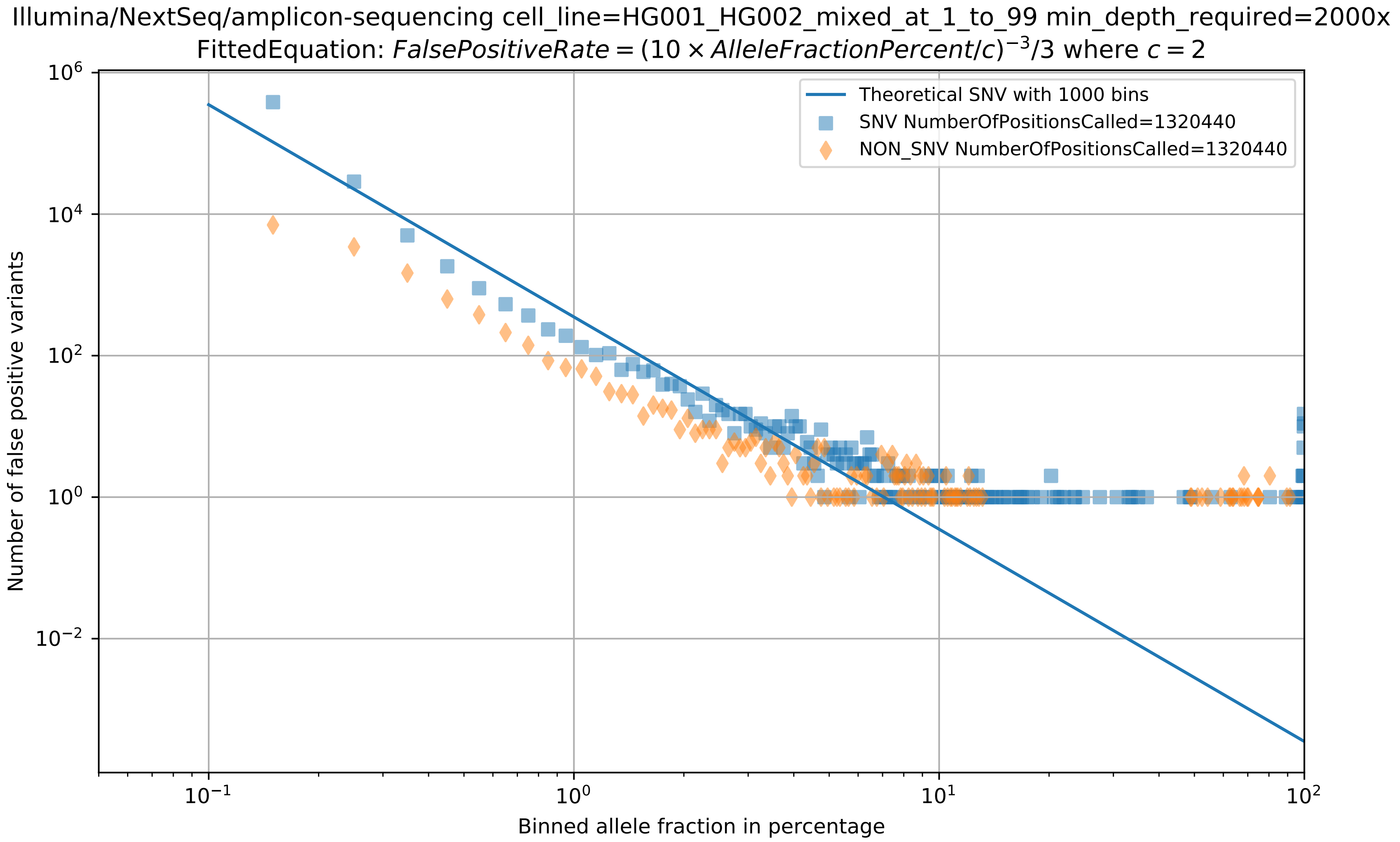

Supplementary Figure 2: Validation of the power-law relationship between the allele fraction of a variant and the probability that the variant is false positive. To generate this figure, the targeted amplicon-sequencing data generated by Illumina NextSeq (accession SRR7526729) were aligned by BWA MEM to the hg19 human reference genome. Here, each targeted region is usually amplified by one single pair of primers, so read-orientation bias and strand bias cannot be detected. Thus, $c = 2$ because the allele fraction would approximately halve if the non-erroneous reads in the other orientation and/or strand were sequenced to the same extent.

#### 1.2 Supplemental evaluation results on WGS datasets for calling germline variants

The HiSeq 300x BAM files at `https://ftp.ncbi.nlm.nih.gov/giab/ftp/data/NA12878/NIST_NA12878_HG001_HiSeq_300x/NHGRI_Illumina300X_novoalign_bams/`, `https://ftp.ncbi.nlm.nih.gov/giab/ftp/data/AshkenazimTrio/HG002_NA24385_son/NIST_HiSeq_HG002_Homogeneity-10953946/NHGRI_Illumina300X_AJtrio_novoalign_bams/`, and `https://ftp.ncbi.nlm.nih.gov/giab/ftp/data/ChineseTrio/HG005_NA24631_son/HG005_NA24631_son_HiSeq_300x/NHGRI_Illumina300X_Chinesetrio_novoalign_bams/` were down-sampled to an average depth of 30x and 60x using samtools.

The MGISEQ 30x FASTQ files (from a total of six runs) at `https://ftp.ncbi.nlm.nih.gov/giab/ftp/data/NA12878/MGISEQ/NA12878_1/`, `https://ftp.ncbi.nlm.nih.gov/giab/ftp/data/AshkenazimTrio/HG002_NA24385_son/MGISEQ/PCR-free/NA24385/`, and `https://ftp.ncbi.nlm.nih.gov/giab/ftp/data/ChineseTrio/HG005_NA24631_son/MGISEQ/PCR-free/NA24631/` were used directly, where each reference sample (HG001, HG002, and HG005) was sequenced in two runs. The first sequencing run/both sequencing runs were used in the evaluation of germline variant calling at 30x/60x average sequencing depths, respectively.

To let UVC generate germline variants, the command-line parameter `-outvar-flag 0x1F` is passed to UVC.

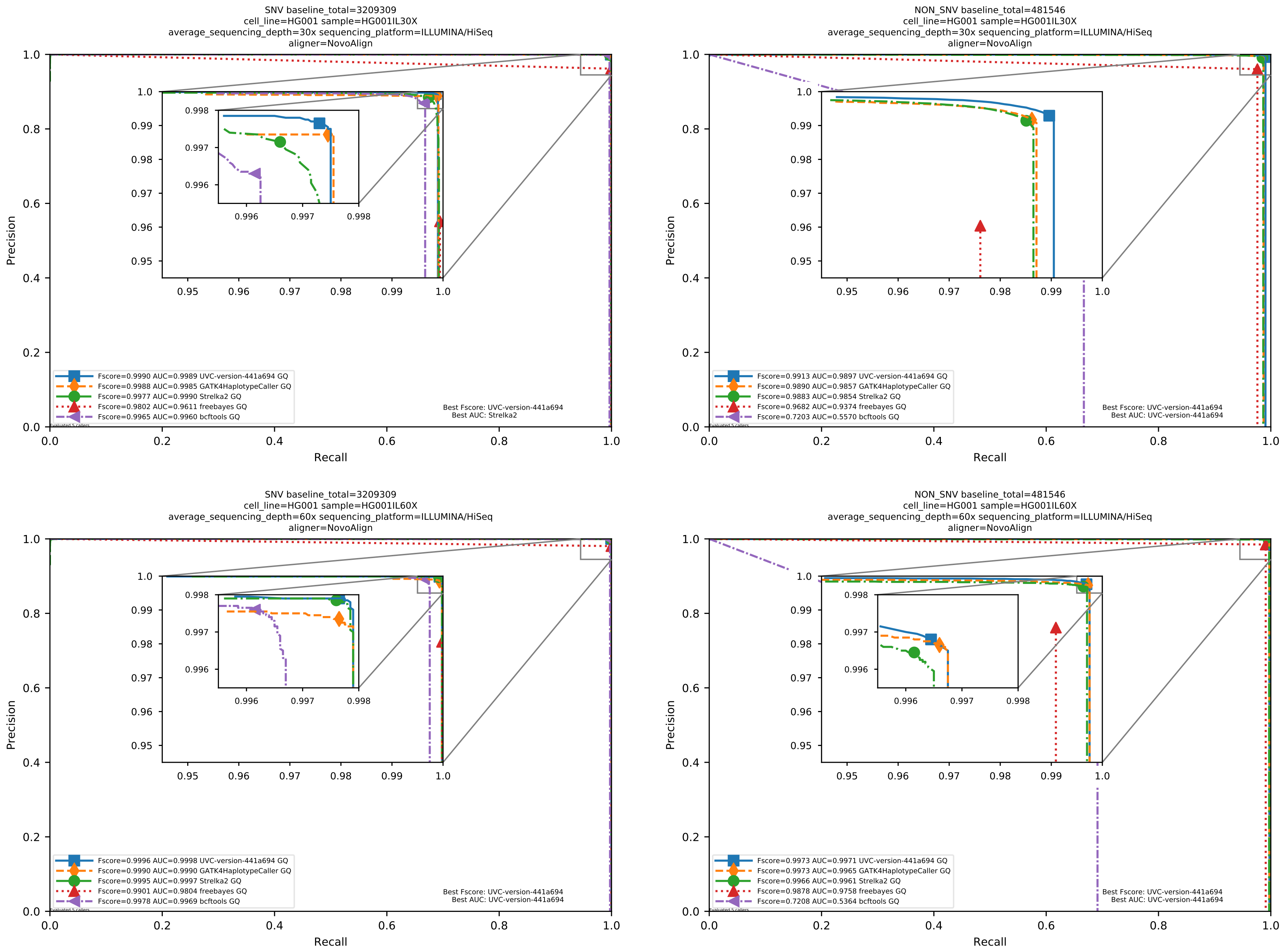

Supplementary Figure 3: Performance of germline variant callers with HiSeq/MGISEQ data pre-aligned/aligned by NovoAlign/BWA MEM to the human reference genome hs37d5.

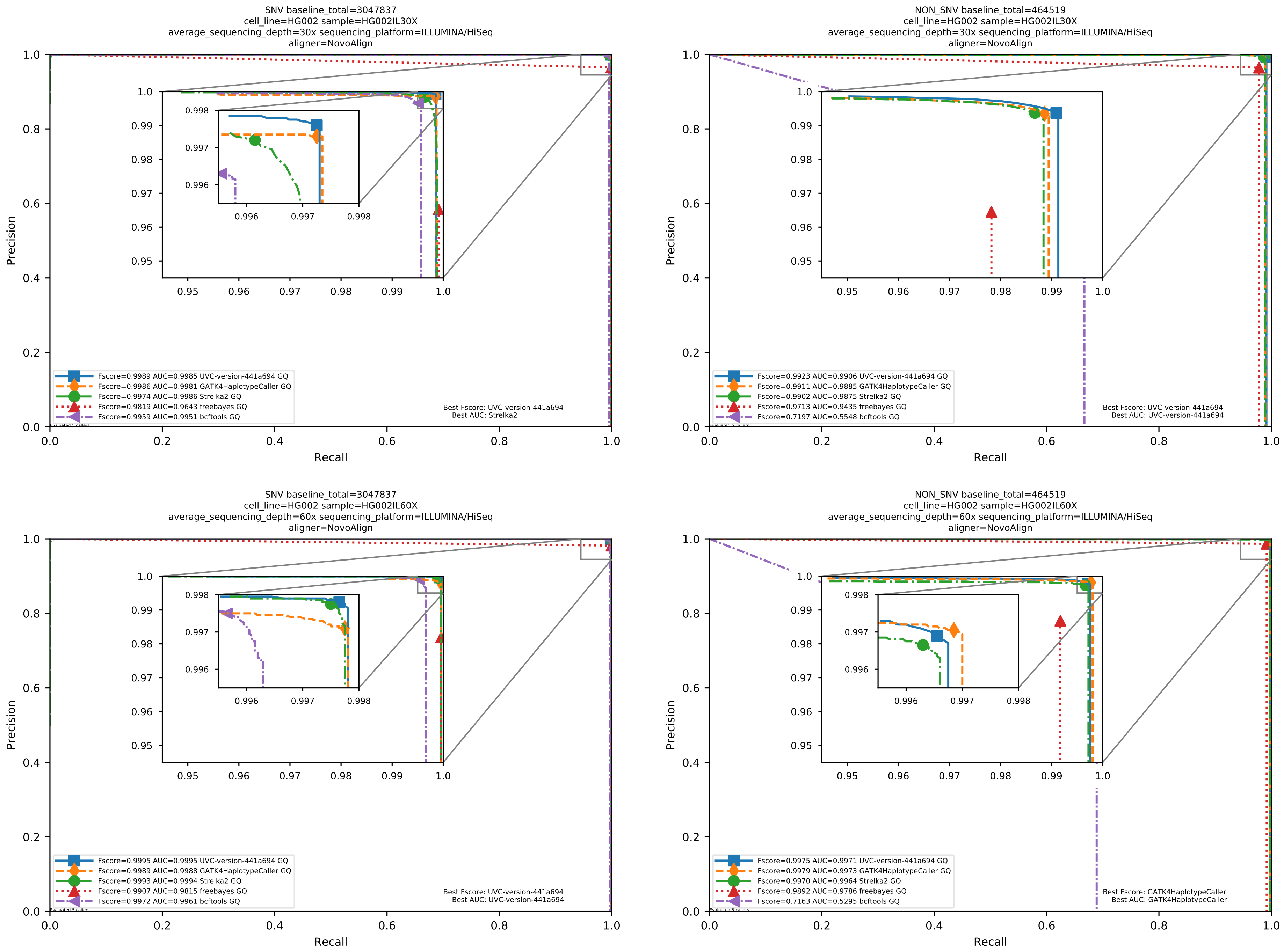

Supplementary Figure 4: Performance of germline variant callers with HiSeq/MGISEQ data pre-aligned/aligned by NovoAlign/BWA MEM to the human reference genome hs37d5.

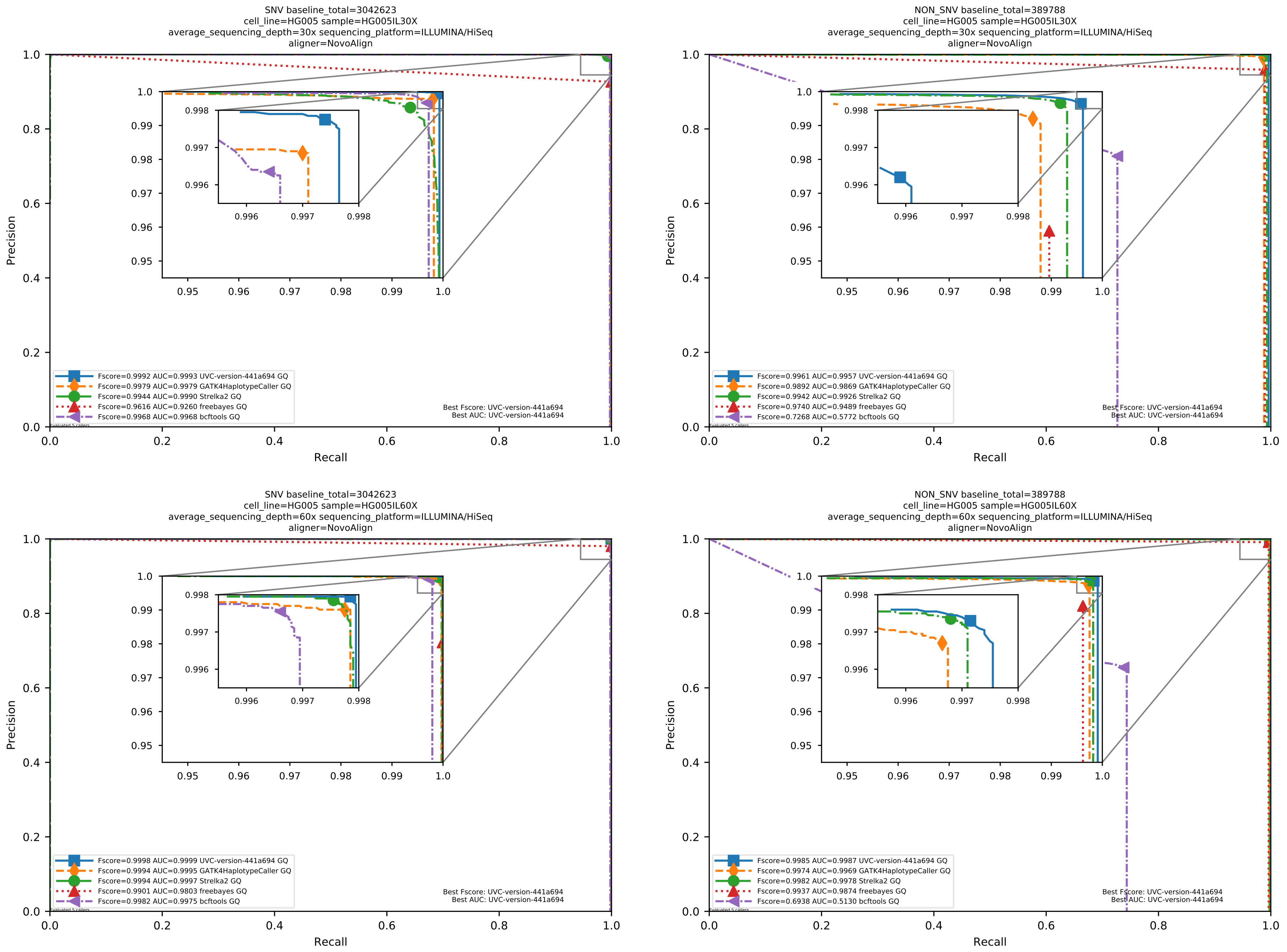

Supplementary Figure 5: Performance of germline variant callers with HiSeq/MGISEQ data pre-aligned/aligned by NovoAlign/BWA MEM to the human reference genome hs37d5.

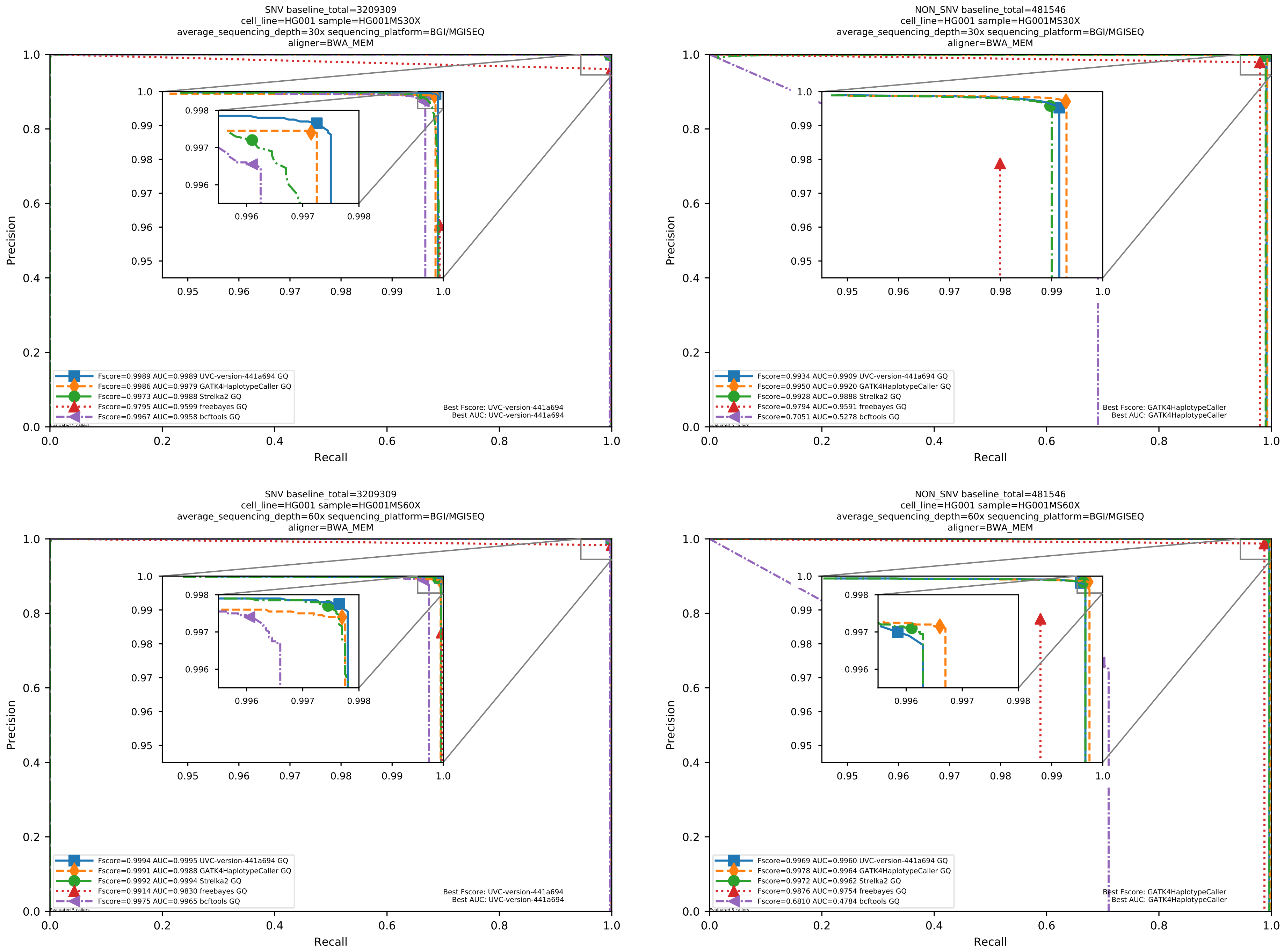

Supplementary Figure 6: Performance of germline variant callers with HiSeq/MGISEQ data pre-aligned/aligned by NovoAlign/BWA MEM to the human reference genome hs37d5.

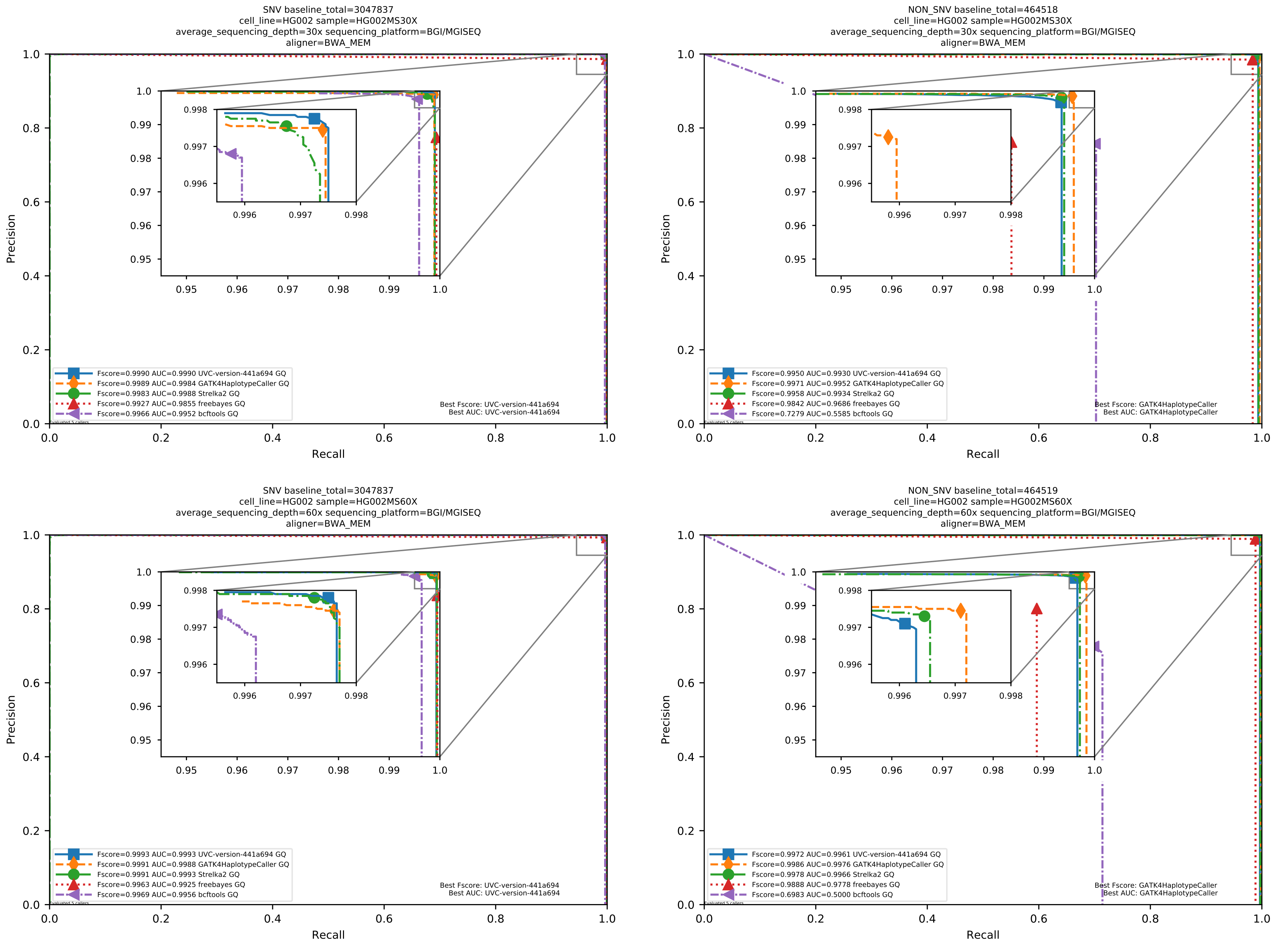

Supplementary Figure 7: Performance of germline variant callers with HiSeq/MGISEQ data pre-aligned/aligned by NovoAlign/BWA MEM to the human reference genome hs37d5.

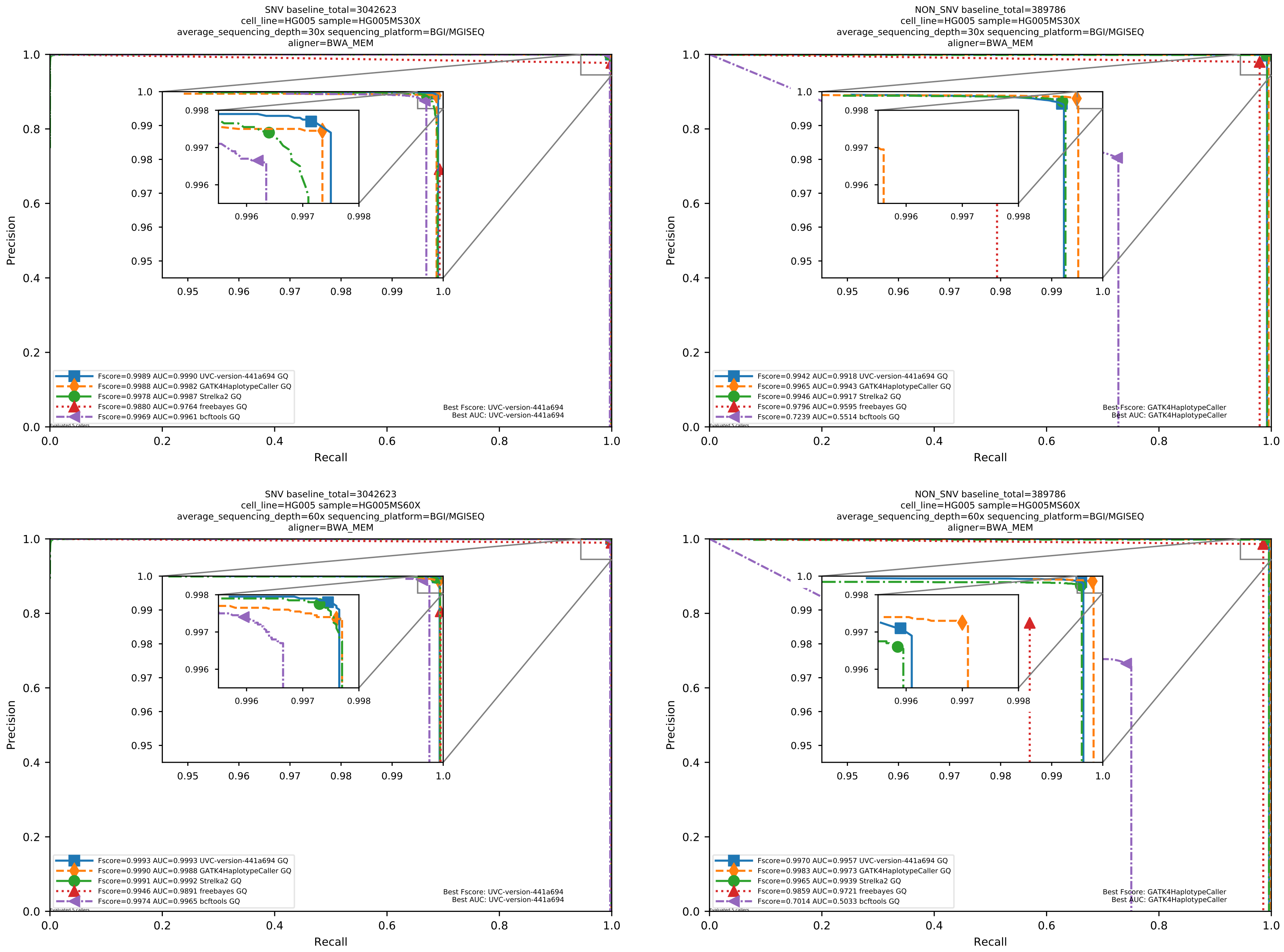

Supplementary Figure 8: Performance of germline variant callers with HiSeq/MGISEQ data pre-aligned/aligned by NovoAlign/BWA MEM to the human reference genome hs37d5.

##### 1.3 Supplemental evaluation results on WGS datasets for calling variants in tumor-only mode, with 64 *in silico* mixtures considering both Illumina and BGI platforms

To generate the Illumina HiSeq *in silico* datasets, we directly used the BAM files pre-aligned by NovoAlign version 3.02.07. More detail about the pre-aligned BAM files is at `https://ftp.ncbi.nlm.nih.gov/giab/ftp/data/AshkenazimTrio/HG002_NA24385_son/NIST_HiSeq_HG002_Homogeneity-10953946/NHGRI_Illumina300X_AJtrio_novoalign_bams/README_NHGRI_Novoalign_bams` and `https://ftp.ncbi.nlm.nih.gov/giab/ftp/data/NA12878/NIST_NA12878_HG001_HiSeq_300x/NHGRI_Illumina300X_novoalign_bams/README`.

To generate the BGI MGISEQ *in silico* datasets, we used the FASTQ files at `https://ftp.ncbi.nlm.nih.gov/giab/ftp/data/NA12878/MGISEQ/NA12878_1/` and `https://ftp.ncbi.nlm.nih.gov/giab/ftp/data/AshkenazimTrio/HG002_NA24385_son/MGISEQ/PCR-free/NA24385/` which were aligned to the hs37d5 human reference genome by BWA MEM.

The raw BAM files used in this subsection were also used to generate the tumor-normal-paired BAM files in Section 1.4. However, samtools cannot directly sub-sample one BAM file into two BAM files which are needed to simulate different tumor purities and normal purities. To perform such sub-sampling, we used the same pseudo-random number generator (PRNG) as the one used by samtools for sub-sampling. In addition, we used the same logic as samtools to sub-sample tumor reads into the tumor BAM and normal reads into the normal BAM: if the integer returned by applying the PRNG on a read name is smaller than the threshold determined by the sub-sampling probability, then the read is kept, an vice versa. To sub-sample tumor reads into the normal BAM (to simulate normal purities) or to sub-sample normal reads into the tumor BAM (to simulate tumor purities), we used the following logic: if the integer returned by applying the PRNG on a read name is bigger than the threshold determined by the sub-sampling probability, then the read is kept, an vice versa. In sum, our sub-sampling method is as similar to the one from samtools as possible.

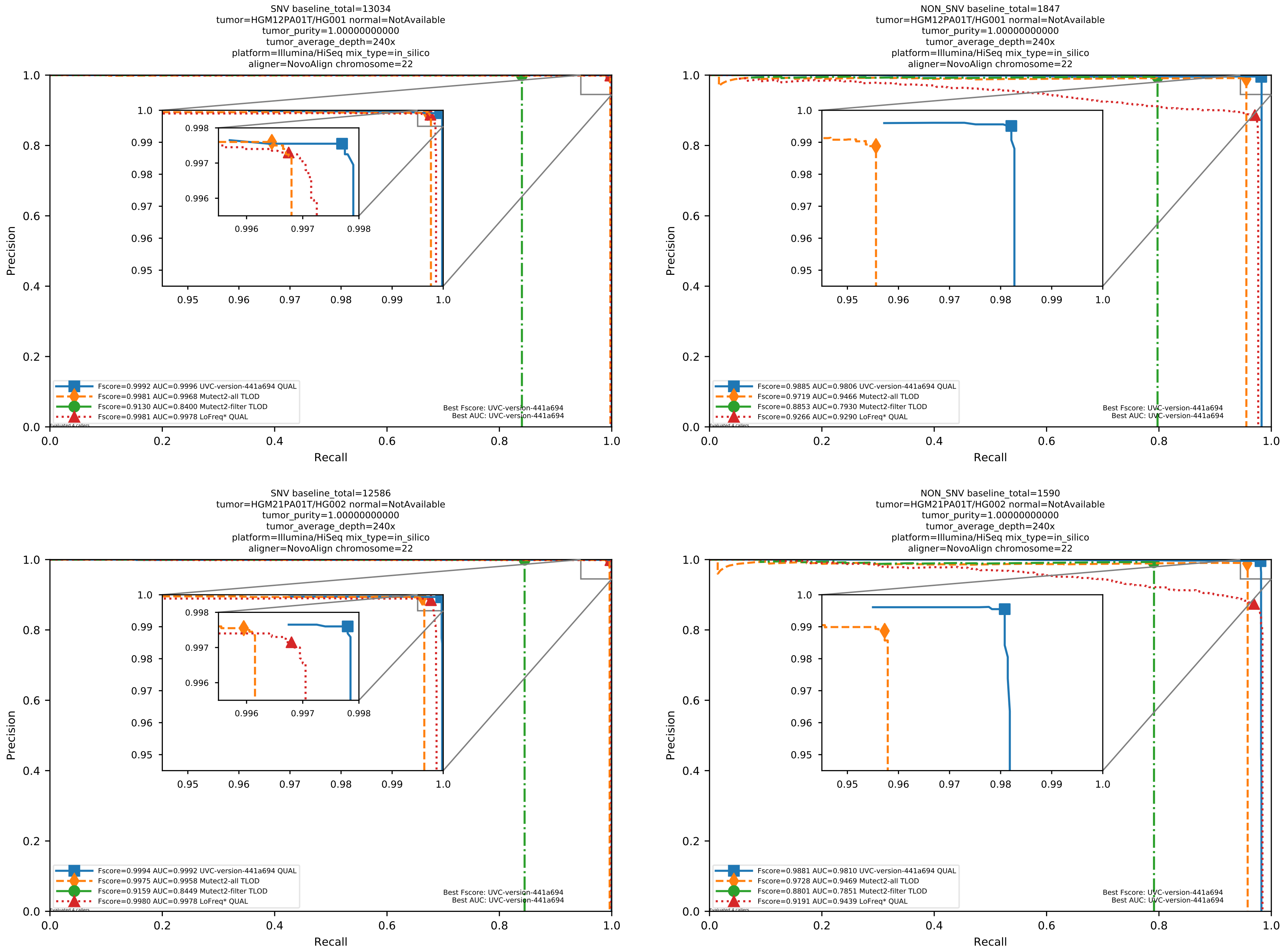

Supplementary Figure 9: Performance of somatic variant callers in tumor-only mode on the *in silico* mixtures with Illumina HiSeq sequencing data pre-aligned with NovoAlign to the human reference genome hs37d5.

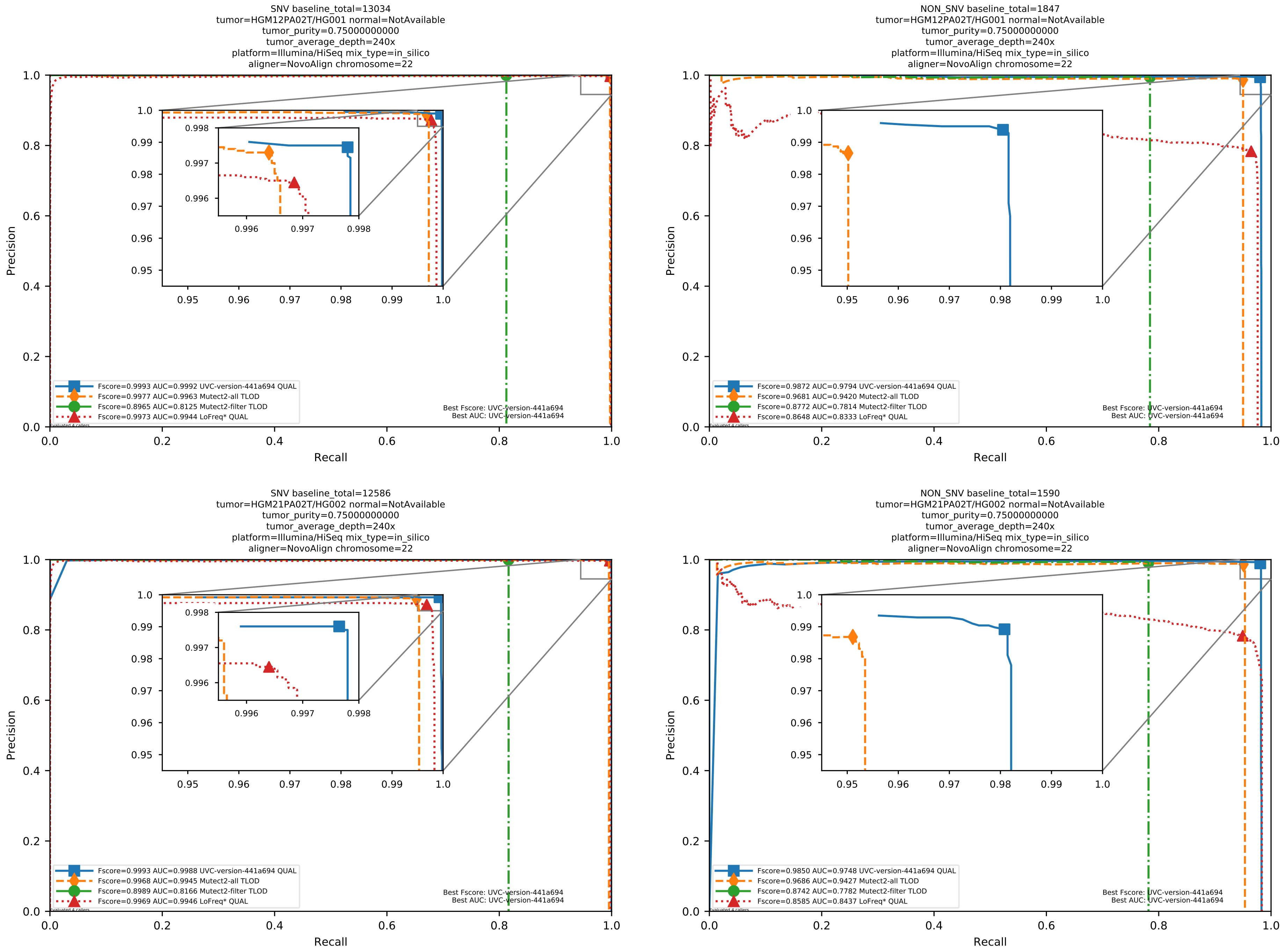

Supplementary Figure 10: Performance of somatic variant callers in tumor-only mode on the *in silico* mixtures with Illumina HiSeq sequencing data pre-aligned with NovoAlign to the human reference genome hs37d5.

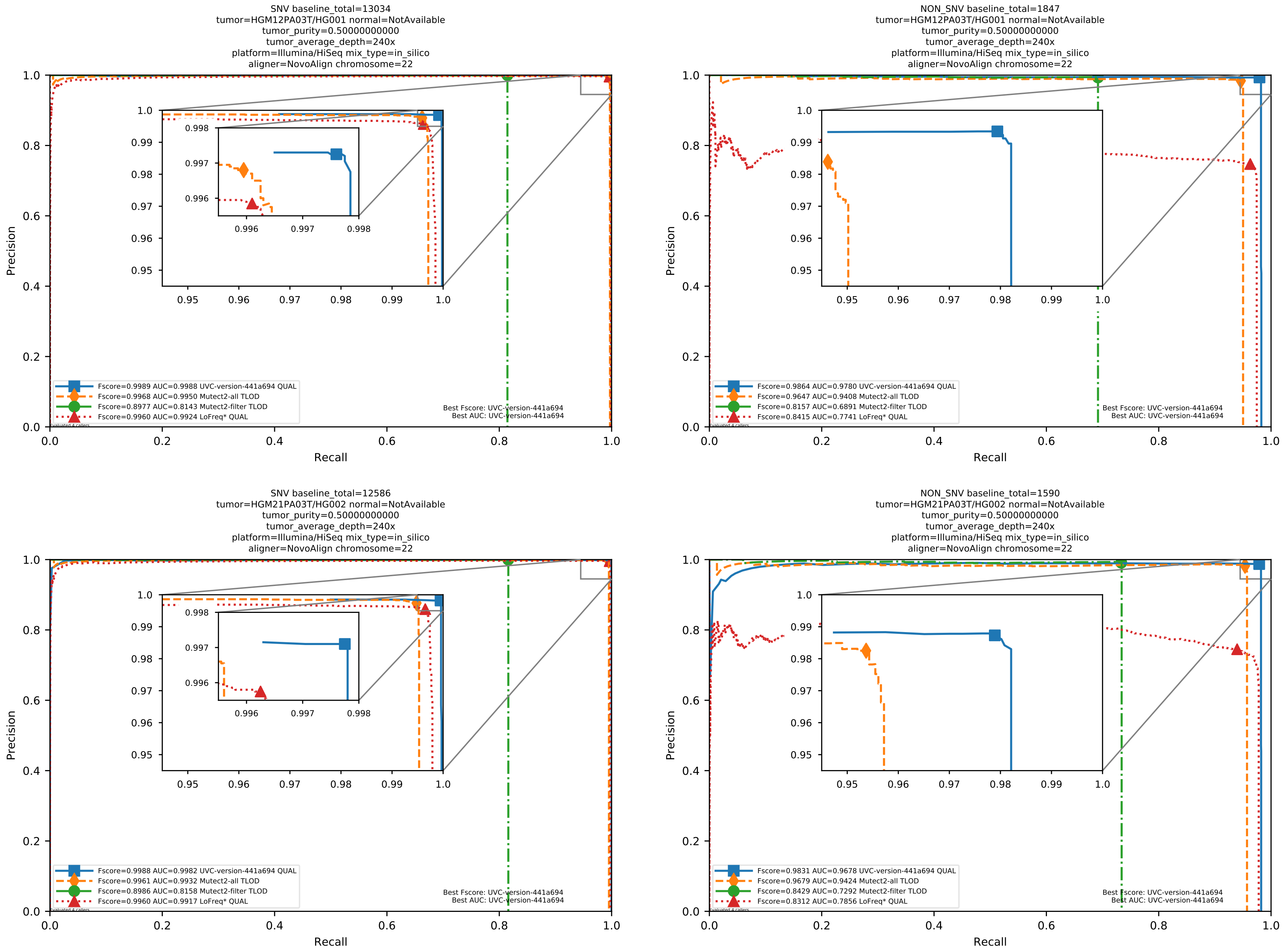

Supplementary Figure 11: Performance of somatic variant callers in tumor-only mode on the *in silico* mixtures with Illumina HiSeq sequencing data pre-aligned with NovoAlign to the human reference genome hs37d5.

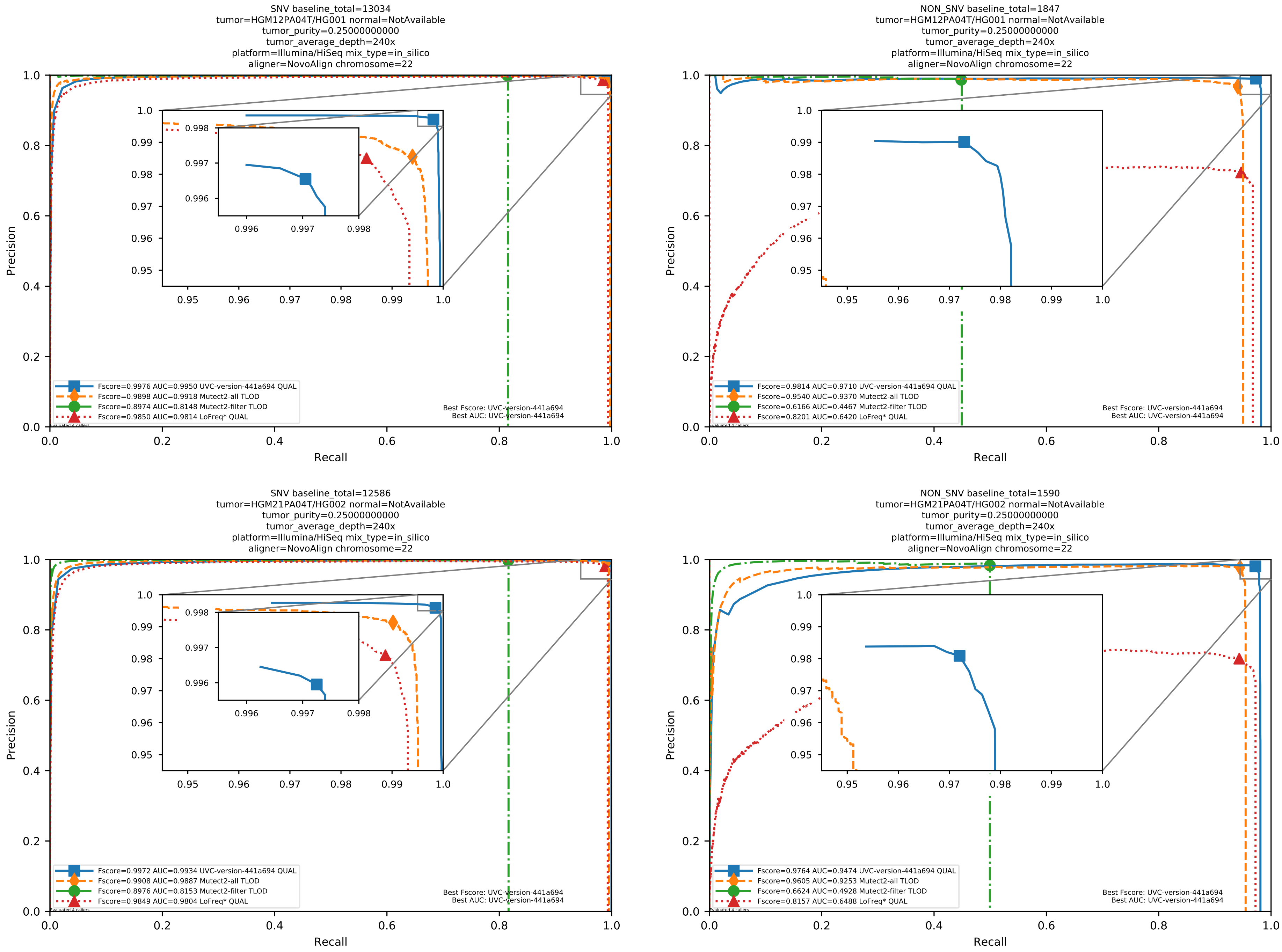

Supplementary Figure 12: Performance of somatic variant callers in tumor-only mode on the *in silico* mixtures with Illumina HiSeq sequencing data pre-aligned with NovoAlign to the human reference genome hs37d5.

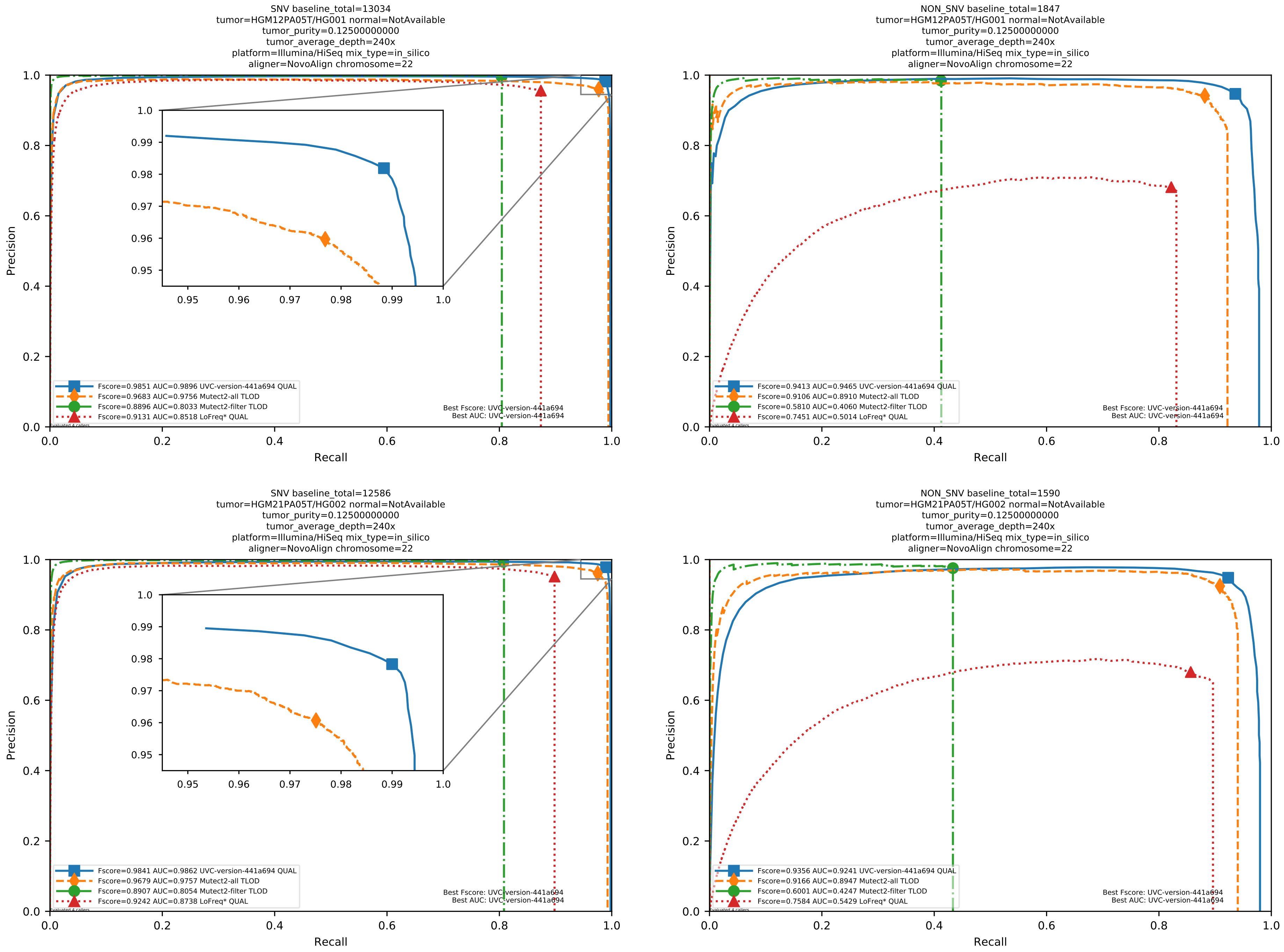

Supplementary Figure 13: Performance of somatic variant callers in tumor-only mode on the *in silico* mixtures with Illumina HiSeq sequencing data pre-aligned with NovoAlign to the human reference genome hs37d5.

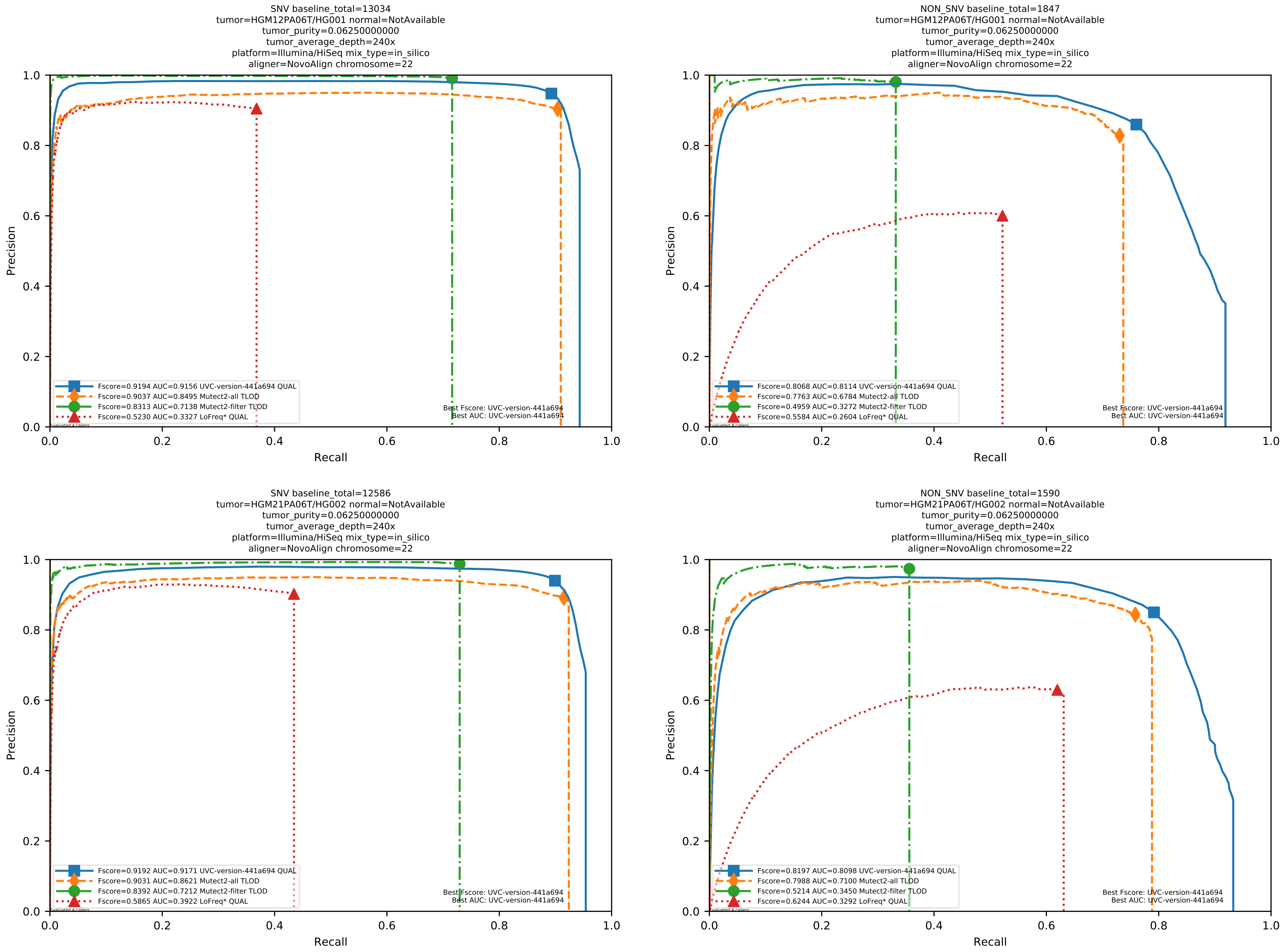

Supplementary Figure 14: Performance of somatic variant callers in tumor-only mode on the *in silico* mixtures with Illumina HiSeq sequencing data pre-aligned with NovoAlign to the human reference genome hs37d5.

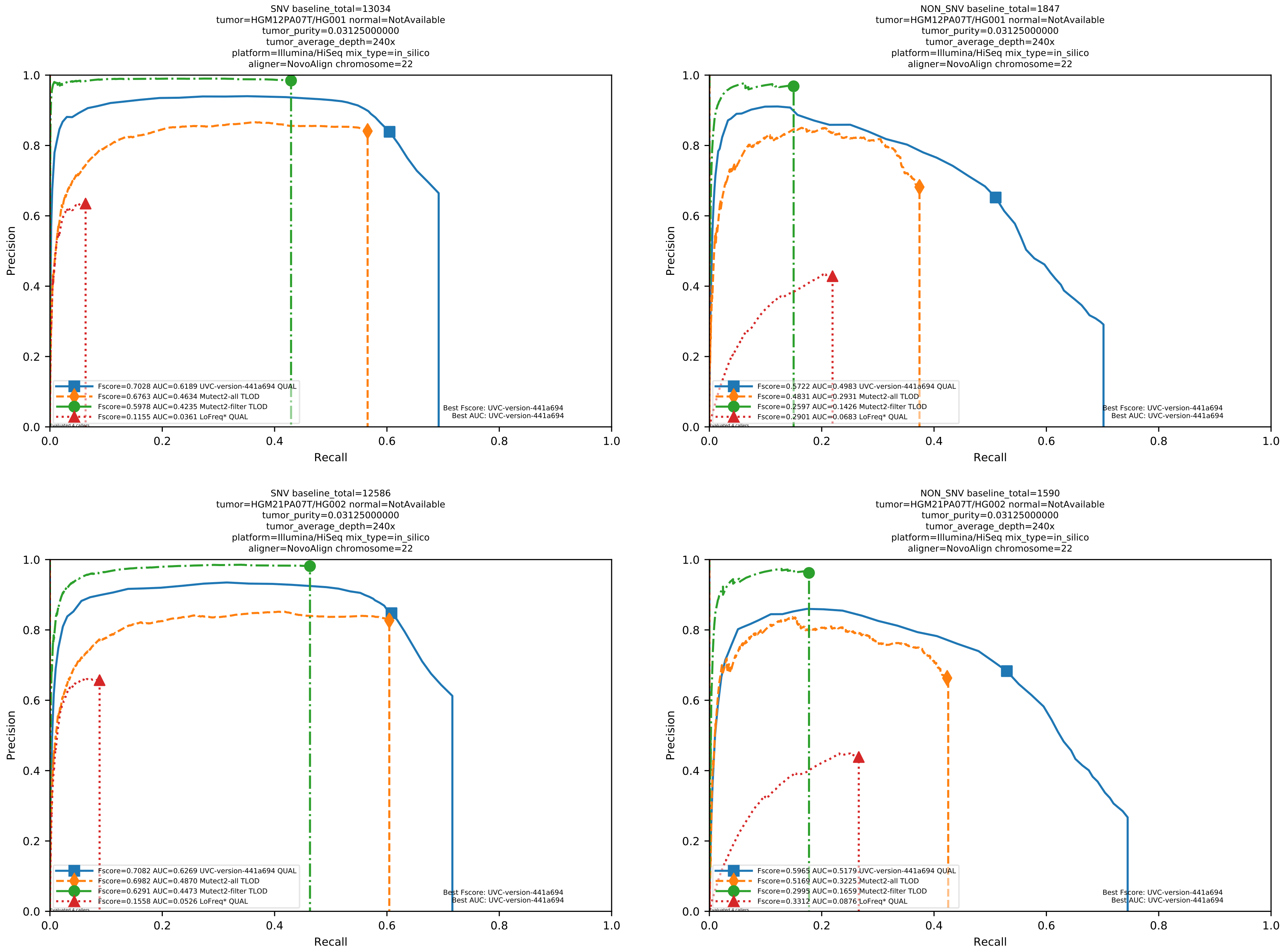

Supplementary Figure 15: Performance of somatic variant callers in tumor-only mode on the *in silico* mixtures with Illumina HiSeq sequencing data pre-aligned with NovoAlign to the human reference genome hs37d5.

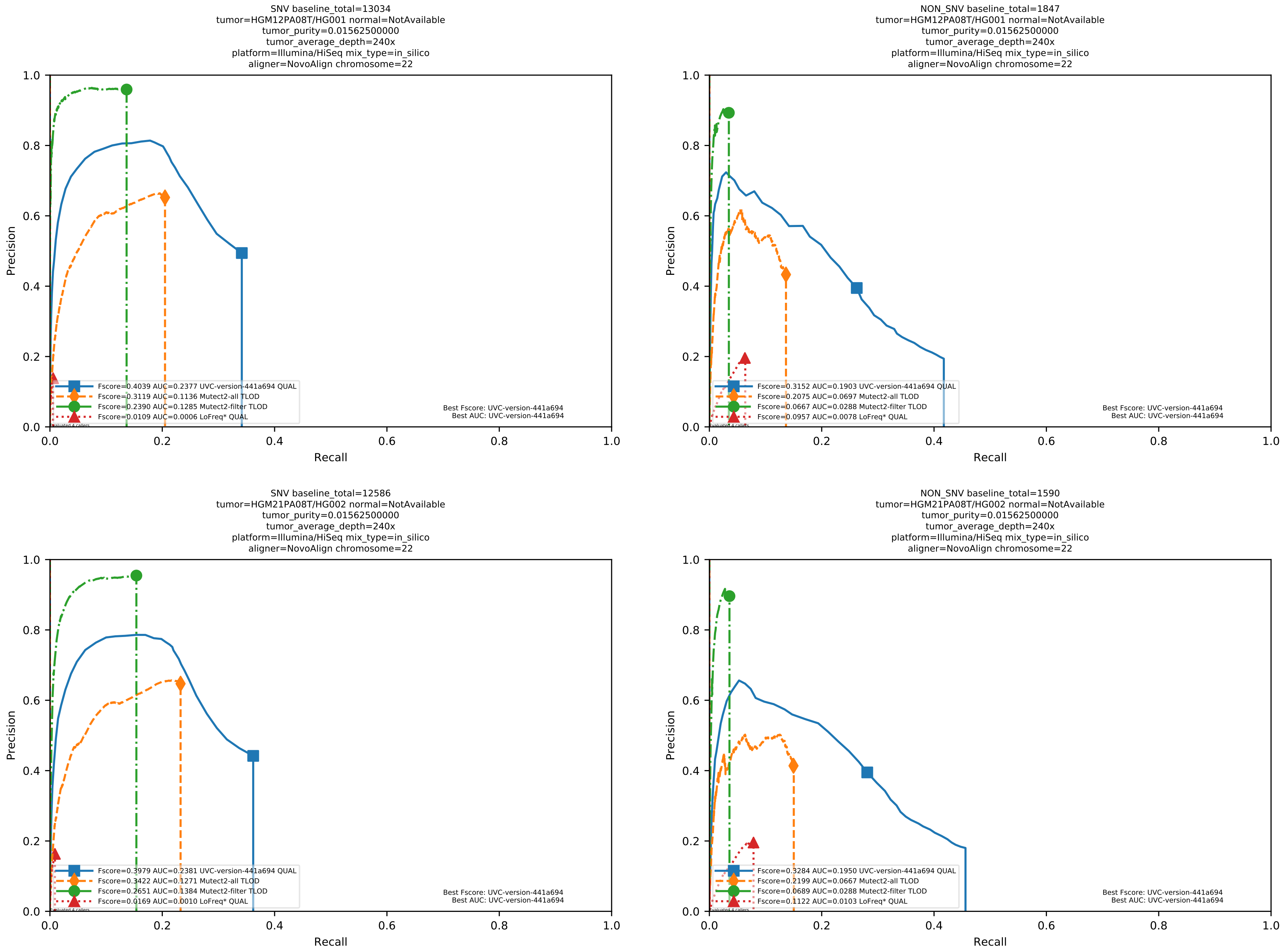

Supplementary Figure 16: Performance of somatic variant callers in tumor-only mode on the *in silico* mixtures with Illumina HiSeq sequencing data pre-aligned with NovoAlign to the human reference genome hs37d5.

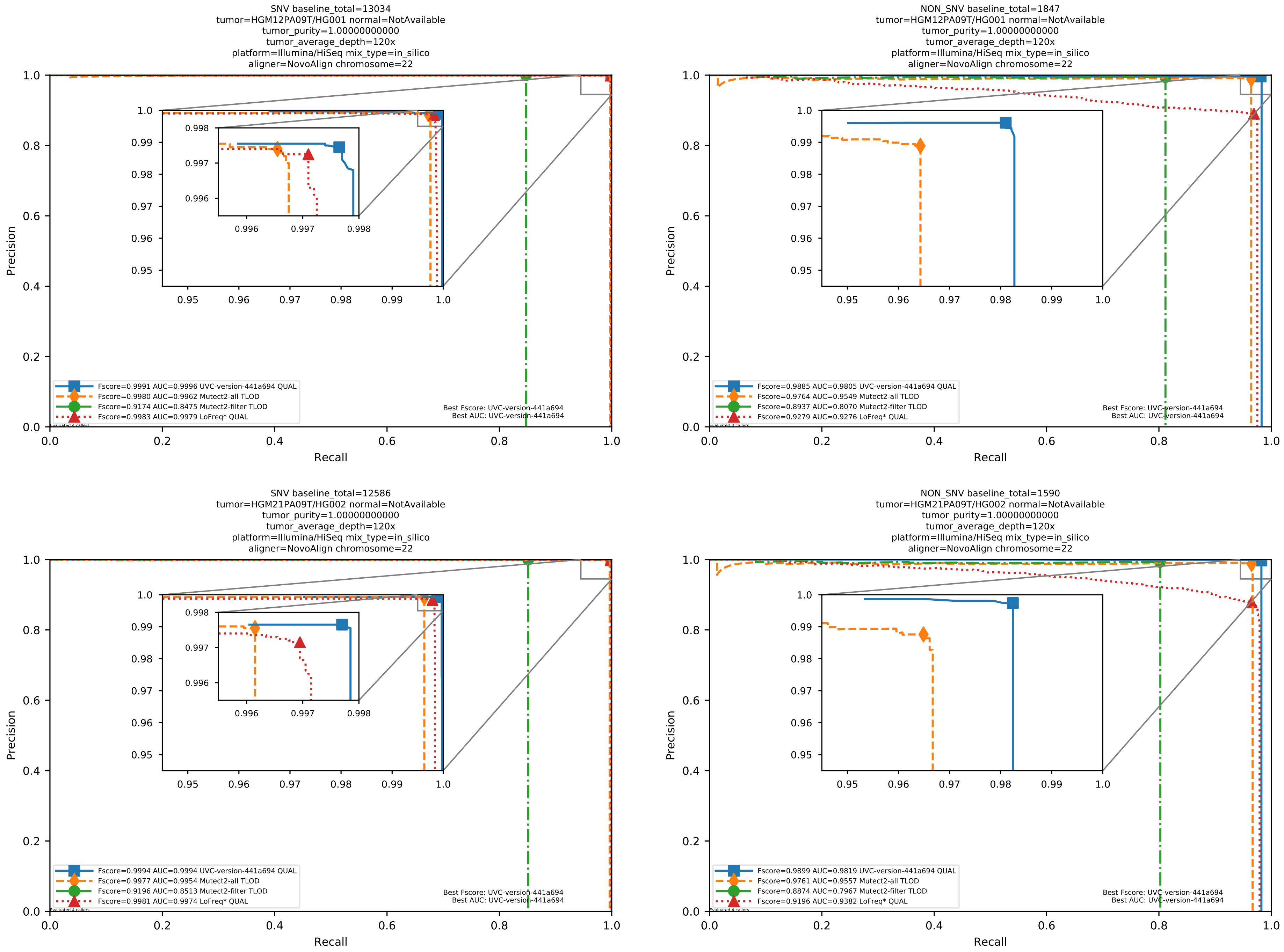

Supplementary Figure 17: Performance of somatic variant callers in tumor-only mode on the *in silico* mixtures with Illumina HiSeq sequencing data pre-aligned with NovoAlign to the human reference genome hs37d5.

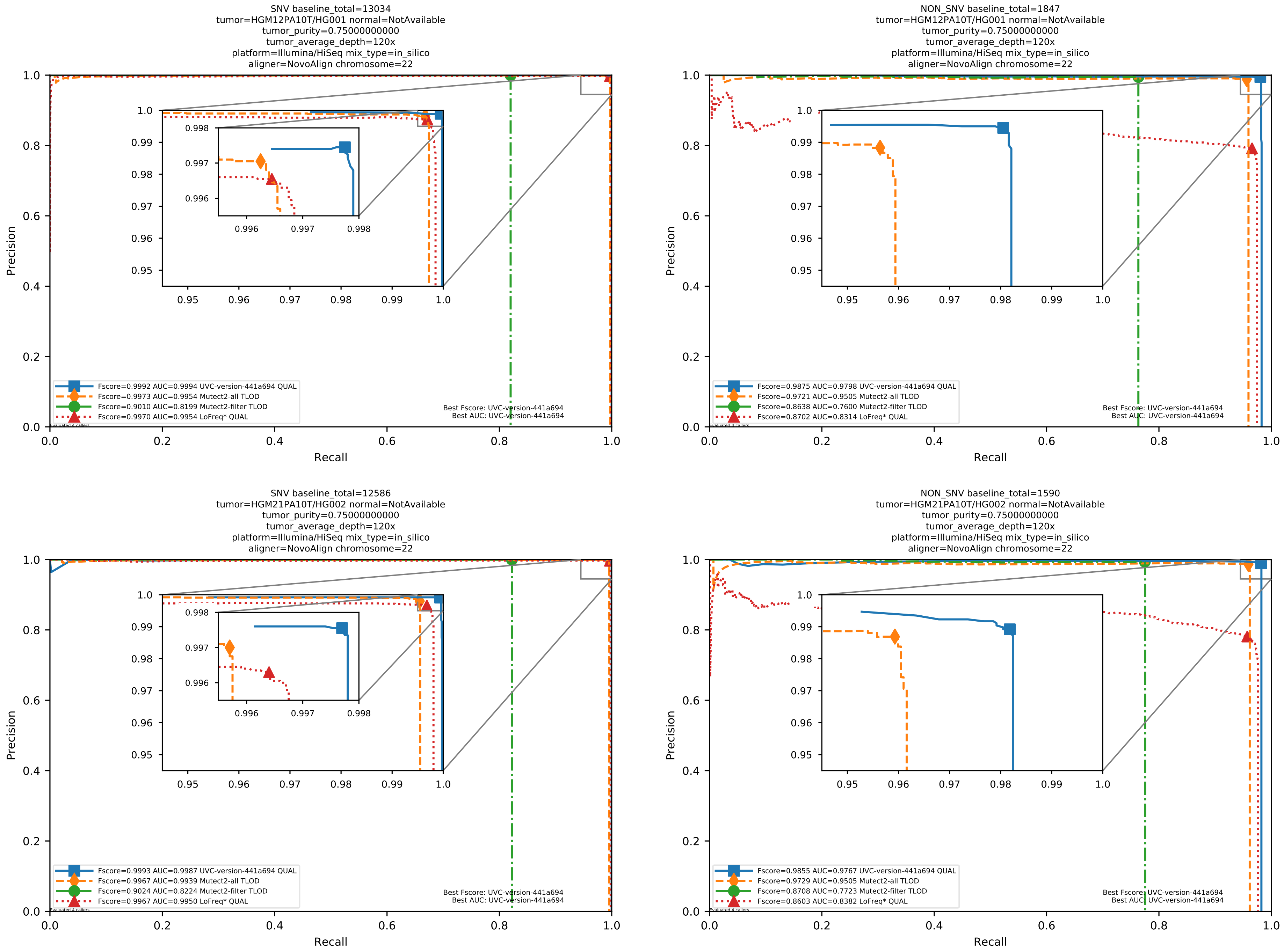

Supplementary Figure 18: Performance of somatic variant callers in tumor-only mode on the *in silico* mixtures with Illumina HiSeq sequencing data pre-aligned with NovoAlign to the human reference genome hs37d5.

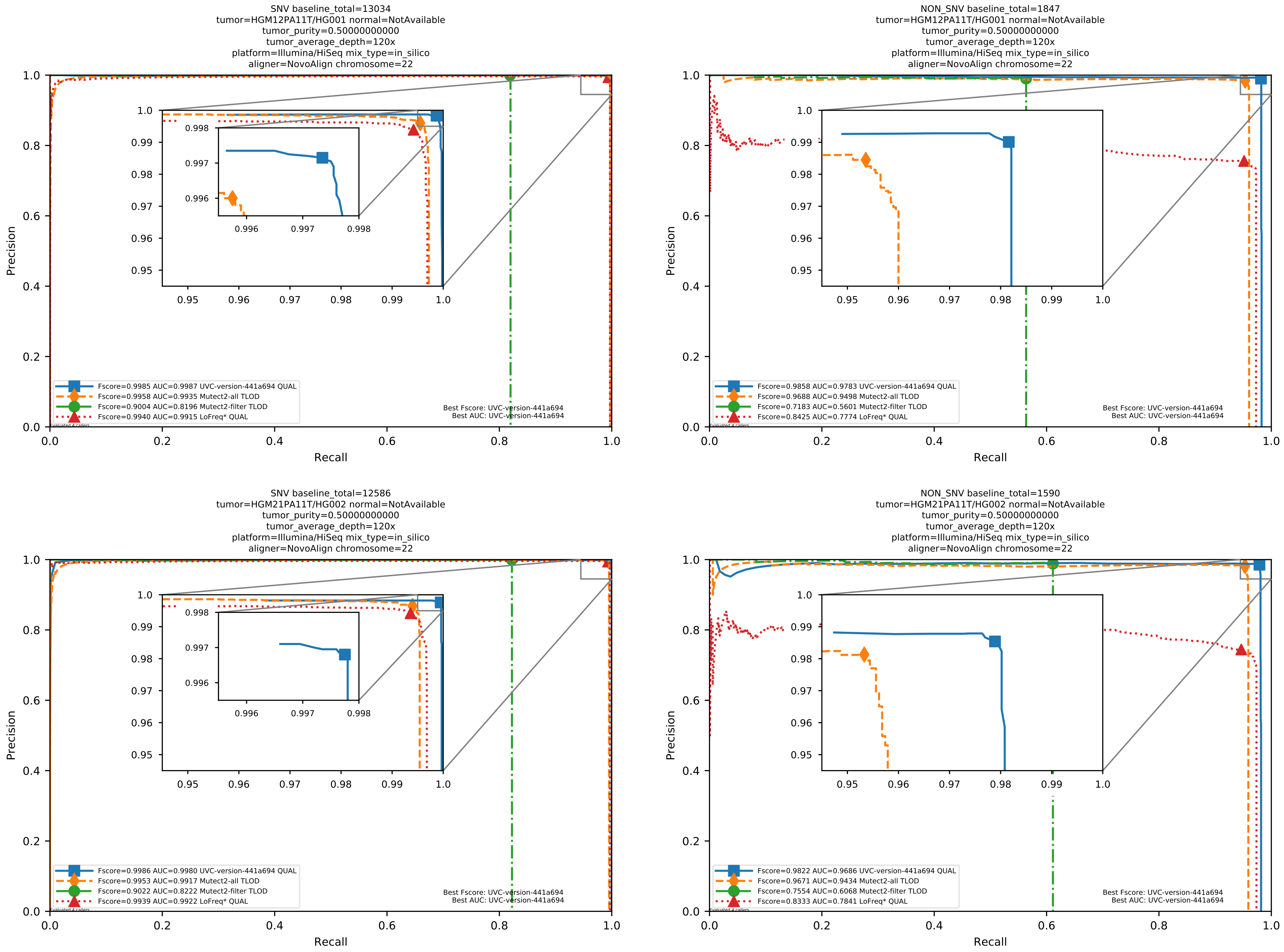

Supplementary Figure 19: Performance of somatic variant callers in tumor-only mode on the *in silico* mixtures with Illumina HiSeq sequencing data pre-aligned with NovoAlign to the human reference genome hs37d5.

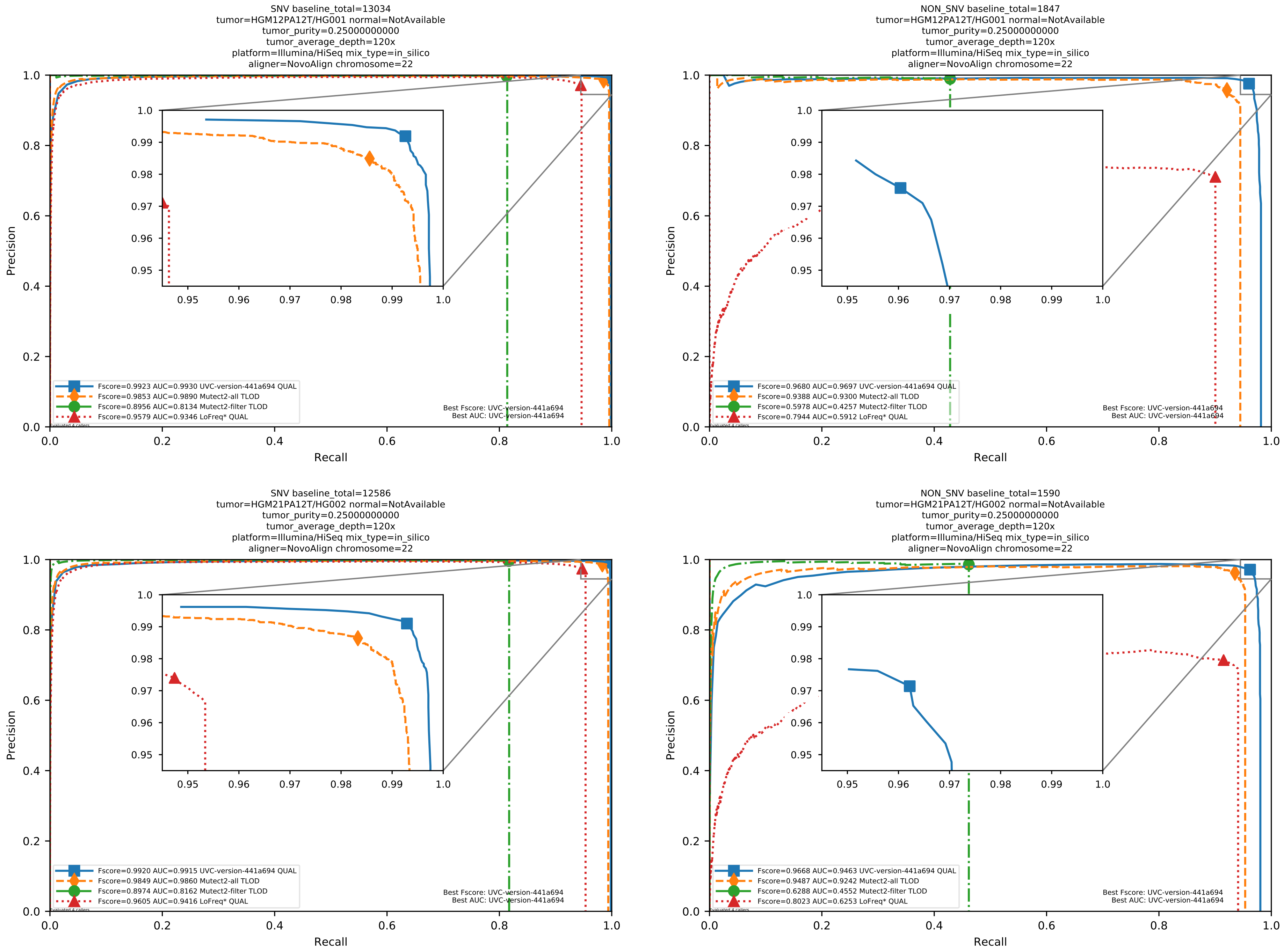

Supplementary Figure 20: Performance of somatic variant callers in tumor-only mode on the *in silico* mixtures with Illumina HiSeq sequencing data pre-aligned with NovoAlign to the human reference genome hs37d5.

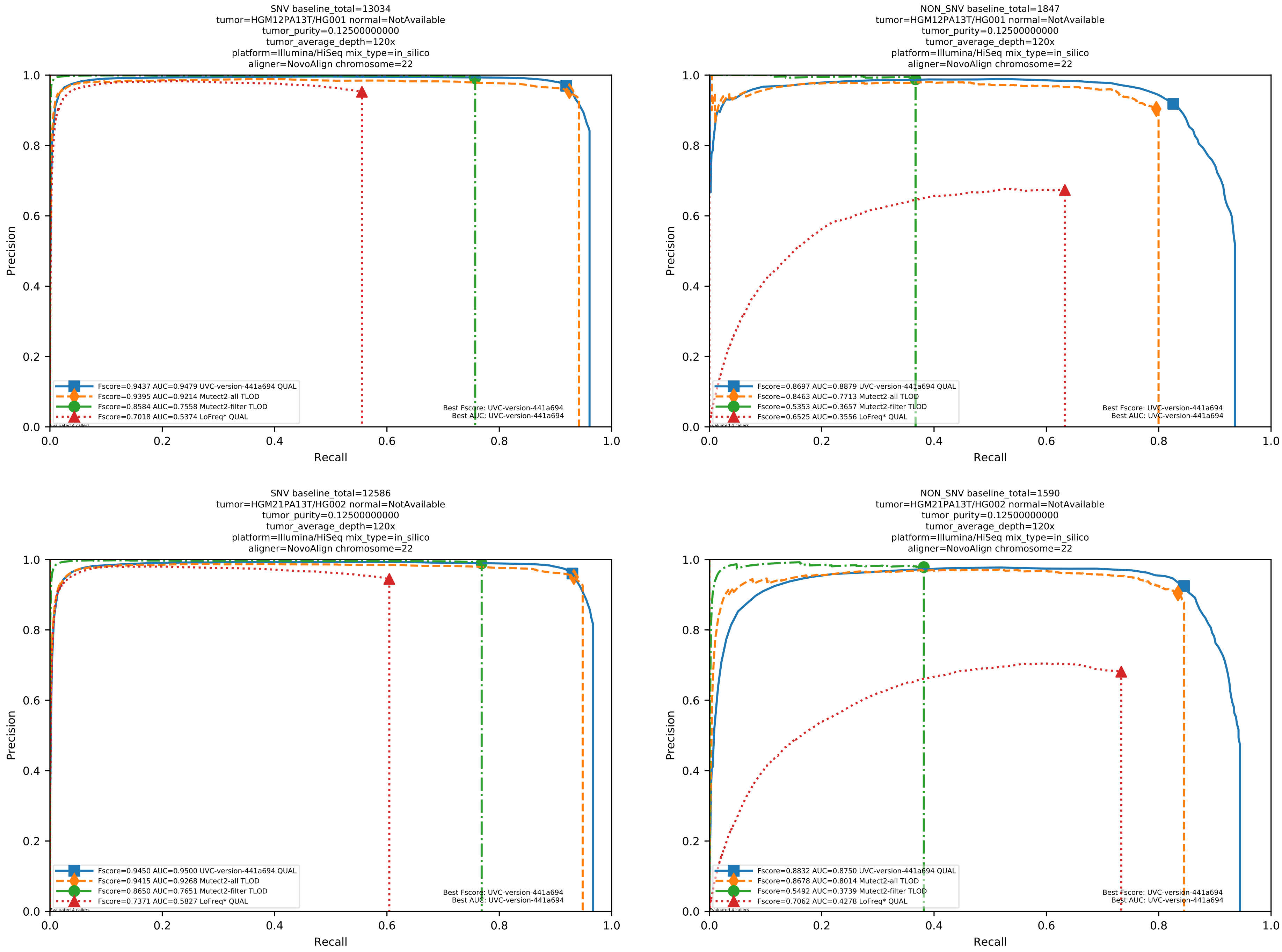

Supplementary Figure 21: Performance of somatic variant callers in tumor-only mode on the *in silico* mixtures with Illumina HiSeq sequencing data pre-aligned with NovoAlign to the human reference genome hs37d5.

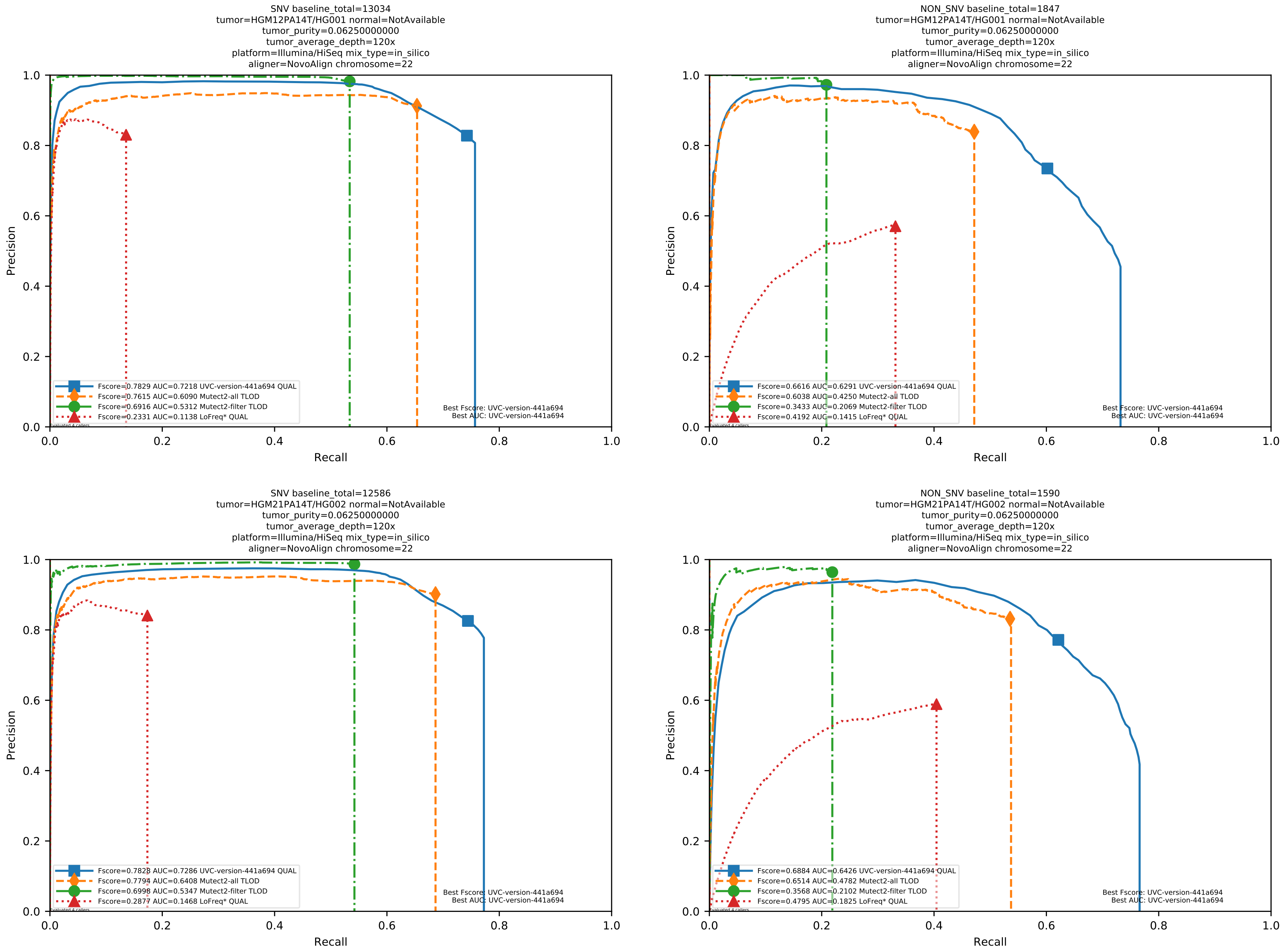

Supplementary Figure 22: Performance of somatic variant callers in tumor-only mode on the *in silico* mixtures with Illumina HiSeq sequencing data pre-aligned with NovoAlign to the human reference genome hs37d5.

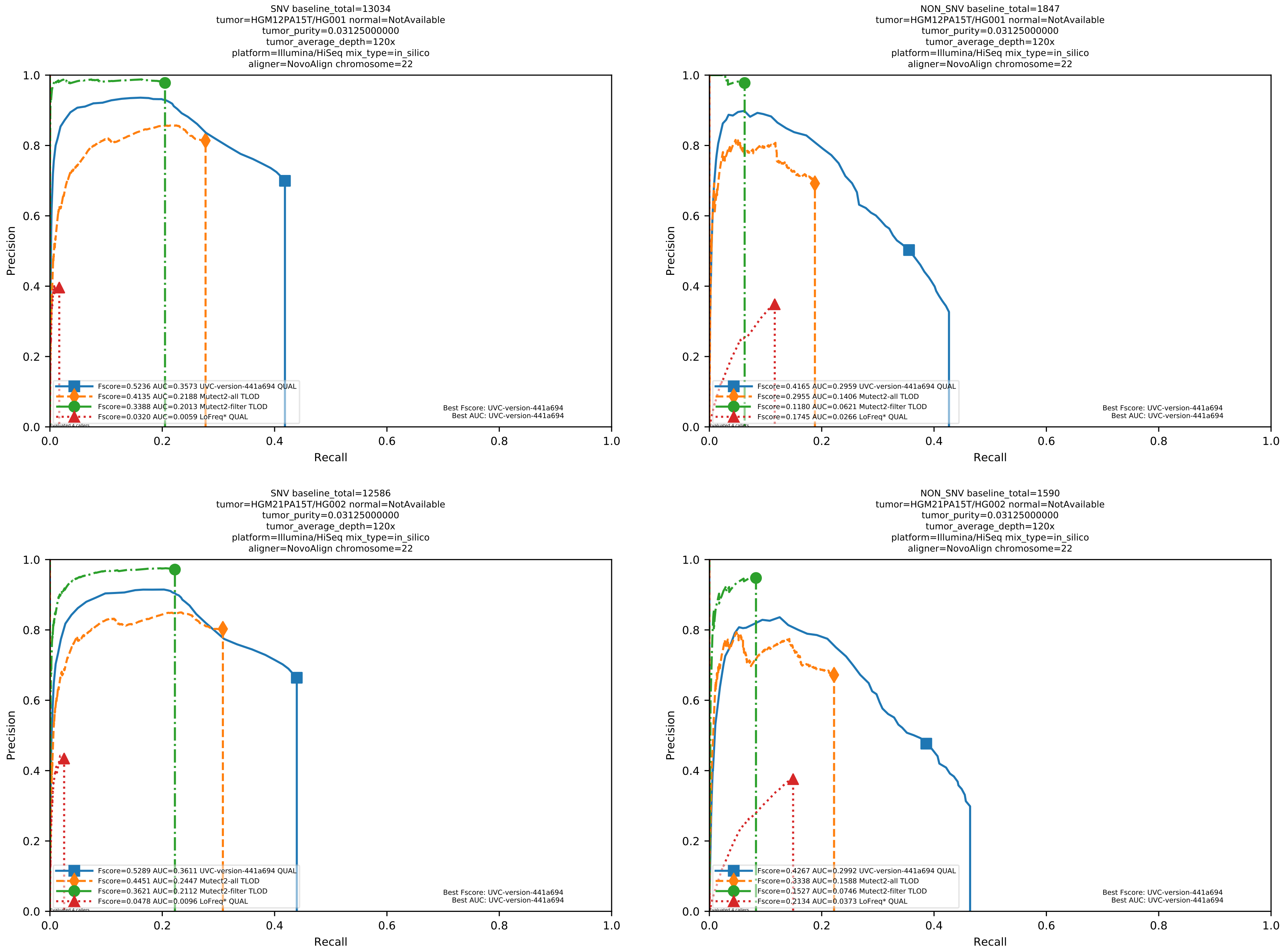

Supplementary Figure 23: Performance of somatic variant callers in tumor-only mode on the *in silico* mixtures with Illumina HiSeq sequencing data pre-aligned with NovoAlign to the human reference genome hs37d5.

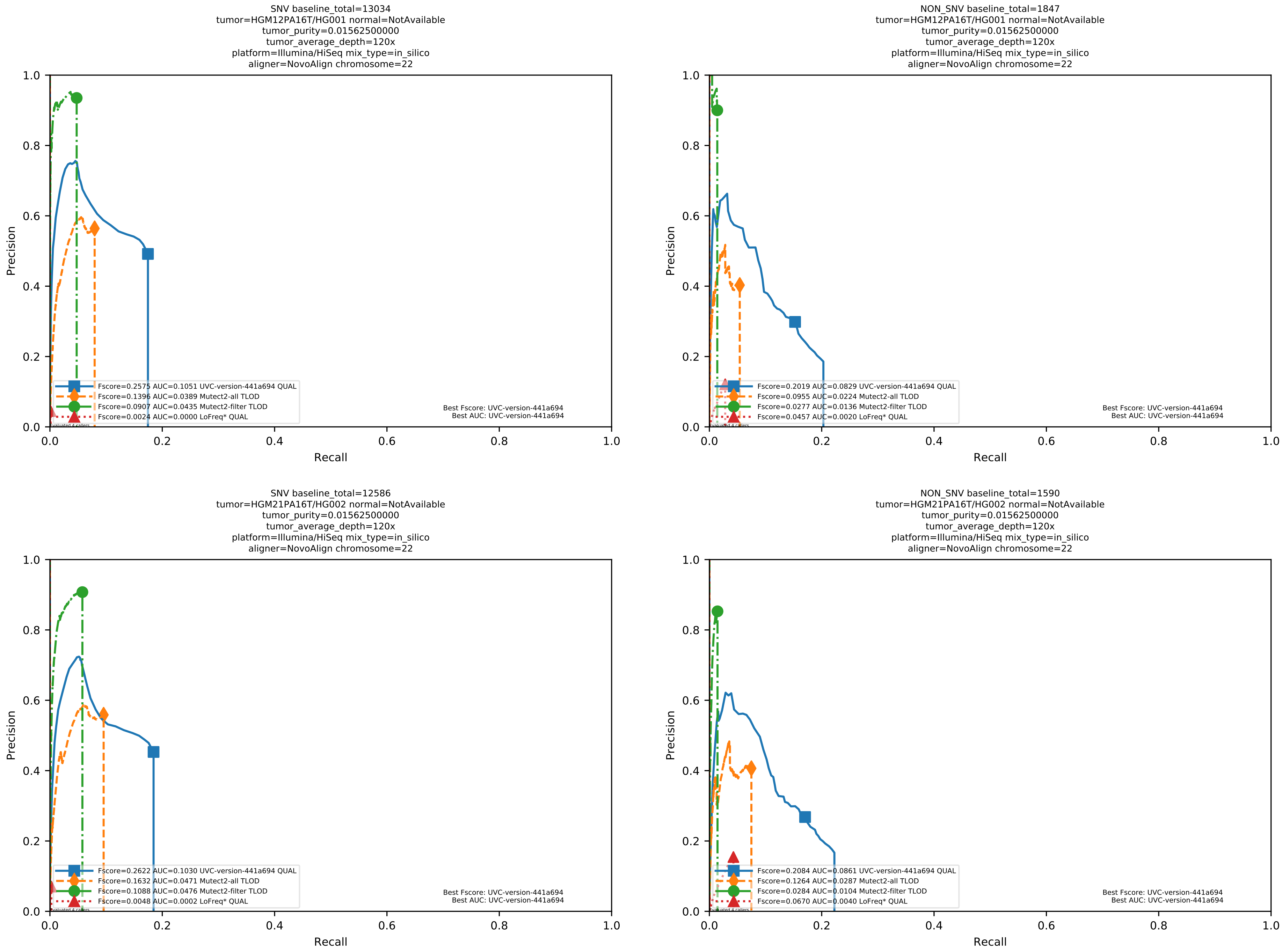

Supplementary Figure 24: Performance of somatic variant callers in tumor-only mode on the *in silico* mixtures with Illumina HiSeq sequencing data pre-aligned with NovoAlign to the human reference genome hs37d5.

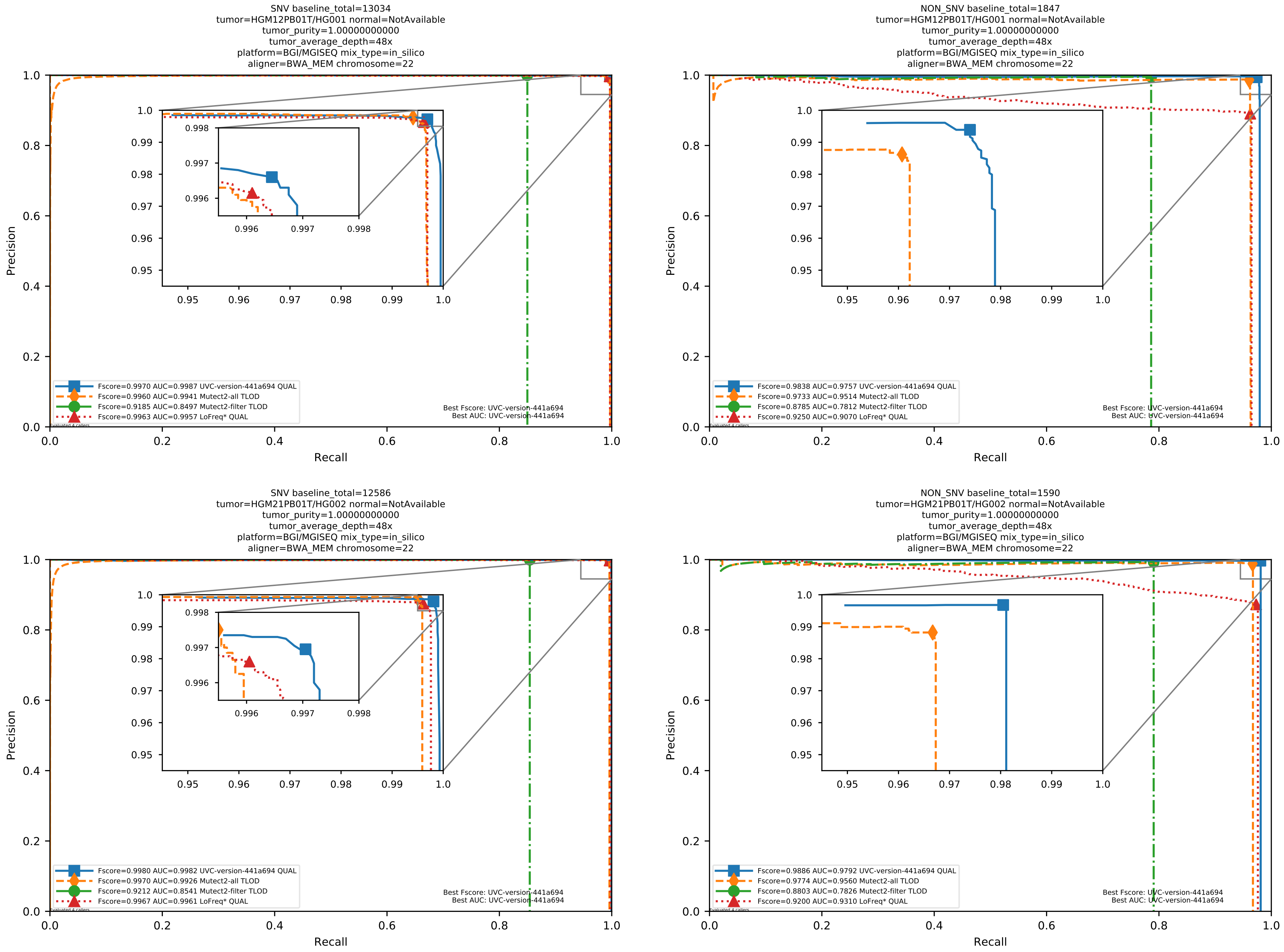

Supplementary Figure 25: Performance of somatic variant callers in tumor-only mode on the *in silico* mixtures with BGI MGISEQ sequencing data aligned with BWA-MEM to the human reference genome hs37d5.

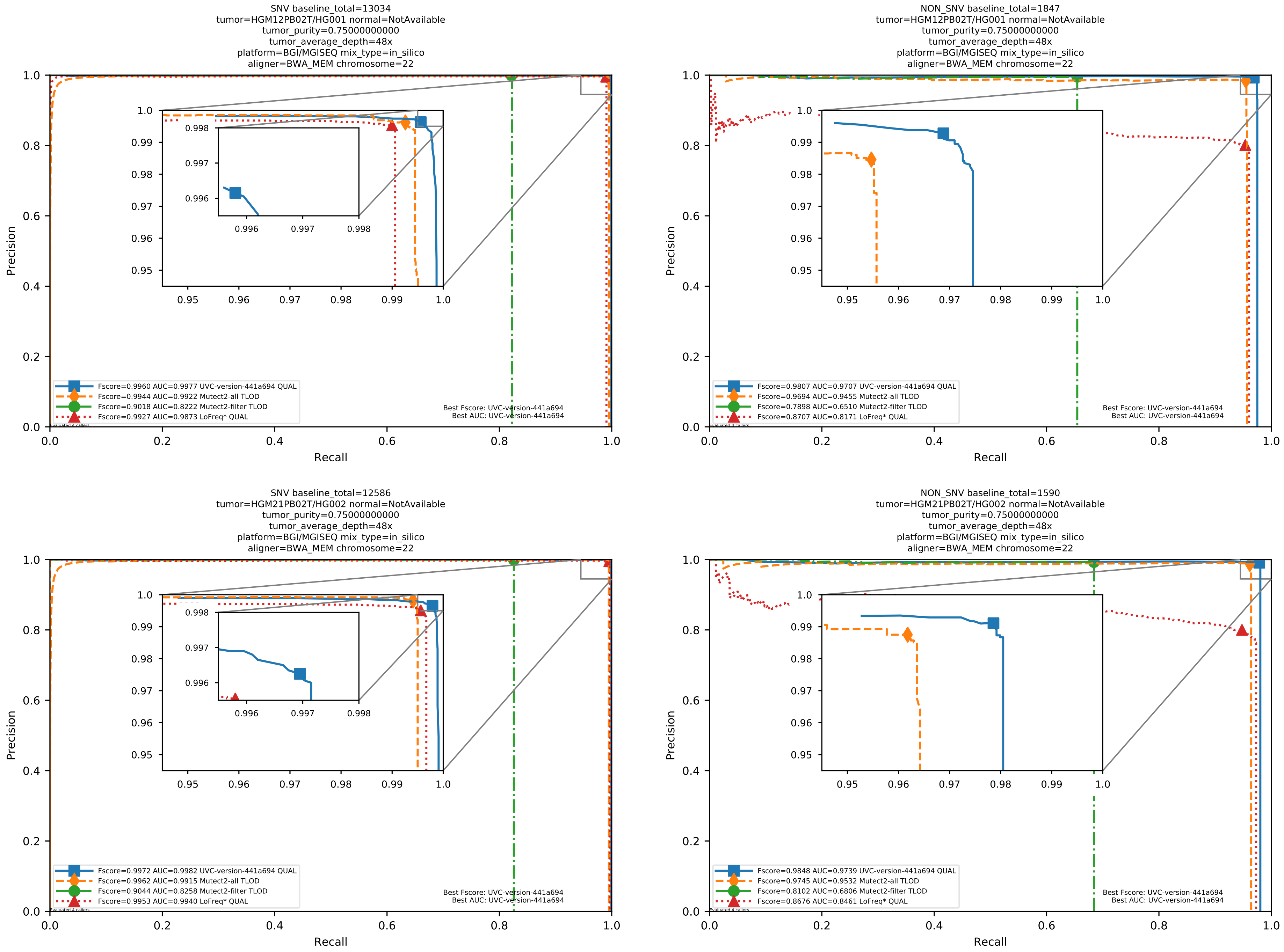

Supplementary Figure 26: Performance of somatic variant callers in tumor-only mode on the *in silico* mixtures with BGI MGISEQ sequencing data aligned with BWA-MEM to the human reference genome hs37d5.

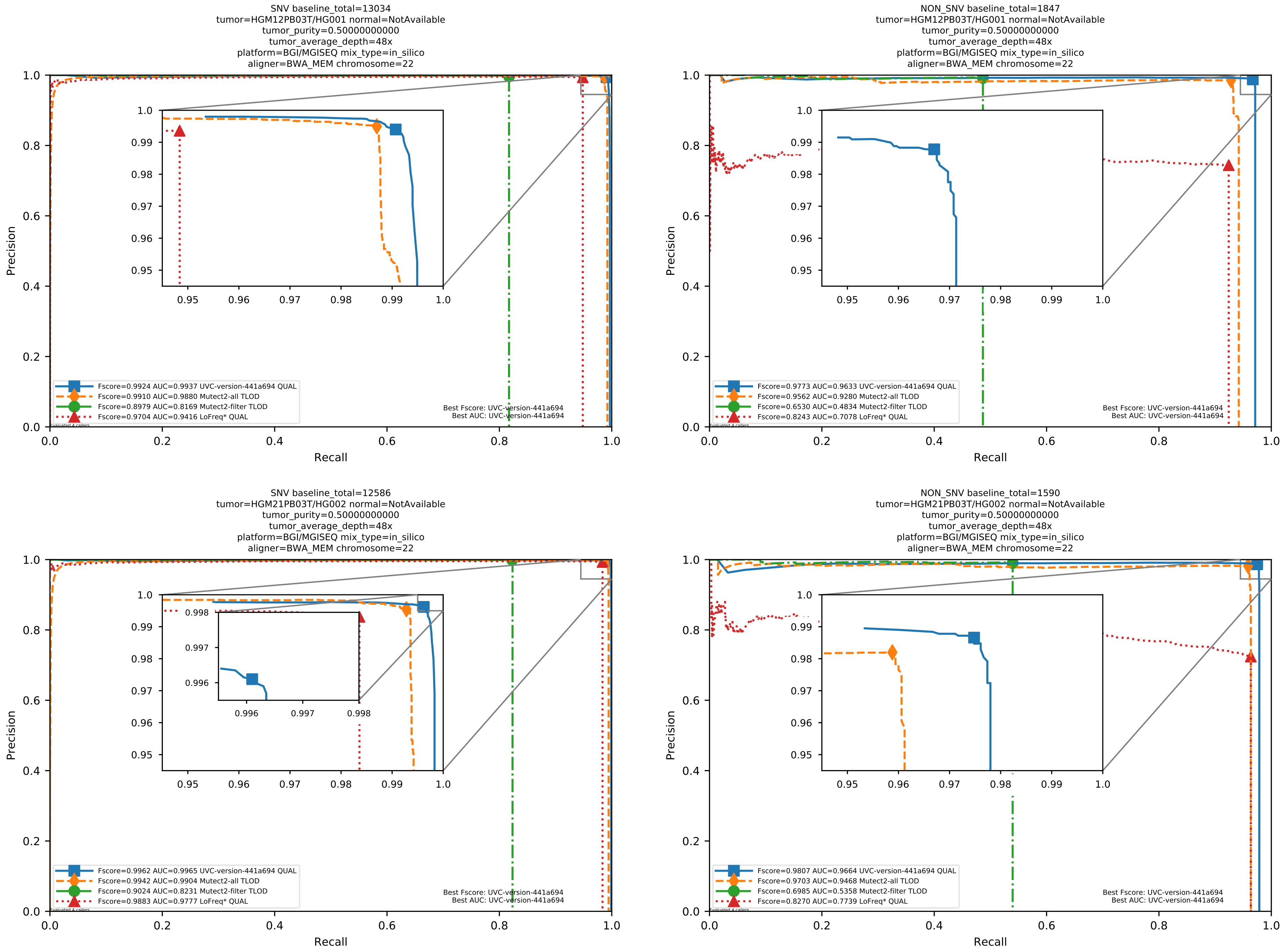

Supplementary Figure 27: Performance of somatic variant callers in tumor-only mode on the *in silico* mixtures with BGI MGISEQ sequencing data aligned with BWA-MEM to the human reference genome hs37d5.

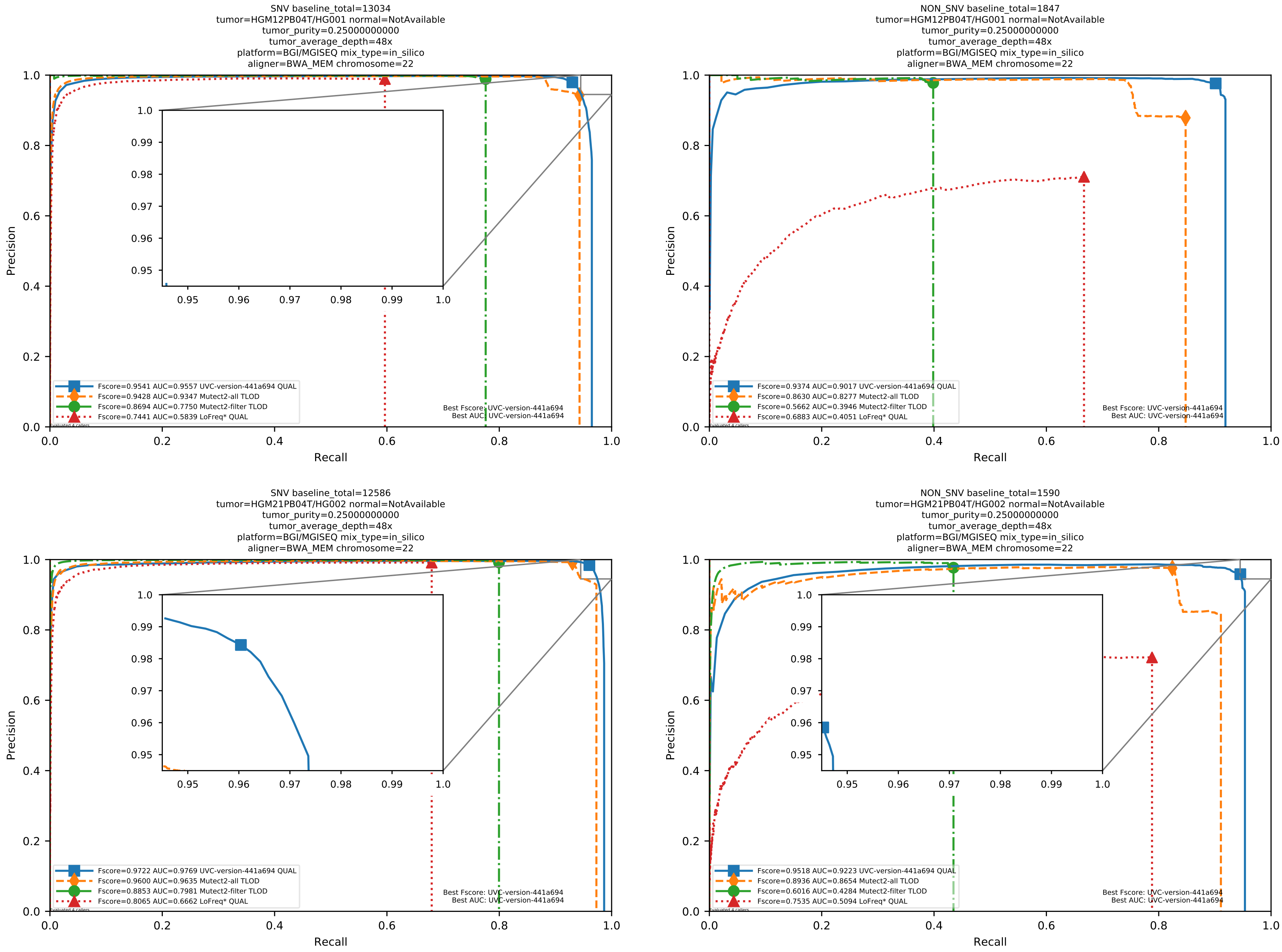

Supplementary Figure 28: Performance of somatic variant callers in tumor-only mode on the *in silico* mixtures with BGI MGISEQ sequencing data aligned with BWA-MEM to the human reference genome hs37d5.

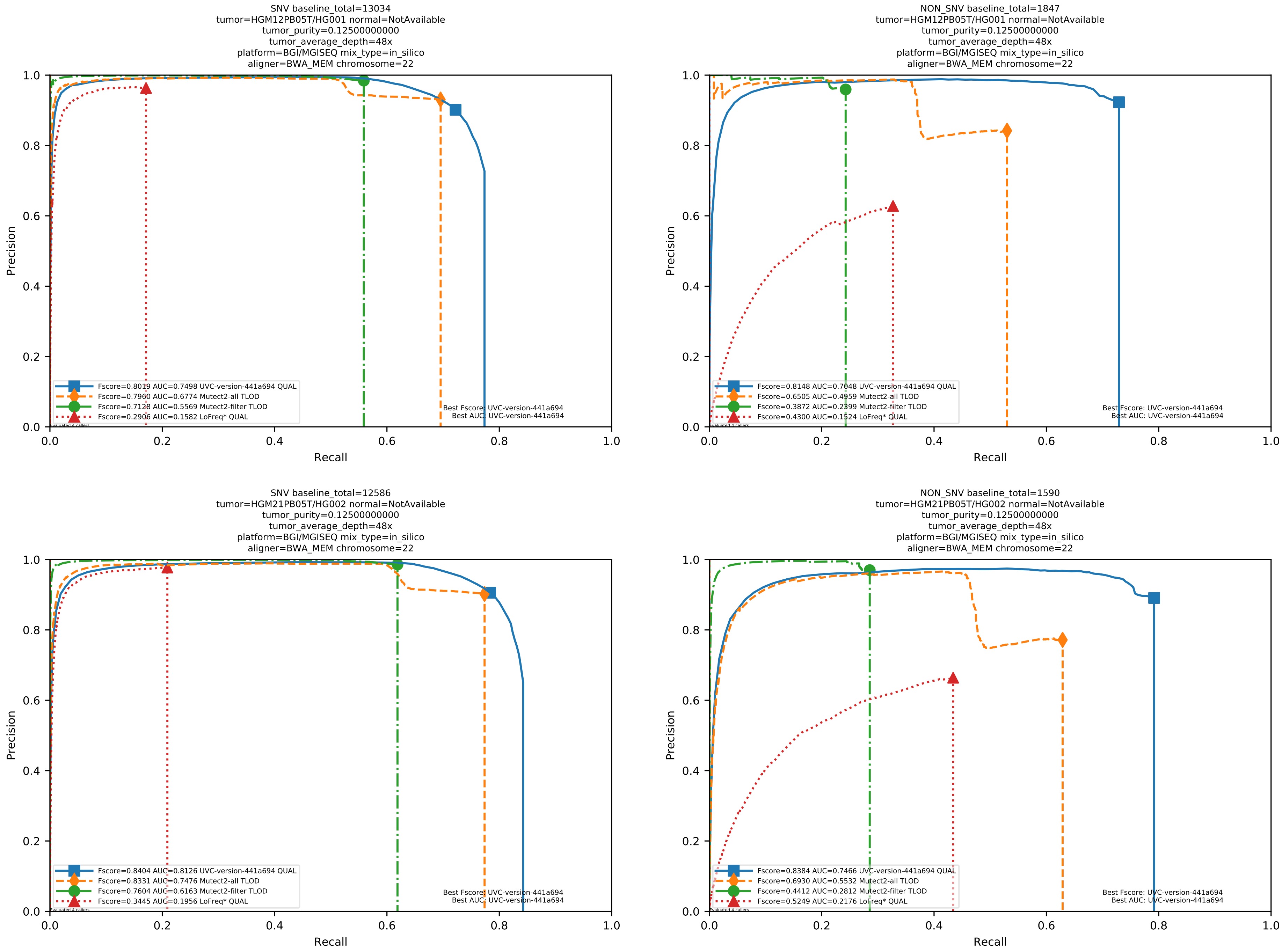

Supplementary Figure 29: Performance of somatic variant callers in tumor-only mode on the *in silico* mixtures with BGI MGISEQ sequencing data aligned with BWA-MEM to the human reference genome hs37d5.

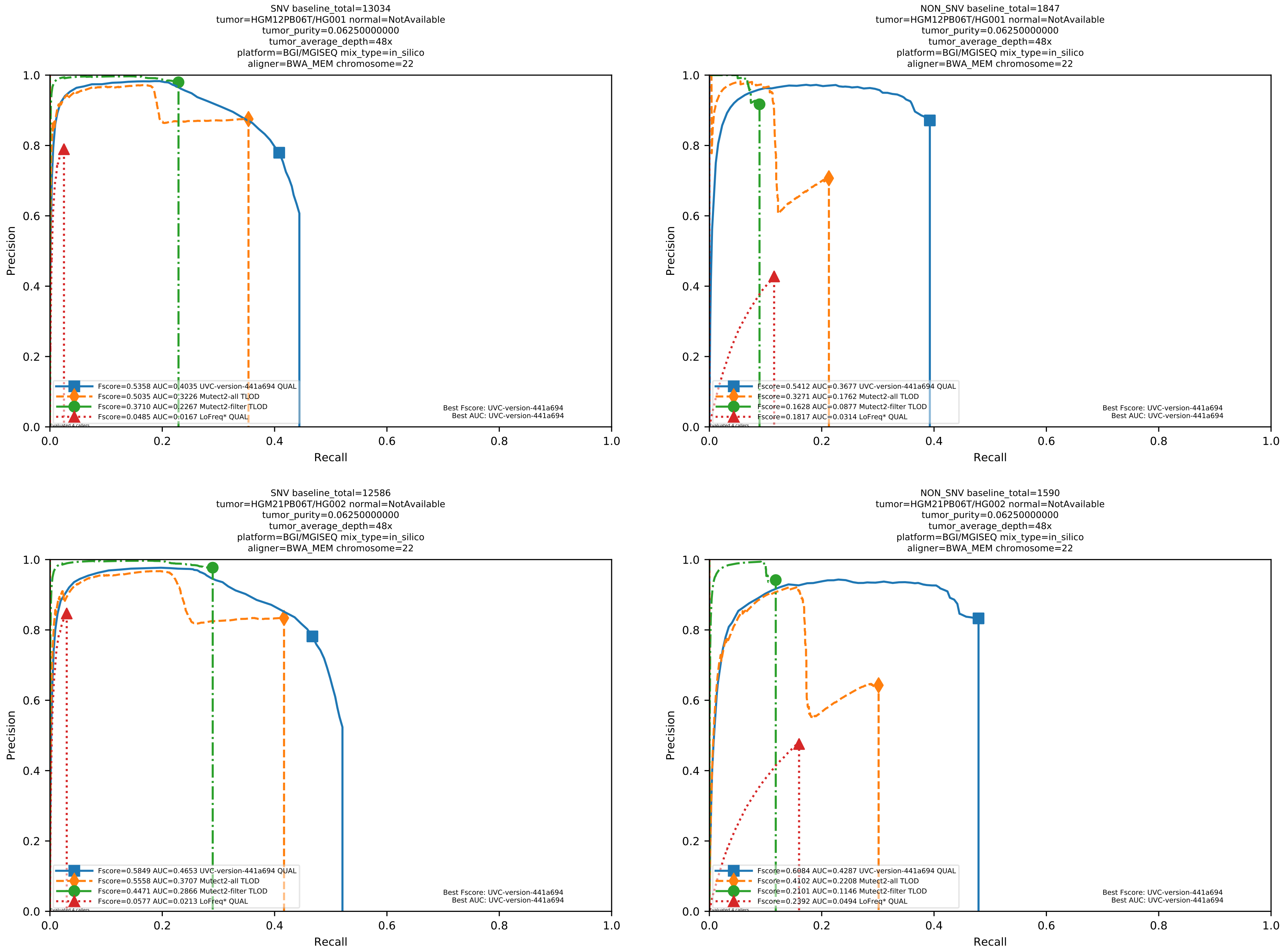

Supplementary Figure 30: Performance of somatic variant callers in tumor-only mode on the *in silico* mixtures with BGI MGISEQ sequencing data aligned with BWA-MEM to the human reference genome hs37d5.

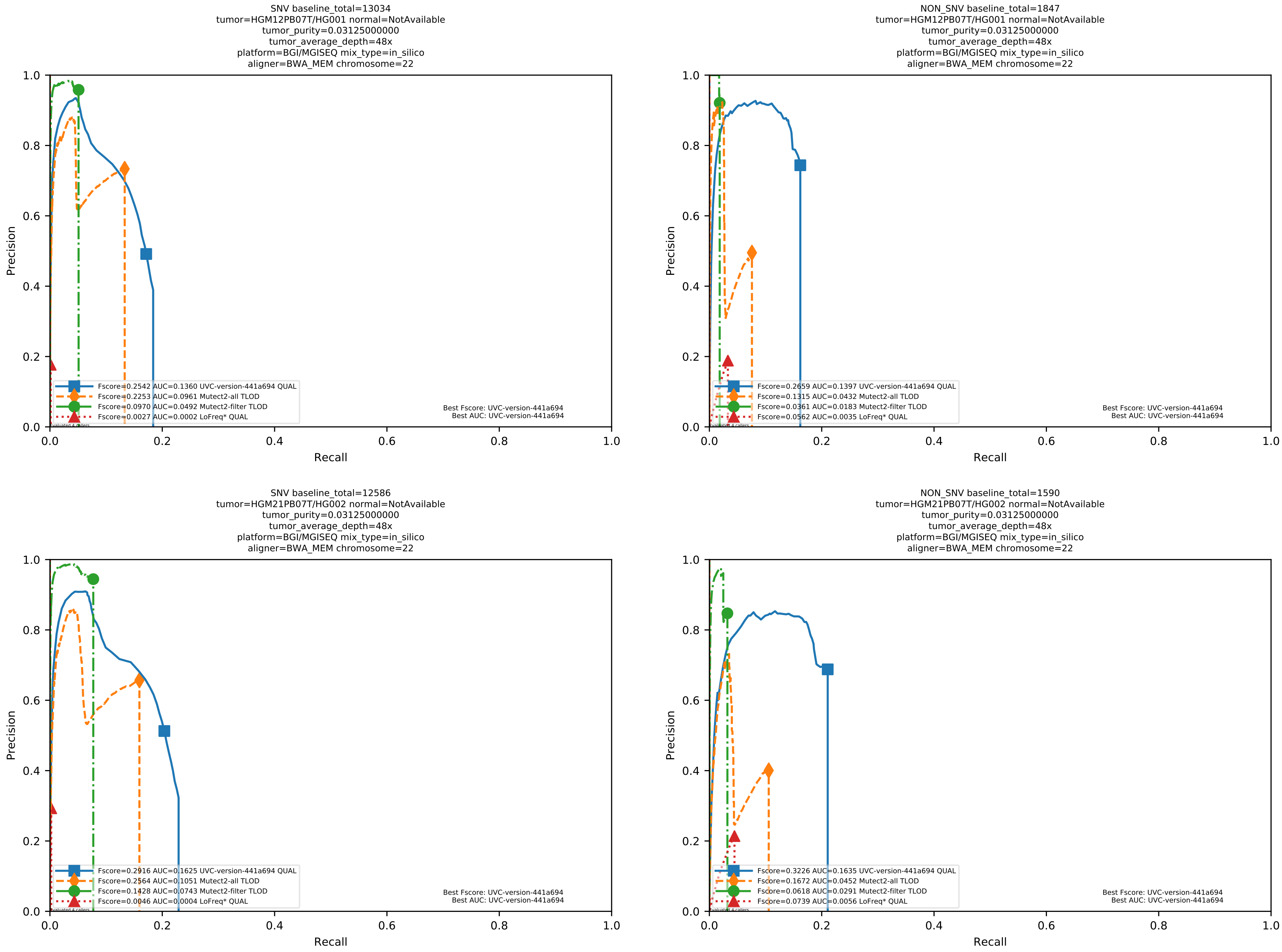

Supplementary Figure 31: Performance of somatic variant callers in tumor-only mode on the *in silico* mixtures with BGI MGISEQ sequencing data aligned with BWA-MEM to the human reference genome hs37d5.

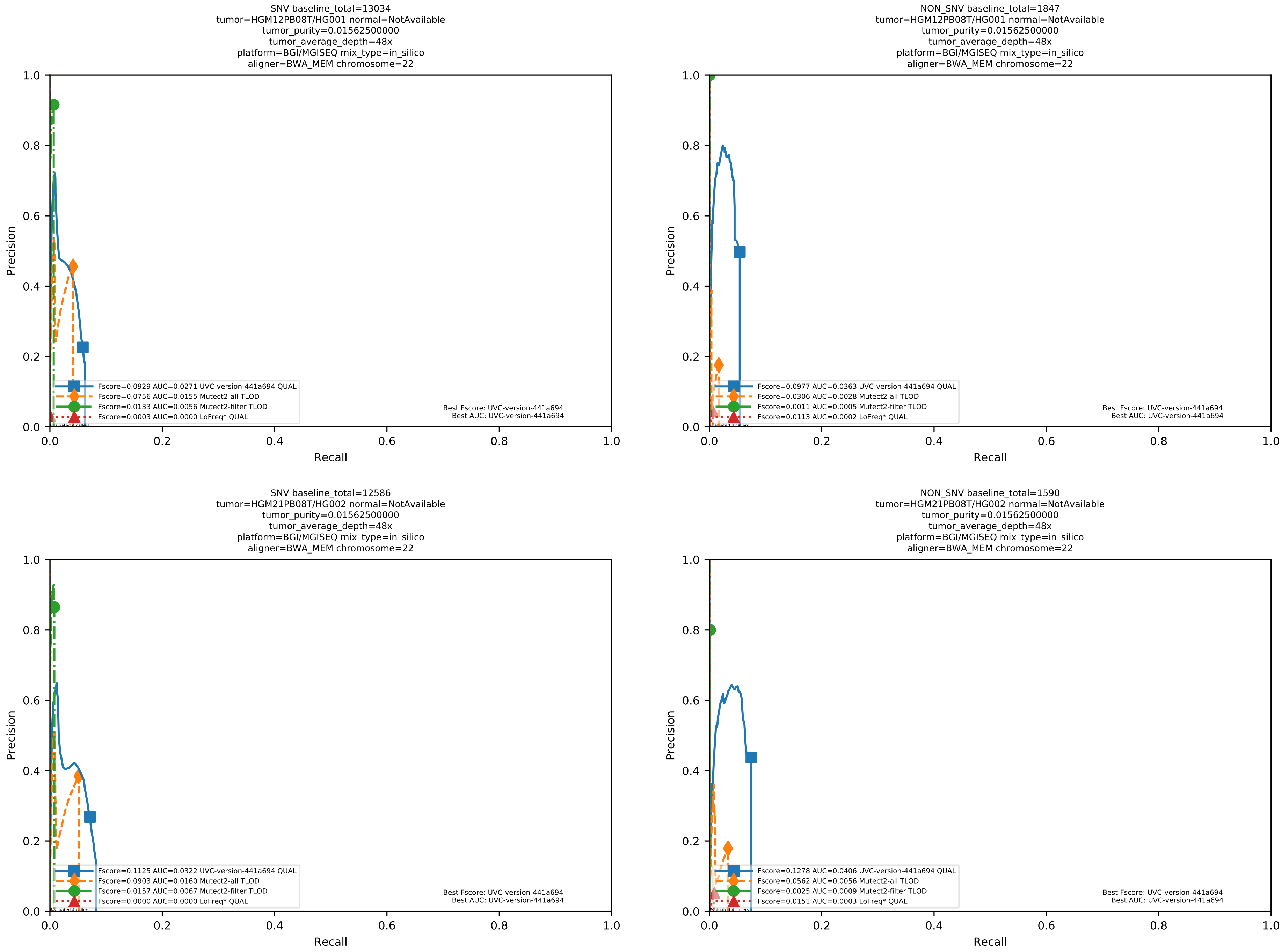

Supplementary Figure 32: Performance of somatic variant callers in tumor-only mode on the *in silico* mixtures with BGI MGISEQ sequencing data aligned with BWA-MEM to the human reference genome hs37d5.

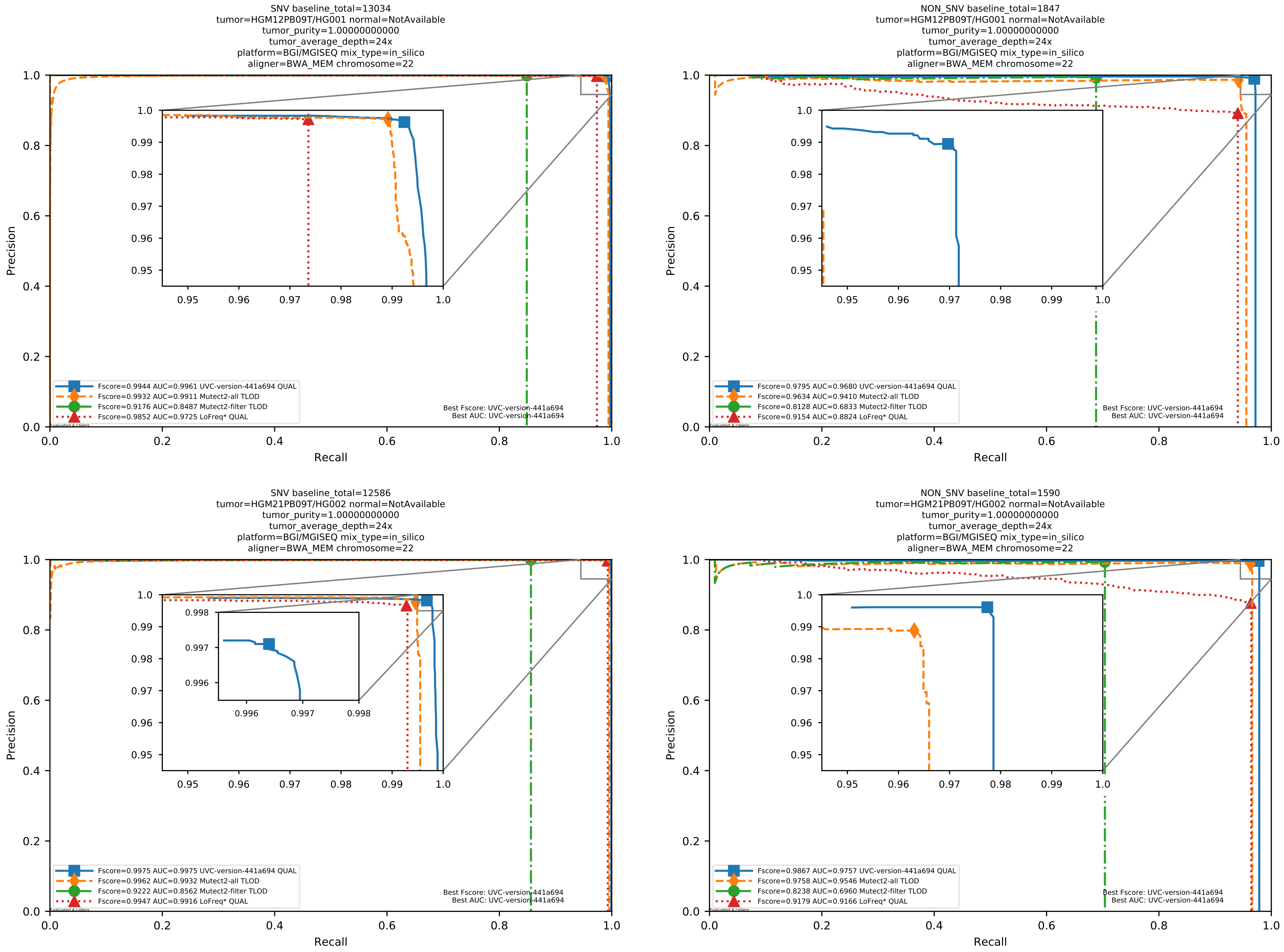

Supplementary Figure 33: Performance of somatic variant callers in tumor-only mode on the *in silico* mixtures with BGI MGISEQ sequencing data aligned with BWA-MEM to the human reference genome hs37d5.

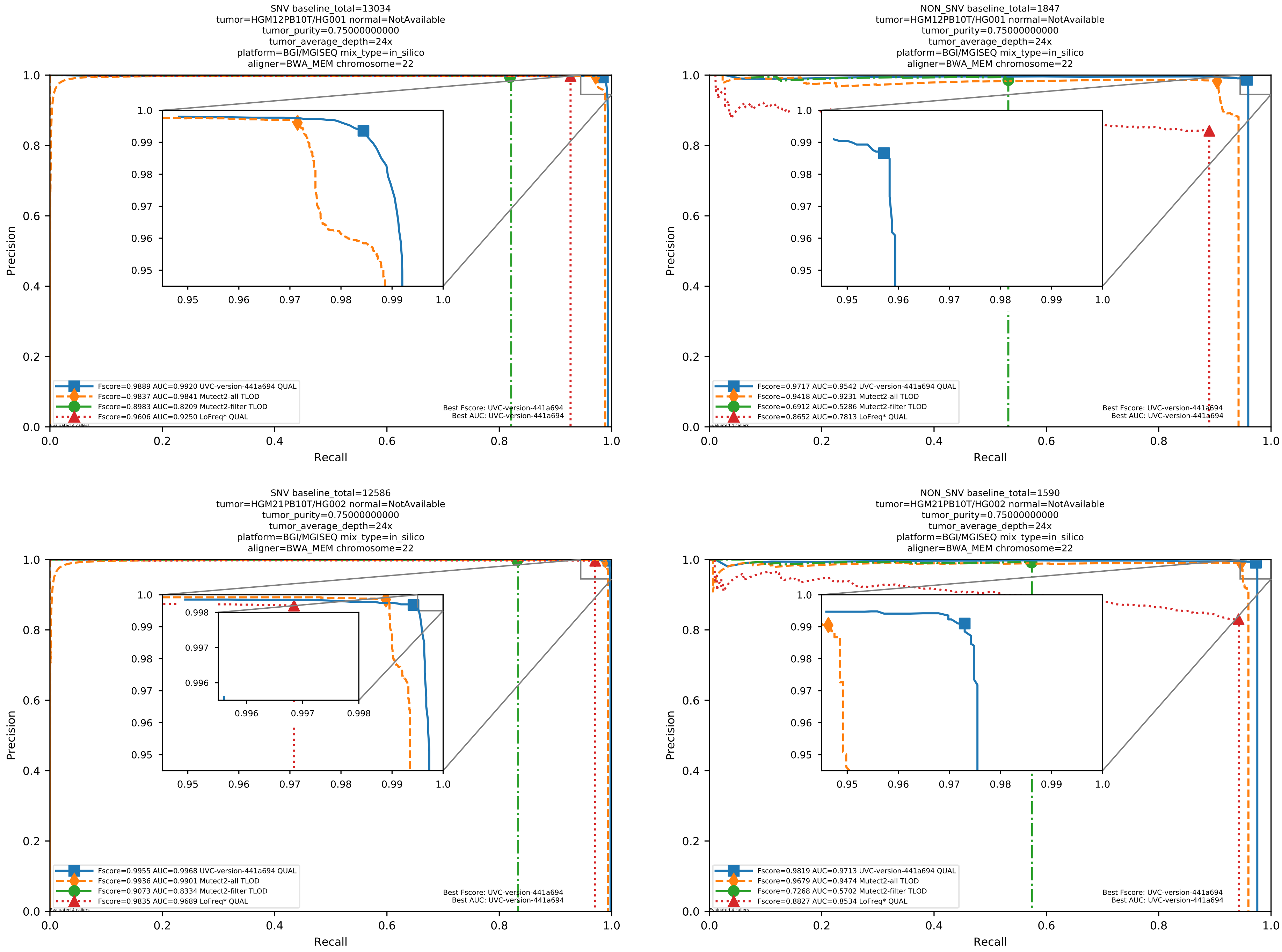

Supplementary Figure 34: Performance of somatic variant callers in tumor-only mode on the *in silico* mixtures with BGI MGISEQ sequencing data aligned with BWA-MEM to the human reference genome hs37d5.

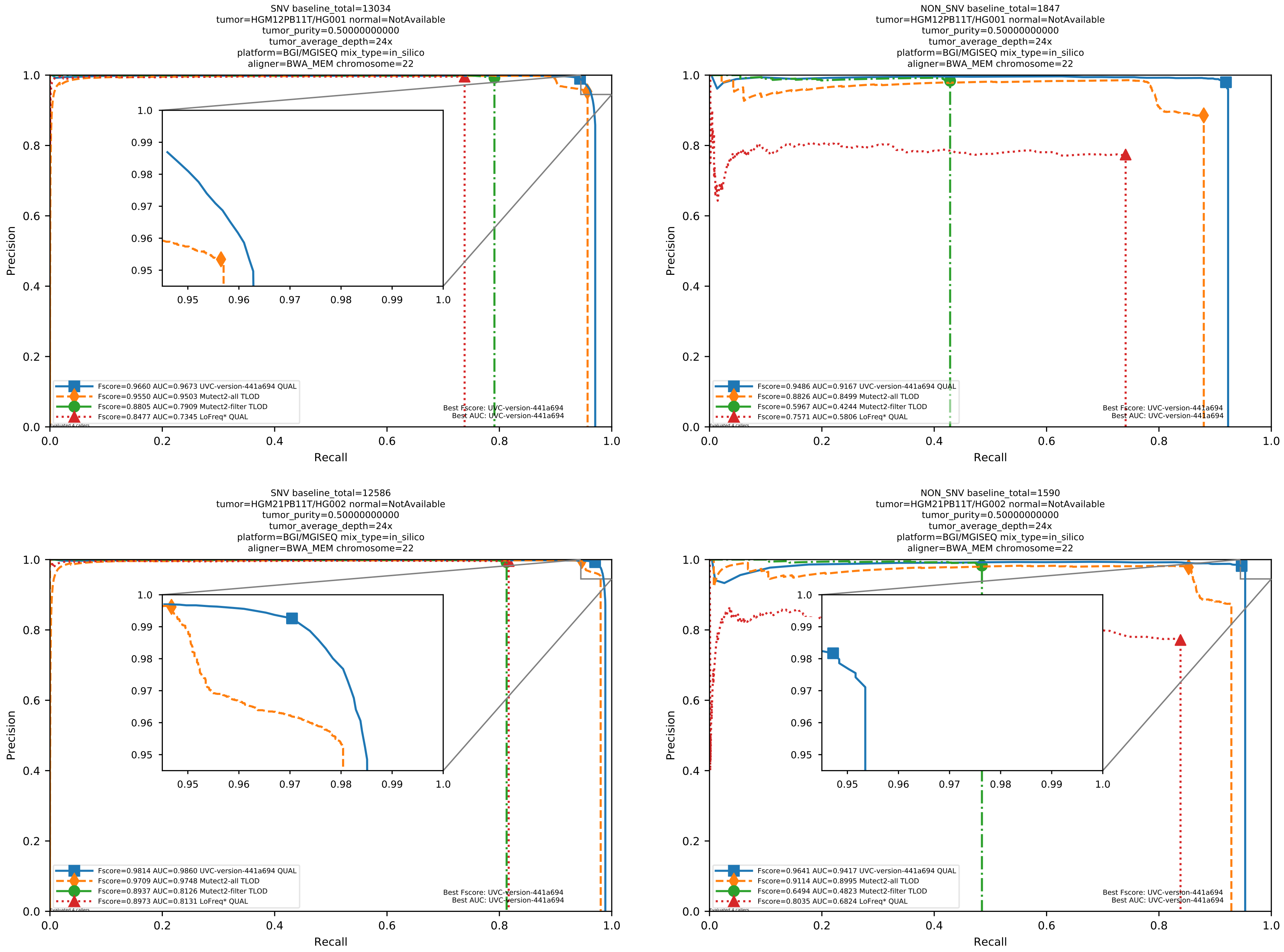

Supplementary Figure 35: Performance of somatic variant callers in tumor-only mode on the *in silico* mixtures with BGI MGISEQ sequencing data aligned with BWA-MEM to the human reference genome hs37d5.

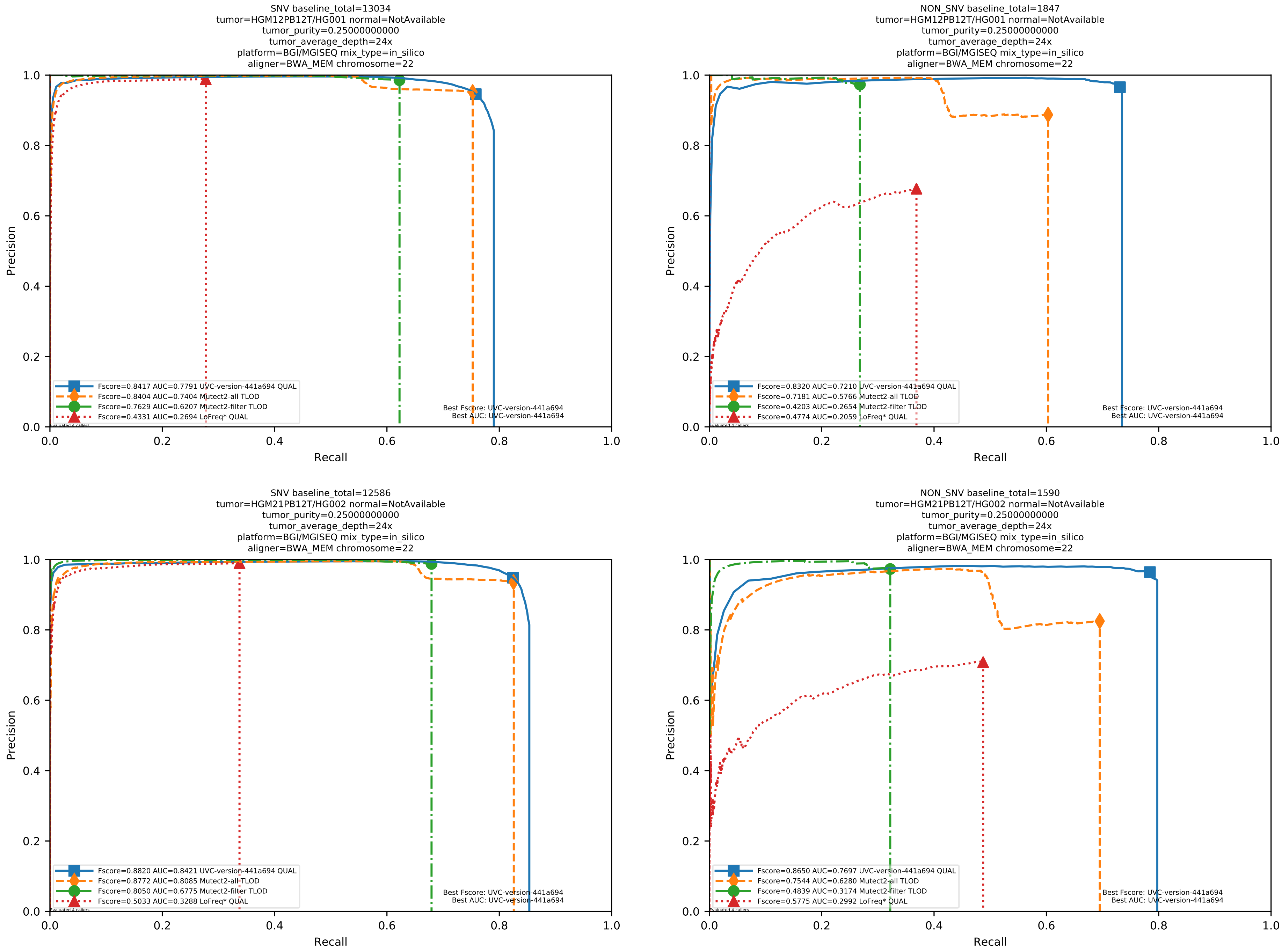

Supplementary Figure 36: Performance of somatic variant callers in tumor-only mode on the *in silico* mixtures with BGI MGISEQ sequencing data aligned with BWA-MEM to the human reference genome hs37d5.

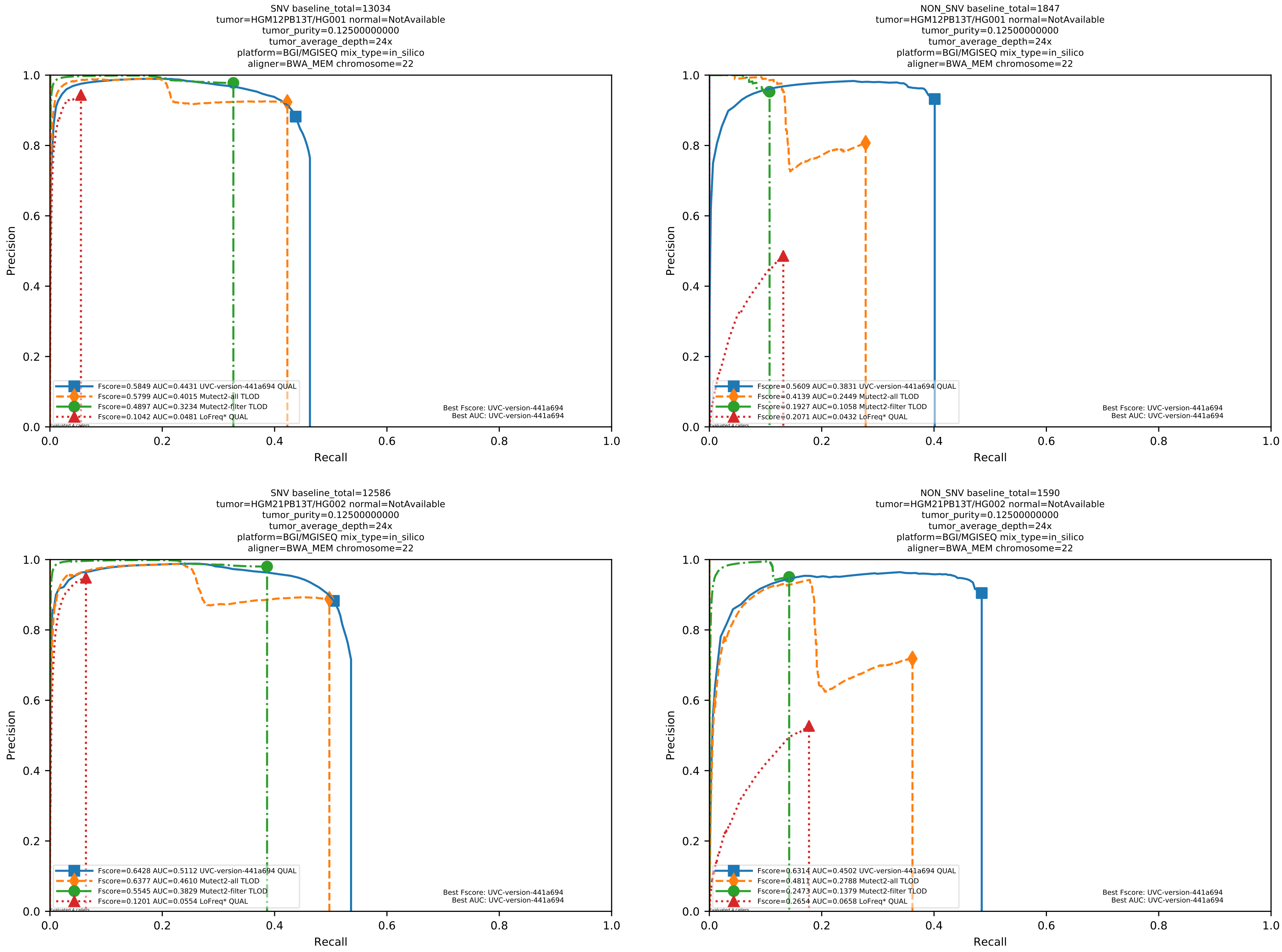

Supplementary Figure 37: Performance of somatic variant callers in tumor-only mode on the *in silico* mixtures with BGI MGISEQ sequencing data aligned with BWA-MEM to the human reference genome hs37d5.

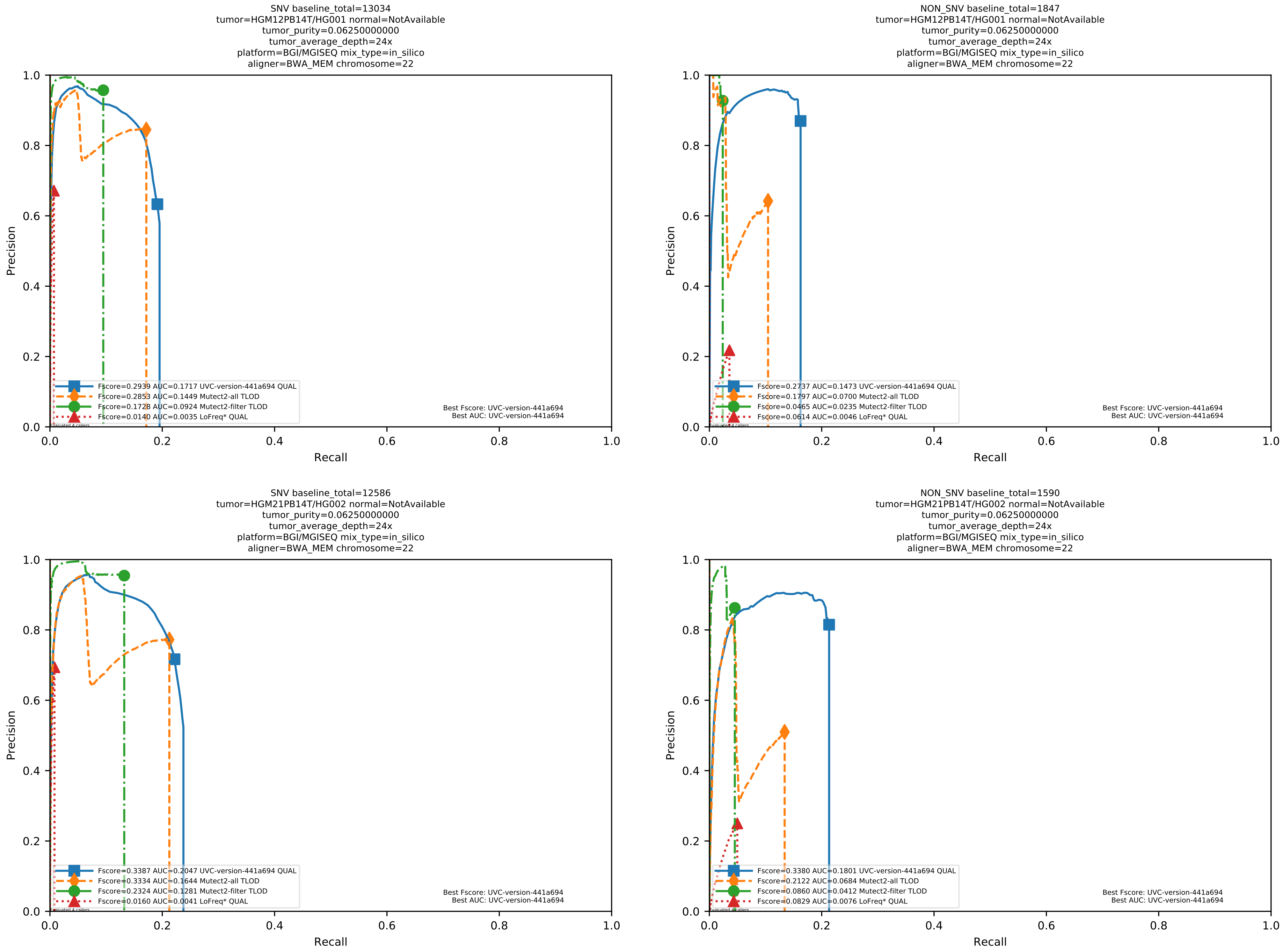

Supplementary Figure 38: Performance of somatic variant callers in tumor-only mode on the *in silico* mixtures with BGI MGISEQ sequencing data aligned with BWA-MEM to the human reference genome hs37d5.

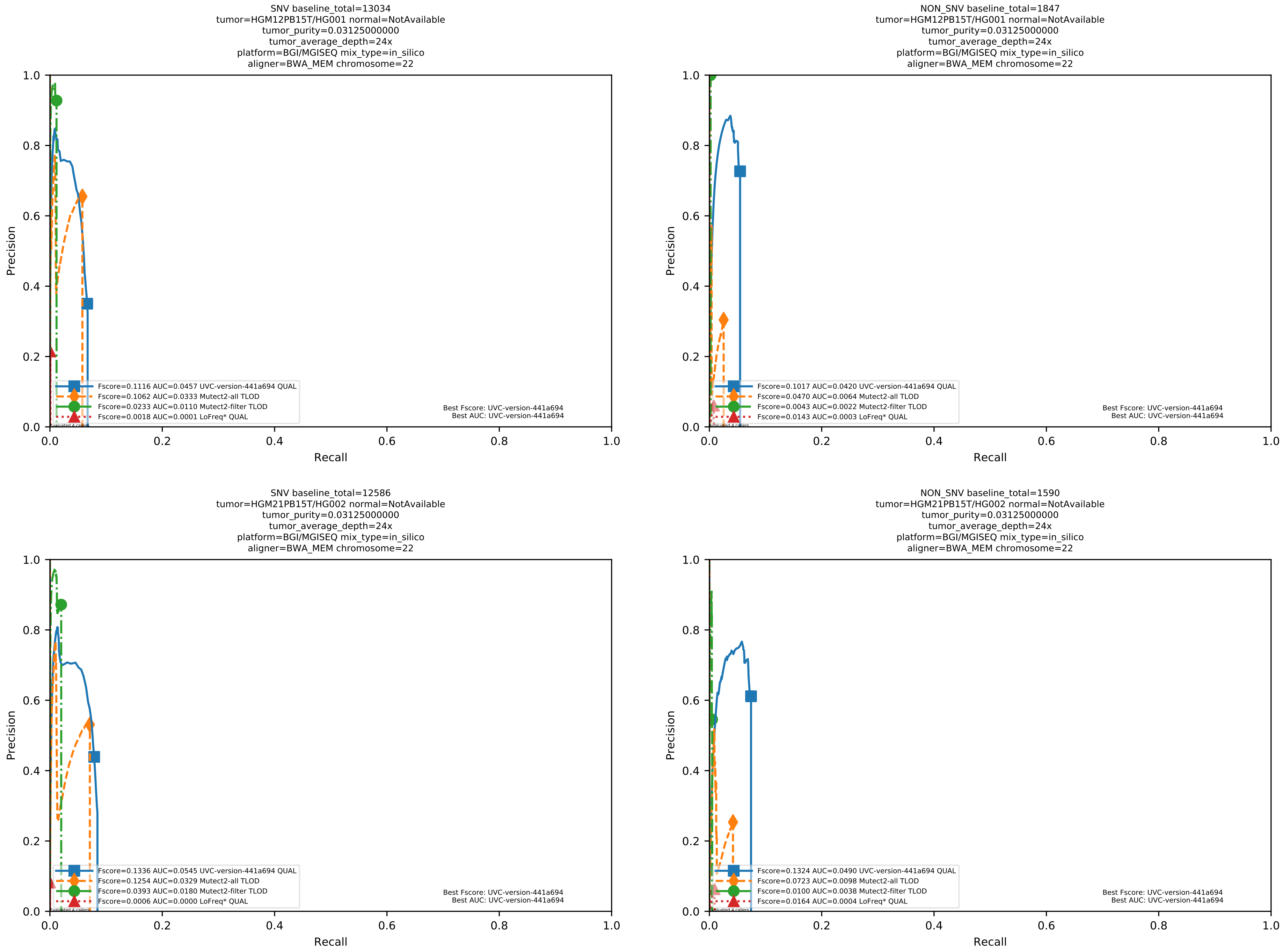

Supplementary Figure 39: Performance of somatic variant callers in tumor-only mode on the *in silico* mixtures with BGI MGISEQ sequencing data aligned with BWA-MEM to the human reference genome hs37d5.

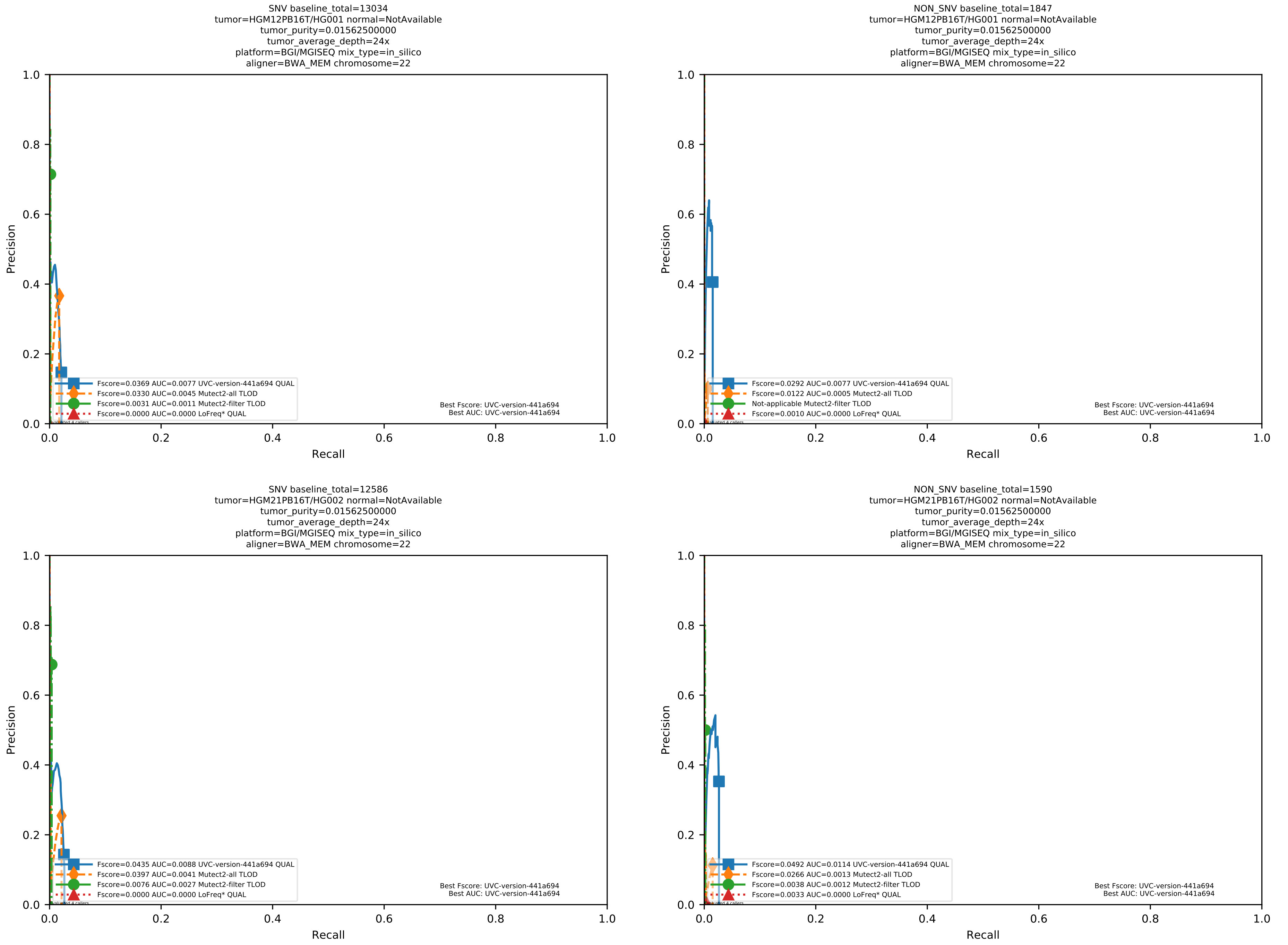

Supplementary Figure 40: Performance of somatic variant callers in tumor-only mode on the *in silico* mixtures with BGI MGISEQ sequencing data aligned with BWA-MEM to the human reference genome hs37d5.

##### 1.4 Supplemental evaluation results on WGS datasets for calling somatic variants from tumor-normal pairs, with 128 *in silico* mixtures considering both Illumina and BGI platforms

The datasets in this subsection were generated in the same way as the datasets in Section 1.3 except that the matched normal BAM file for each tumor BAM file was also generated.

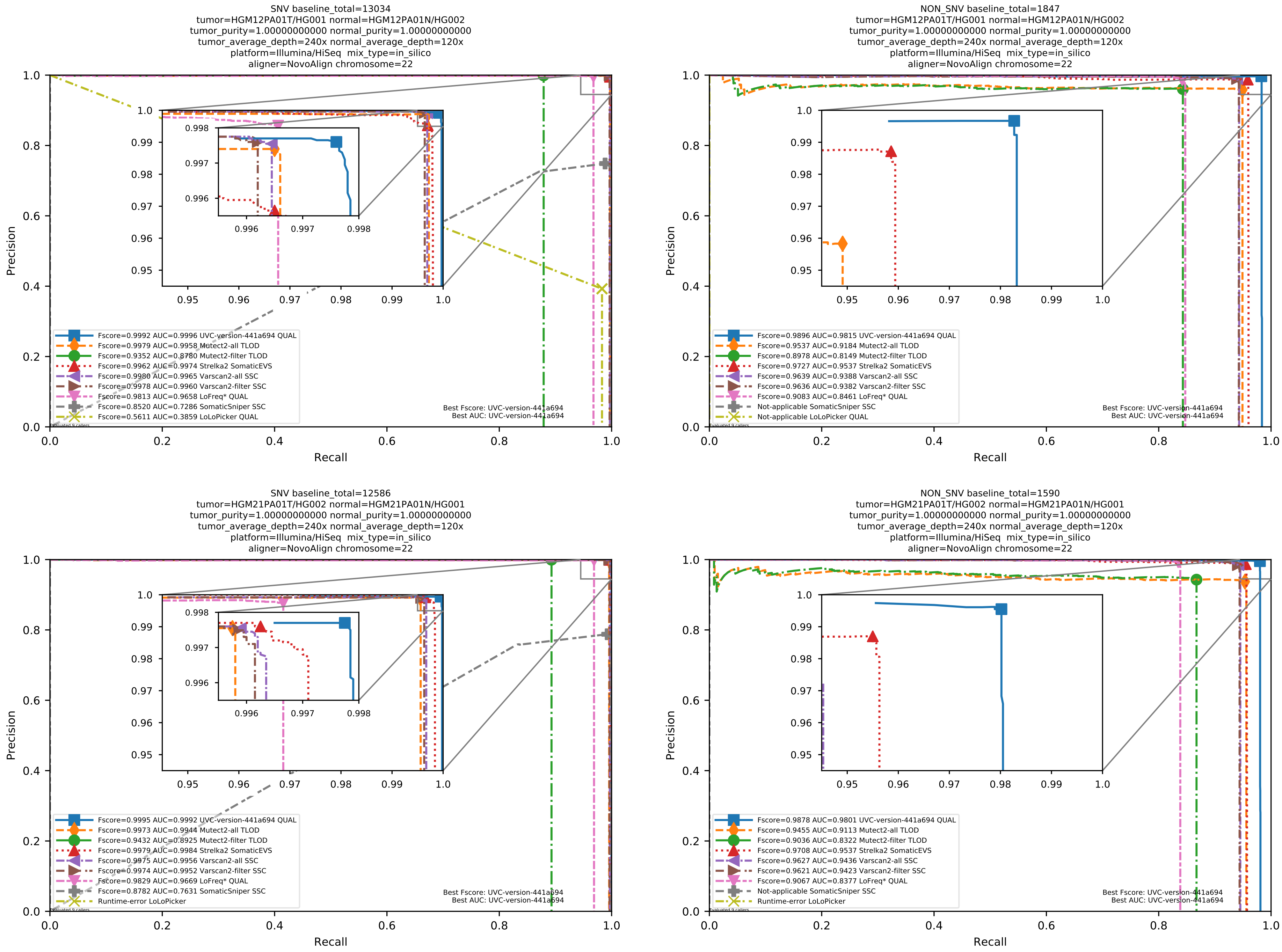

Supplementary Figure 41: Performance of somatic variant callers in tumor-with-normal mode on the *in silico* mixtures with Illumina HiSeq sequencing data pre-aligned with NovoAlign to the human reference genome hs37d5.

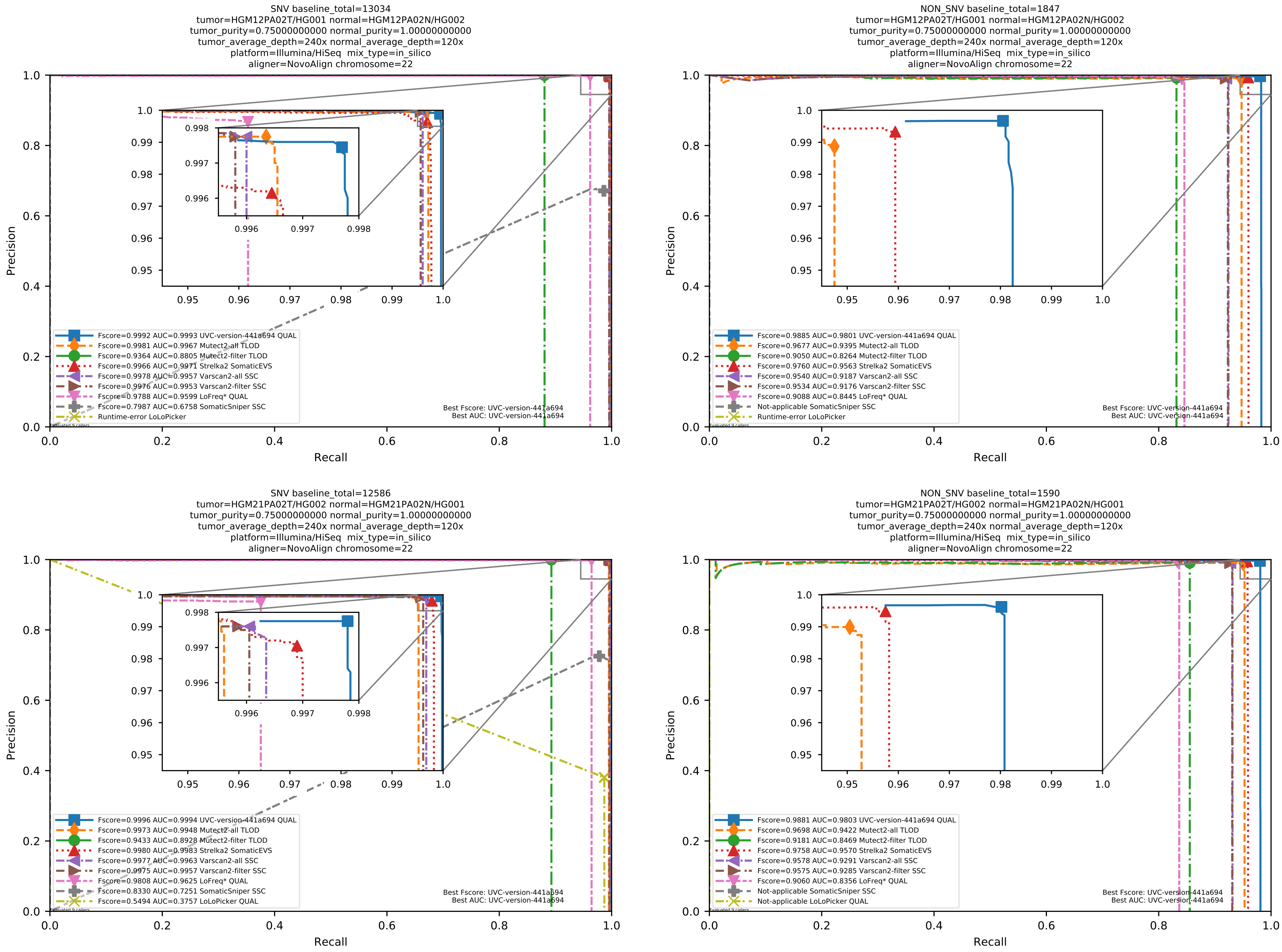

Supplementary Figure 42: Performance of somatic variant callers in tumor-with-normal mode on the *in silico* mixtures with Illumina HiSeq sequencing data pre-aligned with NovoAlign to the human reference genome hs37d5.

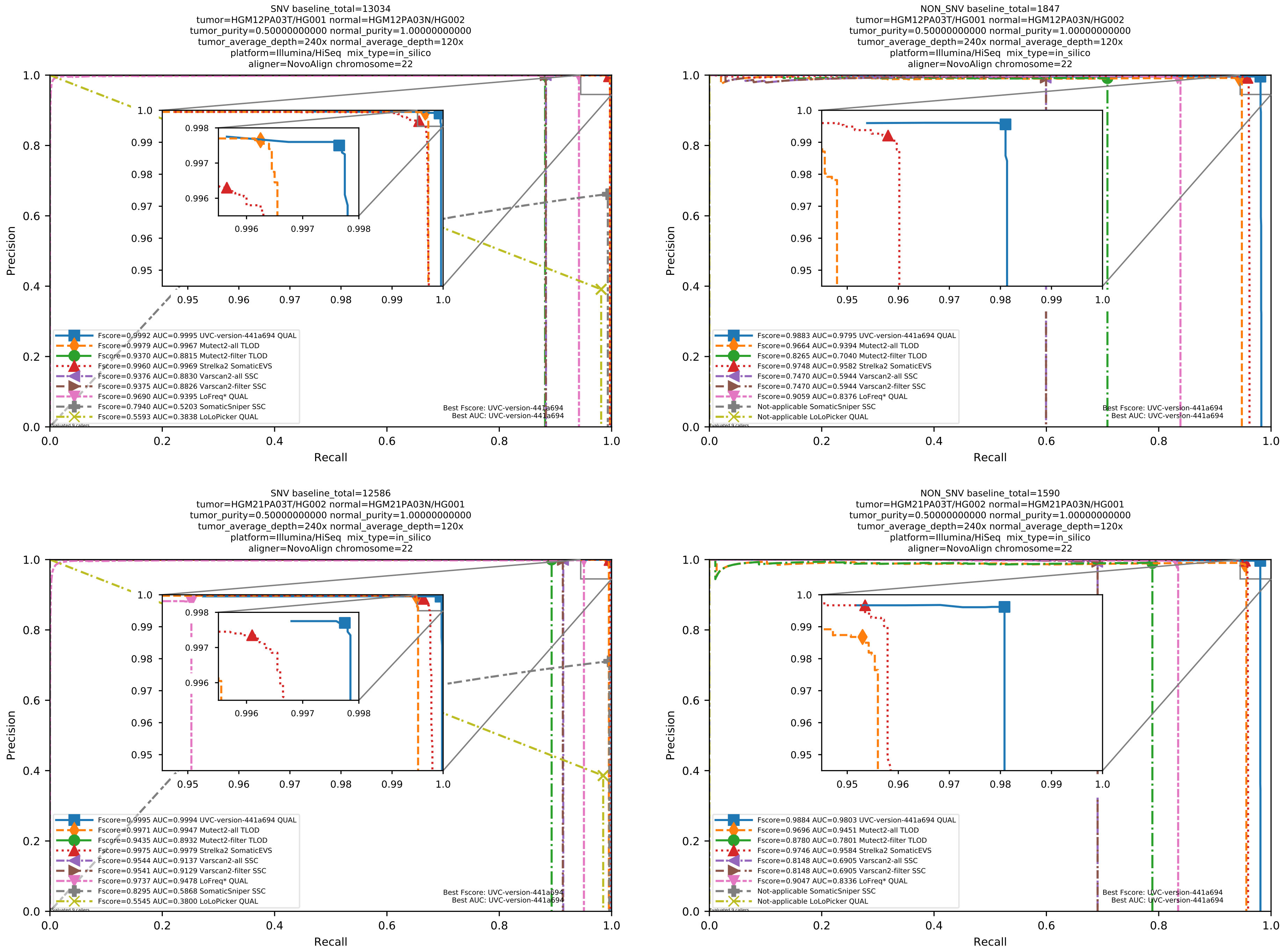

Supplementary Figure 43: Performance of somatic variant callers in tumor-with-normal mode on the *in silico* mixtures with Illumina HiSeq sequencing data pre-aligned with NovoAlign to the human reference genome hs37d5.

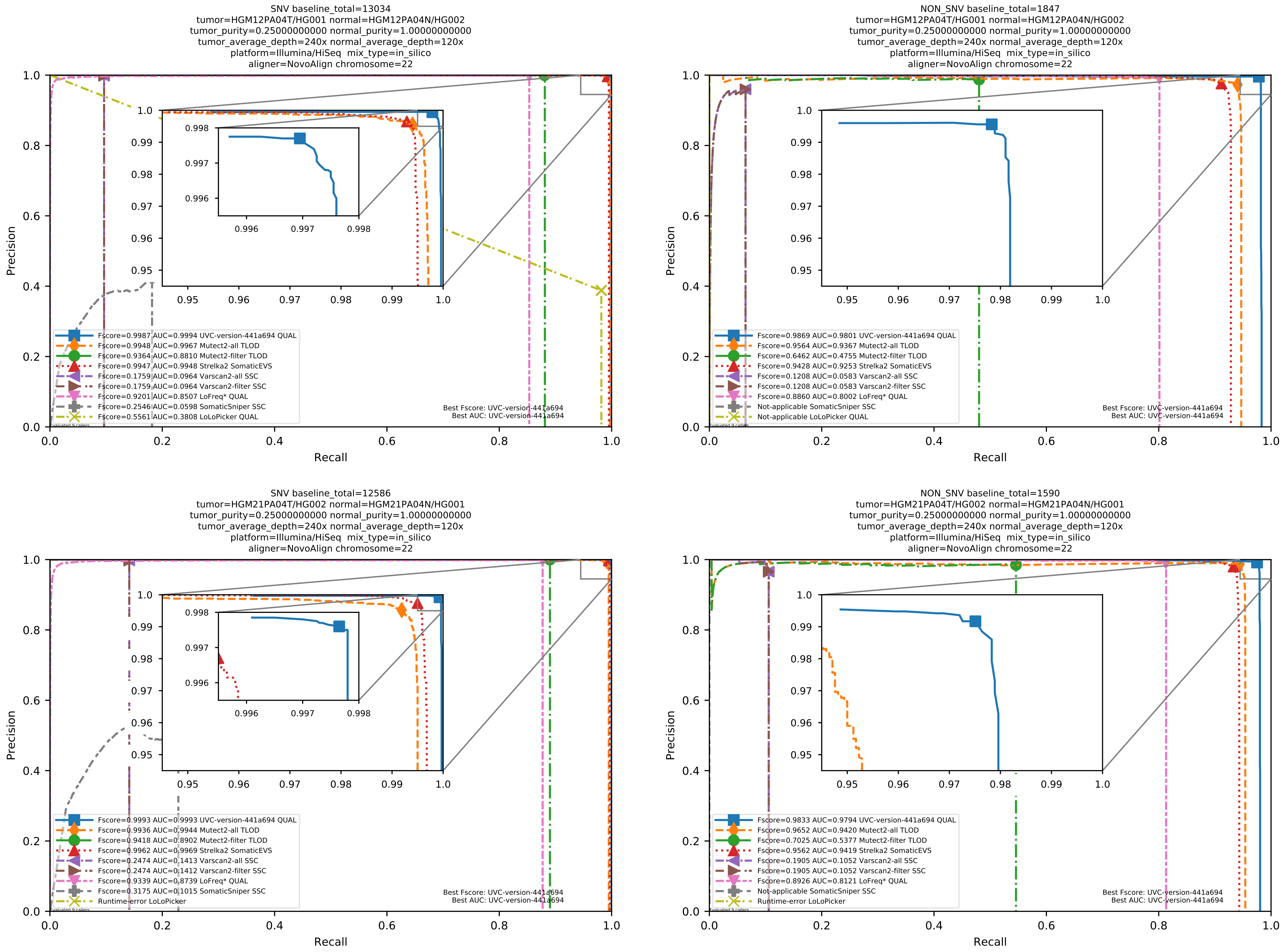

Supplementary Figure 44: Performance of somatic variant callers in tumor-with-normal mode on the *in silico* mixtures with Illumina HiSeq sequencing data pre-aligned with NovoAlign to the human reference genome hs37d5.

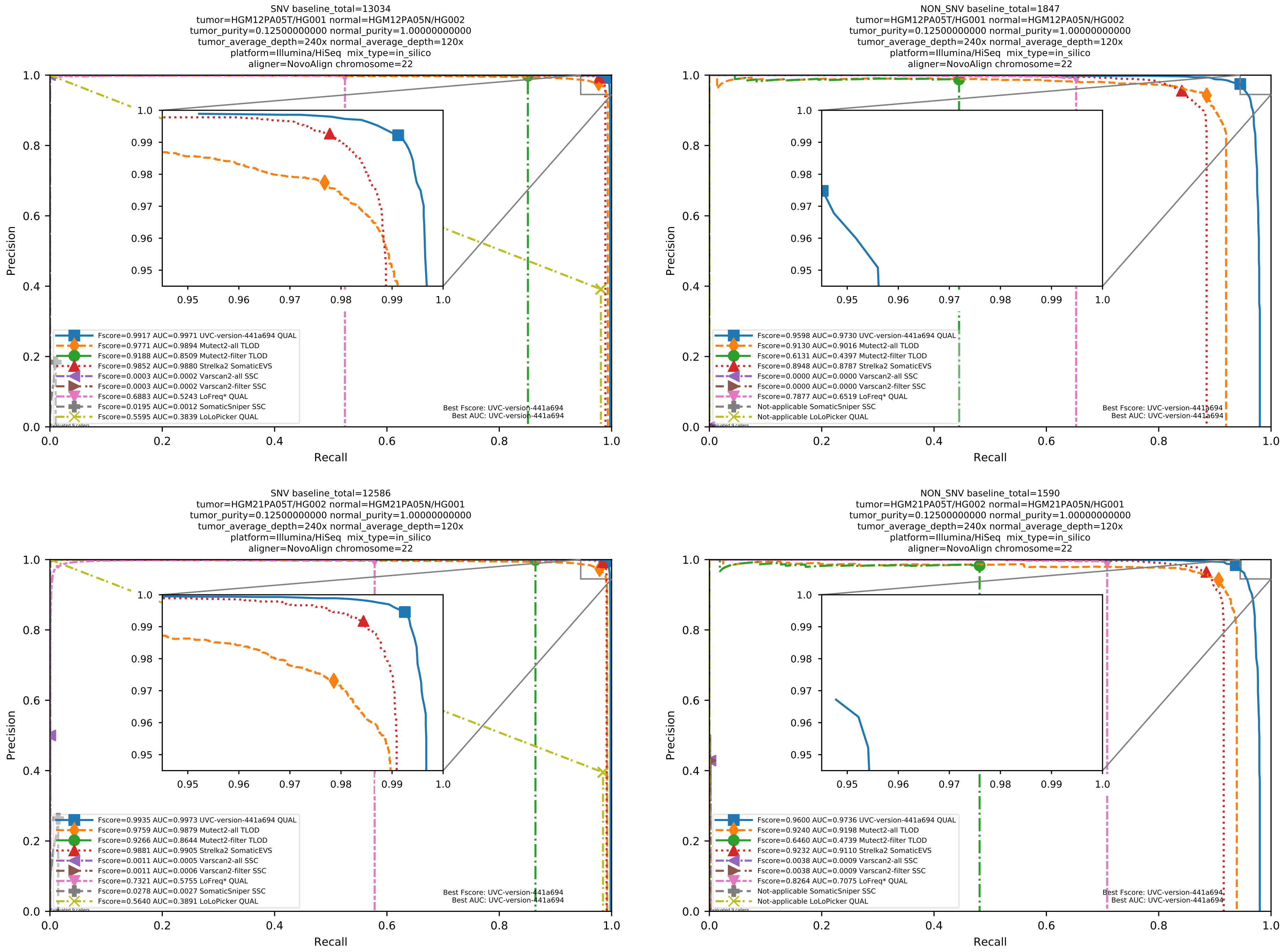

Supplementary Figure 45: Performance of somatic variant callers in tumor-with-normal mode on the *in silico* mixtures with Illumina HiSeq sequencing data pre-aligned with NovoAlign to the human reference genome hs37d5.

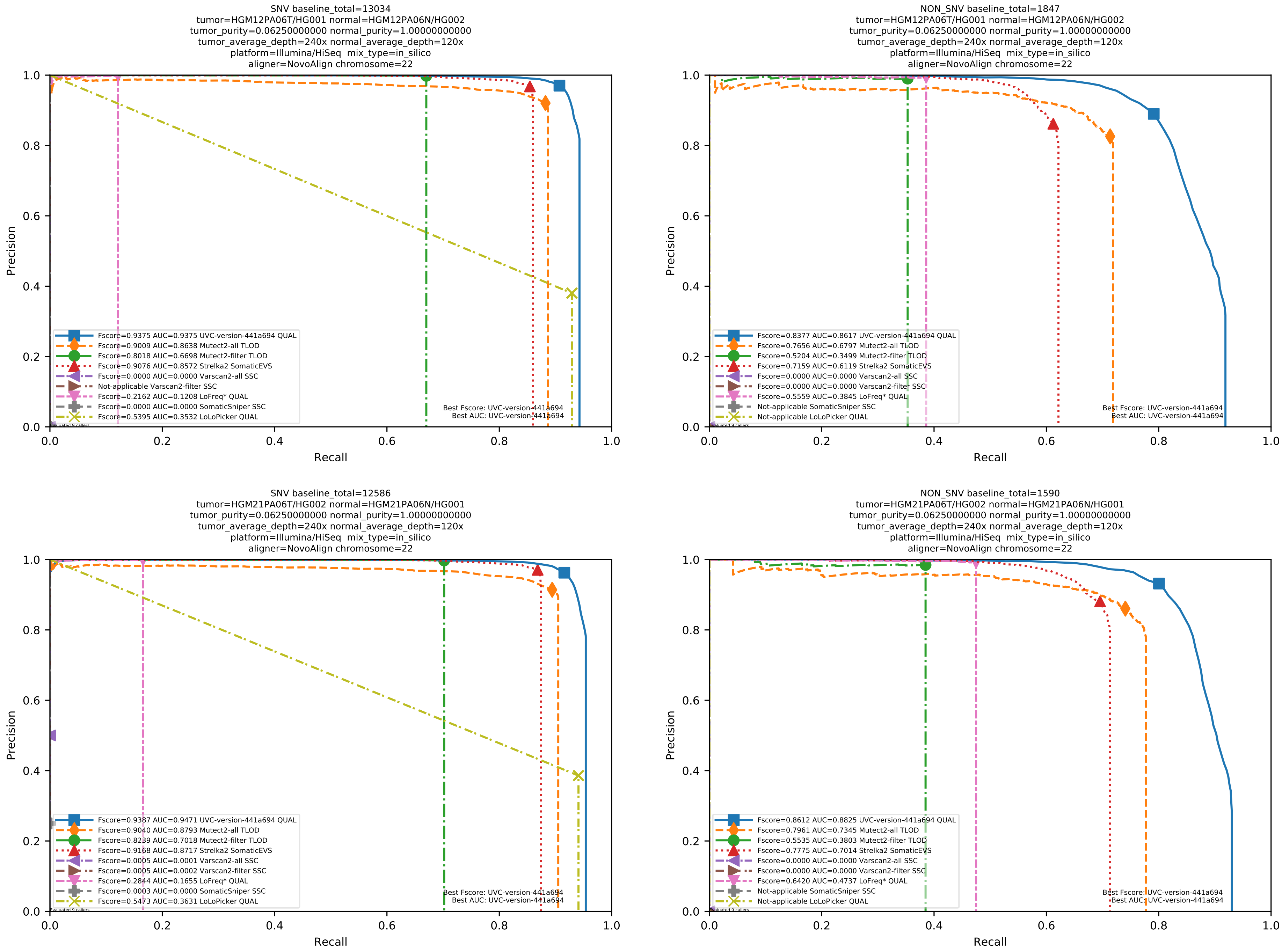

Supplementary Figure 46: Performance of somatic variant callers in tumor-with-normal mode on the *in silico* mixtures with Illumina HiSeq sequencing data pre-aligned with NovoAlign to the human reference genome hs37d5.

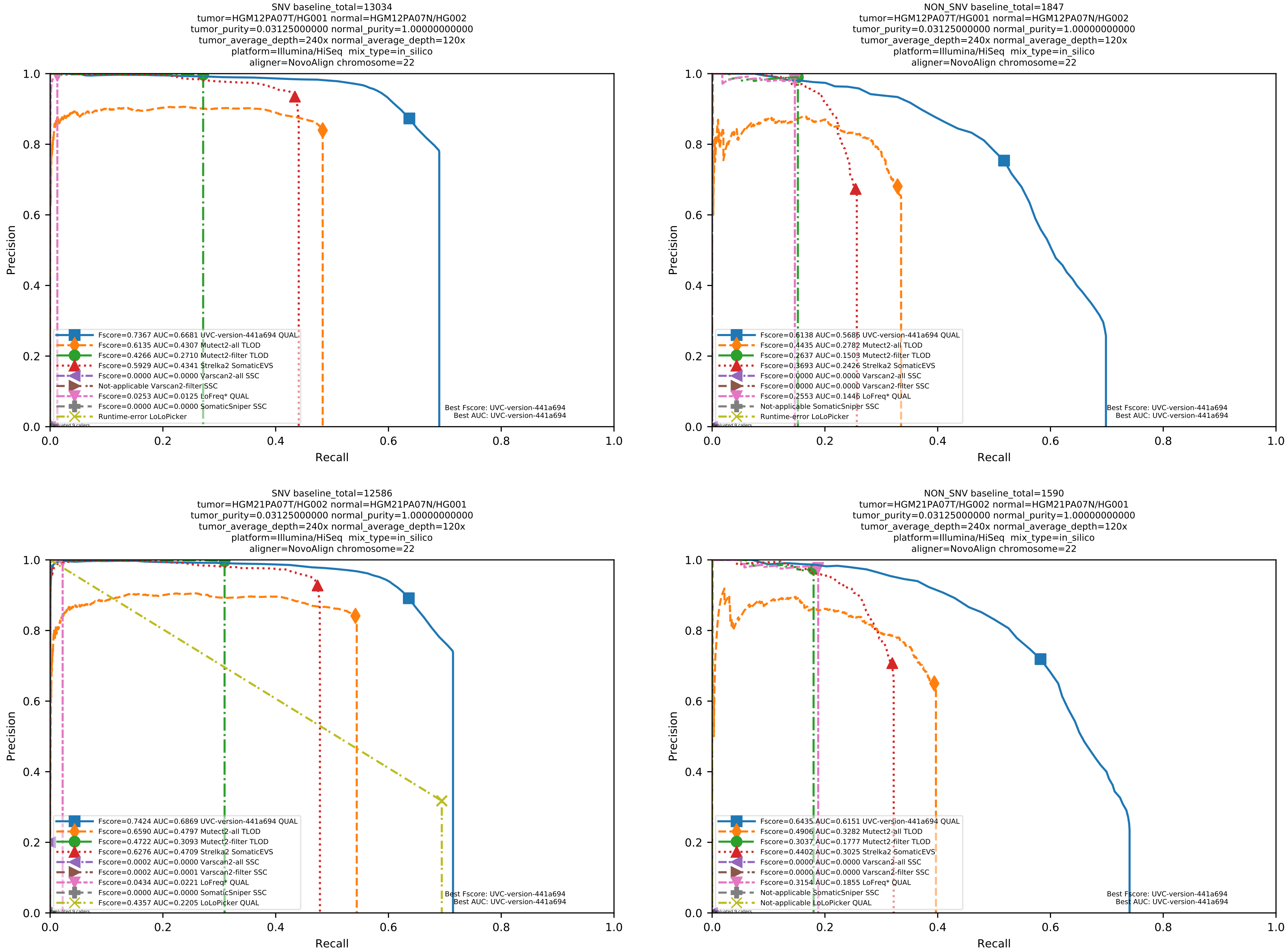

Supplementary Figure 47: Performance of somatic variant callers in tumor-with-normal mode on the *in silico* mixtures with Illumina HiSeq sequencing data pre-aligned with NovoAlign to the human reference genome hs37d5.

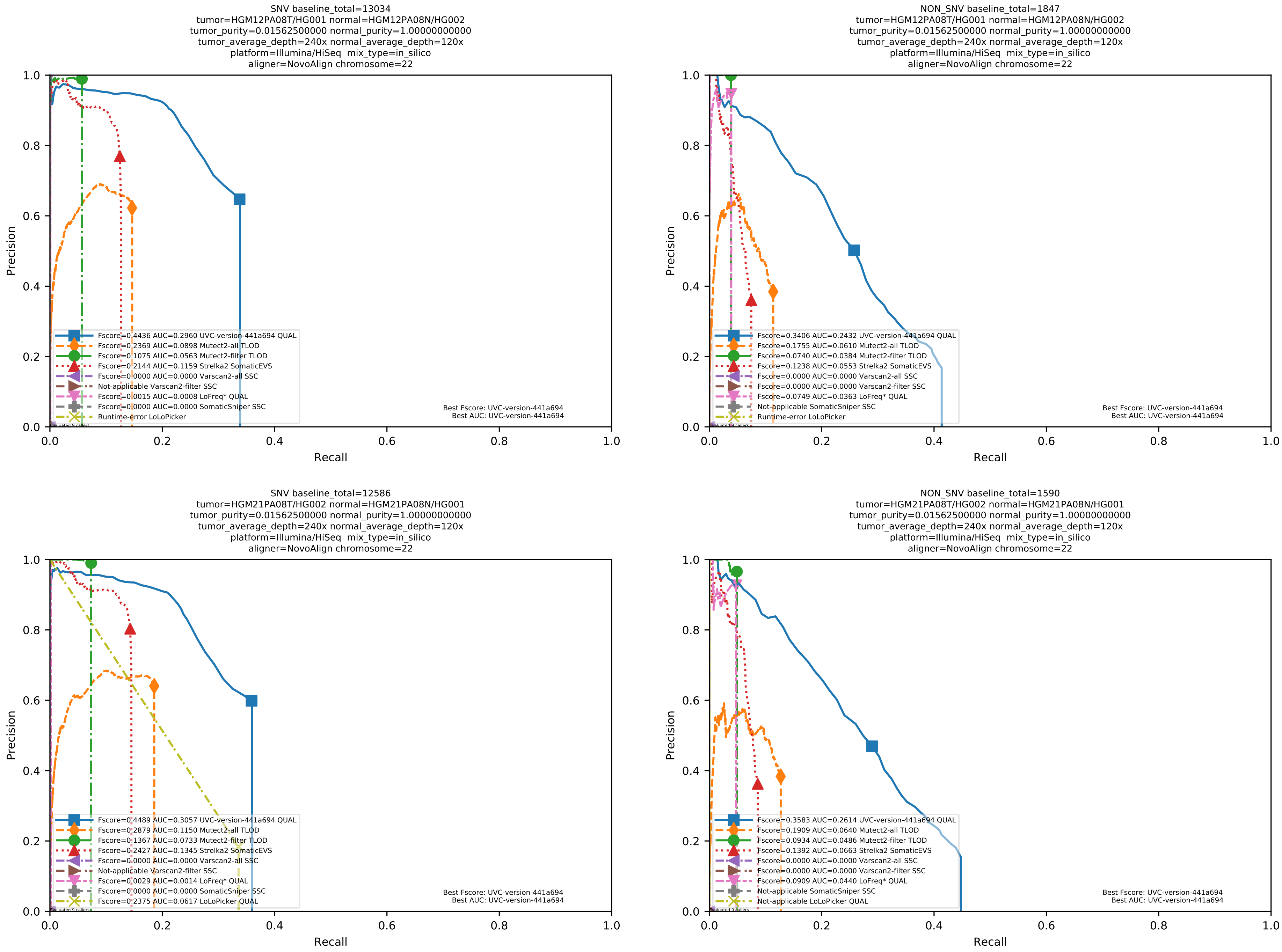

Supplementary Figure 48: Performance of somatic variant callers in tumor-with-normal mode on the *in silico* mixtures with Illumina HiSeq sequencing data pre-aligned with NovoAlign to the human reference genome hs37d5.

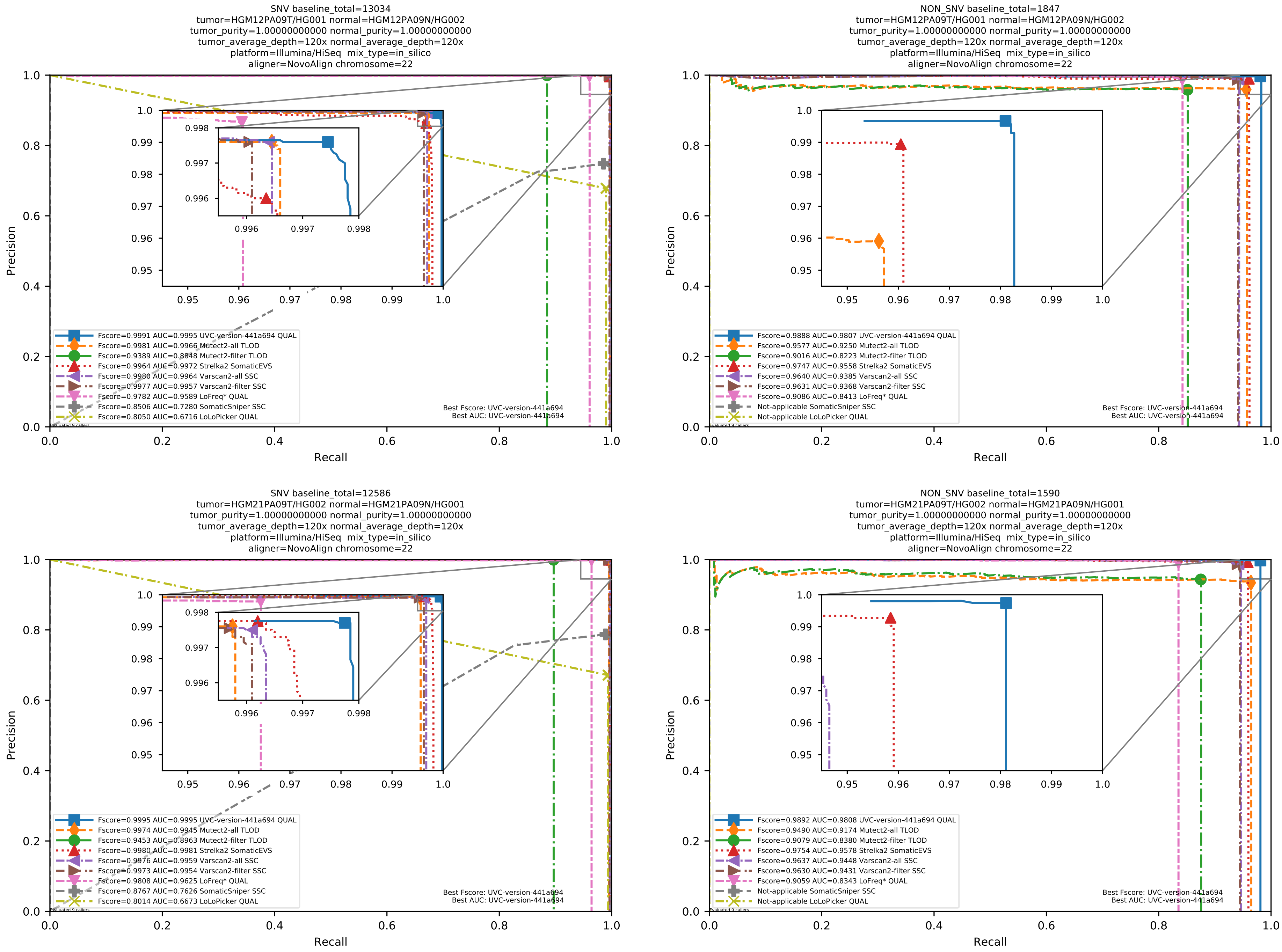

Supplementary Figure 49: Performance of somatic variant callers in tumor-with-normal mode on the *in silico* mixtures with Illumina HiSeq sequencing data pre-aligned with NovoAlign to the human reference genome hs37d5.

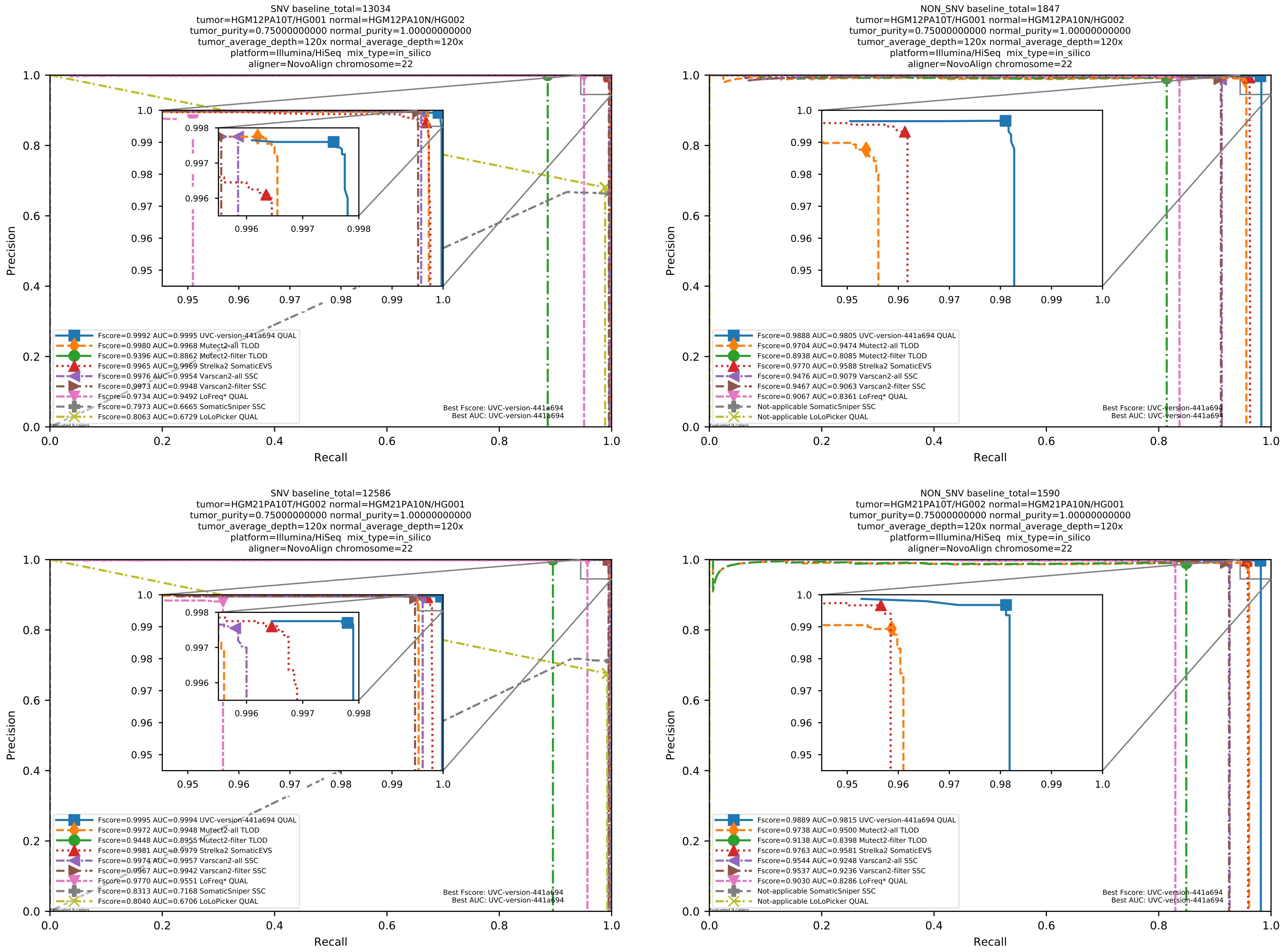

Supplementary Figure 50: Performance of somatic variant callers in tumor-with-normal mode on the *in silico* mixtures with Illumina HiSeq sequencing data pre-aligned with NovoAlign to the human reference genome hs37d5.

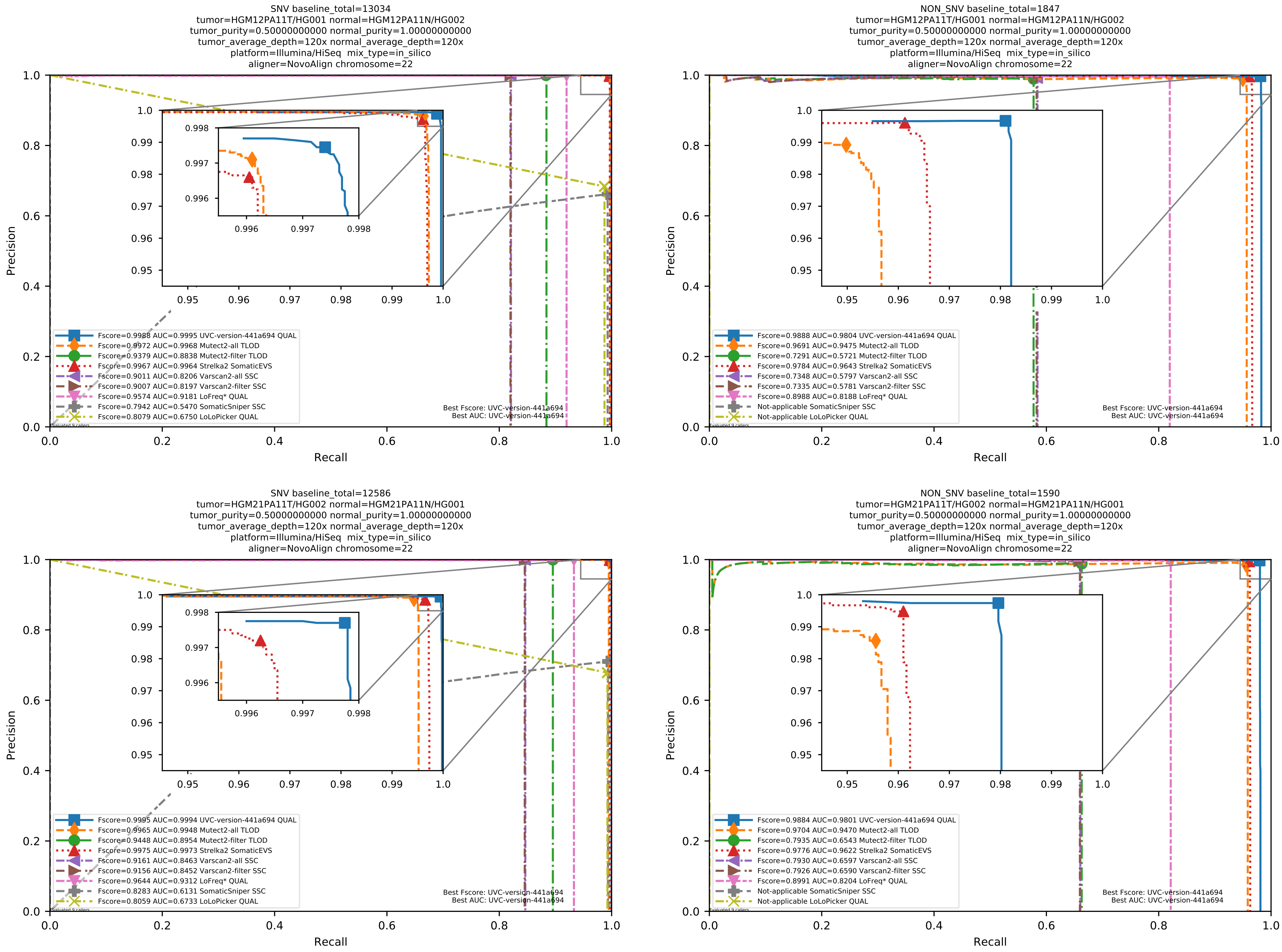

Supplementary Figure 51: Performance of somatic variant callers in tumor-with-normal mode on the *in silico* mixtures with Illumina HiSeq sequencing data pre-aligned with NovoAlign to the human reference genome hs37d5.

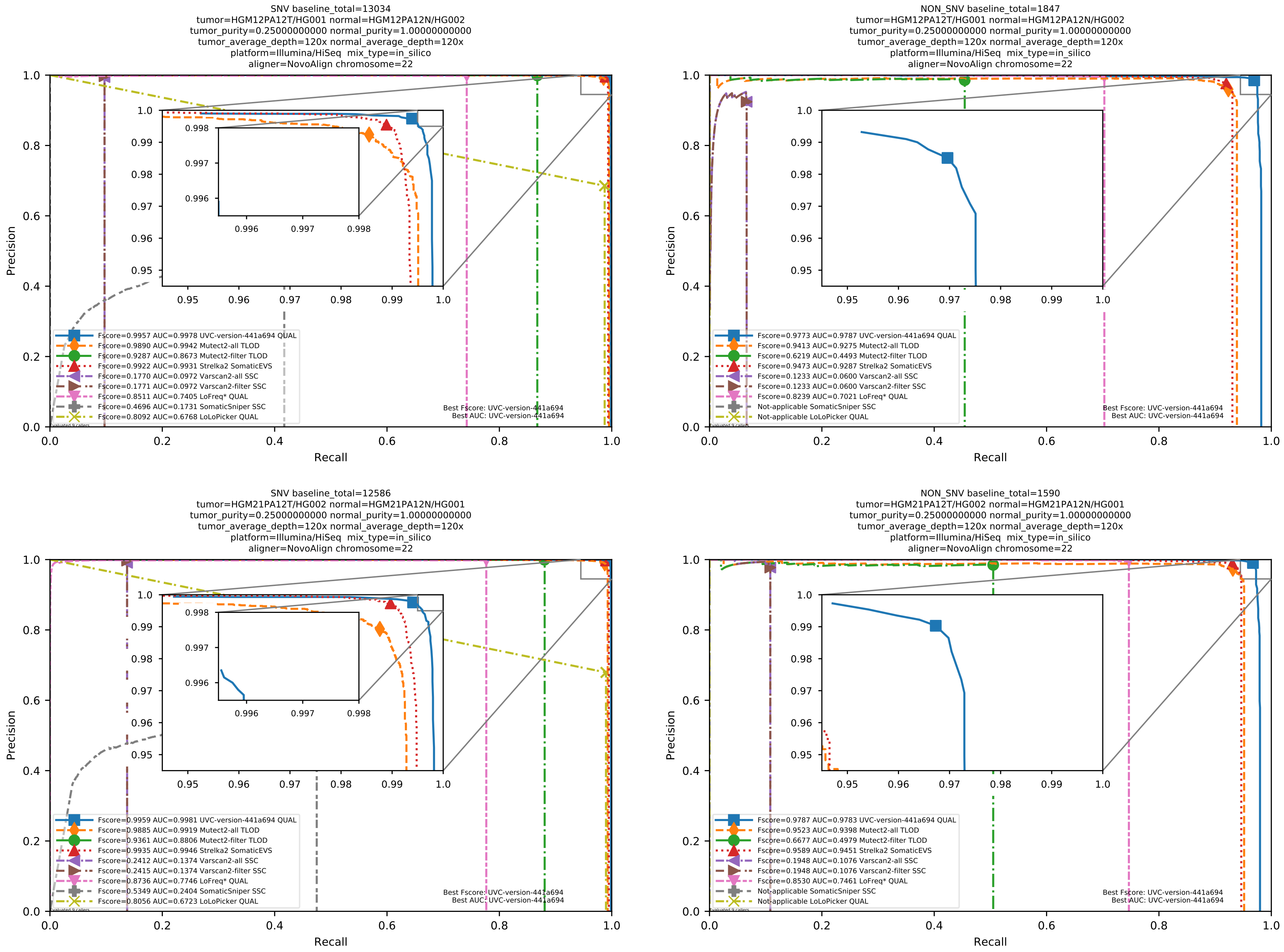

Supplementary Figure 52: Performance of somatic variant callers in tumor-with-normal mode on the *in silico* mixtures with Illumina HiSeq sequencing data pre-aligned with NovoAlign to the human reference genome hs37d5.

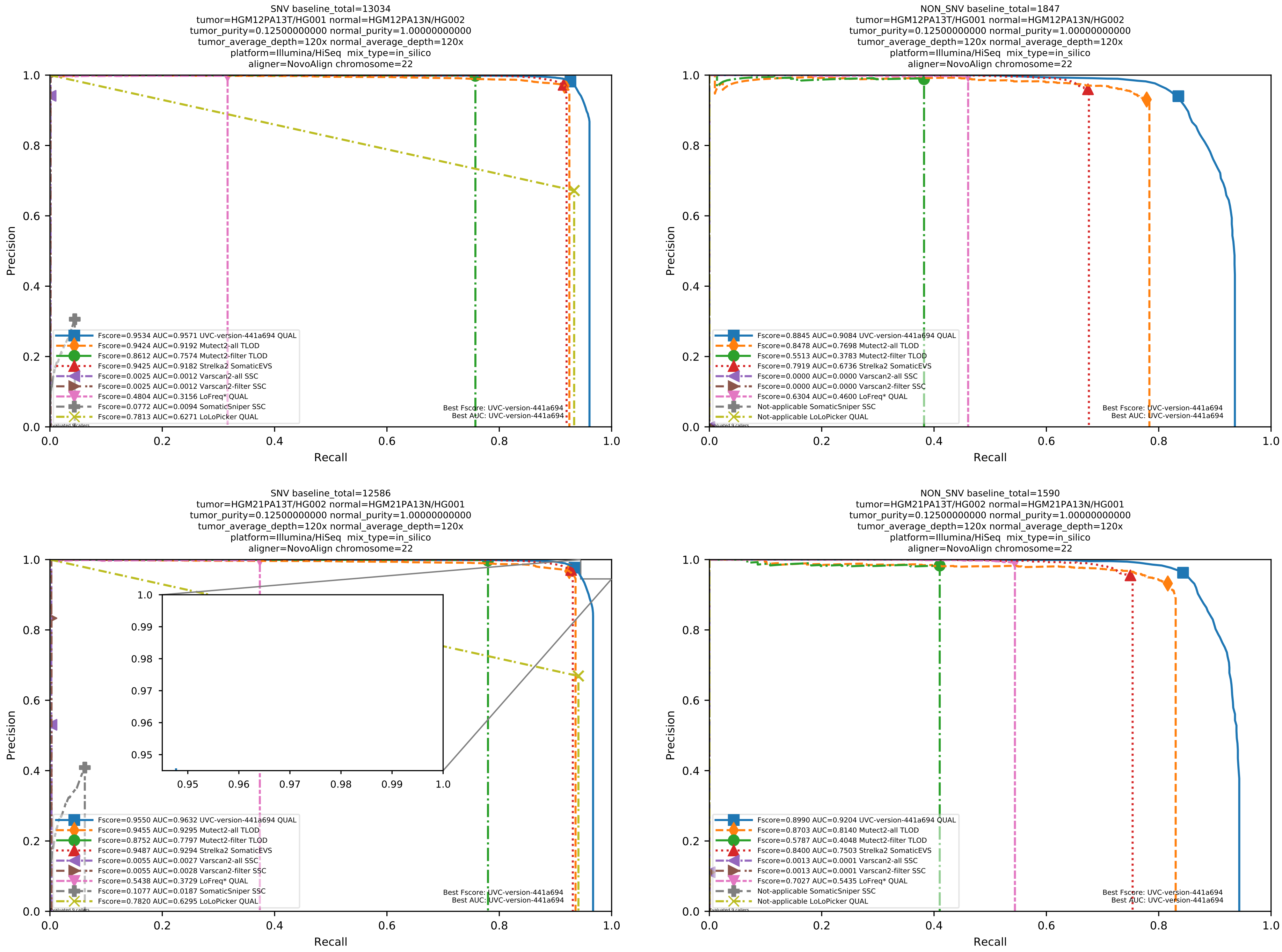

Supplementary Figure 53: Performance of somatic variant callers in tumor-with-normal mode on the *in silico* mixtures with Illumina HiSeq sequencing data pre-aligned with NovoAlign to the human reference genome hs37d5.

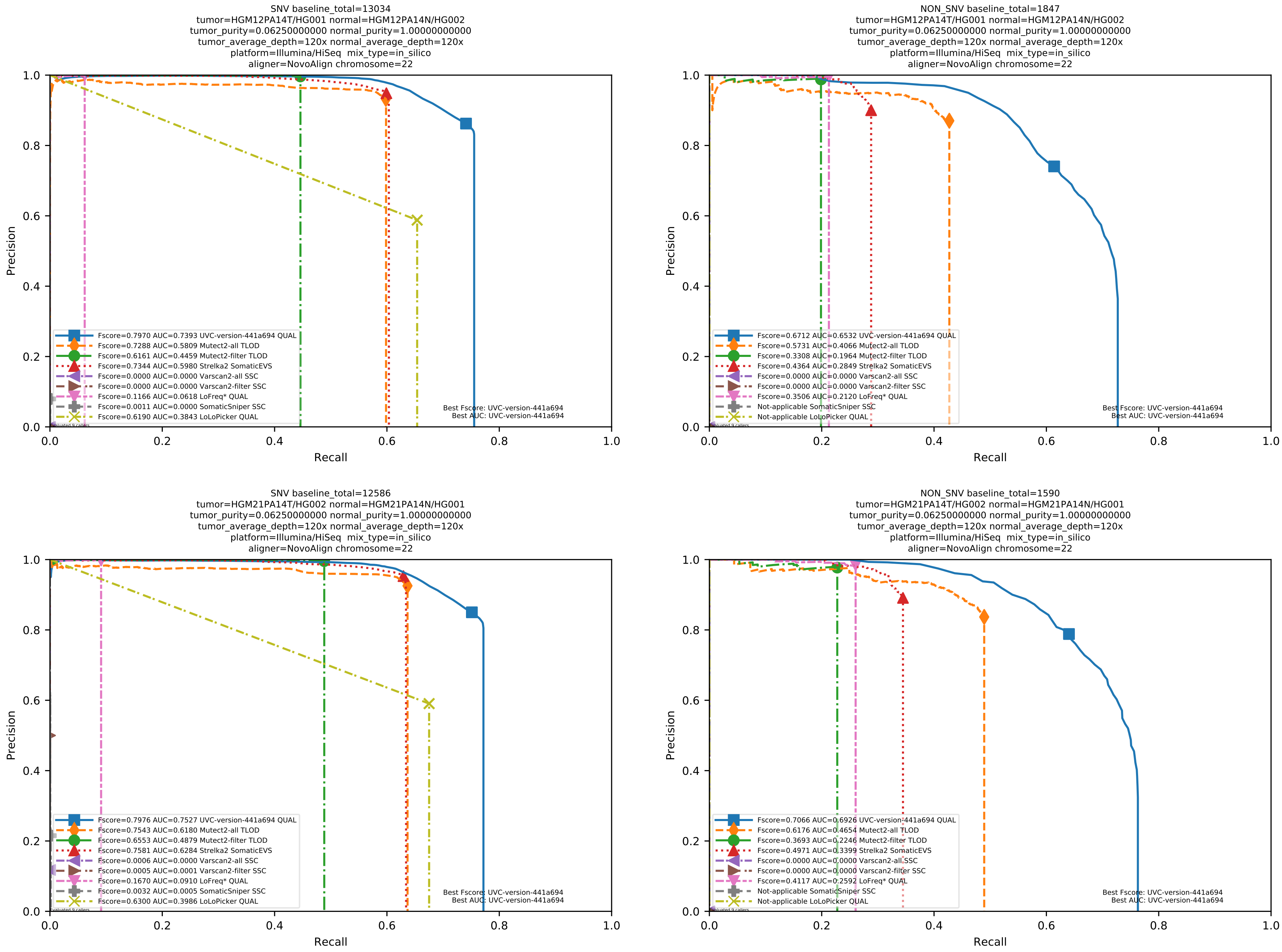

Supplementary Figure 54: Performance of somatic variant callers in tumor-with-normal mode on the *in silico* mixtures with Illumina HiSeq sequencing data pre-aligned with NovoAlign to the human reference genome hs37d5.

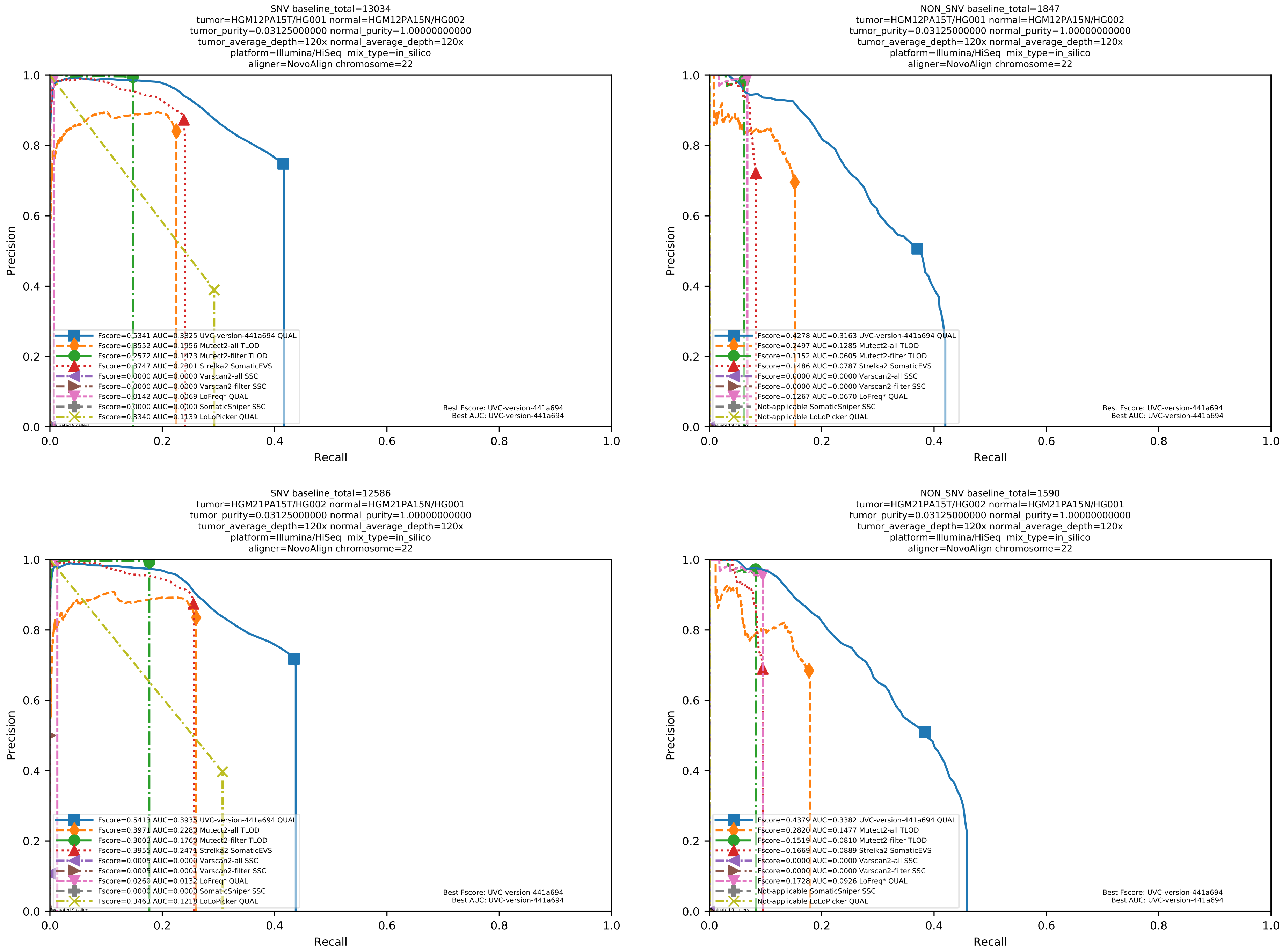

Supplementary Figure 55: Performance of somatic variant callers in tumor-with-normal mode on the *in silico* mixtures with Illumina HiSeq sequencing data pre-aligned with NovoAlign to the human reference genome hs37d5.

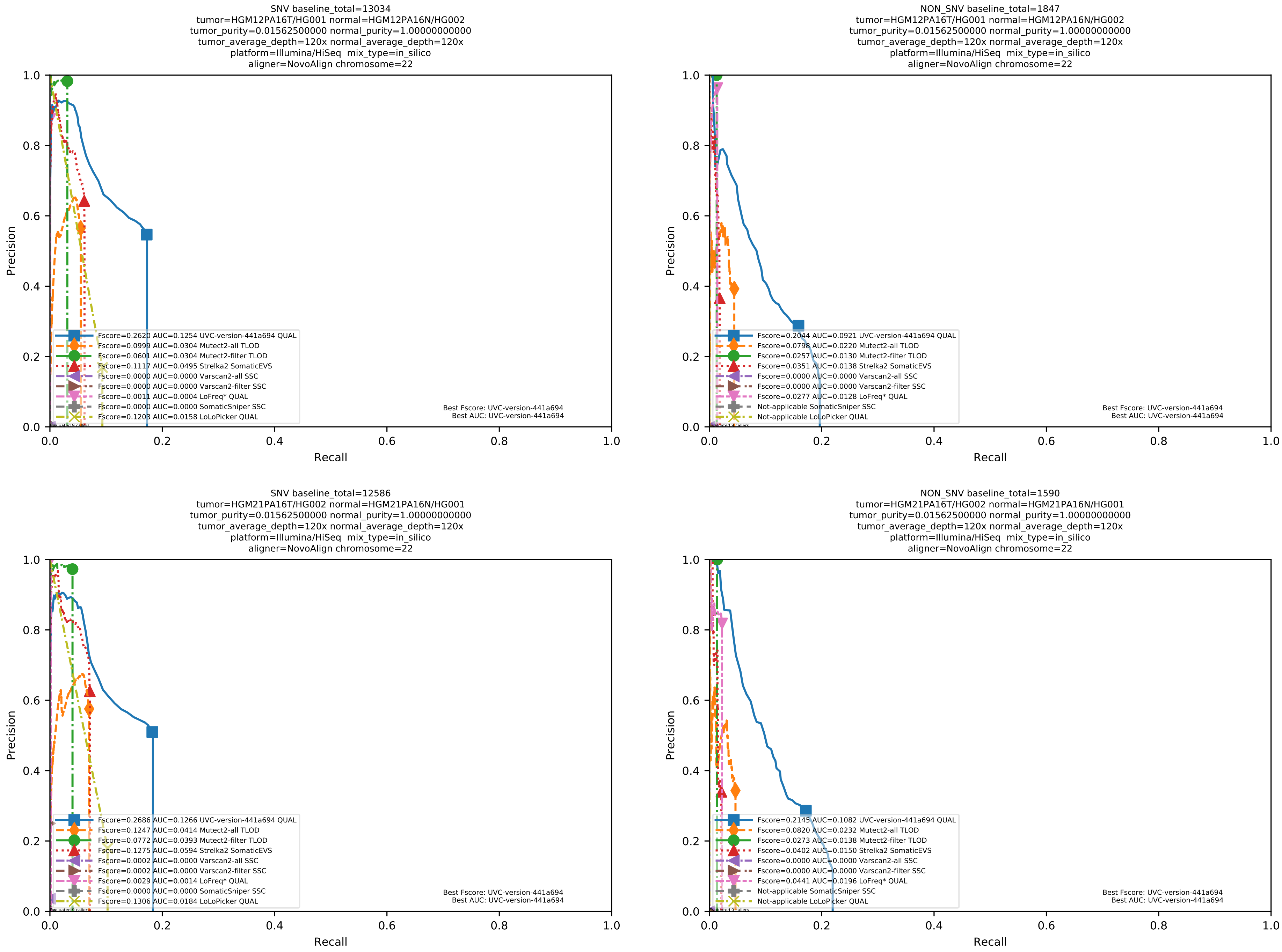

Supplementary Figure 56: Performance of somatic variant callers in tumor-with-normal mode on the *in silico* mixtures with Illumina HiSeq sequencing data pre-aligned with NovoAlign to the human reference genome hs37d5.

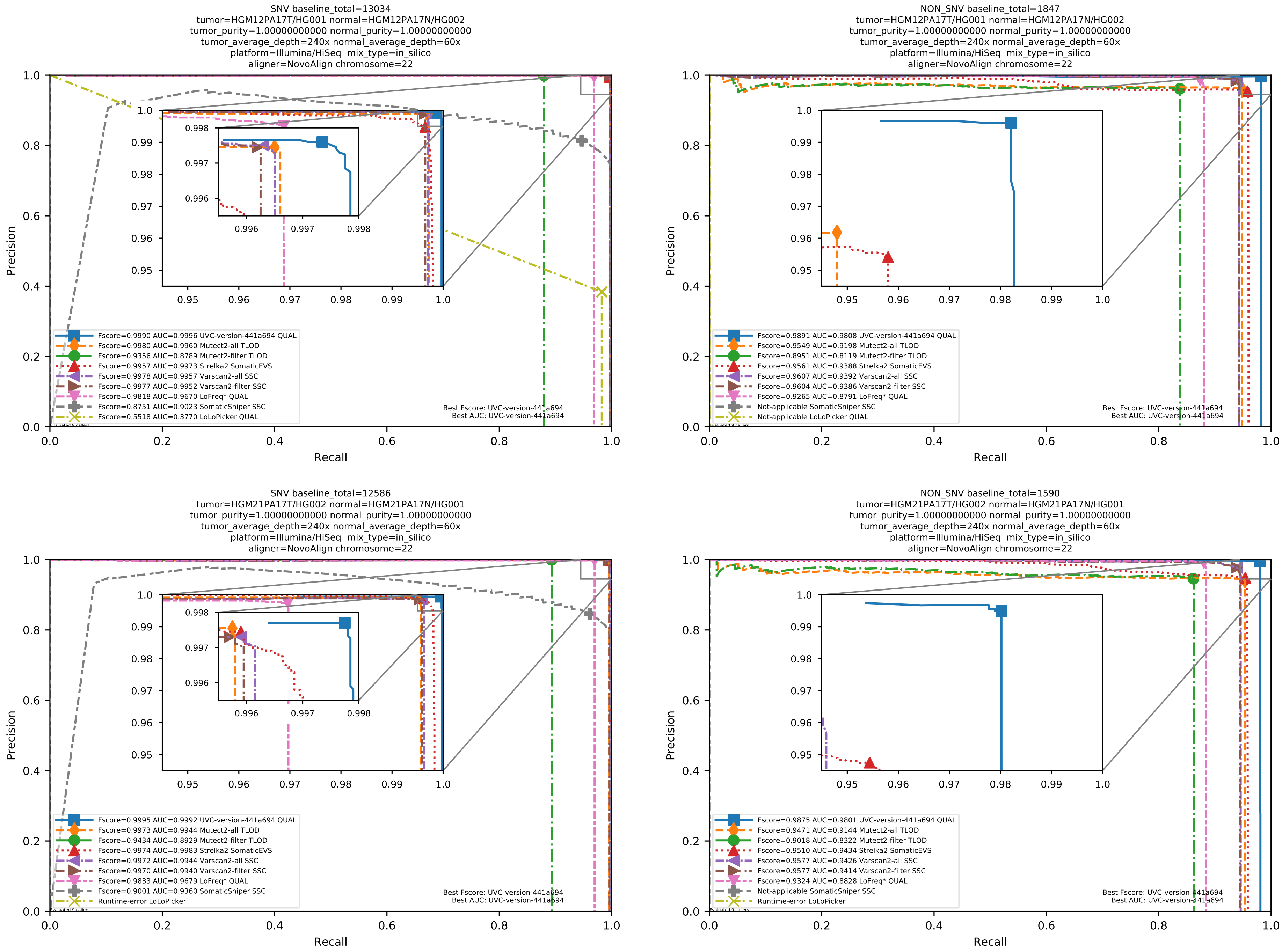

Supplementary Figure 57: Performance of somatic variant callers in tumor-with-normal mode on the *in silico* mixtures with Illumina HiSeq sequencing data pre-aligned with NovoAlign to the human reference genome hs37d5.

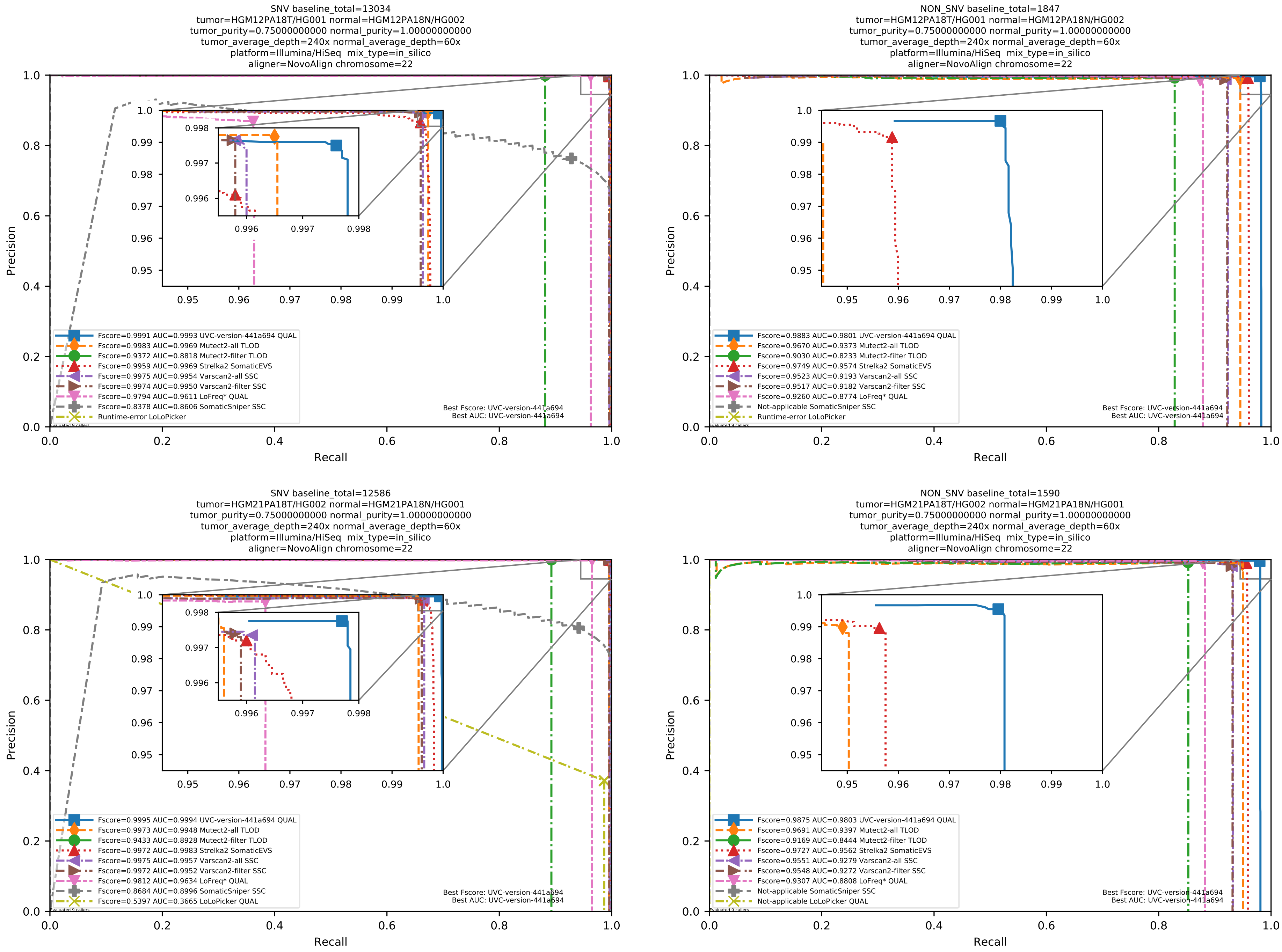

Supplementary Figure 58: Performance of somatic variant callers in tumor-with-normal mode on the *in silico* mixtures with Illumina HiSeq sequencing data pre-aligned with NovoAlign to the human reference genome hs37d5.

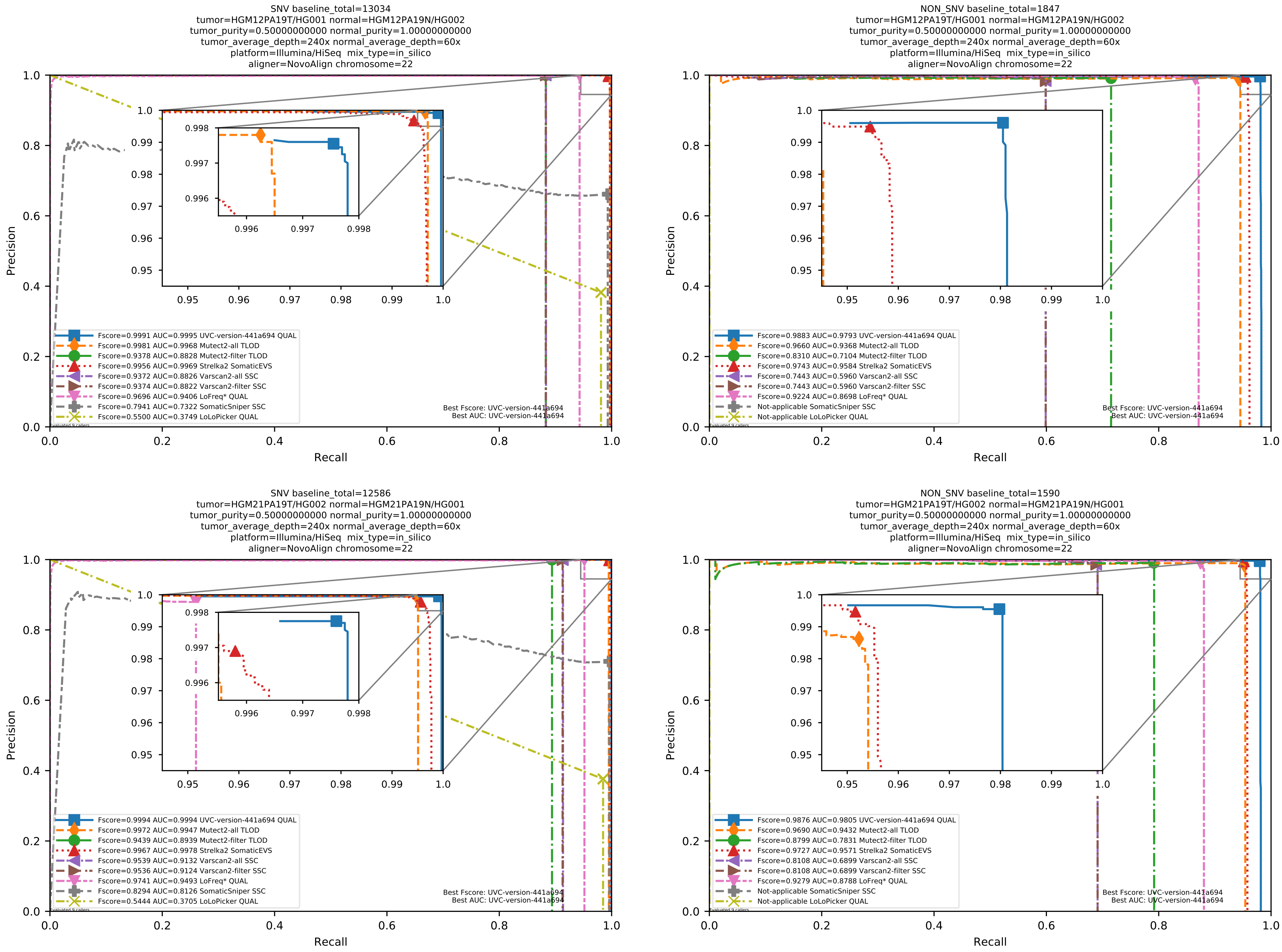

Supplementary Figure 59: Performance of somatic variant callers in tumor-with-normal mode on the *in silico* mixtures with Illumina HiSeq sequencing data pre-aligned with NovoAlign to the human reference genome hs37d5.

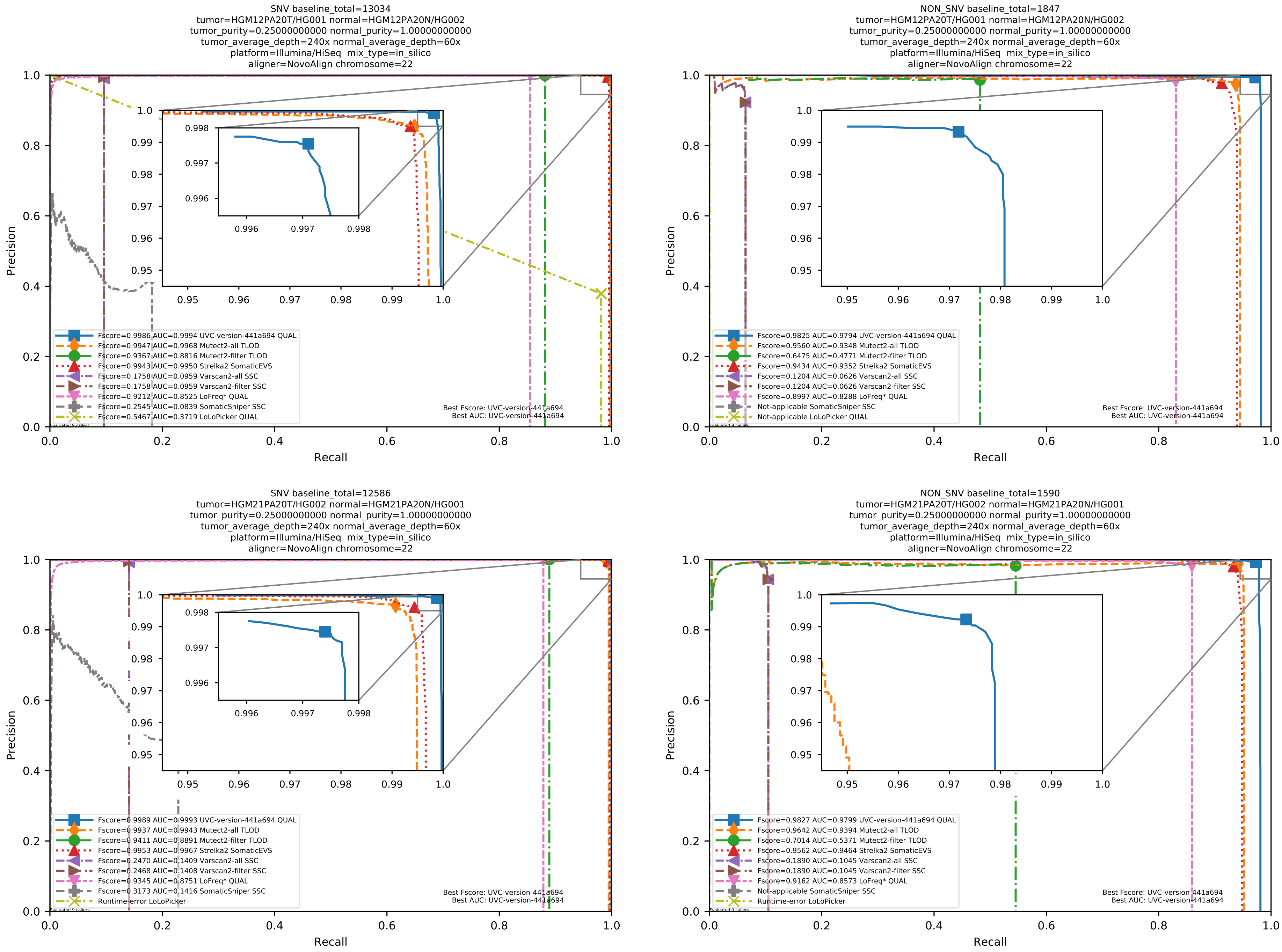

Supplementary Figure 60: Performance of somatic variant callers in tumor-with-normal mode on the *in silico* mixtures with Illumina HiSeq sequencing data pre-aligned with NovoAlign to the human reference genome hs37d5.

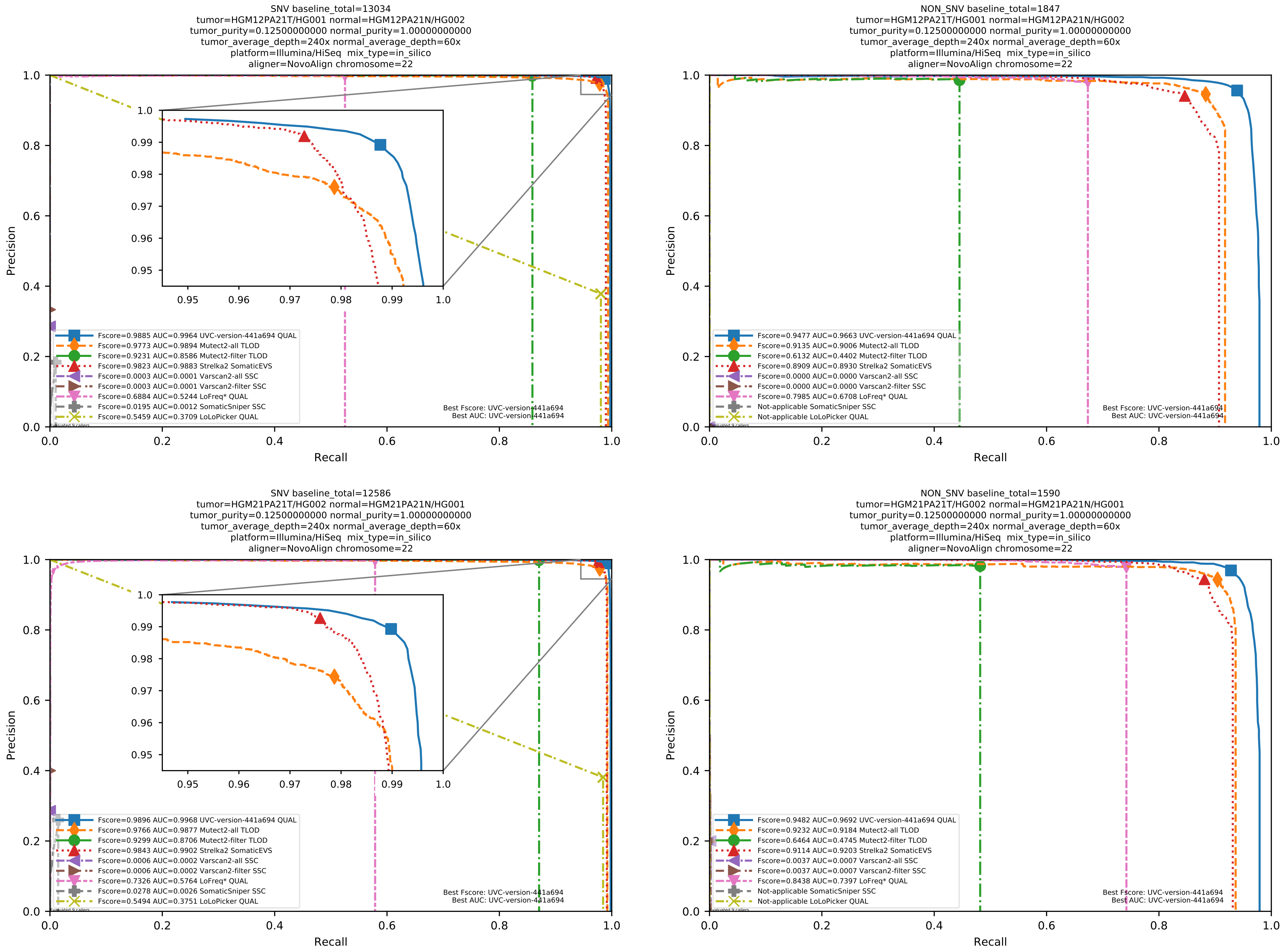

Supplementary Figure 61: Performance of somatic variant callers in tumor-with-normal mode on the *in silico* mixtures with Illumina HiSeq sequencing data pre-aligned with NovoAlign to the human reference genome hs37d5.

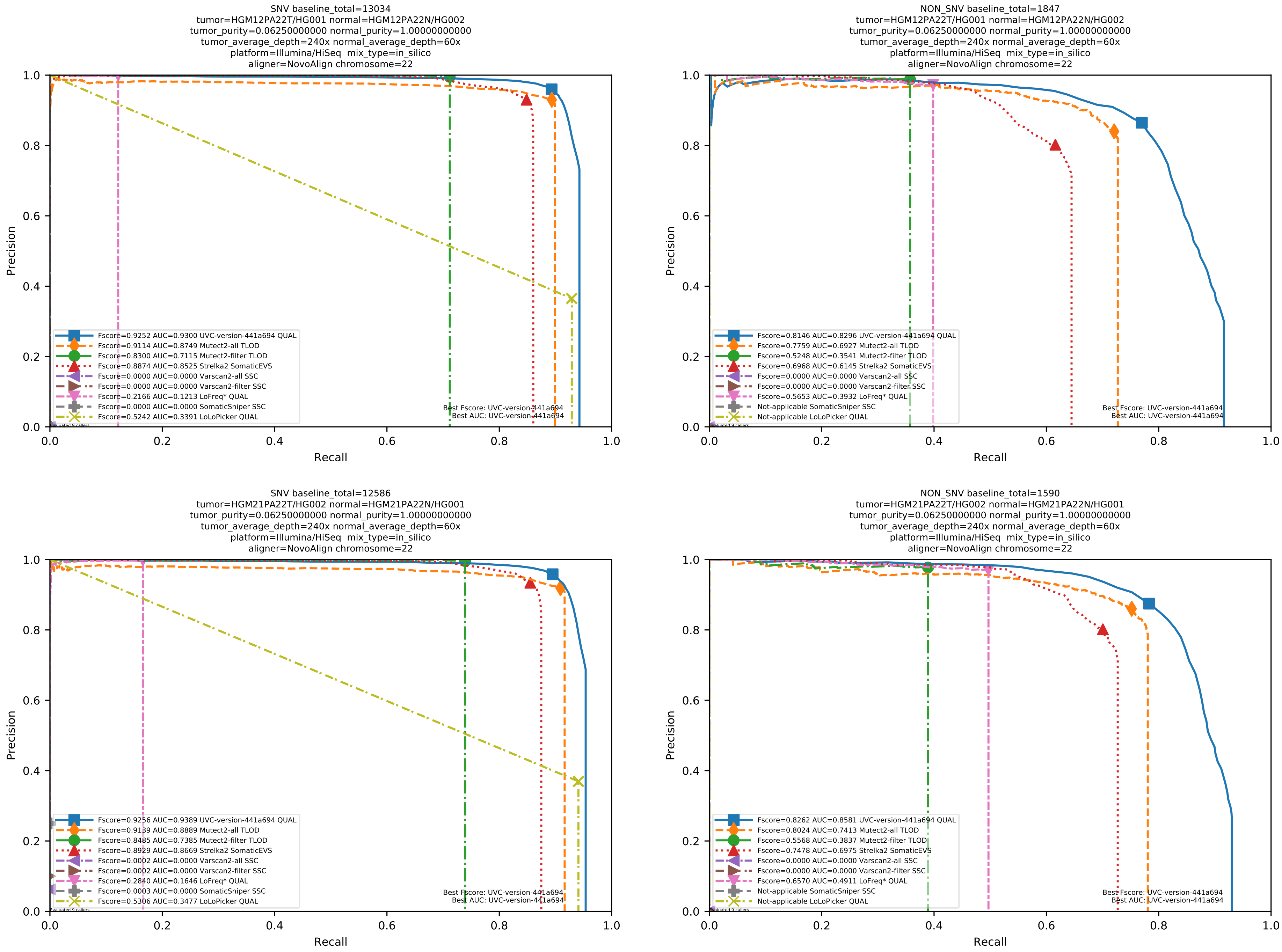

Supplementary Figure 62: Performance of somatic variant callers in tumor-with-normal mode on the *in silico* mixtures with Illumina HiSeq sequencing data pre-aligned with NovoAlign to the human reference genome hs37d5.

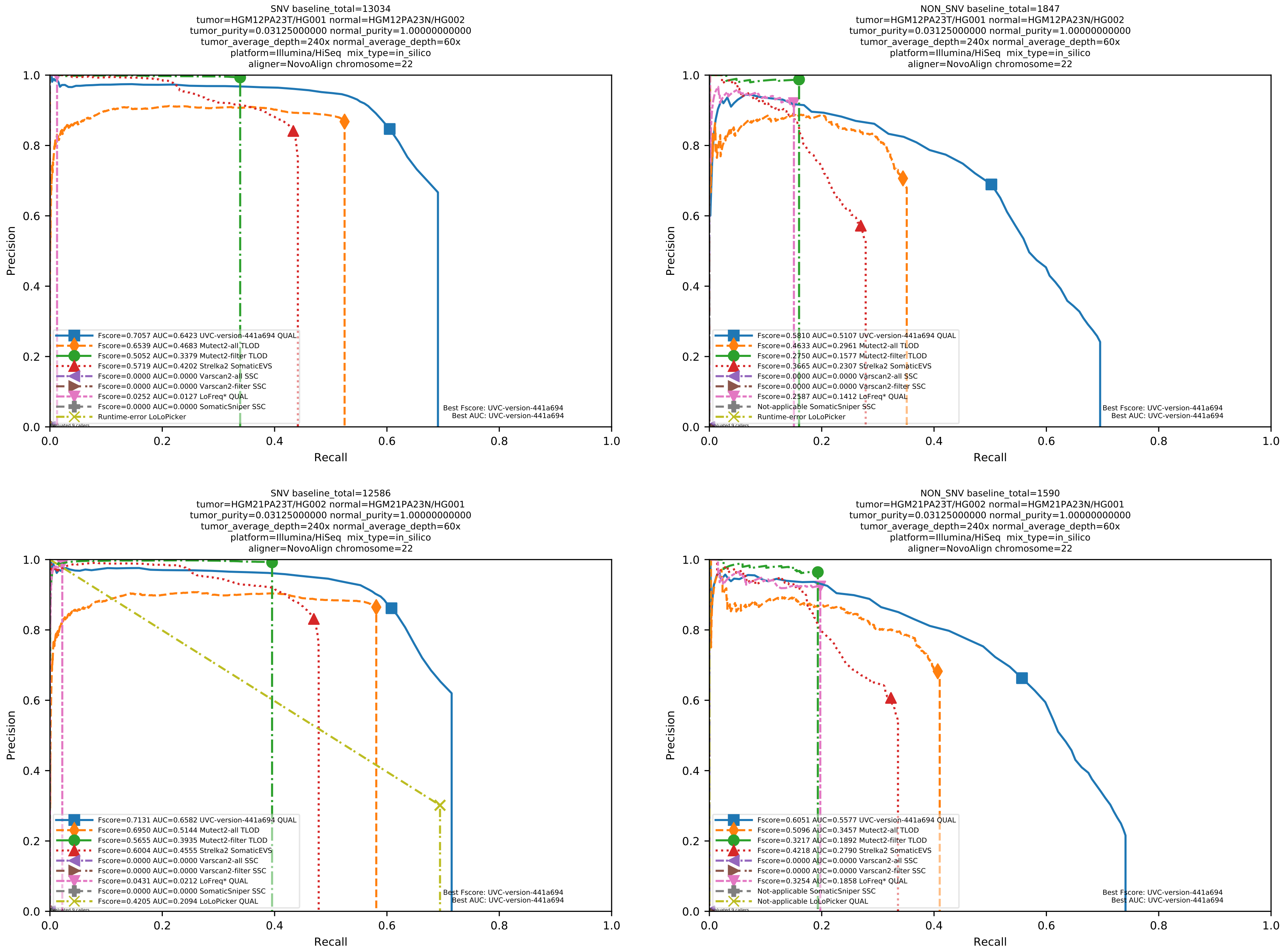

Supplementary Figure 63: Performance of somatic variant callers in tumor-with-normal mode on the *in silico* mixtures with Illumina HiSeq sequencing data pre-aligned with NovoAlign to the human reference genome hs37d5.

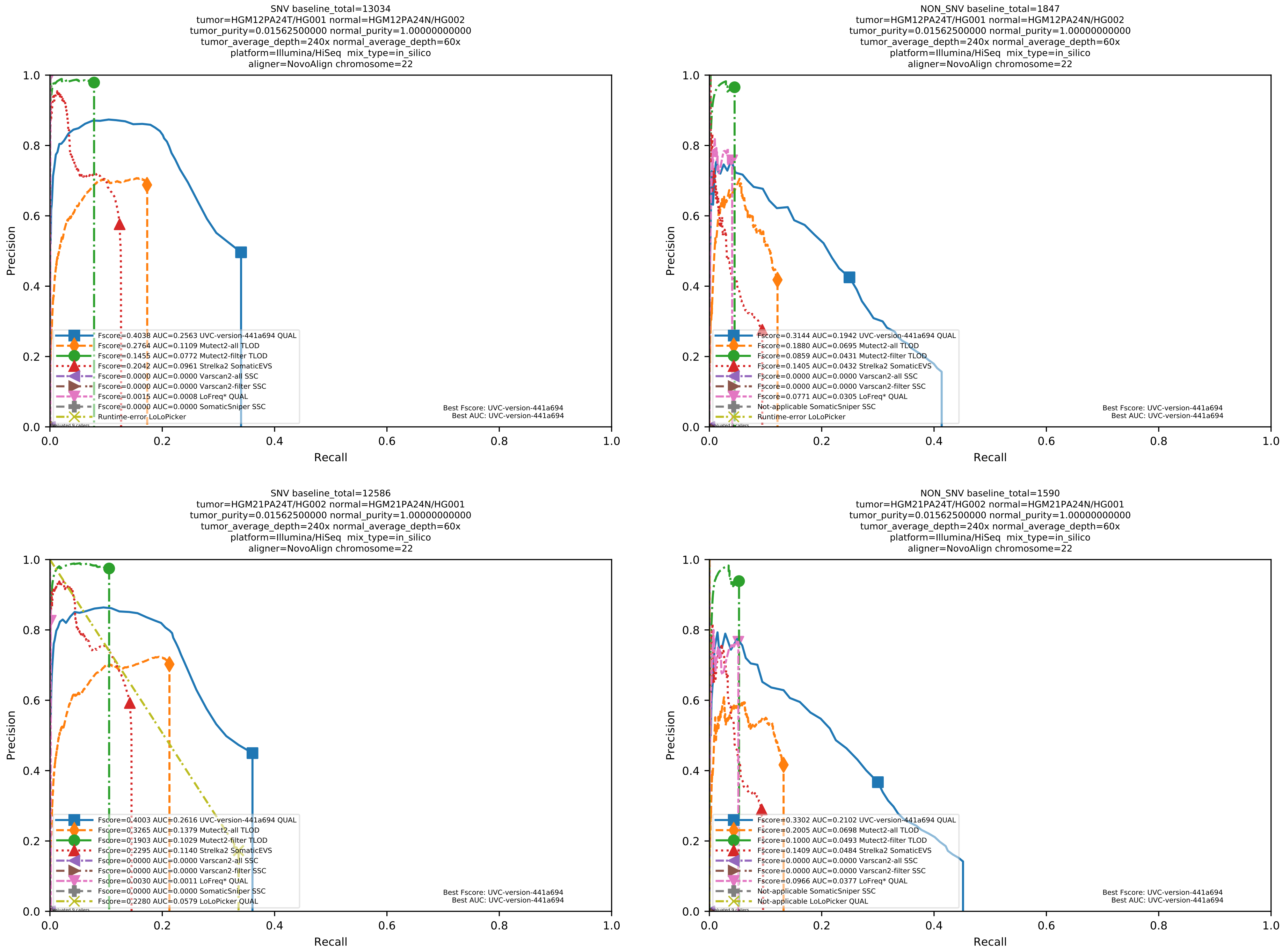

Supplementary Figure 64: Performance of somatic variant callers in tumor-with-normal mode on the *in silico* mixtures with Illumina HiSeq sequencing data pre-aligned with NovoAlign to the human reference genome hs37d5.

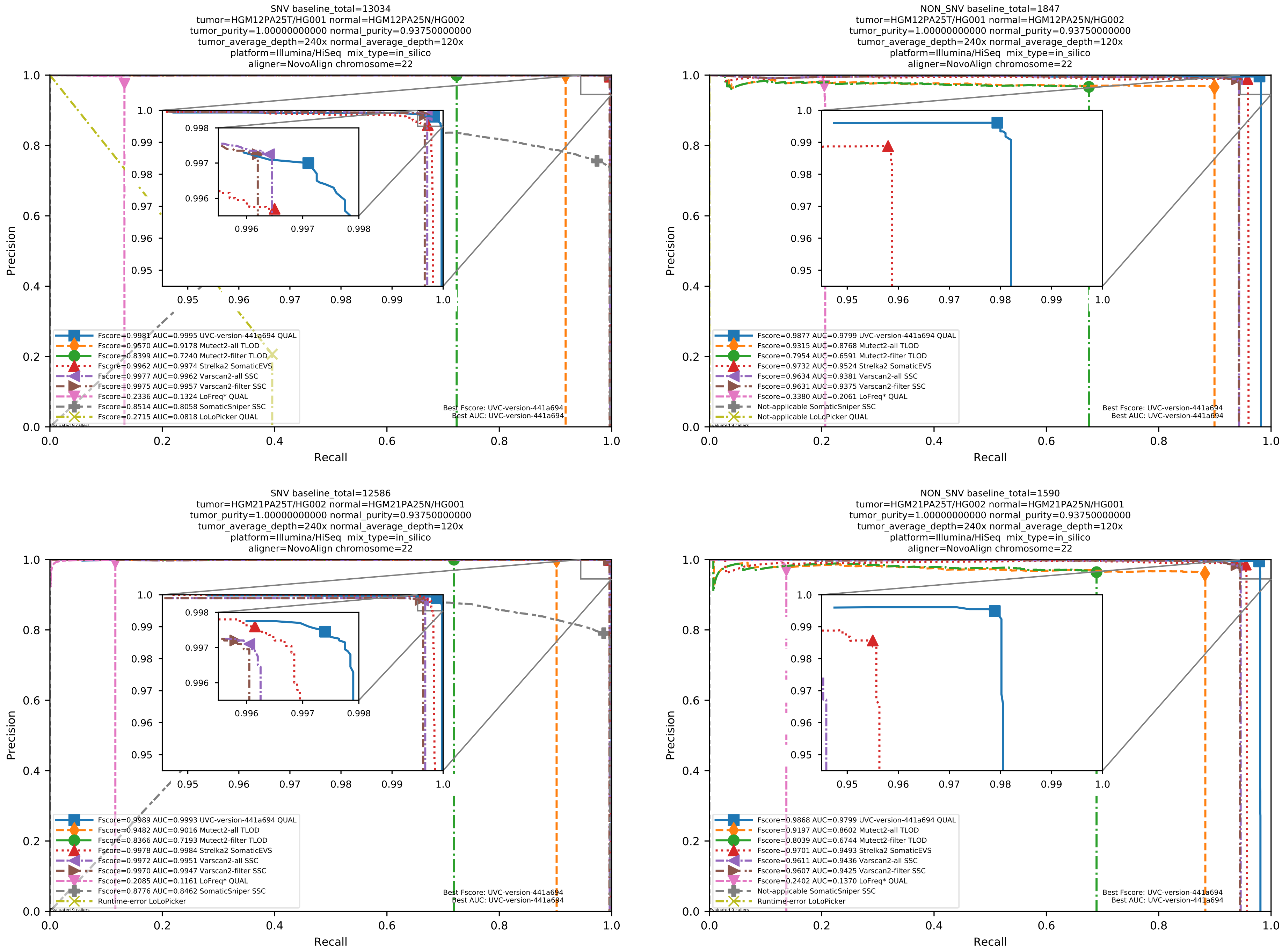

Supplementary Figure 65: Performance of somatic variant callers in tumor-with-normal mode on the *in silico* mixtures with Illumina HiSeq sequencing data pre-aligned with NovoAlign to the human reference genome hs37d5.

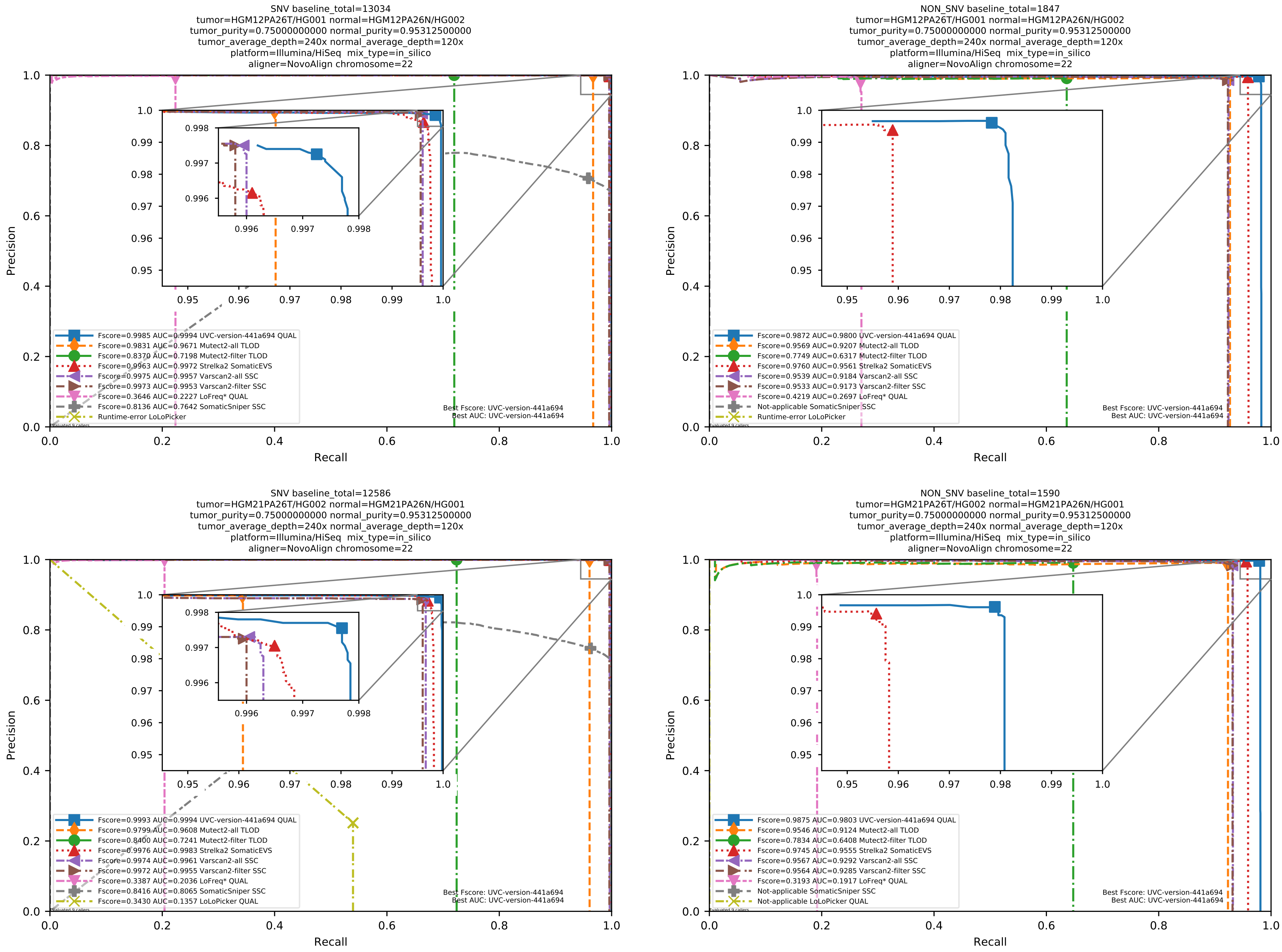

Supplementary Figure 66: Performance of somatic variant callers in tumor-with-normal mode on the *in silico* mixtures with Illumina HiSeq sequencing data pre-aligned with NovoAlign to the human reference genome hs37d5.

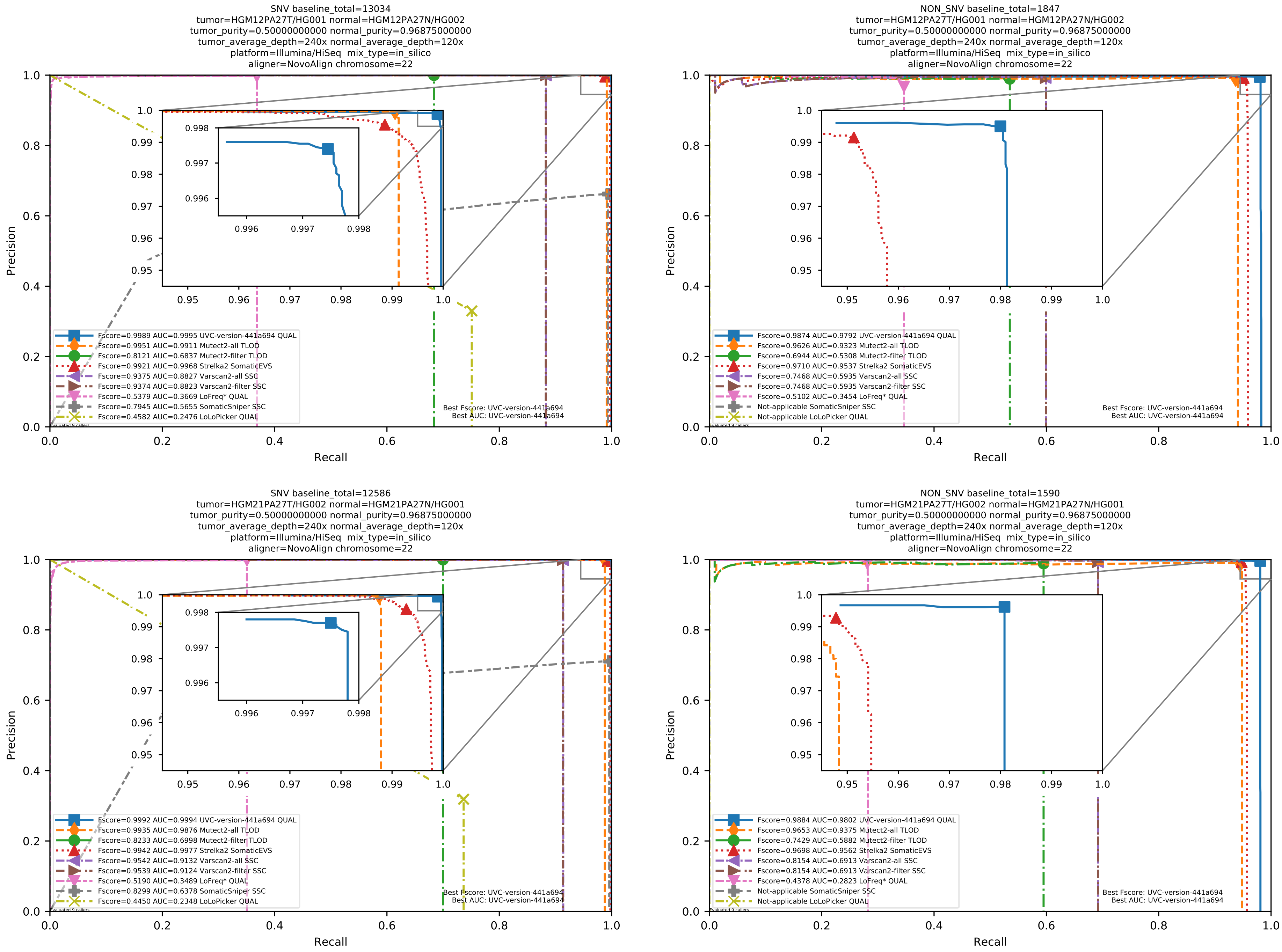

Supplementary Figure 67: Performance of somatic variant callers in tumor-with-normal mode on the *in silico* mixtures with Illumina HiSeq sequencing data pre-aligned with NovoAlign to the human reference genome hs37d5.

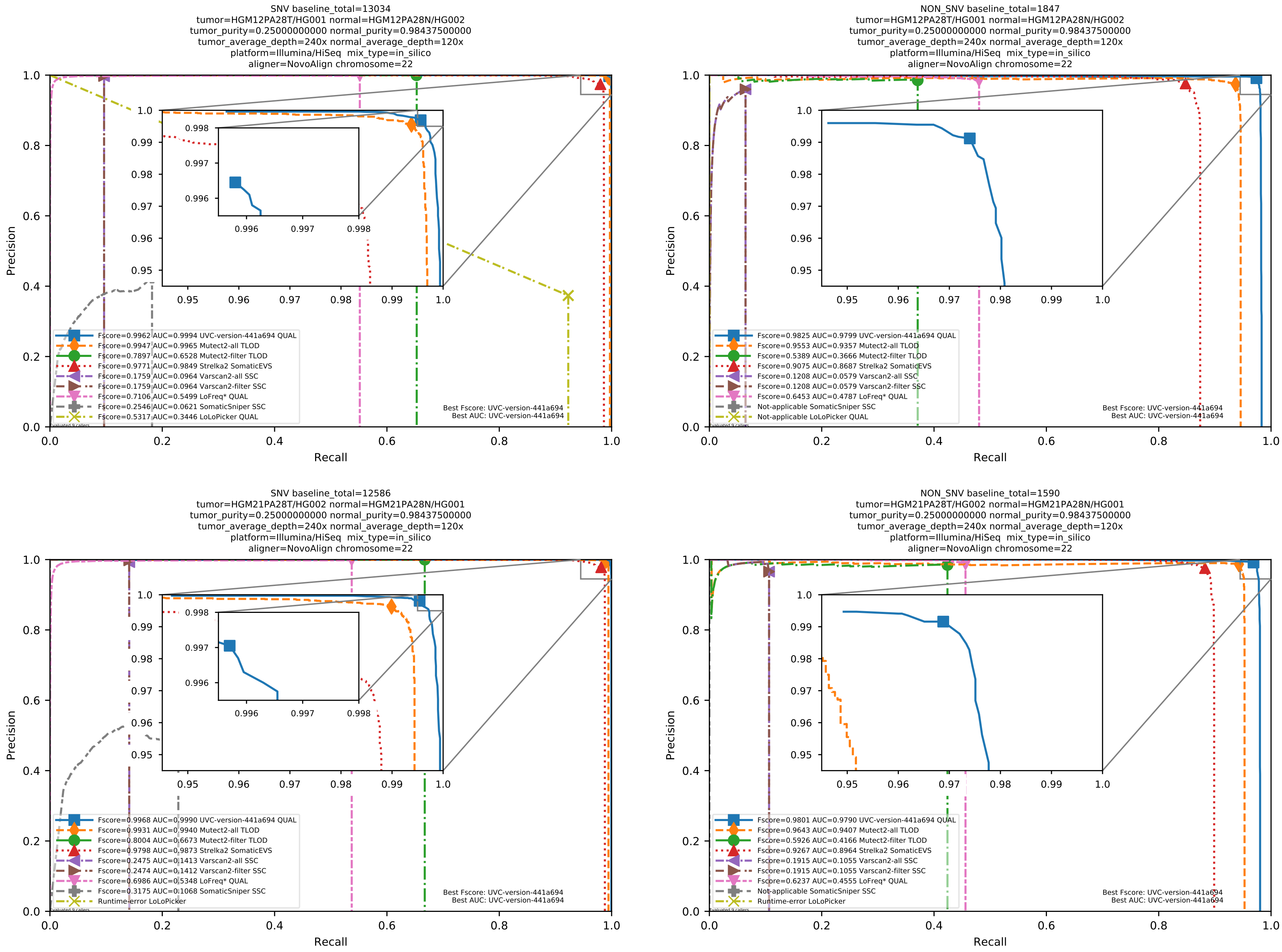

Supplementary Figure 68: Performance of somatic variant callers in tumor-with-normal mode on the *in silico* mixtures with Illumina HiSeq sequencing data pre-aligned with NovoAlign to the human reference genome hs37d5.

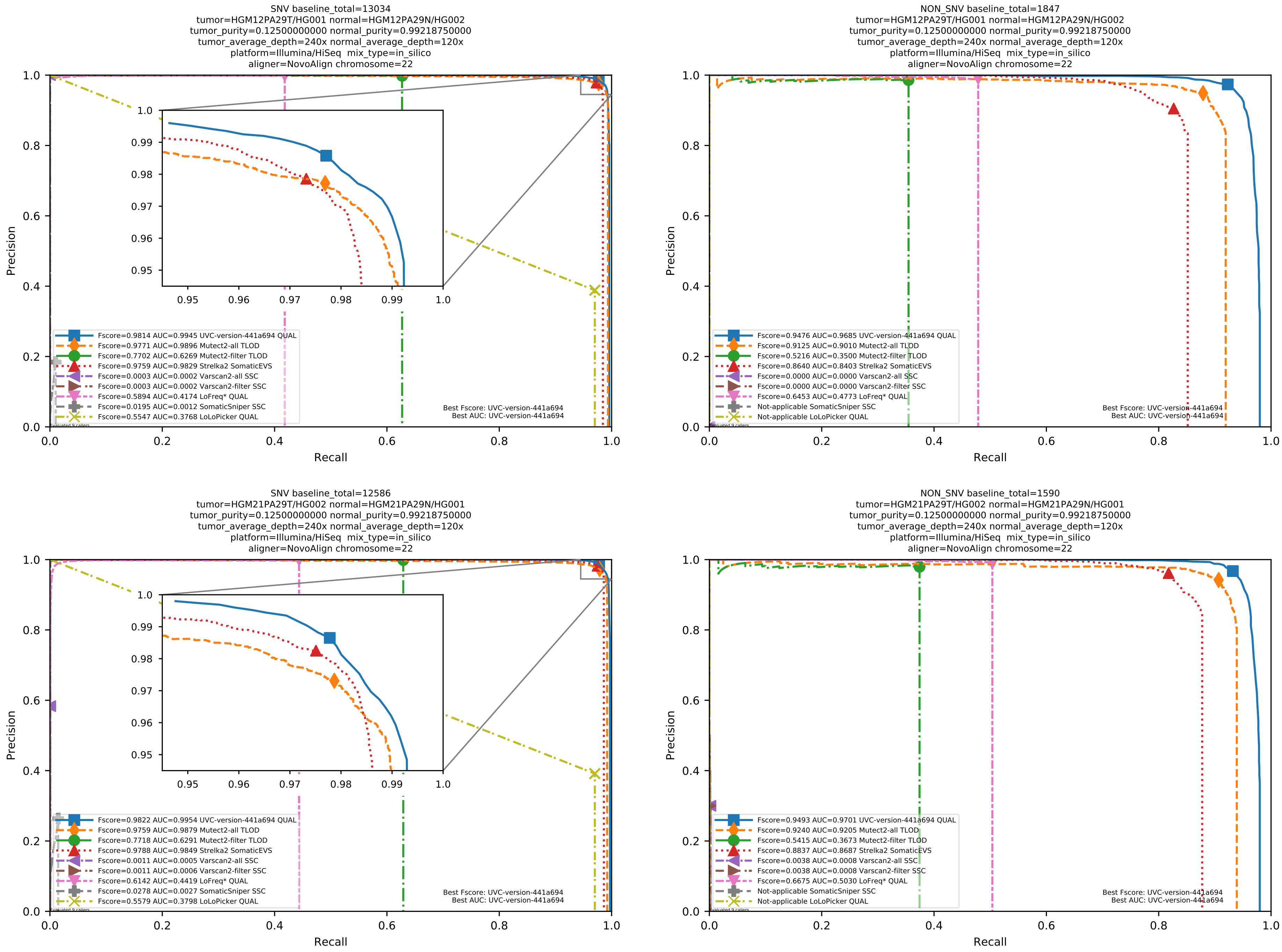

Supplementary Figure 69: Performance of somatic variant callers in tumor-with-normal mode on the *in silico* mixtures with Illumina HiSeq sequencing data pre-aligned with NovoAlign to the human reference genome hs37d5.

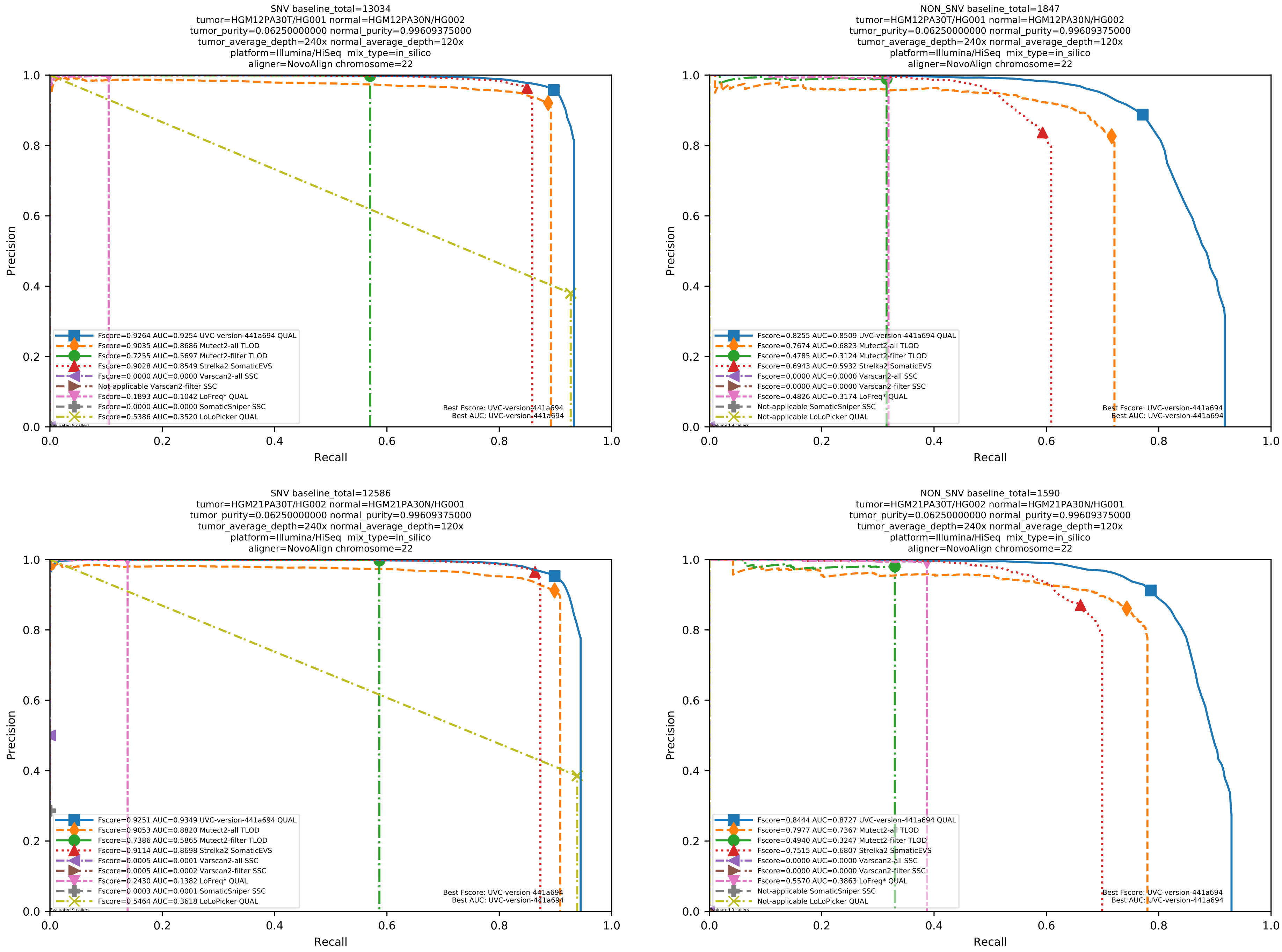

Supplementary Figure 70: Performance of somatic variant callers in tumor-with-normal mode on the *in silico* mixtures with Illumina HiSeq sequencing data pre-aligned with NovoAlign to the human reference genome hs37d5.

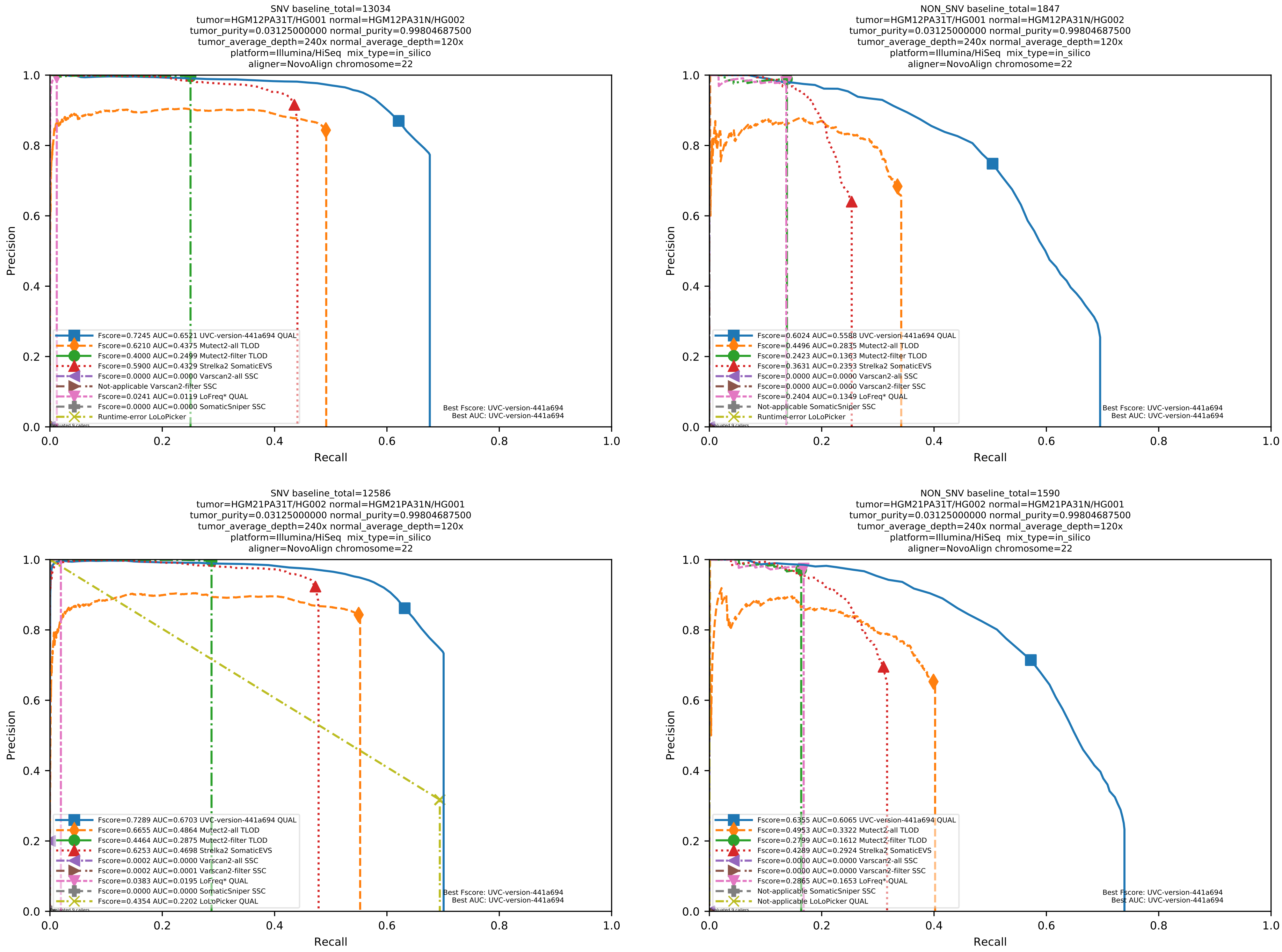

Supplementary Figure 71: Performance of somatic variant callers in tumor-with-normal mode on the *in silico* mixtures with Illumina HiSeq sequencing data pre-aligned with NovoAlign to the human reference genome hs37d5.

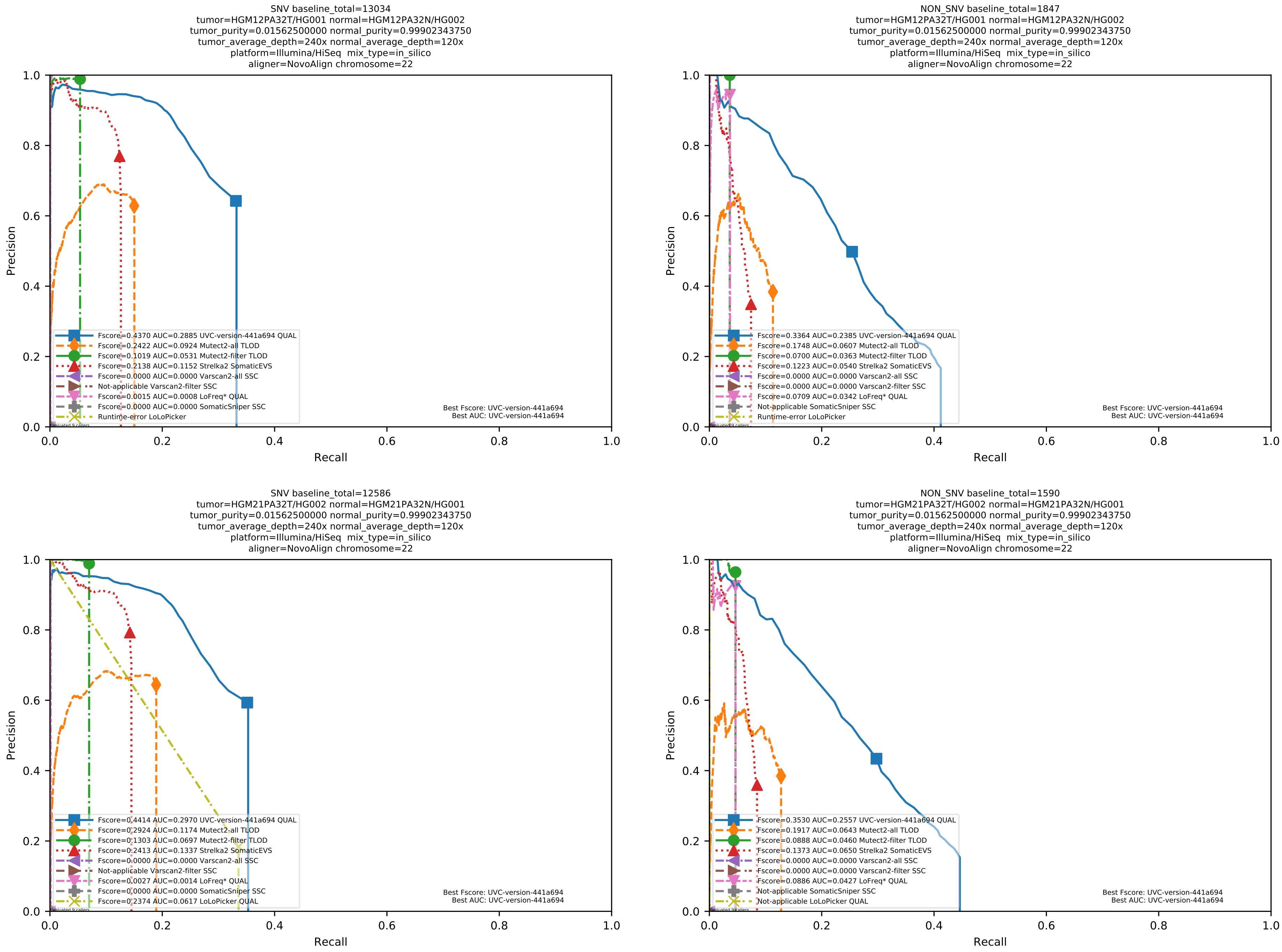

Supplementary Figure 72: Performance of somatic variant callers in tumor-with-normal mode on the *in silico* mixtures with Illumina HiSeq sequencing data pre-aligned with NovoAlign to the human reference genome hs37d5.

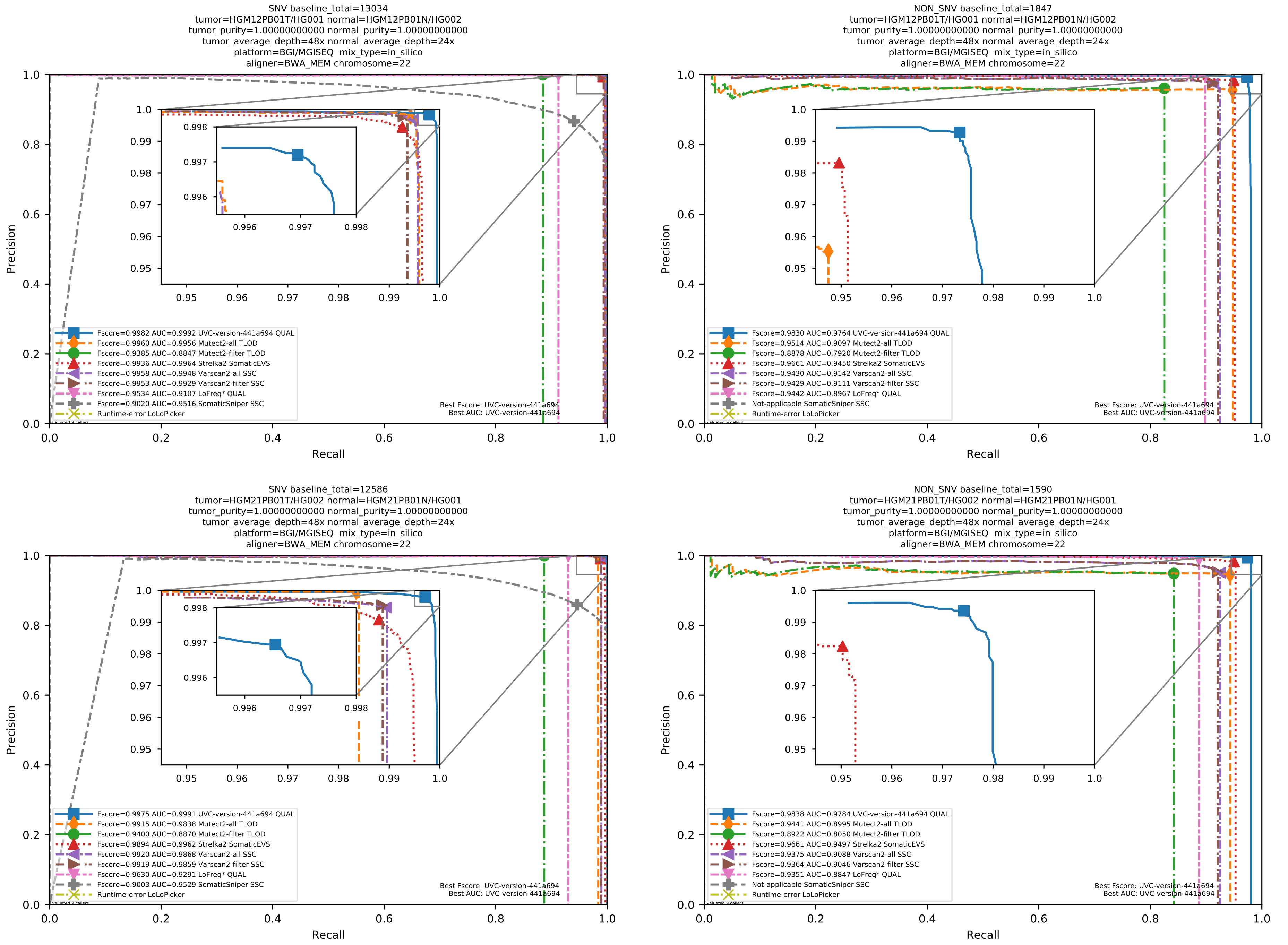

Supplementary Figure 73: Performance of somatic variant callers in tumor-with-normal mode on the *in silico* mixtures with BGI MGISEQ sequencing data aligned with BWA-MEM to the human reference genome hs37d5.

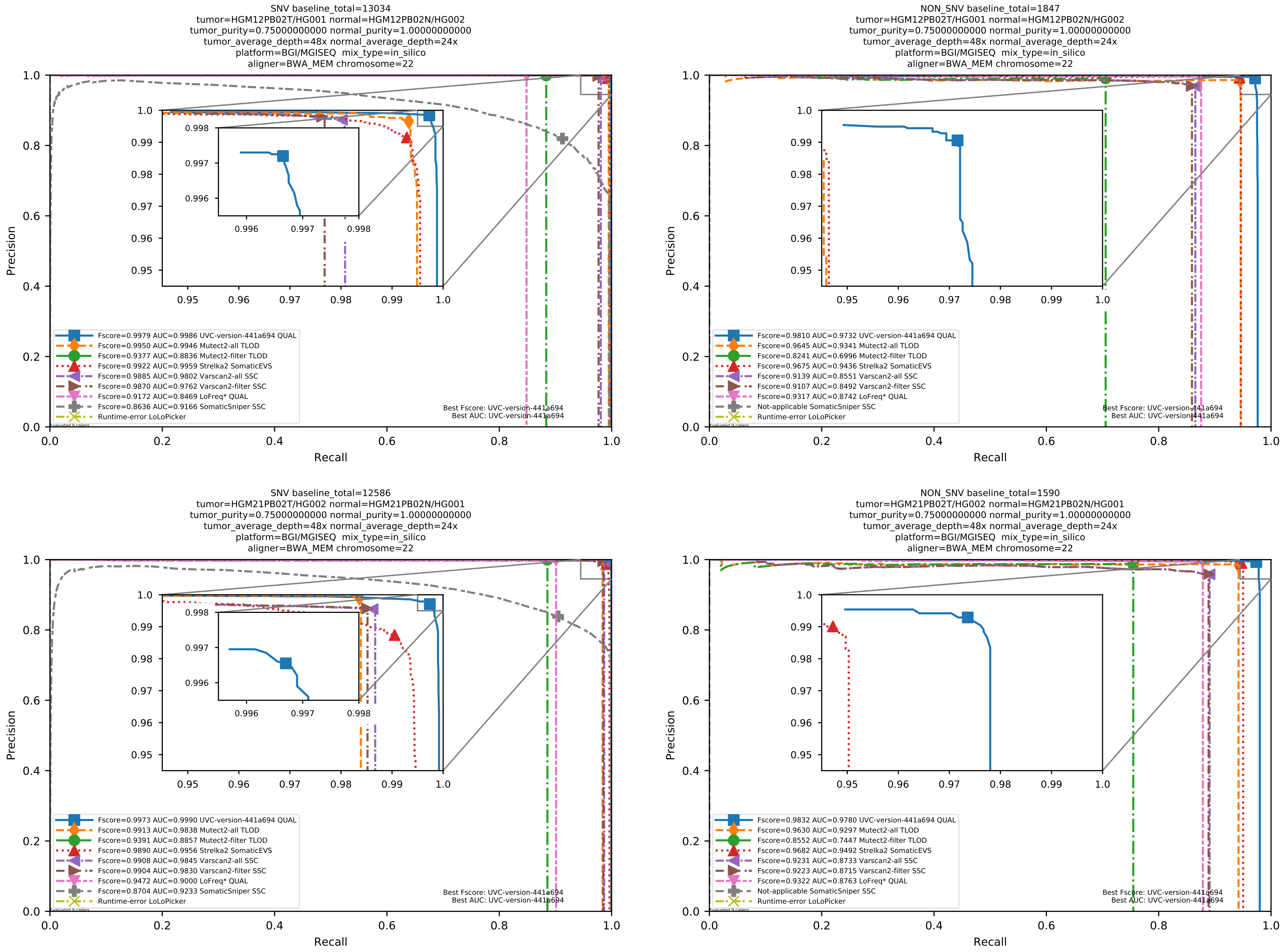

Supplementary Figure 74: Performance of somatic variant callers in tumor-with-normal mode on the *in silico* mixtures with BGI MGISEQ sequencing data aligned with BWA-MEM to the human reference genome hs37d5.

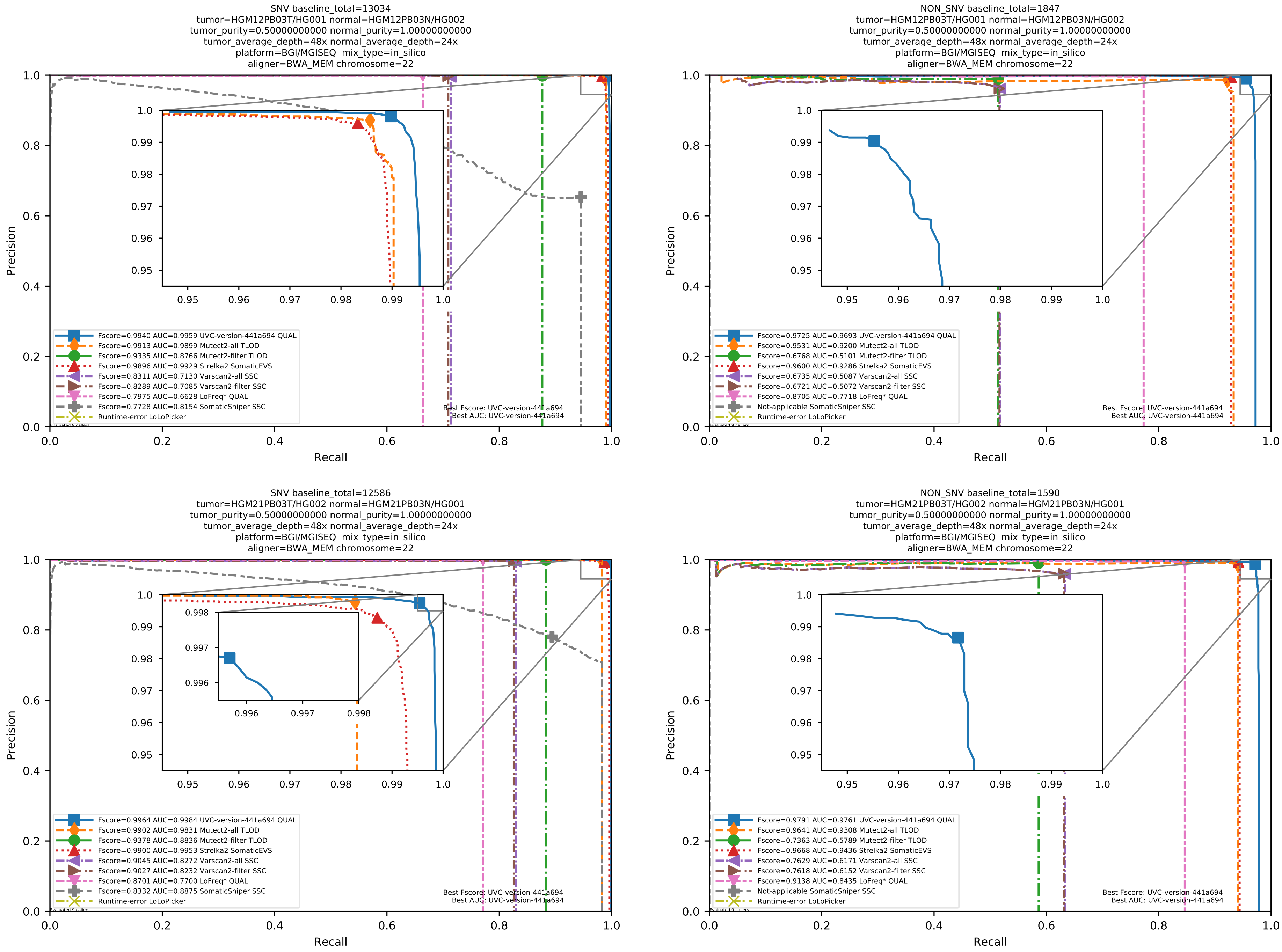

Supplementary Figure 75: Performance of somatic variant callers in tumor-with-normal mode on the *in silico* mixtures with BGI MGISEQ sequencing data aligned with BWA-MEM to the human reference genome hs37d5.

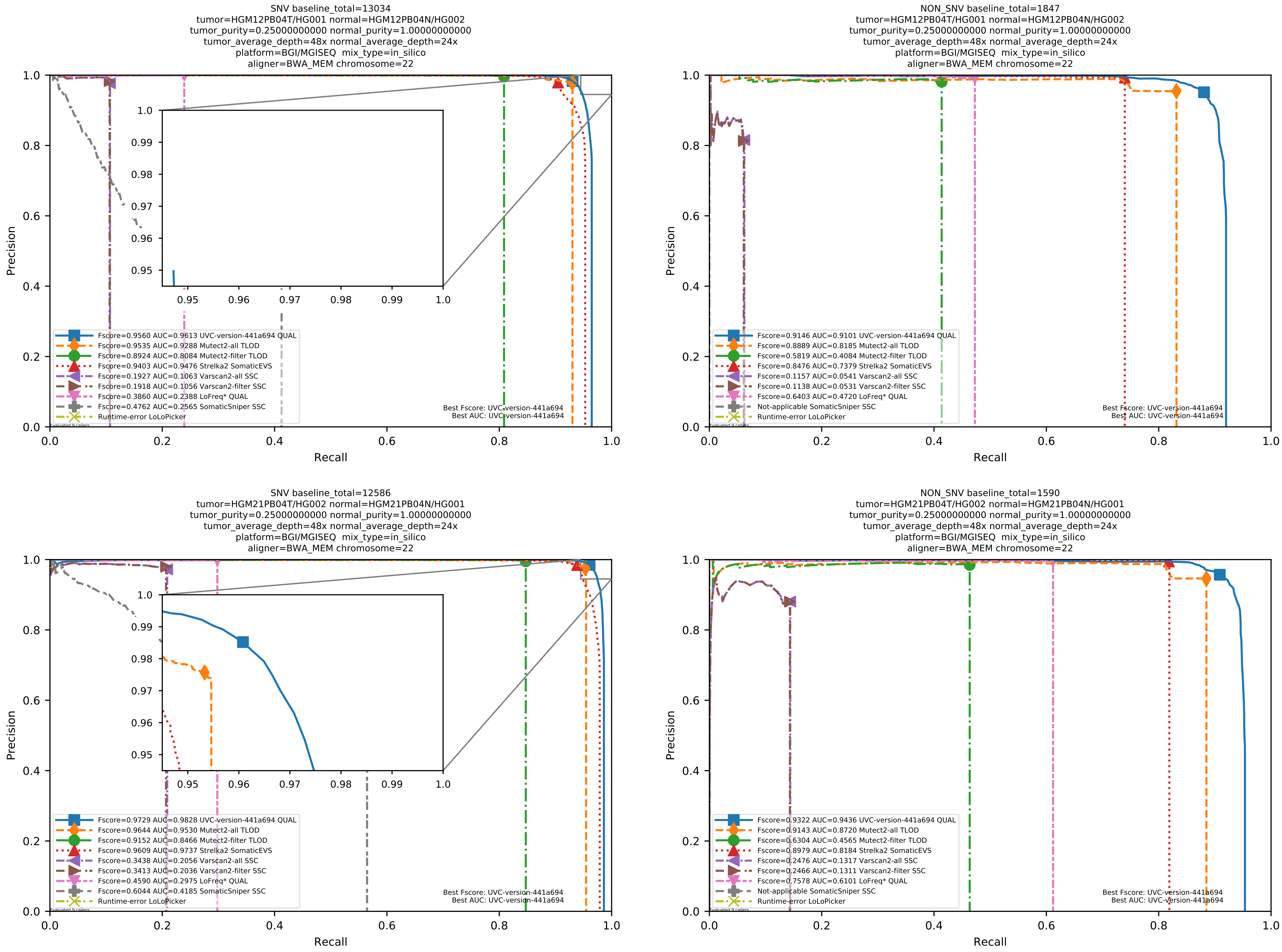

Supplementary Figure 76: Performance of somatic variant callers in tumor-with-normal mode on the *in silico* mixtures with BGI MGISEQ sequencing data aligned with BWA-MEM to the human reference genome hs37d5.

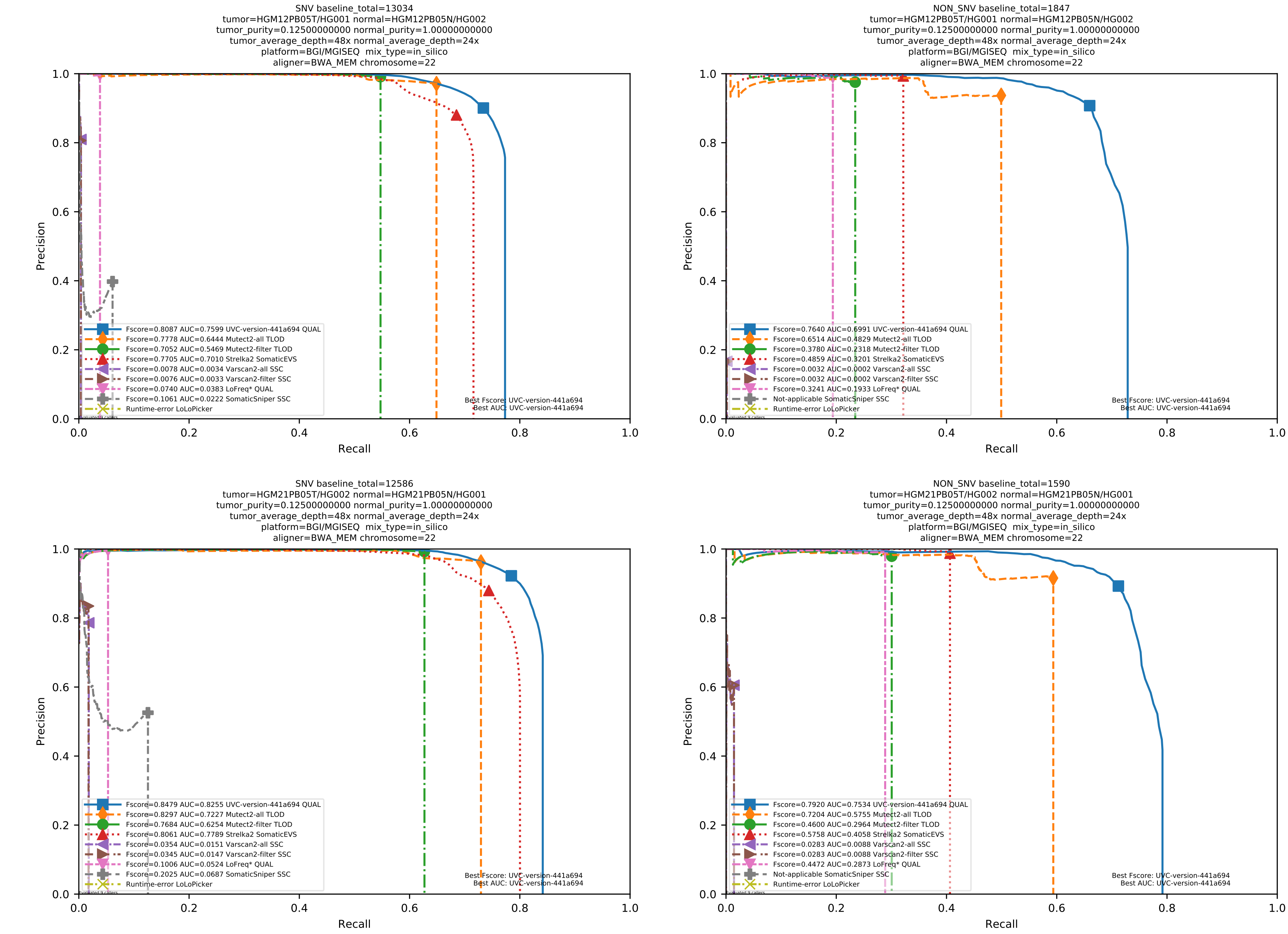

Supplementary Figure 77: Performance of somatic variant callers in tumor-with-normal mode on the *in silico* mixtures with BGI MGISEQ sequencing data aligned with BWA-MEM to the human reference genome hs37d5.

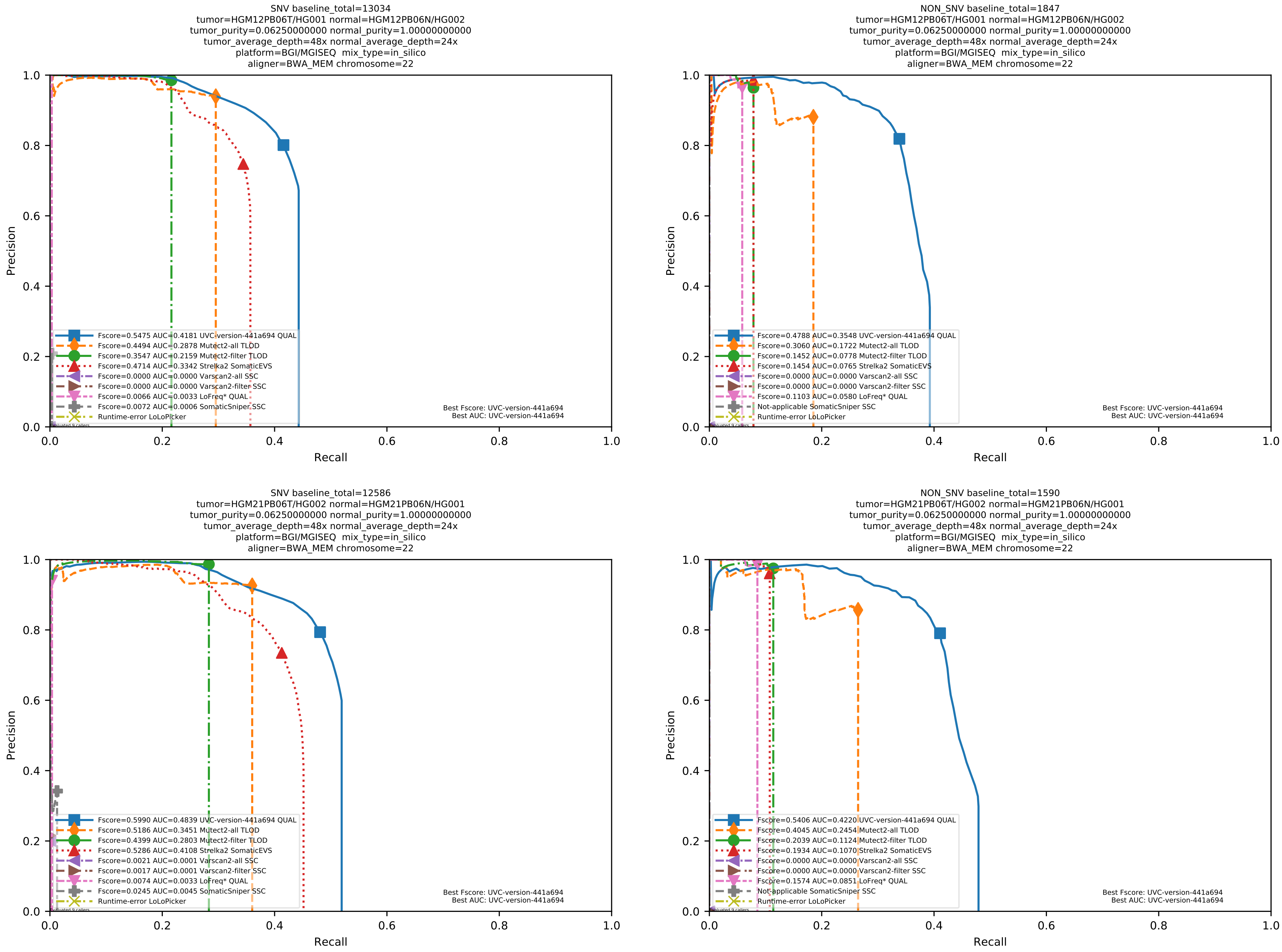

Supplementary Figure 78: Performance of somatic variant callers in tumor-with-normal mode on the *in silico* mixtures with BGI MGISEQ sequencing data aligned with BWA-MEM to the human reference genome hs37d5.

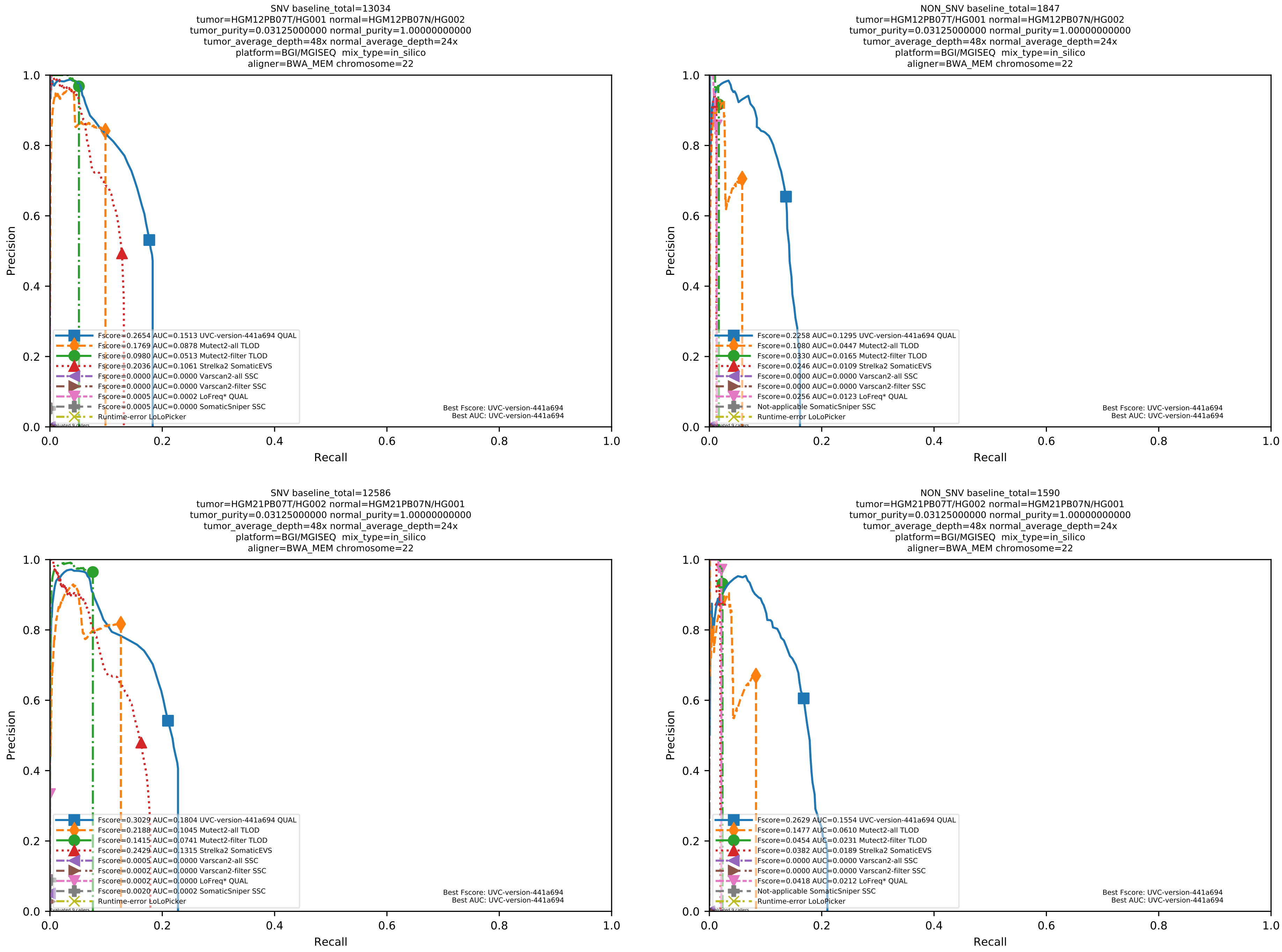

Supplementary Figure 79: Performance of somatic variant callers in tumor-with-normal mode on the *in silico* mixtures with BGI MGISEQ sequencing data aligned with BWA-MEM to the human reference genome hs37d5.

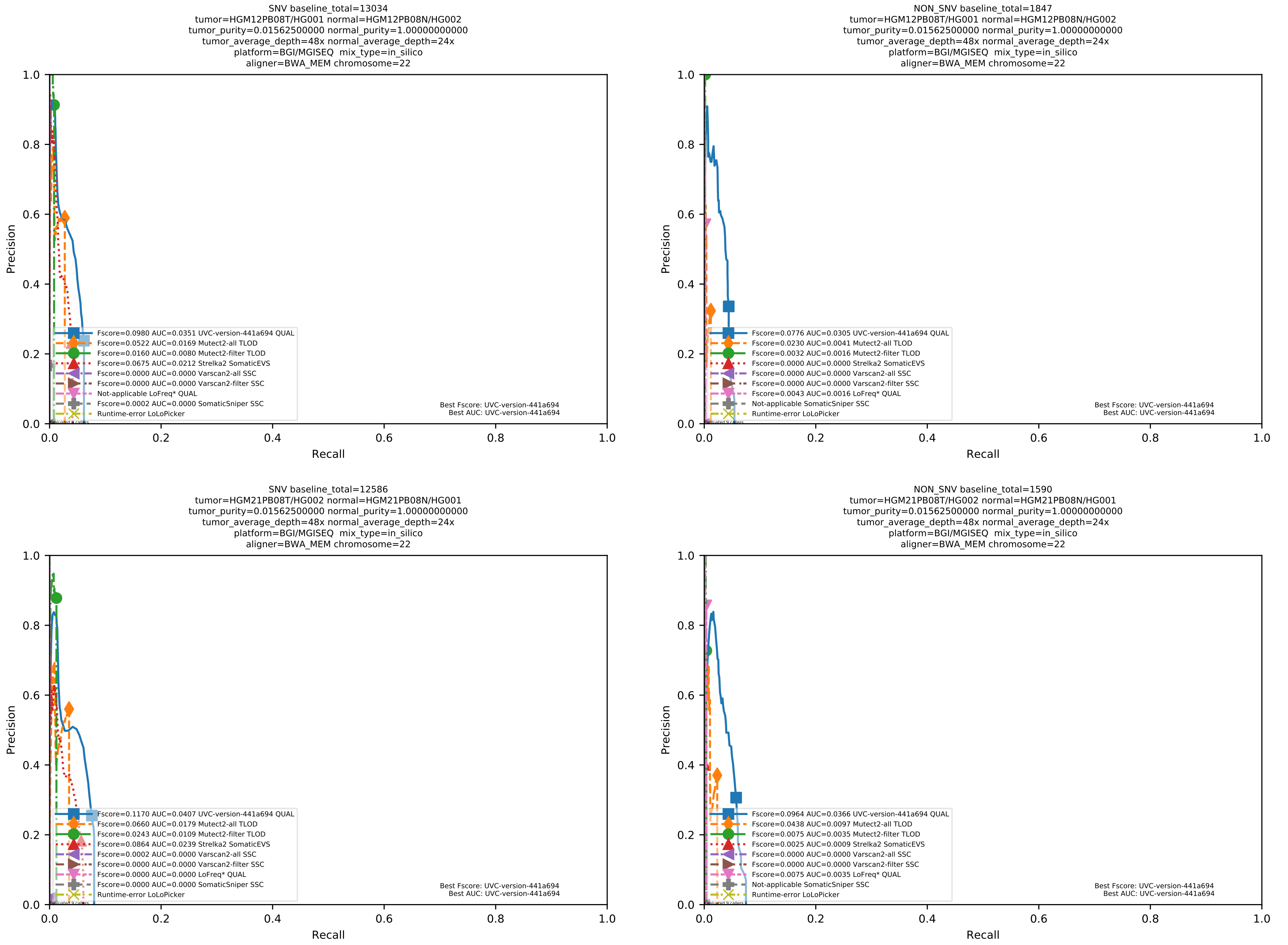

Supplementary Figure 80: Performance of somatic variant callers in tumor-with-normal mode on the *in silico* mixtures with BGI MGISEQ sequencing data aligned with BWA-MEM to the human reference genome hs37d5.

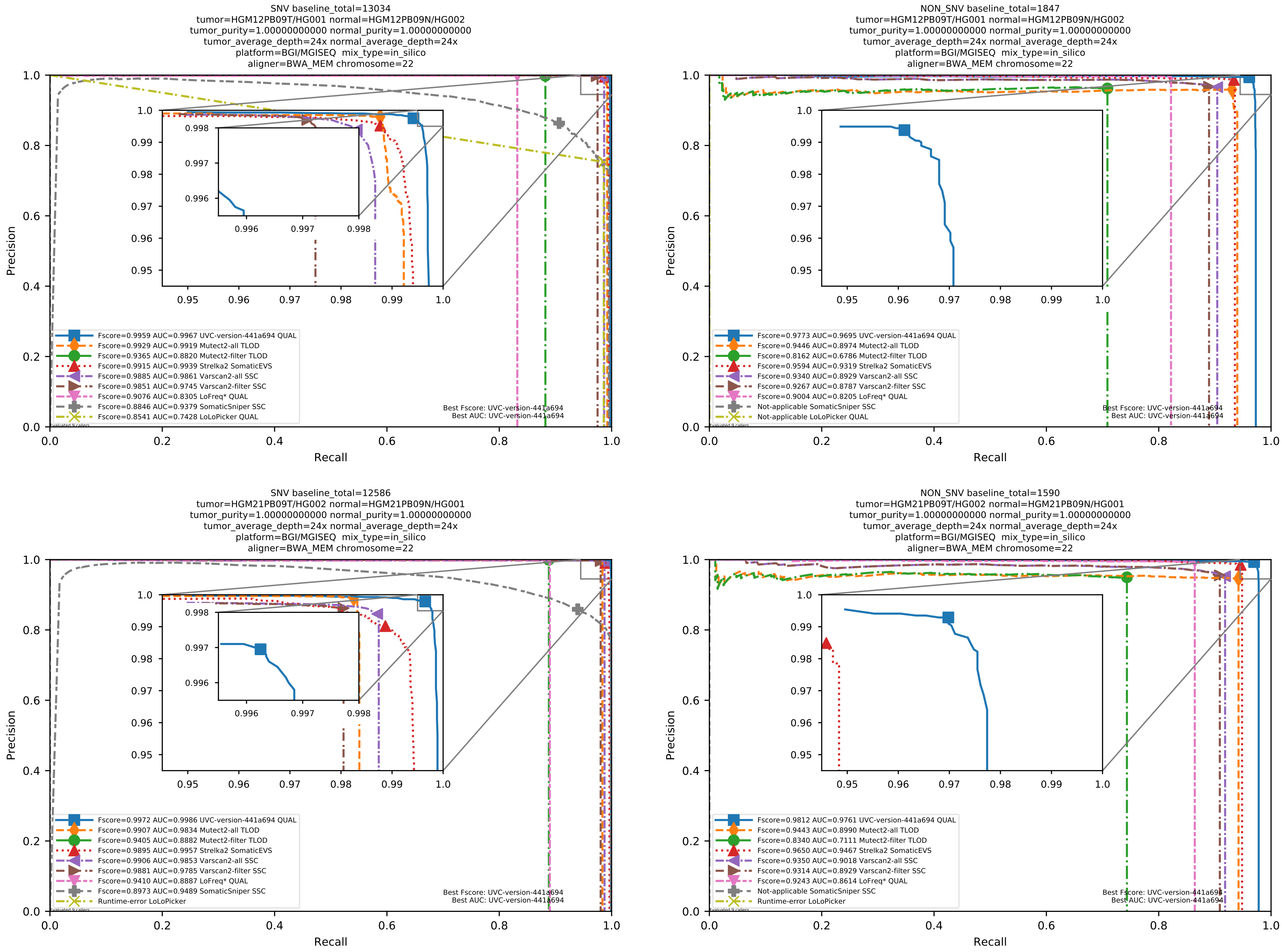

Supplementary Figure 81: Performance of somatic variant callers in tumor-with-normal mode on the *in silico* mixtures with BGI MGISEQ sequencing data aligned with BWA-MEM to the human reference genome hs37d5.

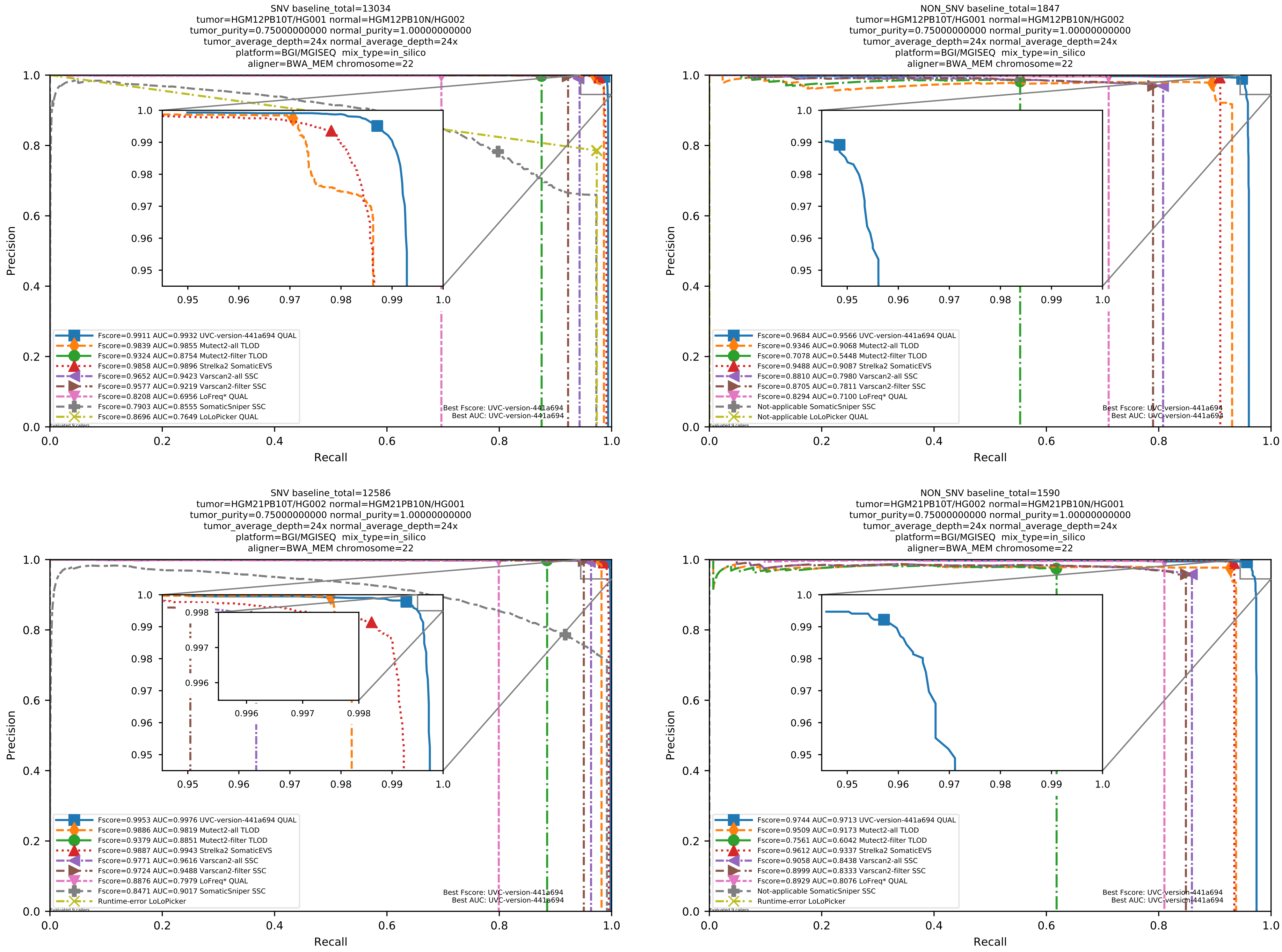

Supplementary Figure 82: Performance of somatic variant callers in tumor-with-normal mode on the *in silico* mixtures with BGI MGISEQ sequencing data aligned with BWA-MEM to the human reference genome hs37d5.

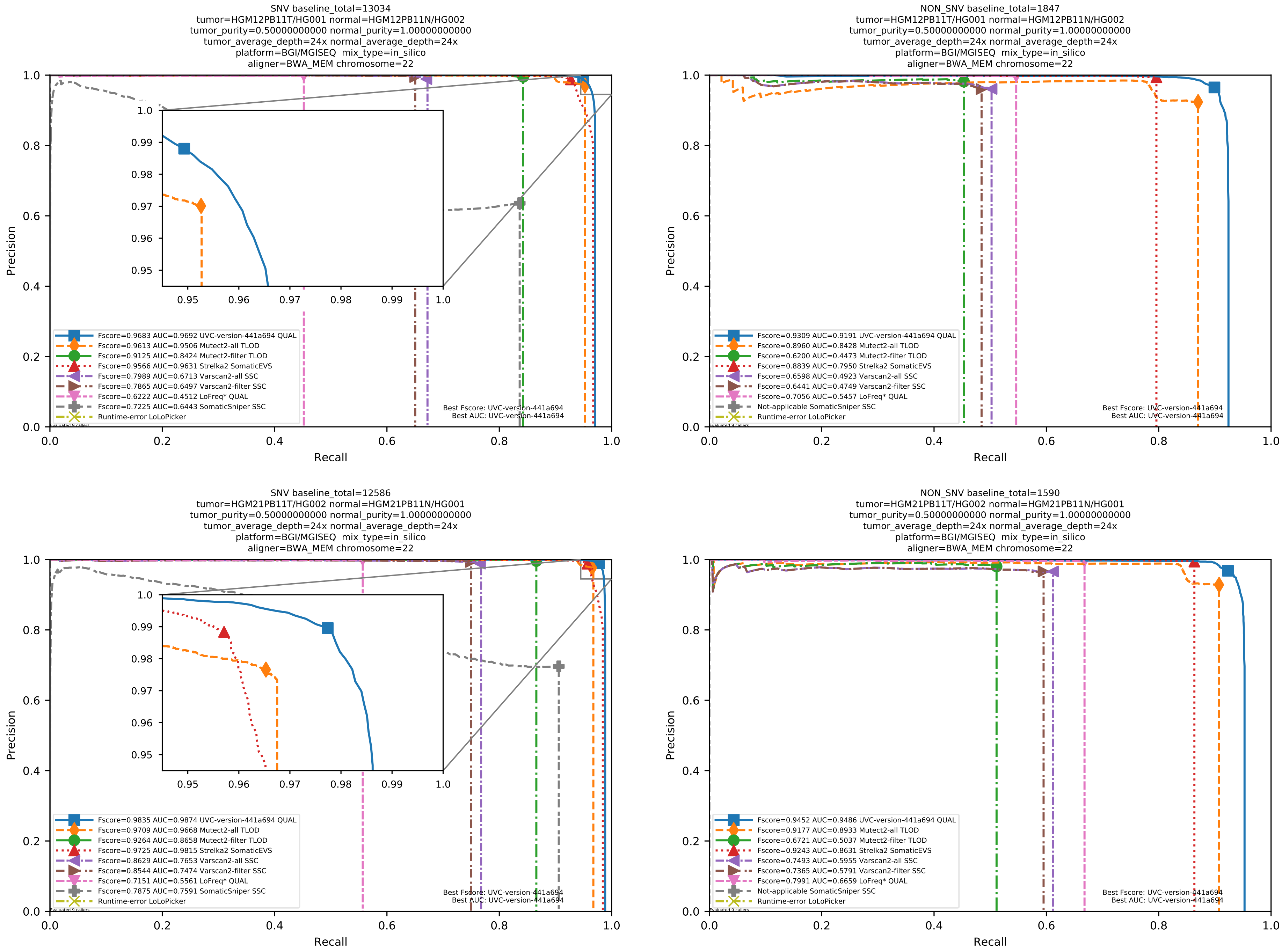

Supplementary Figure 83: Performance of somatic variant callers in tumor-with-normal mode on the *in silico* mixtures with BGI MGISEQ sequencing data aligned with BWA-MEM to the human reference genome hs37d5.

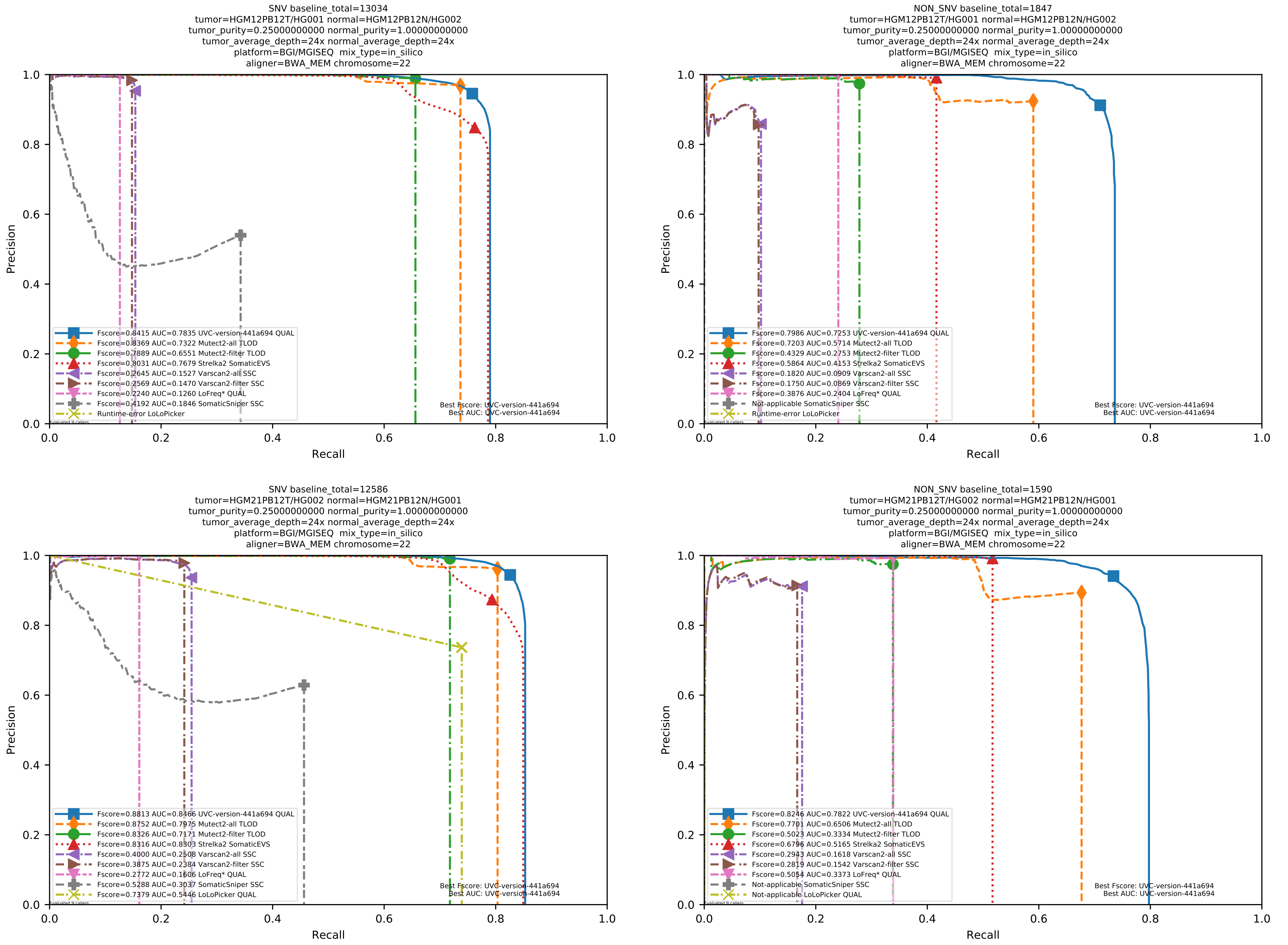

Supplementary Figure 84: Performance of somatic variant callers in tumor-with-normal mode on the *in silico* mixtures with BGI MGISEQ sequencing data aligned with BWA-MEM to the human reference genome hs37d5.

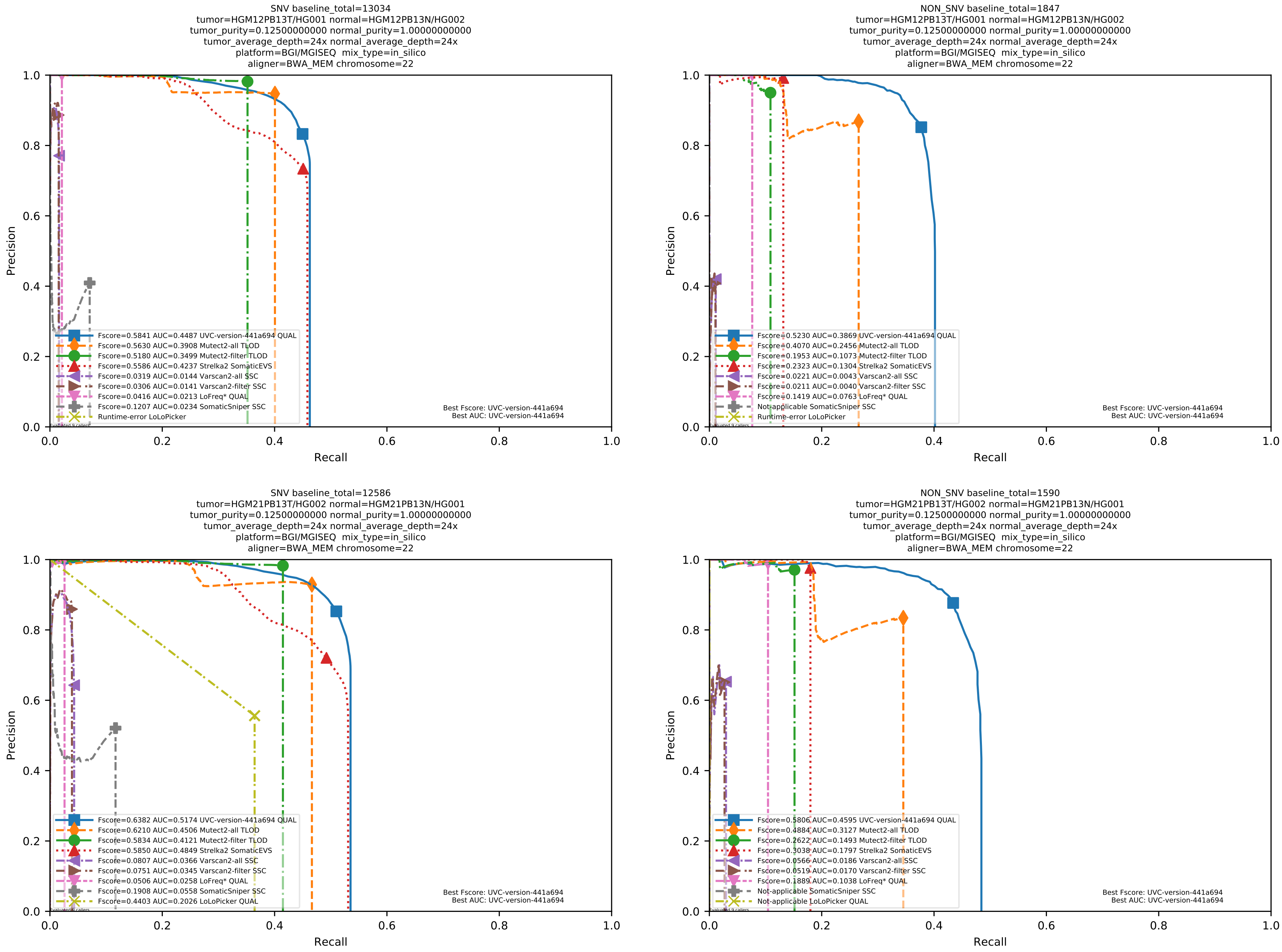

Supplementary Figure 85: Performance of somatic variant callers in tumor-with-normal mode on the *in silico* mixtures with BGI MGISEQ sequencing data aligned with BWA-MEM to the human reference genome hs37d5.

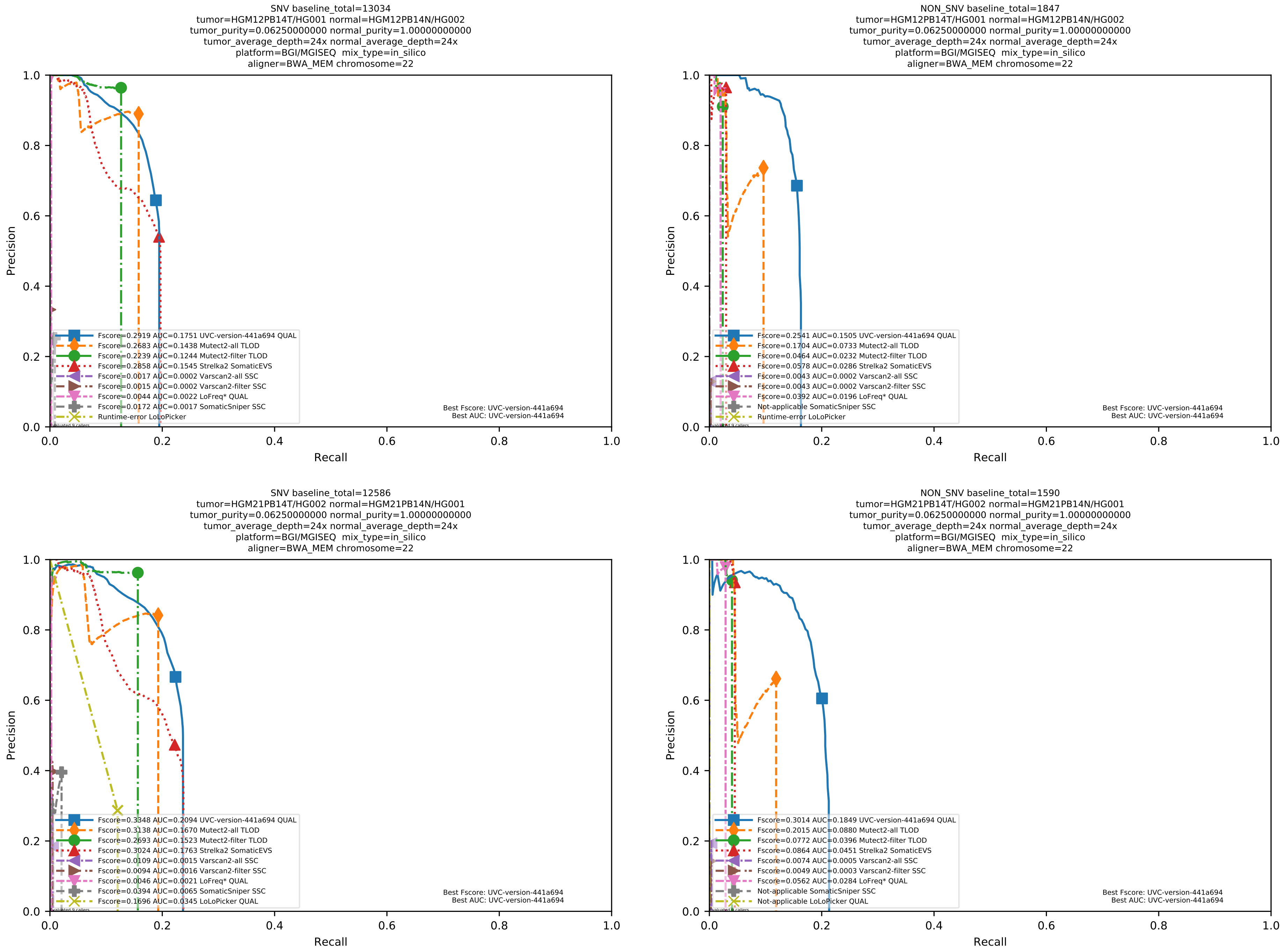

Supplementary Figure 86: Performance of somatic variant callers in tumor-with-normal mode on the *in silico* mixtures with BGI MGISEQ sequencing data aligned with BWA-MEM to the human reference genome hs37d5.

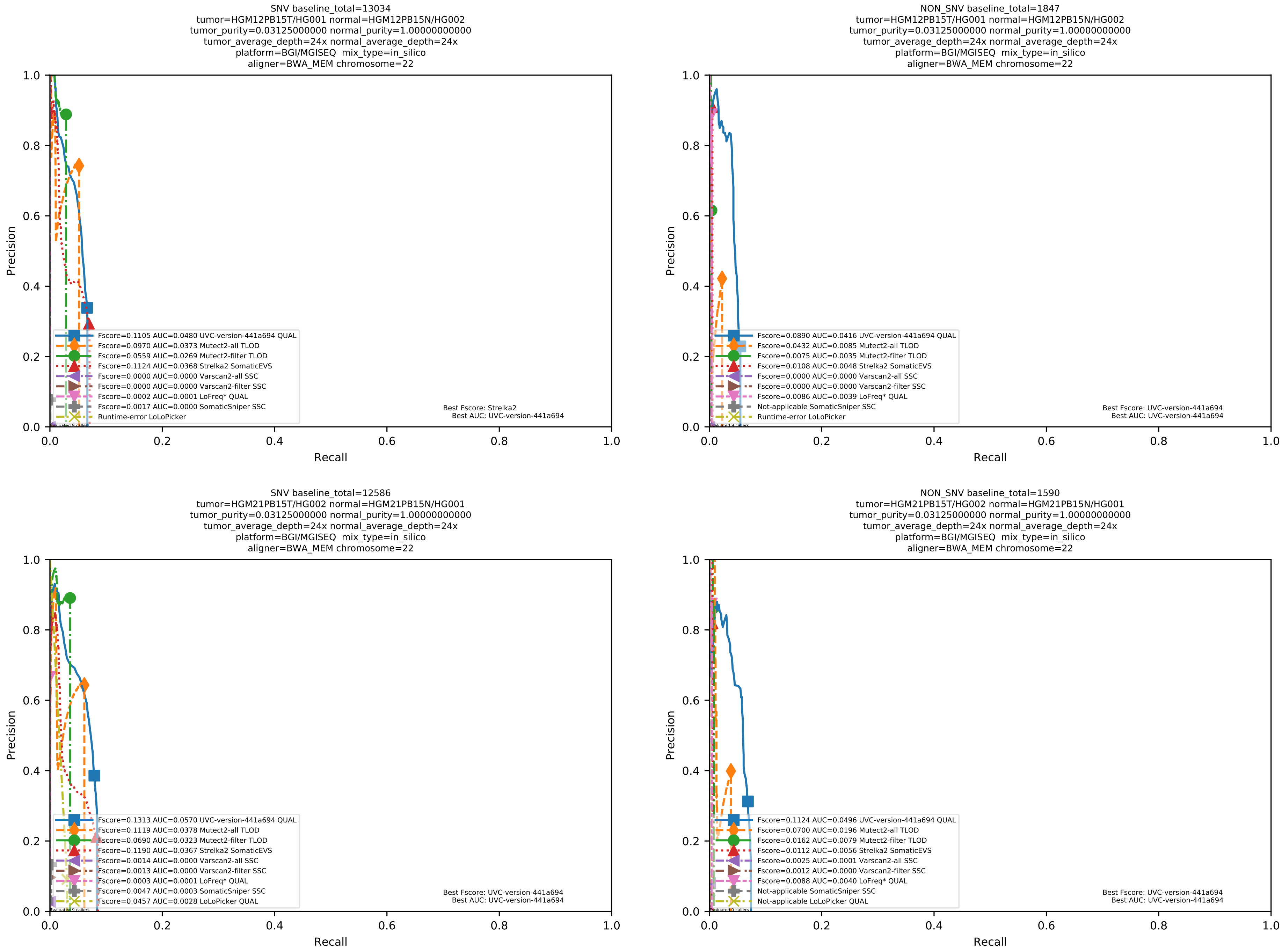

Supplementary Figure 87: Performance of somatic variant callers in tumor-with-normal mode on the *in silico* mixtures with BGI MGISEQ sequencing data aligned with BWA-MEM to the human reference genome hs37d5.

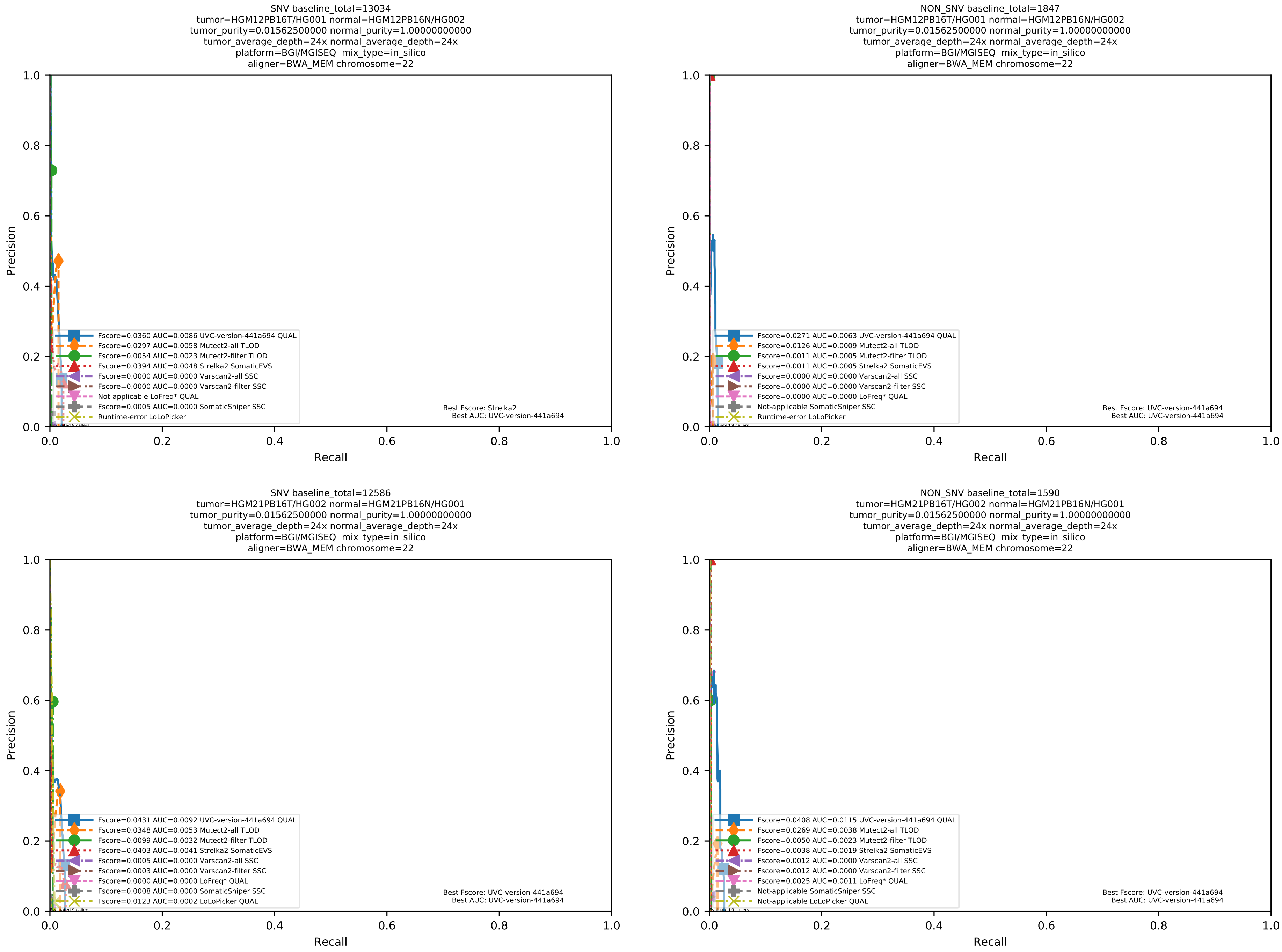

Supplementary Figure 88: Performance of somatic variant callers in tumor-with-normal mode on the *in silico* mixtures with BGI MGISEQ sequencing data aligned with BWA-MEM to the human reference genome hs37d5.

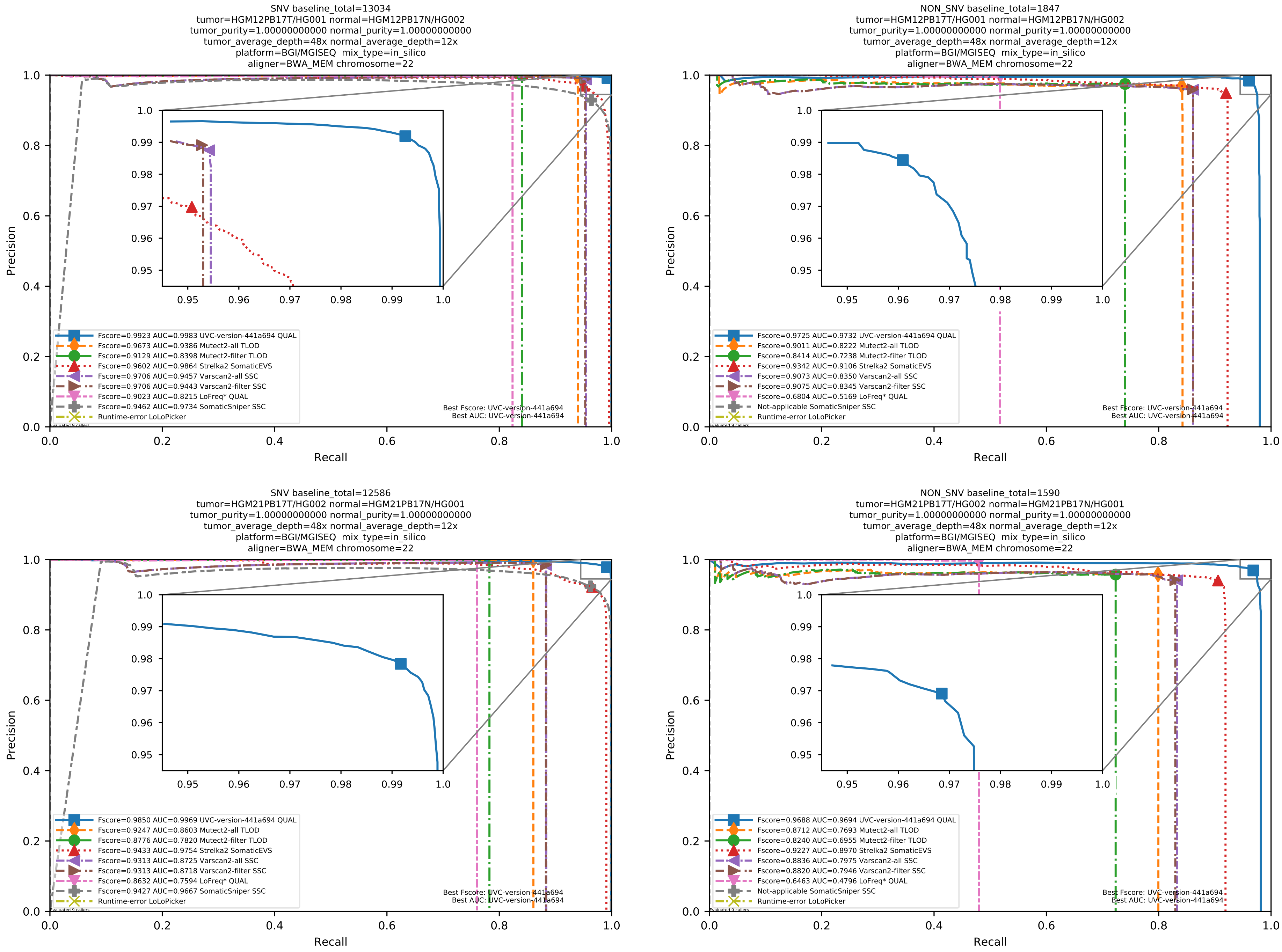

Supplementary Figure 89: Performance of somatic variant callers in tumor-with-normal mode on the *in silico* mixtures with BGI MGISEQ sequencing data aligned with BWA-MEM to the human reference genome hs37d5.

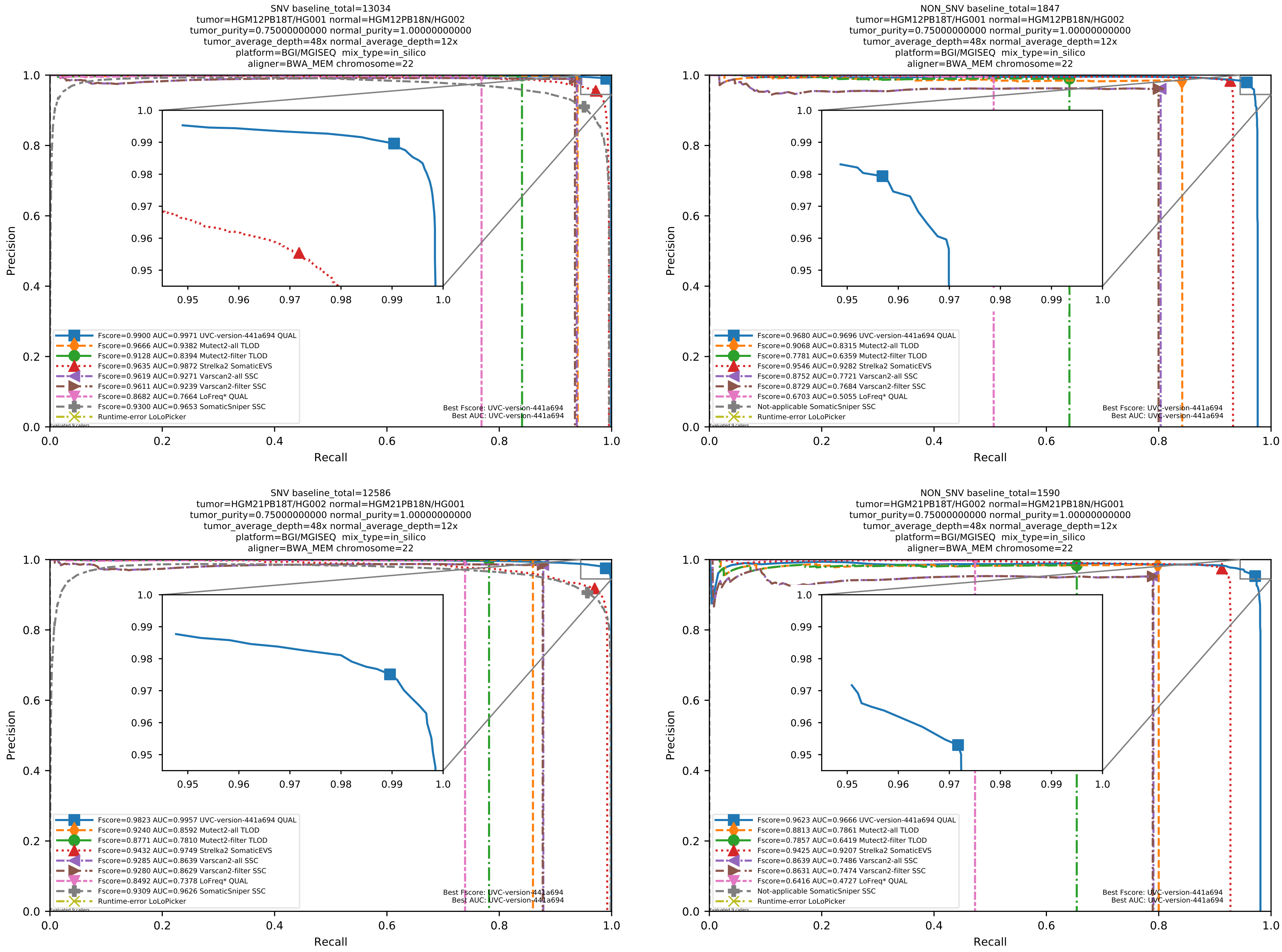

Supplementary Figure 90: Performance of somatic variant callers in tumor-with-normal mode on the *in silico* mixtures with BGI MGISEQ sequencing data aligned with BWA-MEM to the human reference genome hs37d5.

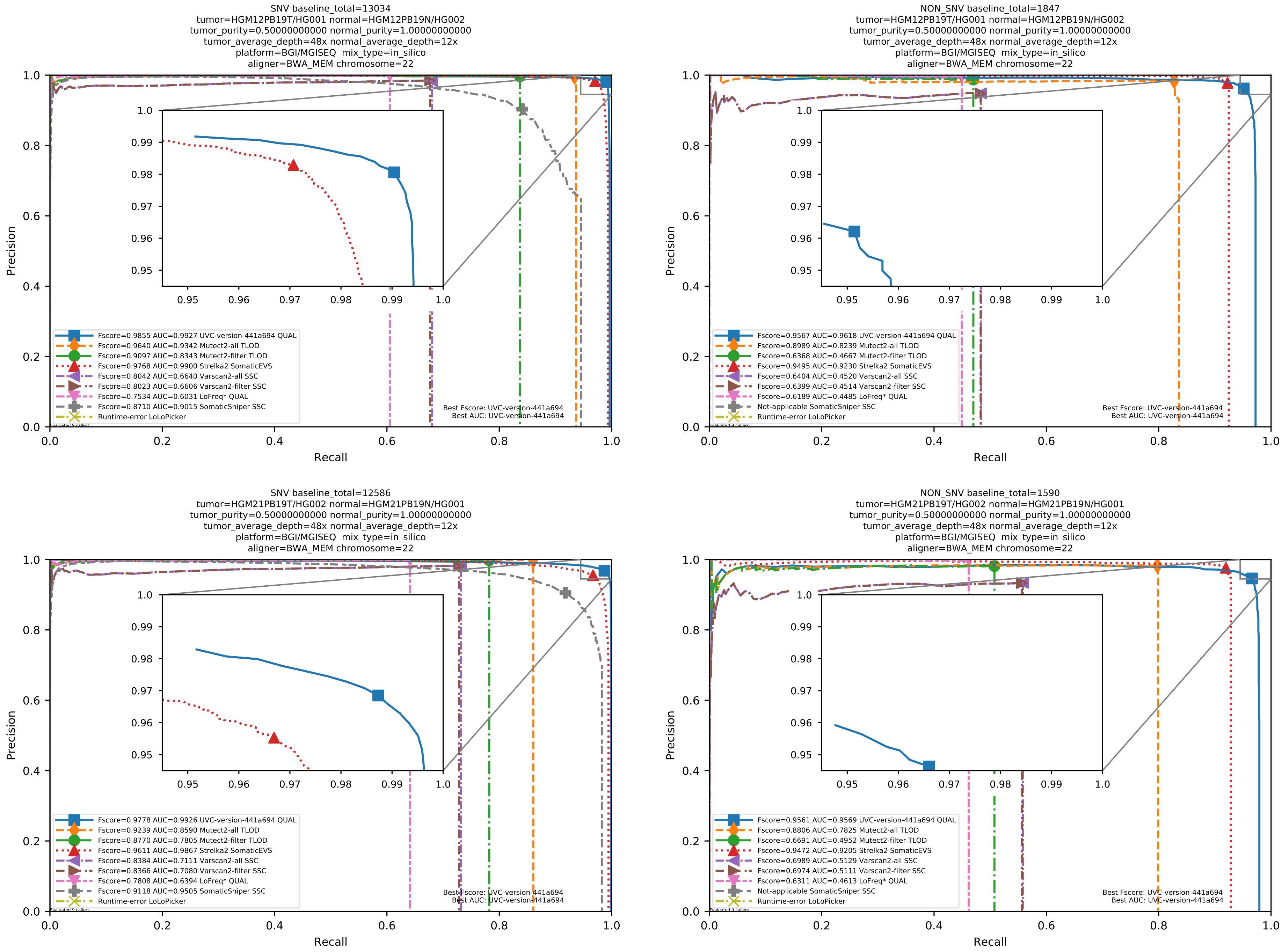

Supplementary Figure 91: Performance of somatic variant callers in tumor-with-normal mode on the *in silico* mixtures with BGI MGISEQ sequencing data aligned with BWA-MEM to the human reference genome hs37d5.

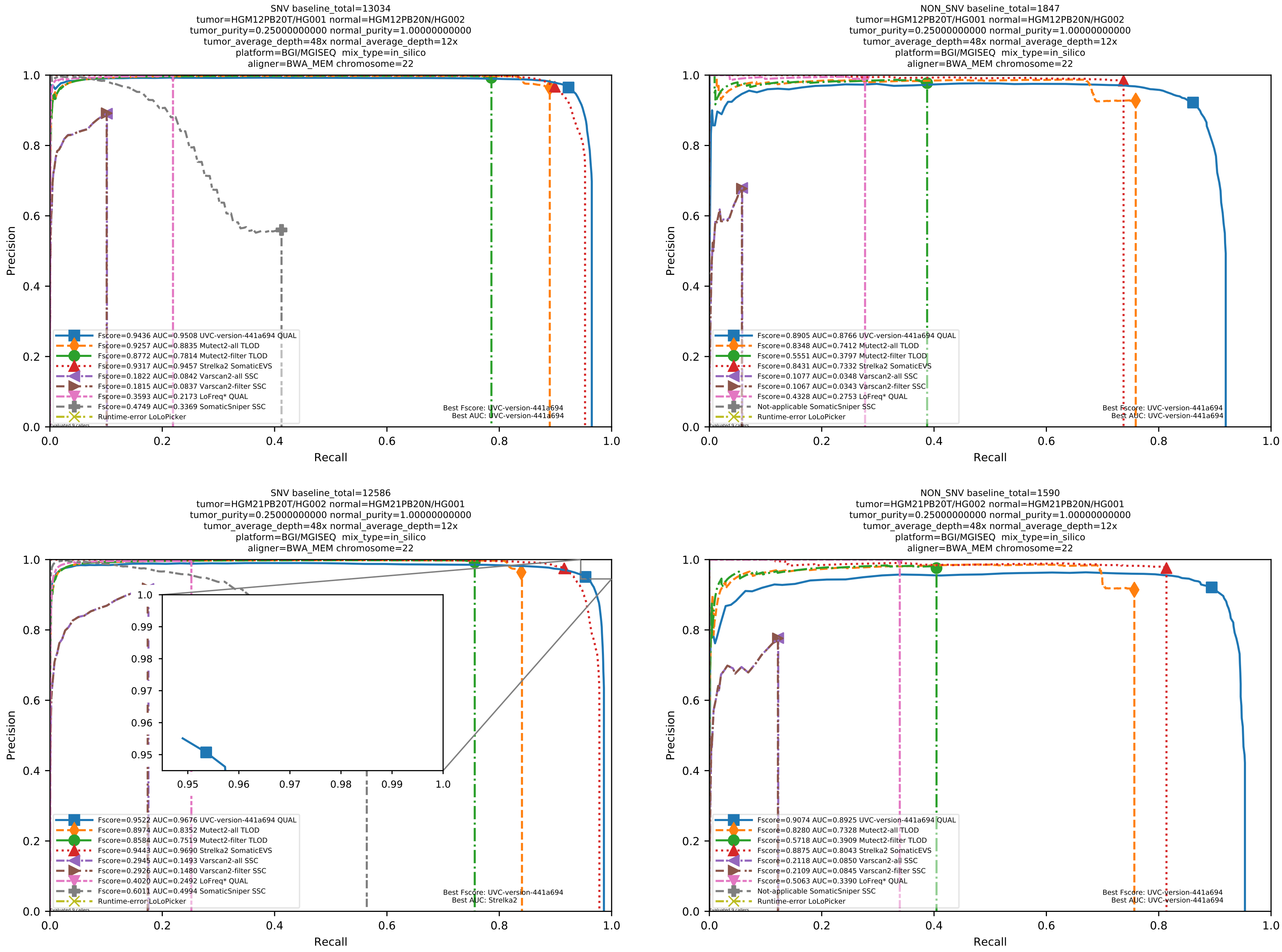

Supplementary Figure 92: Performance of somatic variant callers in tumor-with-normal mode on the *in silico* mixtures with BGI MGISEQ sequencing data aligned with BWA-MEM to the human reference genome hs37d5.

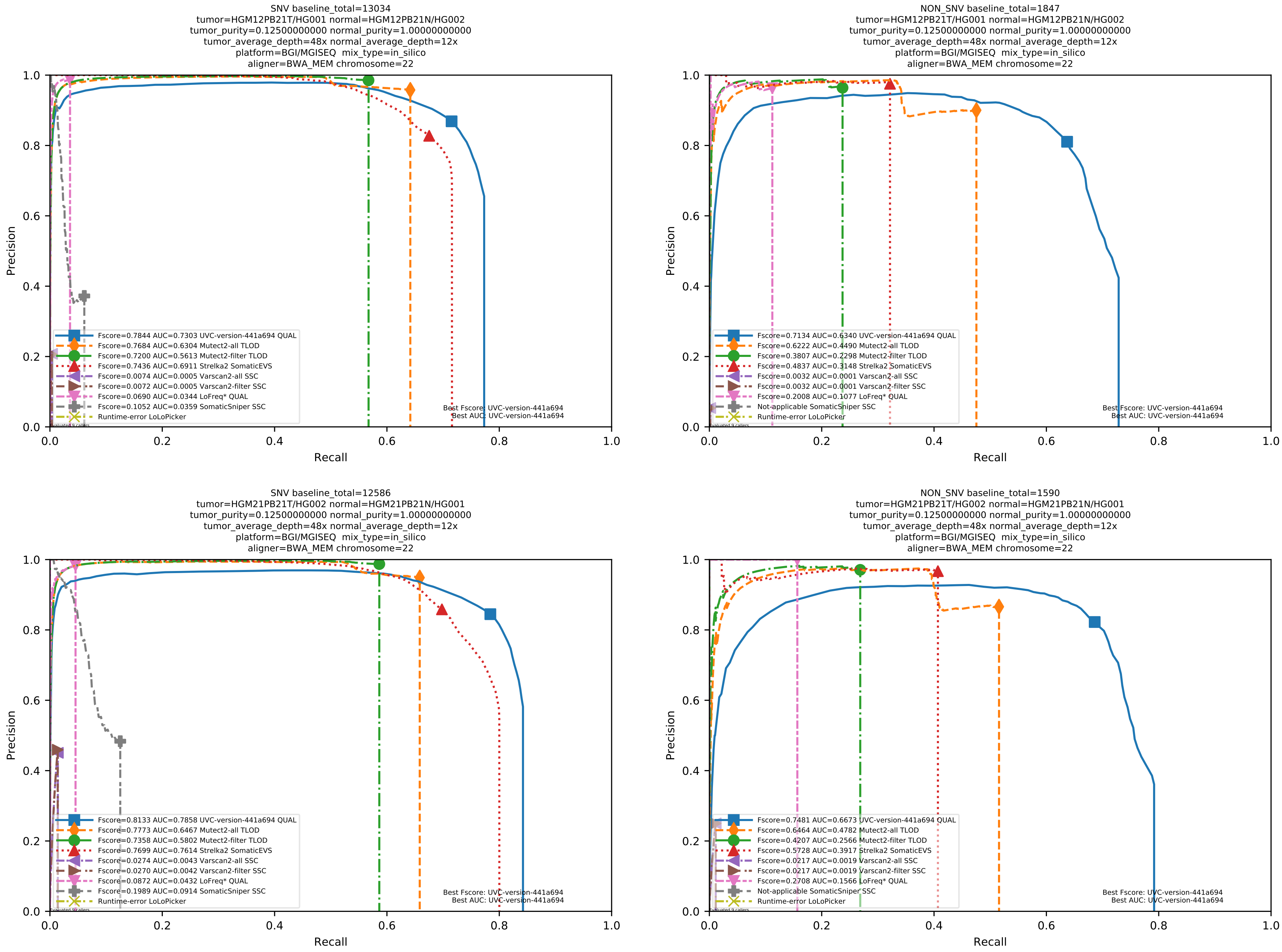

Supplementary Figure 93: Performance of somatic variant callers in tumor-with-normal mode on the *in silico* mixtures with BGI MGISEQ sequencing data aligned with BWA-MEM to the human reference genome hs37d5.

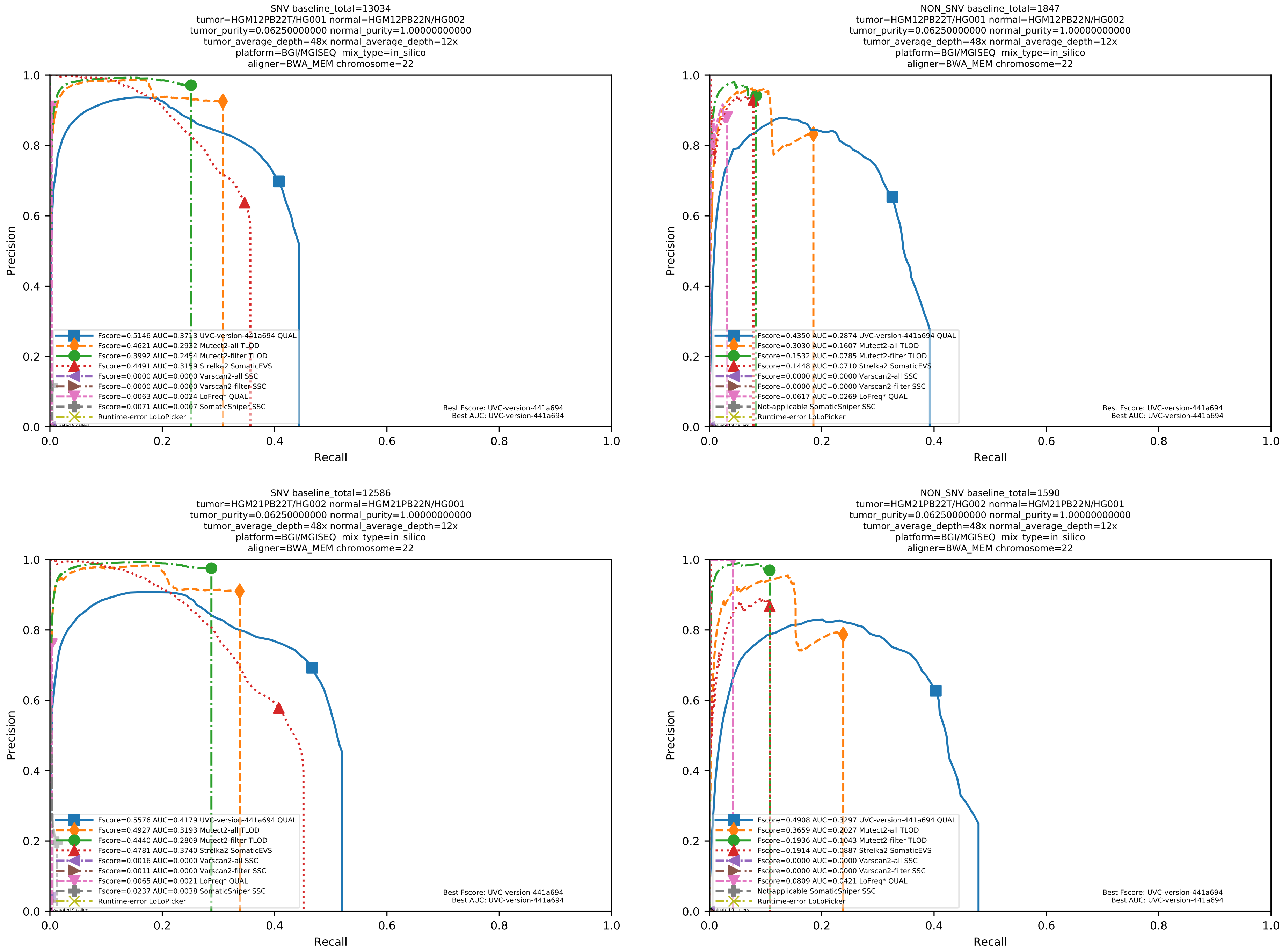

Supplementary Figure 94: Performance of somatic variant callers in tumor-with-normal mode on the *in silico* mixtures with BGI MGISEQ sequencing data aligned with BWA-MEM to the human reference genome hs37d5.

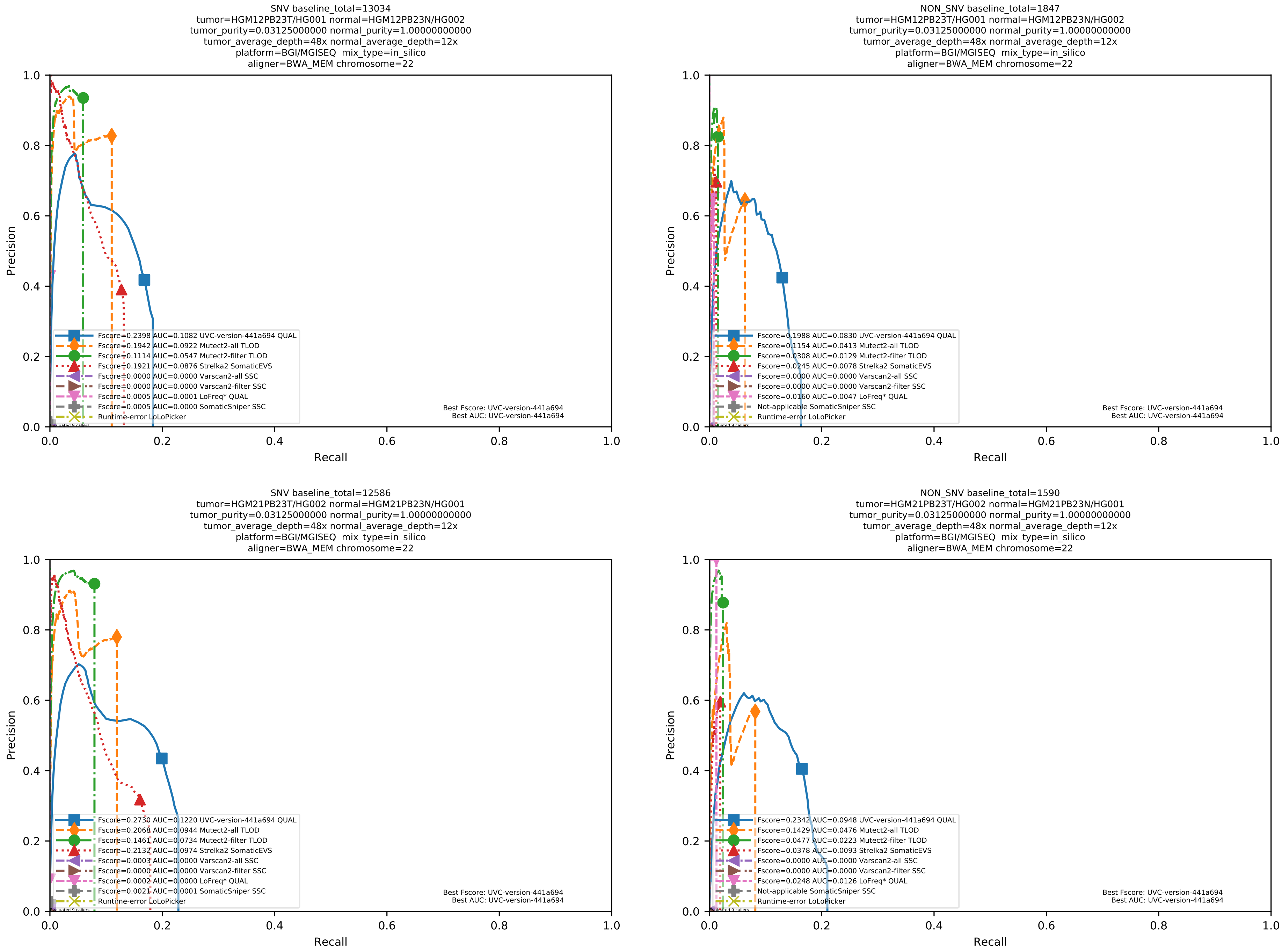

Supplementary Figure 95: Performance of somatic variant callers in tumor-with-normal mode on the *in silico* mixtures with BGI MGISEQ sequencing data aligned with BWA-MEM to the human reference genome hs37d5.

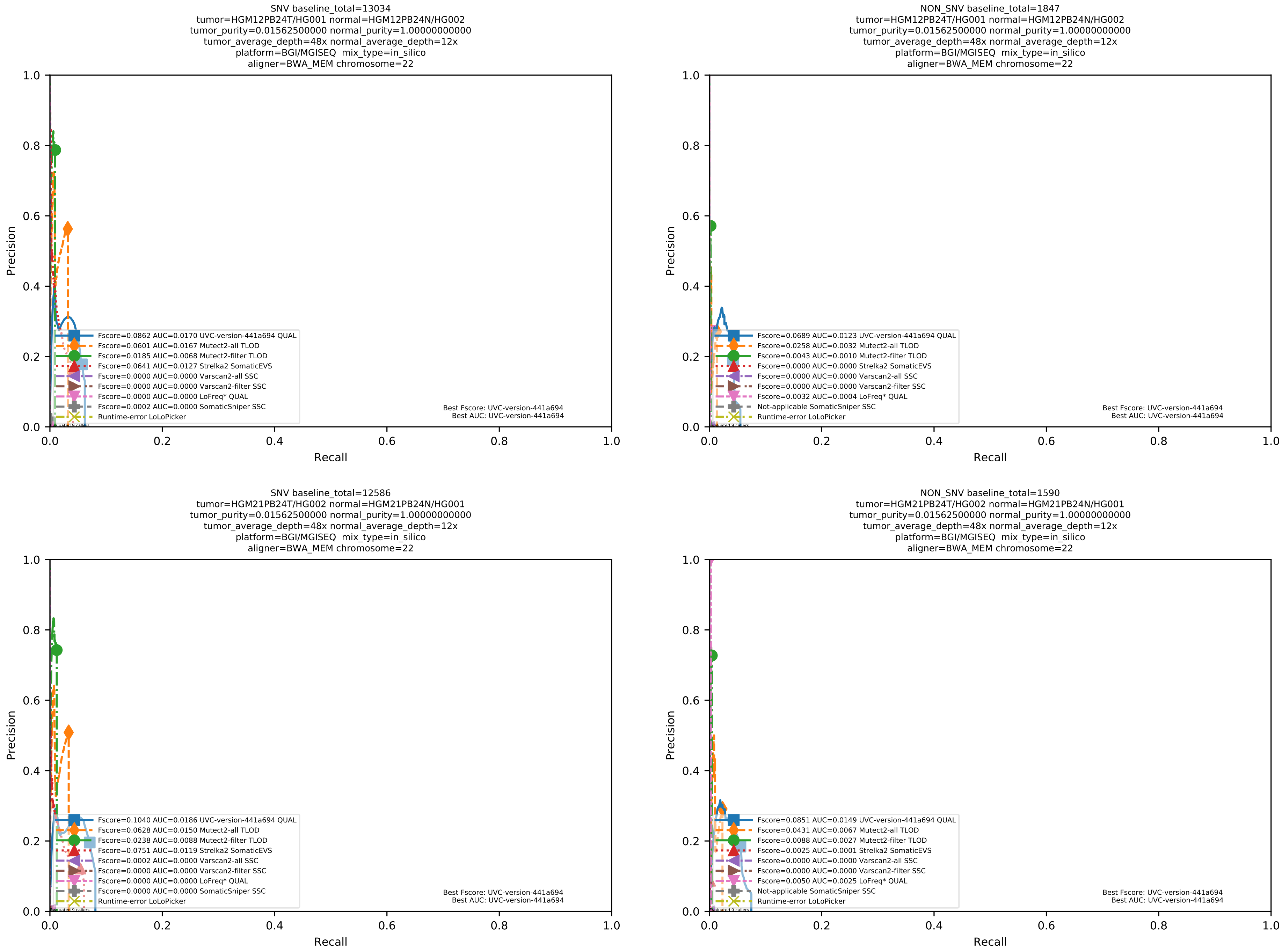

Supplementary Figure 96: Performance of somatic variant callers in tumor-with-normal mode on the *in silico* mixtures with BGI MGISEQ sequencing data aligned with BWA-MEM to the human reference genome hs37d5.

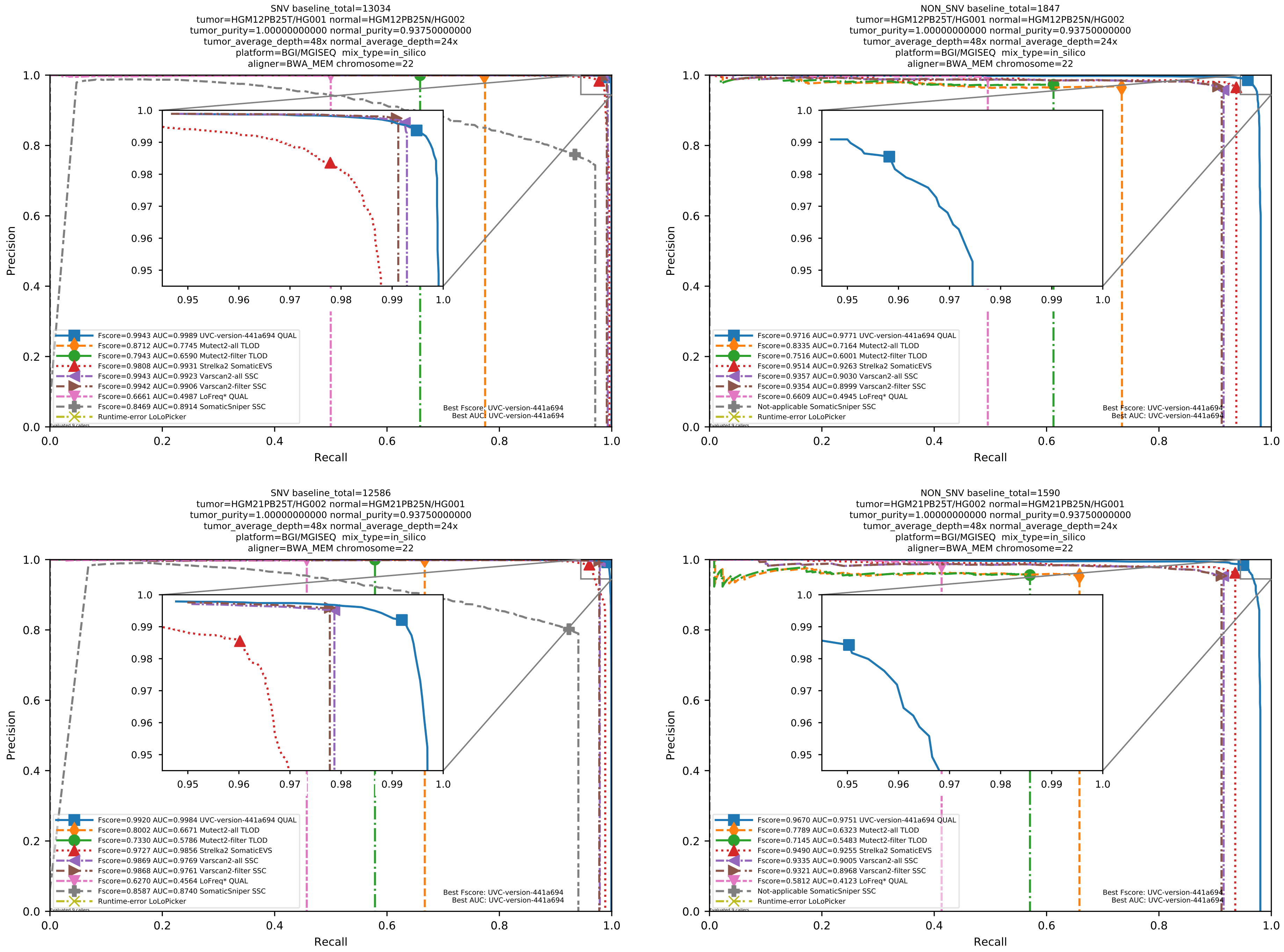

Supplementary Figure 97: Performance of somatic variant callers in tumor-with-normal mode on the *in silico* mixtures with BGI MGISEQ sequencing data aligned with BWA-MEM to the human reference genome hs37d5.

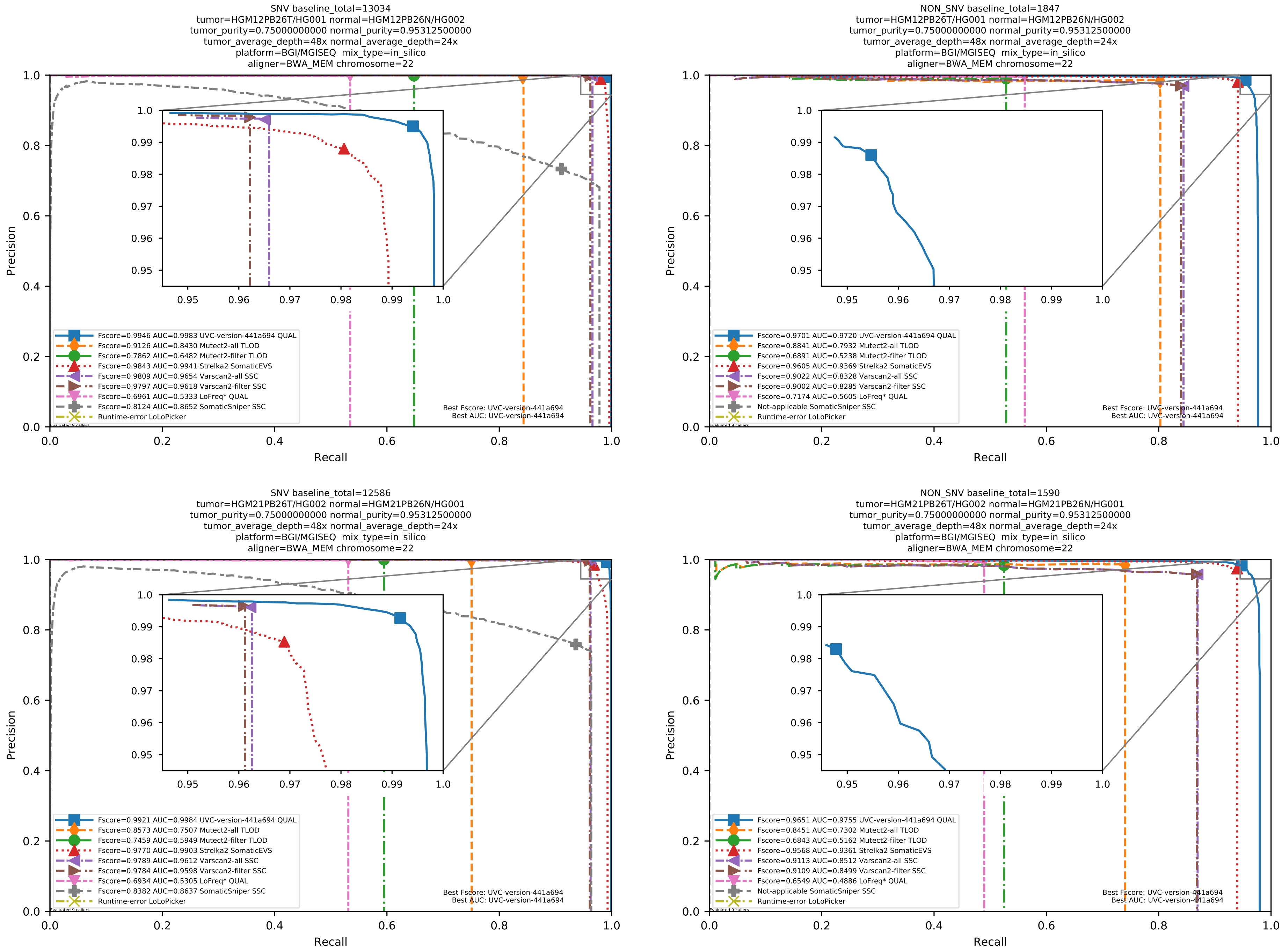

Supplementary Figure 98: Performance of somatic variant callers in tumor-with-normal mode on the *in silico* mixtures with BGI MGISEQ sequencing data aligned with BWA-MEM to the human reference genome hs37d5.

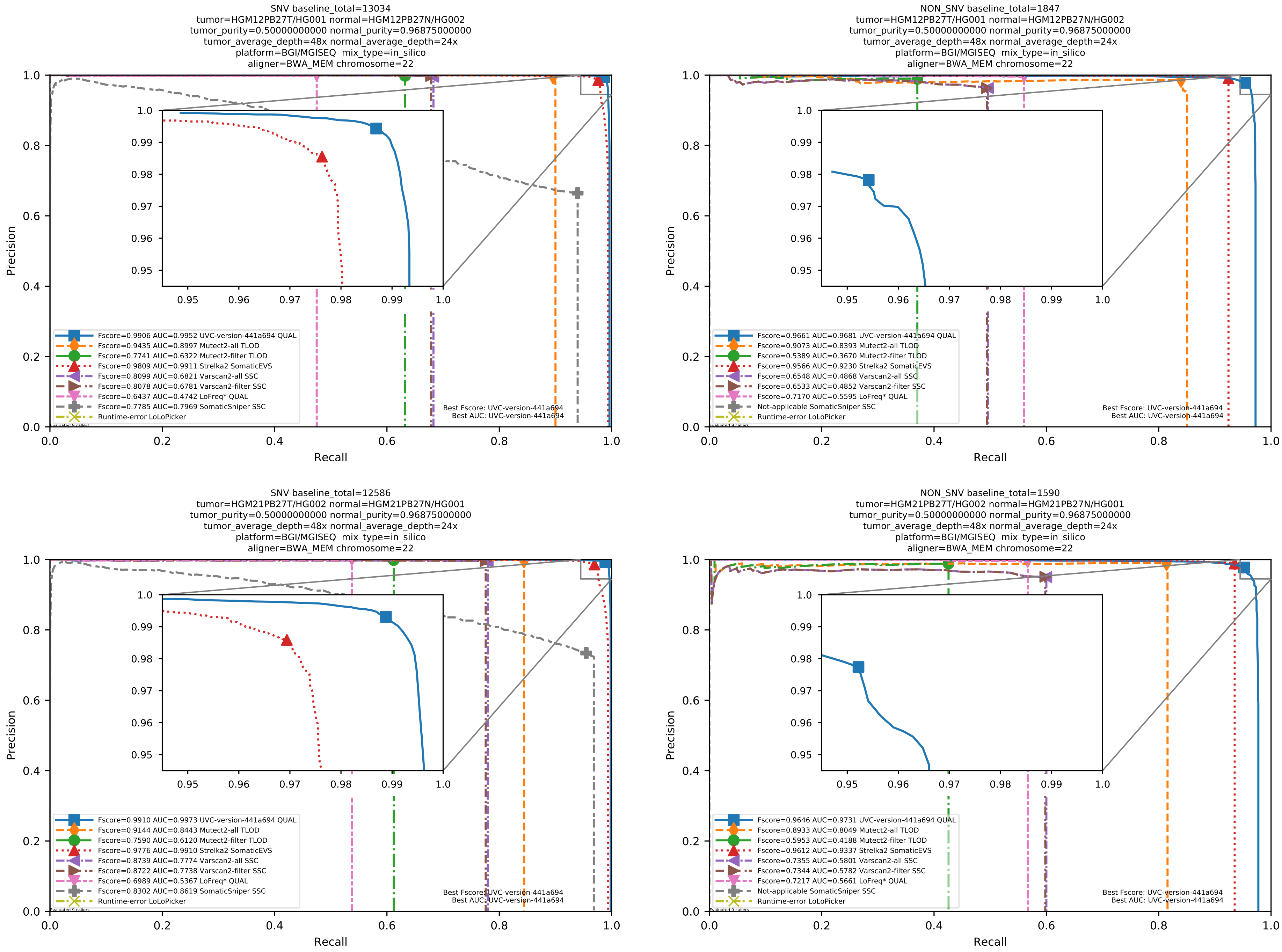

Supplementary Figure 99: Performance of somatic variant callers in tumor-with-normal mode on the *in silico* mixtures with BGI MGISEQ sequencing data aligned with BWA-MEM to the human reference genome hs37d5.

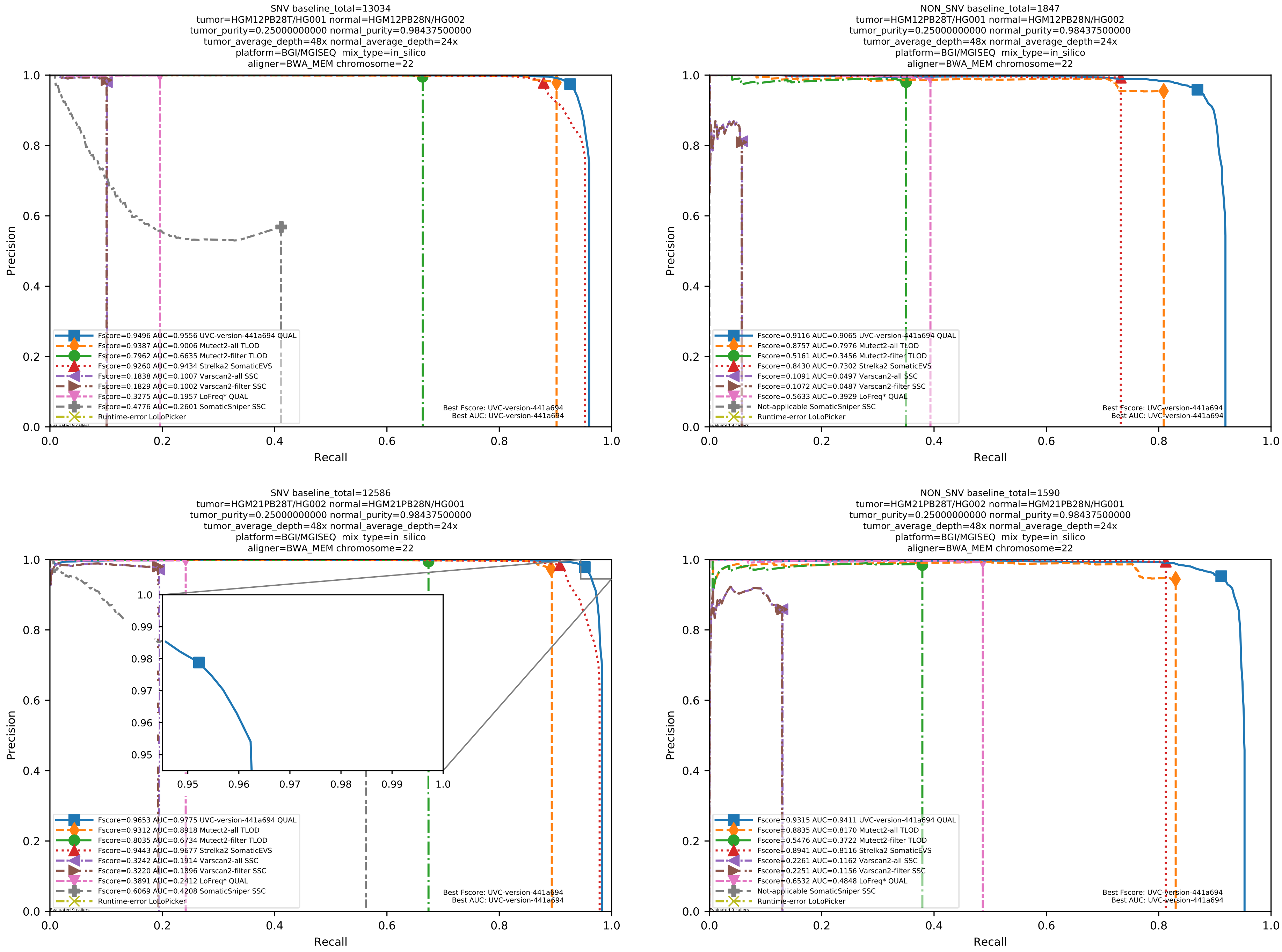

Supplementary Figure 100: Performance of somatic variant callers in tumor-with-normal mode on the *in silico* mixtures with BGI MGISEQ sequencing data aligned with BWA-MEM to the human reference genome hs37d5.

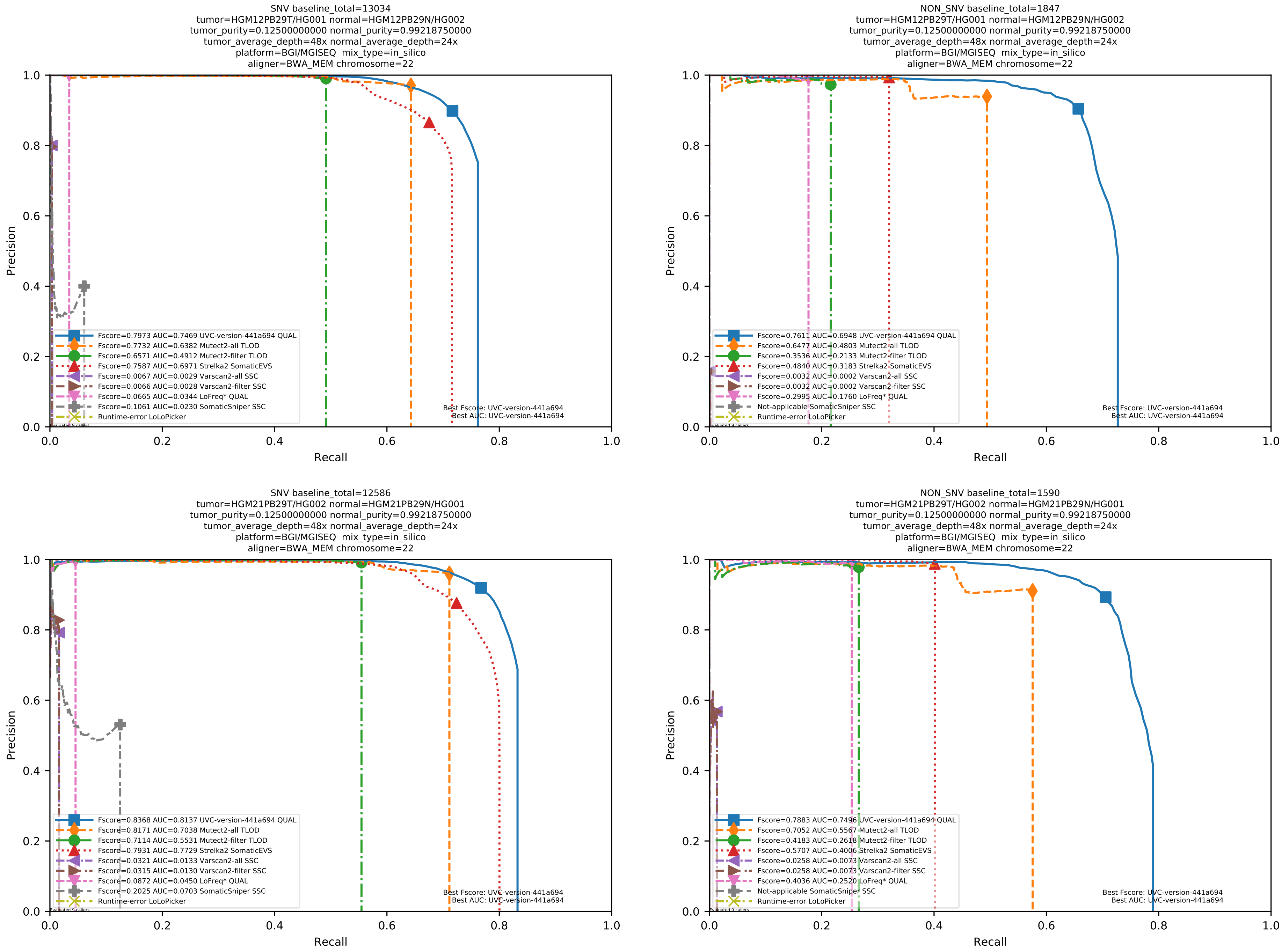

Supplementary Figure 101: Performance of somatic variant callers in tumor-with-normal mode on the *in silico* mixtures with BGI MGISEQ sequencing data aligned with BWA-MEM to the human reference genome hs37d5.

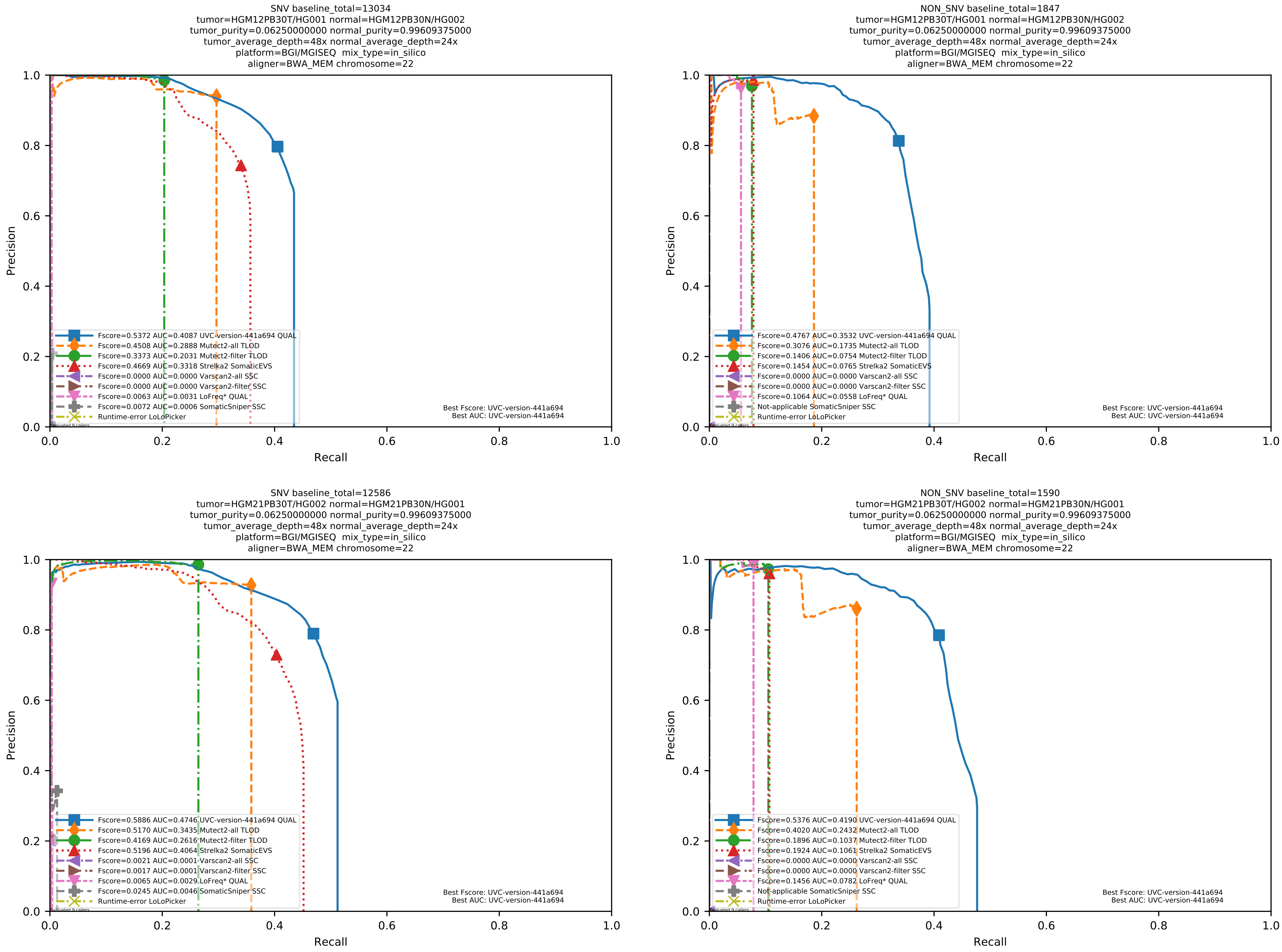

Supplementary Figure 102: Performance of somatic variant callers in tumor-with-normal mode on the *in silico* mixtures with BGI MGISEQ sequencing data aligned with BWA-MEM to the human reference genome hs37d5.

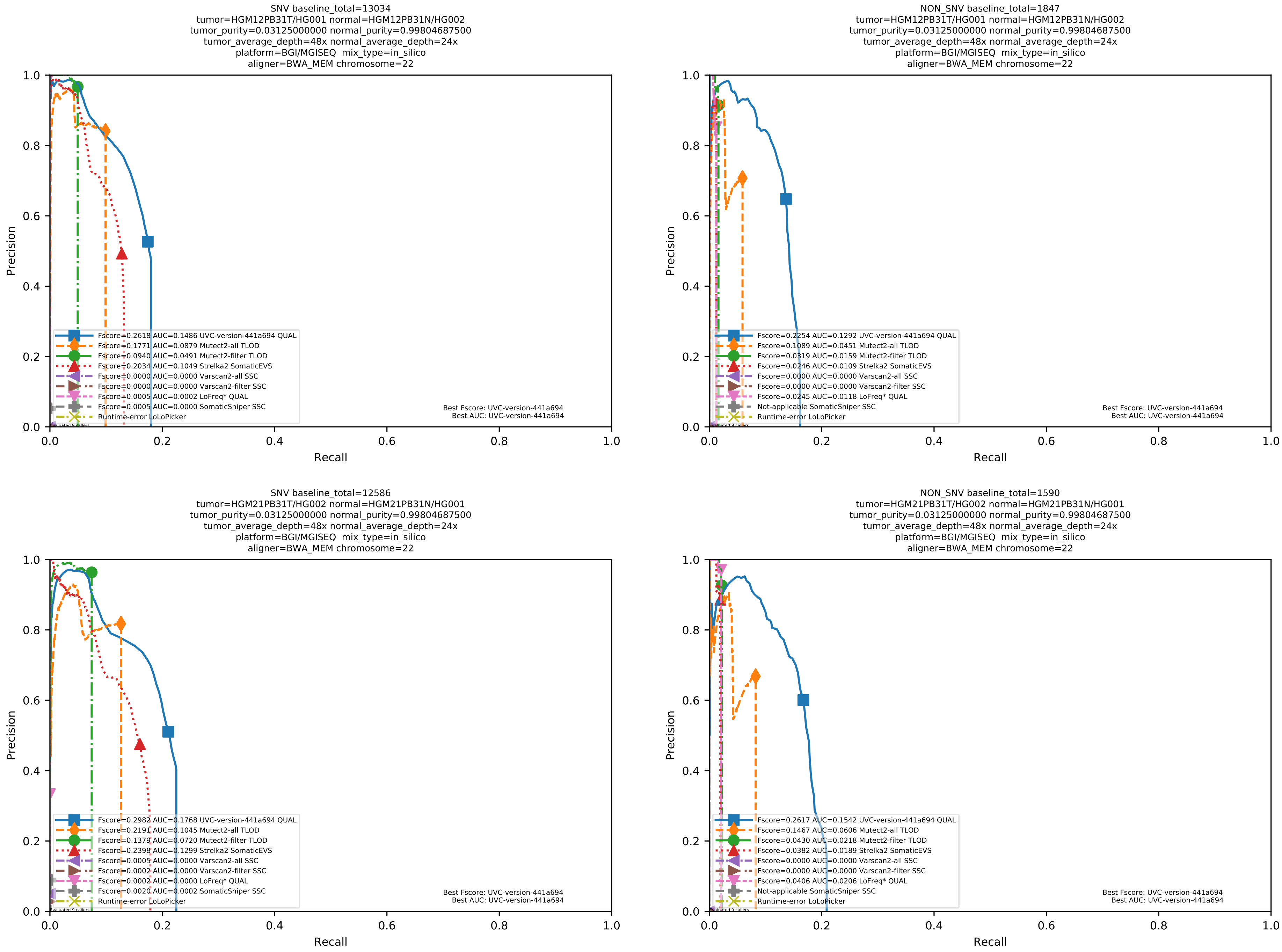

Supplementary Figure 103: Performance of somatic variant callers in tumor-with-normal mode on the *in silico* mixtures with BGI MGISEQ sequencing data aligned with BWA-MEM to the human reference genome hs37d5.

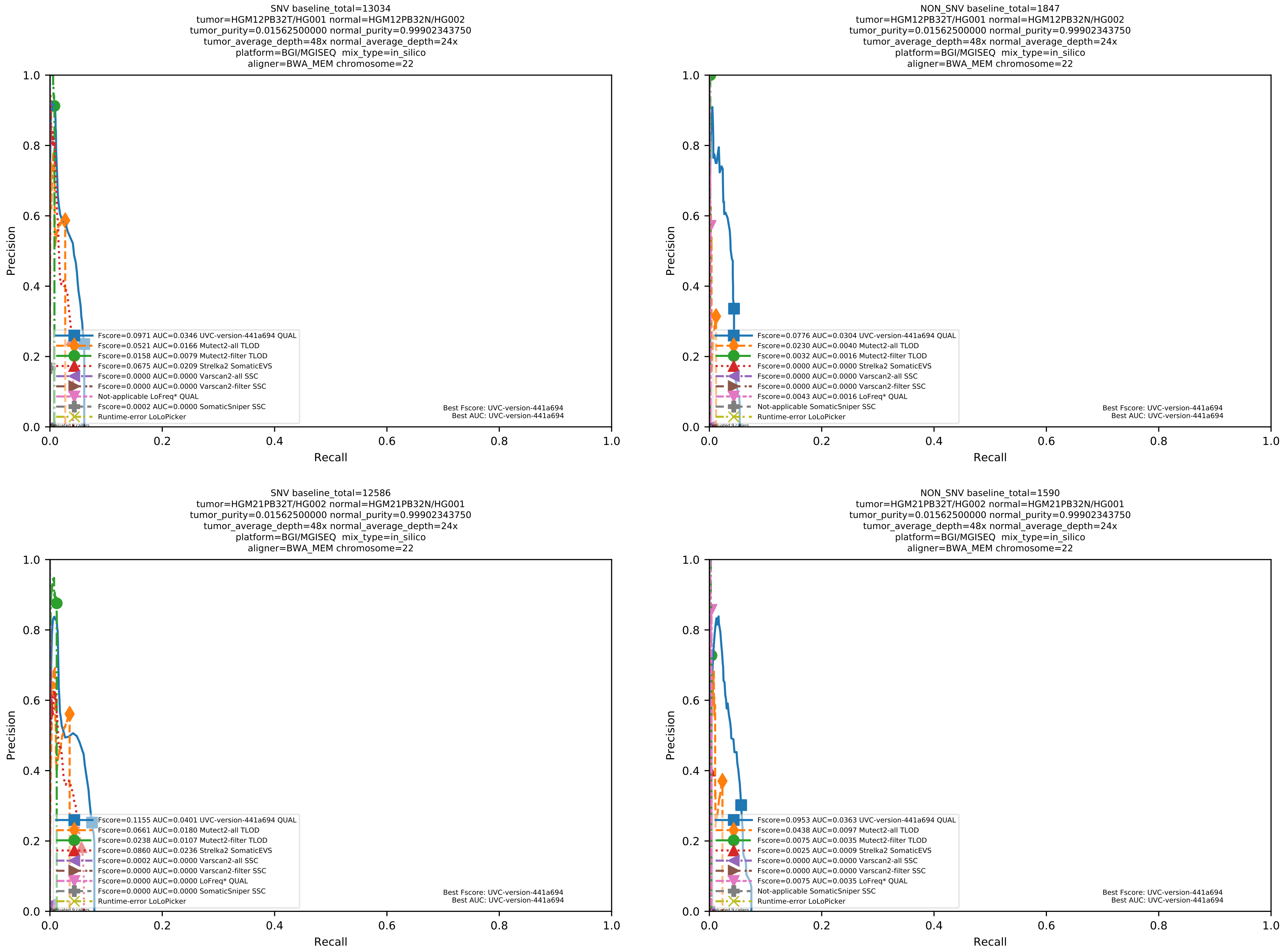

Supplementary Figure 104: Performance of somatic variant callers in tumor-with-normal mode on the *in silico* mixtures with BGI MGISEQ sequencing data aligned with BWA-MEM to the human reference genome hs37d5.

##### 1.5 Supplemental evaluation results on WGS datasets for calling somatic variants from tumor-normal pairs, with physical mixtures

Additional data described in the file at `ftp://ftp-trace.ncbi.nlm.nih.gov/giab/ftp/use_cases/mixtures/UMCUTRECHT_NA12878_NA24385_mixture_10052016/README-NA12878_NA24385_mixture.txt` were used. Evaluation procedures for all applicable variant callers all followed the best practice at `ftp://ftp-trace.ncbi.nlm.nih.gov/giab/ftp/release/NA12878_HG001/NISTv3.3.2/README_NISTv3.3.2.txt`.

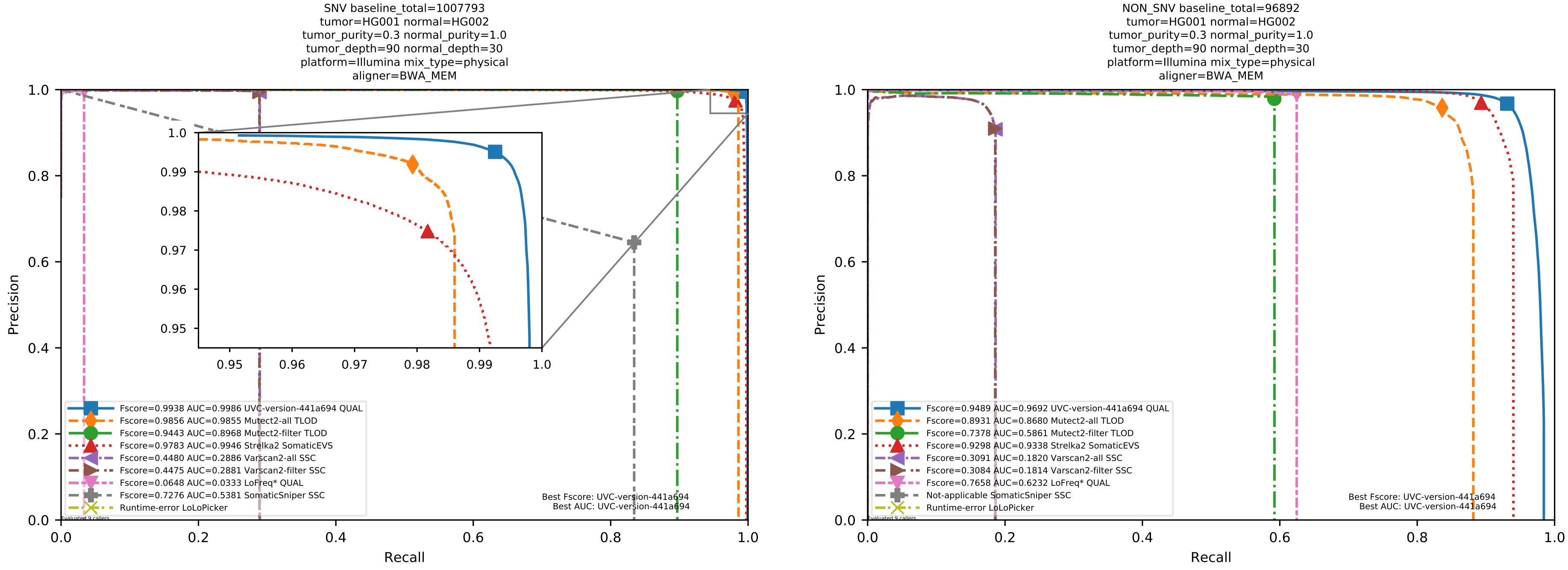

Supplementary Figure 105: Performance of somatic variant callers in tumor-with-normal mode on the physical mixture with Illumina sequencing data aligned with BWA-MEM to the human reference genome hs37d5.

#### 1.6 Supplemental evaluation results on WES and amplicon-sequencing datasets for calling somatic variants from tumor-normal pairs, with the breast-cancer cell-line HCC1395

The datasets used in this subsection were presented in more detail by Fang et al.[2]. GRCh38 is the only human reference genome with respect to which the high-confidence calls were inferred. Hence, we aligned all sequencing data to GRCh38 in this subsection by using BWA MEM.

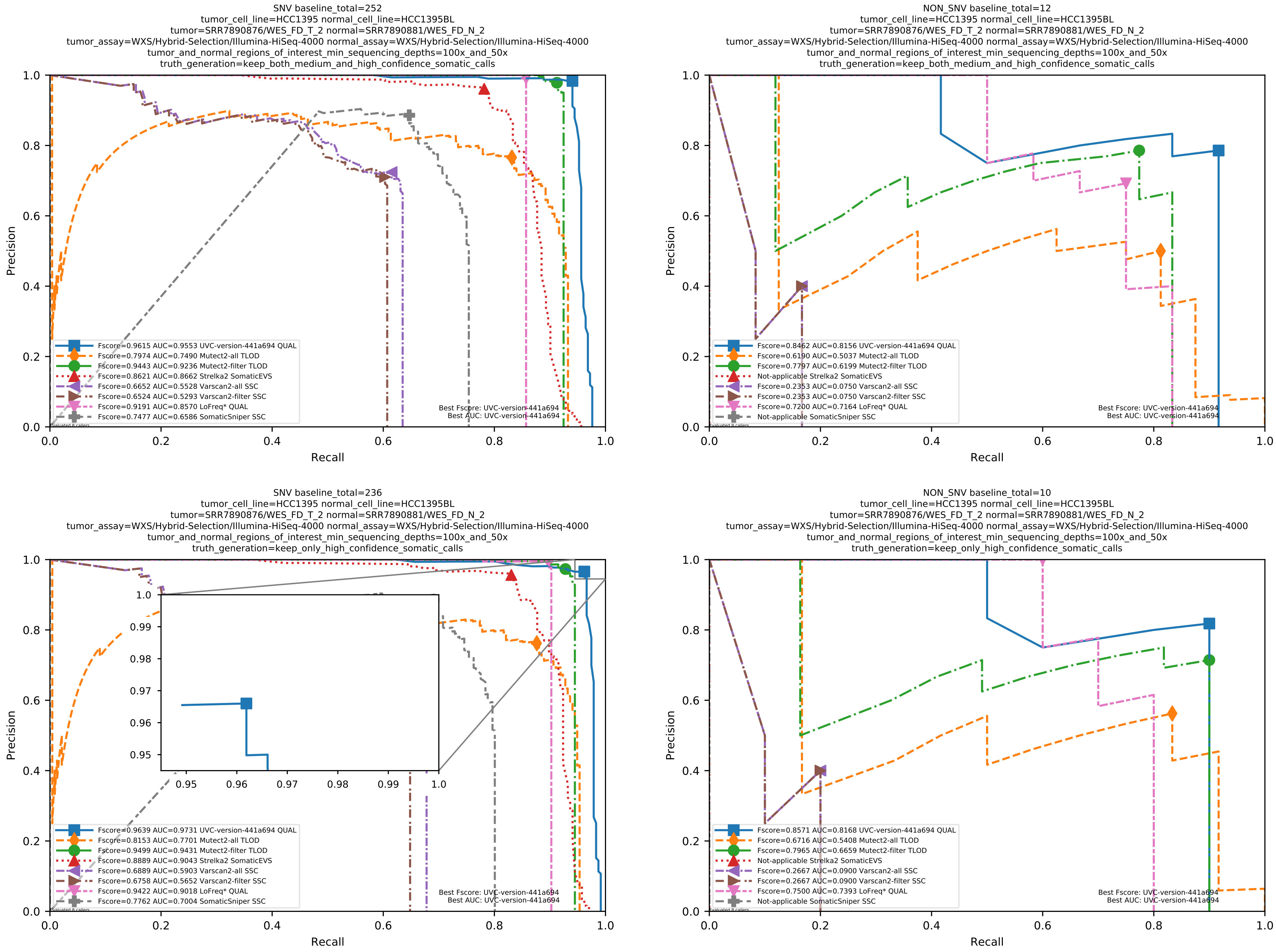

Supplementary Figure 106: Performance of somatic variant callers on the SEQC2 somatic reference sets with Illumina sequencing data aligned with BWA MEM to the GRCh38 human-genome reference.

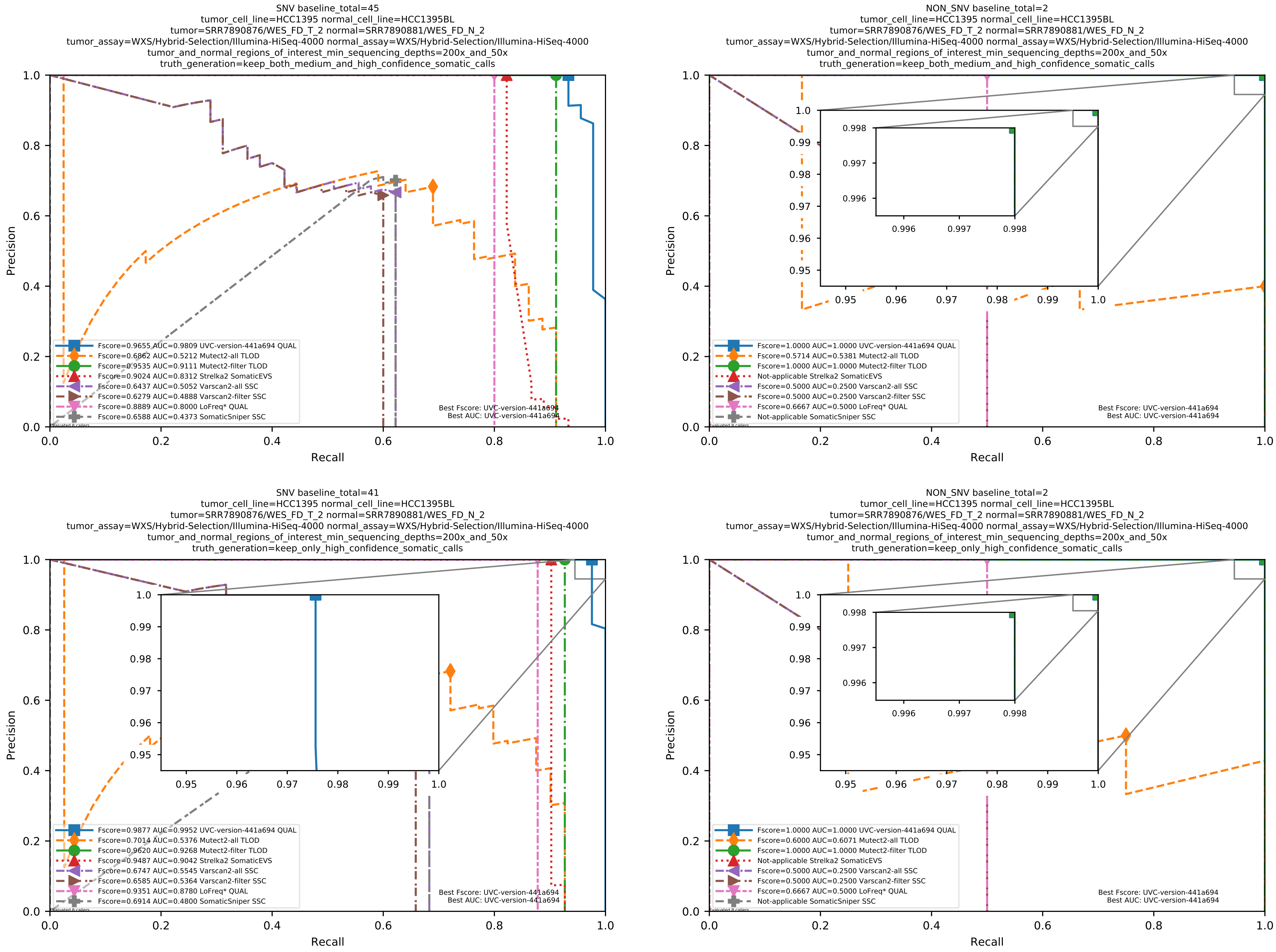

Supplementary Figure 107: Performance of somatic variant callers on the SEQC2 somatic reference sets with Illumina sequencing data aligned with BWA MEM to the GRCh38 human-genome reference.

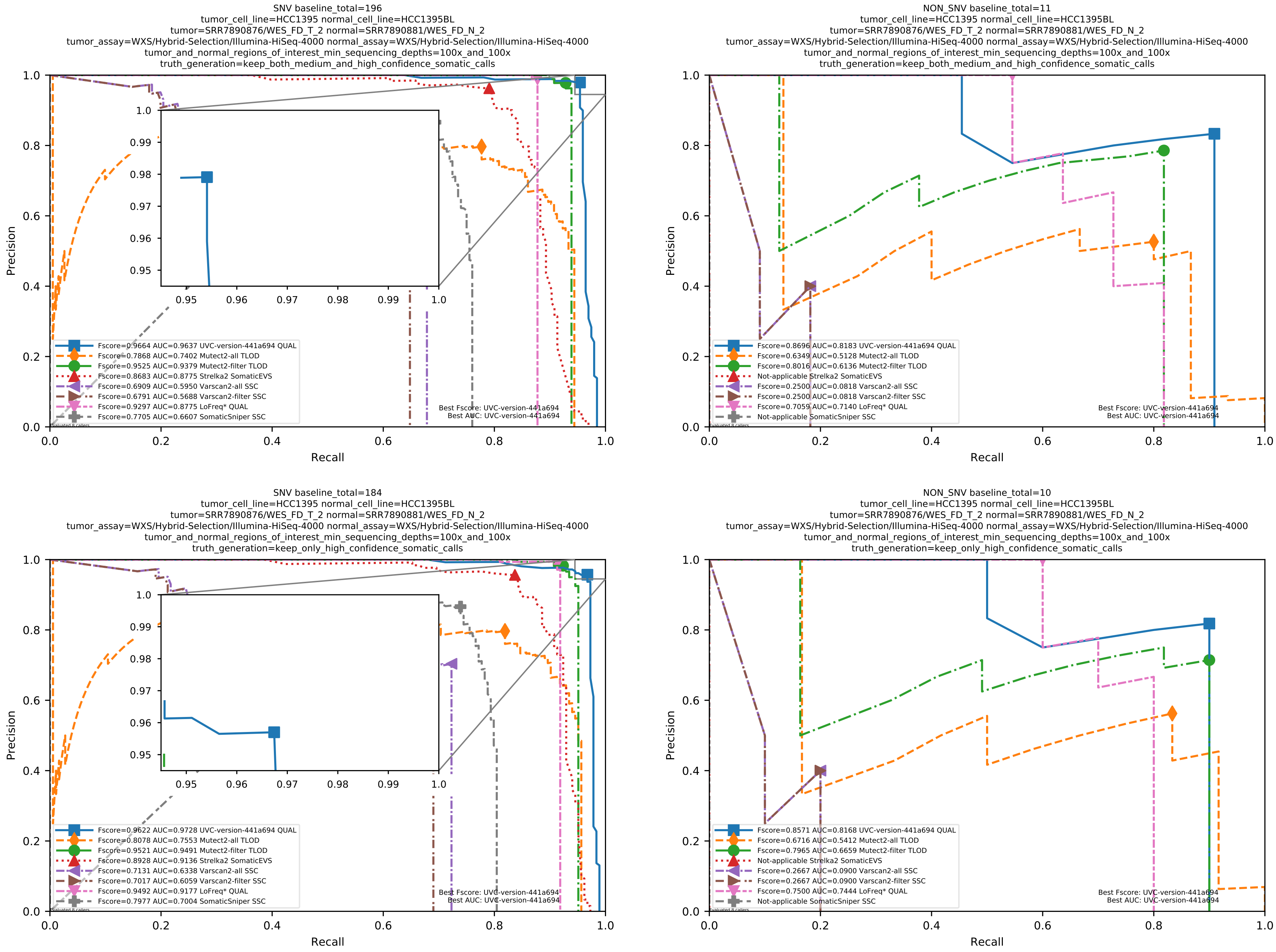

Supplementary Figure 108: Performance of somatic variant callers on the SEQC2 somatic reference sets with Illumina sequencing data aligned with BWA MEM to the GRCh38 human-genome reference.

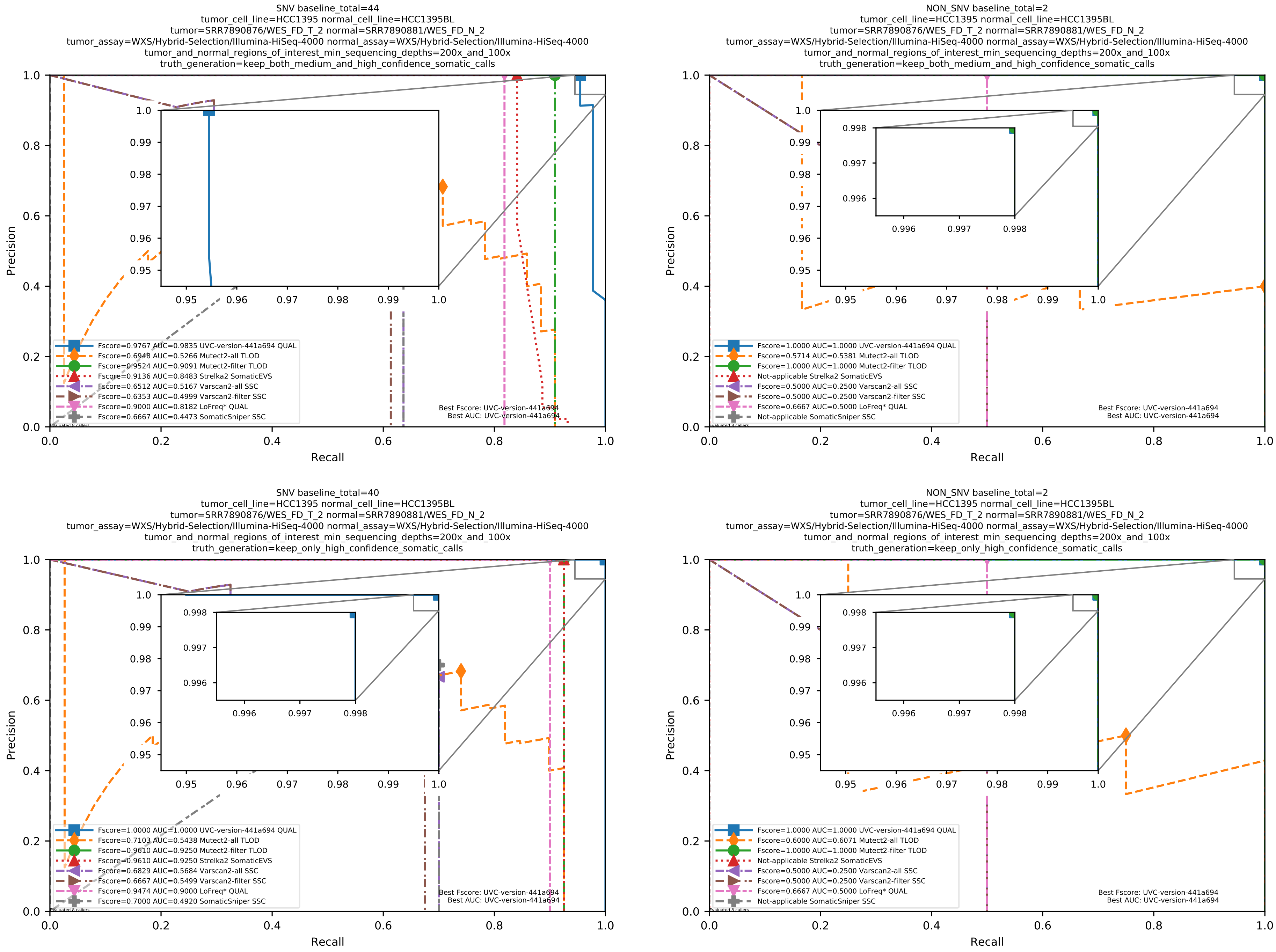

Supplementary Figure 109: Performance of somatic variant callers on the SEQC2 somatic reference sets with Illumina sequencing data aligned with BWA MEM to the GRCh38 human-genome reference.

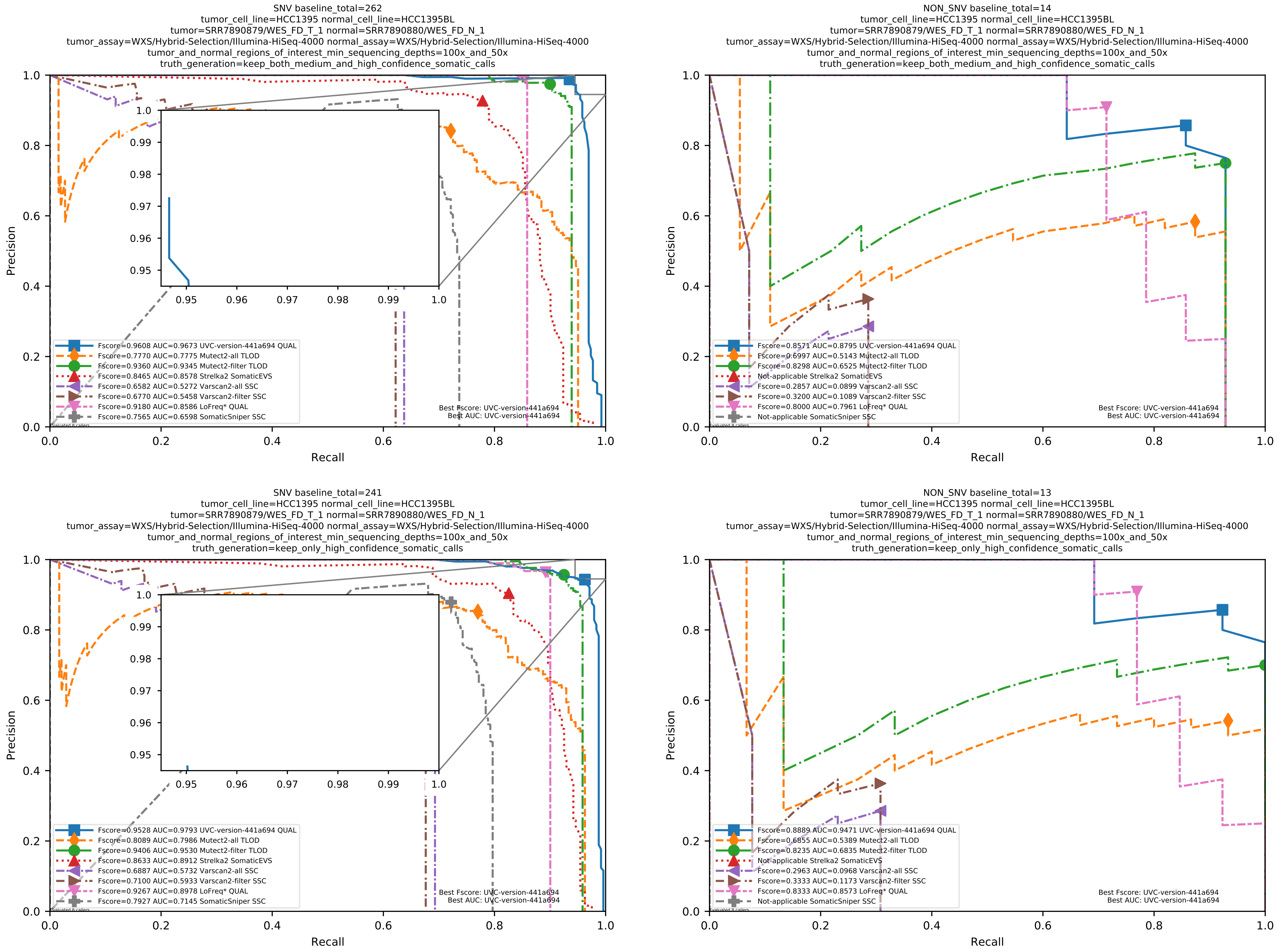

Supplementary Figure 110: Performance of somatic variant callers on the SEQC2 somatic reference sets with Illumina sequencing data aligned with BWA MEM to the GRCh38 human-genome reference.

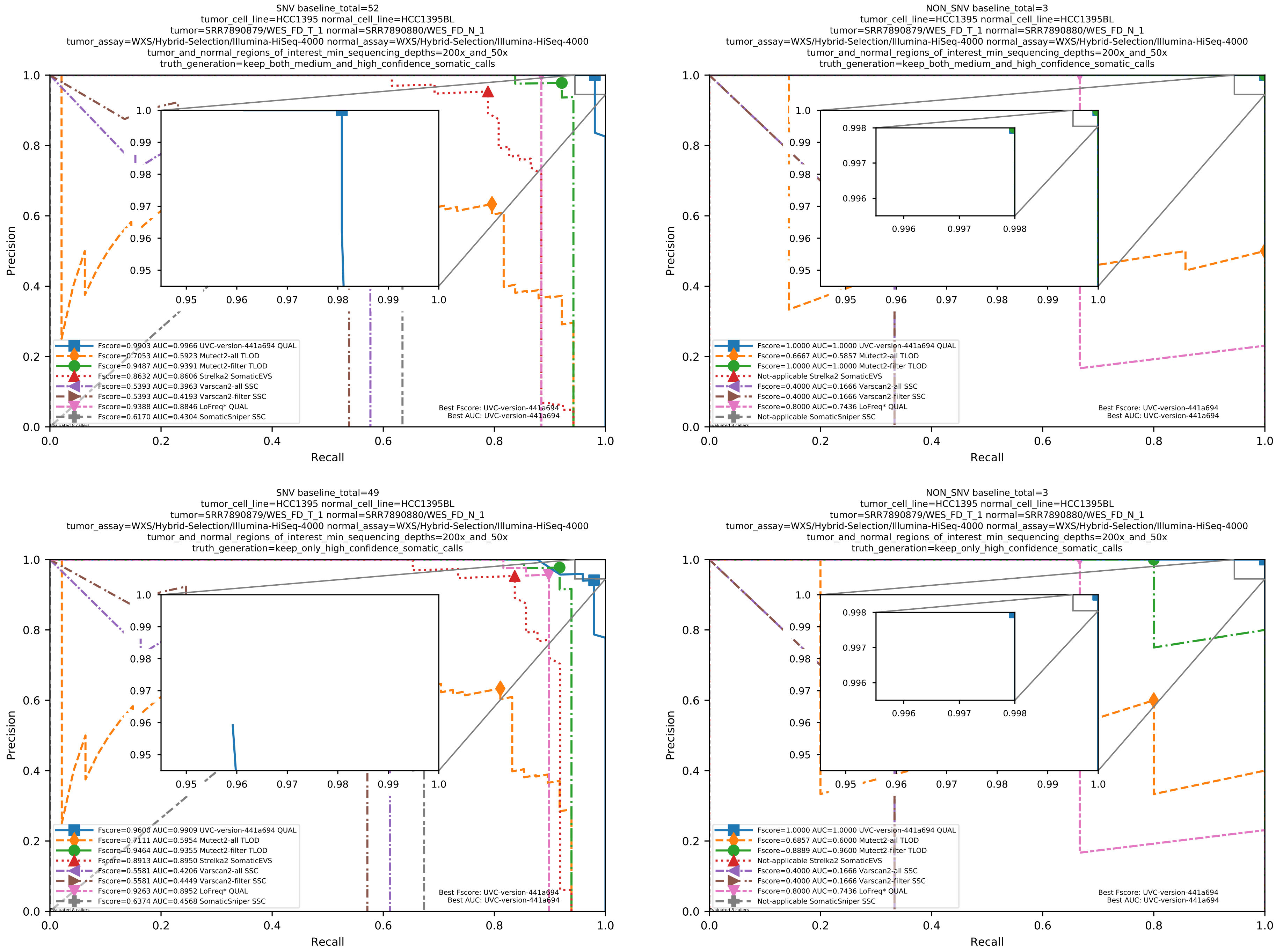

Supplementary Figure 111: Performance of somatic variant callers on the SEQC2 somatic reference sets with Illumina sequencing data aligned with BWA MEM to the GRCh38 human-genome reference.

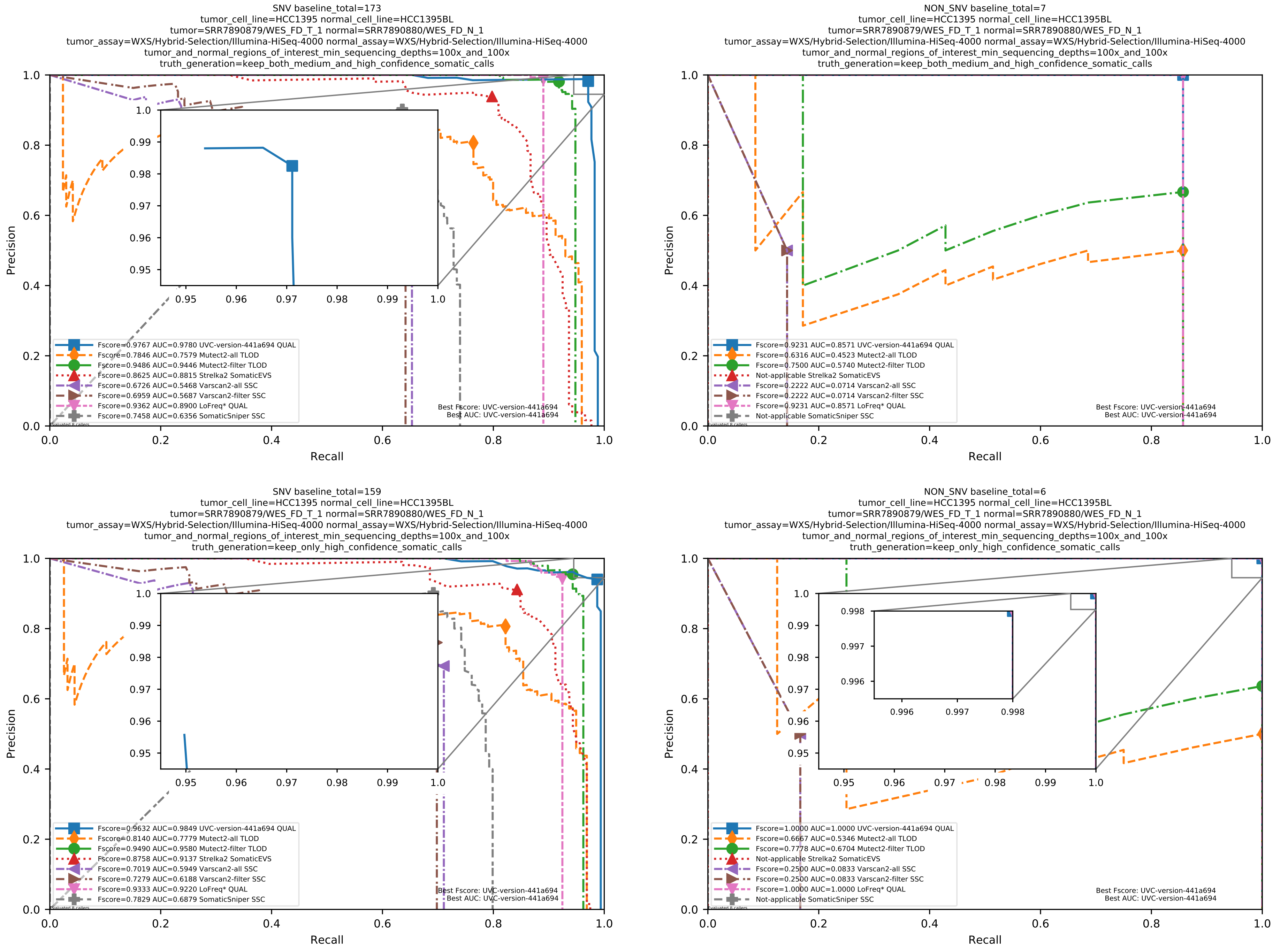

Supplementary Figure 112: Performance of somatic variant callers on the SEQC2 somatic reference sets with Illumina sequencing data aligned with BWA MEM to the GRCh38 human-genome reference.

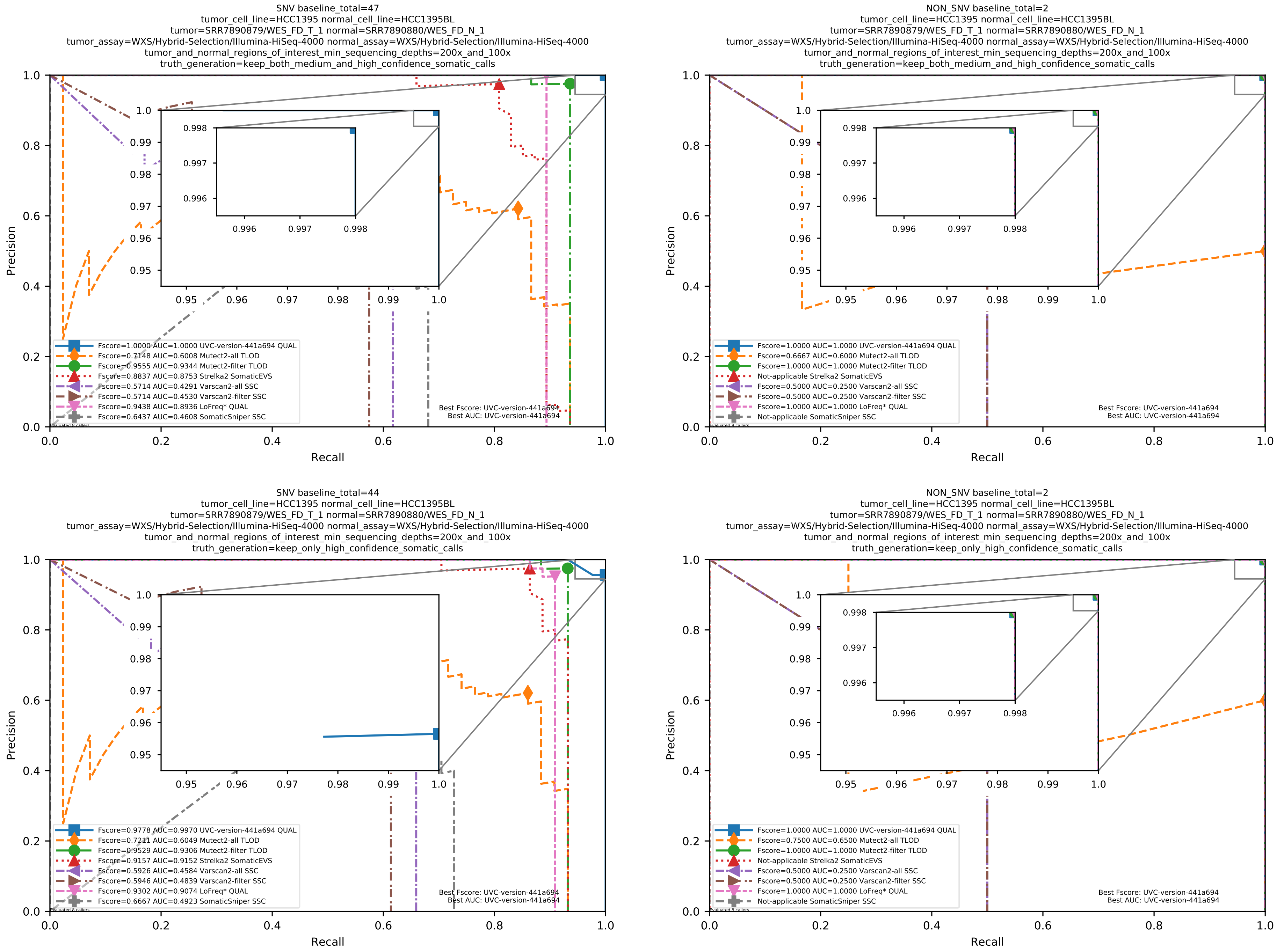

Supplementary Figure 113: Performance of somatic variant callers on the SEQC2 somatic reference sets with Illumina sequencing data aligned with BWA MEM to the GRCh38 human-genome reference.

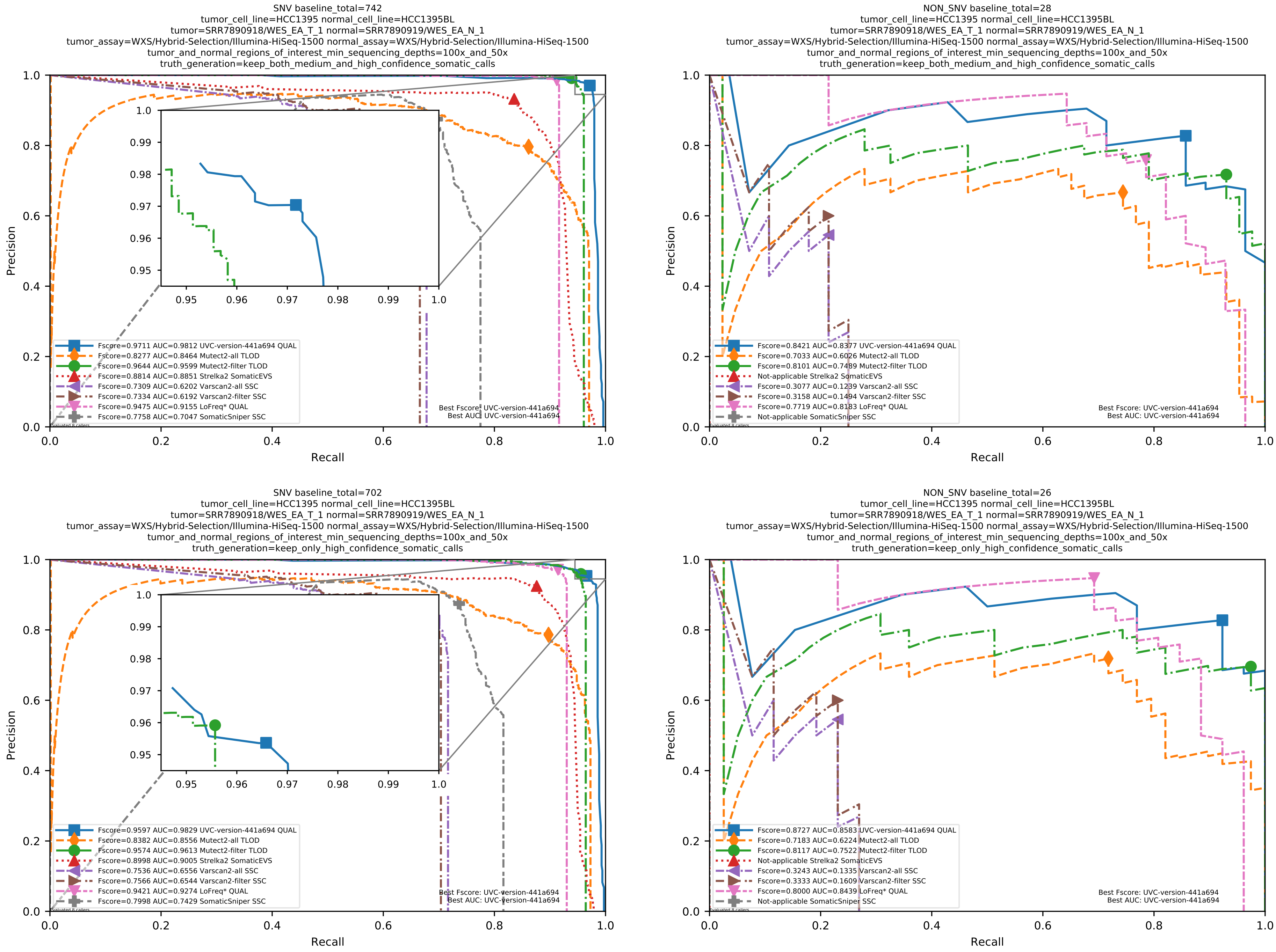

Supplementary Figure 114: Performance of somatic variant callers on the SEQC2 somatic reference sets with Illumina sequencing data aligned with BWA MEM to the GRCh38 human-genome reference.

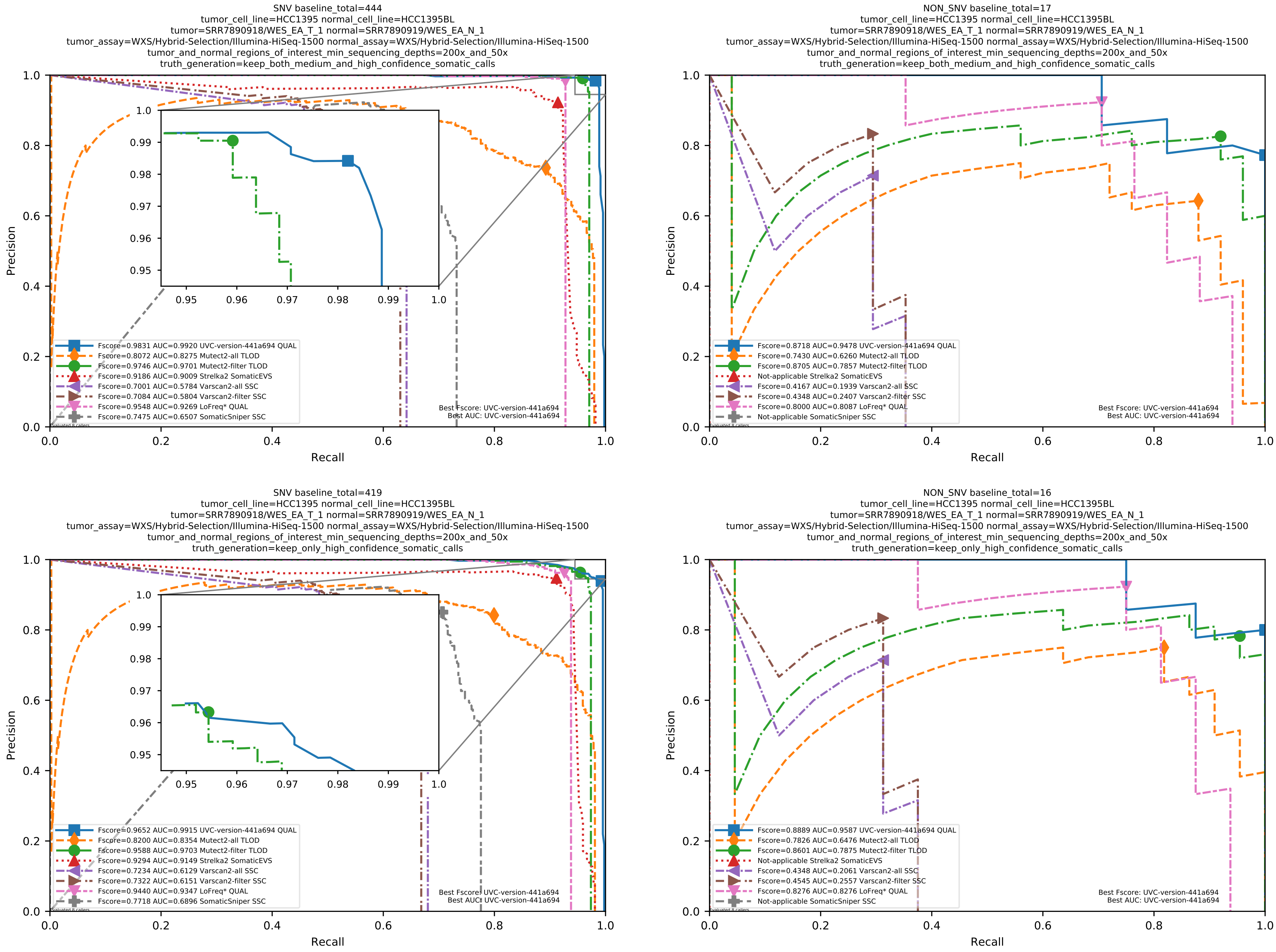

Supplementary Figure 115: Performance of somatic variant callers on the SEQC2 somatic reference sets with Illumina sequencing data aligned with BWA MEM to the GRCh38 human-genome reference.

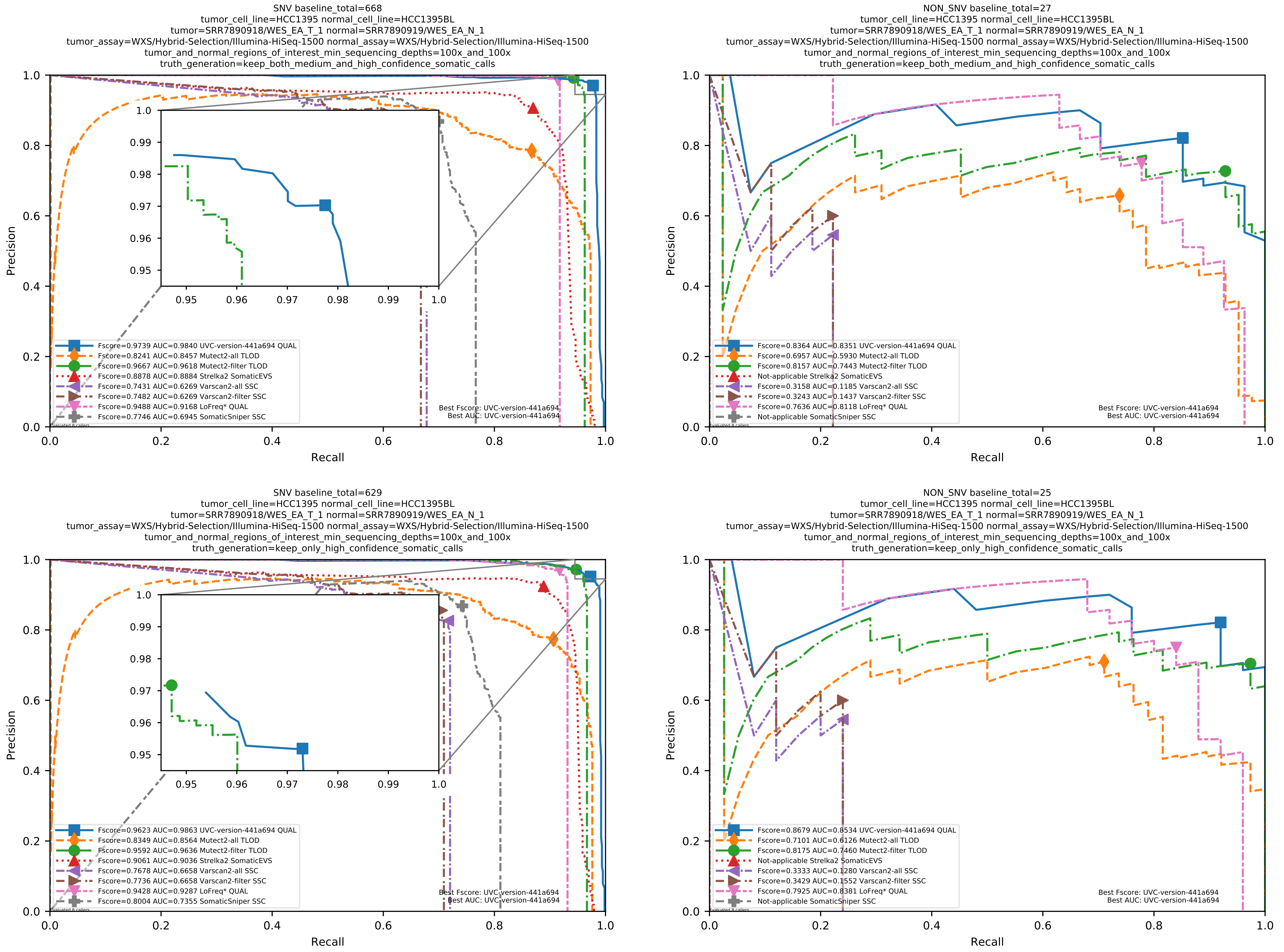

Supplementary Figure 116: Performance of somatic variant callers on the SEQC2 somatic reference sets with Illumina sequencing data aligned with BWA MEM to the GRCh38 human-genome reference.

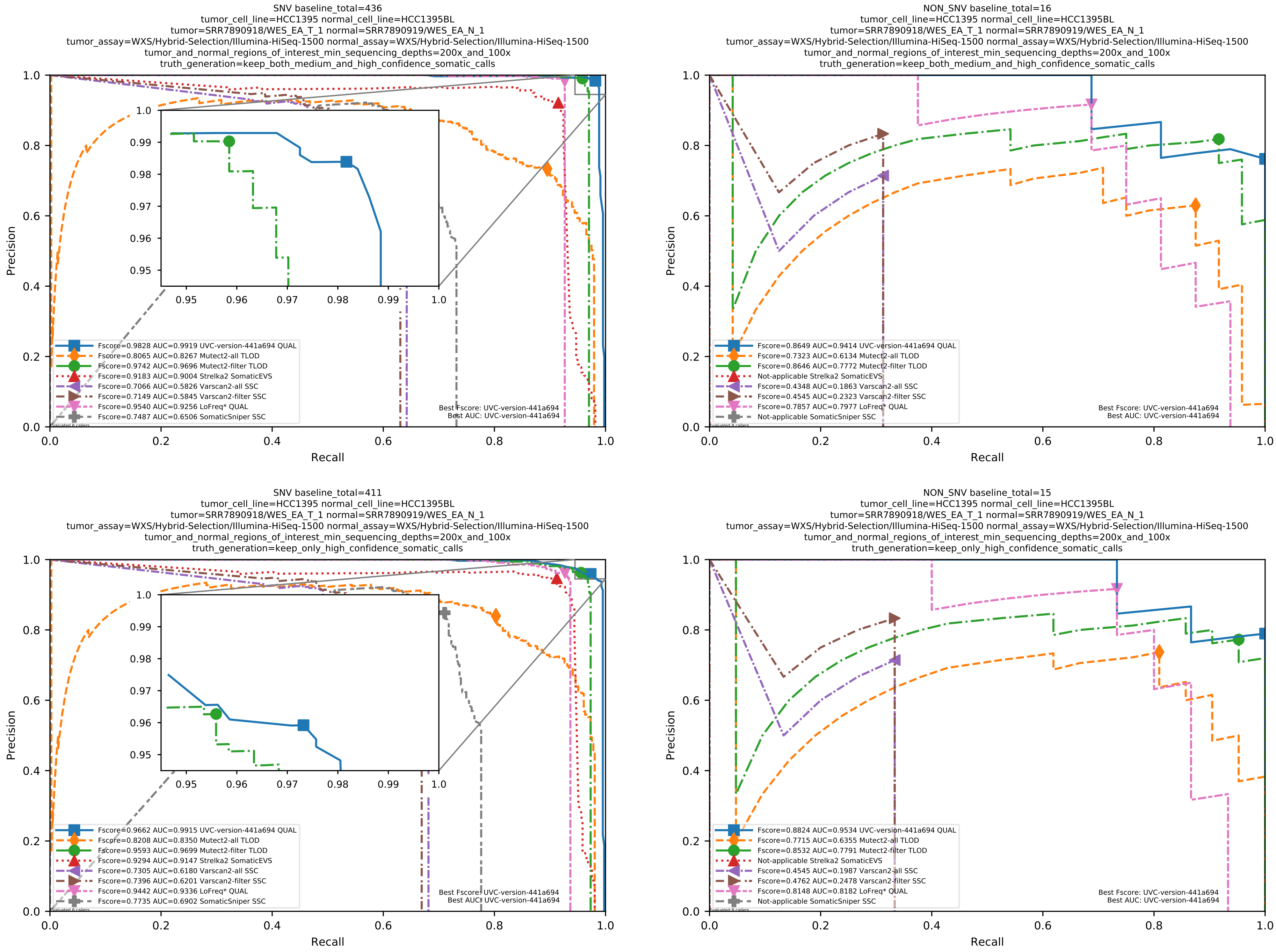

Supplementary Figure 117: Performance of somatic variant callers on the SEQC2 somatic reference sets with Illumina sequencing data aligned with BWA MEM to the GRCh38 human-genome reference.

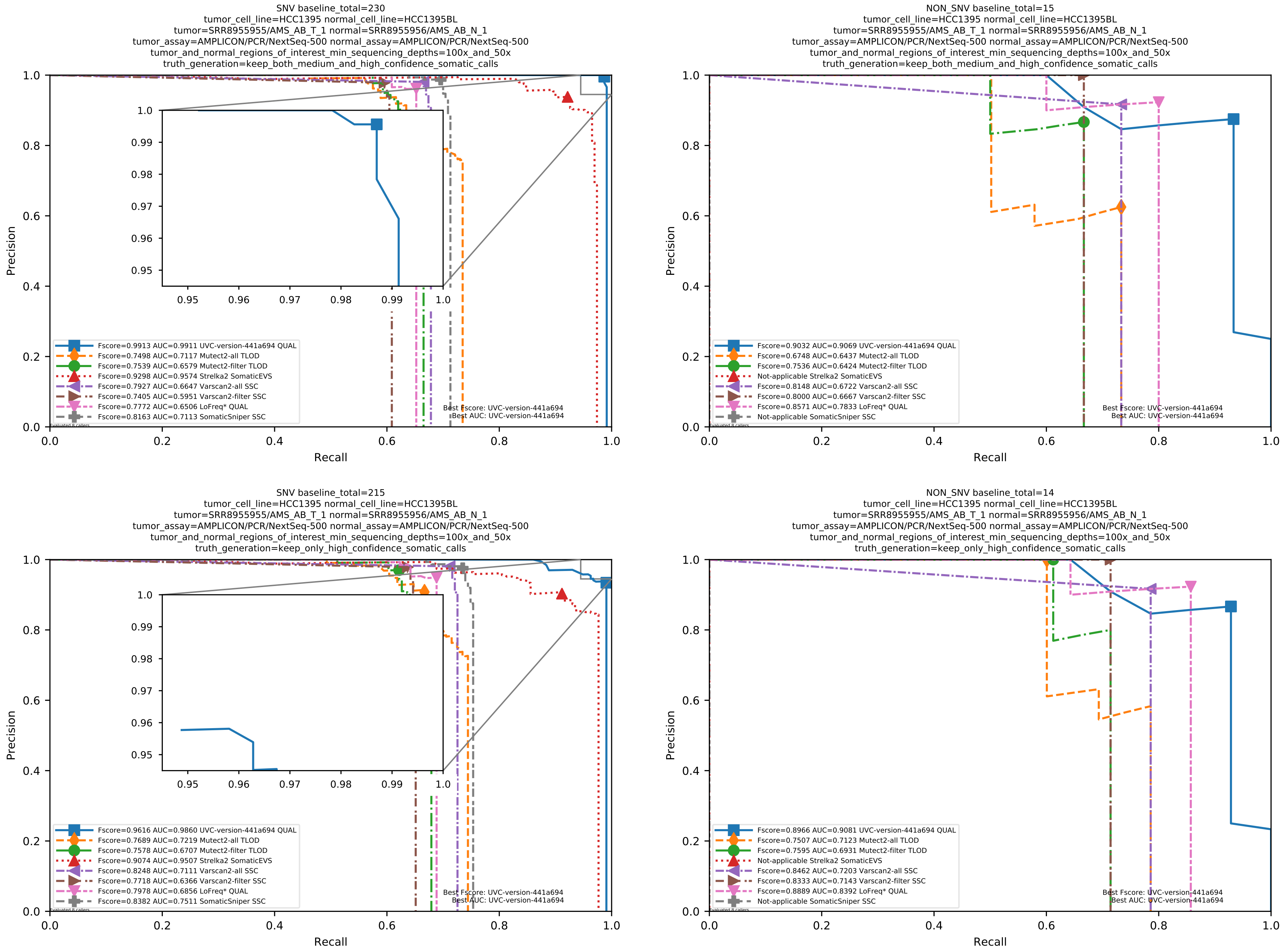

Supplementary Figure 118: Performance of somatic variant callers on the SEQC2 somatic reference sets with Illumina sequencing data aligned with BWA MEM to the GRCh38 human-genome reference.

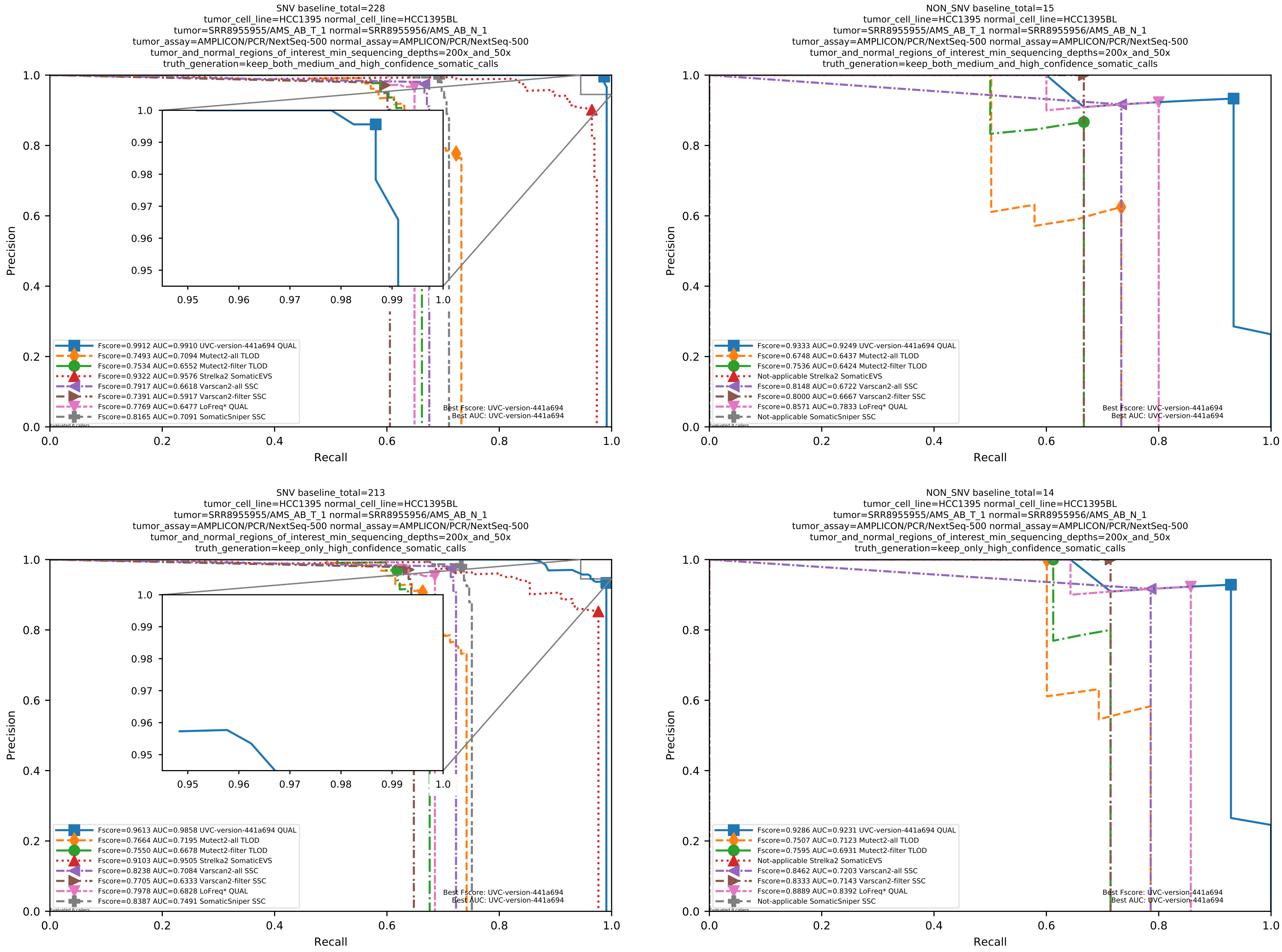

Supplementary Figure 119: Performance of somatic variant callers on the SEQC2 somatic reference sets with Illumina sequencing data aligned with BWA MEM to the GRCh38 human-genome reference.

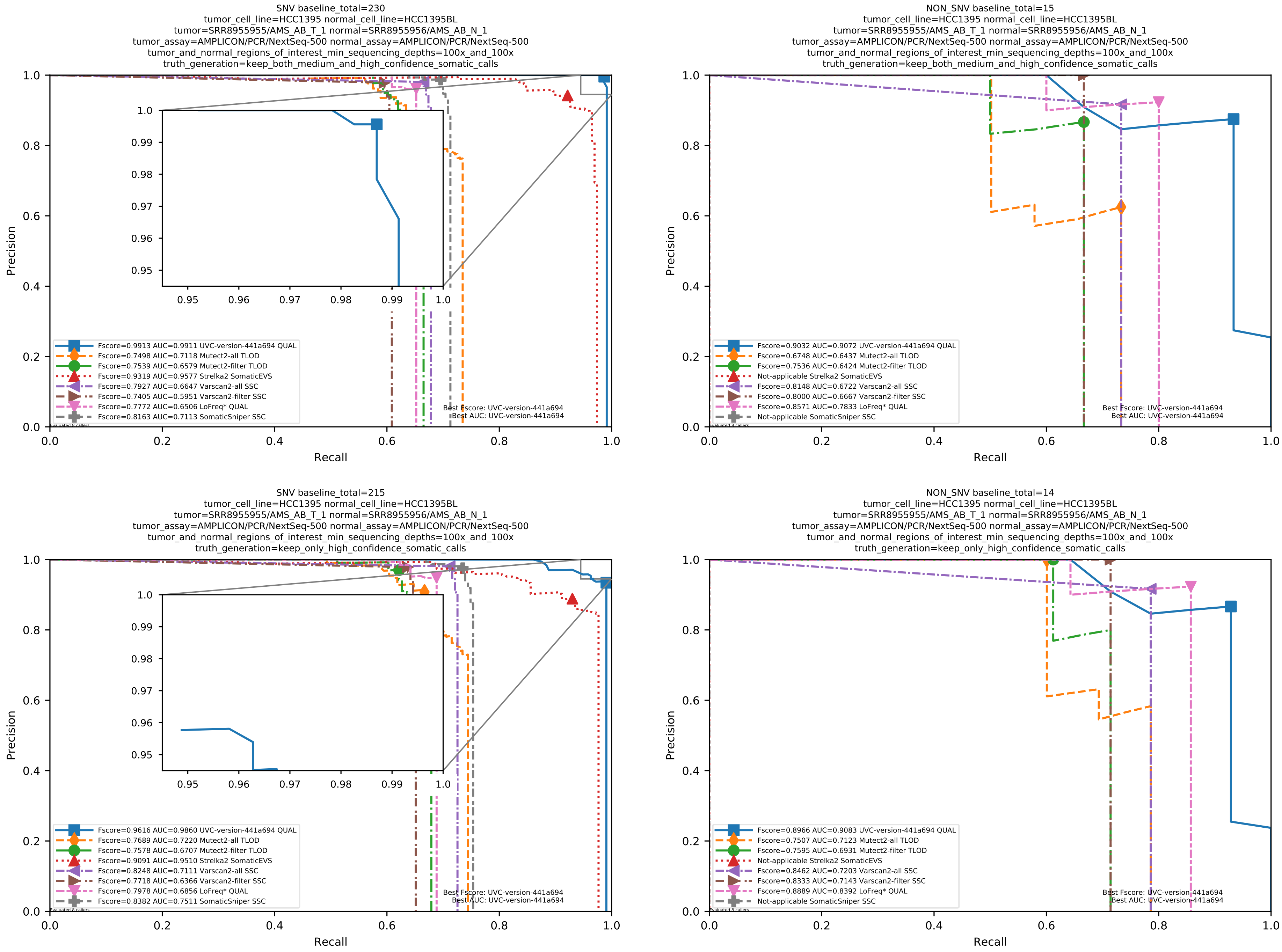

Supplementary Figure 120: Performance of somatic variant callers on the SEQC2 somatic reference sets with Illumina sequencing data aligned with BWA MEM to the GRCh38 human-genome reference.

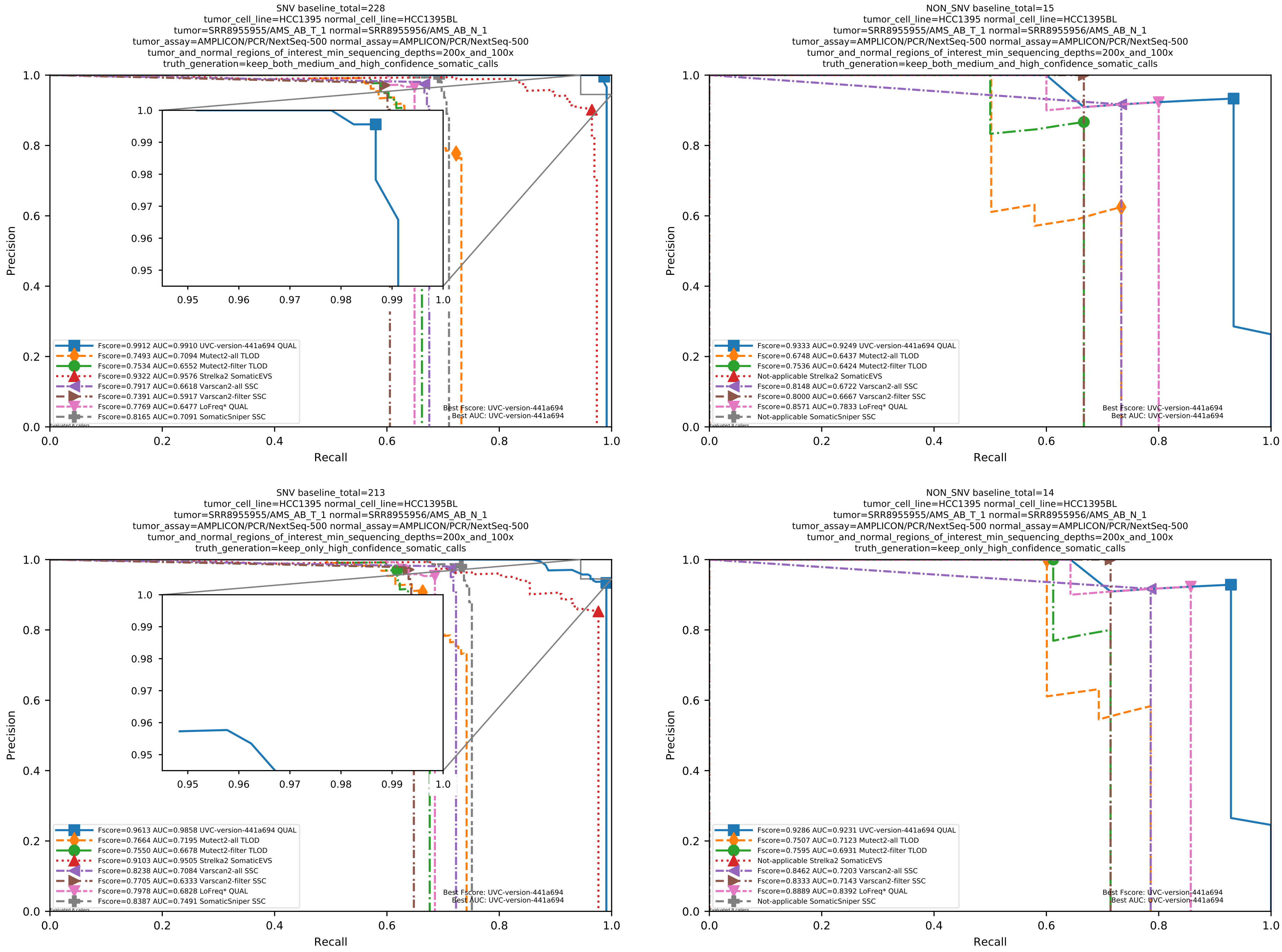

Supplementary Figure 121: Performance of somatic variant callers on the SEQC2 somatic reference sets with Illumina sequencing data aligned with BWA MEM to the GRCh38 human-genome reference.

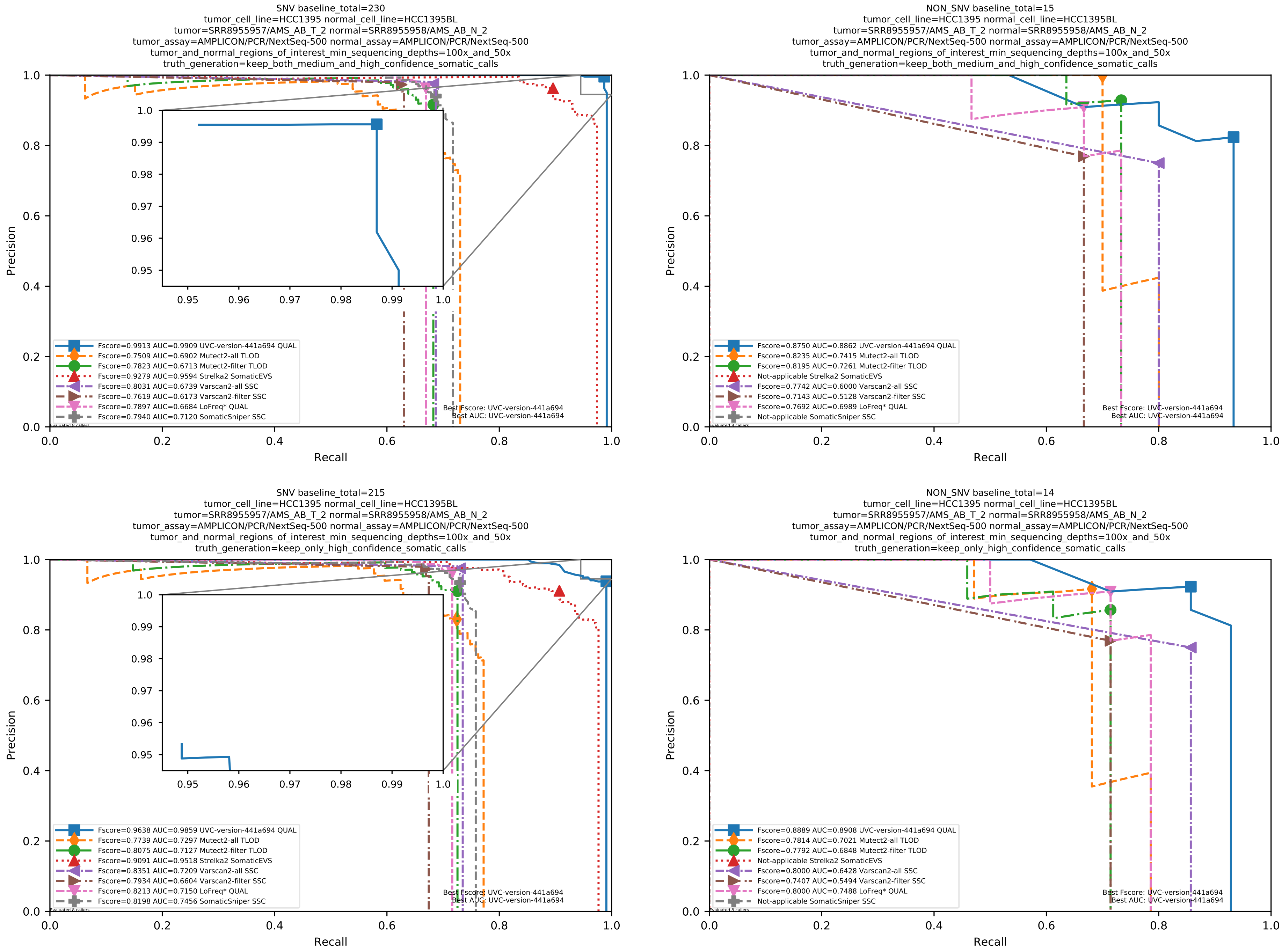

Supplementary Figure 122: Performance of somatic variant callers on the SEQC2 somatic reference sets with Illumina sequencing data aligned with BWA MEM to the GRCh38 human-genome reference.

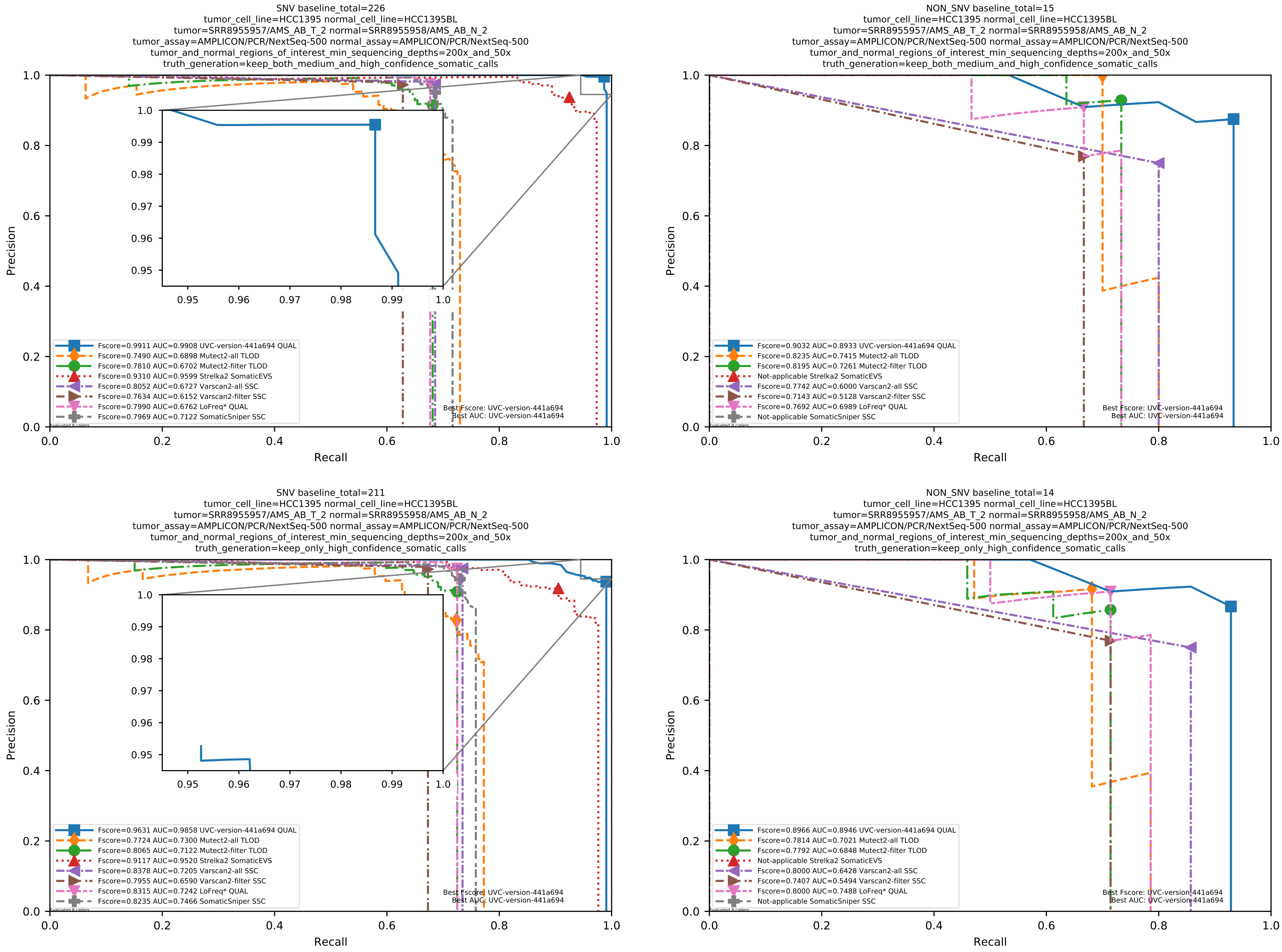

Supplementary Figure 123: Performance of somatic variant callers on the SEQC2 somatic reference sets with Illumina sequencing data aligned with BWA MEM to the GRCh38 human-genome reference.

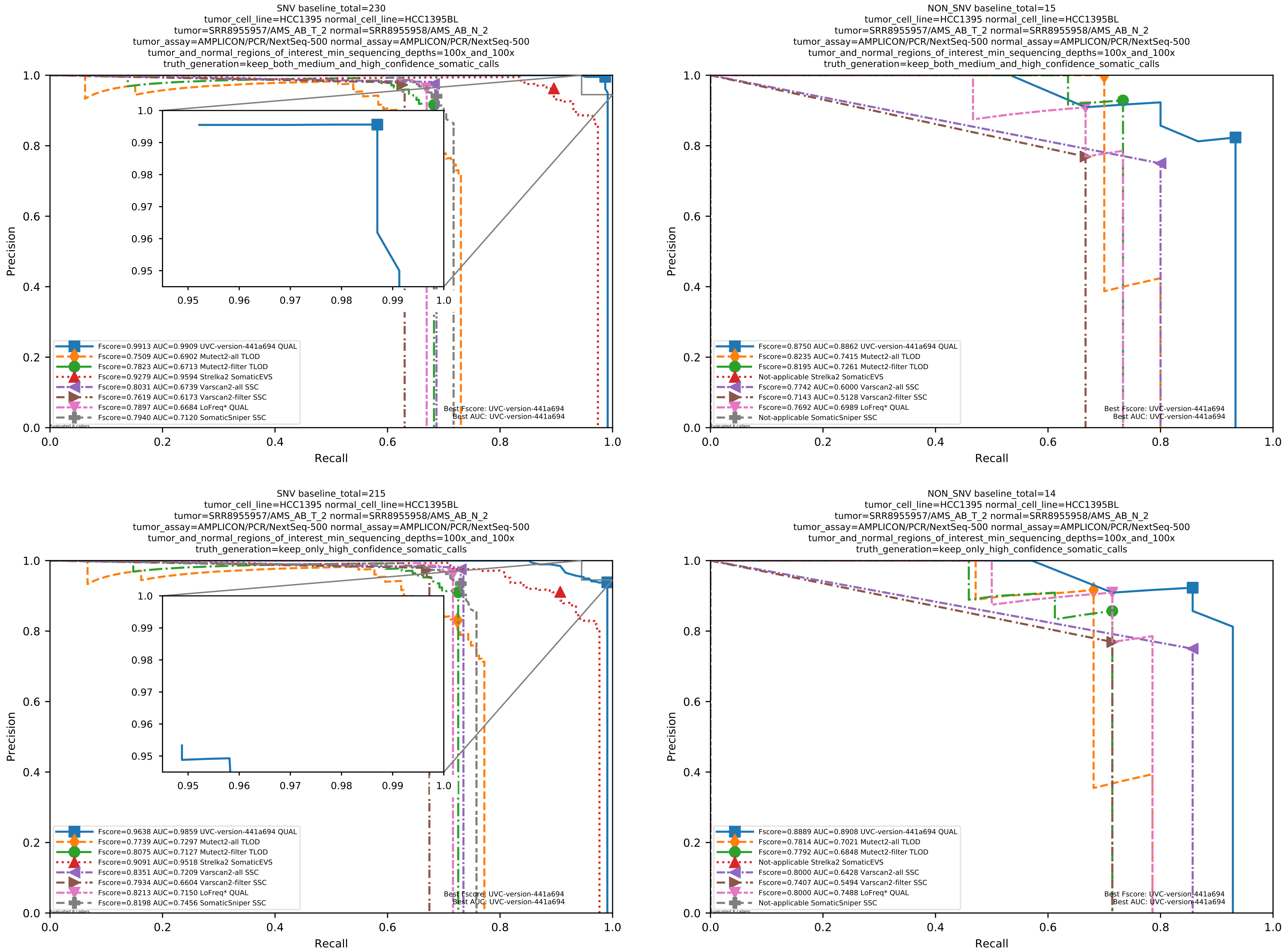

Supplementary Figure 124: Performance of somatic variant callers on the SEQC2 somatic reference sets with Illumina sequencing data aligned with BWA MEM to the GRCh38 human-genome reference.

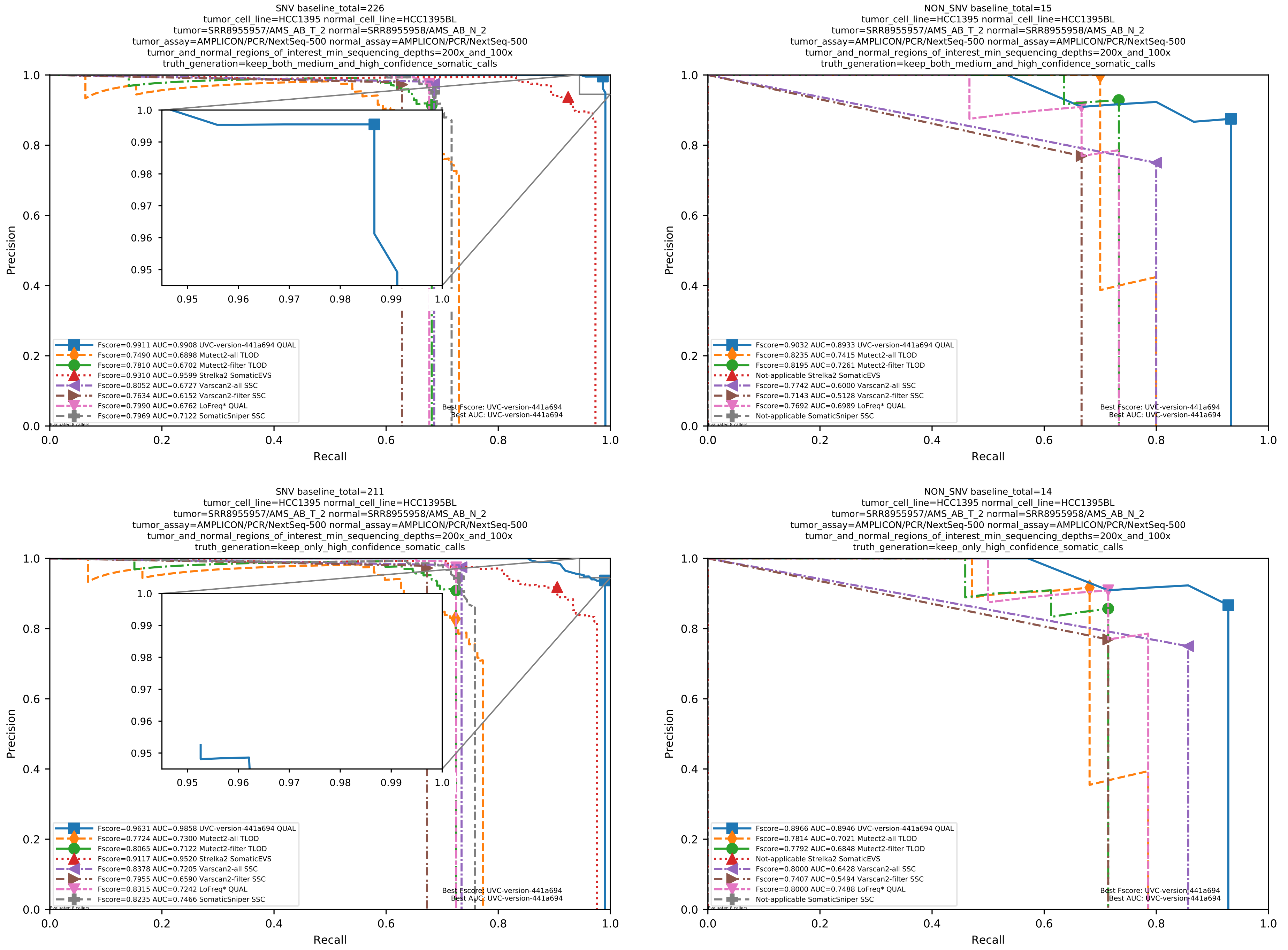

Supplementary Figure 125: Performance of somatic variant callers on the SEQC2 somatic reference sets with Illumina sequencing data aligned with BWA MEM to the GRCh38 human-genome reference.

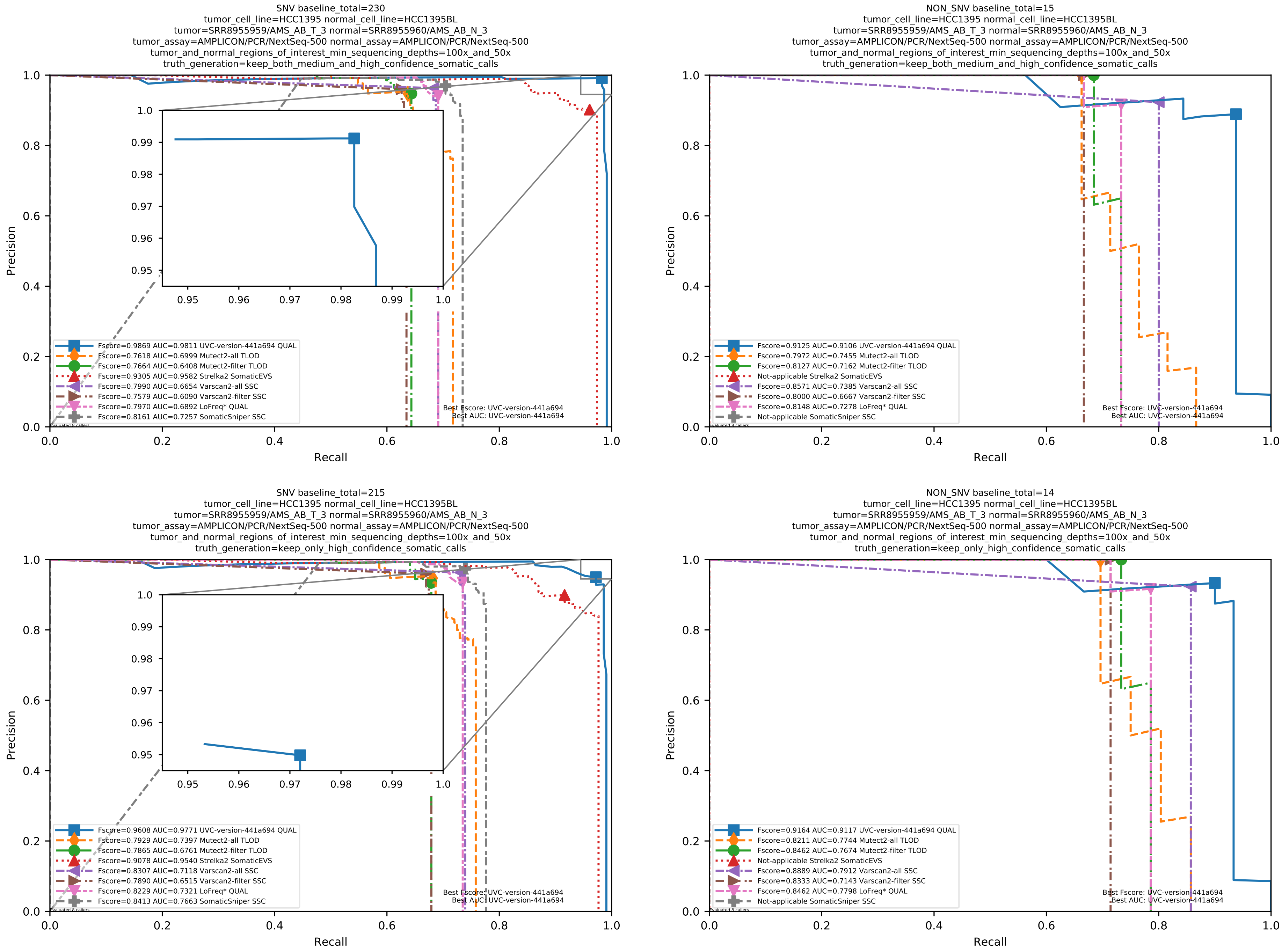

Supplementary Figure 126: Performance of somatic variant callers on the SEQC2 somatic reference sets with Illumina sequencing data aligned with BWA MEM to the GRCh38 human-genome reference.

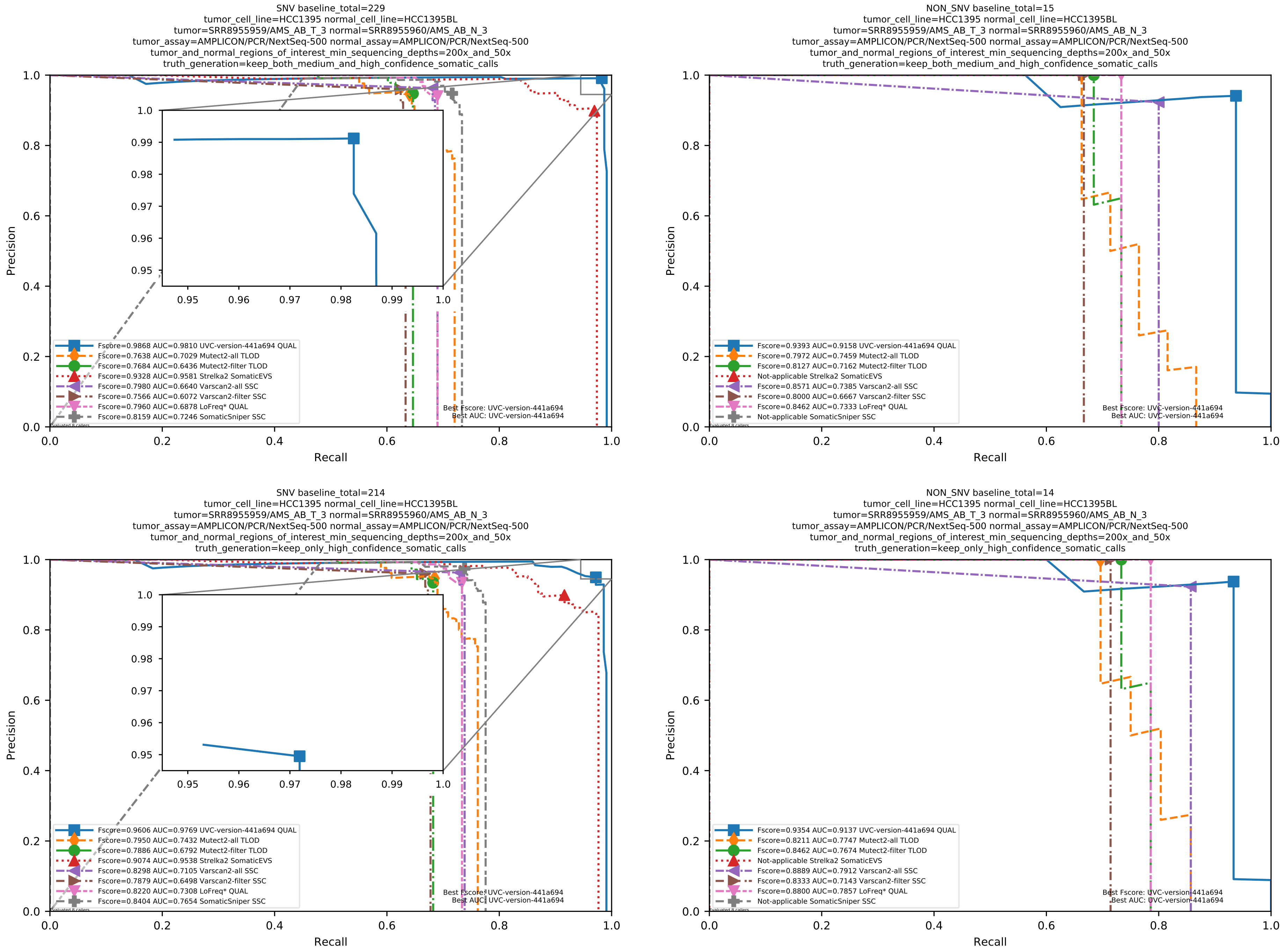

Supplementary Figure 127: Performance of somatic variant callers on the SEQC2 somatic reference sets with Illumina sequencing data aligned with BWA MEM to the GRCh38 human-genome reference.

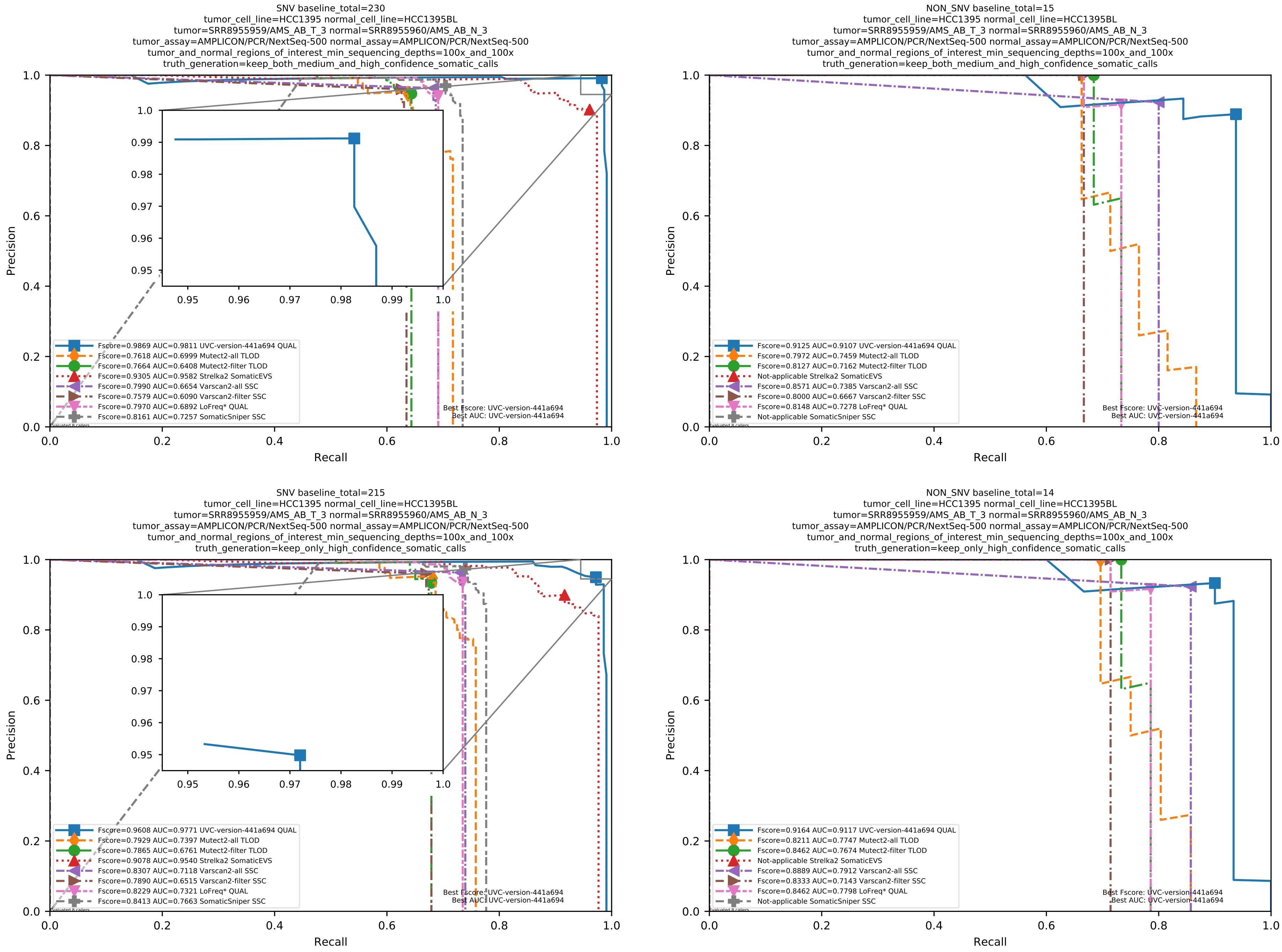

Supplementary Figure 128: Performance of somatic variant callers on the SEQC2 somatic reference sets with Illumina sequencing data aligned with BWA MEM to the GRCh38 human-genome reference.

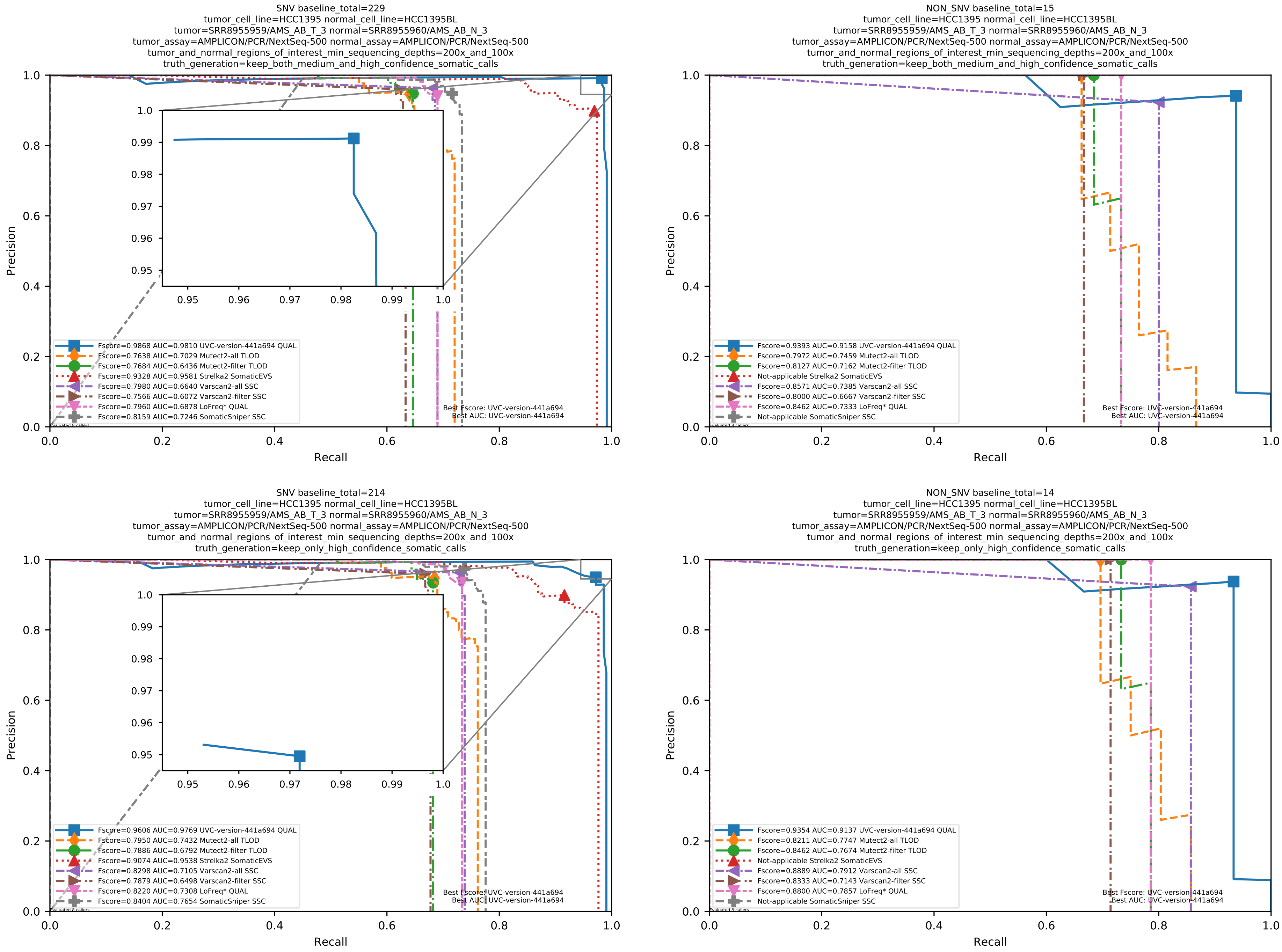

Supplementary Figure 129: Performance of somatic variant callers on the SEQC2 somatic reference sets with Illumina sequencing data aligned with BWA MEM to the GRCh38 human-genome reference.

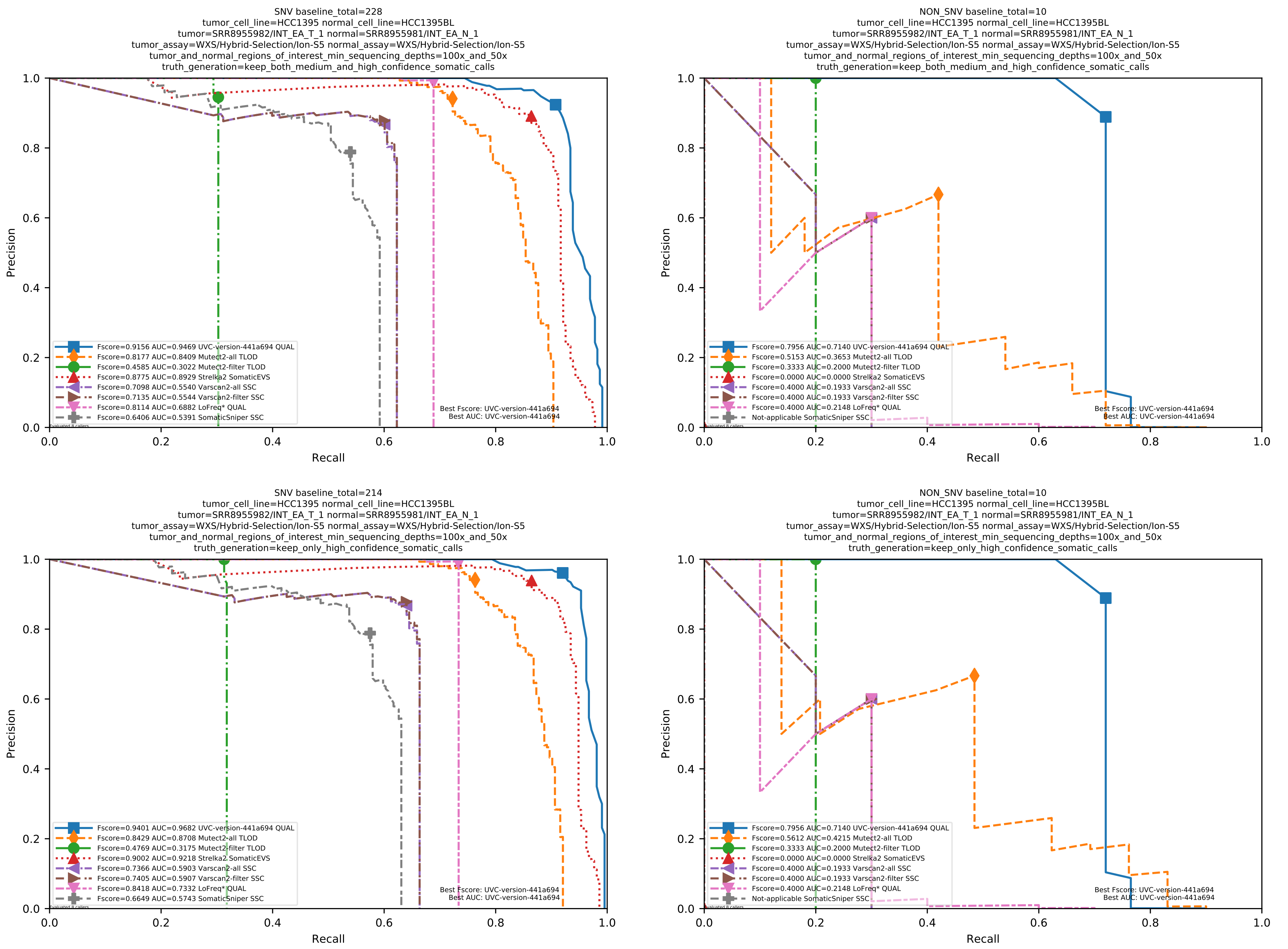

Supplementary Figure 130: Performance of somatic variant callers on the SEQC2 somatic reference sets with Illumina sequencing data aligned with BWA MEM to the GRCh38 human-genome reference.

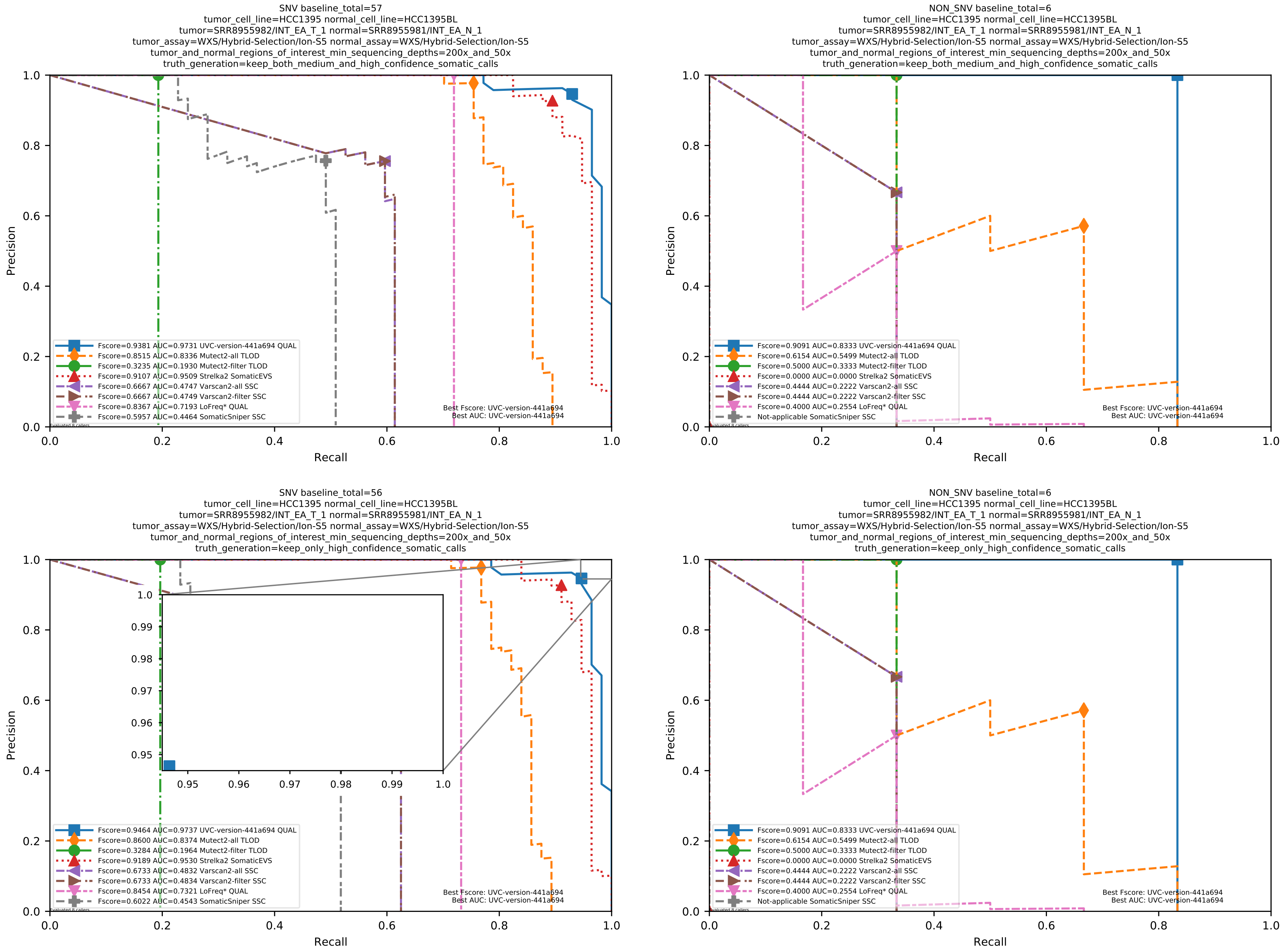

Supplementary Figure 131: Performance of somatic variant callers on the SEQC2 somatic reference sets with Illumina sequencing data aligned with BWA MEM to the GRCh38 human-genome reference.

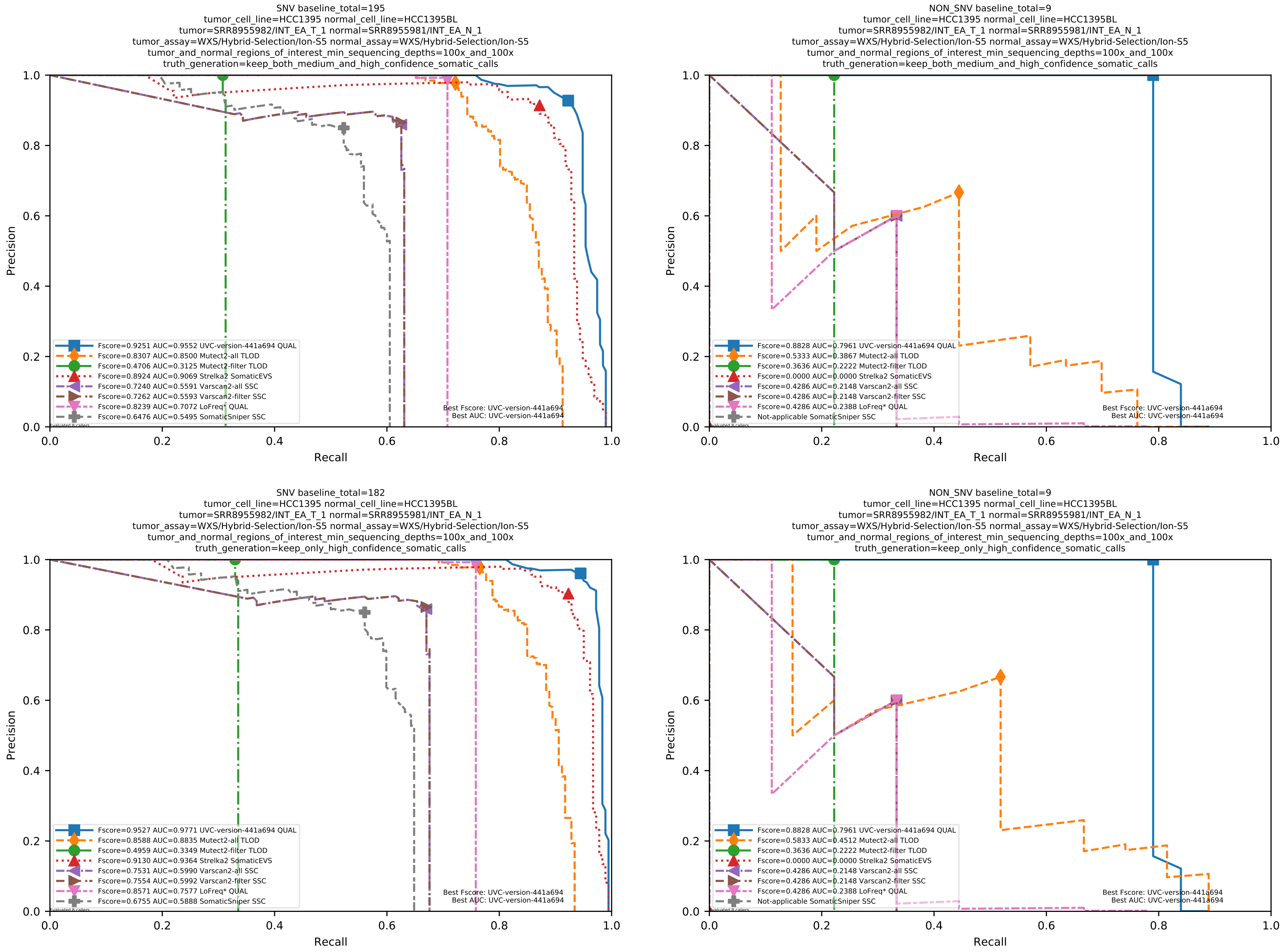

Supplementary Figure 132: Performance of somatic variant callers on the SEQC2 somatic reference sets with Illumina sequencing data aligned with BWA MEM to the GRCh38 human-genome reference.

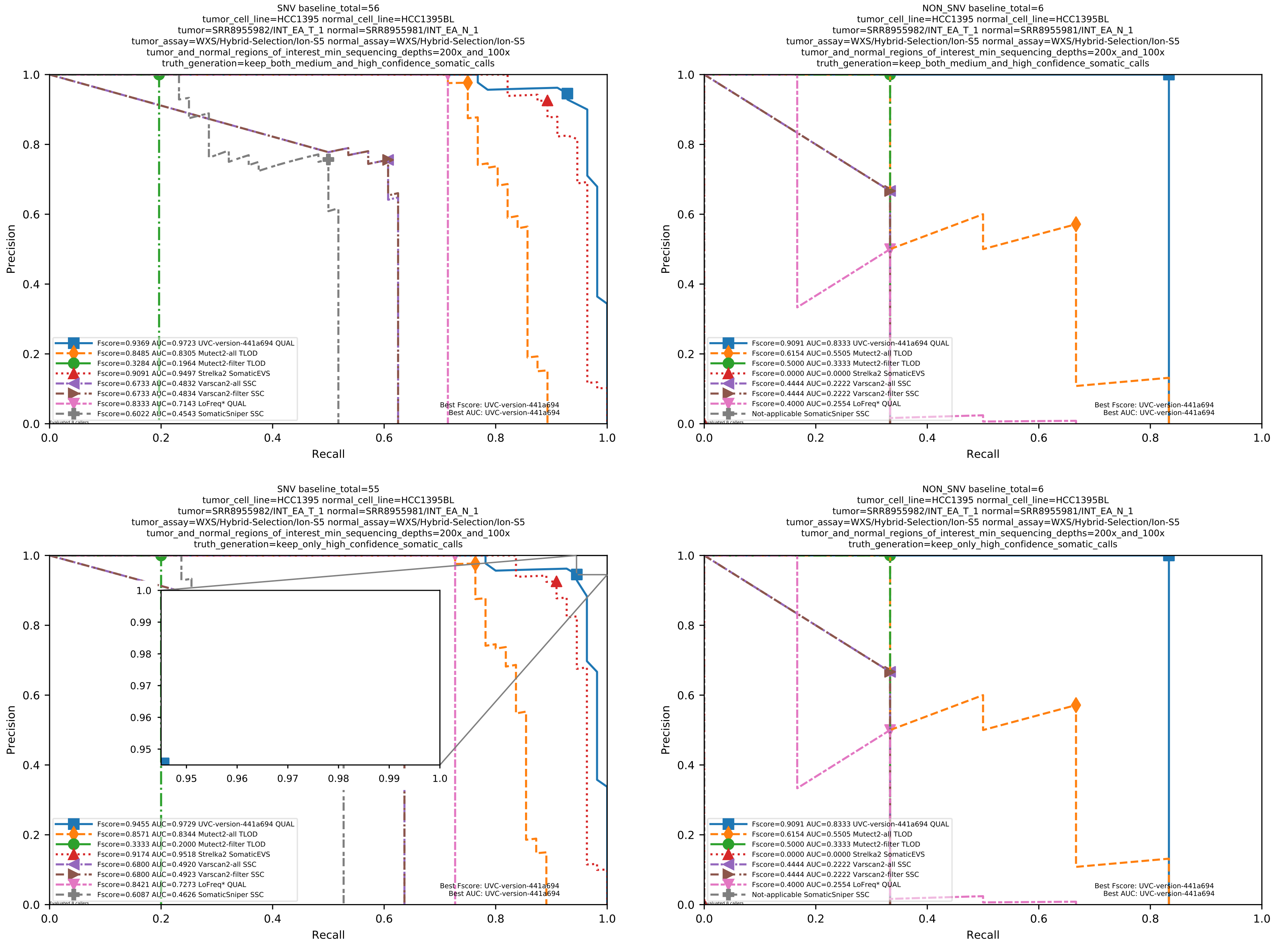

Supplementary Figure 133: Performance of somatic variant callers on the SEQC2 somatic reference sets with Illumina sequencing data aligned with BWA MEM to the GRCh38 human-genome reference.

#### 1.7 Supplemental evaluation results on amplicon-sequencing datasets for calling somatic variants from tumor-normal pairs, with samples from colon-cancer patients

The datasets used in this subsection were presented in more detail and manually reviewed by Dame et al.; Sandmann et al. [1;4].

#### 1.8 Supplemental evaluation results on amplicon-sequencing datasets with UMI for calling variants in tumor-only mode

Mageri version 2.0 and smCounter2 version 2 commit c475863b were used to generate Supplementary Table 1 and Supplementary Fig. 134.

The vcfeval utility from the RTG tools is unable to handle any tri-allelic site properly. Therefore, we wrote a custom script to generate Supplementary Table 1 instead of using vcfeval.

The sequencing data used to generate Supplementary Fig. 134 include PCR-primer sequences. Thus, to exclude false positive variants on primers, we passed the command-line parameter `--primerlen 23` to UVC to ignore read support within 23 base pairs of fragment ends. In Supplementary Fig. 134, two filters were applied to the variant calls generated by UVC. The SNV calls with an average of more than 2 mismatches per 100 bps were filtered out. The InDel calls with UMI-derived basecall qualities of less than 20 were filtered out.

| Variant caller | Number of false positive calls | | | | | | | |
|---|---|---|---|---|---|---|---|---|
| | 0 | 1 | 2 | 3 | 4 | 5 | 6 | 7 |
| UVC | 50 | 52 | 52 | 53 | 53 | 53 | 53 | 53 |
| Mageri | 47 | 47 | 48 | 48 | 48 | 48 | 48 | 49 |

Supplementary Table 1: Number of variants called versus number of false positive calls. This table shows the performance of UVC and Mageri on the HD734 reference standard with Illumina sequencing data aligned with BWA-MEM to the human reference genome UCSC hg19. These data were previously used to validate the trained error model used by Mageri[6].

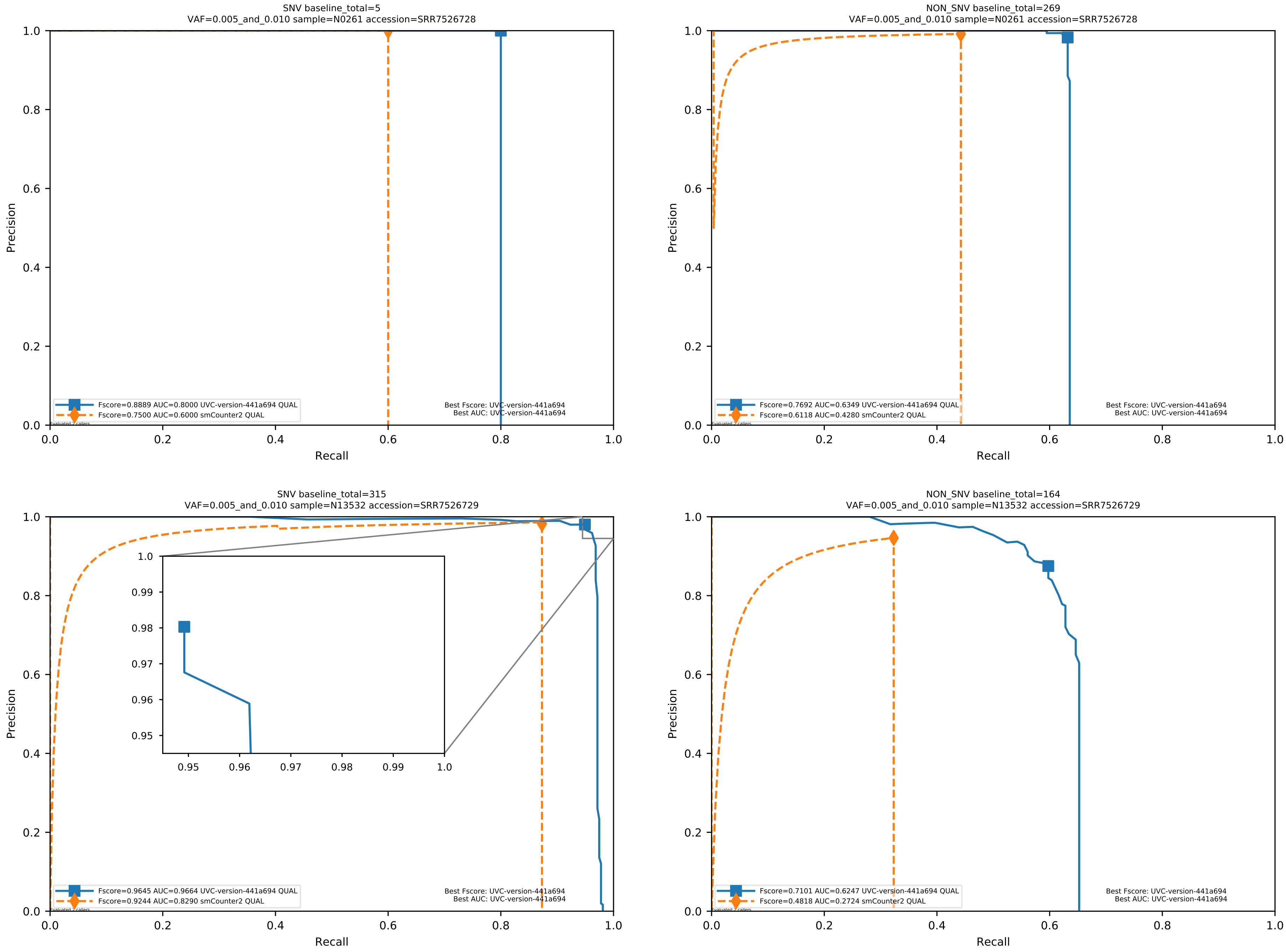

Supplementary Figure 134: Performance of UVC and smCounter2 on the HG001/HG002 physical mixtures with Illumina sequencing data aligned with BWA-MEM to the human reference genome UCSC hg19. These data were previously used to validate the trained error model used by smCounter2[7].

#### 1.9 Supplemental reanalysis results on an amplicon-sequencing dataset with UMI for calling variants in tumor-normal pairs, which provides additional insight about DNA damage repair

For the re-analysis of this SiMSen-Seq dataset (accession: SRP158874)[3], the QUAL threshold was set to 40. Initially, UVC only discovered the *DPH3* -8 variant on the two samples from patients with Cockayne Syndrome, because the *DPH3* -8 variant is characterized by strong signal in these two samples. Then, we manually discovered that the *DPH3* -8 variant is characterized by some weak signal in the two samples from patients with Xeroderma Pigmentosum.

The unique molecular identifier (UMI, which is also known as molecular barcode) family depth of every variant is required to generate Supplementary Fig. 135. Thus, the command-line parameter `-q 0` is passed to UVC to report every variant.

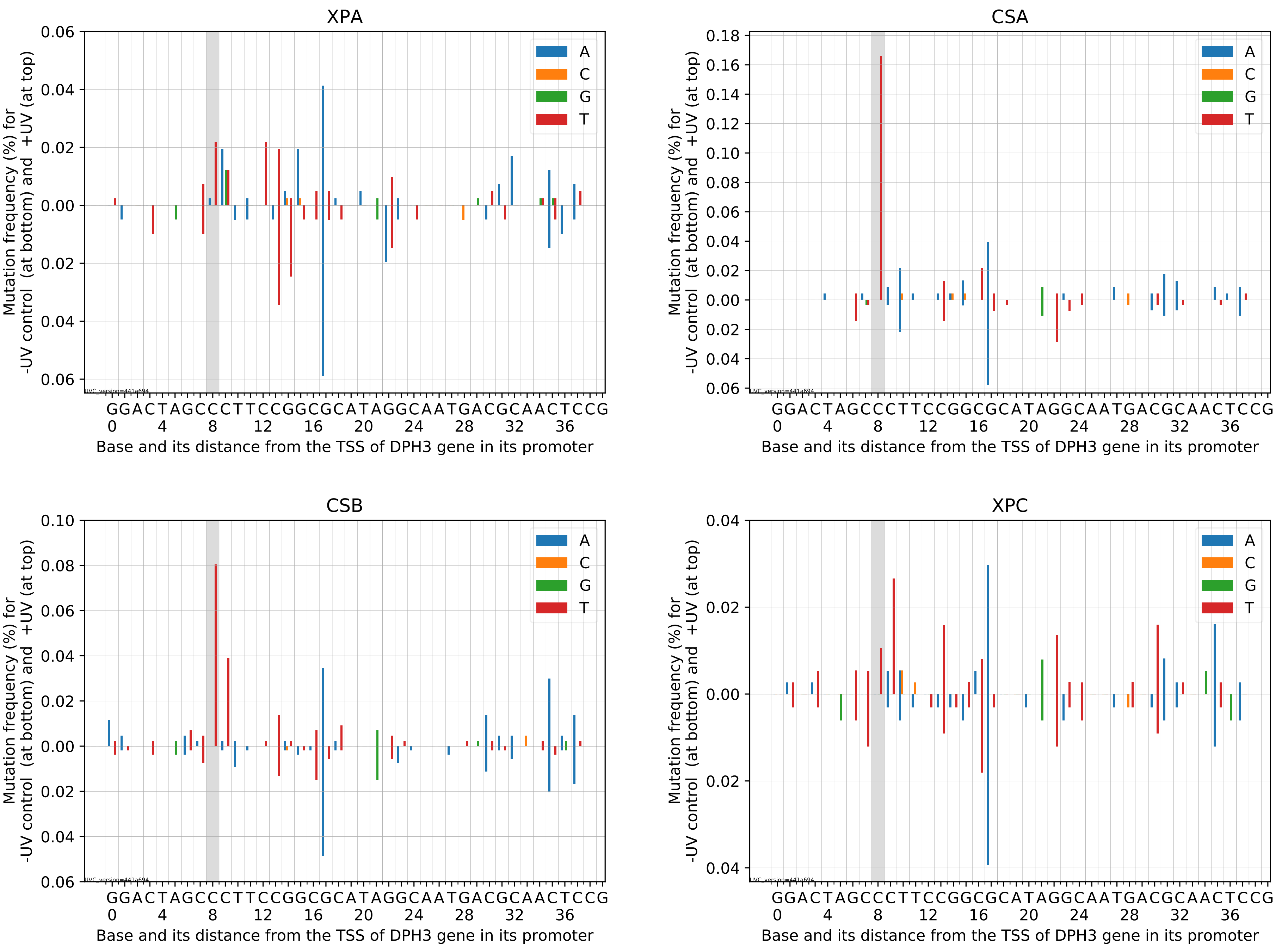

Supplementary Figure 135: Variant allele UMI depths near the `C>T` substitution at the *DPH3* -8 bp `TTCCG` hotspot sites. TSS denotes transcription start site. This figure complements Supplementary Table 2.

| Sample | *RPLA* -116 bp `C>T` | | *DPH3* -8 bp `C>T` | |
|---|---|---|---|---|
| | with UV | without UV | with UV | without UV |
| XPA | 19 / 24928 | 0 / 12670 | 9 / 41271 | 0 / 20422 |
| CSA | 82 / 15164 | 0 / 20286 | 38 / 22887 | 0 / 27881 |
| CSB | 16 / 26007 | 0 / 23822 | 35 / 43512 | 0 / 53855 |
| XPC | 35 / 20047 | 0 / 19003 | 4 / 37702 | 0 / 33215 |

Supplementary Table 2: Variant allele fractions of the `C>T` substitution at the *RPLA* -116 bp and *DPH3* -8 bp `TTCCG` hotspot sites. This table complements Supplementary Fig. 135.

#### 1.10 Computational resources

As a reminder, we tried to run the latest release of each software, with an allocation of 32 logical CPU cores, on a Red Hat 4.8.5-28 server with 128 GB of RAM and 28/56 physical/logical CPU cores (Intel(R) Xeon(R) CPU E5-2680 v4 @ 2.40GHz).

| Software | implementation language | running time (seconds) | | maximum memory (kbytes) |
|---|---|---|---|---|
| | | wall-clock | CPU | |
| BWA MEM | C | 84 512 | 2 675 778 | 41 666 720 |
| UVC | C++ | 32 745 | 831 865 | 84 725 552 |
| GATK4 MarkDuplicates | Java | 47 187 | 263 039 | 4 288 676 |
| GATK4 BaseRecalibrator | Java | 78 404 | 85 764 | 3 310 879 |
| GATK4 ApplyBQSR | Java | 65 742 | 100 598 | 1 494 745 |
| GATK4 Mutect2 | Java | 350 678 | 632 879 | 20 895 232 |
| Strelka2 | C++ | 10 018 | 193 735 | 58 846 704 |
| VarScan2 | Java | 174 896 | 156 086 | 25 340 080 |
| Dindel from LoFreq | C++ | 65 679 | 65 644 | 4 879 584 |
| LoFreq | C++ | 152 619 | 1 532 972 | 14 406 592 |
| SomaticSniper | C++ | 34 429 | 34 410 | 4 879 584 |

Supplementary Table 3: The computational resources consumed by the variant callers listed in Supplementary Fig. 105, as well as the resources consumed by the software packages that these variant callers depend on, on the Illumina-sequenced 30%/70% physical mixture used in Supplementary Fig. 105 in Section 1.5. The resources consumed by these variant callers on other datasets follow the same trend and are therefore not shown. UVC runs at least five times faster than Mageri, smCounter2, and Debarcer (data not shown), which are three variant callers specially designed for data with unique molecular identifiers (UMIs). As shown in this table, the running time of UVC is negligible compared with the commonly used and very fast NGS aligner BWA MEM. Furthermore, the memory consumed by UVC is designed to be approximately 2.5 GB per CPU core. Hence, the usage of UVC is feasible in all settings.
