## Supplementary file 1 for "Calling small variants with universality and Bayesian-frequentist hybridism": supplementar-file-1--861_TandN_R1R2_trimPaired_22_41574679.html


| 22:41574679 | all:2720 A:0 a:1 T:0 t:4 C:2322 c:253 G:2 g:1 Ins:0 ins:0 del:137 n:0 match:2582 mismatch:145 softclip:21 |
| --- | --- |
| Tumor\_Reference | CCTGTTCAGCCCAACCCCATGAGCCCCCAGCAGCATATGCTCCCAAATCAGGCCCAGTCCCCACACCTACAAGGCCAGCAGATCCCTAATTCTCTCTCCAATCAAGTGCGCTCTCCCCAGCCTGTCCCTTCTCCACGGCCACAGTCCCAG\|C\|CCCCCCACTCCAGTCCTTCCCCAAGGATGCAGCCTCAGCCTTCTCCACACCACGTTTCCCCACAGACAAGTTCCCCACATCCTGGACTGGTAGCTGCCCAGGCCAACCCCATGGAACAAGGGCATTTTGCCAGCCCGGACCAGAATTCAA |
| 1:1108:5970:34128#2#R | \| \| |
| MQ=60 isize=-125 | TTTGGCCCAGTCCCCACACCTACAAGgCCAGCAGATCCCTAATTCTCTCTCCAATCAAGTGCGCtCTCCCCAGCCTGTCCCTTCTCCACGGCCACAGTCCCAG\|\_\|CCCCCCACTCCAGTCCTTCCCCAAGanGACTCCAATGTCACGGCACCG |
| 2:2203:17792:35915#2#R | \| \| |
| MQ=60 isize=-121 | gCGTTTTGGCCCAGTCCCCACACCTACAAGGCCAGCAGATCCCTAATTCTCTCTCCAATCAAGTGCGCTCTCCCCAGCCTGTCCCTTCTCCACGGCCACAGTCCCAG\|\_\|CCCCCCACTCCAGTCCTTCCCanGACTCCAATCGTCACCACACT |
| 1:1103:9587:7568#1#F | \| \| |
| MQ=60 isize=166 | GCCCAGTCCCCACACCTaCAaggccAgCAGATcccTAATTCtCTCTcCAATcAAGTGcgCTCTCCCCAGCCTGTCCCTTCtCCACGGCCACAGTCCcaG\|\_\|CCCCCCACTCCagTCCTtccCCAaGGATGCagcCTCAGCCTTctCCACACCa |
| 1:1106:12616:4344#1#F | \| \| |
| MQ=60 isize=177 | GCCCAGTCCCCACACCTACAAGGCCAGCAGATCCCTAATTCTCTCTCCAATCAAGTGCGCTCTCCCCAGCCTGTCCCTTCTCCACGGCCACAGTCCCAG\|\_\|CCCCCCACTCCAGTCCTTCCCCAAGGATGCAGCCTCAGCCTTCTCCACACCa |
| 1:1107:9272:32667#1#F | \| \| |
| MQ=60 isize=268 | GCCCAGTCCCCACACCTACAAGGCCAGCAGATCCCTAATTCTCTCTCCAATCAAGTGCGCTCTCCCCAGCCTGTCCCTTCTCCACGGCCACAGTCCCAG\|\_\|CCCCCCACTCCAGTCCTTCCCCAAGGATGCAGCCTCAGCCTTCTCCACACCa |
| 1:1108:5970:34128#1#F | \| \| |
| MQ=60 isize=125 | GCCCAGTCCCCACACCTACAAGGCCAGCAGATCCCTAATTCTCTCTCCAATCAAGTGCGCTCTCCCCAGCCTGTCCCTTCTCCACGGCCACAGTCCCAG\|\_\|CCCCCCACTCCAGTCCTTCCCCAAGAGGACTCCAATGTCACGGCACCGCTGT |
| 1:1109:18066:5993#1#F | \| \| |
| MQ=60 isize=293 | GCCCAGTCCCCACACCTACAAGGCCAGCAGATCCCTAATTCTCTCTCCAATCAAGTGCGCTCTCCCCAGCCTGTCCCTTCTCCACGGCCACAGTCCCAG\|\_\|CCCCCCACTCCAGTCCTTCCCCAAGGatGCAGCCTCAGCCTTCTCCACACCA |
| 1:1109:5015:21811#1#F | \| \| |
| MQ=60 isize=211 | GCCCAGTCCCCACACCTACAAGGCCAGCAGATCCCTAATTCTCTCTCCAATCAAGTGCGCTCTCCCCAGCCTGTCCCTTCTCCACGGCCACAGTCCCAG\|\_\|CCCCCCACTCCAGTCCTTCCCCAAGGATGCAGCCTCAGCCTTCTCCACACCa |
| 1:1110:19947:28603#1#F | \| \| |
| MQ=60 isize=364 | GCCCAGTCCCCACACCTACAAGGCCAGCAGATCCCTAATTCTCTCTCCAATCAAGTGCGCTCTCCCCAGCCTGTCCCTTCTCCACGGCCACAGTCCCAG\|\_\|CCCCCCACTCCAGTCCTTCCCCAAGGATGCAGCCTCAGCCTTCTCCACACCA |
| 1:1111:1291:6254#1#F | \| \| |
| MQ=60 isize=0 | gcCCAGTCCCCACACCTACAAGGCCAGCAGaTCCCTAATTCTCTCTCCAATCAAGTGCGCTCTCCCCAGCCTGTCCCTTCTCCACGGcCACAGtCCCAG\|\_\|CCcCCcACTCCAGTCCTTCCCCAAGGATGcAGCCTCAGCCTTCTCCACACCa |
| 1:1112:14098:72501#1#F | \| \| |
| MQ=60 isize=243 | GCCCAGTCCCCACACCTaCAAGGCCAGCAGATCCCTAATTCTCTCTCCAATCAAGTGCGCTCTCCCCAGCCTGTCCCTTCTCCACGGCCACAGTCCCAG\|\_\|CCCCCCACTCCAGTCCTTCCCCAAGGATGCAGCCTCAGCCTTCTCCACACCa |
| 1:1113:17323:6882#1#F | \| \| |
| MQ=60 isize=207 | GccCAGTCCCCACACCTACAAGGCCAGCAGATCCCTAATTCTCTCTCCAATCAAGTGCGCTCTCCCCAGCCTGTCCCTTCTCCACGGCCACAGTCCCAG\|\_\|CCCCCCACTCCAgTCCTTCCCCAAGGaTGCAGCCTCAGCCTTCTCCACACCA |
| 1:1114:17042:82311#1#F | \| \| |
| MQ=60 isize=195 | GCCCAGTCCCCACACCTACAAGGCCAGCAGATCCCTAATTCTCTCTCCAATCAAGTGCGCTCTCCCCAGCCTGTCCCTTCTCCACGGCCACAGTCCCAG\|\_\|CCCCCCACTCCAGTCCTTCCCCAAGGATGCAGCCTCAGCCTTCTCCACACCa |
| 1:1115:11463:21490#1#F | \| \| |
| MQ=60 isize=256 | GCCCAGTCCCCACACCTACAAGGCCAGCAGATCCCTAATTCTCTCTCCAATCAAGTGCGCTCTCCCCAGCCTGTCCCTTCTCCACGGCCACAGTCCCAG\|\_\|CCCCCCACTCCAGTCCTTCCCCAAGGATGCAGCCTCAGCCTTCTCCACACCA |
| 1:1115:15178:86872#1#F | \| \| |
| MQ=60 isize=177 | GCCCAGTCCCCACACCTACAAGGCCAGCAGATCCCTAATTCTCTCTCCAATCAAGTGCGCTCTCCCCAGCCTGTCCCTTCTCCACGGCCACAGTCCCAG\|\_\|CCCCCCACTCCAGTCCTTCCCCAAGGATGCAGcCTCAGCCTTCTCCACACCa |
| 1:1202:7953:91713#1#F | \| \| |
| MQ=60 isize=132 | GCCCAGTCCCCACACCTACAAGGCCAGCAGATCCCTAATTCTCTCTCCAATCAAGTGCGCTCTCCCCAGCCTGTCCCTTCTCCACGGCCACAGTCCCAG\|\_\|CCCCCCACTCCAGTCCTTCCCCAAGGATGCAGAGGACTCCAATCGACCCGCc |
| 1:1203:13275:10217#1#F | \| \| |
| MQ=60 isize=378 | GCCCAGTCCCCACACCTACAAGGCCAGCAGATCCCTAATTCTCTCTCCAATCAAGTGCGCTCTCCCCAGCCTGTCCCTTCTCCACGGCCACAGTCCCAG\|\_\|CCCCCCACTCCAGtCCTTCCCCAAGGATGCAGCCTCAgCCTTCTCCACACCc |
| 1:1203:8658:72666#1#F | \| \| |
| MQ=60 isize=268 | GCCCAGTCCCCACACCTACAAGGCCAGCAGATCCCTAATTCTCTCTCCAATCAAGTGCGCTCTCCCCAGCCTGTCCCTTCTCCACgGCCACAGTCCCAg\|\_\|cCcCCCACTCCaGtCCTTCCCCAAGGATGCAGCCTCAGCCTtcTCCACACCC |
| 1:1206:8900:4083#1#F | \| \| |
| MQ=60 isize=174 | GCCCAGTCCCCACACCTACAAGGCCAGCAGATCCCTAATTCTCTCTCCAATCAAGTGCGCTCTCCCCAGCCTGTCCCTTCTCCACGGCCACAGTCCCAG\|\_\|CCCCCCACTCCAGTCCTTCCCCAAGGATGCAGCCTCAGCCTTCTCCACACCc |
| 1:1212:17067:18737#1#F | \| \| |
| MQ=60 isize=212 | GCCCAGTCCCCACACCTACAAGGCCAGCAGATCCCTAATTCTCTCTCCAATCAAGTGCGCTCTCCCCAGCCTGTCCCTTCTCCACGGCCACAGTCCCAG\|\_\|CCCCCCACTCCAGTCCTTCCCCAAGGATGCAgCCTCAgCCTTCTCCACACCa |
| 1:1214:7829:27692#1#F | \| \| |
| MQ=60 isize=256 | GCCCAGTCCCCACACCTACAAGGCCAGCAGATCCCTAATTCTCTCTCCAATCAAGTGCGCTCTCCCCAGCCCGTCCCTTCTCCACGGCCACAGTCCCAG\|\_\|CCCCCCACTCCAGTCCTTCCCCAAGGATGCAGCCTCAGCCTTCTCCACACCa |
| 1:1214:10886:65209#1#F | \| \| |
| MQ=60 isize=207 | GCCCAGTCCCCACACCTACAAGGCCAGCAGATCCCTAATTCTCTCTCCAATCAAGTGCGCTCTCCCCAGCCTGTCCCTTCTCCACGGCCACAGTCCCAG\|\_\|CCCCCCACTCCAGTCCTTCCCCAAGGATGCAGCCTCAGCCTTCTCCACACCa |
| 1:1216:5228:47083#1#F | \| \| |
| MQ=60 isize=181 | GCCCAGTCCCCACACCTACAAGGCCAGCAGATCCCTAATTCTCTCTCCAATCAAGTGCGCTCTCCCCAGCCTGTCCCTTCTCCACGGCCACAGTCCCAG\|\_\|CCCCCCACTCCAGTCCTTCCCCAAGGATGCAGCCTCAGCCTTCTCCACACCa |
| 1:1216:18755:83682#1#F | \| \| |
| MQ=60 isize=207 | GCCCAGTCCCCACACCTACAAGGCCAGCAGATCCCTAATTCTCTCTCCAATCAAGTGCGCTCTCCCCAGCCTGTCCCTTCTCCACGGCCaCAgtCCCAG\|\_\|CCCCCCACTCCAGtCCTTCCCCAAGGATGCAGCCTCAGCCTtctCCaCACCa |
| 1:2102:17757:63250#1#F | \| \| |
| MQ=60 isize=322 | GCCCAGTCCCCACACCTACAAGGCCAGCAGATCCCTAATTCTCTCTCCAATCAAGTGCGCTCTCCCCAGCCTGTCCCTTCTCCACGGCCACAGTCCCAG\|\_\|CCCCCCACTCCAGTCCTTCCCCAAGGATGCAGCCTCAGCCTTCTCCACACCa |
| 1:2104:19213:87926#1#F | \| \| |
| MQ=60 isize=140 | GCCCAGTCCCCACACCTACAAGGCCAGCAGATCCCTAATTCTCTCTCCAATCAAGTGCGCTCTCCCCAGCCTGTCCCTTCTCCACGGCCACAGTCCCAG\|\_\|CCCCCCACTCCAGTCCTTCCCCAAGGATGCAGCCTCAGCCAGGACTCCAATC |
| 1:2106:15871:30572#1#F | \| \| |
| MQ=60 isize=232 | GCCCAGTCCCCACACCTACAAGGCCAGCAGATCCCTAATTCTCTCTCCAATCAAGTGCGCTCTCCCCAGCCTGTCCCTTCTCCACGGCCACAGTCCCAG\|\_\|CCCCCCACTCCAGTCCTTCCCCAAGGATGCAGCCTCAGCCTTCTCCACACCA |
| 1:2109:16208:44268#1#F | \| \| |
| MQ=60 isize=199 | GCCCAGTCCCCACACCTACAAGGCCAGCAGATCCCTAATTCTCTCTCCAATCAAGTGCGCTCTCCCCAGCCTGTCCCTTCTCCACGGCCACAGtCCCAG\|\_\|CCCCCCACTCCAGTCCTTCCCCAAGGATGCAGCCTCAGCCTTCTCCACACCA |
| 1:2111:20764:71366#1#F | \| \| |
| MQ=60 isize=160 | GCCCAGTCCCCACACCTACAAGGCCAGCAGATCCCTAATTCTCTCTCCAATCAAGTGCGCTCTCCCCAGCCTGTCCCTTCTCCACGGCCACAGTCCCAG\|\_\|CCCCCCACTCCAGTCCTTCCCCAAGGATGCAGCCTCAGCCTTCTCCACACCA |
| 1:2111:18577:90395#1#F | \| \| |
| MQ=60 isize=460 | GCCCAGTCCCCACACCTACAAGGCCAGCAGATCCCTAATTCTCTCTCCAATCAAGTGCGCTCTCCCCAGCCTGTCCCTTCTCCACGGCCACAGTCCCAG\|\_\|CCCCCCACTCCAGtCCTTCCCCAAGGATGCAGCCTCAGCCTTCTCCACACCA |
| 1:2113:11275:54733#1#F | \| \| |
| MQ=60 isize=421 | GCCCAGTCCCCACACCTACAAGGCCAGCAGATCCCTAATTCTCTCTCCAATCAAGTGCGCTCTCCCCAGCCTGTCCCTTCTCCACGGCCACAGTCCCAG\|\_\|CCCCCCACTCCAGtCCTTCCCCAAGGATGCAGCCTCAGCCTTCTCCACACCA |
| 1:2113:16409:97435#1#F | \| \| |
| MQ=60 isize=240 | GCCCAGTCCCCACACCTACAAGGCCAGCAGATCCCTAATTCTCTCTCCAATCAAGTGCGCTCTCCCCAGCCTGTCCCTTCTCCACGGCCACAGTCCCAG\|\_\|CCCCCCACTCCAGTTCTTCCCCAAGGATGCAGCCTCAGCCTTCTCCACACCA |
| 1:2114:13171:5774#1#F | \| \| |
| MQ=60 isize=229 | GCCCAGTCCCCACACCTACAAGGCCAGCAGATCCCTAATTCTCTCTCCAATCAAGTGCGCTCTCCCCAGCCTGTCCCTTCTCCACGGCCACAGTCCCAG\|\_\|CCCCCCACTCCAGTCCTTCCCCAAGGATGCAGCCTCAGCCTTCTCCACACCa |
| 1:2115:2799:81460#1#F | \| \| |
| MQ=60 isize=166 | GCCCAGTCCCCACACCTACAAGGCCAGCAGATCCCTAATTCTCTCTCCAATCAAGTGCGCTCTCCCCAGCCTGTCCCTTCTCCACGGCCACAGtCCCAG\|\_\|CCCCCCACTCCAGTCCTTCCCCAAGGATGCAGCCTCAGCCTTCTCCACACCA |
| 1:2116:18225:69787#1#F | \| \| |
| MQ=60 isize=256 | GCCCAGTCCCCACACCTACAAGGCCAGCAGATCCCTAATTCTCTCTCCAATCAAGTGCGCTCTCCCCAGCCTGTCCCTTCTCCACGGCCACAGTCCCAG\|\_\|CCCCCCACTCCAGTCCTTCCCCAAGGATGCAGCCTCAGCCTTCTCCACACCA |
| 1:2204:3514:6197#1#F | \| \| |
| MQ=60 isize=162 | GCCCAGTCCCCACACCTACAAGGCCAGCAGATCCCTAATTCTCTCTCCAATCAAGTGCGCTCTCCCCGGCCTGTCCCTTCTCCACGGCCACAGTCCCAG\|\_\|CCCCCCACTCCAGTCCTTCCCCAAGGATGCAGCCTCAGCCTTCTCCACACCc |
| 1:2204:14133:63410#1#F | \| \| |
| MQ=60 isize=240 | GCCCAGTCCCCACACCTACAAGGCCAGCAGATCCCTAATTCTCTCTCCAATCAAGTGCGCTCTCCCCAGCCTGTCCCTTCTCCcCGGCCACAGTCCCaG\|\_\|CCCCCCaCTCCAGTCCTTCCCCAaGGaTGCAGCCtCAGCCTTCTCCACcCCC |
| 1:2205:8826:3969#1#F | \| \| |
| MQ=60 isize=158 | GCCCAGTCCCCACACCTACAAGGCCAGCAGATCCCTAATTCTCTCTCCAATCAAGTGCGCTCTCCCCAGCCTGTCCCTTCTCCACGGCCACAGTCCCAG\|\_\|CCCCCCACTCCAGTCCTTCCCCAaGGATGCAGCCTCAGCCTTCTCCACACCc |
| 1:2205:4755:25366#1#F | \| \| |
| MQ=60 isize=235 | GCCCAGTCCCCACACCTACAAGGCCAGCAGATCCCTAATTCTCTCTCCAATCAAGTGCGCTCTCCCCAGCCTGTCCCTTCTCCACGGCCACAGTCCCAG\|\_\|CCCCCCACTCCAGTCCTTCCCCAAGGATGCagCCTCagcCTTCTCCacacca |
| 1:2205:3991:78790#1#F | \| \| |
| MQ=60 isize=268 | GCCCAGTCCCCAC\_CCTACAAGGCCAGCAGATCCCTAATTCTCTCTCCAATCAAGTGCGCTCTCCCCAGCCTGTCCCTTCTCCACGGCCACAGTCCCAG\|\_\|CCCCCCACTCCAGTCCTTCCCCAAGGATGCAGCCTCAGCCTTCTCCACACCAC |
| 1:2207:11364:61886#1#F | \| \| |
| MQ=60 isize=208 | GCCCAGTCCCCACACCTACAAGGCCAGCAGATCCCTAATTCTCTCTCCAATCAAGTGCGCTCTCCCCAGCCTGTCCCTTCTCCACGGCCACAGTCCCAG\|\_\|CCCCCCaCTCCAGTCCTTCCCCAAGGATGCAGCCTCAGCCTTCTCCACACCa |
| 1:2210:11748:72868#1#F | \| \| |
| MQ=60 isize=0 | GCCCAGTCCCCACACCTACAAGGCCAGCAGATCCCTAATTCTCTCTCCAATCAAGTGCGCTCTCCCCAGCCTGTCCCTTCTCCACGGCCACAGTCCCAG\|\_\|CCCCCCACTCCAGTCCTTCCCCAAGGATGCAGCCTCAGCCTTCTCCACACCc |
| 1:2212:16753:61055#1#F | \| \| |
| MQ=60 isize=132 | GCCCAGTCCCCACATCTACAAGGCCAGCAGATCCCTAATTCTCTCTCCAATCAAGTGCGCTCTCCCCAGCCTGTCCCTTCTCCACGGCCACAGTCCCAg\|\_\|CCCCCCACTCCAGtcCTTCCCCAAGGAtgcAGGACtCCAattcGCCCCACca |
| 1:2212:16012:63327#1#F | \| \| |
| MQ=60 isize=160 | GCCCAGTCCCCACACCTACAAGGCCAGCAGATCCCTAATTCTCTCTCCAATCAAGTGCGCTCTCCCCAGCCTGTCCCTTCTCCACGGCCACAGTCCCAG\|\_\|CCCCCCACTCCAGTCCTTCCCCAAGGATGCAGCCtCAgCCTTCTCCACACca |
| 1:2214:3697:11043#1#F | \| \| |
| MQ=60 isize=173 | GCCCAGTCCCCACACCTACAAGGCCAGCAGATCCCTAATTCTCTCTCCAATCAAGTGCGCTCTCCCCAGCCTGTCCCTTCTCCACGGCCACAGTCCCAG\|\_\|CCCCCCACTCCAGTCCTTCCCCAAGGATGCAGCCTCAGCCTTCTCCACACCa |
| 1:2216:3741:70157#1#F | \| \| |
| MQ=60 isize=251 | GCCCAGTCCCCACACCTACAAGGCCAGCAGATCCCTAATTcTCTCTCCAATCAAGTGCGCTCTCCCCAGCCtGTCCCTTCTCCACGGCCACAGtCCCAG\|\_\|CCCCCCACTCCAGTCCTTCCCCAaggATGCAGCCTCAGCCTTCTCCACACCA |
| 2:1104:20449:17134#1#F | \| \| |
| MQ=60 isize=166 | GCCCAGTCCCCACACCTACAAGGCCAGCAGATCCCTAATTCTCTCTCCAATCAAGTGCGCTCTCCCCAGCCTGTCCCTTCTCCACGGCCACAGTCCCAG\|\_\|CCCCCCACTCCAGTCCTTCCCCAAGGATGCAGCCTCAGCCTTCTCCACACCa |
| 2:1104:15053:43686#1#F | \| \| |
| MQ=60 isize=133 | GCCCAGTCCCCACGCCTACAAGGCCAGCAGATCCCTAATTCTCTCTCCAATCAAGTGCGCTCTCCCCAGCCTGTCCCTTCTCCACGGCCACAGTCCCAG\|\_\|CCCCCCACTCCAGTCCTTCCCCAAGGATGCAGCAGGACTCCAATCCGCCCTa |
| 2:1104:11531:71184#1#F | \| \| |
| MQ=60 isize=207 | GCCCAGTCCCCACACCTACAAGGCCAGCAGATCCCTAATTCTCTCTCCAATCAAGTGCGCTCTCCCCAGCCTGTCCCTTCTCCACGGCCACAgTCCCAG\|\_\|CCCCCCACTCCAGTCCTTCCCCAAGGATGCAGCCtCAGCCTtcTCCACACCa |
| 2:1106:9961:86872#1#F | \| \| |
| MQ=60 isize=173 A=6 | GCCCAGTCCCCACACCTACAAGGCCAGCAGATCCCTAATTCTCTCTCCAATCAAGTGCGCTCTCCCcAGCCtGTCCCTTCTCCACGgCCACAGtCCCAc\|a\|cccCCCACTCCAgtcCTTCCCCAaggATgCAGCCTCAgccTTCTCCACACC |
| 2:1108:13131:61481#1#F | \| \| |
| MQ=60 isize=174 | GCCCAGTCCCCACACCTACAAGGCCAGCAGATCCCTAATTCTCTCTCCAATCAAGTGCGCTCTCCCCAGCCTGTCCCTTCTCCACGGCCACAGTCCCAG\|\_\|CCCCCCACTCCAGTCCTTCCCCAAGGATGCAGCCTCAGCCTTCTCCACACCa |
| 2:1108:3148:96154#1#F | \| \| |
| MQ=60 isize=174 | gCCCAGTCCCCACACCTACAAGGCCAGCAGATCCCTAATTCTCTCTCCAaTCAAGTGCGCTCTCCCCAGCCTGTCCCTTCTCCACGGCcACAGTCCCag\|\_\|CCCCCaactcCAGTCCTTCCCCaaggatgcagcctcagccttctccacaccc |
| 2:1109:5303:16668#1#F | \| \| |
| MQ=60 isize=242 | GCCCAGTCCCCACACCTACAAGGCCAGCAGATCCCTAATTCTCTCTCCAATCAAGtGCGCTCTCCCCAGCCTGTCCCTTCTCCACGGCCACAgTCCCAG\|\_\|CCCCCCACtCCAGtCCTTCCCCAaGgatgcagCCccAGCCTTCTCCACaCCc |
| 2:1109:20569:81776#1#F | \| \| |
| MQ=60 isize=288 | GCCCAGTCCCCACACCTACAAGGCCAGCAGATCCCTAATTCTCTCTCCAATCAAGTGCGCTCTCCCCAGCCTGTCCCTTCTCCACGGCCACAGTCCCAG\|\_\|CCCCCCACTCCAGTCCTTCCCCAAGGATGCAGCCTCAGCCTTCTCCACACCa |
| 2:1111:19008:61770#1#F | \| \| |
| MQ=60 isize=294 | GCCCAGTCCCCACACCTACAAGGCCAGCAGATCCCTAATTCTCTCTCCAATCAAGTGCGCTCTCCCCAGcCTGTCCCTTCTCcACGgCCAcAgTCCCAG\|\_\|CCCCCCACTCCAGTCCTTCCCCAAGGATgCagCCTCAGCCTtCtCCacaccg |
| 2:1114:8060:48509#1#F | \| \| |
| MQ=60 isize=371 | GCCCAGTCCCCACACCTACAAGGCCAGCAGATCCCTAATTCTCTCTCCAATCAAGTGCGCTCTCCCCAGCCTGTCCCTTCTCCACGGCCACAGTCCCAG\|\_\|CCCCCCACTCCAGTCCTTCCCCAAgGATGCAGCCTCAGCCTTCTCCACACCa |
| 2:1115:18653:32394#1#F | \| \| |
| MQ=60 isize=0 | GCCCAGTCCCCACACCTACAAGGCCAGCAGATCCCTAATTCTCTCTCCAATCAAGTGCGCTCTCCCCAGCCTGTCCCTTCTCCACGGCCACAGTCCCAG\|\_\|CCCCCCACTCCAGTCCTTCCCCAAGGATGCAGCCTCAGCCTTCTCCACACCa |
| 2:1115:6794:86554#1#F | \| \| |
| MQ=60 isize=175 | GCCCAGTCCCCACACCTACAAGGCCAGCAGATCCCTAATTCTCTCTCCAATCAAGTGCGCTCTCCCCAGCCTGTCCCTTCTCCACGGCCACAGTCCCAG\|\_\|CCCCCCACTCCAGTCCTTCCCCAAGGATGCAGCCTCAGCCTTCTCCACACCa |
| 2:1208:19961:18461#1#F | \| \| |
| MQ=60 isize=414 | GCCCAGTCCCCACACCTACAAGGCCAGCAGATCCCTAATTCTCTCTCCAATCAAGTGCGCTCTCCCCAGCCTGTCCCTTCTCCACGGCCACAGTCCCAG\|\_\|CCCCCCACTCCAGTCCTTCCCCAAGGATGCAGCCTCAGCCTTCTCCACACCA |
| 2:1210:9113:40867#1#F | \| \| |
| MQ=60 isize=147 | GCCCAGTCCCCACACCTACAAGGCCAGCAGATCCCTAATTCTCTCTCCAATCAAGTGCGCTCTCCCCAGCCTGTCCCTTCTCCACGGCCACAGTCCCAG\|\_\|CCCCCCACTCCAGTCCTTCCCCAAGGATGCAGCCTCAGCCTTCTCCAGGACT |
| 2:1210:6045:76798#1#F | \| \| |
| MQ=60 isize=132 | GCCCAGTCCCCACACCTACAAGGCCAGCAGATCCCTAATTCTCTCTCCAATCAAGTGCGCTCTCCCCAGCCTGTCCCTTCTCCACGGCCACAGtCCCAG\|\_\|CCCCCCACTCCAGTCCTTCCCCAAGGATGCAGGACTCCAATTCGCCCCACca |
| 2:1210:14334:98366#1#F | \| \| |
| MQ=60 isize=256 | GCCCAGTCCCCACACCTACAAGGCCAGCAGATCCCTAATTCTCTCTCCAATCAAGTGCGCTCTCCCCAGCCTGTCCCTTCTCCACGGCCACAGTCCCAG\|\_\|CCCCCCACTCCAGTCCTTCCCCAAGGATGCAGCCTCAGCCTTCTCCACACCA |
| 2:1212:11395:86189#1#F | \| \| |
| MQ=60 isize=166 | GCCCAGTCCCCACACCTACAAGGCCAGCAGATCCCTAATTCTCTCTCCAATCAAGTGCGCTCTCCCCAGCCTGTCCCTTCTCCACGGCCACAGTCCCAG\|\_\|CCCCCCACTCCAGTCCTTCCCCAAGGATGCAGCCTCAGCCTTCTCCACACCa |
| 2:1213:4831:88470#1#F | \| \| |
| MQ=60 isize=143 | GCCCAGTCCCCACACCTACAAGGCCAGCAGATCCCTAATTCTCTCTCCAATCAAGTGCGCTCTCCCCAGCCTGTCCCTTCTCCACGGCCACAGTCCCAG\|\_\|CCCCCCACTCCAGTCCTTCCCCAAGGATGCAGCCTCAGCCTTCAGGACTCCA |
| 2:2105:16250:81208#1#F | \| \| |
| MQ=60 isize=147 | GCCCAGTCCCCACACCTACAAGGCCAGCAGATCCCTAATTCTCTCTCCAATCAAGTGCGCTCTCCCCAGCCTGTCCCTTCTCCACGGCCACAGTCCCAG\|\_\|CCCCCCACTCCAGTCCTTCCCCaaGGATGCAGcCTCAGCCTTCTCCAGGACt |
| 2:2108:17137:7384#1#F | \| \| |
| MQ=60 isize=234 | GCCCAGTCCCCACACCTACGAGGCCAGCAGATCCCTAATTCTCTCACCAATCAAGTGCGCTCTCCCCAGCCTGTCCCtTCTCCACGGCCACAGTCCCAG\|\_\|CCCCCCACTCCAGTCCTTCCCCAAGGATGCAGCCTCAGCCTTCTCCACACCa |
| 2:2110:5942:7545#1#F | \| \| |
| MQ=60 isize=200 | GCCCAGTCCCCACACCTACAAGGCCAGCAGATCCCTAATTCTCTCTCCAATCAAGTGCGCTCTCCCCAGCCTGTCCCTTCTCCACGGCCACAGtCCCAG\|\_\|CCCCCCaCtCCAGTCCTTCCCCAAGGAtgcagCCTCAgCCTtctccacaccc |
| 2:2114:6099:66257#1#F | \| \| |
| MQ=60 isize=159 | GCCCAGTCCCCACACCTACAAGGCCAGCAGATCCCTAATTCTCTCTCCAATCAAGTGCGCTCTCCCCAGCCTGTCCCTTCTCCACGGCCACAGTCCCAG\|\_\|CCCCCCACTCCAGTCCTTCCCCAAGGATGCAGCCTCAGCCTTCTCCACACCc |
| 2:2116:8782:63600#1#F | \| \| |
| MQ=60 isize=150 | GCCCAGTCCCCACACCTACAAGGCCAGCAGATCcCTAATTCTCTCTCCAATCAAGTGCGCTCTCCCCAGCCTGTCCCTTCTCCACGGCCACAGTCCCAG\|\_\|CCCCCCaCtCCAgtcCTTCCCCAAGGATGCAGCCtCaGccTTCTCCACACAG |
| 2:2203:17792:35915#1#F | \| \| |
| MQ=60 isize=121 | GCCCAGTCCCCACACCTACAAGGCCAGCAGATCCCTAATTCTCTCTCCAATCAAGTGCGCTCTCCcCAGCCTGTCCCTTCTCCACGGCCACAGTCCCAG\|\_\|CCCCCCACTCCAGTCCTTCCCAGGACTCCAATCGTCACCACACTCTGTCTCT |
| 2:2204:7287:62163#1#F | \| \| |
| MQ=60 isize=144 T=6 | GCCCAGTCCCCACACCTACAAGGCCAGCAGATCCCTAATTCTCTCTCCAATCAAGTGCGCTCTCCCCAGCCTGTCCCTTCTCCACGGCCACAGTCCCAG\|t\|cCCCCCACTCCAGTCCTTCCCCAAGGATGCAGCCTCAGCCTTCTAGGACTC |
| 2:2204:5367:82689#1#F | \| \| |
| MQ=60 isize=200 T=6 | GCCCAGTCCCCACACCTACAAGGCCAGCAGATCCCTAATTCTCTCTCCAATCAAGTGCGCTCTCCCCAGCCTGTCCCTTCTCCACGGCtaCAGTCCCAG\|t\|cccCCCaCTCCAGTCCTTCCCCAAGGATGCAGCCTCAGCCTTCTCCACACC |
| 2:2204:4764:100134#1#F | \| \| |
| MQ=60 isize=138 T=6 | GCCCAGTCCCCACACCTACAAGGCCAGCAGATCCCTAATTCTCTCTCCAATCAAGTGCGCTCTCCCCAGCCTGTCCCTTCTCCACGGCCACAGTCCCAG\|t\|cccCCCACTCCAGTCCTTCCCCAAGGATGCAGCCTCAGAGGACTCCAATCC |
| 2:2206:6058:49540#1#F | \| \| |
| MQ=60 isize=159 | GCCCAGTCCCCACACCTACAAGGCCAGCAGATCCCTAATTCTCTCTCCAATCAAGTGCGCTCTCCCCAGCCTGTCCCTTCTCCACGGCCACAGtCCCAG\|\_\|CCCCCCACTCCAGtCCTTCCCCAAGGATGCAGCCTCAGCCTTCTCCACACCa |
| 2:2208:13688:36300#1#F | \| \| |
| MQ=60 isize=201 | GCCCAGTCCCCACACCTACAAGGCCAGCAGATCCCTTATTCTCTCTCCAATCAAGTGCGCTCTCCCCAGCCTGTCCCTTCTCCACGGCCACAGTCCCAG\|\_\|CCCCCCACTCCAGTCCTTCCCCAAGGATGCAGCCTCAGCCTTCTCCACACCA |
| 2:2209:8293:48119#1#F | \| \| |
| MQ=60 isize=147 | GCCCAGTCCCCACACCTACAAGGCCAGCAGATCCCTAATTCTCTCTCCAATCAAGTGCGCTCTCCCCAGCCTGTCCCTTCTCCACGGCCACAGTCCCAG\|\_\|CCCCCCaCtCCAGtCcTtCCCcaaggatgcagcctcagcctcccccgggccc |
| 2:2211:14069:89663#1#F | \| \| |
| MQ=60 isize=240 | GCCCAGTCCCCACACCTACAAGGCCAGCAGATCCCTAATTCTCTCTCCAATCAAGTGCGCTCTCCCCAGCCTGTCCCTTCTCCACGGCCACAGTCCCAG\|\_\|CCCCCCACTCCAGTCCTTCCCCAAGGATGCAGCCTCAGCCTTCTCCACACCa |
| 2:2114:11737:99936#1#F | \| \| |
| MQ=60 isize=158 | CCCAGTCCCCACACCtACaAGGCCAGCAGATCCCtAATTCTCTCTCCAATCAAGtgCGCTCTCCCCAgCCTGTCCCTTCTcCACGGCCACAgtcCCAg\|\_\|CCCCCCaCtcCagTCCtTcCCCaAgGATGCAGCCtcaGCCttctccacaccac |
| 1:2212:16753:61055#2#R | \| \| |
| MQ=60 isize=-132 | cCCagtCCCCACAtCtacaaGGCCAgcaGATCCCTAattctctCtCCAATCAAGTgCGCTCTCCCCAGCCTGtcCCTTCTCCACGGCCACAGTCCCAG\|\_\|CCCCCCACtCCAGTCCTTCCCCAAGGATGCAGGACTCCAATTCGCCCCACCAC |
| 2:1210:6045:76798#2#R | \| \| |
| MQ=60 isize=-132 | CCCaGtCcCCacaCCtaCAAGGCCAGCAGATCCCTAATTCtCTCcCCAATCAAGTGCgCTCTCCCCAGCCtGTCCCTTCTCCACGGCcaCAGtCCCAG\|\_\|CCCCCCACTCCAGTCCTTCCCCAAGGATGCAGGACTCCAATTCGCCCCACCAC |
| 1:1202:7953:91713#2#R | \| \| |
| MQ=60 isize=-132 | CAGTCCCcaCaCCTACAAGGCCAGCAGATCCCTAAtTCTCTCTCCAATCaAGTGCGCTCTCCCCAGCCTGTCCCTTCTCcACGGCCACAGTCCCAG\|\_\|CCCCCCACTCCAGTCCTTCCCCAAGGATGCAGAGGACTCCAATCGACCCGCTGCC |
| 2:1104:15053:43686#2#R | \| \| |
| MQ=60 isize=-133 | cgtcCCCaCGCCTaCAAGGCCaGCAGATCCCTAATTCTCTCTCcAATCAAGTGcGCTCTCCCcAGCctGTCCCTtctCcACGGCCACAGTCCCAG\|\_\|cCCCCcACTCCAGTCCTTCCCCAAGGATGCAGCAGGACTCCAATCCGCCCTAGGAA |
| 1:2104:19213:87926#2#R | \| \| |
| MQ=60 isize=-140 | aCACCTACAAGGCCAGCAGaTCCCTAATTCTCTCTCCAATCAAGTGCGCtCTCCCCAGCCTGTCCCTTCTCCACGGCCACAGTCCCAG\|\_\|CCCCCCACTCCAGTCCTTCCCCAAGGATGCAGCCTCAGCCanGACTCCAATCCGTTTCCGATA |
| 2:1213:4831:88470#2#R | \| \| |
| MQ=60 isize=-143 | CcTACAAGGCCAGCAGAtCCCTAATtCTCTCTCCAATCaAGTGCgCtCTCCCCAgCCTGTCCCTTCTCCACGGCCACAGTCCCAG\|\_\|CCCCCCACTCCAGTCCTTCCCCAAGGATGCAGCCTCAGCCTTCAGGACTCCAATAAACCAGTAAGA |
| 2:1210:9113:40867#2#R | \| \| |
| MQ=60 isize=-147 | aCAAGGCCAGCAGATCCCTAATTCtCTCTCCAATCAAGTGCGCTCTCCCCAGCCTGTCCCTTCTCCACGGCCACAGTCCCAG\|\_\|CCCCCCACTCCAGTCCTTCCCCAAGGATGCAGCCTCAGCCTTCTCCAGGACTCCAATTCTATCAAGGGT |
| 2:2105:16250:81208#2#R | \| \| |
| MQ=60 isize=-147 | ccaaggccagcagatccctaatcctctctccaatcaagtgcgctctccccagCcTGTCCCTtCtCCACGGCCACAgtCCcag\|\_\|CcccCCACTCCAGTCCTTCCCCAAGGATGCAGCCTCAGCCTTCTCCanGACTCCAATAGGCGGGCTGCG |
| 2:2209:8293:48119#2#R | \| \| |
| MQ=60 isize=-147 | cccaggccagcaggtccctaattcccccccccatcaagtgcgctcccCCcAGCCTgtCCCTTctCCccgGccacagTCCCaG\|\_\|CCCCCCaCTCCAGTCCTTCCCCAAGGATGCAGCCTCAGCCTTCTCCanGACTCCAATTCTATCAAGGGT |
| 2:2116:8782:63600#2#R | \| \| |
| MQ=60 isize=-150 | ggccagcagatccctaattctctctCCAATCaAGTGCgCcCTCCCCAGCctGTCcCTTCtcCACGGCCACAGTCCCAG\|\_\|CCCCCCACTCCAGtCCTTCCCCAAGGATGCAGCCTCAGCCTTCTCCACACAgnACTCCAAtCTCCGCATTCCC |
| 2:1214:4187:53633#2#F | \| \| |
| MQ=60 isize=235 | GTTAGAGGGGCGATTGGAGTCCTGCCAGCAGATCCCTAATTCTCTCTCCAATCAAGTGCGCTCTCCCCAGCCTGTCCCTTCTCCACggcCACAGtCCCAG\|\_\|CCCcCCACTCCAGTCCTTCCCCaagGATgcagCCTCAGcCTTCTCCaCACC |
| 1:2205:8826:3969#2#R | \| \| |
| MQ=60 isize=-158 | gtcCCTAATTctCtccCCAATCaAGTGcgctCtCCCCAGCCTGTCCCTTCTCCACGGCCACAGTCCCAG\|\_\|CCCCCCaCTCCAGTCCTTCCCCAAGGATGCAGCCTCAGCCTTCTCCACACCACGTTTcaanGACTCCAATGCCCAAATCCCC |
| 2:2114:6099:66257#2#R | \| \| |
| MQ=60 isize=-159 | gtCCCTAATTCtCtccCCAATCAAGtGCGCtCtCCCCAGCCTGTCCCTTCtCCACGGCCACAGTCCCAG\|\_\|CCCCCCACTCCAGTCCTTCCCCAAGGAtGCAGCCTCAGCCTTCTCCACACCACGTTTCCAgnACTCCAATTATACGTTAGCT |
| 2:2114:11737:99936#2#R | \| \| |
| MQ=60 isize=-158 | aTCCCTAAttctCTCTCCAaTCAAGTGCGCTCTCCCCAGCCTGtCCCTTCTCCACGGCCACAGTCCCAG\|\_\|CCCCCCACTCCAGTCCTTCCCCAAGGATGCAGCCTCAGCCTTCTCCACACCACGTTTCCAgnACTCCAATTATACGTTAGCT |
| 2:2206:6058:49540#2#R | \| \| |
| MQ=60 isize=-159 | atCCCTaAtTCTCTCTCCAAtCAAGTGCgCtCTCCCCaGCCTGTCCCTTctcCACGGCCACAGTCCCaG\|\_\|CCCCcCaCTCCAGTCCTTCCCCAAGGATGCAGCCTCAGCCTTCTCCACACCACGTTTCCanGACTCCAATTATACGTTAGCT |
| 1:2111:20764:71366#2#R | \| \| |
| MQ=60 isize=-160 | tCCCTAAtTCtCtCtCCaATCAaGTGCGccctCCCCAGCCTGTCCCTTCTCCACGGCCACAGtcCCAG\|\_\|CCCCCCaCTCCAGTCCTTCCCCAAGGATGCAGCCTCAGCCTTCTCCACACCACGTTTCCCAGGACTCCAATAACAAAACATCC |
| 1:2212:16012:63327#2#R | \| \| |
| MQ=60 isize=-160 | cccctaattctccctCCAATCAAgtgCGCtCTCCCCAGCCTGTCCCTTCtCCACGGCCaCAGTCCCaG\|\_\|CCCCCCACTCCAGTCCTTCCCCAAGGATGCAGCCTCAGCCTTCTCCACACCACGTTTCCCAGGACTCCAATCTTCACAAGTGT |
| 1:2204:3514:6197#2#R | \| \| |
| MQ=60 isize=-162 | CCCTAaTTCTCtCTCCAATCaAGtGCGCtCTCCCCGGCCTGTCCCTTCTACACGGCCACAGTCcCAG\|\_\|CCCCCCACTCCAGTCCTTCCCCAAGGATGCAGCCTCAGCCTTCTCCACACCACGTTTCCCCAGGACTCCAATAACACAAGCGGT |
| 1:1204:5672:37361#2#F | \| \| |
| MQ=60 isize=223 | GAGTTGGCCGGAATTGGAGTCntCTAATTCTCTCTCCAATCAAGTGCGCTCTCCCCAGCCTGTCCCTTCTCCACGGCCACAGTCCCAG\|\_\|CCCCCCACTCCAGTCCTTCCCCAAGGATGCAGCCTCAGCCTTCTCCACACCACgtTTCCCCaC |
| 1:1213:13632:55762#2#F | \| \| |
| MQ=60 isize=223 | CGCCCCTCAAGTATTGGAGTncTCTAATTCTCTCTCCAATCAAGTGCGCTCTCCCCAGCCTGTCCCTTCTCCACGGCCACAGTCCCAG\|\_\|CCCCCCACTCCAGTCCTTCCCCAAGGATGCAGCCTCAGCCTTCTCCACACCcCGTTTCCCCAC |
| 2:2210:9595:85284#2#F | \| \| |
| MQ=60 isize=223 | CGCCCCTCAAGTATTGGAGTCCTCTAATTCTCTCTCCAATCAAGTGCGCTCTCCCCAGCCTGTCCCTTCTCCACGGCCACAGTCCCAG\|\_\|CCCCCCACTCCAGtCCTTCCCCAAGGATGCAGCCTCAGCCTTCTCCACaCCACgtttcCCCaC |
| 1:2115:2799:81460#2#R | \| \| |
| MQ=60 isize=-166 | AATtCTCTCTccAAtCAAGTGcgcccTCCCCAGCCcGTCCCTTCTcCAcgGCCACAGTCCCAG\|\_\|CCCCCCaCTCCAGTCCTTCCCCAAGGATGCAGCCTCAGCCTTCTCCACACCACGTTTCCCCACAGAGGACTCCAATCATCCTAGcTAG |
| 2:1104:20449:17134#2#R | \| \| |
| MQ=60 isize=-166 | aattctctctccaatcaagtgcGctCTCCCCAGCCTGTCCCTTCTCCACGGCCAcAGTCCCAG\|\_\|CCCCCCACTCCAGTCCTTCCCCAAGGATGCAGCCTCAGCCTTCTCCACACCACGTTTCCCCACAGAGGACTCCAATGCCTTTATTTCT |
| 2:1212:11395:86189#2#R | \| \| |
| MQ=60 isize=-166 | aattctctCTCCAATCAAGTGCGCTCtCCCCAGCCTGTCCCTTCTCCACGGCCcCAGTCCCAG\|\_\|CCCCCCACTCCAGTCCTTCCCCAAGGATGCAGCCTCAGCCTTCTCCACACCACGTTTCCCCACAGAGGACTCCAATGCCTTTATTTCT |
| 1:1103:9587:7568#2#R | \| \| |
| MQ=60 isize=-166 | aagcctctctccaatcaagtgcgctctccccagcctgtcccttctcctcGGCCacaGTCCCAG\|\_\|CCccCCgCTCCAGtCCTTCCCCAAGgaTGCAGCCtCaGCCTtCtCCAcAcCACgTTTCCCCaCAGAGGACTCCAATGCGAGATAGCGT |
| 1:1102:6025:59235#2#F | \| \| |
| MQ=60 isize=214 | AGAGGGGTATAGATTGGAGTCnttCTCCAATCAAGTGCgCTCTCCCCAGCCTGTCCCTTCTCCACGGCCACAGTCCCAG\|\_\|CCCCcCACTCCAGcCCTTCCCCaaGgAtgcaGCCTCAGcCTTCTCCACACCACGtttCcCcACAgaCAaGTt |
| 1:2214:3697:11043#2#R | \| \| |
| MQ=60 isize=-173 | CTCCAATCAAgTGCGCTCTCCCCAGCCTGTCCCTTCTCCACGGCCACAGTCCCAG\|\_\|CCCCCCaCTCCAGTCCTTCCCCAAGGATGCAGCCTCAGCCTTCTCCACACCACGTTTCCCCACAGACAAGTTCAgnACTCCAATGCGACTCCCTGG |
| 1:1206:8900:4083#2#R | \| \| |
| MQ=60 isize=-174 | cCCAATCAAGTGCGCTCTCCCCAGCCTGTCCCTTCTCCACGGCCACAGTCCCAG\|\_\|CCCCCCACTCCAGTCCTTCCCCAAGGATGCAGCCTCAGCCTTCTCCACACCACGTTTCCCCACAGACAAGTTCCAGGACTCCAATCACGTGAAAGTG |
| 2:1108:13131:61481#2#R | \| \| |
| MQ=60 isize=-174 | cCCAATCAagtGCGCTccCCCCAGCCTGTCCCTTCTCCACGGCCACAGtCCCAG\|\_\|CCCCCCactCCAGTCCTTCCCCAAGGATGCAGCCTCAGCCTTCTCCACACCACGTTTCCCCACAGACAAGTTCcanGACTCCAATCACGTGAAAGTG |
| 2:1115:6794:86554#2#R | \| \| |
| MQ=60 isize=-175 | CCAAtccAGTGCgCtctCCCCAGCCTGTCCCTTCtCCACGGCCACAGTCCCAG\|\_\|CCCCCCACTCCAGTCCTTCCCCAAGGATGCAGCCTCAGCCTTCTCCACACCACGTTTCCCCACAGACAAGTTCCCAgnACTCCAATTTATGCGCTCCG |
| 1:1106:12616:4344#2#R | \| \| |
| MQ=60 isize=-177 | CAATCAAGTGCgCtCTCCCCAGCCTGTCCCttCtCCACGGCCACAGTCCCAG\|\_\|CCCCCCaCtCCAGTCCTTCCCCAAGGATGCAGCCTCAGCCTTCTCCACACCACGTTTCCCCACAGACAAGTTCCCCanGACTCCAATGGCGCCCACGCC |
| 1:1115:15178:86872#2#R | \| \| |
| MQ=60 isize=-177 | CAATCAAGtGCGCTCTCCCCAGCCTGTCCCTTCTCCaCGGCCACAGTCCCAG\|\_\|CCCCCCaCTCCAGTCCTTCCCCAAGGATGCAGCCTCAGCCTTCTCCACACCACGTTTCCCCACAGACAAGTTCCCCAGGACTCCAATTCAAACAGGGCG |
| 1:1216:5228:47083#2#R | \| \| |
| MQ=60 isize=-181 | AAGTGCGCTCTCCCCAGCCTGTCCCTTCTCCACGGCCACAGTCCCAG\|\_\|CCCCCCACTCCAGTCCTTCCCCAAGGATGCAGCCTCAGCCTTCTCCACACCACGTTTCCCCACAGACAAGTTCCCCACATCAgnACTCCAATCCCCGCCCTCGC |
| 1:1105:3799:98934#2#F | \| \| |
| MQ=60 isize=202 | ACTTTAGTAGCAATTGGAGTCntGCGCTCTCCCCAGCCTGTCCCTTCTCCACGGCCACAGTCCCAG\|\_\|CCCCCCACTCCAGTCCTTCCCCAAGGATGCAGCCTCAGCCTTCTCCACACCACGTTCCCCCACAGaCAAGtTCCCCACATCCTGG |
| 1:2113:20520:36039#2#F | \| \| |
| MQ=60 isize=202 | AGgCTTATTGgtATTGGAGTnntgCGCTCTCCCCAGCCTGTCCCTTCTCCACGGCCACAGTCCCAG\|\_\|CCCCCCACTCCAGTCCTTCCCCAAGGATgCAGCCtCAgCCTTCTCCACACCACGTTTCCCCACAGACAAGtTCCCCACATCCTGG |
| 1:2216:16664:87382#2#F | \| \| |
| MQ=60 isize=202 | ACTTTAGTAGCAATTGGAGTncTGCGCTCTCCCCAGCCTGTCCCTTCTCCACGGCCACAGTCCCAG\|\_\|CCCCCCACTCCAGTCCTTCCCCAAGGATGCAgCCTcAGCCtTCTCCaCACCcCGTTTCCCCacagacAAGTTCCCCACAtCCTGG |
| 2:1213:5567:34019#2#F | \| \| |
| MQ=60 isize=202 | AGGCTTATTGGTATTGGAGTCCTGCGCTCTCCCCAGCCTGTCCCTTCTCCACGGCCACAGTCCCAG\|\_\|CCCCCCACTCCAGTCCTTCCCCAAGGATGCAGCCTCAGCCTTCTCCACACCaCGTTTCCCCACAGACAAGTTCCCCACATCCTGG |
| 2:2205:6084:20740#2#F | \| \| |
| MQ=60 isize=202 | AGGCTTATTGGTATTGGAGTCntGCGCTCTCCCCAGCCTGTCCCTTCTCCACGGCCACAGTCCCAG\|\_\|CCCCCCACTCCAGTCCTTCCCCAAGGATGCAGCCTCAGCCTTCTCCACACCACGTTTCCCCACAGACAAGTTCCCCACATCCTGG |
| 2:1206:13247:35678#2#F | \| \| |
| MQ=60 isize=199 G=37 | TTTAATCTAACTATTGGAGTCntGCTCTCCCCAGCCTGTCCCTTCTCCACGGCCACAGTCCCAG\|G\|CCCCCCACTCCAGTCCTTCCCCAAGGATGCAGCCTCaGCCTTCTCCACACCACGtTTCCCCACaGACAAGTTCCCCACATCCtggc |
| 2:1211:18391:55330#2#F | \| \| |
| MQ=60 isize=199 G=22 | TTTAATCTAACTATTGGAGTCCtGCTCTCCCCAgCCTGTCCCTTCTCCACgGCCACAGTCCCAG\|G\|CCCCCCACTCCAGTCCTTCCCCAAGGaTGCAGCCTCAGCCTTCTCCACACCaCGTTTCCCCACAgACAAGTTCCCCACatCCtgga |
| 2:2116:3921:80393#2#F | \| \| |
| MQ=60 isize=197 | TAATTAGGTGCAAtTGGAGTncTTCTCCCCAGCCTGTCCCTTCTCCACGGCCACAGTCCCAG\|\_\|CCCCCCACTCCAGTCCTTCCCCAAGGAtgcAgCCtcagCCTTctCCACaCCccgtTtCCCCacagacaagTcCCCcacaacctggactg |
| 2:2111:20589:59571#2#R | \| \| |
| MQ=60 isize=-155 T=2 | cagattcccagttcgctcccccaacacctacgccccccccagcctgtccttacgcctcggccccagtcccac\|t\|ccccccaccccagtccttccccaaggatgcagccgcaGCCttCTcCaCacCACGTAGGACTCCAAtCTcTGCGACtAa |
| 2:1103:3370:92957#2#F | \| \| |
| MQ=60 isize=324 | GGGGGGCGCGAGATTGGAGTCCTCAGCCTGTCCCTTCTCCACGGCCACAGTCCCAG\|\_\|CCCCCCaCTCCAGTCCTTCCCCaaggatgcagccccagccttccccccaccccgtttccccccaaccaagttccccccccccggacctggacgcc |
| 1:1114:17042:82311#2#R | \| \| |
| MQ=60 isize=-195 | CAGCCTGTCCCTTCtCCACGGCCACAGTCCCAG\|\_\|CCCCCCACTCCAGTCCTTCCCCAAGGATGCAGCCTCAGCCTTCTCCACACCACGTTTCCCCACAGACAAGTTCCCCACATCCTGGACTGGTAGCTAGGACTCCAATGGACTGGTGTGT |
| 2:2208:13688:36300#2#R | \| \| |
| MQ=60 isize=-201 | tGtCCCTTCTCCACGGCCACAGTCCCAG\|\_\|CCCcCCaCtccAGTCCTTCCCCAAGGATGCAGCCTCaGCCTTCTCCACACCacGTTTCCCCACAGACAAGTTCCCCACATCCTGGACTGGTAGCTGCCCAgnGGACTCCAATACGGAATGACC |
| 1:2109:16208:44268#2#R | \| \| |
| MQ=60 isize=-199 | GTCCCTTCtCCACGGCCACAGTCCCAG\|\_\|CCCCCCaCTCCAGTCCTTCCCCAAGGATGCagaCTCAGCCTTCTCcaCACCACGTTTCCCCACAGACAAGTTCCCCACATCCTGGACTGGTAGCTGCCCGganGACcCCAATAGTAGTGGTCTA |
| 2:1211:19609:50962#2#F | \| \| |
| MQ=60 isize=184 | GGATCCTGTCCCATTGGAGTCCTCCCTTCTCCACGGCCACAGTCCCAG\|\_\|CCCCCCACTCCAGtCCTTCCCCAAGGATGCAgCCTCAGCCtTCTCCACACCACGTTTCCCCACAGACAAGTTCCCCACATCCTGGACTGgtagCTGCCCAggC |
| 2:1104:2451:60232#2#R | \| \| |
| MQ=60 isize=-187 G=2 | cctgccgcagagtgccccttctacacgggcaccgtcccag\|g\|ccccccactccaggccttcccccaGGgTGCAGcCTCAGCctTCTccaccCcacgtTTCcCCccagACAAGTTCCCCaCATCCTGGACAGGACTCCAATCCgCGCCCTACT |
| 2:1102:8141:31258#2#F | \| \| |
| MQ=60 isize=316 | AGGGCCGCGGGTATTGGAGTCCTCCTTCTCCACGGCCACAGTCCCAG\|\_\|CCCCCCACTCCaGtCCTtcCCCAAGGAtgcagcctcagcctcccccccaccccgttcccccccagacaagtcccccacccccgggcctggcagctgcccggccc |
| 2:2112:5790:32376#2#F | \| \| |
| MQ=60 isize=287 | AGGGCCGCGGGTATTGGAGTncTCCTTCTCCACGGCCACAGTCCCAG\|\_\|CCCCCCACTCCAGTCCTTCCCCAAGGATGcaGCCTCAGCCTTCtCCaCACCaCGTTTCCCCaCAGacAagTtCCCCacaccctggactggtaccgccccaggca |
| 2:2208:9939:68079#2#F | \| \| |
| MQ=60 isize=182 | AGGGCCGCGGGTATTGGAGTCntCCTTCTCCACGGCCACAGTCCCAG\|\_\|CCCCCCACTCCAGTCCTTCCCCAAGGATGCAGCCTCAGCCTTCTCCACACCACGTTtCCCCACAGACAAGTTCCCCaCATCCTGGaCTGGtAGCTGCCCAGGCC |
| 1:1106:2654:5818#2#F | \| \| |
| MQ=60 isize=181 | ACTTCTGTGGTAATTGGAGTCntCTTCTCCACGGCCACAGTCCCAG\|\_\|CCCCCCACTCCAGTCCTTCCCCAAGGATGCAGCCTCAGCCTTCTCCACACCACGTTTCCCCACAGACAAGTTCCCCACATCCTGGACTGGTAGCTGCCCAGGCCA |
| 2:1215:13398:25834#2#F | \| \| |
| MQ=60 isize=181 | ACTTCTGTGGTAATTGGAGTncTCTTCTCCACGGCCACAGTCCCAG\|\_\|CCCCCCACTCCAGTCCTTCCCCAAGGATGCAGCCTCAGCCTTCTCCACACCACGTTTCcCCACAGACAAGTTCCCCACATCCtgGACTGGTAGCTGCCCAGGCCa |
| 2:2105:9677:41928#2#F | \| \| |
| MQ=60 isize=175 | TATACTTCAAATATTcGAGTCntCTCCACGGCCACAGTCCCAG\|\_\|CCCCCCACTCCAGTCCTTCCCCAAGGAtGCAGCCTCAgcCTtctccccaccccgtttccccccagacaagttcccccaaccctggactggacgcgccccagcccaacc |
| 2:2215:17794:25914#2#F | \| \| |
| MQ=60 isize=179 | TATACTTCAAATATTGGAGTCCTCTCCACGGCCACAGTCCCAG\|\_\|CCCCCCACTCCAGTCCTTCCCCAAGGATGCAGCCTCAGCCTTCTCCACACCACGTTTCCCCACAGACAAGTTCCCCACATCCTGGACTGGTAGCTGCCCAGGCCAACC |
| 1:1113:17323:6882#2#R | \| \| |
| MQ=60 isize=-207 | cctccacggccaCAGTCCcag\|\_\|CCCCCcaCTCCAgTCCttCcccAAGGATGCaGCctCAgccttctCCACaCCACGtttccCCACAGaCAAGTTCCCccCaTCCTGGAcTGGTAGCTGCCCAGGCCAACAGGACtCCAATTAGCATGACCCG |
| 1:1214:10886:65209#2#R | \| \| |
| MQ=60 isize=-207 | TCtCCACGGCCACAGtCCCAG\|\_\|CCCCCcaCTCCAGTCCTTCCCCAAGGATGCAGCCTCAGCCTTCTCCACACCACGTTTCCCCACAGACAAGTTCCCCACATCCTGGACTGGTAGCTGCCCAGGCCAACAgnACTCCAatCGCACTCCACGC |
| 1:2207:11364:61886#2#R | \| \| |
| MQ=60 isize=-208 | ctccacggccacagtcccag\|\_\|ccccccactccagtccttccccaaggatgcagcctcagccttctccacaccacgtttccccACAGACAAGTTCCCCACATCCTGGACTGGTAGCTGCCCAGGCCAACCanGtCTCCAATACACACGGCCAA |
| 1:1109:5015:21811#2#R | \| \| |
| MQ=60 isize=-211 | CCACGGCCACAGTCCCAG\|\_\|CCCCCCACTCCAGTCCTTCCCCAAGGATGCAGCCTCAGCCTTCTCCaCaCCACGTTTCCCCACAGACAAGTTCCCCACATCCTGGACTGGTAGCTGCCCAGGCCAACCCCAGGACTCCAATTAAGGCGAGCTA |
| 2:1114:6375:84122#2#F | \| \| |
| MQ=60 isize=280 | CTTTGCCATCTGATTGGAGTncTCACGGCCACAGTCCCAG\|\_\|CCCCCCACTCCAGTCCTTCCCCAAGGATGCAGCCTCAGCCTTCTCCACACCACGTTTCCCCaCAGACAAGTTCCCCACATCCTGGACTGGTAgCTGCCCAGGCCAACCCCa |
| 1:2104:14195:58688#2#F | \| \| |
| MQ=60 isize=174 | GGCGTTGCCCCATTGGAGTCCnacGGcCACagTCCCAg\|\_\|CCCCCCacTCCcgtCCTTcCCCaAGGATGCAGCCtcagCCtTCtccaccccacgtttccccacagacaagttccccacctcctggactggtagctgccccggccaaccccatg |
| 1:1212:17067:18737#2#R | \| \| |
| MQ=60 isize=-212 | ccggcCACAGTCCCAG\|\_\|CCCCCCACTCCAGtCCtTCCCCAAGGATGCAGCCTCAGCCTTCTCCacACCACGTTTCCCCACAGACAAGTTCCCCaCATCCTGGACTGGTAGCTGCCCAGGCCAACCCCATAGGACTCCAATCGTGGGAATTTC |
| 1:1107:15911:62731#2#F | \| \| |
| MQ=60 isize=172 | GGGAATTGGGAAATTGGAGTCCTGGCCACAGTCCCAG\|\_\|CCCCCCACTCCAGTCCTTCCCCAAGGATGCAGCCTCAGCCTTCTCCACACCACGTTTCCCCACAGACAAGTTCCCCACATCCTGGACTGGTAGCTGCCCAGGCCAACCCCATGG |
| 1:1211:18991:79822#2#F | \| \| |
| MQ=60 isize=172 | GGGAATTGGGAAATTGGAGTCCTGGCCACAGTCCCAG\|\_\|CCCCCCACTCCAGTCCTTCCCCAAGGATGCAGCCTCAGCCTTCTCCACACCACGTTTCCCCACAGACAAGTTCCCCACATCCTGGACTGGtAGCTGCCCAGGCCAACCCCATGg |
| 2:1201:5494:19762#2#F | \| \| |
| MQ=60 isize=172 | AAGCCCGAGTAAATTGGAGTCCTGGCCACAGTCCCAG\|\_\|CCCCCCACTCCAGTCCTTCCCCAAGGATGCAGCCTCAGCCTTCTCCACACCACGTTTCCCCACAgaCAAGTTCCCCACATCCTGgACTGGTAGCTGCCCaGgCCAACCCCATGg |
| 2:2205:12182:56823#2#F | \| \| |
| MQ=60 isize=304 | AGTTCAGATCAGATTGGAGTCntcCACAGTCCCAG\|\_\|CCcCCCACTCCAGTCCTTCCCCAAGGATGCAGCCTCAGCCTTCTCCACACCACGTTTCCCCACagACAAGTTCCCCACATCctGGACtgGtaGCTGCCCaGgCCAacCCCATGGAA |
| 2:2108:5390:79428#2#F | \| \| |
| MQ=60 isize=168 | GCTAGTTTCGTGATTGGAGTCCTACAGTCCCAG\|\_\|CCCCCCACTCCAGTCCTTCCCCAAGGATGCAGCCTCAGCCTTCTCCaCACCaCgtTTCCCCacAgacaAGTTCCCcacaccctggccgggtagctgccccggccaaccccatggaaca |
| 2:2202:10156:98187#2#R | \| \| |
| MQ=60 isize=-435 | cctgttcagcccagccccaggagcccccagcagcatatgctcccaaatcaggcccagtccccacacctacaaggccagcaggtcccgaattctctctccaatcaagtgcgctCTCccCAGcCTGtCcCAGGACTCCAAtatGtCAaTGgA\|T\| |
| 1:1112:11349:37561#2#R | \| \| |
| MQ=60 isize=-452 | CAtGaGCCCCCAGCAGCATATGCTCCCAAATCAGGCCCAGTCCCCACACCTACAAGGCCAGCAGATCCCTAATTCTCTCTCCAATCAAGTGCGCTCTCCCCAGCCTGTCCCTTCTCCACGGCCACAGTAgnAC\|T\|CCAATTCCTGGAAACAA |
| 2:2216:1778:6381#2#F | \| \| |
| MQ=60 isize=107 | AAGCCCGTGTGCATTGGAGTCCTCAAATCAGGCCCAGTCCCCACACCTACAAGGCCAGCAGATCCCTAATTCTCTCTCCAATCAAGTGCGCTCTCCCCAGCCTGTCCCTTCTCCACGGCCaCAGTCCCAG\|G\|TCCgTGGCGCTTGCACCATG |
| 2:2216:1778:6381#1#R | \| \| |
| MQ=60 isize=-107 | GTCCTCAAATCAGGCCCAGTCCCCACACCTACAAGGCCAGCAGATCCCTAATTCTCTCTCCAATCAAGTGCGCTCTCCCCAGCCTGTCCCTTCTCCACGGCCACAGTCCCAG\|G\|TCCGTGGCGCTTGCACCATGGAGGTTTGCCATGCCTGG |
| 1:1116:9833:36204#1#F | \| \| |
| MQ=60 isize=0 | GCCCAGTCCCCACACCTACAAGGCCAGCAGATCCCTAATTCTCTCTCCAATCAAGTGCGCTcTCCCCAGCCTGTCCCTTCTCCACTGCCACCACGCcCA\|G\|CTGATGTCTTTGTTtGAAagttATCACTCATGGTCGGgCGCGGTGGctcAC |
| 1:1214:7067:94036#1#F | \| \| |
| MQ=60 isize=0 | GCCCAGTCCCCACACCTACAAGGCCAGCAGATCCCTAATTCTCTCTCCAATCAAGTGCGCTCTCCCCAGCCTGTCCCTTCTAGGACTCCAATCATTCCA\|T\|GCAT |
| 1:2104:3516:51252#1#F | \| \| |
| MQ=60 isize=0 | GCCCAGTCCCCACACCTACAAGGCCAGCAGATCCCTAATTCTCTCTCCAATCAAGTGCGCTCTCCCCAGCCTGTCCCTTCTCCACGGCCACAGTCCCCC\|T\|TTTGAGTCCATTCTATATGATTCCATTCCATTCAAATCCATTACATTTGGT |
| 1:2107:8511:31421#1#F | \| \| |
| MQ=60 isize=0 | GCCCAGTCCCCACACCTACAAGGCCAtCAGATCCCTAATTCTCTCTCCAATCAAGTGCGCTCTCCCCAGCCTGTCCCTTCTCCACGGCCAAGACCAGCC\|T\|GGGCAACATGGCAAAATCCCATCTCTACAAAAAATACAAAAAAATTAGgtG |
| 1:2113:16738:14825#1#F | \| \| |
| MQ=60 isize=0 | GCCCAGTCCCCACACCTACAAGGCCAGCAGATCCCTAATTCTCTCTCCAATCAAGTGCGCTCTCCCCAGCCTGTCCCTTCTCCACGGCCACAGTCCCAG\|G\|GTCCTTCTCATTTTCACCAACGAGAAAACCAAGTTATAAAGATGGTAAGTA |
| 1:2205:1386:89465#1#F | \| \| |
| MQ=60 isize=0 | GCCCAGTCCCCACACCTACAAGGCCAGCAGATCCCTAATTCTCTCTCCAATCAAGTGCGCTCTCCCCAGCCTGTCCCTTCTCCACGGCCACAGTCCCAG\|G\|ACTCCAATAGTGAAAATaA |
| 1:2213:8953:49538#1#F | \| \| |
| MQ=60 isize=0 | GCCCAGTCCCCACACCTACAAGGCCAGCAGATCCCTAATTCTCTCTCCAATCAAGTGCGCTCTCCCCAGCCTGTCCCTTCTCCACGGCCACAGTCCCAT\|A\|AAAATCTGACTTATAGTAACTCCATATTTTAGAGATAAAGATTTAGGTAGG |
| 2:1103:19397:4244#1#F | \| \| |
| MQ=60 isize=0 | GCCCAGTCCCCACACCTACAAGGCCAGCAGATCCCTAATTCTCTCTCCAATCAAGTGCGCTCTCCCCAGCCTGTCCCTTCTCCACGGCCACAGTCCCAT\|A\|AAAATCTGACTTATAGTAACTCCATATTTTAGAGATAAaGaTTTAGGTAGG |
| 2:1106:15112:84760#1#F | \| \| |
| MQ=60 isize=0 | GCCCAGTCCCCACACCTACAAGGCCAGCAGATCCCTAATTCTCTCTCCAATCAAGTGCGCTCTCCCCAGCCTGTCCCCCTATGACAGAACTACATAAAA\|T\|TCAAGaAAGTAAAAGCTAGCAAAAACATTGTCTAAAACACACAATATTTTT |
| 2:1113:8646:56500#1#F | \| \| |
| MQ=60 isize=0 | GCCCAGTCCCCACACCTACAAGGCCAGCAGATCCCTAATTCTCTCTCCAATCAAGTGCGCTCTCCCCAGCCTGTCCCTTCTCCACGGCCACAGTCCCAT\|A\|AAAATCTGACTTATAGTAACTCCATATTTTAGAGATAAAGATTTAGGTAGG |
| 2:1115:1677:3732#1#F | \| \| |
| MQ=60 isize=0 | GCCCAGTCCCCACACCTACAAGGCCAGCAGATCCCTAATTCTCTCTCCAATCAAGTGCGCTCTCCCCAGCCTGTCCCTTCTCCACGGCCACGGGACTCC\|A\|AtCCCCGCCcCAAG |
| 2:1201:14532:62675#1#F | \| \| |
| MQ=60 isize=0 | GCCCAGTCCCCACACCTACAAGGCCAGCAGATCCCTAATTCTCTCTCCAATCAAGTGCGCTCTCCCCAGCCTGTCCCTTCTGCAAATTTCTCTCTCTCC\|A\|TTGATGTCCCCAGTAATTTCTGTCCAGAAAGCTGAAGATTTCATGATCCTT |
| 2:1212:15005:66995#1#F | \| \| |
| MQ=60 isize=0 | GCCCAGTCCCCACACCTACAAGGCCAGCAGATCCCTAATTCTCTCTCCAATCAAGTGCGCTCTCCCCAGCCTGTCCCTTCTCCACGGCAGGACTCCAAT\|T\|AAGGGCCGCG |
| 2:1214:11427:53383#1#F | \| \| |
| MQ=60 isize=0 | gCCCAgtcccCACACCTACAAGGCCAgCAGaTCCCTAATTCTCTCTCCAATCAAGtgCGcTCTCcCCaGCCTGTCCCtTCTCCACGGCCACAGTCCCAG\|A\|GgACTCCAAtgttcACGtCGAGCTGTTtATACACATCTCcGagCccACgag |
| 2:2114:18205:4692#1#F | \| \| |
| MQ=60 isize=77 | GCCCAGTCCCCACACCTACAAGGCCAGCAGATCCCTAATTCTCTCTCCAATCAAGTGCGCTCTCCCCAGCCTGTCCCCTGAGAATTCTTCTGTCTAGCA\|G\|AATATGAAGAAATCCCGTTTCCAACGAAAGCCTCAAAAATGTCAGGACTCC |
| 1:2111:19549:12269#1#F | \| \| |
| MQ=60 isize=0 | CCCAGTCCCCACACCTACAAGGCCAGCAGATCCCTAATTCTCTCTCCAATCAAGTGCGCTCTCCCCAGCCTGTCCCTTCAGGACTCCAATGTCCGTAC\|T\|T |
| 2:2114:18205:4692#2#R | \| \| |
| MQ=60 isize=-77 | CTACAAGgCCAGCagaTccCTAATTcTCtctccaaTCAaGtGCGCtcTCCCCAGCCTGtCCcCTGAGAATTCTTCTGTCTAGCA\|G\|AATATGAAGAAATCCCGTTTCCAACGAAAGCCTCAAAAATGTCAgnACTCCAATGGCGCCCGCCGA |
| 2:1204:21163:58203#2#F | \| \| |
| MQ=60 isize=284 C=33 | CGGAGCGGCAGCATTGGAGTCntcCCCAGCAGCATATGCTCCCAAATCAGGCCCAGTCCCCACACCTACAAGGCCAGCAGATCCCTAATTCTCTCTCCAATCAAGTGCGCTCTACCCAGCCTGTCCCTTCTCCACGGCCACAGTCCCAG\|C\|C |
| 1:2109:9918:44399#2#F | \| \| |
| MQ=60 isize=283 C=2 | CCCCCCATAACAATTGGAGTCntCCCAGCAGCATATGCTCCCAAATCAggcCCaGTCCCCaCACCTACAAGgCCAGCAGatCCCTAattCTCtCTCCAATCaAgTGCGCcCTCCCCAGCCTgtCCCTTCTCCaCGgCCACAGtcCCAg\|c\|cc |
| 2:2202:8349:62754#2#F | \| \| |
| MQ=60 isize=282 C=2 | GGTTCGCATCTGATTGGaGTCTTCCAGCAGCATATGCTCCCAAATCAGGCCCAGTCCCCACACCTACAaGGCCAGCAGAtCCCTAATTCTCtCTCCAAtCAagcgCGCtCtCCCCAgcCTgtCCCttCtCCaCGGCCACagtcccag\|c\|ccc |
| 1:2101:1518:51121#2#F | \| \| |
| MQ=60 isize=279 C=33 | CGAGGGGGGCCCATTGGAGTCCTGCAGCATATGCCCCCGAATCAGGCCCAGTCCCCACACCTACAAGGCCAGCAGATCCCTAATTCTCTCTCCAATCAAGTGCGCTCTCCCCAGCCTGTCCCTTCTCCACGGCCACAGTCCCAG\|C\|CCCCCC |
| 2:1108:2798:33428#2#F | \| \| |
| MQ=60 isize=279 C=2 | TCTAGTCCAtGCATTGGAGTCntGCAGCATATGCTCCCAAATCAGGCCCAGTCCCCACACCtACAAGgccagcagatccctaattctctctccaatcaagtgcgctctccccagccggcccctcctccacggccacagtcccac\|c\|cccccc |
| 1:2110:15009:51556#2#F | \| \| |
| MQ=60 isize=278 C=37 | CAAATTCCAAAGATTGGAGTCCTCAGCATATGCTCCCAAATCAGGCCCAGTCCCCACACCTACAAGGCCAGCAGATCCCTAATTCTCTCTCCAATCAAGTGCGCTCTCCCCAGCCTGTCCCTTCTCCACGGCCACAGTCCCAG\|C\|CCCCCCc |
| 2:2208:4246:97496#2#F | \| \| |
| MQ=60 isize=271 C=2 | TTCGGTCCCTTCATTGGAGTCntGcTCCCAAATCAGGCCCAGTCCCCACACCTACAAGgCCAGCAGATCCCTAATTCTCTCTCCaatcaagtgcgctctccccaccctgtcccttctccacggccacagtcccag\|c\|cccccccctccaccc |
| 1:2114:7756:79539#2#F | \| \| |
| MQ=60 isize=269 C=33 | AGCGTATGCAGCATTGGAGTCCTCCCAAATCAGGCCCAGTCCCCACACCTACAAGGCCAGCAGATCCCTAATTCTCTCTCCAATCAAGTGCGCTCTCCCCAGCCTGTCCCTTCTCCACGGCCACAGTCCCAG\|C\|CCCCCCACTCCAGtCCTt |
| 1:2213:16457:95367#2#F | \| \| |
| MQ=60 isize=269 C=2 | AGGTTtcGagGtATTGGAGtncTCCcAAAtCAGGCCCAGTCCCCaCAcCTacAaggCCAGCAgatccctaattctctctccaatcaagcgcgctctccccagcctgtcccttctccacggccacagtcccac\|c\|ccccccactccagtcctc |
| 2:2109:11896:24166#2#F | \| \| |
| MQ=60 isize=269 C=2 | CCGcCTCCCAcCATTGGAGTCCTCCCCCACCTTAGccTCCCAAATCaGGCccAGTCCCCACAcCTACAAggcCaGCAGATCcCTAATtctCTCtccaatcaagtgcgctctccccagcctgtcccttctccacgcccacaggcccag\|c\|ccc |
| 2:2214:8436:54248#2#F | \| \| |
| MQ=60 isize=269 C=22 | ACGGGGCTATCGATTGGAGTCCTCCCAAATCAGGCCCAGTCCCCACACCTACAAGGCCAGCAGATCCCTAATTCTCACtCCAATCAAGTgcgCtCtCCCCAGCCTGtCCCTtCTCCACggCCACAGTCCCag\|C\|CCCCCCACTCCAGtCCtt |
| 1:2113:14113:10332#2#F | \| \| |
| MQ=60 isize=268 C=2 | AGGTTGCGATGTATTGGAGTnntcCCAAATCAGGCCCAGTCCCCACACCTACAAGGCCAGCAGATCCCTAATTCTCTCTCcaatcaagtgcgctctccccagcctgtcccttctccacggccacagtcccag\|c\|ccccccactccagtcctt |
| 2:1204:19206:2706#2#F | \| \| |
| MQ=60 isize=267 C=33 | AAGAGCCGGGTAtTGGAGTCCncCCAAATCAGGCCCAGTCCCCACACCTACAAGGCCAGCAGATCCCTAATTCTCTCTCCAATCAAGTGCGCTCTCCCCAGCCTGTCCCTTCTCCACGGCCACAGTCCCAG\|C\|CCCCCCACtCCAGTCCTTC |
| 1:1110:6170:11066#2#F | \| \| |
| MQ=60 isize=265 C=33 | GGGTGCGAGTGAATTGGAGTCCTCAAATCAGGCCCAGTCCCCACACCTACAAGGCCAGCAGATCCCTAATTCTCTCTCCAATCAAGTGCGCTCTCCCCAgCCTGTCCCTTCTCCACGGCCACAGTCCCAg\|C\|CCCCCCACtCCAGtCCTTCC |
| 1:1204:15694:66876#2#F | \| \| |
| MQ=60 isize=262 C=33 | AGGATGGGTCTCATTGGAGTCntATCAGGCCCAGTCCCCACACCTACAAGGCCAGCAGATCCCTAATTCTCTCTCCAATCAAGTGCGCTCTCCCCAGCCTGTCCCTTCTCCACGGCCACAGTCCCAG\|C\|CCCCCCaCtCCAGtCCTTCCCCa |
| 2:1203:3681:97839#2#F | \| \| |
| MQ=60 isize=262 C=33 | AGGATGGGTCTCATTGGAGTCCTATCAGGCCCAGTCCCCACACCTACAAGGCCAGCAGATCCCTAATTCTCTCTCCAATCAAGtGCGCTCTCCCCAGCCTGTCCCTTCTCCACGgCCACAGtCCCAG\|C\|CCCCCCaCTCCAGTCCTtCCCCA |
| 1:1115:18659:94720#2#F | \| \| |
| MQ=60 isize=261 C=22 | CTCGCACCGTGCATTGGAGTCCTCAGGCCCAGTCCCCACACCTACAAGGCCAGCAGATCCCTAATTCTCTCTCCAATCAAGTGCGcTCTCCCCAGCCtgTCCCTTCTCCaCGgCCACAGTCCCAG\|C\|CCCCCCaCtCcAgtCCtTcCCCAAa |
| 1:1215:19964:72454#2#F | \| \| |
| MQ=60 isize=395 C=27 | GCAGACAAACCGATTGGAGTCCTCAGgCCCAGTCCCCACACCTACAAGGCCAGCAGATCCCTAATTCCCTCTCCAAtCAAGtGCgctcTCCCCAccCTgtccCTTCtCCaCggCCACAGTCCCAg\|C\|CCCCCCactccagcccttccccacc |
| 2:1210:18746:36404#2#F | \| \| |
| MQ=60 isize=260 C=37 | CACAGTCGTTTAATTGGAGTCCCAGGCCCAGTCCCCACACCTACAAGGCCAGCAGATCCCTAATTCTCTCTCCAATCAAGTGCGCTCTCCCCAGCCTGTCCCTTCTCCACGGCCACAGTCCCAG\|C\|CCCCCCACTCCAGTCCTTCCCCAAGG |
| 2:2114:7810:51415#2#R | \| \| |
| MQ=60 isize=-123 C=37 | tGGCCCAgtCCCCACcCCTACAAGGCCAGCAGATCCCTAATTCtCtCtCCAATCAAGTGCgCtCtCCCCAGCCTGTCCCTtCTCCACGGCCACAGTCCCAG\|C\|CCCCCCACTCCAGTCCTTCCCCAtGGAgnACTCCAAtCAGTTGCAGCGC |
| 1:1101:2725:34073#1#F | \| \| |
| MQ=60 isize=425 C=33 | GGCCCAGTCCCCACACCTACAAGGCCAGCAGATCCCTAATTCTCTCTCCAATCAAGTGCGCTCTCCCCAGCCTGTCCCTTCTCCACGGCCACAGTCCCAG\|C\|CCCCCCACTCCAGTCCTTCCCCAAGGATGCAGCCTCAGCCTTCTCCACAC |
| 2:1107:8031:71310#1#F | \| \| |
| MQ=60 isize=186 C=2 | TGCCCAGTCCCCACACCTACAAGGCCAGCAGATCCCTAATTCTCTCTCCAATCAAGTGCGCTCTCCCCAGCCTGTCCCTTCTCCACGGCCACAGTCCCag\|c\|ccccccaccccagccctcccccaaggatgcaacctcagccttctccacac |
| 2:2215:11484:97693#1#F | \| \| |
| MQ=60 isize=234 C=37 | GGCCCAGTCCCCACACCTACAAGGCCAGCAGATCCCTAATTCTCTCTCCAATCAAGTGCGCTCTCCCCAGCCTGTCCCTTCTCCACGGCCACAGTCCCAG\|C\|CCCCCCACTCCAGTCCTTCCCCAAGGATGCAGCCTCAGCCTTCTCCACAC |
| 1:1107:2704:45421#2#R | \| \| |
| MQ=60 isize=-127 C=37 | TTTTGGCCCAGTCCCCACACCTACAAGGCCAGCAGATCCCTAAttCtCTCTCCAATCAAGTGCGCTCTCCCCAGCCTGTCCCTTCTCCACGGCCACAGTCCCAG\|C\|CCCCCCACTCCAGTCCTTCCCCAAGGACTCCAATGCTGCATAGAAA |
| 1:1109:8182:46018#2#R | \| \| |
| MQ=60 isize=-121 C=37 | CGTTTTGGCCCAGTCCCCACaCCTACAAGGCCAGCAGATCCCTAATTCTCTCTCCAATCAAGTGCGCTCTCCCCAGCCTGTCCCTTCTCCACGGCCACAGTCCCAG\|C\|CCCCCCACTCCAGTCCTTCCCAGGACTCCAATTCTGACAAGCTA |
| 1:1114:4244:28468#2#R | \| \| |
| MQ=60 isize=-123 C=37 | gTtttGGCCCaGTCCCCaCACCTACAAGcCCAGCAGATCCCTAattctCTCTCCAatCaagTGCGCtCTCCCCAGCCTGTCCCttCtCCACGGCCACaGTCCCAG\|C\|CCCCCCACTCCAGTCCTTCCCCAGGACTCCAATGCTCCCCTACAC |
| 1:1201:14181:89606#2#R | \| \| |
| MQ=60 isize=-120 C=33 | tcgttttggccccgtccccacacccacaaggccagcagatccctaattctctcccccatccagggccctctcccccgcctgtcccttccccccggccaaaggcCCAG\|C\|CCCCCCACTCCAGTCCTTCCanGACTCCAATCACGAGGACCCC |
| 1:1205:7607:97269#2#R | \| \| |
| MQ=60 isize=-125 C=2 | ttggcccagcccccacatctacaagcccagcagctccctaatcccctccccaatcaagtgcgctctccccagcctgtcccttctccacggccacagtcccag\|c\|ccccccactccagtcCTTCCCCAAganGActccaagtcaCTTTGCTTA |
| 1:2103:18709:27958#2#R | \| \| |
| MQ=60 isize=-125 C=2 | ttggcccagtccccacccctacaaggccaccacatccctaattctctctccaatcaagtgcgccccccccagcctgtcccttctccacggccacagtcccag\|c\|ccccccactccagtcctaccccAAgAGGACTCCAATaCagTCaacaag |
| 1:2111:8443:48443#2#R | \| \| |
| MQ=60 isize=-125 C=37 | TTGGCCCagTCCCCAcaCCTACAAGGCCaGCAgaTCCCTAATTCTCTCTCCAATCAAgTGcgCTCTCCCCAGCCTGTCCCTtCTCCACGGCCACAGTCCCAG\|C\|CCCCCCACTCCAGTCCTTCCCCAAGAGGACTCCAATGGCGCGTGGATG |
| 1:2204:18578:100026#2#R | \| \| |
| MQ=60 isize=-124 C=37 | TTTGGCCCAGtCcccacaCCTaCAAGGCCAGCAGATCCCTAATtCtCtcTCCAATCAAGTgCGCTCTCCCCAGCCTGTCCCTTCTCCACGGCCACAGTCCCAG\|C\|CCCCCCACTCCAGTCCTTCCCCAAAGGACTCCAATTAATGCCCTGCC |
| 1:2207:5364:60522#2#R | \| \| |
| MQ=60 isize=-123 C=33 | gTTTTggcCCAGTCCCcACACCTACAAGGCCAGCAGATCCCTAATTCTCTCTCCAATCAAGTGCGCTCTCCCCAGCCTGTCCCttCTCCACGGCCACAGTCCCAG\|C\|CCCCCCACTCCAGTCCTTCCCcanGACTCCAATACAGTCTTGCCG |
| 1:2216:4954:11018#2#R | \| \| |
| MQ=60 isize=-125 C=22 | TTgGCCcAgtCCCCACACCTACAAGgCCAGCAGaTCCCTAATtCtCTCTCCAATCAAGTGCGCTCtCCCCAGCCTGTCCCTTCTCcaCGGCCACAGTCCCAG\|C\|CCCCCCACTCCAGTCCTTCCCCAAGAgnACTCCAATGTGAAAAACTCG |
| 2:1106:21271:80357#2#R | \| \| |
| MQ=60 isize=-123 C=22 | cttttggcccagtccccaCACCTaCAAgcCcagCAgatCCctaattCTctCtCCaatCAagtGCgCtCTCCCCAGCCTGTCCCttCtCCACGGCCACAgtCCCAG\|C\|CCCCCCACTCCAGTCCTTCCCcanGACTCCAATAAACGACACTTA |
| 2:1110:17771:14057#2#R | \| \| |
| MQ=60 isize=-125 C=27 | TTGGCcCaGTcCCcccACCTACAAGGCCAGCAGATCcCTAattCtctcTccAATCaaGtgCGCtCTCCCCAGCCTGTCCCTTCTCCACGGCCACAGTCCcAG\|C\|CCCCCCACTCCAGTCCTTCCCCAAGAGGACTCCAATCTGAAATGTAAA |
| 2:1116:15265:36054#2#R | \| \| |
| MQ=60 isize=-125 C=33 | TTGGCCCAGTCcccACACCTACAAGGCCAGCAGaTccCTAATTCTCTCTCCAATCAAGTGCGCTCTCCCCAGCCTGTCCCTTCTCCACGGCCACAGTCCCAG\|C\|CCCCCCACTCCAGTCCTTCCCCAAGAGGACTCCAATACAGTCAACAAG |
| 2:1203:17037:49995#2#R | \| \| |
| MQ=60 isize=-121 C=37 | CGTTTTGGCCCAGTCCCCACACCTACAAGGCCAGCAGATCCCTAATTCTCTCTCCAATCAAGTGCGCTCTCCCCAGCCTGTCCCTTCTCCACGGCCACAGTCCCAG\|C\|CCCCCCACTCCAGTCCTTCCCAGGACTCCAATCGTCACCACACT |
| 2:1205:3304:69946#2#R | \| \| |
| MQ=60 isize=-121 C=2 | ggttttggcccagtccccccccctacaaggccagcagacccttaattccctctccaatcaagtcccctctccccaccctgccccttctccacgcccacagcccccg\|c\|ccccccactccagtccttcccangactcaaaTTTTCCCGCCTAT |
| 2:1212:4486:9387#2#R | \| \| |
| MQ=60 isize=-123 C=37 | gtTTTGGCCCAGTCCCCaCACCTACAAGGCCAGCAGATCCCTAATTCTCTCTCCAATCAAGTGCGCtCTCCCCAGCCTGTCCCTTCTCCACGGCCACAGTCCCAG\|C\|CCCCCCACTCCAGTCCTTCCCCAGGACTCCAATGTCAGGACCTTA |
| 2:1213:6079:39542#2#R | \| \| |
| MQ=60 isize=-123 C=37 | gtTTTGGCCCAGTCCCCACaCCTACAAGGCCAGCaGaTCCCTAATTCTCTCTCCAATCAAGTGCgctCTCCCCAGCctgTCCCTTCTCCACGGCCACAGTCCCAG\|C\|CCCCCCaCTCCAGTCCTTCCCCAGGACTCCAATCTGCTGTTTCAG |
| 2:2107:20767:55222#2#R | \| \| |
| MQ=60 isize=-125 C=37 | TTGGCCCaGtCCCCacaCCtaCAaggccAGCAGaTCCCTAAttCTCTCTCCAATCAAGTGCGCTCTCCCCAGCCTGTCCCTTCTCCACGGCCACAGTCCCAG\|C\|CCCCCCACTCCAGTCCTTCCCCAAGAGGACTCCAATCCTCAAGAGCGC |
| 2:2207:11493:42022#2#R | \| \| |
| MQ=60 isize=-124 C=27 | tttggcccagtccccacaCCTaCaAGgCCAGCAGATCCCTAattCTCTCaCCAATCAAGTGCGCtCTCCCCAGCCTGTCCCTTCTCCACGGCCACAGTCCCAG\|C\|CCCCCCACTCCAGTCCttCCCCAaanGActCCAATCCATGCCGCCCG |
| 2:2213:14390:68804#2#R | \| \| |
| MQ=60 isize=-121 C=37 | CGTTTTGGCCCAgcCCCCAcaCCTACAAGGCCAGCAGATCCCtAAttCtctCtCCAATCAAGTGCGctCtCCCCAGCCTGtCCCttCtCCACGGCCACAGTCCCAG\|C\|CCCCCCACTCCAGTCCTTCCCAGGACtCCAATGTTAGAACCATG |
| 1:1101:1457:16610#1#F | \| \| |
| MQ=60 isize=139 C=33 | GCCCAGTCCCCACACCTACAAGGCCAGCAGATCCCTAATTCTCTCTCCAATCAAGTGCGCTCTCCCCAGCCTGTCCCTTCTCCACGGCCACAGTCCCAG\|C\|CCCCCCACTCCAGTCCTTCCCCAAGGATGCAGCCTCAGCAGGACTCCAATG |
| 1:1101:3471:34116#1#F | \| \| |
| MQ=60 isize=173 C=33 | GCCCAGTCCCCACACCTACAAGGCCAGCAGATCCCTAATTCTCTCTCCAATCAAGTGCGCTCTCCCCAGCCTGTCCCTTCTCCACGGCCACAGTCCCaG\|C\|CCCCCCACTCCAGTCCTTCCCCAAGGATGCAGCCTCAGCCTTCTCCACACC |
| 1:1101:2642:96222#1#F | \| \| |
| MQ=60 isize=132 C=37 | GCCCAGTCCCCACACCTACAAGGCCAGCAGATCCCTAATTCTCTCTCCAATCAAGTGCGCTCTCCCCAGCCTGTCCCTTCTCCACGGCCACAGTCCCAG\|C\|CCCCCCACTCCAGTCCTTCCCCAAGGATGCAGAGGACTCCAATCCCCATTA |
| 1:1102:17557:31313#1#F | \| \| |
| MQ=60 isize=262 C=37 | GCCCAGTCCCCACACCTACAAGGCCAGCAGATCCCTAATTCTCTCTCCAATCAAGTGCGCTCTCCCCAGCCTGTCCCTTCTCCACGGCCACAGTCCCAG\|C\|CCCCCCACTCCAGTCCTTCCCCAAGGATGCAGCCTCAGCCTTCTCCACaCC |
| 1:1102:6000:37275#1#F | \| \| |
| MQ=60 isize=250 C=37 | GCCCAGTCCCCACACCTACAAGGCCAGCAGATCCCTAATTCTCTCTCCAATCAAGTGCGCTCTCCCCAGCCTGTCCCTTCTCCACGGCCACAGTCCCAG\|C\|CCCCCCACTCCAGTCCTTCCCCAAGGATGCAGCCTCAGCCTTCTCCACaCC |
| 1:1102:2860:51157#1#F | \| \| |
| MQ=60 isize=201 C=37 | GCCCAGTCCCCACACCTACAAGGCCAGCAGATCCCTAATTCTCTCTCCAATCAAGTGCGCTCTCCCCAGCCTGTCCCTTCTCCACGGCCACAGTCCCAG\|C\|CCCCCCACTCCAGTCCTTCCCCAAGGATGCAGCCTCAGCCTTCTCCACACC |
| 1:1103:11093:11985#1#F | \| \| |
| MQ=60 isize=139 C=33 | GCCCAGTCCCCACACCTACAAGGCCAGCAGATCCCTAATTCTCTCTCCAATCAAGTGCGCTCTCCCCAGCCTGTCCCTTCTCCACGGCCACAGTCCCAG\|C\|CCCCCCACTCCAGtCCTTCCCCAAGGaTGCAGCCTCAGCAGGACTCCAATa |
| 1:1103:16405:34667#1#F | \| \| |
| MQ=60 isize=302 C=37 | GCCCAGTCCCCACACCTACAAGGCCAGCAGATCCCTAATTCTCTCTCCAATCAAGTGCGCTCTCCCCAGCCTGTCCCTTCTCCACGGCCACAGTCCCAG\|C\|CCCCCCACTCCAGTCCTTCCCCAAGGATGCAGCCTCAGCCTTCTCCACACC |
| 1:1103:6427:79626#1#F | \| \| |
| MQ=60 isize=236 C=33 | GCCCAGTCCCCACACCTACAAGGCCAGCAGATCCCTAATTCTCTCTCCAATCAAGTGCGCTCTCCCCAGCCTGTCCCTTCTCCACGGCCACAGTCCCAG\|C\|CCCCCCACTCCAGTCCTTCCCCAAGGATGCAGCCTCAGCCTTCTCCACACC |
| 1:1104:16727:47588#1#F | \| \| |
| MQ=60 isize=144 C=37 | GCCCAGTCCCCACACCTACAAGGCCAGCAGATCCCTAATTCTCTCTCCAATCAAGTGCGCTCTCCCCAGCCTGTCCCTTCTCCACGGCCACAGTCCCAG\|C\|CCCCCCACTCCAGTCCTTCCCCAAGGATGCAGCCTCAGCCTTCTAGGACTC |
| 1:1104:19066:60446#1#F | \| \| |
| MQ=60 isize=158 C=33 | GCCCAGTCCCCACACCTACAAGGCCAGCAGATCCCTAATTCTCTCTCCAATCAAGTGCGCTCTCCCCAGCCTGTCCCTTCTCCACGGCCACAGTCCCAG\|C\|CCCCCCACTCCAGTCCTTCCCCAAGGATGCAGCCTCAGCCTTCTCCACACC |
| 1:1104:16898:69060#1#F | \| \| |
| MQ=60 isize=192 C=37 | GCCCAGTCCCCACACCTACAAGGCCAGCAGATCCCTAATTCTCTCTCCAATCAAGTGCGCTCTCCCCAGCCTGTCCCTTCTCCACGGCCACAGTCCCAG\|C\|CCCCCCACTCCAGTCCTTCCCCAAGGATGCAGCCTCAGCCTTCTCCACACC |
| 1:1104:19111:77348#1#F | \| \| |
| MQ=60 isize=129 C=37 | GCCCAGTCCCCACACCTACAAGGCCAGCAGATCCCTAATTCTCTCTCCAATCAAGTGCGCTCTCCCCAGCCTGTCCCTTCTCCACGGCCACAGTCCCAG\|C\|CCCCCCACTCCAGTCCTTCCCCAAGGATGAGGACTCCAATAGGAAGCGAAG |
| 1:1104:4360:95929#1#F | \| \| |
| MQ=60 isize=160 C=37 | GCCCAGTCCCCACACCTACAAGGCCAGCAGATCCCTAATTCTCTCTCCAATCAAGTGCGCTCTCCCCAGCCTGTCCCTTCTCCACGGCCACAGTCCCAG\|C\|CCCCCCACTCCAGTCCTTCCCCAAGGATGCAGCCTCAGCCTTCTCCACACC |
| 1:1105:20661:46888#1#F | \| \| |
| MQ=60 isize=285 C=37 | GCCCAGTCCCCACACCTACAAGGCCAGCAGATCCCTAATTCTCTCTCCAATCAAGTGCGCTCTCCCCAGCCTGTCCCTTCTCCACGGCCACAGTCCCAG\|C\|CCCCCCACTCCAGtCCTTCCCCAAGGATGCAGCCTCAGCCTTCTCCACACC |
| 1:1105:7126:74009#1#F | \| \| |
| MQ=60 isize=177 C=33 | GCCCAGTCCCCACACCTACAAGGCCAGCAGATCCCTAATTCTCTCTCCAATCAAGTGCGCTCTCCCCAGCCTGTCCCTTCTCCACGGCCACAGTCCCAG\|C\|CCCCCCACTCCAGTCCTTCCCCAAGGATGCAGCCTCAGCCTTCTCCACACC |
| 1:1106:19795:22811#1#F | \| \| |
| MQ=60 isize=186 C=33 | GCCCAGTCCCCACACCTACAAGGCCAGCAGATCCCTAATTCTCTCTCCAATCAAGTGCGCTCTCCCCAGCCTGTCCCTTCTCCACGGCCACAGTCCCAG\|C\|CCCCCCACTCCAGTCCTTCCCCAAGGATGCAGCCTCAGCCTtcTCCaCACC |
| 1:1106:7735:56565#1#F | \| \| |
| MQ=60 isize=202 C=33 | GCCCAGTCCCCACACCTACAAGGCCAGCAGATCCCTAATTCTCTCTCCAATCAAGTGCGCTCTCCCCAGCCTGTCCCTTCTCCACGGCCACAGTCCCAG\|C\|CCCCCCACTCCAGTCCTTCCCCAAGGATGCAGCCTCAGCCTTCTCCACACC |
| 1:1106:13117:79295#1#F | \| \| |
| MQ=60 isize=221 C=37 | GCCCAGTCCCCACACCTACAAGGCCAGCAGATCCCTAATTCTCTCTCCAATCAAGTGCGCTCTCCCCAGCCTGTCCCTTCTCCACGGCCACAGTCCCAG\|C\|CCCCCCACTCCAGTCCTTCCCCAAGGATGCAGCCTCAGCCTTCTCCACACC |
| 1:1107:12928:11322#1#F | \| \| |
| MQ=60 isize=144 C=37 | GCCCAGTCCCCACACCTACAAGGCCAGCAGATCCCTAATTCTCTCTCCAATCAAGTGCGCTCTCCCCAGCCTGTCCCTTCTCCACGGCCACAGTCCCAG\|C\|CCCCCCACTCCAGTCCTTCCCCAAGGATGCAGCCTCAGCCTTCTAGGACTC |
| 1:1107:11338:20494#1#F | \| \| |
| MQ=60 isize=177 C=33 | gCCCAGTCCCCACACCTACAAGGCCAGCAGATCCCTAATTCTCTCTCCAATCAAGTGCGCTCTCCCCAGCCTGTCCCTTCTCCACGGCCACAGTCCCAG\|C\|CCCCCCACTCCAGTCCTTCCCCAAGGATGCAGCCTCAGCCTTCTCCACACC |
| 1:1107:21155:65033#1#F | \| \| |
| MQ=60 isize=170 C=33 | GCCCAGTCCCCACACCTACAAGGCCAGCAGATCCCTAATTCTCTCTCCAATCAAGTGCGCTCTCCCCAGCCTGTCCCTTCTCCACGGCCACAGTCCCAG\|C\|CcCCCcACTCCAGTCCTTCCCCAAGGATGCAGCCTCAGCCTTCTCCACACC |
| 1:1108:16808:4941#1#F | \| \| |
| MQ=60 isize=0 C=37 | GCCCAGTCCCCACACCTACAAGGCCAGCAGATCCCTAATTCTCTCTCCAATCAAGTGCGCTCTCCCCAGCCTGTCCCTTCTCCACGGCCACAGTCCCAG\|C\|CCCCCCACTCCAGTCCTTCCCCAAGGATGCAGCCTCAGCCTTCTCCACACC |
| 1:1108:6180:35901#1#F | \| \| |
| MQ=60 isize=192 C=33 | CCCCAGTCCCCACACCTACAAGGCCAGCAGATCCCTAATTCTCTCTCCAATCAAGTGCGCTCTCCCCAGCCTGTCCCTTCTCCACGGCCACAGTCCCAG\|C\|CCCCCCACTCCAGtCCTTCCCCAAGGATGCAGCCTCAGCCTTCTCCACACC |
| 1:1108:5323:39155#1#F | \| \| |
| MQ=60 isize=131 C=37 | GCCCAGTCCCCACACCTACAAGGCCAGCAGATCCCTAATTCTCTCTCCAATCAAGTGCGCTCTCCCCAGCCTGTCCCTTCTCCACGGCCACAGTCCCAG\|C\|CCCCCCACTCCAGTCCTTCCCCAAGGATGCAGGACTCCAATCACTTATATC |
| 1:1108:16871:61225#1#F | \| \| |
| MQ=60 isize=292 C=37 | GCCCAGTCCCCACACCTACAAGGCCAGCAGATCCCTAATTCTCTCTCCAATCAAGTGCGCTCTCCCCAGCCTGTCCCTTCTCCACGGCCACAgTCCCAG\|C\|CCCCCCACTCCAGTCCTTCCCCAAGGATGCAGCCTCAGCCTTCTCCACACC |
| 1:1108:5778:92966#1#F | \| \| |
| MQ=60 isize=195 C=37 | GCCCAGTCCCCACACCTACAAGGCCAGCAGATCCCTAATTCTCTCTCCAATCAAGTGCGCTCTCCCCAGCCTGTCCCTTCTCCACGGCCACAGTCCCAG\|C\|CCCCCCACTCCAGTCCTTCCCCAAGGATGCAGCCTCaGCCTTCTCCACACC |
| 1:1108:13029:99582#1#F | \| \| |
| MQ=60 isize=0 C=33 | GCCCAGTCCCCACACCTACAAGGCCAGCAGATCCCTAATTCTCTCTCCAATCAAGTGCGCTCTCCCCAGCCTGTCCCTTCTCCACGGCCACAGTCCCAG\|C\|CCCCCCACTCCAGTCCTTCCCCAAGGATGCAGCCTCAGCCTTCTCCACACC |
| 1:1109:17500:22398#1#F | \| \| |
| MQ=60 isize=123 C=37 | GCCCAGTCCCCACACCTACAAGGCCAGCAGATCCCTAATTCTCTCTCCAATCAAGTGCGCTCTCCCCAGCCTGTCCCTTCTCCACGGCCACAGTCCCAG\|C\|CCCCCCACTCCAGTCCTTCCCCAGGACTCCAATACAGTCTTGCCGCTGTCT |
| 1:1109:13191:59852#1#F | \| \| |
| MQ=60 isize=145 C=15 | GCCCAGTCCCCACACCTACAAGGCCAGCAGATCCCTAATTCTCTCTCCAATCAAGTGCGCTCTCCCCAGCCTgTCCCTTCTCCACGGCCACAGTCCCAg\|c\|CcCCCCACTCCAGTCCTTCCCCAAgGATGCatcCTCAGCCtTCTCaggacc |
| 1:1109:6555:78796#2#F | \| \| |
| MQ=60 isize=257 C=37 | TAATAGGAAAGCATTGGAGTCCTGCCCAGTCCCCACACCTACAAGGCCAGCAGATCCCTAATTCTCTCTCCAATCAAGTGCgCTCTCCCCAGCCTGTCCCTTCTCCACGGCCACAGTCCCAG\|C\|CCCCCCACTCCAgtCCTtCCCCAAGGAt |
| 1:1110:12229:15928#1#F | \| \| |
| MQ=60 isize=0 C=37 | GCCCAGTCCCCACACCTACAAGGCCAGCAGATCCCTAATTCTCTCTCCAATCAAGTGCGCTCTCCCCAGCCTGTCCCTTCTCCACGGCCACAGTCCCAG\|C\|CCCCCCACAGGACTCCAATTCGCAGCCTGGA |
| 1:1110:14985:46283#1#F | \| \| |
| MQ=60 isize=149 C=37 | GCCCAGTCCCCACACCTACAAGGCCAGCAGATCCCTAATTCTCTCTCCAATCAAGTGCGCTCTCCCCAGCCTGTCCCTTCTCCACGGCCACAGTCCCAG\|C\|CCCCCCACTCCAGTCCTTCCCCAAGGATGCAGCCTCAGCCTTCTCCACAGG |
| 1:1110:5299:86368#1#F | \| \| |
| MQ=60 isize=211 C=33 | GCCCAGTCCCCACACCTACAAGGCCAGCAGATCCCTAATTCTCTCTCCAATCAAGTGCGCTCTCCCCAGCCTGTCCCTTCTCCAcGGCCACAGTCCCAG\|C\|CCCCCCACTCCAGTCCTTCCCCAAGGATGCAGCCTCAGCCtTCTCCACACC |
| 1:1110:18353:96221#1#F | \| \| |
| MQ=60 isize=120 C=33 | GCCCAGTCCCCACACCTACAAGGCCAGCAGATCCCTAATTCTCTCTCCAATCAAGTGCGCTCTCCCCAGCCTGTCCCTTCTCCACGGCCACAGTCCCAG\|C\|CCCCCCACTCCAGTCCTTCCAGGACTCCAATCATCACTCTCACCTGTCTCT |
| 1:1111:17398:73213#1#F | \| \| |
| MQ=60 isize=218 C=37 | GCCCAGTCCCCACACCTACAAGGCCAGCAGATCCCTAATTCTCTCTCCAATCAAGTGCGCTCTCCCCAGCCTGTCCCTTCTCCACGGCCACAGTCCCAG\|C\|CCCCCCACTCCAGTCCTTCCCCAAGGATGCAGCCTCAGCCTTCTCCACACC |
| 1:1111:5284:77785#1#F | \| \| |
| MQ=60 isize=175 C=33 | GCCCAGTCCCCACACCTACAAGGCCAGCAGATCCCTAATTCTCTCTCCAATCAAGTGCGCTCTCCCCAGCCTGTCCCTTCTCCACGGCCACAGTCCCAG\|C\|CCCCCCACTCCAGTCCTTCCCCAAGGATGCAGCCTCAGCCTTCTCCACACC |
| 1:1111:10984:91915#1#F | \| \| |
| MQ=60 isize=147 C=37 | GCCCAGTCCCCACACCTACAAGGCCAGCAGATCCCTAATTCTCTCTCCAATCAAGTGCGCTCTCCCCAGCCTGTCCCTTCTCCACGGCCACAGTCCCAG\|C\|CCCCCCACTCCAGTCCTTCCCCAAGGATGCAGCCTCAGCCTTCTCCAGGAC |
| 1:1112:2524:33394#1#F | \| \| |
| MQ=60 isize=145 C=37 | GCCCAGTCCCCACACCTACAAGGCCAGCAGATCCCTAATTCTCTCTCCAATCAAGTGCGCTCTCCCCAGCCTGTCCCTTCTCCACGGCCACAGTCCCAG\|C\|CCCCCCACTCCAGTCCTTCCCCAAGGATGCAGCCTCAGCCTTCTCAGGACT |
| 1:1112:17481:36083#1#F | \| \| |
| MQ=60 isize=179 C=33 | GCCCAGTCCCCACACCTACAAGGCCAGCAGATCCCTAATTCTCTCTCCAATCAAGTGCGCTCTCCCCAGCCTGTCCCTTCTCCACGGCCACAGTCCCAG\|C\|CCCCCCATTCCAGtCCTTCCCCAAGGATGCAGCCTCAGCCTTCTCCACACC |
| 1:1112:13726:55512#1#F | \|C\| |
| MQ=60 isize=208 C=33 | GCCCAGTCCCCACACCTACAAGGCCAGCAGATCCCTAATTCTCTCTCCAATCAAGTGCGCTCTCCCCAGCCTGTCCCTTCTCCACGGCCACAGTCCCAI\|C\|CCCCCCATTCCAGTCCTTCCCCAAGGATGCAGCCTCAGCCTTCTCCACAC |
| 1:1113:17926:5966#1#F | \| \| |
| MQ=60 isize=322 C=37 | GCCCAGTCCCCACACCTACAAGGCCAGCAGATCCCTAATTCTCTCTCCAATCAAGTGCGCTCTCCCCAGCCTGTCCCTTCTCCACGGCCACAGTCCCAG\|C\|CCCCCCACTCCAGTCCTTCCCCAAGGATGCAGCCTCAGCCTTCTCCACACC |
| 1:1113:13421:40391#1#F | \| \| |
| MQ=60 isize=366 C=37 | GCCCAGTCCCCACACCTACAAGGCCAGCAGATCCCTAATTCTCTCTCCAATCAAGTGCGCTCTCCCCAGCCTGTCCCTTCTCCACGGCCACAGTCCCAG\|C\|CCCCCCACTCCAGTCCTtCCCCAAGgaTGCAGCCTCAGCCTTCTCCACACC |
| 1:1113:9491:76232#1#F | \| \| |
| MQ=60 isize=221 C=37 | GCCCAGTCCCCACACCTACAAGGCCAGCAGATCCCTAATTCTCTCTCCAATCAAGTGCGCTCTCCCCAGCCTGTCCCTTCTCCACGGCCACAGTCCCAG\|C\|CCCCCCACTCCAGTCCTTCCCCAAGGATGCAGCCTCAGCCTTCTCCACACC |
| 1:1114:18023:5083#1#F | \| \| |
| MQ=60 isize=239 C=27 | GCCCAGTCCCCACACCTACAAGGCCAGCAGATCCCTAATTCTCTCTCCAATCAAGTGCGCTCTCCCCAGCCTGTCCCTTCTCcacggCCACAGtCCCAG\|C\|CcCCCCactCCAGTcCTTCCCCAaggATGCAGCCtCagccttctccacacc |
| 1:1114:16073:12881#1#F | \| \| |
| MQ=60 isize=165 C=33 | GCCCAGTCCCCACATCTACAAGGCCAGCAGATCCCTAATTCTCTCTCCAATCAAGTGCGCTCTCCCCAGCCTGTCCCTTCTCCACGGCCACAGTCCCAG\|C\|CCCCCCACTCCAGTCCTTCCCCAAGGATGCAGCCTCAGCCTTCtCCACACC |
| 1:1114:4244:28468#1#F | \| \| |
| MQ=60 isize=123 C=33 | GCCCAGTCCCCACACCTACAAGGCCAGCAGATCCCTAATTCTCTCTCCAATCAAGTGCGCTCTCCCCAGCCTGTCCCTTCTCCACGGCCACAGtCCCAG\|C\|CCCCCCACtCCAgtcCtTCCCCagGaCTCCAATGCTCCCCtaCaCCtgcct |
| 1:1114:18853:77429#1#F | \| \| |
| MQ=60 isize=177 C=33 | GCCCAGTCCCCACACCTACAAGGCCAGCAGATCCCTAATTCTCTCTCCAATCAAGTGCGCTCTCCCCAGCCTGTCCCTTCTCCACGGCCACAGTCCCAG\|C\|CCCCCCACTCCAGTCCTTCCCCAAGGATGCAGCCTCAGCCTTCTCCACACC |
| 1:1115:3112:15302#1#F | \| \| |
| MQ=60 isize=266 C=37 | GCCCAGTCCCCACACCTACAAGGCCAGCAGATCCCTAATTCTCTCTCCAATCCAGTGCGCTCTCCCCAGCCTGTCCCTTCTCCACGGCCACAGTCCCAG\|C\|CCCCCCACTCCAGTCCTTCCCCAAGGATGCAGCCTCAGCCTTCTCCACACC |
| 1:1115:16969:52164#1#F | \| \| |
| MQ=60 isize=205 C=33 | GCCCAGTCCCCACACCTACAAGGCCAGCAGATCCCTAATTCTCTCTCCAATCAAGTGCGCTCTCCCCAGCCTGTCCCTTCTCCACGGCCACAGTCCCAG\|C\|CCCCCCACTCCAGtcCTTCCCCAAGGATGCAGCCTCAGCCTTCTCCACACC |
| 1:1115:10303:100545#1#F | \| \| |
| MQ=60 isize=149 C=33 | GCCCAGTCCCCACACCTACAAGGCCAGCAGATCCCTAATTCTCTCTCCAATCAAGTGCGCTCTCCCCAGCCTGTCCCTTCTCCACGGCCACAGTCCCAG\|C\|CCCCCCACTCCAGTCCTTCCCCAAGGATGCAGCCTCAGCCTTCTCCACAGG |
| 1:1116:5589:47430#1#F | \| \| |
| MQ=60 isize=170 C=33 | GCCCAGTCCCCACACCTACAAGGCCAGCAGATCCCTAATTCTCTCTCCAATCAAGTGCGCTCTCCCCAGCCTGTCCCTTCTCCACGGCCACAGTCCCAG\|C\|CCCCCCACTCCAGtCCTTCCCCAAGGATGCAGCCTCAGCCTTCTCCACACC |
| 1:1116:11704:67221#1#F | \| \| |
| MQ=60 isize=132 C=37 | GCTCAGTCCCCACACCTACAAGGCCAGCAGATCCCTAATTCTCTCTCCAATCAAGTGCGCTCTCCCCAGCCTGTCCCTTCTCCACGGCCACAGTCCCAG\|C\|CCCCCCACTCCAGTCCTTCCCCAAGGATGCAGGACTCCAATTATCGACCTT |
| 1:1116:14964:87680#1#F | \| \| |
| MQ=60 isize=408 C=37 | GCCCAGTCCCCACACCTACAAGGCCAGCAGATCCCTAATTCTCTCTCCAATCAAGTGCGCTCTCCCCAGCCTGTCCCTTCTCCACGGCCACAGTCCCAG\|C\|CCCCCCACTCCAGtCCTTCCCCAAGGATGCAGCCTCAGCCTTCTCCACACC |
| 1:1201:3099:19081#1#F | \| \| |
| MQ=60 isize=244 C=22 | GCCCAGTCCCCACACCTACAGGGCCAGCAGATCCCTAATTCTCTCTCCAATCAAGTGCGCTCTCCCCAGCCTGTCCCTTCTCCACGGCCACAGTCCCAG\|C\|cCCCCCACTCCAGTCCTTCCCCaAGGATGCAGCCTCAGCCTTCTCCaCACC |
| 1:1201:7165:28572#1#F | \| \| |
| MQ=60 isize=123 C=33 | GCCCAGTCCCCACACCTACAAGGCCAGCAGATCCCTAATTCTCTCTCCAATCAAGTGCGCTCTCCCCAGCCTGTCCCTTCTCCACGGCCACAGTCCCAG\|C\|CcCCCCaCTCCAGTCCTTCCCCAGGACTCCAATCTACATATCTCTCTgTCt |
| 1:1201:11894:31574#1#F | \|C\| |
| MQ=60 isize=208 C=37 | GCCCAGTCCCCACACCTACAAGGCCAGCAGATCCCTAATTCTCTCTCCAATCAAGTGCGCTCTCCCCAGCCTGTCCCTTCTCCACGGCCACAGTCCCAI\|C\|CCCCCCacTCCAGtCCTTCCCCAAGGATgCAgCCTCAGCCTtctccacac |
| 1:1201:13609:62892#1#F | \| \| |
| MQ=60 isize=338 C=37 | GCCCAGTCCCCACACCTACAAGGCCAGCAGATCCCTAATTCTCTCTCCAATCAAGTGCGCTCTCCCCAGCCTGTCCCTTCTCCACGGCCACAGTCCCAG\|C\|CCCCCCACTCCAGTCCTTCCCCAAGGATGCAGCCTCAGCCTTCTCCACACC |
| 1:1201:14181:89606#1#F | \| \| |
| MQ=60 isize=120 C=15 | GCCCAGTCCCCACACCTACAAGGCCAGCAGATCCCTAATTCTCTCTCCAATCAAGTGCGCTCTCCCCAGCCTGTCCCTTCTCCACGGCCACAGTCCCac\|c\|CCCCCccccccagcctttccaggctcccaacaacagggaccccctgtcctt |
| 1:1202:5960:12947#1#F | \| \| |
| MQ=60 isize=166 C=37 | GCCCAGTCCCCACACCTACAAGGCCAGCAGATCCCTAATTCTCTCTCCAATCAAGTGCGCTCTCCCCAGCCTGTCCCTTCTCCACGGCCACAGTCCCAG\|C\|CCCCCCACTCCAGTCCTTCCCCAAGGATGCAGCCTCAGCCTTCTCCACACC |
| 1:1202:7474:20391#1#F | \| \| |
| MQ=60 isize=170 C=33 | GCCCAGTCCCCACACCTACAAGGCCAGCAGATCCCTAATTCTCTCTCCAATCAAGTGCGCTCTCCCCAGCCTGTCCCTTCTCCACGGCCACAGTCCCAG\|C\|CCCCCCACTCCAGTCCTTCCCCAAGGATGCAGCCTCAGCCTTCTCCACACC |
| 1:1202:19260:40197#1#F | \| \| |
| MQ=60 isize=198 C=22 | GCCCAGTCCCCACACCTACAAGGCCAGCAGATCCCTAATTCTCTCTCCAATCAAgTGCGCTCTCCCCAGCCTGTCCCTTCTCCACGGCcACAGTCCcAg\|C\|CCCCCCACTCCAGtCCTTCCCCAAGGaTGCAGCCTCAGcCTTCTcCaCACC |
| 1:1202:11783:46982#1#F | \| \| |
| MQ=60 isize=188 C=37 | GCCCAGTCCCCACACCTACAAGGCCAGCAGATCCCTAATTCTCTCTCCAATCAAGTGCGCTCTCCCCAGCCTGTCCCTTCTCCACGGCCACAGTCCCAG\|C\|CCCCCCACTCCAGTCCTTCCCCAAGGATGCAGCCTCAGCCTTCTCCACACC |
| 1:1202:16246:61915#1#F | \| \| |
| MQ=60 isize=203 C=33 | GCCCAGTCCCCACACCTACAAGGCCAGCAGATCCCTAATTCTCTCTCCAATCAAGTGCGCTCTCCCCAGCCTGTCCCTTCTCCACGGCCACAGTCCCAG\|C\|CCCCCCACTCCAGTCCTtCCCCAAGGAtGCAGCCTCAGCCTTCTCCacaCC |
| 1:1202:18983:89374#1#F | \| \| |
| MQ=60 isize=213 C=33 | GCCCAGTCCCCACACCTACAAGGCCAGCAGATCCCTAATTCTCTCTCCAATCAAGTGCGCTCTCCCCAGCCTGTCCCTACTCCACGGCCACAGTCCCAG\|C\|CCCCCCACTCCAGTCCTTCCCCAAGGATGCAgCgTCAGCCTTCTCCACACC |
| 1:1203:5267:41633#1#F | \| \| |
| MQ=60 isize=232 C=33 | GCCCAGTCCCCACACCTACAAGGCCAGCAGATCCCTAATTCTCTCTCCAATCAAGTGCGCTCTCCCCAGCCTGTCCCTTCTCCACGGCCACAGTCCCAG\|C\|CCCCCCACTCCAGTCCTTCCCCAAGGATGCAGCCTCAGCCTTCTCCACACC |
| 1:1203:19837:75495#1#F | \| \| |
| MQ=60 isize=184 C=37 | ACCCAGTCCCCACACCTACAAGGCCAGCAGATCCCTAATTCTCTCTCCAATCAAGTGCGCTCTCCCCAGCCTGTCCCTTCTCCACGGCCACAGTCCCAG\|C\|CCCCCCACTCCAGTCCTTCCCCAAGGATGCAGCCTCAGCCTTCTCCACACC |
| 1:1204:14520:7143#1#F | \| \| |
| MQ=60 isize=207 C=33 | GCCCAGTCCCCACACCTACAAGGCCAGCAGATCCCTAATTCTCTCTCCAATCAAGtGCGCTCTCCCCAGCCTGTCCCTTCTCCACGGCCACAGTCCCAG\|C\|CCCCCcaCtCCAGtCCTTCcccaaggatgcagcctcagccttctccacacc |
| 1:1204:4920:13567#1#F | \| \| |
| MQ=60 isize=388 C=33 | GCCCAGTCCCCACACCTACAAGGCCAGCAGATCCCTAATTCTCTCTCCAATCAAGTGCGCTCTCCCCAGCCTGTCCCTTCTCCACGGCCACAGTCCCAG\|C\|CCCCCCACTCCAGTCCTTCCCCAAGGATGCAGCCTCAGCCTTCTCCACACC |
| 1:1204:6213:89174#1#F | \| \| |
| MQ=60 isize=147 C=37 | GCCCAGTCCCCACACCTACAAGGCCAGCAGATCCCTAATTCTCTCTCCAATCAAGTGCGCTCTCCCCAGCCTGTCCCTTCTCCACGGCCACAGTCCCAG\|C\|CCCCCCACTCCAGTCCTTCCCCAAGGATGCAGCCtCAGCCTTCTCCAGGAC |
| 1:1205:20553:36672#1#F | \| \| |
| MQ=60 isize=177 C=37 | GCCCAGTCCCCACACCTACAAGGCCAGCAGATCCCTAATTCTCTCTCCAATCAAGTGCGCTCTCCCCAGCCTGTCCCTTCTCCACGGCCACAGTCCCAG\|C\|CCCCCCACTCCAGTCCTTCCCCAAGGATGCAGCCTCAGCCTTCTCCACACC |
| 1:1205:10704:50798#1#F | \| \| |
| MQ=60 isize=140 C=33 | GCCCAGTCCCCACACCTACAAGGCCAGCAGATCCCTAATTCTCTCTCCAATCAAGTGCGCTCTCCCCAGCCTGTCCCTTCTCCACGGCCACAGTCCCAG\|C\|CCCCCCACTCCAGTCCTTCCCCAAGGATGCAGCCtCaGCCAGGACTCCAAT |
| 1:1205:4044:61322#1#F | \| \| |
| MQ=60 isize=140 C=33 | GCCCAGTCCCCACACCTACAAGGCCAGCAGATCCCTAATTCTCTCTCCAATCAAGTGCGCTCTCCCCAGCCTGTCCCTTCTCCACGGCCACAGtcCCAG\|C\|CCCCCCACTCCAGTCCTTCCCCAAGGATGCAGCCTCAGCCAGGACTCCAAT |
| 1:1205:13886:94297#1#F | \| \| |
| MQ=60 isize=268 C=33 | GCCCAGTCCCCACACCTACAAGGCCAGCAGATCCCTAATTCTCTCTCCAATCAAGTGCGCTCTCCCCAGCCTGTCCCTTCTCCACGGCCACAGTCCCAG\|C\|CCCCCCACTCCAGTCCTTCCCCAAGGATgCAGCCTCAGCCTTCTCCACACC |
| 1:1206:19516:16739#1#F | \| \| |
| MQ=60 isize=153 C=33 | GCCCAGCCCCCACACTTACAAGGCCAGCAGATCCCTAATTCTCTCTCCAATCAAGTGCGCTCTCCCCAGCCTGTCCCTTCTCCACGGCCACAGTCCCAG\|C\|CCCCCCACTCCAGTCCTTCCCCAAGGATGCAGCCTCAGCCTTCTCCACACC |
| 1:1206:13124:24637#1#F | \| \| |
| MQ=60 isize=211 C=22 | GCCCAGTCCCCACACCTACAAGGCCAGCAGATCCCTAATTCTCTCTCCAATCAAGTGCGCTCTCCCCAGCCTGTCCCTTCTCCACGGCCACAGTCCCAg\|C\|CCCCCCACTCCAGtCCTTCCCCAAGGATGCAGCCtCAGCCTtctccacacc |
| 1:1206:20776:38957#1#F | \| \| |
| MQ=60 isize=233 C=27 | GCCCAGTCCCCACACCTACAAGGCCAGCAGATCCCTAATTCTCTCTCCAATCAAGTGCGCTCTCCCCAGCCTGTCCCTTCTCCACGGCCACAGTCCCAG\|C\|CCCCCCACTCCAGTCCTTCCCCAAGGATGCAGCCTCAGCCTTCTCCACACC |
| 1:1206:18404:58961#1#F | \| \| |
| MQ=60 isize=139 C=33 | GCCCAGTCCCCACACCTACAAGGCCAGCAGATCCCTAATTCTCTCTCCAATCAAGTGCGCTCTCCCCAGCCTGTCCCTTCTCCACGGCCACAGTCCCAG\|C\|CCCCCCACTCCAGTCCTTCCCCAAGGATGCAGCCTCAGCAGGACTCCAATa |
| 1:1207:19260:13278#1#F | \| \| |
| MQ=60 isize=147 C=27 | GCCCAgTCCCCACACCTACAAGGCCAGCAGATCCCTAATTCTCTCTCCAATCAAGTGCGCTCTCCCCAgCCTGTCCCTTCTCCACGgcCACAGTCCCAG\|C\|cccCCCACTCCAGTCCTTCCCCAAGGATGCAGCCtCAGCCTTCTCCAGGAC |
| 1:1207:5173:42104#1#F | \| \| |
| MQ=60 isize=147 C=33 | GCCCAGTCCCCACACCTACAAGGCCAGCAGATCCCTAATTCTCTCTCCAATCAAGTGCGCTCTCCCCAGCCTGTCCCTTCTCCACGGCCACAGTCCCAG\|C\|CCCCCCACTCCAGTCCTTCCCCaAGGATGCAGCCTCAGCCTTCTCCAgGAC |
| 1:1207:19200:60627#1#F | \| \| |
| MQ=60 isize=289 C=33 | GCCCAGTCCCCACACCTACAAGGCCAGCAGATCCCTAATTCTCTCTCCAATCAAGTGCGCTCTCCCCAGCCTGTCCCTTCTCCACGGCCACAGTCCCAG\|C\|CCCCCCACTCCAGTCCTTCCCCAAGGATGCAGcCTCAGCCTTCTCCACACC |
| 1:1207:8622:85928#1#F | \| \| |
| MQ=60 isize=232 C=37 | GCCCAGTCCCCACACCTACAAGGCCAGCAGATCCCTAATTCTCTCTCCAATCAAGTGCGCTCTCCCCAGCCTGTCCCTTCTCCACGGCCACAGTCCCAG\|C\|CCCCCCACTCCAGTCCTTCCCCAAGGATGCAGCCTCAGCCTTCTCCACACC |
| 1:1208:14540:19330#1#F | \| \| |
| MQ=60 isize=203 C=33 | GCCCAGTCCCCACACCTACAAGGCCAGCAGATCCCTAATTCTCTCTCCAATCAAGTGCGCTCTCCCCAGCCTGTCCCTTCTCCACGGCCACAGTCCCAG\|C\|CCCCCCACTCCAGTCCTTCCCCAAGGATGCAGCCTCAGCCTTCTCCACACC |
| 1:1208:18681:45414#1#F | \| \| |
| MQ=60 isize=258 C=37 | GCCCAGTCCCCACACCTACAAGGCCAGCAGATCCCTAATTCTCTCTCCAATCAAGTGCGCTCTCCCCAACCTGTCCCTTCTCCACGGCCACAGTCCCAG\|C\|CCCCCCACTCCAGTCCTTCCCCAAGGATGCAGCCTCAGCCTTCTCCACACC |
| 1:1208:14155:64039#1#F | \| \| |
| MQ=60 isize=206 C=33 | GCCCAGTCCCCACACCTACAAGGCCAGCAGATCCCTAATTCTCTCTCCAATCAAGTGCGCTCTCCCCAGCCTGTCCCTTCTCCaCGGCCaCaGTCCCAG\|C\|CCCCCCaCTCCAGTCCTTCCCCAAGGATGCAGCCTCAGCCTTCtCCacccc |
| 1:1209:17199:18033#1#F | \| \| |
| MQ=60 isize=277 C=33 | GCCCAGTCCCCACACCTACAAGGCCAGCAGATCCCTAATTCTCTCTCCAATCAAGTGCGCTCTCCCCAGCCTGTCCCTTCTCCACGGCCACAGTCCCAG\|C\|CCCCCCACTCCAGTCCTTCCCCAAGGATGCAGCCTCAGCCTTCTCCACACC |
| 1:1209:6702:60916#1#F | \| \| |
| MQ=60 isize=197 C=33 | GCCCAGTCCCCACACCTACAAGGCCAGCAGATCCCTAATTCTCTCTCCAATCAAGTGCGCTCTCCCCAGCCTGTCCCTTCTCCACGGCCACAGTCCCAG\|C\|CCCCCCACTCCAGTCCTTCCCCAAGGATGCAGCCTCAGCCTTCTccACACC |
| 1:1209:9786:82295#1#F | \| \| |
| MQ=60 isize=129 C=33 | GCCCAGTCCCCACACCTACAAGGCCAGCAGATCCCTAATTCTCTCTCCAATCAAGTGCGCTCTCCCCAGCCTGTCCCTTCTCCACGGCCACAGTCCCAG\|C\|CCCCCCACTCCAGTCCTTCCCCAAGGATGAGGACTCCAATAGGAAGCGAAG |
| 1:1209:11798:93639#1#F | \| \| |
| MQ=60 isize=198 C=37 | GCCCAGTCCCCACACCTACAAGGCCAGCAGATCCCTAATTCTCTCTCCAATCAAGTGCGCTCTCCCCAGCCTGTCCCTTCTCCACGGCCACAGTCCCAG\|C\|CCCCCCACTCCAGTCCTTCCCCAAGGATGCAGCCTCAGCCTTCTCCACACC |
| 1:1210:19394:47312#1#F | \| \| |
| MQ=60 isize=0 C=6 | gCcCAGTCCCCACACCTACAAGGCCAGCAGATCCCTAaTTCTCTCTCCAATCAAGTGCGCTCTCCCCAgCCtGTCCCtTCTCCACGcCCaCAgtCCCAg\|c\|CCcCCCACtCCAgtCcTTCCCcaatgatgcagcctcaaccttctccacacc |
| 1:1210:9599:87774#1#F | \| \| |
| MQ=60 isize=139 C=37 | GCCCAGTCCCCACACCTACAAGGCCAGCAGATCCCTAATTCTCTCTCCAATCAAGTGCGCTCTCCCCAGCCTGTCCCTTCTCCACGGCCACAGTCCCAG\|C\|CCCCCCACTCCAGTCCTTCCCCAAGGATGCAGCCTCAGCAGGACTCCAATC |
| 1:1210:10360:92374#1#F | \| \| |
| MQ=60 isize=205 C=37 | GCCCAGTCCCCACACCTACAAGGCCAGCAGATCCCTAATTCTCTCTCCAATCAAGTGCGCTCTCCCCAGCCTGTCCCTTCTCCACGGCCACAGTCCCAG\|C\|CCCCCCACTCCAGTCCTTCCCCAAGGATGCAGCCTCAGCCTTCTCCACACC |
| 1:1211:20977:17310#1#F | \| \| |
| MQ=60 isize=209 C=22 | GCCcAGTCCCCACACCTACAAGGCCAGCAGATCCCTAATTcTCTCTCCAATCAAGTGCGCTCTCCCCAGCCTGTCCCTTCTCCACGGCCACAGTCCCAG\|C\|CCCCCCacTCCAGtCCTTCCCCAAGGATGCAGCCtCAGCCtTCtCCACAcc |
| 1:1211:15704:22009#1#F | \| \| |
| MQ=60 isize=251 C=33 | GCCCAGTCCCCACACCTACAAGGCCAGCAGATCCCTAATTCTCTCTCCAATCAAGTGCGCTCTCCCCAGCCTGTCCCTTCTCCACGGCCACAGTCCCAG\|C\|CCCCCCACTCCAGTCCTTCCCCAAGGATGcAGCCTCAGCCTTCTCCACACC |
| 1:1211:12484:35217#1#F | \| \| |
| MQ=60 isize=187 C=33 | GCCCAGTCCCCACACCTACAAGGCCAGCAGATCCCTAATTCTCTCTCCAATCAAGTGCGCTCTCCCCAGCCTGTCCCTTCTCCACGGCCACAGTCCCAG\|C\|CCCCCCACTCCAGtCCTTCCCCAAGGATGCAGCCTCAGCCTTCtCCacaCC |
| 1:1211:4011:72019#1#F | \| \| |
| MQ=60 isize=207 C=27 | GCCCAGTCCCCACACCTACAAGGCCAGCAGATCCCTAATTCTCTCTCCAATCAAGTGCGCTCTCCCCAGCCTGTCCCTTCTCCACgGCCACAGTCCCAG\|C\|CCCCCCACTCCAGTCCTTCCCCAAGGATGCAGCCTCAGCCTTCTCCACACC |
| 1:1212:9247:21355#1#F | \| \| |
| MQ=60 isize=175 C=33 | GCCCAGTCCCCACACCTACAAGGCCAGCAGATCCCTAATTCTCTCTCCAATCAAGTGCGCTCTCCCCAGCCTGTCCCTTCTCCACGGCCACAGTCCCAG\|C\|CCCCCCaCTCCAGTCCTTCCCCAAGGATGCAGCCTCAGCCTTCTCCACACC |
| 1:1212:10461:39792#1#F | \| \| |
| MQ=60 isize=133 C=33 | GCCCAGTCCCCACACCTACAAGGCCAGCAGATCCCTAATTCTCTCTCCAATCAAGTGCGCTCTCCCCAGCCTGTCCCTTCTCCACGGCCACAGTCCCAG\|C\|CCCCCCACTCCAGTCCTTCCCCAAGGATGCAGCAGGACTCCAATTCTTCGG |
| 1:1213:13721:35644#1#F | \| \| |
| MQ=60 isize=132 C=37 | GCCCAGTCCCCACACCTACAAGGCCAGCAGATCCCTAATTCTCTCTCCAATCAAGTGCGCTCTCCCCAGCCTGTCCCTTCTCCACGGCCACAGTCCCAG\|C\|CCCCCCACTCCAGTCCTTCCCCAAGGATGCAGAGGACTCCAATGAACTTCa |
| 1:1213:7418:63630#1#F | \| \| |
| MQ=60 isize=147 C=37 | GCCCAGTCCCCACACCTACAAGGCCAGCAGATCCCTAATTCTCTCTCCAATCAAGTGCGCTCTCCCCAGCCTGTCCCTTCTCCACGGCCACAGTCCCAG\|C\|CCCCCCACTCCAGTCCTTCCCCAAGGATGCAGCCTCAGCCTTCTCCAGGAC |
| 1:1213:10931:89776#1#F | \| \| |
| MQ=60 isize=177 C=2 | GCCCAGTCCCCAcACCTACAAGGCCAGCAGATcCCTAATTCTcTCTCCAATCAAGTGCGCTCTCCCCAgcCTGTCCCTTCTCCACGGccacAGtcccag\|c\|ccccccactccagtccttccccaaggatgcagcctcagccctctccacacc |
| 1:1214:17316:5733#1#F | \| \| |
| MQ=60 isize=162 C=33 | GCCCAGTCCCCACACCTACAAGGCCAGCAGATCCCTAATTCTCTCTCCAATCAAGTGCGCTCTCCCCAGCCTGTCCCCTCTCCACGGCCaCAGtCCCAG\|C\|CCCCCCACtCCagtCCTtCCCCAAGGttgCagcctcagccttctccacccc |
| 1:1214:18757:25245#1#F | \| \| |
| MQ=60 isize=162 C=33 | GCCCAGTCCCCACACCTACAAGGCCAGCAGATCCCTAATTCTCTCTCCAATCAAGTGCGCTCTCCCCAGCCTGTCCCTTCTCCACGGCCACAGTCCCAG\|C\|CCCCCCACtCCAGTCCTTCCCCAAGGATGCAGCCTCAGCCTTCtCCACACC |
| 1:1214:18559:41342#1#F | \| \| |
| MQ=60 isize=134 C=33 | GCCCAGTCCCCACACCTACAAGGCCAGCAGATCCCTAATTCTCTCTCCAATCAAGTGCGCTCTCCCCAGCCTGTCCCTTCTCCACGGCCACAGTCCCAG\|C\|CCCCCCACTCCAGTCCCTCCCCAAGGATGCAGCCAGGaCTCCAAtgGCGCG |
| 1:1215:8643:21495#1#F | \| \| |
| MQ=60 isize=172 C=37 | GCCCAGTCCCCACACCTACAAGGCCAGCAGATCCCTAATTCTCTCTCCAATCAAGTGCGCTCTCCCCAGCCTGTCCCTTCTCCACGGCCACAGTCCCAG\|C\|CCCCCCACTCCAGTCCTTCCCCAAggatGCAGCCTCAGCCTTCTCCACACC |
| 1:1215:19930:43615#1#F | \| \| |
| MQ=60 isize=172 C=2 | gCcCAgTCCCCACACCTACaAGGCCAGCAGATCCCTAATTCTCTCTCCAAtCAAGTGCgcTCTCCCCAgCCTGtCCCTTCTCCACGgCCACAgtCCCAc\|c\|ccccccactcccgtccttccccaaggatacaccctcacccttctccaaacc |
| 1:1215:12143:60200#1#F | \| \| |
| MQ=60 isize=139 C=22 | GCCCAGTCCCCACACCTACAAGGCCAGCAGATCCCTAATTCTCTCTCCAATCAAGTGCGCTCTCCCCAgCCtcTCCCTTCTCCACggCCACAGTCCCAG\|C\|CCCCCCaCtCCAGTCctTCCCCaAGGaTGCAccCTCAGCagGACTCCAATa |
| 1:1215:14289:77539#1#F | \| \| |
| MQ=60 isize=165 C=6 | GCCCAGTCCCCACACTTACAAGGCCAGCAGATCCCTAATTCTCTCTCCAATCAAGTGCGCTCTCCCCAGCCTGTCCCTTCTCCACGGCCACAgtcCCAG\|c\|CCCCCCAcTCCAGTCCTTCCCCaAGGATGCAGCCTCagccttctccccccc |
| 1:1215:20915:93345#1#F | \| \| |
| MQ=60 isize=134 C=37 | GCCCAGTCCCCACACCTACAAGGCCAGCAGATCCCTAATTCTCTCTCCAATCAAGtGCGCTCTCCCCAGCCTGTCCCTTCTCCACGGCCACAGTCCCAG\|C\|CCCCCCACTCCAGTCCTTCCCCAAGGATGCAGCCAGGaCtCcAATTGCGCc |
| 1:1216:9525:15996#1#F | \| \| |
| MQ=60 isize=233 C=33 | GCCCAGTCCCCACACCTACAAGGCCAGCAGATCCCTAATTCTCTCTCCAATCAAGTGCGCTCTCCCCAGCCTGTCCCTTCTCCACGGCCACAGTCCCAG\|C\|CCCCCCACTCCAGTCCTTCCCCAAGGATGCAGCCTCAGcCTTCTCCACACC |
| 1:1216:10754:52174#1#F | \| \| |
| MQ=60 isize=318 C=33 | GCCCAGTCCCCACACCTACAAGGCCAGCAGATCCCTAATTCTCTCTCCAATCAAGTGCGCTCTCCCCAGCCTGTCCCTTCTCCACGGCCACAGTCCCAG\|C\|CCCCCCACTCCAGTCCTTCCCCAAGGATGCAGCCTCAGCCTTCTCCACACC |
| 1:1216:13828:60086#1#F | \| \| |
| MQ=60 isize=293 C=33 | GCCCAGTCCCCACACCTACAAGGCCAGCAGATCCCTAATTCTCTCTCCAATCAAGTGCGCTCTCCCCAGCCTGTCCCTTCTCCACGGCCACAGTCCCAG\|C\|CCCCCCACTCCAGtCCTTCCCCAAGGATGCagCCTCAGcCTTCTCCACACC |
| 1:1216:5876:95364#1#F | \| \| |
| MQ=60 isize=140 C=33 | GCCCAGTCCCCACACCTACAAGGCCAGCAGATCCCTAATTCTCTCTCCAATCAAGTGCGCTCTCCCCAGCCTGTCCCTTCTCCACGGCCACAGtCCCAG\|C\|CCCCCCACTCCAGTCCTTCCCCAAGGATGcAGCCTCAGCCAgGACTCCAAt |
| 1:2101:9408:32369#1#F | \| \| |
| MQ=60 isize=128 C=37 | GCCCAGTCCCCACACCTACAAGGCCAGCAGATCCCTAATTCTCTCTCCAATCAAGTGCGCTCTCCCCAGCCTGTCCCTTCTCCACGGCCACAGTCCCAG\|C\|CCCCCCACTCCAGTCCTTCCCCAAGGATAGGACTCCAATGTTTAAGTCGAT |
| 1:2101:2365:81115#1#F | \| \| |
| MQ=60 isize=193 C=33 | GCCCAGTCCCCACACCTACAAGGCCAGCAGATCCCTAATTCTCTCTCCAATCAAGTGCGCTCTCCCCAGCCTGTCCCTTCTCCACGGCCACAGTCCCAG\|C\|CCCCCCACTCCAGTCCTTCCCCAAGGATGCAGCCTCAGCCTTCTCCACACC |
| 1:2101:15895:88428#1#F | \| \| |
| MQ=60 isize=344 C=33 | GCCCAGTCCCCACACCTACAAGGCCAGCAGATCCCTAATTCTCTCTCCAATCAAGTGCGCTCTCCCCAGCCTGTCCCTTCTCCACGGCCACAGTCCCAG\|C\|CCCCCCACTCCAGTCCTTCCCCAAGGATGCAGCCTCAGCCTTCTCCACACC |
| 1:2102:20543:52880#1#F | \| \| |
| MQ=60 isize=236 C=33 | GCCCAGTCCCCACACCTACAAGGCCAGCAGATCCCTAATTCTCTCTCCAATCAAGTGCGCTCTCCCCAGCCTGTCCCTTCTCCACGGCCACAGtCCCAG\|C\|CCCCCCACTCCAGTCCTTCCCCAAGGATGCAGCCTCAGCCTTCTCCACACC |
| 1:2102:12482:68206#1#F | \| \| |
| MQ=60 isize=160 C=37 | GCCCAGTCCCCACACCTACAAGGCCAGCAGATCCCTAATTCTCTCTCCAATCAAGTGCGCTCTCCCCAGCCTGTCCCTTCTCCACGGCCACAGTCCCAG\|C\|CCCCCCACTCCAGTCCTTCCCCAAGGATGCAGCCTCAGCCTTCTCCACACC |
| 1:2103:3599:9077#1#F | \| \| |
| MQ=60 isize=132 C=37 | GCCCAGTCCCCACACCTACAAGGCCAGCAGATCCCTAATTCTCTCTCCAATCAAGTGCGCTCTCCCCAGCCTGTCCCTTCTCCACGGCCACAGTCCCAG\|C\|CCCCCCACTCCAGTCCTTCCCCAAGGATGCAGGACTCCAATGCTGGCTGTC |
| 1:2103:18709:27958#1#F | \| \| |
| MQ=60 isize=125 C=33 | GCCCAGTCCCCACACCTACAAGGCCAGCAGaTCCCTAATTCTCTCTCCAATCAAGTGCGCTCTCCCCAGCCTGTCCCTTCTCCACGGCCACAGTCCCAG\|C\|CCCCCCACtCCAGtcCTTCCCCAaGagGACtccAAtACAGtcaACAagctc |
| 1:2103:16703:69540#1#F | \| \| |
| MQ=60 isize=233 C=33 | GCCCAGTCCCCACACCTACAAGGCCAGCAGATCCCTAATTCTCTCTCCAATCAAGTGCGCTCTCCCCAGCCTGTCCCTTCTCCACGGCCACAGTCCCAG\|C\|CCCCCCACTCCAGTCCTTCCCCAAGGATGCAGCCTCAGCCTTCTCCACACC |
| 1:2104:21036:20185#1#F | \| \| |
| MQ=60 isize=177 C=37 | GCCCAGTCCCCACACCTACAAGGCCAGCAGATCCCTAACTCTCTCTCCAATCAAGTGCGCTCTCCCCAGCCTGTCCCTTCTCCACGGCCACAGTCCCAG\|C\|CCCCCCACTCCAGTCCTTCCCCAAGGATGCAGCCTCAGCCTTCTCCACACC |
| 1:2104:19934:39228#1#F | \| \| |
| MQ=60 isize=208 C=33 | GCCCAGTCCCCACACCTACAAGGCCAGCAGATCCCTAATTCTCTCTCCAATCAAGTGCGCTCTCCCCAGCCTGTCCCTTCTCCACGGCCACAGTCCCAG\|C\|CCCCCCACTCCaGtCCTTCCCCAAGGATgcagCCtCAGCCTTCTCCACACC |
| 1:2104:6385:60141#1#F | \| \| |
| MQ=60 isize=236 C=33 | GCCCAGTCCCCACACCTACAAGGCCAGCAGAtCCCTAATTcTCTCTCCAATCAAGTGCGCTCTCCCCagCCTGTCCCTTCTcCACGgCCACAGtcCcAG\|C\|CCCCCCACTCCAGTCCTTCCCCAAGGAtgCAgCCTCAgcCTtCTCCACACC |
| 1:2105:11646:18839#1#F | \| \| |
| MQ=60 isize=280 C=33 | GCCCAGTCCCCACACCTACAAGGCCAGCAGATCCCTAATTCTCTCTCCAATCAAGTGCGCTCTCCCCAGCCTGTCCCTTCTCCACGGCCACAGTCCCAG\|C\|CCCCCCACTCCAGTCCTTCCCCaAGGATGCAGCCTCAGCCTTCTcCACACC |
| 1:2105:13216:28166#1#F | \| \| |
| MQ=60 isize=249 C=37 | GCCCAGTCCCCACACCTACAAGGCCAGCAGATCCCTAATTCTCTCTCCAATCAAGTGCGCTCTCCCCAGCCTGTCCCTTCTCCACGGCCACAGTCCCAG\|C\|CCCCCCACTCCAGTCCTTCCCCAAGGATGCAGCCTCAGCCTTCTCCACATC |
| 1:2105:17959:73044#1#F | \| \| |
| MQ=60 isize=268 C=37 | GCCCAGTCCCCACACCTACAAGGCCAGCAGATCCCTAATTCTCTCTCCAATCAAGTGCGCTCTCCCCAGCCTGTCCCTTCTCCACGGCCACAGTCCCAG\|C\|CCCCCCACTCCAGTCCTTCCCCAAGGATGCAGCCTCAGCCTTCTCCACACC |
| 1:2105:6418:98776#1#F | \| \| |
| MQ=60 isize=233 C=33 | GCCCAGtCCCCACACCTACAAgGCCAGCAGATCCCTAATCCTCTCTCCAATCAAGTGCGCTCTCCCCAGCCTGTCCCTTCTCCACGGCcacAGTCCCAG\|C\|CCCCCCACTCCAGTCCttCCCCAaGGATgCAGCCTCAgcCTtctCCACACC |
| 1:2106:5701:66006#1#F | \| \| |
| MQ=60 isize=139 C=37 | GCCCAGTCCCCACACCTACAAGGCCAGCAGATCCCTAATTCTCTCTCCAATCAAGTGCGCTCTCCCCAGCCTGTCCCTTCTCCACGGCCACAGTCCCAG\|C\|CCCCCCACTCCAGTCCTTCCCCAAGGATGCAGCCTCAGCAGGACTCCAATC |
| 1:2106:14436:77968#1#F | \| \| |
| MQ=60 isize=170 C=33 | GCCCAGTCCCCACACCTACAAGGCCAGCAGATCCCTAATTCTCTCTCCAATCAAGTGCGCTCTCCCCAGCCTGTCCCTTCTCCACGGCCACAGTCCCAG\|C\|CCCCCCACTCCAGTCCTTCCCCAAGGATGCAGCCTCAGCCTTCTCCACACC |
| 1:2107:17297:7043#1#F | \| \| |
| MQ=60 isize=140 C=37 | GCCCAGTCCCCACACCTACAAGGCCAGCAGATCCCTaATTCTCTCTCCAATCAAGTGCGCTCTCCCCAGCCTGTCCCTTCTCCACGGCCACAGTCCCAG\|C\|CCCCCCACTCCAGTCCTTCCCCAAGGATGCAGCCTCAGCCAGGACTCCAAt |
| 1:2107:20471:24237#1#F | \| \| |
| MQ=60 isize=177 C=33 | GCCCAGTCCCCACACCTACAAGGCCAGCAGATCCCTAATTCTCTCTCCAATCAAGTGCGCTCTCCCCAGCCTGTCCCTTCTCCACGGCCACAGTCCCAG\|C\|CcCCCCACtCCAGTCCTTCCCCAAGGATGCAgCCTCAGCCTTCTCCACACC |
| 1:2107:16871:82467#1#F | \| \| |
| MQ=60 isize=233 C=33 | GCCCAGTCCCCACACCTACAAGGCCAGCAGATCCCTAATCCTCTCTCCAATCAAGTGCGCTCTCCCCAGCCTGTCCCTTCTCCACGGCCACAGTCCCAG\|C\|CCCCCCACTCCAGTCCTTCCCCAAGGATGCAGCCTCAGCCTTCTCCACACC |
| 1:2107:3424:91715#1#F | \| \| |
| MQ=60 isize=177 C=37 | GCCCAGTCCCCACACCTACAAGGCCAGCAGATCCCTaATTCTCTCTCCAATCAAGTGCGCTCTCCCCAGCCTGTCCCTTCTCCACGGCCACAGTCCCAG\|C\|CCCCCCACTCCAGTCCTTCCCCAAGGATGCAGCCTCAGCCTTCTCCACACC |
| 1:2108:8561:3065#1#F | \| \| |
| MQ=60 isize=185 C=33 | GCCCAGTCCCCACACCTACAAGGCCAGCAGATCCCTAATTCTCTCTCCAATCAAGTGCGCTCTCCCCAGCCTGTCCCTTCTCCACGGCCACAGTCCCAG\|C\|CCCCCCACTCCAGTCCTTCCCCAAGGATGCAGCCTCAGCCTTCTCCACACC |
| 1:2108:8761:35354#1#F | \| \| |
| MQ=60 isize=213 C=33 | GCCCAGTCCCCACACCTACAAGGCCAGCAGATCCCTAATTCTCTCTCCAATCAAGTGCGCTCTCCCCAGCCTGTCCCTTCTCCACGGCCACAGTCCCAG\|C\|CCCCCCACTCCAGTCCTTCCCCAAGGATGCAGCCTCAGCCTTCTCCACACC |
| 1:2108:4505:87736#1#F | \| \| |
| MQ=60 isize=137 C=33 | GCCCAGTCCCCACACCTACAAGGCCAGCAGATCCCTAATTCTCTCTCCAATCAAGTGCGCTCTCCCCAGCTTGTCCCTTCTCCACGGCCACAGTCCCAG\|C\|CCCCCCACTCCAGTCCTTCCCCAAGGATGCAGCCTCAGGACTCCAATATGC |
| 1:2109:1277:17799#1#F | \| \| |
| MQ=60 isize=0 C=37 | GCCCAGTCCCCACACCTACAAGGCCAGCAGATCCCTAATTCTCTCTCCAATCAAGTGCGCTCTCCCCAGCCTGTCCCTTCTCCACGGCCACAGTCCCAG\|C\|CCCCCCACTCCAGTCCTTCCCCAAGGATGCAGCCTCAGCCTTCTCCAGGAC |
| 1:2109:15122:40321#1#F | \| \| |
| MQ=60 isize=169 C=33 | GCCCAGTCCCCACACCTACAAGGCCAGCAGATCCCTAATTCTCTCTCCAATCAAGTGCGCTCTCCCCAGCCTGTCCCTTCTCCACGGCCACAGTCCCAG\|C\|CCCCCCACTCCAGTCCTTCCCCAAGGATGCAGCCTCAGCCTTCTCCACACC |
| 1:2109:5304:96055#1#F | \| \| |
| MQ=60 isize=269 C=33 | GCCCAGTCCCCACACCTACAAGGCCAGCAGATCCCTAATTCTCTCTCCAATCAAGTGCGCTCTCCCCAGCCTGTCCCTTCTCCACGGCCACAGTCCCAG\|C\|CCCCCCACTCCAGTCCTTCCCCAAGGATGCAGCCTCAGCCTTCTCCACACC |
| 1:2110:13371:10110#1#F | \| \| |
| MQ=60 isize=269 C=37 | GCCCAGTCCCCACACCTACAAGGCCAGCAGATCCCTAATTCTCTCTCCAATCAAGTGCGCTCTCCCCAGCCTGTCCCTTCTCCACGGCCACAGTCCCAG\|C\|CCCCCCACTCCAGTCCTTCCCCAAGGATGCAGCCTCAGCCTTCTCCACACC |
| 1:2110:19393:48120#1#F | \| \| |
| MQ=60 isize=139 C=33 | GCCCAGTCCCCACACCTACAAGGCCAGCAGATCCCTAATTCTCTCTCCAATCAAGTGCGCTCTCCCCAGCCTGTCCCTTCTCCACGGCCACAGtCCCAG\|C\|CCCCCCACTCCAGTCCTTCCCCAAGGATGCAGCCTCAGCAGGACTCCAATC |
| 1:2110:20797:73676#1#F | \| \| |
| MQ=60 isize=276 C=33 | GCCCAGTCCCCACACCTACAAGGCCAGCAGATCCCTAATTCTCTCTCCAATCAAGTGCGCTCTCCCCAGCCTGTCCCTTCTCCACGGCCACAGTCCCAG\|C\|CCCCCCACTCCAGCCCTTCCCCAAGGATGCAGCCTCAGCCTTCTCCACACC |
| 1:2111:2936:19602#1#F | \| \| |
| MQ=60 isize=189 C=33 | GCCCAGTCCCCACACCTACAAGGCCAGCAGATCCCTAATTCTCTCTCCAATCAAGTGCGCTCTCCCCAGCCTGTCCCTTCTCCACGGCCACAGTCCCAG\|C\|CCCCCCACTCCAGTCCTTCCCCAAGGATGCAGCCTCAGCCTTCTCCACACC |
| 1:2111:17936:40889#1#F | \| \| |
| MQ=60 isize=147 C=27 | GCCCAGTCCCCACACCTACAAGGCCAGCAGATCCCTAATTCTCTCTCCAATCAAGTGCGCTCTCCCCAGCCTGTCCCTTCTCCACGGCCACAGTCCCAg\|C\|CCCCCCaCTCCAGTCCTTCCCCaaGGaTgCagCCtCAGCCTTCTCCAAGGa |
| 1:2111:8443:48443#1#F | \| \| |
| MQ=60 isize=125 C=37 | GCCCAGTCCCCACACCTACAAGGCCAGCAGATCCCTAATTCTCTCTCCAATCAAGTGCGCTCTCCCCAGCCTGTCCCTTCTCCACGGCCACAGTCCCAG\|C\|CCCCCCACgcCAGtCCTTCCCCAAGAGGACTCCAATGGCGCGTGGATGCTG |
| 1:2112:2145:2368#1#F | \| \| |
| MQ=60 isize=140 C=37 | GCCCAGTCCCCACACCTACAAGGCCAGCAGATCCCTAATTCTCTCTCCAATCAAGTGCGCTCTCCCCAGCCTGTCCCTTCTCCACGGCCACAGTCCCAG\|C\|CCCCCCACTCCAgTCCTTCCCCAAGGATGCAGCCTCAGCCAGGACTCCAAt |
| 1:2112:16227:22462#1#F | \| \| |
| MQ=60 isize=239 C=33 | GCCCAGTCCCCACACCTACAAGGCCAGCAGATCCCTAATTCTCTCTCCAATCAAGTGCGCTCTCCCCAGCCTGTCCCTTCTCCACGGCCACAGTCCCAG\|C\|CCCCCCACTCCAGTCCTTCCCCAAGGATGCAGCCTCAGCCTTCTCCACACC |
| 1:2112:4752:52767#1#F | \| \| |
| MQ=60 isize=140 C=37 | GCCCAGTCCCCACACCTACAAGGCCAGCAGATCCCTAATTCTCTCTCCAATCAAGTGCGCTCTCCCCAGCCTGTCCCTTCTCCACGGCCACAGTCCCAG\|C\|CCCCCCACTCCAGTCCTTCCCCAAGGATGCAGCCTCAGCCAGGACTCCAAT |
| 1:2112:16109:94126#1#F | \| \| |
| MQ=60 isize=261 C=33 | GCCCAGTCCCCACACCTACAAGGCCAGCAGATCCCTAATTCTCTCTCCAATCAAGTGCGCTCTCCCCAGCCTGTCCCTTCTCCACGGCCACAGTCCCAG\|C\|CCCCCCACTCCAGTCCTTCCCCAAGGATGCAGCCTCAGCCTTCTCCACACC |
| 1:2113:20818:19380#1#F | \| \| |
| MQ=60 isize=187 C=33 | GCCCAGTCCCCACACCTACAAGGCCAGCAGATCCCTAATTCTCTCTCCAATCAAGTGCGCTCTCCCCAGCCTGTCCCTTCTCCACGGCCACAGtCCCAG\|C\|CCCCCCAcTCCAGTCCTTCCCCAAGGaTGCAGCCTCAGCCTTCTCCACACC |
| 1:2113:7687:36316#1#F | \| \| |
| MQ=60 isize=131 C=33 | GCCCAGTCCCCACACCTACAAGGCCAGCAGATCCCTAATTCTCTCTCCAATCAAGTGCGCTCTCCCCAGCCTGTCCCTTCTCCACGGCCACAGTCCCAG\|C\|CCCCCCACtCCAGtCCTTCCCCAAGGATGCAAGGACTCCAAtACCTCCCtg |
| 1:2113:7767:79325#1#F | \| \| |
| MQ=60 isize=187 C=2 | GCCCAGTCCCCACACCTACAAGGCCAGCAGATCCCTAATTCTCTCTCCAATCAAGTGCGCTCTCCCCAGCCTGtccCtTCTCCACGGCCacagtcccag\|c\|cccccccccccacccctttcccaaagatgcacccccagccttctccacacc |
| 1:2114:19290:9568#1#F | \| \| |
| MQ=60 isize=258 C=37 | GCCCAGTCCCCACACCTACAAGGCCAGCAGATCCCTAATTCTCTCTCCAATCAAGTGCGCTCTCCCCAGCCTGTCCCTTCTCCACGGCCACAGTCCCAA\|C\|CCCCCCACTCCAGTCCTTCCCCAAGGATGCAGCCTCAGCCTTCTCCACACC |
| 1:2114:7386:38386#1#F | \| \| |
| MQ=60 isize=207 C=33 | GCCCAGTCCCCACACCTACAAGGCCAGCAGATCCCTAATTCTCTCTCCAATCAAGTGCGCTCTCCCCAGCCTGTCCCTTCTCCACGGCCACAGTCCCAG\|C\|CCCCCCACTCCAGTCCTTCCCCAAGGATGCAGCCTCAGCCTTCTCCACACC |
| 1:2114:10872:82725#1#F | \| \| |
| MQ=60 isize=162 C=33 | GCCCAGTCCCCACaCCTACAAGGCCAGCAGATCCCTAATTCTCTCTCCAATCAAGTGCGCTCTCCCCAGCCTGTCCCTTCTCCACGGCCACAGTCCCAG\|C\|CCCCCCACTCCAGTCCTTCCCCAAGGaTGCAGCCTCAGCCTTCTCCACACC |
| 1:2114:6243:98771#1#F | \| \| |
| MQ=60 isize=466 C=33 | GCCCAGTCCCCACACCTACAAGGCCAGCAGATCCCTAATTCTCTCTCCAATCAAGTGCGCTCTCCCCAGCCTGTCCCTTCTCCACGGCCACAGTCCCAG\|C\|CCCCCCaCTCCAGTCCTTCCCCAAGGAtGCAGCCTCAGCCTTCTCCACacC |
| 1:2115:16363:18577#1#F | \| \| |
| MQ=60 isize=149 C=37 | GCCCAGTCCCCACACCTACAAGGCCAGCAGATCCCTAATTCTCTCTCCAATCAAGTGCGCTCTCCCCAGCCTGTCCCCTCTCCACGGCCACAGTCCCAG\|C\|CCCCCCACTCCAGTCCTTCCCCAAGGATGCAGCCTCAGCCTTCTCCACAGG |
| 1:2115:6857:20729#1#F | \| \| |
| MQ=60 isize=256 C=37 | GCCCAGTCCCCACACCTACAAGGCCAGCAGATCCCTAATTCTCTCTCCAATCAAGTGCGCTCTCCCCAGCCTGTCCCTTCTCCACGGCCACAGTCCCAG\|C\|CCCCCCACTCCAGTCCTTCCCCAAGGATGCAGCCTCAGCCTTCTCCACACC |
| 1:2115:17881:61456#1#F | \| \| |
| MQ=60 isize=170 C=33 | GCCCAGTCCCCACACCTACAAGGCCAGCAGATCCCTAATTCTCTCTCCAATCAAGTGCGCTCTCCCCAGCCTGTCCCTTCTCCACGGCCACAGTTCCAG\|C\|CCCCCCACTCCAGTCCTTCCCCAAGGATGCAGCCTCAGCCTTCTCCACACC |
| 1:2115:17660:75745#1#F | \| \| |
| MQ=60 isize=174 C=33 | GCCCAGTCCCCACACCTACAAGGCCAGCAGATCCCTAATTCTCTCTCCAATCAAGTGCGCTCTCCCCAGCATGTCCCTTCTCCACGGCCACAGTCCCAG\|C\|CCCCCCACTCCAGTCCTTCCCCAAGGATGCAGCCTCAGCCTTCTCCACACC |
| 1:2115:14597:90359#1#F | \| \| |
| MQ=60 isize=277 C=33 | GCCCAGTCCCCACACCTACAAGGCCAgCAGATCCCTAATTCTCTCTCCAATCAAGTGCGCTCTCCCCAGCCTGTCCCTTCTCCACGGCCACAGTCCCAG\|C\|CCCCCCACTCCAGTCCTTCCCCAAGGAcGcAGCCTCAGCCTTCTCCACACC |
| 1:2116:18026:36685#1#F | \| \| |
| MQ=60 isize=160 C=37 | GCCCAGTCCCCACACCTACAAGGCCAGCAGATCCCTAATTCTCTCTCCAATCAAGTGCGCTCTCCCCAGCCTGTCCCTTCTCCACGGCCACAGTCCCAG\|C\|CCCCCCACTCCAGTCCTTCCCCAAGGATGCAGCCTCAGCCTTCTCCACACC |
| 1:2116:3892:69432#1#F | \| \| |
| MQ=60 isize=302 C=33 | GCCCAGTCCCCACACCTACAAGGCCAGCAGATCCCTAATTCTCTCTCCAATCAAGTGCGCTCTCCCCAGCCTGTCCCTTCTCCACGGCCACAGTCCCAG\|C\|CCCCCCACTCCAGTCCTTCCCCAaGGATGCAGCCTCAGCCTTCTCCACACC |
| 1:2116:15784:86857#1#F | \| \| |
| MQ=60 isize=131 C=33 | GCCCAGTCCCCACACCTACAAGGCCAGCAGATCCCTAATTCTCTCTCCAATCAAGTGCGCTCTCCCCAGCCTGTCCCTTCTCCACGGCCACAGtCCCAG\|C\|CCCCCCaCTCCAGTCCTTCCCCAAGGATGCAGGACTCCAATGGACGTTTAA |
| 1:2201:14032:5999#1#F | \| \| |
| MQ=60 isize=293 C=33 | GCCCAGTCCCCACACCTACAAGGCCAGCAGATCCCTAATTCTCTCTCCAATCAAGTGCGCTCTCCCCAGCCTGTCCCTTCTCCACGGCCACagtCCCAG\|C\|CCCCCCactCCAGTCCTTCCCCAaGGATGCAgCCtcAGCCTtctccacacc |
| 1:2201:17789:17314#1#F | \| \| |
| MQ=60 isize=278 C=33 | GCCCAGTCCCCACACCTACAAGGCCAGCAGATCCCTAATTCTCTCTCCAATCAAGTGCGCTCTCCCCAGCCTGTCCCTTCTCCACGGCCACAGTCCCAG\|C\|CCCCCCaCtCCAGtcCTTCCCCAAGGATGCAGCctcAgCCttctccacacc |
| 1:2201:10290:45814#1#F | \| \| |
| MQ=60 isize=177 C=33 | GCCCAGTCCCCACACCTACAAGGCCAGCAGATCCCTAATTCTCTCTCCAATCAAGTGCGCTCTCCCCAGCCTGTCCCTTCTCCACGGCCACAGTCCCAg\|C\|CCCCCCACTCCAgTCCTTCCCCAAGgaTgCagCctCAgcCTtctccccacc |
| 1:2202:10924:13514#1#F | \| \| |
| MQ=60 isize=165 C=2 | GCCCAgTCCCCACACCTACAAGGCCAGCAGaTCcctaattCTCTCTCCAAtCAAGtgCGCACTCcCcAgcctgtcccttctccacggccacagtcccag\|c\|ccccccactccaggccttccccaaggatgcagcctcagccttctccacacc |
| 1:2202:9715:34020#1#F | \| \| |
| MQ=60 isize=162 C=33 | GCCCAGT\_CCCACACCTACAAGGCCAGCAGATCCCTAATTCTCTCTCCAATCAAGTGCGCTCTCCCCAGCCTGTCCCTTCTCCACGGCCACAGTCCCAG\|C\|CCCCCCACTCCAGTCCTTCCCCAAGGATGCAGCCTCAgCCTTCTCCacACcc |
| 1:2202:15404:34450#1#F | \| \| |
| MQ=60 isize=236 C=33 | GCCCAGTCCCCACACCTACAAGGCCAGCAGATCCCTAATTCTCTCTCCAATCAAGTGCGCTCTCCCCAGCCTGTCCCTTCTCCACGGCCACAGtCCcAG\|C\|CCCCCCACTCCAgtCCTTCCCCAAGGATgcagcctcagccttctccacacc |
| 1:2202:10673:68770#1#F | \| \| |
| MQ=60 isize=138 C=33 | GCCCAGTCCCCACACCTACAAGGCCAGCAGATCCCTAATTCTCTCTCCAATCAAGTGCGCTCTCCCCAGCCTGTCCCTTCTCCACGGCCACAGTCCCAG\|C\|CCCCCCACTCCAGTCCTTCCCCAAGGATGCAGCCtCagAGGACTCCAATCC |
| 1:2202:19940:87181#1#F | \| \| |
| MQ=60 isize=133 C=33 | CCCCAGTCCCCACACCTACAAGGCCAGCAGATCCCTAATTCTCTCTCCAATCAAGTGCGCTCTCCCCAGCCTGTCCCTTCTCCACGGCgACAGTCCCAG\|C\|CCCCCCACTCCAGTCCTTCCCCAAGGATGCAGCAGGACTCCAATCTCTGCT |
| 1:2203:2076:15227#1#F | \| \| |
| MQ=60 isize=221 C=37 | GCCCAGTCCCCACACCTACAAGGCCAGCAGATCCCTAATTCTCTCTCCAATCAAGTGCGCTCTCCCCAGCCTGTCCCTTCTCCACGGCCACAGTCCCAG\|C\|CCCCCCACTCCAGTCCTTCCCCAAGGATGCAGCCTCAGCCTTCTCCACACC |
| 1:2203:11825:59476#1#F | \| \| |
| MQ=60 isize=158 C=2 | GCCCAGTCCCCACACCTACAAGGCCAGCAGATCCCTAATTCTCtCTCCAATCAAGTgcgctccccccaccctgtcccttctccccggccccagtcccag\|c\|ccccccaccccagcccttccccaacgttgcaacccaccccttccccacacc |
| 1:2203:7570:78710#1#F | \| \| |
| MQ=60 isize=228 C=27 | GCCCAGTCCCCACACCTACAAGGCCAGCAGATCCCTAATTCTCTCTCCAATCAAGTGCGCTCCCCCCAGCCTGTCCCTTCTCCACgGCCACAGTCCCAG\|C\|CCCCCCACTCCAGtcCTTCCCCAAGGaTGCAGCCTCAGCCTtCTCCaCACC |
| 1:2204:7677:53050#1#F | \| \| |
| MQ=60 isize=233 C=37 | GCCCAGTCCCCACACCTACAAGGCCAGCAGATCCCTAATTCTCTCTCCAATCAAGTGCGCTCTCCCCAGCCTGTCCCTTCTCCACGGCCACAGTCCCAG\|C\|CCCCCCACTCCAGTCCTTCCCCAAGGATGCAGCCTCAGCCTTCTCCACACC |
| 1:2204:6567:86873#1#F | \| \| |
| MQ=60 isize=132 C=37 | GCCCAGTCCCCACACCTACAAGGCCAGCAGATCCCTAATTCTCTCTCCAATCAAGTGCGCTCTCCCCAGCCTGTCCCTTCTCCACGGCCACAGTCCCAG\|C\|CCCCCCACTCCAGTCCTTCCCCAAGGATGCAGAGGACTCCAATGATACACt |
| 1:2204:18578:100026#1#F | \| \| |
| MQ=60 isize=124 C=33 | GCCCAGTCCCCACACCgaCAAGGCCAGCAGATgCCTAATTCTCTCTCCAATCAAGTGCGCTCTCCCCAGCCTGTCCCTTCTCCACGGCCACAGTCCCAG\|C\|CCCCCCACTCCAGTCCTTCCCCAAAGGACTCCAATTAATGCCCTGCCCTGt |
| 1:2205:19853:34657#1#F | \| \| |
| MQ=60 isize=230 C=33 | GCCCAGTCCCCACACCTACAAGGCCAGcAGATCCCTAATTCTCTCTCCAATCAActGCGCTCTCCCCAGCCTGTCCCTTCTCCACGGCCACAGTCCCAG\|C\|CCCCCCACTCCAGTCCTTCCCCaAGGATGCAGCCTCAGCCTTCTCctcacC |
| 1:2205:13274:41591#1#F | \| \| |
| MQ=60 isize=147 C=2 | GCcCAGTCCCCACACCTACAAGGCCAGCAGATCCCTAATTCTCTCTCCAATCAAGTGcGCTCtCCCcAGCCTGTCCCTTCTCCACGGCCACagtccaag\|c\|ccccccactccagtccttccccaaggatgcagcctcagccttctccaggac |
| 1:2205:10726:75062#1#F | \| \| |
| MQ=60 isize=131 C=37 | GCCCAGTCCCCACACCTACAAGGCCAGCAGATCCCTAATTCTCTCTCCAATCAAGTGCGCTCTCCCCAGCCTGTCCCTTCTCCACGGCCACAGTCCCAG\|C\|CCCCCCACTCCAGTCCTTCCCCAAGGATGCAGGACTCCAATatAGCACAAt |
| 1:2205:8400:89480#1#F | \| \| |
| MQ=60 isize=125 C=33 | GCCCAGTCCCCACACCTACAAGGCCAGCAGATCCCTAATTCTCTCTCCAATCAAGTGCGCTCTCCCCAGCCTGTCCCTTCTCCACGGCCACAGTCCCAG\|C\|CCCCCCACTCCAGTCCTTCCCCAAGAGGACTCCAATGCGGGTGCacGtctg |
| 1:2205:16002:96749#1#F | \| \| |
| MQ=60 isize=278 C=33 | GCCCAGTCCCCACACCTACAAGGCCAGCAGATCCCTAATTCTCTCTCCAATCAAGTGCGCTCTCCCCAGCCTGTCCCTTCTCCACGGCCACAGTCCCAG\|C\|CCCCCCACTCCAGTCCTTCCCCAAGGAggcAGCCTCAgCCTtctccacccc |
| 1:2206:14292:11144#1#F | \| \| |
| MQ=60 isize=232 C=37 | GCCCAGTCCCCACACCTACAAGGCCAGCAGATCCCTAATTCTCTCTCCAATCAAGTGCGCTCTCCCCAGCCTGTCCCTTCTCCACGGCCACAGTCCCAG\|C\|CCCCCCACTCCAGTCCTTCCCCAAGGATGCAGCCTCAGCCTTCTCCAcACC |
| 1:2206:15991:18790#1#F | \| \| |
| MQ=60 isize=207 C=33 | GCCCAGTCCCCACACCTACAAGGCCAGCAGATCCCTAATTCTCTCTCCAATCAAGTGCGCTCTCCCCAGCCTGTCCCTTCTCCACGGCCACAGTCCCAG\|C\|CcCCCCaccccagccctcccccaaggatgcacccccagctttctccacacc |
| 1:2206:13969:48222#1#F | \| \| |
| MQ=60 isize=121 C=33 | GCCCAGTCCCCACACCTACAAGGCCAGCAGATCCCTAATTCTCTCTCCAATCAAGTGCGCTCTCCCCAGCCTGTCCCTTCTCCACGGCCACAGTCCCAG\|C\|CCCCCCACTCCAGtCCTTCCCAGGACTCCAATGTTAGAACCATGCTGTCTc |
| 1:2206:11951:90268#1#F | \| \| |
| MQ=60 isize=307 C=33 | GCCCAGTCCCCACACCTACAAGGCCAGCAGATCCCTAATTCTCTCTCCAATCAAGTGCGCTCTCCCCAGCCTGTCCCTTCTCCACGGCCACAGTCCCAg\|C\|CcCCCCACTCCAGTCCTTCCCCAAGGaTGCAGCCtcAGCCTTctCCACaCC |
| 1:2207:8281:60663#1#F | \| \| |
| MQ=60 isize=155 C=33 | GCCCAGTCCCCACACCTACAAGGCCAGCAGATCCCTAATTCTCTCTCCAATCAAGTGCGCTCTCCCCAGCCTGTCCCTTCTCCACGGCCACAGTCCCAG\|C\|CCCCCCACTCCAGTCCTTCCCCAAGGATGCAGCCTCAGCCTTCTCCACACC |
| 1:2208:19102:30656#1#F | \| \| |
| MQ=60 isize=266 C=37 | GCCCAGTCCCCACACCTACAAGGCCAGCAGATCCCTAATTCTCTCTCCAATCAAGTGCGCTCTCCCCAGCCTGTCCCTTCTCCACGGCCACAGTCCCAG\|C\|CCCCCCACTCCAGTCCTTCCCCAAGGATGCAGCCTCAGCCTTCTCCACACC |
| 1:2208:4655:43528#1#F | \| \| |
| MQ=60 isize=250 C=33 | GCCCAGTCCCCACACCTACAAGGCCAGCAGATCCCTAATTCTCTCTCCAATCAAGTGCGCTCTCCCCAGCCTGTCCCTTCTCCACGGCCACAGTCCCAG\|C\|CCCCCCACTCCAGTCCTTCCCCAAGGATGCAGCCtCAGCCTTCTCCACACC |
| 1:2208:5656:73326#1#F | \| \| |
| MQ=60 isize=203 C=27 | GCCCAGTCCCCACACCTACAAGGCCAGCAGATCCCTAATTCTCTCTCCAATCAAGTGCGCTCTCCCCAGCCTGTCCCTTCTCCACGGCCACAGTCCCAG\|C\|CCCCCcaCtCCAGTCCTTCCCCAAGGATGCAGcCTCAGCCTTCTCCACACC |
| 1:2208:16160:90874#1#F | \| \| |
| MQ=60 isize=213 C=33 | GCCCAGTCCCCACACCTACAAGGCCAGCAGATCCCTAATTCTCTCTCCAATCAAGTGCGCTCTCCCCAGCCTGTCCCTTCTCCACGGCCACAGTCCCAG\|C\|CCCCCCACTCCAGTCCTTCCCCAAGGATGcAGCCtcAGCCTTCTCCACACC |
| 1:2209:8525:51070#1#F | \| \| |
| MQ=60 isize=150 C=33 | GCCCAGTCCCCACACCTACAAGGCCAGCAGATCCCTAATTCTCTCTCCAATCAAGTGCGCTCTCCCCAGCCTGTCCCTtCTCCACGGCCACAGtCCcAG\|C\|CCCCCCaCTCCAGtCCTTCCCCaAGgatccaGCCtCAGCCTTCTCCACACA |
| 1:2209:19259:80099#1#F | \| \| |
| MQ=60 isize=0 C=33 | GCCCAGTCCCCACACCTACAAGGCCAGCAGATCCCCAATTCTCTCTCCAATCAAGTGCGCTCTCCCCAGCCTGTCCCTTCTCCACGgCCACAGtcCCAG\|C\|CCCCCCACTCCAGTCCTTCCCCAAGGAtGCAGCCTCAGCCCTAaAAttttg |
| 1:2210:19124:17117#1#F | \| \| |
| MQ=60 isize=162 C=27 | gcCCAGTCcCCACACCTACAAGGCCAGCAGATCCCTAATTCTCTCTCCAATCAAGTGCgCtCTCCCCAGCCTGTCCCTTCTCCACGgCCACAGTCCCAG\|C\|CCCCCCaCTCCAGTCCTTCCCCAAGgAtgCagcCtCAGCCTTCTCCACACC |
| 1:2210:17336:66493#1#F | \| \| |
| MQ=60 isize=186 C=22 | GCCCAGTCCCCACACCTACAAGGCCAGCAGATCCCTAATTCTCTCTCCAATCAAGTGCGCTCTCCCCAGCCTGTCCCTTCTCCACGGCCACAGTCCCAG\|C\|CCCCCCACTCCAGTCCTTCCCCAAGGATGCAGCCTCAGCCTTCTccacacc |
| 1:2210:10725:80195#1#F | \| \| |
| MQ=60 isize=0 C=33 | GCCCAGTCCCCACACCTACAAGGCCAGCAGATCCCTAATTCTCTCTCCAATCAAGTGCGCTCTCCCCAGCCTGTCCCTTCTCCACGGCCACAGTCCCAG\|C\|CCCCCCACTCCAGTCCTTCCCCAAGGAtgCAGCCTCAGCCTTCTCCACACC |
| 1:2211:15018:19792#1#F | \| \| |
| MQ=60 isize=269 C=37 | GCCCAGTCCCCACACCTACAAGGCCAGCAGATCCCTAATTCTCTCTCCAATCAAGTGCGCTCTCCCCAGCCTGTCCCTTCTCCACGGCCACAGTCCCAG\|C\|CCCCCCACTCCAGTCCTTCCCCAAGGATGCAGCCtCAgCCTTCTCCACACC |
| 1:2211:10149:58383#1#F | \| \| |
| MQ=60 isize=203 C=33 | GCCCAGTCCCCACACCTACAAGGCCAGCAGATCCCTAATTCTCTCTCCAATCAAGTGCGCTCTCCCCAGCCTGTCCCTTCTCCACGGCCACAGTCCCAG\|C\|CCCCCCACTCCAGTCCTTCCCCAAGGATGCAGCCTCAGCCTTCTCCACACC |
| 1:2211:6706:82014#1#F | \| \| |
| MQ=60 isize=222 C=37 | GCCCAGTCCCCACACCTACAAGGCCAGCAGATCCCTAATTCTCTCTCCAATCAAGTGCGCTCTCCCCAGCCTGTCCCTTCTCCACGGCCACAGTCCCAG\|C\|CCCCCCACTCCAGTCCTTCCCCAAGGATGCAGCCTCAGCCTTCTCCACACC |
| 1:2212:21069:17697#1#F | \| \| |
| MQ=60 isize=221 C=33 | GCCCAGTCCCCACACCTACAAGGCCAGCAGATCCCTAATTCTCTCTCCAATCAAGTGCGCTCTCCCCAGCCTGTCCCTTCTCCACGGCCACAGTCCCAG\|C\|CCCCCCaCTCCAGTCCTTCCCCAAGGATGCAGcctCAGCCTtCTCCACACC |
| 1:2212:19056:22205#1#F | \| \| |
| MQ=60 isize=245 C=27 | GCCCAGTCCCCACACCTACAAGGCCAGCAGATCCCTAATTCTCCCTCCAATCAAGTGCGCTCTCCCCAGCCTGTCCCTTCTCCACGGCCACAGtCCCag\|C\|CCCCCCACtCCAgtcCTtCCCCAAGGaTGCagCctcAgcCTTctccacacc |
| 1:2212:9125:31349#1#F | \| \| |
| MQ=60 isize=199 C=33 | GCCCAGTCCCCACACCTACAAGGCCAGCAGATCCCTAATTCTCTCTCCAATCAAGTGCGCTCTCCCCAGCCTGTCCCTTCTCCACGGCCACAGtCCCAG\|C\|CCCCCCACTCCAGTCCTTCCCCAAGGATGCAGcCTcaGCCTTCTCcACACC |
| 1:2212:7911:41090#1#F | \| \| |
| MQ=60 isize=156 C=33 | GCCCAGTCCCCACACCTACAAGGCCAGCAGATCCCTAATTCTCTCTCCAATCAAGTGCGCTCTCCCCAGCCTGTCCCTTCTCCACGGCCACAGTCCCAG\|C\|CCCCCCACTCCAGTCCTTCCCCAAGGATGCAGCCTCAGCCTTCTCCACACC |
| 1:2212:13887:69972#1#F | \| \| |
| MQ=60 isize=190 C=33 | GCCCAGTCCCCACACCTACAAGGCCAGCAGATCCCTAATTCTCTCTCCAATCAAGTGCGCTCTCCCCAGCCTGTCCCTTCTCCACGGCCACAGtCCCAG\|C\|CCCCCCaCTCCAGtCCTTCCCCAAGGATGCAgCCtCAgCCTTctcCACACC |
| 1:2212:19910:77257#1#F | \| \| |
| MQ=60 isize=302 C=33 | GCCCAGTCCCCACACCTACAAGGCCAGCAGATCCCTAATTCTCTCTCCAATCAAGTGCGCTCTCCCCAGCCTGTCCCTTCTCCACGGCCACAGTCCCAG\|C\|CCCCCCACTCCAGTCCTTCCCCAAGGATGCAGCCTCAGCCTTCtCCACACC |
| 1:2212:11357:98700#1#F | \| \| |
| MQ=60 isize=144 C=22 | GCCCAGTCCCCACACCTACAAGGCCAGCAGATCCCTAATTCTCTCTCCAATCAAGTGCGCTCTCCCCAGCCTGTCCCTTCTCCACGGCCACAGTCCCAg\|C\|CcCCCCACTCCaGTCCTTcCCCAAGGATGCAGCCTCAGCCTtCTAGGACTC |
| 1:2213:11032:53662#1#F | \| \| |
| MQ=60 isize=217 C=33 | GCCCAGTCCCCACACCTACAAGGCCAGCAGATCCCTAATTCTCTCTCCAATCAAGTGCGCTCTCCCCAGCCTGTCCCTTCTCCACGGCCACAGTCCCAG\|C\|CCCCCCACTCCAGTCCTTCCCCAAGGATGCAGCCTCAGCCTTCTCCACACC |
| 1:2213:15594:81260#1#F | \| \| |
| MQ=60 isize=186 C=33 | GCCcAGTCCCCACACCTACAAGGCCAGCAGATCCCTAATTCTCTCTCCAATCAAGTGCGCTCTCCCCAGCCTGTCCCTTCTCCACGgcCACAGtccCAG\|C\|CCCCCCActCCAGTCCTTCCCCAAGgATGCaGCCtcagcCTTCCCCACACC |
| 1:2213:12607:100374#1#F | \| \| |
| MQ=60 isize=192 C=37 | GCCCAGTCCCCACACCTACAAGGCCAGCAGATCCCTAATTCTCTCTCCAATCAAGTGCGCTCTCCCCAGCCTGTCCCTTCTCCACGGCCACAGTCCCAG\|C\|CCCCCCACTCCAGTCCTTCCCCAAGGATGCAGCCTCAGCCTTCTCCaCACC |
| 1:2214:14025:33487#1#F | \| \| |
| MQ=60 isize=235 C=37 | GCCCAGTCCCCACACCTACAAGGCCAGCAGATCCCTAATTCTCTCTCCAATCAAGTGCGCTCTCCCCAGCCTGTCCCTTCTCCACGGCCACAGTCCCAG\|C\|CCCCCCACTCCAGTCCTTCCCCAAGGATGCAGCCTCAGCCTTCTCCACACC |
| 1:2214:13290:70750#1#F | \| \| |
| MQ=60 isize=192 C=37 | GCCCAGTCCCCACACCTACAAGGCCAGCAGATCCCTAATTCTCTCTCCAATCAAGTGCGCTCTCCCCAGCCTGTCCCTTCTCCACGGCCACAGTCCCAG\|C\|CCCCCCACTCCAGTCCTTCCCCAAGGATGCAGCCTCAGCCTTCTCCACACC |
| 1:2215:8349:16882#1#F | \| \| |
| MQ=60 isize=139 C=37 | GCCCAGTCCCCACACCTACAAGGCCAGCAGATCCCTAATTCTCTCTCCAATCAAGTGCGCTCTCCCCAGCCTGTCCCTTCTCCACGGCCACAGTCCCAG\|C\|CCCCCCACTCCAGTCCTTCCCCAAGGATGCAGCCTCAGCAGGACTCCAATA |
| 1:2215:11623:40497#1#F | \| \| |
| MQ=60 isize=173 C=33 | GCCCAGTCCCCACACCTACAAGGCCAGCAGATCCCTAATTCTCTCTCCAATCAAGTGCGCTCTCCCCAGCCTGTCCCTTCTCCACGGCCACAGTCCCAG\|C\|CCCCCCACTCCaGTCCtTCCCCAAGGATGCAGCCTCAGCCTTCTCCACACC |
| 1:2215:6743:53160#1#F | \| \| |
| MQ=60 isize=231 C=33 | GCCCAGTCCCCACACCTACAAGGCCAGCAGATCCCTAATTCTCTCTCCAATCAAGTGCGCTCTCCCCAGCCTGTCCCTTCTCCACGGCCACAGTCCCAG\|C\|CCCCCCACTCCAGTCCTTCCCCAAGGATGCAGCCTCAGCCTTCTCCACACC |
| 1:2215:1818:63964#1#F | \| \| |
| MQ=60 isize=258 C=33 | GCCCAGTCCCCACACCTACAAGGCCAGCAGATCCCTAATTCTCTCTCCAATCAAGTGCGCTCTCCCCAGCCTGTCCCTTCTCCACGGCCACAGTCCCAG\|C\|CCCCCCACTCCAGTcCTTCCCCaAGGATGCAGCCTCAGCCTTCTCCACACC |
| 1:2215:16111:67955#1#F | \| \| |
| MQ=60 isize=150 C=33 | GCCCAGTCCCCACACCTACAAGGCCAGCAGATCCCTAATTCTCTCTCCAATCAAGTGCGCTCTCCCCAGCCTGtCCCTTCTCCACGGCCACCGtcCCAG\|C\|CCCCCCACtCcaGtCCTtCCCCAaGGAtgCAGCCtcagCCttctccacaca |
| 1:2216:4954:11018#1#F | \| \| |
| MQ=60 isize=125 C=33 | GCCCAGTCCCCACACCTACAAGGCCAGCAGATCCCTAATTCTCTCTCCAATCAAGTGCGCTCTCCCCAGCCTGTCCCTTCTCCACGGCCACAGTCCCAG\|C\|CCCCCCACTCCAGTCCTTCCCCAagAGGACTCCAATGTGAAAAACTCGCTG |
| 1:2216:15935:21450#1#F | \| \| |
| MQ=60 isize=162 C=33 | GCCCAGTCCCCACACCTACAAGGCCAGCAGATCCCTAATTCTCTCTCCAATCAAGTGCGCTCTCCCCAGCCTGTCCCTTCTCCACGGCCACAGTCCCAG\|C\|CCCCCCACTCCaGTCCTTCCCCAAGGATGCAGCCtcagCCTTCtCCACACC |
| 1:2216:4612:47327#1#F | \| \| |
| MQ=60 isize=0 C=33 | GCCCAGTCCCCACACCTACAAGGCCAGCAGATCCCTAATTCTCTCTCCAATCAAGTGCGCTCTCCCCAGCCTGTCCCTTCtcCACGGCCACAGtCCCAG\|C\|CCCCCCACTCCAGTCCTtcCCCAaGGATGCAGCCTCAGCCTTCTCCACCAG |
| 1:2216:11366:71888#1#F | \| \| |
| MQ=60 isize=229 C=33 | GCCCAGTCCCCACACCTACAAGGCCAGCAGATCCCTAATTCTCTCTCCAATCAAGTGCGCTCTCCCCAGCCTGTCCCTCCTCCACGGCCACAGTCCCAG\|C\|CCCCCCACTCCAGTCCTTCCCCAAGGATGCAGCCTCAGCCTTCTCCACACC |
| 1:2216:2137:89013#1#F | \| \| |
| MQ=60 isize=169 C=33 | GCCCAGTCCCCACACCTACAAGGCCAGCAGATCCCTAATTCTCTCTCCAATCAAGTGCGCTCTCCCCAGCCTGTCCCTTCTCCACGGCCACAGTCCCAG\|C\|CCCCCCACTCCAGTCCTTCCCCaagGATGCAGCCTCAGCCTTCTCCACACC |
| 2:1101:6582:2630#1#F | \| \| |
| MQ=60 isize=153 C=37 | GCCCAGTCCCCACACCTACAAGGCCAGCAGATCCCTAATTCTCTCTCCAATCAAGTGCGCTCTCCCCAGCCTGTCCCTTCTCCACGGCCAcAGTCCCAG\|C\|CCCCCCACTCCAGtCCTTCCCcaAGGAtgCagCCtcAgCcTTCtCCACACC |
| 2:1101:7586:52466#1#F | \| \| |
| MQ=60 isize=138 C=33 | GCCCAGTCCCCACACCTACAAGGCCAGCAGATCCCTAATTCTCTCTCCAATCAAGTGCGCTCTCCCCAGCCTGTCCCTTCTCCACGGCCACAGTCCCAG\|C\|CCCCCCACTCCAGTCCTTCCCCAAGGatgcagcCtCAGGaCTCCAAtcgCC |
| 2:1101:2125:59581#1#F | \| \| |
| MQ=60 isize=177 C=33 | GCCCAGTCCCCACACCTACAAGGCCAGCAGATCCCTAATTCTCTCTCCAATCAAGTGcgCTCTCCCCAGCCTGTCCCTTCTCCccGgCCacAGtCCCAG\|C\|CcCCCcaccccagtccttccccaaggatgcagccccgccctcctccacccc |
| 2:1101:13566:99885#1#F | \| \| |
| MQ=60 isize=174 C=37 | GCCCAGTCCCCACACCTACAAGGCCAGCAGATCCCTAATTCTCTCTCCAATCAAGTGCGCTCTCCCCAGCCTGTCCCTTCTCCACGGCCACAGTCCCAG\|C\|CCCCCCACTCCAGTCCTTCCCCAAGGATGCAGCCTCAGCCTTCTCCACACc |
| 2:1102:13522:26533#1#F | \| \| |
| MQ=60 isize=260 C=37 | GCCCAGTCCCCACACCTACAAGGCCAGCAGATCCCTAATTCTCTCTCCAATCAAGTGCGCTCTCCCCAGCCTGTCCCTTCTCCACGGCCACAGTCCCAG\|C\|CCCCCCACTCCAGTCCTTCCCCAAGGATGCAGCCTCAGCCTTCTCCACACC |
| 2:1102:1857:38785#1#F | \| \| |
| MQ=60 isize=229 C=22 | GCCCAGTCCCCACACCTACAAGGCCAGCAGATCCCTAATTCTCTCTCCAATCAAGTGCGCTCTCCCCAGCCTGTCCCTTCTCCacGgCCaCagtCCCag\|C\|CCCCCCactCCggtcctTCCCCAaggtgccgcccccgcccttctccacacc |
| 2:1102:14955:49267#1#F | \| \| |
| MQ=60 isize=134 C=37 | GCCCAGTCCCCACACCTACAAGGCCAGCAGATCCCTAATTCTCTCTCCAATCAAGTGCGCTCTCCCCAGCCTGTCCCTTCTCCACGGCCACAGTCCCAG\|C\|CCCCCCACTCCAGTCCTTCCCCAAGGATGCAGCCAGGACTCCAATCCGAGA |
| 2:1102:10424:51241#1#F | \| \| |
| MQ=60 isize=237 C=33 | GCCCAGTCCCCACACCTACAAGGCCAGCAGATCCCTAATTCTCTCTCCAATCAAGTGCGCTCTCCCCAGCCTGTCCCTTCTCCACGGCCACAGTCCCAG\|C\|CCCCCCACTCCagTCCTTCCCCAAGGATGCAGCCtCAGCCTTCTCCACACC |
| 2:1103:14677:6090#1#F | \| \| |
| MQ=60 isize=198 C=15 | GCCCAGTCCCCACACCTACAAGGCCAGCAGATCCCTAATTCTCTCTCCAATCAAGTGCGCTCTCCCCAGCCTGTCCCTTCTCCACGgccACAGtCCCAG\|c\|CCCCcCACTCCAGTCCTTcCCCAAgGATGCAGcCTCAGCCctCTCCaCAcC |
| 2:1103:5327:73546#1#F | \| \| |
| MQ=60 isize=161 C=33 | GCCCAGTCCCCACACCTACAAGGCCAGCAGATCCCTAATTCTCTCTCCAATCAAGTGCGCTCTCCCCAGCCTGTCCCTTCTCCACGGCCACAGTCCCAG\|C\|CCCCCCACtCCAGtCCTTCCCCAAGGAtgCAgCCtcagCCttctccaaccc |
| 2:1103:17518:88448#1#F | \| \| |
| MQ=60 isize=175 C=33 | GCCCAGTCCCCACACCTACAAGGCCAGCAGATCCCTAATTCTCTCTCCAATCAAGTGCGCTCTCCCCAGCCTGTCCCTTCTCCACGGCCACAGTCCCAG\|C\|CCCCCCACTCCAGTCCTTCCCCAAgGATGCAGCCTCAGCCTTCTCCACACC |
| 2:1104:13834:21084#1#F | \| \| |
| MQ=60 isize=252 C=37 | GCCCAGTCCCCACACCTACAAGGCCAGCAGATCCCTAATTCTCTCTCCAATCAAGTGCGCTCTCCCCAGCCTGTCCCTTCTCCACGGCCACAGTCCCAG\|C\|CCCCCCACTCCAGTCCTTCCCCAAGGATGCAGCCtCAGCCTTCTCcACACC |
| 2:1104:12457:39879#1#F | \| \| |
| MQ=60 isize=351 C=37 | GCCCAGTCCCCACACCTACAAGGCCAGCAGATCCCTAATTCTCTCTCCAATCAAGTGCGCTCTCCCCAGCCTGTCCCTTCTCCACGGCCACAGTCCCAG\|C\|CCCCCCACTCCaGTCCTTCCCCAAGGATGCAGCCTCAGCCTTCTCCACACC |
| 2:1104:14439:55584#1#F | \| \| |
| MQ=60 isize=190 C=27 | GCCCAGTCCCCACACCTACAAGGCCAGCAGATCCCTAATTCTCTCTCCAATCAAGTGCGCTCTCCCCAGCCTGTCCCTTCTCCACGGCCACAGTCCCAG\|C\|CCCCCCACTCCAGTCCTTCCCCAAGGATGCAGCCtcAGCCTTCTCCAcccc |
| 2:1104:17382:61201#1#F | \| \| |
| MQ=60 isize=288 C=22 | GCCCAGTCCCCACACCTACAAGGCCAGCAGATCCCTAATTCTCTCTCCAATCAAGTGCGCTCTCCCCAGCCTGTCCCTTCTCCACGGCCACAGTCCCAG\|C\|CcCCCCACTCCAGTCCTTCCCCAAGGATGCAGCCTCAGCCTTttcCACacC |
| 2:1104:13082:82825#1#F | \| \| |
| MQ=60 isize=227 C=37 | GCCCAGTCCCCACACCTACAAGGCCAGCAGATCCCTAATTCTCTCTCCAATCAAGTGCGCTCTCCCCAGCCTGTCCCTTCTCCACGGCCACAGTCCCAG\|C\|CCCCCCACTCCAGTCCTTCCCCAAGGATGCAGCCTCAGCCTTCTCCACACC |
| 2:1104:21059:95595#1#F | \| \| |
| MQ=60 isize=140 C=33 | GCCCAGTCCCCACACCTACAAGGCCAGCAGATCCCTAATTCTCTCTCCAATCAAGTGCGCTCTCCCCAGCCTGTCCCTTCTCCACGGCCACAGTCCCAG\|C\|CCCCCCACTCCAGTCCTTCCCCAAGGATGCAGCCTCAGCCAGGACTccAAT |
| 2:1105:17963:14165#1#F | \| \| |
| MQ=60 isize=203 C=33 | GCCCAGTCCCCACACCTACAAGGCCAGCAGATCCCTAATTCTCTCTCCAATCAAGTGCGCTCTCCCCAGCCTGTCCCTTCTCCACGGCCACAGTCCCAG\|C\|CCCCCCACTCCAGTCCTTCCCCAAGGATGCAGCCTCAGCCTTCTCCACACC |
| 2:1105:14388:54255#1#F | \| \| |
| MQ=60 isize=138 C=33 | GCCCAGTCCCCACACCTACAAGGCCAGCAGATCCCTAATTCTCTCTTCAATCAAGTGCGCTCTCCCCAGCCTGTCCCTTCTCCACGGCCACAGTCCCAG\|C\|CCCCCCACTCCAGTCCTTCCCCaAGGATGCAGCCTCAGAGGACTCCaATCT |
| 2:1105:10671:65346#1#F | \| \| |
| MQ=60 isize=232 C=37 | GCCCAGTCCCCACACCTACAAGGCCAGCAGATCCCTAATTCTCTCTCCAATCAAGTGCGCTCTCCCCAGCCTGTCCCTTCTCCACGGCCACAGTACCAG\|C\|CCCCCCACTCCAGTCCTTCCCCAAGGATGCAGCCTCAGCCTTCTCCACaCC |
| 2:1105:6866:88820#1#F | \| \| |
| MQ=60 isize=177 C=22 | GCCCAGTCCCCACACCTACAAGGCCAGCAGATCCCTAATTCTCTCTCCAATCAAGTGCGCTCTCCCCAGCCTGTCCCTTCTCCACGGCCACAGtCCcAG\|C\|CCCCCCACTCCAGtCCTTCCCCAAGGaTgCAGCCTCAgCCTTCTCCAcacc |
| 2:1106:17904:17444#1#F | \| \| |
| MQ=60 isize=192 C=33 | GCCCAGTCCCCACACCTACAAGGCCAGCAGATCCCTAATTCTCTCTCCAATCAAGTGCGCTCTCCCCAGCCTGTCCcTTCTCCACGGCCACAGTCCCAG\|C\|CCCCCCACTCCAGTCCTTCCCCAaGGATGCAGCCTCAGcCTTCTCCACACC |
| 2:1106:13577:40760#1#F | \| \| |
| MQ=60 isize=232 C=33 | GCCCAGTCCCCACACCTACAAGGCCAGCAGATCCCTAATTCTCTCTCCAATCAAGTGCGCTCTCCCCGGCCTGTCCCTTCTCCACGGCCACAGTCCCAG\|C\|CCCCCCACTCCAGTCCTTCCCCAAGGATGCAGCCTCAGCCTTCTCCACACC |
| 2:1106:21271:80357#1#F | \| \| |
| MQ=60 isize=123 C=22 | GCCCAGTCCCCACACCTACAAGGCCAGCAGATCCCTAATTCTCTCTCCAATCAAGTGCGCTCTCCCCAGCCTGTCCCTTCTCCACGGCCACAGTCCCAG\|C\|CCcCCCACTCCAGTCCTTCCCCAGGACTCCaATAAACGACACTTACTGTCT |
| 2:1106:14744:93067#1#F | \| \| |
| MQ=60 isize=147 C=33 | GCCCAGTCCCCACACCTACAAGGCCAGCAGATCCCTAATTCTCTCTCCAATCAAGTGCGCTCTCCCCAGCCTGTCCCTTCTCCACGGCCACAGTCCCAG\|C\|CCCCCCACTCCAGTCCTTCCCCAAGGATGCAGCCTCAGCCTTCTCCAGGAC |
| 2:1107:7563:31730#1#F | \| \| |
| MQ=60 isize=170 C=33 | GCCCAGTCCCCACACCTACAAGGCCAGCAGATCCCTAATTCTCTCTCCAATCAAGTGCGCTCTCCCCAGCCTGTCCCTTCTCCACGGCCACAGTCCCAG\|C\|CACCCCACTCCAGTCCTTCCCCAAGGaTGCAGCCTCAGCCTTCTCCacaCC |
| 2:1107:10358:60315#1#F | \| \| |
| MQ=60 isize=200 C=6 | GCCCAGtCCCCACACCTACAaGGCCAgCAGATCCCTAATTCTCTCTCCAATCAAGTGCGCTCTCCCCAGCCTGTCcCTTCTCCACGgCcaCAgTCCCAg\|c\|cccCCCACTCCagTcCTTCccCAAggATACAGCctcagccttttccacacc |
| 2:1107:19518:74391#1#F | \| \| |
| MQ=60 isize=175 C=22 | GCCCAGTCCCCACACCTACAAGGCCAGCAGATCCCTAATTCTCTCTCCAATCAAGTGTGCTCTCCCCAGCCTGTCCCTTCTCCACGGCCACAGTCCCAG\|C\|CcCCCCACTCCcgtCctTcCCCAAGGAagCagCCTCAgCCtTCTcCaCaCC |
| 2:1107:10084:93634#1#F | \| \| |
| MQ=60 isize=175 C=33 | GCCCAGTCCCCACACCTACAAGGCCAGCAGATCCCTAATTCTCTCTCCAATCAAGTGCGCTCTCCCCAGCCTGTCCCTTCTCCACGGCCACAGtCCCAG\|C\|CCCCCCACTCCAGTCCTTCCCCAAGGATGCAGCCTCAGCCTTCTCCACACC |
| 2:1108:14068:2864#1#F | \| \| |
| MQ=60 isize=147 C=33 | GCCCAGTCCCCACACCTACAAGGCCAGCAGATCCCTAATTCTCTCTCCAATCAAGTGCGCTCTCCCCAGCCTGTCCCTTCTCCACGGCCaCAGTCCCAG\|C\|CcCCCCACTCCAGTCCTTCCCCAAGGATGCAGCCTCAGCCTTCTCCaGAAC |
| 2:1108:5994:26459#1#F | \| \| |
| MQ=60 isize=208 C=22 | GCCCAGTCCCCACACCTACAAGGCCAGCAGATCCCTAATTCTCTCTCCAATCAAGTGCGCTCTCCCCAGCCTGTCCCTTCTCCACGGCCaCaGTCCCag\|C\|CCCCCCACTCCAGTCCTtCCCCAAGGAtgCAgCCtcAgCCTtCTccccacc |
| 2:1108:7726:42211#1#F | \| \| |
| MQ=60 isize=245 C=33 | GCCcAGTCCCCACACCTACAAGGCCAGCAGATCCCTAATTCTCTCTCCAATCAAGTGCGCTCTCCCCAGCCTGTCCCTTCTCCACGGCCACAGTCCCAG\|C\|CCCCCCACTCCAGTCCTTCCCCAAGGATGCAGCCTcagCCTTCTCCacaCC |
| 2:1108:8231:99252#1#F | \| \| |
| MQ=60 isize=169 C=33 | GCCCAGTCCCCACACCTACAAGGCCAGCAGATCCCTAATTCTCTCTCCAATCAAGTGCGCTCTCCCCAGCCTGTCCCTTCTCCACGGCCaCAGtcCCAG\|C\|CCCCCcactcCaGcCCTTCCCCAaGGaTGCAGCCtCAgCCtTCTCCACaCC |
| 2:1109:17221:31295#1#F | \| \| |
| MQ=60 isize=132 C=37 | GCCCAGTCCCCACACCTACAAGGCCAGCAGATCCCTAATTCTCTCTCCAATCAAGTGCGCTCTCCCCAGCCTGTCCCTTCTCCACGGCCACAGTCCCAG\|C\|CCCCCCACTCCAGTCCTTCCCCAAGGATGCAGAGGACTCCAatAGATtCGt |
| 2:1109:9491:37962#1#F | \| \| |
| MQ=60 isize=164 C=33 | GCCCAGTCCCCACACCTACAAGGCCAGCAGATCCCTAATTCTCTCTCCAATCAAGTGCGCTCTCCCCAGCCTGTCCCTTCTCCACGGCCACAGTCCCAG\|C\|CCCCCCACTCCAGTCCTTCCCCAAGGATGCAGCCTCAGCCTTCTCCACACC |
| 2:1109:8225:46314#1#F | \| \| |
| MQ=60 isize=256 C=33 | GCCCAGTCCCCACACCTACAAGGCCAGCAGATCCCTAATTCTCTCTCCAATCAAGTGCGCTCTCCCCAGCCTGTCCCTTCTCCACGGCCACAGTCCCAG\|C\|CCCCCCACTCCAGTCCTTCCCCAAGGAggCagCCTCAGCCTTCTCCACACC |
| 2:1109:7041:47650#1#F | \| \| |
| MQ=60 isize=149 C=33 | GCCCAGTCCCCACACCTACAAGGCCAGCAGATCCCTAATTCTCTCTCCAATCAAGTGCGCTCTCCCCAGCCTGTCCCTTCTCCACGGCCACAGTCCCAg\|C\|CCCCCCACTCCAGTCCTTCCCCAAGGATGCAGCCTcaGCCTtCtCCACAAG |
| 2:1109:3717:64359#1#F | \| \| |
| MQ=60 isize=132 C=33 | GCCCAGTCCCCACACCTACAAGGCCAGCAGATCCCTAATTCTCTCTCCAATCAAGTGCGCTCTCCCCAGCCTGTCCCTTCTCCACGGCCACAGtCCCAG\|C\|CCCCCCactCCAGtCCTTCCCCAAGGatgcAGAGGaCtCCAATTAgaCCTc |
| 2:1109:12625:81855#1#F | \| \| |
| MQ=60 isize=147 C=33 | GCCCAGTCCCCACACCTACAAGGCCAGCAGATCCCTAATTCTCTCTCCAATCAAGTGCGCTCTCCCCAGCCTGTCCCTTCTCCACGGCCACAGTCCCAG\|C\|CCCCCCACTCCAGTCCTTCCCCAAGGATGCAGCCTCAGCCTTCTCCAGGaC |
| 2:1109:5196:88572#1#F | \| \| |
| MQ=60 isize=166 C=37 | GCCCAGTCCCCACACCTACAAGGCCAGCAGATCCCTAATTCTCTCTCCAATCAAGTGCGCTCTCCCCAGCCTGTCCCTTCTCCACGGCCACAGTCCCAG\|C\|CCCCCCACTCCAGtCCTTCCCCAAGGATGCAGCCtCAgCCTTctCCACacc |
| 2:1110:17771:14057#1#F | \| \| |
| MQ=60 isize=125 C=33 | GCCCAGTCCCCACACCTACAAGGCCAGCAGATCCCTAATTCTCTCTCCAATCAAGTGCGCTCTCCCCAGCCTGTCCCTTCTCCACGGCCACAGTCCCAG\|C\|CCCCCCACTCCAGTCCTTCCCCAAGAGGACTCCAATCTGAAATGTAAaCTg |
| 2:1110:5130:21864#1#F | \| \| |
| MQ=60 isize=212 C=22 | GcCCAGTCCCCACACCTACAAGGCCAGCAGATCCCTAATTCTCTCTCCAATCAAGTGCGCTCTCCCCAGCCTGTCCCTTCTCCACGGCcacAGTCCCAg\|C\|CCCCCCACTCCagtCCTTCCCcAAGgaTgcaGCCtcAgcCtTCtCCaCaCC |
| 2:1110:13512:66481#1#F | \| \| |
| MQ=60 isize=0 C=33 | GCCCAGTCCCCACACCTACAAGGCCAGCAGATCCCTAATTCTCTCTCCAATCAAGTGCGCTCTCCCCAGCCTGTCCCTTCTCCACGGCCACAGTCCCAG\|C\|CCCCCCACTCCAGTCCTTCCCCAAGGATGCAGCCTCAgCCTTCTCCACACC |
| 2:1110:10643:97536#1#F | \| \| |
| MQ=60 isize=175 C=33 | GCCCAGTCCCCACACCTACAAGGCCAGCAGATCCCTAATTCTCTCTCCAATCAAGTGCGCTCTCCCCAGCCTGTCCCTCCTCCACGGCCACAGTCCAAG\|C\|CCCCCCACTCCAGTCCTTCCCCAAGGATGCAGCCTCAGCCTTCTCCACACC |
| 2:1111:6217:54415#1#F | \| \| |
| MQ=60 isize=221 C=33 | GCCCAGTCCCCACACCTACAAGGCCAGCAGATCCCTAATTCTCTCTCCAATCAAGTGCGCTCTCCCCAGCCTGTCCCTTCTCCACGGccACAGtCCCAG\|C\|CCCCCCACTCCAGTCCCTCCCCAAGgATGCAGCCtCAGCCTTCTCCaCaCC |
| 2:1112:10744:13835#1#F | \| \| |
| MQ=60 isize=140 C=33 | GCCCAGTCCCCACACCTACAAGGCCAGCAGATCCCTAATTCTCTCTCCAATCAAGTGCGCTCTCCCCAGCCTGTCCCTTCTCCACGGCCACAGTCCCAG\|C\|CCCCCCACTCCAGTCCTTCCCCAAGGATGCAGCCTCAGCCAGGACTCCAAt |
| 2:1112:12335:83569#1#F | \| \| |
| MQ=60 isize=143 C=33 | GCCCAGTCCCCACACCTACAAGGCCAGCAGATCCCTAATTCTCTCTCCAATCAAGTGCGCTCTCCCCAGCCTGTCCCTTCTCCACGGCCACAGTCCCAG\|C\|CCCCCCACTCCAGTCCTTCCCCAAGGATGCAGCCTCAGCCTTCAGGACTCC |
| 2:1112:2895:94502#1#F | \| \| |
| MQ=60 isize=233 C=33 | GCCCAGTCCCCACACCTACAAGGCCAGCAGATCCCTAATTCTCTCTCCAATCAAGTGCGCTCTCCCCAGCCTGTCCCTTCTCCACGGCCACAGTCCCAG\|C\|CCCCCCaCtCCAGtCCTTCCCCAAGGAtgCAgCCcCaGCCTtctccacacc |
| 2:1113:13340:8689#1#F | \| \| |
| MQ=60 isize=229 C=33 | GCCCAGTCCCCACACCTACAAGGCCAGCAGATCCCTAATTCTCTCTCCAATCAAGTGCGCTCTCCCCAGCCTGTCCCTTCTCCACGGCCACAGTCCcAG\|C\|CCCCCCACTCCAGTCCTTCCCCAAGGATGCAGCCTCAGCCTtCTCCACACC |
| 2:1113:15517:34936#1#F | \| \| |
| MQ=60 isize=225 C=33 | GCCCAGTCCCCACACCTACAAGGCCAGCAGATCCCTAATTCTCTCTCCAATCAAGtGCGCTCTCCCCAGCCTGTCCCTTCTCCACGGCCACAGTCCCAG\|C\|CCCCCCACTCCAGTCCTTCCCCAagGatGCAGCCTCAGCCtTCTCCACACC |
| 2:1113:8258:58532#1#F | \| \| |
| MQ=60 isize=233 C=15 | GCcCAGTCCCCACACCTACAAGGCCAGCAGATCCCTAaTTCTCTCTCCAATCAAGTGCGCTCTCCCcAGCCTGTCCcTTCTCCACGGCCACAGtCCCAG\|c\|CcCCCCaCTCCAGtCctTCCCCAAGGATGCAGCCTCAGCCtTCTccacacc |
| 2:1113:16914:95085#1#F | \| \| |
| MQ=60 isize=174 C=33 | GCCCAGTCCCCACACCTACAAGGCCAGCAGATCCCTAATTCTCTCTCCAATCAAGTGCGCTCTCCCCAGCCTGTCCCTTCTCCACGGCCACAGTCCCAG\|C\|CCCCCCACTCCAGTCCTTCCCCAAGGATGCAGCCTCAGCCTTCTCCACACC |
| 2:1114:19870:8134#1#F | \| \| |
| MQ=60 isize=121 C=37 | GCCCAGTCCCCACACCTACAAGGCCAGCAGATCCCTAATTCTCTCTCCAATCAAGTGCGCTCTCCCCAGCCTGTCCCTTCTCCACGGCCACAGTCCCAG\|C\|CCCCCCACTCCAGTCCTTCCCAGGACTCCAATAATCTTTTGTTACTGTCTC |
| 2:1114:20256:16931#1#F | \| \| |
| MQ=60 isize=138 C=33 | GCCCAGTCCCCACACCTACAAGGCCAGCAGATCCCTAATTCTCTCTCCAATCAAGTGCGCTCTCCCCAGCCTGTCCCTTCTCCACGGCCACAGTCCCAG\|C\|CCCCCCACTCCAGTCCTTCCCCAAGGATGCAGCCTCAGAGGACTCCAATTT |
| 2:1114:14890:80836#1#F | \| \| |
| MQ=60 isize=169 C=33 | GCCCAGTCCCCACACCTACAAGGCCAGCAGATCCCTAATTCTCTCTCCAATCAAGTGCGCTCTCCCCAGCCTGTCCCTTCTCCACGGCCACAGTCCCAG\|C\|CCCCCCaCTCCAGTCCTTCCCCAAGGATGCAGCCTCAGCCTTCTCCACACC |
| 2:1115:2064:39960#1#F | \| \| |
| MQ=60 isize=153 C=33 | GCCCAGTCCCCACACCTACAAGGCCAGCAGATCCCTAATTCTCTCTCCAATCAAGTGCGCTCTCCCCAGCCTGTCCCTTCTCCACGgCCACAGTCCCAG\|C\|CCCCCCacTCCAgtCCTTCCCCAAGgATgCaccCtCAgCCTTCTCcaCACC |
| 2:1115:3977:48244#1#F | \| \| |
| MQ=60 isize=0 C=37 | GCCCAGTCCCCACACCTACAAGgCCAGCAGATCCCTAATTCTCTCTCCAATCAAGTGCGCTCTCCCCAGCCTGTCcCTTCTCCACGGCCACAGtCCcAG\|C\|CCCCCCaCTCCAgtCCTtCCCCAagGatgCagCcTCAgCCTTCTCCACACC |
| 2:1115:6720:77928#1#F | \| \| |
| MQ=60 isize=323 C=33 | GCCCAGTCCCCACACCTACAAGGCCAGCAGATCCCTAATTCTCTCTCCAATCAAGTGCGCTCTCCCCAGCCTGTCCCTTCTCCACGGCCACAGTcCCAG\|C\|CCCCCCACTCCAGTCCTTCCCCAAGGATGCAGCCTCAGCCTTCTCCACACC |
| 2:1116:3693:3969#1#F | \| \| |
| MQ=60 isize=277 C=33 | GCCCAGTCCCCACACCTACAAGGCTAGCAGATCCCTAATTCTCTCTCCAATCAAGTGCGCTCTCCCCAGCCTGTCCCTTCTCCACGGCCACaGTCCCAG\|C\|CCCCCCACTCCAGtCCTTCCCCAAGGATGCAGCCTCAGCCTTCTCCACACC |
| 2:1116:19303:17858#1#F | \| \| |
| MQ=60 isize=198 C=37 | GCCCAGTCCCCACACCTACAAGGCCAGCAGATCCCTAATTCTCTCTCCAATCAAGTGCGCTCTCCCCAGCCTGTCCCTTCTCCACGGCCACAGTCCCAG\|C\|CCCCCCACTCCAGTCCTTCCCCAAGGATGCAGCCTCAGCCTTCTCCACACC |
| 2:1116:2803:25815#1#F | \| \| |
| MQ=60 isize=0 C=33 | GCCCAGTCCCCACACCTACAAGGCCAGCAGATCCCTAATTCTCTCTCCAATCAAGTGCGCTCTCCCCAGCCTGTCCCTTCTCCACGGCCACAGTCCCAG\|C\|CCCCCCACtCCAGTCCTTCCCCAAGGATGCAGCCTCagCCTTCTCCacaCC |
| 2:1116:19569:48179#1#F | \| \| |
| MQ=60 isize=250 C=33 | GCCCAGTCCCCACACCTACAAGGCCAGCAGATCCCTAATTCTCTCTCCAATCAAGTGCGCTCTCCCCAGCCTGTCCCTTCTCCACGGCCACAGTCCCAG\|C\|CCCCCCACTCCAGTCCTTCCCCAAGGATGCAGCCTCAGCCTTCTCCACACC |
| 2:1116:16394:55767#1#F | \| \| |
| MQ=60 isize=228 C=22 | GCCCAGTCCCCACACCTACAAGGCCAGCAGATCCCTAATTCTCTCTCCAaTCAAGtGCGCTCTCCCCAGCCTGTCCCTTCTCCACGGCCACAGtCCCAG\|C\|CCCCCCACTCCAGtCCTTCCCCAAGGATGCAGCCTcAGCcTTCtCCACACC |
| 2:1116:20846:75736#1#F | \| \| |
| MQ=60 isize=159 C=22 | GCCCAGTCCCCACACCTACAAGGCCAGCAGATCCCTAATTCTCTCTCCAATCAAGTGCGCTCTTCCCAGCCTGTCCCTTCTCCACGGCCACAGTCCCAG\|C\|CCCCCCACTCCAGTCCTTCCCCAAGGATGCAGCCTCAGCCTTCTCCACACC |
| 2:1116:11398:91751#1#F | \| \| |
| MQ=60 isize=144 C=33 | GCCCAGTTCCCACACCTACAAGGCCAGCAGATCCCTAATTCTCTCTCCAATCAAGTGCGCTCTCCCCAGCCTGTCCCTTCTCCACGGCCACAGTCCCAG\|C\|CCCCCCACTCCAGTCCtTCCCCAAGGATGCAgCCTCAGCCTTCTAGGACTC |
| 2:1201:1542:36819#1#F | \| \| |
| MQ=60 isize=198 C=33 | GCCCAGTCCCCACACCTACAAGGCCAGCAGATCCCTAATTCTCTCTCCAATCAAGTGCGCTCTCCCCAGCCTGTCCCTTCTCCACGGCCACAGTCCCAG\|C\|CCCCCCACTCCAGTCCTTCCCCAAGGATGCAGCCTTAGCCTTCTCCACACC |
| 2:1201:11115:68061#1#F | \| \| |
| MQ=60 isize=147 C=33 | gCCCAGTCCCcAcACCTACAAgGCCAGCAGATCCCTAATTCTCTCTCCAATCAAGTGCGCTCtCCCCAGCCTGTCCCTTCTCCACGGCCACAgtccCAG\|C\|CcCCCCACTCCAGTCCTtcCCCAaggatgcAGCCTCaGCCttctcCAGGaC |
| 2:1201:6205:74125#1#F | \| \| |
| MQ=60 isize=166 C=22 | GCCCAGTCCCCACACCTACAAGGCCAGCAGaTCCCTAATTcTCTCTCCAATCAAGTgCGCTCTCCCCAGCCTGTCCCTTCTCCACGGCCACaGTCCCAG\|C\|CCCCCCaCtCcaGtCCTTCCCCAAAGATGCAGcCTCAGCcTTCTCCACACC |
| 2:1202:9898:41051#1#F | \| \| |
| MQ=60 isize=251 C=37 | GCCCAGTCCCCACACCTACAAGGCCAGCAGATCCCTAATTCTCTCTCCAATCAAGTGCGCTCTCCCCAGCCTGTCCCTTCTCCACGGCCACAGTCCCAG\|C\|CCCCCCACTCCAGTCCTTCCCCAAGGATGCAGCCTCAGCCTTCTCCACACC |
| 2:1202:15938:63383#1#F | \| \| |
| MQ=60 isize=262 C=22 | GCCCAGTCCCCACACCTACAAGGCCAGCAGATCCCTAATTCTCTCTCCAATCAAGTGCGCTCTCCCCAGCCTGTCCCTTCTCCACGGCCACAGTCCCAG\|C\|CCCCCCACTCCAGTCCtTCCCCAAGGATGCAGCCTCAGCCTTCTCCACACC |
| 2:1202:3450:86967#1#F | \| \| |
| MQ=60 isize=0 C=33 | GCCCAGTCCCCACACCTACAAGGCCAGCAGATCCCTAATTCTCTCTCCAATCAAGTGCGCTCTCCCCAGCCTGTCCCTTCTCCACGGCCACAGTCCCAG\|C\|CCCCCCACTCCAGTCCTTCCCCAAGGATGCAGCCTCAGCCTTCTCCACACC |
| 2:1202:7620:100113#1#F | \| \| |
| MQ=60 isize=159 C=33 | GCCCAGTCCCCACACCTACAAGGCCAGCAGATCCCTAATTCTCTCTCCAATCAAGTGCGCTCTCCCCAGCCTGTCCCTTCTCCACGGCCACAGTCCCAG\|C\|CCCCCCACTCCAGTCCTTCCCCAAGGATGCAGCCTCAGCCTTCTACACACC |
| 2:1203:17225:35953#1#F | \| \| |
| MQ=60 isize=179 C=37 | GCCCAGTCCCCACACCTACAAGGCCAGCAGATCCCTAATTCTCTCTCCAATCAAGTGCGCTCTCCCCAGCCTGTCCCTTCTCCACGGCCACAGTCCCAG\|C\|CCCCCCACTCCAGTCCTTCCCCAAGGATGCAGCCTCAGCCTTCTCCACACC |
| 2:1203:15277:77102#1#F | \| \| |
| MQ=60 isize=132 C=37 | GCCCAGTCCCCACACCTACAAGGCCAGCAGATCCCTAATTCTCTCTCCAATCAAGTGCGCTCTCCCCAGCCTGTCCCTTCTCCACGGCCACAGTCCCAG\|C\|CCCCCCACTCCAGTCCTTCCCCAAGGATGCAGAGGACTCCAATAGGAGACT |
| 2:1203:17008:89798#1#F | \| \| |
| MQ=60 isize=175 C=33 | GCCCAGTCCCCACACCTACAAGGCCAGCAGATCCCTAATTCTCTCTCCAATCAAGTGCGCTCTCCCCAGCCTGTCCCTTCTCCACGGCCACAGTCCCAG\|C\|CCCCCCACTCCAGTCCTTCCCCAAGGATGCAGCCTCAGCCTTCTCCACACC |
| 2:1204:16893:30023#1#F | \| \| |
| MQ=60 isize=131 C=33 | GCCCAGTCCCCACACCTACAAGGCCAGCAGATCCCTAATTCTCTCTCCAATCAAGTGCGCTCtCCCCAGCCTGTCCCTTCTCCACGGCCACAGTCCCAG\|C\|CCCCCCACTCCAGTCCTTCCCCAAGGATGCAGGACTCCAATAAATAGtAGC |
| 2:1205:7030:25691#1#F | \| \| |
| MQ=60 isize=121 C=37 | GCCCAGTACCCACACCTACAAGGCCAGCAGATCCCTAATTCTCTCTCCAATCAAGTGCGCTCTCCCCAGCCTGTCCCTTCTCCACGGCCACAGTCCCAG\|C\|CCCCCCACTCCAGTCCTTCCCAGGACTCCAATGTTAGAACCATGCTGTCTC |
| 2:1205:19814:33623#1#F | \| \| |
| MQ=60 isize=297 C=33 | GCCCAGTCCCCACACCTACAAGGCCAGCAGATCCCTAATTCTCTCTCCAATCAAGTGCGCTCTCCCCAGCCTGTCCCTTCTCCACGGCCACAGTCCCAg\|C\|CCCCCCACTCCAgTCCTTCCCCAAGGATGcAGCCTCagcCTTCTCCACACC |
| 2:1205:7701:49034#1#F | \| \| |
| MQ=60 isize=272 C=37 | GCCCAGTCCCCACACCTACAAGGCCAGCAGATCCCTAATTCTCTCTCCAATCAAGTGCGCTCTCCCCAGCCTGTCCCTTCTCCACGGCCACAGTCCCAG\|C\|CCCCCCACTCCAGTCCTTCCCCAAGGATGCAGCCTCAGCCTTCTCCACACC |
| 2:1205:4346:61254#1#F | \| \| |
| MQ=60 isize=187 C=33 | GCCCAGTCCCCACACCTACAAGGCCAGCAGATCCCTAATTCTCTCTCCAATCAAGTGCGCTCTCCCCAGCCTGTCCCTTCTCCACGGCCACAGTCCCAG\|C\|CCCCCCACTCCAGTCCTTCCCCAAGGATGCAGCCTCAGCCTTCTCCACACC |
| 2:1205:17586:69841#1#F | \| \| |
| MQ=60 isize=149 C=37 | GCCCAGTCCCCACACCTACAAGGCCAGCAGATCCCTAATTCTCTCTCCAATCAAGTGCGCTCTCCCCAGCCTGTCCCTTCTCCACGGCCACAGTCCCAG\|C\|CCCCCCACTCCAGTCCTTCCCCAAGGATGCAGCCTCAGCCTTCTCCACAGG |
| 2:1205:10317:95070#1#F | \| \| |
| MQ=60 isize=131 C=33 | GCCCAGTCCCCACACCTACAAGGCCAGCAGATCCCTAATTCTCTCTCCAATCAAGTGCGCTCTCCCCAGCCTGTCCCTTCTCCACGGCCACAGTCCCAG\|C\|CCCCCCaCTCCAGTCCTTCCCCAAGGATGCAGGACTCCAATGCGCTTCGAA |
| 2:1206:11446:16080#1#F | \| \| |
| MQ=60 isize=144 C=37 | GCCCAGTCCCCACACCTACAAGGCCAGCAGATCCCTAATTCTCTCTCCAATCAAGTGCGCTCTCCCCAGCCTGTCCCTTCTCCACGGCCACAGTCCCAG\|C\|CCCCCCACTCCAGTCCTTCCCCAAGGATGCAGCCTCAGCCTTCTAGGACTC |
| 2:1206:14383:53171#1#F | \| \| |
| MQ=60 isize=203 C=33 | GCCCAGTCCCCACACCTACAAGGCCAGCAGATCCCTAATTCTCTCTCCAAACAAGTGCGCTCTCCCCAGCCTGTCCCTTCTCCACGGCCACAGTCCCAG\|C\|CCCCCCACTCCAGTCCTTCCCCAAGGATGCAGCCTCAGCCTTCTCCACACC |
| 2:1207:17573:81114#1#F | \| \| |
| MQ=60 isize=253 C=37 | GCCCAGTCCCCACACCTACAAGGCCAGCAGATCCCTAATTCTCTCTCCAATCAAGtGCGCTCTCCCCAGCCTGTCCCTTCTCCACGGCCACAGTCCCAG\|C\|CCCCCCACTCCAGTCCTTCCCCAAGGATGCAGCCTCAGCCTTCTCCACACC |
| 2:1208:5942:36365#1#F | \| \| |
| MQ=60 isize=194 C=37 | GCCCAGTCCCCACACCTACAAGGCCAGCAGATCCCTAATTCTCTCTCCAATCAAGTGCGCTCTCCCCAGCCTGTCCCTTCTCCACGGCCACAGTCCCAG\|C\|CCCCCCACTCCAGTCCTTCCCCAAGGATGCAGCCTCAGCCTTCTCCACACC |
| 2:1208:11976:67031#1#F | \| \| |
| MQ=60 isize=133 C=33 | GCCCAGTCCCCACACCTACAAGGCCAGCAGATCCCTAATTCTCTCTCCAATCAAGTGCGCTCTCCCCAGCCTGTCCCTTCTCCACGGCCACAGTCCCAG\|C\|CCCCCCACTCCAGTCCTTCCCCAAGGATGCAGCAGGACTCCAATGCGCACA |
| 2:1209:4889:7972#1#F | \| \| |
| MQ=60 isize=189 C=33 | GCCCAGTCCCCACACCTACAAGGCCAGCAGATCCCTAATTCTCTCTCCAATCAAGTGCGCTCTCCCCAgCCTGTCCCTTCTCCACGGCCACAGTCCCAG\|C\|CCCCCCACTCCAGTCCTTCCCCAAGGATGCAGCCTCAGCCTTCTCCACACC |
| 2:1209:8075:29043#1#F | \| \| |
| MQ=60 isize=170 C=33 | GCCCAGTCCCCACACCTACAAGGCCAGCAGATCCCTAATTCTCTCTCCAATCAAGTGCGCTCTCCCCAGCCTGTCCCTTCTCCACGGCCACAGTCCCAG\|C\|CCCCCCACTCCAGTCCTTCCCCAAGGAtGCAGCCTCAGCCTTCTCCACACC |
| 2:1209:11658:41425#1#F | \| \| |
| MQ=60 isize=205 C=33 | GCCCAGTCCCCACACCTACAAGGCCAGCAGATCCCTAATTCTCTCTCCAATCAAGTGCGCTCTCCCCAGCCTGTCCCTTCTCCACGGCCACAGTCCCAG\|C\|CCCCCCACTCCAGTCCTTCCCCAAGGATGCAGCCTCAGCCTTCTCCACACC |
| 2:1209:5781:72494#1#F | \| \| |
| MQ=60 isize=221 C=33 | GCCCAGT\_CCCACACCTaCAAGGCCAGCAGATCCCTAATTCTCTCTCCAATCAAGTGCGCTCTCCCCAGCCTGTCCCTTCTCCACGGCCACAGtCCCAG\|C\|CCCCCCACTCCAGTCCTTCCCCAAGGATgCAGCCTCAGCCTTCTCCACAcca |
| 2:1209:6559:79942#1#F | \| \| |
| MQ=60 isize=209 C=37 | GCCCAGTCCCCACACCTACAAGGCCAGCAGATCCCTAATTCTCTCTCCAATCAAGTGCGCTCTCCCCAGCCTGTCCCTTCTCCACGGCCACAGTCCCAG\|C\|CCCCCCACTCCAGTCCTTCCCCAAGGATGCAGCCTCAGCCTTCTCCACACC |
| 2:1210:6548:11943#1#F | \| \| |
| MQ=60 isize=244 C=33 | GcCCAGTCCCCACACCTACAAGGCCAGCAGATCCCTAATTCTCTCTCCAATCAAGTGCgCTCTCCCCAGCCTGTCCCTtCTCCACGGCCACAGtcCCAG\|C\|CCCCCCACtCCAgtCCTtCCCCAAGGATGCaGCCTCAgCcttctccacacc |
| 2:1210:17230:51125#1#F | \| \| |
| MQ=60 isize=250 C=33 | GCCCAGTCCCCACACCTACAAGGCCAGCAGATCCCTAATTCTCTCTCCAATCAAGTGCGCTCTCCCCAGCCTGTCCCTTCTCCACGGCCACAGTCCCAG\|C\|CCCCCCACTCCAGTCCTTCCCCAAGGATGCAGCCTCAGCCTTCTCCACACC |
| 2:1210:5262:85604#1#F | \| \| |
| MQ=60 isize=132 C=33 | GCCCAGTCCCCACACCTACAAGGCCAGCAGATCCCTAATTCTCTCTCCAATCAAGTGCGCTCTCCCCAGCCTGTCCCTTCTCCACGGCCACAGTCCCAG\|C\|CCCCCCACtCCAGtCCTTCCCCAAGGAtGCAGGactCCAATTgCCgGCcCC |
| 2:1211:12983:16351#1#F | \| \| |
| MQ=60 isize=230 C=33 | GCCCAGTCCCCACACCTACAAGGCCAGCAGATCCCTAATTCTCTCTCCAATCAAGTGCGCTCTCCCCAGCCTGTCCCTTCTCCACGGCCACAGTCCCAG\|C\|CCCCCCACTCCAGTCCTTCCCCAAGGATGCAGCCTCAGCCTTCTCCACACC |
| 2:1211:17102:19160#1#F | \| \| |
| MQ=60 isize=208 C=33 | GCCCAgTCCCCACACCTACAAGGCCAgCAGATCCCTAATTCTCTCTCCAATCAAGTGCGCTCTCCCCAGCCTGTCCCTTCTCCACGGCtACAGtCcCAG\|C\|CCCCCCACTCCAGTCCTTCCCCAAGGaTGCAGCCTCAGCCTTCTCCACACC |
| 2:1211:18259:35142#1#F | \| \| |
| MQ=60 isize=132 C=37 | GCCCAGTCCCCACACCTACAAGGCCAGCAGATCCCTAATTCTCTCTCCAATCAAGTGCGCTCTCCCCAGCCTGTCCCTTCTCCACGGCCACAGTCCCAG\|C\|CCCCCCACTCCAGTCCTTCCCCAAGGATGCAGAGGACTCCAATGCAGCCGG |
| 2:1211:4116:50194#1#F | \| \| |
| MQ=60 isize=301 C=37 | GCCCAGTCCCCACACCTACAAGGCCAGCAGATCCCTAATTCTCTCTCCAATCAAGTGCGCTCTCCCCAGCCTGTCCCTTCTCCACGGCCACAGTCCCAG\|C\|CCCCCCACTCCAGtCCTTCCCCAAGGATGCAGCCTCAGCCTTCTCCACACC |
| 2:1211:7939:87612#1#F | \| \| |
| MQ=60 isize=205 C=33 | GCCCAGTCCCCACACCTACAAGGCCAGCAGATCCCTAATTCTCTCTCCAATCAAGTGCGCTCTCCCCAGCCTGTCCCTTCTCCACGGCCACAGTCCCAG\|C\|CCCCCCACTCCAGTCCTTCCCCAAGGATGCAGCCTCAGCCTTCTCCACACC |
| 2:1212:8747:10241#1#F | \| \| |
| MQ=60 isize=450 C=33 | GCCCAGTCCCCACACCTACAAGGCCGGCAGATCCCTAATTCTCTCTCCAATCAAGTGCGCTCTCCCCAGCCTGTCCCTTCTCCACGGCCACAGTCCCAG\|C\|CCCCCCACTCCagtcCTTCCCCAAGGAtGCAGCCtcAGCCTtCTCCACaCC |
| 2:1212:5092:28942#1#F | \| \| |
| MQ=60 isize=229 C=22 | GCCCAGTCCCCACACCTACAAGGCCAGCAGATCCCTAATTCTCTCTCCAATCAAGTGCGCTCTCCCCAGCCTGTCCCTTCTCCACGGCCACAGTCCCAg\|C\|CCCCCCACTCCAGTCCTTCCCCAAGGATGCAGCCTCAGCCTTCTCCACACC |
| 2:1212:12693:39325#1#F | \| \| |
| MQ=60 isize=218 C=33 | GCCCAGTCCCCACACCTACAAGGCCAGCAGATCCCTAATTCTCTCTCCAaTCAAGTGCGCTCTCCCCAGCCTGTCCCTTCTCCACGcCCACAgcCCCAG\|C\|CccCCCACTCCAGtcCttCCCCAAGGATgCAGCCTCAGCCtTCtCCacACC |
| 2:1212:14738:45581#1#F | \| \| |
| MQ=60 isize=209 C=27 | GCCCAGTCCCCACACCTACAAGGCCAGCAGATCCCTAATTCTCTCTCCAATCAAGTGCGCTCTCCCCAGCCTGTCCCTTCTCCACGGCCACAGTCCCAG\|C\|CCCCCCACTCCAGTCCTTCCCCAAGGATGCAGCCTCAGCCTtCTCCACACC |
| 2:1212:6586:94382#1#F | \| \| |
| MQ=60 isize=304 C=33 | GCCCAGTCCCCACACCTACAAGGCCAGCAGATCCcTAATTCTCTCTCCAATCAAGTGCGCTCTCCCCAGCCTGTCCCTTCTCCACGGCCACAGtccCAG\|C\|CCCCCCACTCCAGTCCTTCCCCAAGGatGCagCCTCAgccttcTCCACACC |
| 2:1213:5809:25691#1#F | \| \| |
| MQ=60 isize=261 C=22 | GccCAGTCCCCACACCTACAAGGCCAGCAGATCCCTAATTCTCTCTCCAATCAaGTGCGCTCTCCCCAGCCTGTCCCTTCTCCACGgCCaCAGtCCCAg\|C\|CcCCCCACTCCAGTCCTtCCCCAAGGaTGCaGCCTCAGCCTTCTCCACACC |
| 2:1213:16003:45156#1#F | \| \| |
| MQ=60 isize=272 C=33 | GCCCAGTCCCCACACCTACAAGGCCAGCAGATCCCTAATTCTCTCTCCAATCAAGTGCGCTCTCCCCAGCCTGTCCCTTCTCCACGGCCACAGTCCCAG\|C\|CCCCCCACTCCAGTCCtTCCCCAAGGATGCAGCCTCAgCCTTCTCCACACC |
| 2:1214:12923:26337#1#F | \| \| |
| MQ=60 isize=139 C=37 | GCCCAGTCCCCACACCTACAAGGCCAGCAGATCCCTAATTCTCTCTCCAATCAAGTGCGCTCTCCCCAGCCTGTCCCTTCTCCACGGCCACAGTCCCAG\|C\|CCCCCCACTCCAGTCCTTCCCCAAGGATGCAGCCTCAGCAGGACTCCAata |
| 2:1214:2414:52436#1#F | \| \| |
| MQ=60 isize=127 C=27 | GCCCAGTCCCCACACCTACAAGGCCAGCAGATCCCTAATTCTCTCTCCAATCAAGTGCGCTCTCCCCAGCCTGTCCCTTCTCCACGGCCACAGTCCCAG\|C\|CCCCCCACTCCAGTCCTTCCCCAAGGAGGACTCCAATCAGTTGCAGCCCCT |
| 2:1214:9857:81920#1#F | \| \| |
| MQ=60 isize=149 C=33 | GCCCAGTCCCCACACCTACAAGGCCAGCAGATCCCTAATTCTCTCTCCAATCAAGTGCGCTCTCCCCAGCCTGTCCCTTCTCCACGGCCACAGTCCCAG\|C\|CCCCCCaCTCCAgTCCTTCCCCaAGGATGCAGCCTCAGCCTTCTCCACAGG |
| 2:1215:2603:8240#1#F | \| \| |
| MQ=60 isize=175 C=33 | GCCCAGTCCCCACACCTACAAGGCCAGCAGATCCCTAATTCTCTCTCCAATCAAGTGCGCTCTCCCCAGCCTGTCCCTTCTCCACGGCCACAGTCCCAG\|C\|CCCCCCACtCCAGTCCTTCCCCAaGGATGCAGCCTCAGCCTTCTCCACACC |
| 2:1215:17901:60784#1#F | \| \| |
| MQ=60 isize=239 C=37 | GCCCAGTCCCCACACCTACAAGGCCAGCAGATCCCTAATTCTCTCTCCAATCAAGTGCGCTCTCCCCAGCCTGTCCCTTCTCCACGGCCACAGTCCCAG\|C\|CCCCCCACTCCAGTCCTTCCCCAAGGATGCAGCCTCAGCCTTCTCCACACC |
| 2:1215:9402:77111#1#F | \| \| |
| MQ=60 isize=202 C=37 | GCCCAGTCCCCACACCTACAAGGCCAGCAGATCCCTAATTCTCTCTCCAATCAAGTGCGCTCTCCCCAGCCTGTCCCTTCTCCACGGCCACAGTCCCAG\|C\|CCCCCCACTCCAGTCCTTCCCCAAGGATGCAGCCTCAGCCTTCTCCACACC |
| 2:1215:18269:91516#1#F | \| \| |
| MQ=60 isize=260 C=37 | GCCCAGTCCCCACACCTACAAGGCCAGCAGATCCCTAATTCTCTCTCCAATCAAGTGCGCTCTCCCCAGCCTGTCCCTTCTCCACGGCCACAGTCCCAG\|C\|CCCCCCACTCCAGTCCTTCCCCAAGGATGCAGCCTCAGCCTTCTCCACACC |
| 2:1216:19785:36237#1#F | \| \| |
| MQ=60 isize=125 C=33 | GCCCAGTCCCCACACCTACAAGGcCAgcAGATCCCtAATTCTCTCTCCAATCAAGtGCGCTCTCCCCAGCCTGTCCCtTCTCCACGgCCACAgTCCCAG\|C\|CCCCCCACTCCAGtCCTtCCCCAAGAGGACTCCAATGTGaAAAACTCGCTG |
| 2:1216:2406:65558#1#F | \| \| |
| MQ=60 isize=170 C=33 | GCCCAGTCCCCACACCTACAAGGCCAGCAGaTCCCTAATTCTCTCTCCAATCAAGTGCGCTCTCCCCAGCCTGTCCCTTCTCCACGGCCACAGtCCCAG\|C\|CCCCCCACTCCGGtCCTTCCCCAagGATGCAGCcTCaGCCTTCtCCACACC |
| 2:2101:3996:17600#1#F | \| \| |
| MQ=60 isize=198 C=22 | GCCCAGTCCCCACACCTACaAGGCCAGCAGATCCCTAATTCTCTCTCCAATCAAGTGCGCTCTCCCCAgCcTGtCCCTTCTCCaCGgCCACAgtcCcAg\|C\|CCCCCCacTCCAGtccTtCCCCAaggaggcacccccaccctcctccacccc |
| 2:2101:8151:41193#1#F | \| \| |
| MQ=60 isize=179 C=33 | GCCCAGTCCCCACACCTACAAGGCCAGCAGATCCCTAATTCTCTCTCCAATCAAGTGCGCTCTCCCCAGCCTGTCCCTTCTCCACGGCCACAGTCCCAG\|C\|CCCCCCaCtCCAGTCCTTCCCCAAGgaTgCagcCTCAgCCTtctccccacc |
| 2:2101:1630:49743#1#F | \| \| |
| MQ=60 isize=132 C=27 | GCCCAGTCCCCACACCTACAAGGCCAGCAGATCCCTAATTCTCTCTCCAATCAAGtGCGCTCTCCCCAGCCTGTCCCTTCTCCACGgCCaCAGtCCCAg\|C\|CCCCCCcctCCAGtCCTTCCCcaaggatccggaggcctccaatgatccacg |
| 2:2102:5657:51955#1#F | \| \| |
| MQ=60 isize=208 C=27 | gCCCAGTCCCCACACCTACAAGGCCAGCAGATCCCTAaTTCTCTCTCCAATCAAGTGCGCTCTCCCCAGCCTGTCCCTTCTCCaCGgcCACAGtCCCag\|C\|CccCCCaCtCCagcccTTCCCCAAGgatgccgcctcagccttctccccccc |
| 2:2102:19167:57199#1#F | \| \| |
| MQ=60 isize=140 C=37 | GCCCAGTCCCCACACCTACAAGGCCAGCAGATCCCTAATTCTCTCTCCAATCAAGTGCGCTCTCCCCAGCCTGTCCCTTCTCCACGGCCACAGTCCCAG\|C\|CCCCCCACTCCAGTCCTTCCCCAAGGATGCAGCCTCAGCCAGGACTCCAAt |
| 2:2103:20418:9721#1#F | \| \| |
| MQ=60 isize=198 C=33 | GCCCAGTCCCCACACCTACAAGGCCAGCAGATCCCTAATTCTCTCTCCAATCAAGTGCGCTCTCCCCAGCCTGTCCCTTCTCCACGGCCACAGTCCCAG\|C\|CCCCCCACTCCAGTCCTTCCCCAAGGATGCAGCCTCAGCCTTCTCCACACC |
| 2:2103:13128:53412#1#F | \| \| |
| MQ=60 isize=150 C=33 | GCCCAGTCCCCACACCTACAAGGCCAGCAGATCCCTAATTCTCTCTCCAATCAAGTGCGCTCTCCCCAGCCTGTCCCTTCTCCACGGCCACAGTCCCAG\|C\|CCCCCCACTCCAGTCCTTCCCCAAGGATGCAGCCtCAGCCTTCTCCACACa |
| 2:2103:5819:95658#1#F | \| \| |
| MQ=60 isize=189 C=33 | GCCCAGTCCCCACACCTACAAGGCCAGCAGATCCCTAATTCTCTCTCCAATCAAGTGCGCTCTCCCCAGCCTGTCCCTTCTCCACGGCCaCaGtCCCAG\|C\|CCCCCCACTCCggtccTtCCCCaaggaggcgccctcacccttctccacacc |
| 2:2104:17624:12158#1#F | \| \| |
| MQ=60 isize=158 C=37 | GCCCAGTCCCCACACCTACAAGGCCAGCAGATCCCTAATTCTCTCTCCAATCAAGTGCGCTCTCCCCAGCCTGTCCCTTCTCCACGGCCACAGTCCCAG\|C\|CCCCCCACTCCAGTCCTTCCCCAAGGATGCAGCCTCAGCCTTCTCCACACC |
| 2:2104:3264:56169#1#F | \| \| |
| MQ=60 isize=203 C=22 | GCCCAGTCCCCACACCTACAAGGCCAGCAGATCCCTAATTCTCTCTCCAATCAAGTGCGCTCTCCCCAGCCTGTCCCTTCTCCaCGgCCACAGTCCCAg\|C\|CCCCCCACtCCAGtCCTTCCCCAAGgatccaccctcagccttccccacacc |
| 2:2105:16395:62282#1#F | \| \| |
| MQ=60 isize=164 C=15 | GCCCAGTCCCCACACCTACAAGGCCAGCAGATCCCTAATTCTCTCTCCAATCAAGtGCGCTCTCCCCAGCCTGtCCCTTCTCCACGGCCaCagtCccAG\|c\|CCacCCACTCCcGtCCTTCCCCAAGGatGcAGCCTCagcCTTCtCCAcacc |
| 2:2106:14935:5406#1#F | \| \| |
| MQ=60 isize=147 C=33 | GCCCAGTCCCCACACCTACAAGGCCAGCAGATCCCTAATTCTCTCTCCAATCAAGTGCGCTCTCCCCAGCCTGTCCCTTCTCCACGGCCACAGTCCCAG\|C\|CCCCCCACTCCAGTCCTTCCCCAAGGATGCAGCCTCAGCCTTCTCCAGGAC |
| 2:2106:11992:27454#1#F | \| \| |
| MQ=60 isize=123 C=2 | GCCCAGTCCCCACACCTACAAGGCCAGCAGATCCCTAaTTcTCTCTCCAATCAAGTGCGCTCTCCCCAGCCTGTCCCTTCTCCACGGCCACagtcccag\|c\|ccccccacccccatccctcccccggaccccaatcacccccacccccccgcc |
| 2:2106:18604:43605#1#F | \| \| |
| MQ=60 isize=338 C=33 | GCCCAGTCCCCACACCTACAAGGCCAGCAGATCCCTAATTCTCTCTCCAATCAAGTGCGCTCTCCCCAGCCTGTCCCTTCTCCACGGCCACAGTCCCAG\|C\|CCCCCCACTCCAGTCCTTCCCCAAGGATGCAGCCTCAGCCTTCTCCACACC |
| 2:2106:3638:98107#1#F | \| \| |
| MQ=60 isize=0 C=27 | GCCCAGTCCCCACACCTACAAGGCCAGCAGATCCCTAATTCTCTCTCCAATCAAGTGCGCTCTCCCCAgCCTGTCCCTTCTCCccggCCaCAGcCCCAg\|C\|CCCCCCAcccCaGcCcTtCCCcaaggatccacccccagccttctccccccc |
| 2:2107:11842:51369#1#F | \| \| |
| MQ=60 isize=235 C=37 | GCCCAGTCCCCACACCTACAAGGCCAGCAGATCCCTAATTCTCTCTCCAATCAAGTGCGCTCTCCCCAGCCTGTCCCTTCTCCACGGCCACAGTCCCAG\|C\|CCCCCCACTCCAGTCCTTCCCCAAGGATGCAGCCTCAGCCTTCTCcacacc |
| 2:2107:9475:77122#1#F | \| \| |
| MQ=60 isize=207 C=33 | GCCCAGTCCCCACACCTACAAGGCCAGCAGATCCCTAATTCTCTCTCCAATCAAGTGCGCTCTCCCCAGCCTGTCCCTTCTCCACGGCCACAGTCCCAG\|C\|CCCCCCACTCCAGTCCTTCCCCAaGGaTGCAGCCccaGCCTTctCCACACc |
| 2:2108:10546:19617#1#F | \| \| |
| MQ=60 isize=206 C=33 | GCCCAGTCCCCACACCTACAAGGCCAGCAGATCCCTAATTCTCTCTCCAATCAAGTGCGCTCTCCCCAGCCTGTCCCTTCTCCACGGCCACAGTCCCAG\|C\|CCCCCCACTCCAGTCCTTCCCCAAGGATGCAGCCTCAGCCTTCTCCACACC |
| 2:2108:18073:26462#1#F | \| \| |
| MQ=60 isize=262 C=33 | GCcCAGTCCCCACACCTACAAGGCCAGCAGATCCCTAaTTCTCTCTCCAATCAAGTGCGCTCTCCCCAGCCTGTCCCTTCTCCACGGCCACAGTCCCAG\|C\|CCcCCCACTCCagtCCTTCCCcAAGGATGCAGCCTcAGCCTTCTCCACACc |
| 2:2108:5671:43256#1#F | \| \| |
| MQ=60 isize=181 C=37 | GCCCAGTCCCCACACCTACAAGGCCAGCAGATCCCTAATTCTCTCCCCAATCAAGTGCGCTCTCCCCAGCCTGTCCCTTCTCCACGGCCACAGACCCAG\|C\|CcCCCCaCTCCAgtCCTtCCCCaAGgatgcAGCCTCAGCCTTctccacacc |
| 2:2109:8822:4014#1#F | \| \| |
| MQ=60 isize=132 C=33 | GCCCAGTCCCCACACCTACAAGGCCAGCAGATCCCTAATTCTCTCTCCAATCAAGTGCGCTCTCCCCAGCCTGTCCCTTCTCCACGGCCACAGTCCCAG\|C\|CCCCCCACTCCAGTCCTTCCCCAAGGATGCAGAGGaCTCCAATCGCGTTAT |
| 2:2109:7868:57543#1#F | \| \| |
| MQ=60 isize=134 C=33 | GCCCAGTCCCCACACCTACAAGGCCAGCAGATCCCTAATTCTCTCTCCAATCAAGTGCGCTCTCCCCAGCCTGTCCCTTCTCCACGGCCACAGTCCCAG\|C\|CCCCCCACTCCAGtCCTTCCCCAAGGATGCAGCCAGGACTCCAATTACACA |
| 2:2109:5517:73892#1#F | \| \| |
| MQ=60 isize=134 C=33 | GCCCAGTCCCCACACCTACAAGGCCAGCAGATCCCTAATTCTCTCTCCAATCAAGTGCGCTCTCCCCAGCCTGTCCCTTCTCCACGGCCaCAGTCCCAG\|C\|CCCCCCaCtCCAGtCCTTCCCCAAGGATGCAgCCaggacTCCAATtactgt |
| 2:2109:8660:82689#1#F | \| \| |
| MQ=60 isize=145 C=37 | GCCCAGTCCCCACACCTACAAGGCCAGCAGATCCCTAATTCTCTCTCCAATCAAGTGCGCTCTCCCCAGCCTGTCCCTTCTCCACGGCCACAGTCCCAG\|C\|CCCCCCACTCCAGTCCTTCCCCAAGGATGCAGCCTCaGCCTTCTCAGGacc |
| 2:2110:9096:4652#1#F | \| \| |
| MQ=60 isize=245 C=33 | GCCCAGTCCCCACACCTACAAGGCCAGCAGATCCCTAATTCTCTCTCCAATCAAGTGCGCTCTCCCCAGCCTGTCCCTTCTCCACGGCCACAGTCCCAG\|C\|CCCCCCACTCCAGTCCTTCCCCAAGGATGCAGCCtCAGCCTtctCCACACC |
| 2:2110:4078:23503#1#F | \| \| |
| MQ=60 isize=187 C=15 | GCCCAGTCCCCACACCTACAAGGCCAGCAGATCCCTAATTCTCTCTCCAATCAAGTGCGCTCTCCCCAgCCTGtCCCTTCTCCACGGCCACaGTCCCAG\|c\|CCCCCCaCtCCAGtCCTtCCCCAaGGatgcagccccggccttctccacccc |
| 2:2110:13444:84277#1#F | \| \| |
| MQ=60 isize=190 C=33 | GCCCAGTCCCCACACCTACAAGGCCAGCAGATCCCTAATTCTCTCTCCAATCAAGTGCGCTCTCCCCAGCCTGTCCCTTCTCCACGGCCACAGTCccAG\|C\|CCCCCCACTCCAGTCCTTCCCCAAGGATGCAGCCTCAGCCTTCTCCACACC |
| 2:2111:3376:6280#1#F | \| \| |
| MQ=60 isize=0 C=33 | GCCCAGTCCCCACACCTACAAGGCCAGCAGATCCCTAATTCTCTCTCCAATCAAGTGCGCTCTCCCCGgCCTGTCCCTTCTCCACGgCCACAGTCCCAG\|C\|CCCCCCaCTCCAgtCCTTCCCCAAGGAtgcggcctcagccttctccccccc |
| 2:2111:4089:22803#1#F | \| \| |
| MQ=60 isize=277 C=33 | GCCCAGTCCCCACACCTACAAGGCCAGCAGATCCCTAATTCTCTCTCCAATCAAGTGCGCTCTCCCCAGCCTGTCCCTTCTCCACGGCCACAGTCCCAG\|C\|CCCCCCaCccCAGtCcTtCCCCAaGgatgcagcctcagccttctccacccc |
| 2:2111:8762:42409#1#F | \| \| |
| MQ=60 isize=206 C=37 | GCCCAGTCCCCACACCTACAAGGCCAGCAGATCCCTAATTCTCTCTCCAATCAAGTGCGCTCTCCCCAGCCTGTCCCTTCTCCACGGCCACAGTCCCAG\|C\|CCCCCCACTCCAGtCCTTCCCCAAGGATGCAGCCTCAGCCTTCTCCACACC |
| 2:2111:18851:61483#1#F | \| \| |
| MQ=60 isize=144 C=37 | GCCCAGTCCCCACACCTACAAGGCCAGCAGATCCCTAATTCTCTCTCCAATCAAGTGCGCTCTCCCCAGCCTGTCCCTTCTCCACGGCCACAGTCCCAG\|C\|CCCCCCACTCCAGTCCTTCCCCAAGGATGCAGCCTCAGCCTTCTAGGACTC |
| 2:2111:5698:98880#1#F | \| \| |
| MQ=60 isize=200 C=33 | GCCCAGTCCCCACACCTACAAGGCCAGCAGATCCCTAATTCTCTCTCCAATCAAGTGCGCTCTCCCCAGCCTGTCCCTTCTCCACGGCCACAGTCCCAG\|C\|CCCCCCccTCCAGtCCTTCCCCAAGGATGCAGCCTCAGCCTTCTCCACACC |
| 2:2112:20711:16329#1#F | \| \| |
| MQ=60 isize=160 C=37 | GCCCAGTCCCCACACCTACAAGGCCAGCAGATCCCTAATTCTCTCTCCAATCAAGTGCGCTCTCCCCAGCCTGTCCCTTCTCCACGGCCACAGTCCCAG\|C\|CCCCCCACTCCAGTCCTTCCCCAAGGATGCAGCCTCAGCCTTCTCCACACC |
| 2:2112:16087:44938#1#F | \| \| |
| MQ=60 isize=205 C=33 | GCCCAGTCCCCACACCTACAAGgCCAGCAGaTCCCTAaTTCtCTCTCCAATCAAGTGCGcTCTCCCCAGCCTGTCCCTTCTCCACGgccACAGTCCCAG\|C\|CCCCCCACTCCAGTCCTTCCCCAAGGAtGCAGCCTcaGCCTTCTCcACAcc |
| 2:2112:19488:62434#1#F | \| \| |
| MQ=60 isize=138 C=37 | GCCCAGTCCCCACACCTACAAGGCCAGCAGATCCCTAATTCTCTCTCCAATCAAGTGCGCTCTCCCCAGCCTGTCCCTTCTCCACGGCCACAGTCCCAG\|C\|CCCCCCACTCCAGTCCTTCCCCAAGGATGCAGCCTCAGAGGACTCCAATTG |
| 2:2112:15737:74412#1#F | \| \| |
| MQ=60 isize=246 C=33 | GCCCAGTCCCCACACCTACAAGGCCAGCAGATCCCTAATTCTCTCTCCAATCAAGTGCGCTCTCCCCAGCCTGTCCCTTCTCCACGGCCACAGTCCCAG\|C\|CCCCCCACTCCAGTCCTTCCCCAAGGATGCAGCCTCAGCCTTCTCCACACC |
| 2:2113:7125:10840#1#F | \| \| |
| MQ=60 isize=201 C=37 | GCCCAGTCCCCACACCTACAAGGCCAGCAGATCCCTAATTCTCTCTCCAATCAAGTGCGCTCTCCCCAGCCTGTCCCTTCTCCACGGCCACAGTCCCAG\|C\|CCCCCCACTCCAGTCCTTCCCCaAGGaTGCAGCCTCAgCCTTCTCCACACc |
| 2:2113:4928:54280#1#F | \| \| |
| MQ=60 isize=132 C=33 | GCCCAGTCCCCACACCTACAAGGCCAGCAGATCCCTAATTCTCTCTCCAATCAAGTGCGCTCTCCCCAGCCTGTCCCTTCTCCACGGCCACAGTCCCAG\|C\|CCCCCCACTCCAGtCCTTCCCCAAGGATGCagAGGACtCCAATGATaCACt |
| 2:2113:17287:90016#1#F | \| \| |
| MQ=60 isize=230 C=27 | GCCCAGTCCCCACACCTACAAGGCCAGCAGATCCCTAATTCTCTCTCCAATCAAGTGCGCTCTCCCCAGCCTGTCCCTTCTCCACGGCCACAGTCCCAG\|C\|CCCCCCACTCCAGTCCTTCCCCAAGGATGCAGCCTCAGCCTTCTCCACACC |
| 2:2114:7810:51415#1#F | \| \| |
| MQ=60 isize=123 C=33 | GCCCAGTCCCCACACCTACAAGGCCAGCAGATCCCTAATTCTCTCTCCAATCAAGTGCGCTCTCCCCAGCCTGTCCCTTCTCCACGGCCACAGTCCCAG\|C\|CCCCCCACTCCAGTCCTTCCCCAaGGAGGACTCCAATCAGtTGCAGCGCCt |
| 2:2114:15521:65576#1#F | \| \| |
| MQ=60 isize=283 C=37 | GCCCAGTCCCCACACCTACAAGGCCAGCAGATCCCTAATTCTCTCTCCAATCAAGTGCGCTCTCCCCAGCCTGTCCCTTCTCCACGGCCACAGTCCCAG\|C\|CCCCCCACTCCAGTCCTTCCCCAAGGATGCAGCCTCAGCCTTCTCCACACC |
| 2:2114:10330:97507#1#F | \| \| |
| MQ=60 isize=233 C=37 | GCCCAGTCCCCACACCTACAAGGCCAGCAGATCCCTAATTCTCTCTCCAATCAAGTGCGCTCTCCCCAGCCTGTCCCTTCTCCACGGCCACAGTCCCAG\|C\|CCCCCCACTCCAGTCCTTCCCCAAGGATGCAGCCTCAGCCTTCTCCACACC |
| 2:2115:4265:2103#1#F | \| \| |
| MQ=60 isize=212 C=37 | ncCCAGTCCCCACACCTACAAGGCCAGCAGATCCCTAATTCTCTCTCCAATCAAGTGCGCTCTCCCCAGCCTGTCCCTTCTCCACGGCCACAGTCCCAG\|C\|CCCCCCACtCCAGTCCTTCCCCAAGGATtcaGCCTCaGCCTTCTCCACACC |
| 2:2115:1506:25394#1#F | \| \| |
| MQ=60 isize=132 C=33 | GCCCAGTCCCCACACCTACAAGGCCAGCAGATCCCTAATTCTCTCTCCAATCAAGTGCGCTCTCCCCAGCCTGTCCCTTctCCACGGCCccAgTCCCAg\|C\|CCCCCCactCCAgtCcTTCCCCAaggaggcaggactccaatattaacaccg |
| 2:2115:14383:30663#1#F | \| \| |
| MQ=60 isize=209 C=33 | GcCCAGTCCCCACACCTACAAGGCCAGCAGATCCCTAATTCTCTCTCCAATCAAGTGCGCTCTCCCCAGCCTGtCCCTTCTCCACGGCCACAGTCCCAG\|C\|CcCTCCACTcCAGTCCTTCCCCAAGGATGCAGCCTCAGCCTTCTCCACACc |
| 2:2115:11701:68588#1#F | \| \| |
| MQ=60 isize=147 C=33 | GCCCAGTCCCCACACCTACAAGGCCAGCAGATCCCTAATTCTCTCTCCAATCAAGTGCGCTCTCCCCAGCCTGTCCCTTCTCCACGGCCACAGTCCCAG\|C\|CCCCCCACTCCAGTCCTTCCCCAAGGATGCAGCCTCAGCCTTCTCCAGGAC |
| 2:2116:2701:2859#1#F | \| \| |
| MQ=60 isize=174 C=33 | GCCCAGTCCCCACACCTACAAGGCCAGCAGATCCCTAATTCTCTCTCCAATCAAGTGCGCTCTCCCCAGCCTGTCCCTTCtCCACGGCCACAGtCCCAG\|C\|CCCCCCACtCCagtCcttcCCCAaggatgcagcctcagccttctccacacc |
| 2:2116:15977:51515#1#F | \| \| |
| MQ=60 isize=149 C=33 | GCCCAGTCCCCACACCTACAAGGCCAGCAGATCCCTAATTCTCTCTCCAATCAAGTGCGCTCTCCCCAGCCTGTCCCTTCTCCACGGCCACAGTCCCAG\|C\|CCCCCCACTCCAGTCCTTCCCCAAGGATGCAGCCTCAGCCTTCTCCACAGG |
| 2:2116:18590:86459#1#F | \| \| |
| MQ=60 isize=239 C=37 | GCCCAGTCCCCACACCTACAAGGCCAGCAGATCCCTAATTCTCTCTCCAATCAAGTGCGCTCTCCCCAGCCTGTCCCTTCTCCACGGCCACAGTCCCAG\|C\|CCCCCCACTCCAGTCCTTCCCCAAGGATGCAGCCTCAGCCTTCTCCACACC |
| 2:2201:18236:26983#1#F | \| \| |
| MQ=60 isize=177 C=37 | GCCCAGTCCCCACACCTACAAGGCCAGCAGATCCCTAATTCTCTCTCCAATCAAGTGCGCTCTCCCCAGCCTGTCCCTTCTCCACGGCCACAGTCCCAG\|C\|CCCCCCACTCCAGTCCTTCCCCAAGGATGCAGCCTCAGCCTTCTCCACacc |
| 2:2201:14484:64378#1#F | \| \| |
| MQ=60 isize=160 C=33 | GCCCAGTCCCCACACCTACAAGGCCAGCAGATCCCTAATTCTCTCTCCAATCAAGTGCGCTCTCCCCAGCCTGTCCCTTCTCCACGGCCACAGTCCCAG\|C\|CCCCCCACTCCAGTCCTTCCCCAAGGATGCAGCCTCAGCCTTCTCCACACC |
| 2:2202:12041:22341#1#F | \| \| |
| MQ=60 isize=137 C=37 | GCCCAGTCCCCACACCTACAAGGCCAGCAGATCCCTAATTCTCTCTCCAATCAAGTGCGCTCTCCCCAGCCTGTCCCTTCTCCACGGCCACAGTCCCAG\|C\|CCCCCCACTCCAGTCCTTCCCCAAGGATGCAGCCTCAAGGACTCCAATACg |
| 2:2202:4973:44987#1#F | \| \| |
| MQ=60 isize=205 C=37 | GCCCAGTCCCCACACCTACAAGGCCAGCAGATCCCTAATTCTCTCTCCAATCAAGTGCGCTCTCCCCAGCCTGTCCCTTCTCCACGGCCACAGTCCCAG\|C\|CCCCCCACTCCAGTCCTTCCCCAAGGATGCAGCCTCAGCCTTCTCCACACC |
| 2:2202:12301:62809#1#F | \| \| |
| MQ=60 isize=307 C=33 | GCCCAGTCCCCACACCCACAAGGCCAGCAGATCCCTAATTCTCTCTCCAATgaAGTGCGCTCTCCCCAGCCTGTCCCTTCTCCACGGCCACAGTCCCAG\|C\|CCCCCCACTCCAGTCCTTCCCCAAGGATGCAGCCTCAGCCTTCTCCACGCC |
| 2:2202:15863:76980#1#F | \| \| |
| MQ=60 isize=193 C=33 | GCCCAGTCCCCACACCTACAAGGCTAGCAGATcCCTAATTCTCTCTCCAATCAAGTGCGCTCTCCCCAGCCTGTCCCTTCTCCACGGCCACAGTCCCAG\|C\|cCCCCCACTCCAGTCCTTCCCCAAGGATGCAGCCTCaGcCTTCTCCACACC |
| 2:2202:12505:96125#1#F | \| \| |
| MQ=60 isize=244 C=37 | GCCCAGTCCCCACACCTACAAGGCCAGCAGATCCCTAATTCTCTCTCCAATCAAGTGCGCTCTCCCCAGCCTGTCCCTTCTCCACGGCCACAGTCCCAG\|C\|CCCCCCACTCCAGTCCTTCCCCAAGGATGCAGCCTCAGCCTTCTCCACACC |
| 2:2203:14945:18147#1#F | \| \| |
| MQ=60 isize=268 C=33 | GCCCAGTCCCCACACCTACAAGGCCAGCAGATCCCTAATTCTCTCTCCAATCAAGTGCGCTCTCCCCAGCCTGTCCCTTCTCCACGGCCACAGTCCCAG\|C\|CCCCCCACTCCAGTCCTTCCCCAAGGATGCAGCCTCAGCCTTCTCCACACC |
| 2:2203:13170:32362#1#F | \| \| |
| MQ=60 isize=175 C=37 | GCCCAGTCCCCACACCTACAAGGCCAGCAGATCCCTAATTCTCTCTCCAATCAAGTGCGCTCTCCCCAGCCTGTCCCTTCTCCACGGCCACAGTCCCAG\|C\|CCCCCCACTCCAGTCCTTCCCCAAGGATGCAGCCTCAGCCTTCTCCACACC |
| 2:2203:8962:56545#1#F | \| \| |
| MQ=60 isize=271 C=37 | GCCCAGTCCCCACACCTACAAGGCCAGCAGATCCCTAATTCTCTCTCCAATCAAGTGCGCTCTCCCCAGCCTGTCCCTTCTCCACGGCCACAGTCCCAG\|C\|CCCCCCACTCCAGTCCTTCCCCAAGGATGCAGCCTCAGCCTTCTCCACACC |
| 2:2203:4204:85254#1#F | \| \| |
| MQ=60 isize=246 C=33 | GCCCAGTCCCCACACCTACAAGGCCAGCAGATCCCTAATTCTCTCTCCAATCAAGTGCGCTCTCCCCAGCCTGTCCCTTCTCCACGGCcACAGtCCCAG\|C\|CCCCCCACTCCAgtCCTTCCCCAAGGATGCAGCCtcagCCTTCTCCACaCC |
| 2:2204:5396:19236#1#F | \| \| |
| MQ=60 isize=161 C=33 | GCCCAGTCCCCACACCTACAAGGCCAGCAGATCCCTAATTCTCTCTCCAATCAAGTGCGCTCTCCCCAGCCTGTCCCTTCTCCACGGCCACAGTCCCAG\|C\|CCCCCCACTCCAGTCCTTCCCCAAGGATGCAGCCTCAGCCTTCTCCACACC |
| 2:2204:3657:33353#1#F | \| \| |
| MQ=60 isize=285 C=33 | GCCCAGTCCCCACACCTACAAGGCCAGCAGATCCCTAATTCTCTCTCCAATCAAGTGCGCTCTCCCCAGCCTGTCCCTTCTCCACGGCCACAGTCCCAG\|C\|CCCCCCACTCCAGTCCTTCCCCAAGGATGCAGCCTCAGCCTTCTCCACACC |
| 2:2204:14595:76151#1#F | \| \| |
| MQ=60 isize=132 C=33 | GCCCAGTCCCCACACCTACAAGGCCAGCAGATCCCTAATTCTCTCTCCAATCAAGTGCGCTCTCCCCAGCCTGTCCCTTCTCCACGGCCACAGTCCCTG\|C\|CCCCCCACTCCAGTCCTTCCCCAAGGATGCAGAGGACTCCAATCGCGGTGG |
| 2:2204:20398:95911#1#F | \| \| |
| MQ=60 isize=192 C=33 | GCCCAGTCCCCACAcCTACAAGGCCAGCAGATCCCTAATTCTCTCTCCAATCAAGTGCGCTCTCCCCAGCCTGTCCCTTCTCCACGGCtACAGTCCCAG\|C\|CCCCCCACTCCAGtCCtTCCCCAAGGAtGCAGCCTCAGCCTTCTCCACACC |
| 2:2205:18951:10568#1#F | \| \| |
| MQ=60 isize=140 C=33 | GCCCAGTCCCCACACCTACAGGGCCAGCAGATCCCTAATTCTCTCTCCAATCAAGTGCGCTCTCCCCAGCCTGTCCCTTCTCCACGGCCACAGTCCCAG\|C\|CCCCCCACTCCAGTCCTTCCCCAAGGATGCAGCCTCAGCCAGGACTCCAAT |
| 2:2205:20047:69020#1#F | \| \| |
| MQ=60 isize=149 C=33 | GCCCAGTCCCCACACCTACAAGGCCAGCAGATCCCTAATTCTCTCTCCAATCAAGTGCGCTCTCCCCAGCCTGTCCCTTCTCCACGGCCACAGTCCCAG\|C\|CCCCCCACTCCAGTccttccCCAAGGATGCAGCCTCAGCCTTCTCCACAGG |
| 2:2205:19293:81887#1#F | \| \| |
| MQ=60 isize=262 C=33 | GCCCAGTCCCCACACCTACAAGGCCAGCAGATCCCTAATTCTCTCTCCAATCAAGTGCGCTCTCCCCAGCCTGTCCCTTCTCCACGGCCACAGTCCCAG\|C\|CCCCCCACTCCAGTgcttccccaagGATGCAGCCTCAGCCTTCTCCACACC |
| 2:2206:18445:8540#1#F | \| \| |
| MQ=60 isize=187 C=33 | GCCCAGtCcCCACACCTACAAGGCCAGCAGATCCCTAATTCTCTCTCCAATCAAGTGCGCTCTCCCCAGCCTGTCCCTTCTCCACGGCCACAGtCCCAG\|C\|CCCCCCACTCCAGtCCTTCCCCAAGGATGCAgcCTCAGCCTTCTCCACACC |
| 2:2206:15384:20804#1#F | \| \| |
| MQ=60 isize=467 C=33 | GCCCAGTCCCCACACCTACAAGGCCAGCAGATCCCTAATTCTCTCTCCAATCAAGTGCGCTCTCCCCAGCCTGTCCCTTCTCCACGGCCACAGTCCCAG\|C\|CCCCCCACTCCAGTCCTTCCCCAAGGATGCAgcCTCAGCCTTCTCCACACC |
| 2:2206:3212:43751#1#F | \| \| |
| MQ=60 isize=254 C=33 | GCCCAGTCCCCACACCTACAAGGCCAGCAGATCCCTAATTCTCTCTCCAATCAAGTGCGCTCTCCCCAGCCTGTCCCTTCTCCACGGCCACAGTCCCAG\|C\|CCCCCCACTCCAGTCCTTCCCCAAGGATGCAGCCTCAGCCTTCTCCACACC |
| 2:2206:6054:73469#1#F | \| \| |
| MQ=60 isize=226 C=33 | GCCCAGTCCCCACACCTACAAGGCCAGCAGATCCCTAATTCTCTCTCCAATCAAGTGCGCTCTCCCCAGCCTGTCCCTTCTCCACGGCCACAGTCCCAG\|C\|CCCCCCACTCCAGTCCTTCCCCAAGGATGCAGCCTCAGCCTTCTCCACACC |
| 2:2206:12224:88059#1#F | \| \| |
| MQ=60 isize=263 C=33 | GCCCAGTCCCCACACCTACAAGGCCAGCAGATCCCTAATTCTCTCTCCAATCAAGTGCGCTCTCCCCAGCCTGTCCCTTCTCCACGGCCACAGTCCCAG\|C\|CCCCCCACTCCAGTCCTTCCCCAAGGATGCAGCCGCAGCCTTCTCCACACC |
| 2:2207:6337:48800#1#F | \| \| |
| MQ=60 isize=185 C=33 | GCCCAGTCCCCACACCTACAAGGCCAGCAGATCCCTAATTCTCTCTCCAATCAAGTGCGCTCTCCCCAGCCTGTCCCTTCTCCACGGCCACAGtCCCAG\|C\|CCCCCCACTCCAGtCCTTCCCCAAGGATGCAGCCTCAGCCTTCTCCACacc |
| 2:2207:11454:69556#1#F | \| \| |
| MQ=60 isize=175 C=33 | GCCCAGTCCCCACACCTACAAGGCCAGCAGATCCCTAATTCTCTCTCCAATCAAGTGCGCTCTCCCCAGCCTGTCCCTTCTCCACGGCCACAGTCCCAG\|C\|CCCCCCACTCCAGTCCTTCCCCAAGGATGCAGCCTCAGCCTTCTCCACACC |
| 2:2208:13727:10451#1#F | \| \| |
| MQ=60 isize=144 C=33 | GCCCAGTCCCCACACCTACAAGGCCAGCAGATCCCTAATTCTCTCTCCAATCAAGTGCGCTCTCCCCAGCCTGTCCCTTCTCCATGGCCACAGtCCCAG\|C\|CCCCCCACTCCAGTCCTTCCCCAAGGATGCAGCCTCAGCCTTCTaGGACTC |
| 2:2208:12749:44029#1#F | \| \| |
| MQ=60 isize=120 C=37 | GCCCAGTCCCCACACCTACAAGGCCAGCAGATCCCTAATTCTCTCTCCAATCAAGTGCGCTCTCCCCAGCCTGTCCCTTCTCCACGGCCACAGTCTCAG\|C\|CCCCCCACTCCAGTCCTTCCAGGACTCCAATCCCTAACCCTCCCTGTCTCT |
| 2:2208:10450:77939#1#F | \| \| |
| MQ=60 isize=162 C=27 | GCCCAGTCCCCACACCTACAAGGCCAGCAGATCCCTAATTCTCTCTCCAATCAAGTGCGCTCGCCCCAGCCTGTCCCTTCTCCACGGCCACAGTCCCAg\|C\|CCCCCCACTCCAgTCCTTCCCCAAgGATGCaGCCTCagCCttcTCCACACC |
| 2:2209:1302:48088#1#F | \| \| |
| MQ=60 isize=0 C=15 | GCCCAGTCCCCACACCTACAAGGCCAGCAGATCCCTAATTCTcTCTCCAATCAAGTGCGCTCTCCCCAGCCTgTCCCTTCTCCACGgcCACAGTCCCAG\|c\|cccccCACTCCAGTCCTTCCCCAAGGATGCAGCCtCAGCCTTCTCCACACC |
| 2:2209:10594:66565#1#F | \| \| |
| MQ=60 isize=213 C=33 | GCCCAGTCCCCACACCTACAAGGCCAGCAGATCCCTAATTCTCTCTCCAATCAAGTGCGCTCTCCCCAGCCTGTCCCTTCTCCACGGCCACAGTCCCAG\|C\|CCCCCCACTCCAGTCCTTCCCCAAGGATGCAGCCTCAGCCTTCTCCACACC |
| 2:2210:8616:23282#1#F | \| \| |
| MQ=60 isize=172 C=33 | GCCCAGTCCCCACACCTACAAGGCCAGCAGATCCCTAATTCTCTCTCCAATCAAGTGCGCTCTCCCCAGCCTGTCCCTTCTCCACGGCCACAGTCCCAG\|C\|CCCCCCACTCCAGTCCTTCCCCAAGGATGCAGCCTCAGCCTTCTCCACACC |
| 2:2210:5196:39897#1#F | \| \| |
| MQ=60 isize=233 C=33 | GCCCAGTCCCCACACCTACAAGGCCAGCAGATCCCTAATTCTCTCTCCAATCAAGTGCGCTCTCCCCAGCCTGTCCCTTCTCCACGGCCACAGTCCCAG\|C\|CCCCCCACTCCAGTCCTTCCCCAAGGATGCAGCCTCAGCCTTCTCCaCACC |
| 2:2210:9479:86829#1#F | \| \| |
| MQ=60 isize=132 C=33 | GCCCAGTCCCCACACCTACAAGGCCAGCAGATCCCTAATTCTCTCTCCAATCAAGTGCGCTCTCCCCAGCCTGTCCCTTCTCCACGGCCACAGTCCCAG\|C\|CCCCCCACTCCAGTCCTTCCCCAAGGATGCAGAGGACTCCAATCGCGGTGG |
| 2:2211:4469:16637#1#F | \| \| |
| MQ=60 isize=132 C=37 | GCCCAGTCCCCACACCTACAAGGCCAGCAGATCCCTAATTCTCTCTCCAATCAAGTGCGCTCTCCCCAGCCTGTCCCTTCTCCACGGCCACAGTCCCAG\|C\|CCCCCCACTCCAGTCCTTCCCCAAGGATGCAGGACTCCAATCTACGCTAGa |
| 2:2211:18349:27866#1#F | \| \| |
| MQ=60 isize=162 C=33 | GCCCAGTCCCCACACCTACAAGGCCAGCAGATCCCTAATTCTCTCTCCAATCAAGTGCGCTCTCCCCAGCCTGTCCCTTCTCCACGGCCACAGTCCCAG\|C\|CCCCCCACTCCAGTCCTTCCCCAAGGATGCAGcCTCAGCCTTCTCCACACC |
| 2:2211:16910:82807#1#F | \| \| |
| MQ=60 isize=132 C=33 | GCCCAGTCCCCACACCTACAAGGCCAGCAGATCCCTAATTCTCTCTCCAATCAAGTGCGCTCTCCCCAGCCTGTCCCTTCTCCACGGCCACAGTCCCAG\|C\|CCCCCCACTCCAGTCCTTCCCCAAGGATGCAGAGGACTCCAATAACTGCGC |
| 2:2211:6060:89929#1#F | \| \| |
| MQ=60 isize=164 C=27 | GCCCAGTCCCCACACCTACAAGGCCAGCAGATCCCTAATTCTCTCTCCAATCAAGTGCGCTCTCCCCAGCCTGTCCCTTCTCCACgGCCACAGTCCCAG\|C\|CCCCCCACTCCAGTCCTTCCCCAAGGATGCAGCCTCAGCCTTCTCCACACC |
| 2:2212:18470:12227#1#F | \| \| |
| MQ=60 isize=205 C=37 | GCCCAGTCCCCACACCTACAAGGCCAGCAGATCCCTAATTCTCTCTCCAATCAAGTGCGCTCTCCCCAGCCTGTCCCTTCTCCACGGCCACAGTCCCAG\|C\|CCCCCccctccagtCcttCCCCaAGgagccagctccagcctcttccccacc |
| 2:2212:11946:38567#1#F | \| \| |
| MQ=60 isize=221 C=33 | GCCCAGTCCCCACACCTACAAGGCCAGCAGATCCCTAATTCTCTCTCCAATCAAGTGCGCTCTCCCCAGCCTGTCCCTTCTCCACGGCCACAGTCCCAG\|C\|CCCCCCACTCCAGTCCTTCCCCAAGGATGCAGCCTCAGCCTTCTCCACACC |
| 2:2212:13940:58715#1#F | \| \| |
| MQ=60 isize=352 C=33 | GCcCAGTCCCCACACCTACAAGGCCAGCAGATCCCTAATTCTCTCTCCAATCAAGTGCGCTCTCCCCAGCCTGTCCCTTCTCCACggCCACAGTCCCAG\|C\|CCCCCCACTCCAGtCctTCCCCAAGGATGCAGCCTCaGCCTTCTCCACACC |
| 2:2212:17026:96115#1#F | \| \| |
| MQ=60 isize=233 C=33 | GCCCAGTCCCCACACCTACAAGGCCAGCAGATCCCTAATTCTCTCTCCAATCAAGTGCGCTCTCCCCAGCCTGTCCCTTCTCCACGGCCACAGTCCCAG\|C\|CCCCCCACTCCAGTCCTTCCCCAAGGATGCAGCCTCAGCCTTCTCCACACC |
| 2:2213:4036:4627#1#F | \| \| |
| MQ=60 isize=138 C=37 | GCCCAGTCCCCACACCTACAAGGCCAGCAGATCCCTAATTCTCTCTCCAATCAAGTGCGCTCTCCCCAGCCTGTCCCTTCTCCACGGCCACAGTCCCAG\|C\|CCCCCCaCtCCAGTCCTTCCCCAAGGATGCAGCCTCAGAGGaCTCCAATCa |
| 2:2213:13557:37324#1#F | \| \| |
| MQ=60 isize=318 C=33 | GCCCAGTCCCCACACCTACAAGGCCAGCAGATCCCTAATTCTCTCTCCAATCAAGTGCGCTCTCCCCAGCCTGTCCCTTCTCCACGGCCACAGTCCCAG\|C\|CCCCCCACTCCaGTCCTTCCCCAAGGATGCAGCCTCAGCCTTCTCCACACC |
| 2:2213:7126:47743#1#F | \| \| |
| MQ=60 isize=186 C=37 | GCCCAGTCCCCACACCTACAAGGCCAGCAGATCCCTAATTCTCTCTCCAATCAAGTGCGCTCTCCCCAGCCTGTCCCCTCTCCACGGCCACAGTCCCAG\|C\|CCCCCCACTCCAGTCCTTCCCCAAGGATGCAGCCTCAGCCTTCTCCACACC |
| 2:2213:14390:68804#1#F | \| \| |
| MQ=60 isize=121 C=33 | GCCCAGTCCCCACACCTACAAGGCCAGCAGATCCCTAATTCTCTCTCCAATCAAGTGCGCTCTCCCCAGCCTGTCCCTTCTCCACGGCCACAGTCCCAG\|C\|CCCCCCACTCCAGTCCTTCCCAGGACTCCAATGTTAGAACCATGCTGTCTC |
| 2:2213:17249:85880#1#F | \| \| |
| MQ=60 isize=179 C=33 | GCCCAGTCCCCACACCTACAAGGCCAGCAGATCCCTAATTCTCTCTCCAATCAAGTGCGCTCTCCCCAGCCTGTCCCTTCTCCACGGCCACAGTCCCAG\|C\|CCCCCCACTCCAGTCCTTCCCCAAGGATGCAGCCtCAGCCTTCTCCACACC |
| 2:2214:14212:20930#1#F | \| \| |
| MQ=60 isize=132 C=37 | GCCCAGTCCCCACACCTACAAGGCCAGCAGATCCCTAATTCTCTCTCCAATCAAGTGCGCTCTCCCCAGCCTGTCCCTTCTCCACGGCCACAGTCCCAG\|C\|CCCCCCACTCCAGTCCTTCCCCAAGGATGCAGAGGACTCCAATGATACACT |
| 2:2214:5108:56225#1#F | \| \| |
| MQ=60 isize=144 C=33 | GCCCAGTCCCTACACCTACAAGGCCAGCAGATCCCTAATTCTCTCTCCAATCAAGTGCGCTCTCCCCAGCCTGTCCCTTCTCCACGGCCACAGTCCCAG\|C\|CCCCCCACTCCAGTCCTTCCCCAAGGATGCaGCCTCAGCCTTCtaGGaCTC |
| 2:2214:19698:61114#1#F | \| \| |
| MQ=60 isize=132 C=22 | GccCAGTCCCCACACCTACAAGGCCAGCAGATCCCTAATTCTCTCTCCAATCAAGTGcGctctCCCCAgCCTGtcCCTTctCCACGGCCACAGTCCCAg\|C\|CcCCCCACTCCAGTCCTTCCCCAAGgAtGcAGgACTCCAATCgATattAAC |
| 2:2214:4106:71822#1#F | \| \| |
| MQ=60 isize=189 C=37 | GCCCAGTCCCCACACCTACAAGGCCAGCAGATCCCTAATTCTCTCTCCAATCAAGTGCGCTCTCCCCAGCCTGTCCCTTCTCCACGGCCACAGTCCCAG\|C\|CCCCCCACTCCAGTCCTTCCCCAAGGATGCAGCCTCAGCCTTCTCCACACC |
| 2:2214:7777:83746#1#F | \| \| |
| MQ=60 isize=200 C=37 | GCCCAGTCCCCACACCTACAAGGCCAGCAGATCCCTAATTCTCTCTCCAATCAAGTGCGCTCTCCCCAGCCTGTCCCTTCTCCACGGCCACAGTCCCAG\|C\|CCCCCCACTCCAGTCCTTCCCCAAGGATGCAGCCTCAGCCTTCTCCACACC |
| 2:2215:10048:22999#1#F | \| \| |
| MQ=60 isize=145 C=33 | gcCCAGtCCCCACACCTACAAGGCCaGcAGaTCCCTAaTTCTCTcTCCAATCAAGTGCGCTCTCCCCAGCCTGTCCCtTCTCCACGGCCACAGTcccAG\|C\|CcCCCCACTCCagtCCTTCCCCAAGGATGcAGCCTCAGCCtTCTCAGGACT |
| 2:2215:18972:53510#1#F | \| \| |
| MQ=60 isize=145 C=33 | GCCCAGTCCCCACACCTACAAGGCCAGCAGATCCCTAATTCTCTCTCCAATCAAGTGCGCTCTCCCCAGCCTGTCCCTTCTCCACGGCCACAGTCCCAG\|C\|CCCCCCACTCCAGTCCTTCCCCAAGGaTGCAGCCTCAGCCTTCTCAGGACT |
| 2:2215:16761:67222#1#F | \| \| |
| MQ=60 isize=254 C=37 | GCCCAGTCCCCACACCTACAAGGCCAGCAGATCCCTAATTCTCTCTCCAATCAAGTGCGCTCTCCCCAGCCTGTCCCTTCTCCACGGCCACAGTCCCAG\|C\|CCCCCCACTCCAGTCCTTCCCCAAGGATGCAGCCTCAGCCTTCTCCACACC |
| 2:2215:20981:90327#1#F | \| \| |
| MQ=60 isize=205 C=22 | GCCCAGTCCCCACACCTACAAGGCCAGCAGATcCCTAATTCTCTCTCCAATCAAGtGCGCTCTCCCCAGCTTGTCCCTTCTCCACGGCCACAGTCCCAG\|C\|CccCCCACTcCAGTCCTTCCCCAaGGatgCAgCcTCAGCCTTCTCCACACC |
| 2:2216:11681:6512#1#F | \| \| |
| MQ=60 isize=207 C=33 | GCCCAGTCCCCACACCTACAAGGCCAGCAGATCCCTAATTCTCTCTCCAATCAAGTGCGCTCTCCCCAGCCTGTCCCTTCTCCACGGCCACAGtCCCAG\|C\|CCCCCCACTCCAGTCCTTCCCCAAGGATGCAGCCTCAGCCTTCTCCACACC |
| 2:2216:10851:27216#1#F | \| \| |
| MQ=60 isize=132 C=27 | GCCCAGTCCCCACACCTACAAGGCCAGCAGATCCCTAATTCTCTCTCCAATCAAGTGCGCTCTCCCCAGCCTGtCCCTTCTCCaCGGCCACAGtccCAG\|C\|CCCCCCaCTCCaGTCCTTCCCCAcgGAtgcagGactcCAATCAACCgctgc |
| 2:2216:16653:53291#1#F | \| \| |
| MQ=60 isize=140 C=22 | GCCCAGTCCCCACACCTACAAGGCCAGCAGATCCCTAATTCTCTCTCCAATCAAGTGCGCTCTCCCCAGCCTGTCCCTTCTCCACgGCCACAGtCCCAg\|C\|CCCCCCaCTCCagtCCTTcCCCAAGGATGCAGCCTCAGCCAGGACtCCaat |
| 2:2216:3077:80666#1#F | \| \| |
| MQ=60 isize=179 C=33 | GCCCAGTCCCCACACCTACAAGGCCAGCAGATCCCTAATTCTCTCTCCAATCAAGTGCGCTCTCCCCAGCCTGTCCCTTCTCCACGGCCACAGTCCCAG\|C\|CCCCCCACTCCAGTCCTTCCCCAAGGATGCAGCCTCAGCCTTCTCCACACC |
| 1:2202:15133:34148#2#R | \| \| |
| MQ=60 isize=-128 C=2 | gcccagtcccctctcctacacggccagcagatccctaattctctctccaatcaagtgcgctctccccagcctgtaccttcgccacggccacagtcccag\|c\|ccccccactccagtccttccccaaggatanGACTCCAATTtCGACCCAAAA |
| 1:1108:7767:5295#2#F | \| \| |
| MQ=60 isize=256 C=27 | GGCCATTTCTTGATTGGAaTCntCCCAGTCCCCACACCTACAAGGCCAGCAGATCCCTAATTCTCTCTCCAATCAAGTGCGCTCTCCCCAGCCTGTCCCTTCTCCACGGCCACAGTCCCAG\|C\|CCCCCCACTCCAGtCCtTCCccaaggatg |
| 1:1213:20592:66249#1#F | \| \| |
| MQ=60 isize=137 C=2 | CCCagTCCCCACACCTACAAGGCCAgCaGaTCCCTCATTTtCTCtCCAAtCAAGTGCGCtCTcCCCaacccttcccttctccacggccacagtccccg\|c\|ccccccactcccgtccttccccaaggatgcagcctcaggactccaatgagcc |
| 1:2209:18378:48392#1#F | \| \| |
| MQ=60 isize=206 C=33 | CCCAGTCCCCACACCTACAAGGCCAGCAGATCCCTAATTCTCTCTCCAATCAAGtGCGCTCTCCCCAGCCTGTCCCTTCTCCACGGCCaCAGTCCCAG\|C\|CCCCCCcctCCAGTCCTTCCCCAagGATGCAGCCTCAccCTTCtCCaCACCa |
| 2:1105:4988:38112#1#F | \| \| |
| MQ=60 isize=136 C=27 | CCCAGT\_CCCACACCTACAAGGCCAGCAGATCCCTAATTCTCTCTCCAATCAAGTGcGCTCTCCCCAgCCTGTCCCTTCcCCACGGCcacAGtCCCAg\|C\|CCCCcccccccatcccttccccagggaggcagccccgggcctccaatacccac |
| 2:1207:5637:84950#1#F | \| \| |
| MQ=60 isize=144 C=37 | CCCAGTCCCCACACCTACAAGGCCAGCAGATCCCTAATTCTCTCTCCAATCAAGTGCGCTCTCCCCAGCCTGTCCCTTCTCCACGGCCACAGTCCCAG\|C\|CCCCCCACTCCAGTCCTTCCCCAAGGATGCAGCCTCAGCCTTCTCAGGACTC |
| 2:1215:12948:29055#1#F | \| \| |
| MQ=60 isize=186 C=15 | CCCAGTCCCCACACCTACAAgGCCAGCAGAtCCCTAATTCTCTCTCCaATCAAGtgCgCtCTCCCCAgCCtGtCCCtTCTCCACgGCCACaGtCCCAg\|c\|CCCCCCactcCagtCCtTCCCCaagGatgCaGCCtcagccttctccacaccc |
| 1:1103:20658:33370#2#R | \| \| |
| MQ=60 isize=-132 C=37 | CCCaGcCCCCACa\_CTACAAggCcaGCaGAtCCCTaatTCtCtCtCCAATCAaGtGCgCTCtCCCCAGCCTGTCCCTtCtCCACGGCCACAGTCCCAG\|C\|CCCCCCACTCCAGTCCTTCCCCAAGGATGCAGGACTCCAATCCAGACTCATAC |
| 1:1209:9786:82295#2#R | \| \| |
| MQ=60 isize=-129 C=37 | CCCAGTCCCCACACCTACAAGGCCAGCAGATCCCTAATTCTCTCTCCAATCAAGTGCGCTCTCCCCAGCCTGTCCCTTCTCCACGGCCACAGTCCCAG\|C\|CCCCCCACTCCAGTCCTTCCCCAAGGATGanGACTCCAATAGGAAGCGAAGT |
| 2:2109:10741:61525#2#R | \| \| |
| MQ=60 isize=-129 C=33 | cccagtccccacccctacaaggccagcagaTCCCTAATTCtCTCtCCAATCAAGTGCGCtCTCCCCAGCCTGTCCCTTCTCCACGGCCACAGTCCCAG\|C\|CCCCCCACTCCAGTCCTTCCCCAAGGATGAGGACTCCAATCTTAGTAGTGCC |
| 2:1214:18809:10546#2#F | \| \| |
| MQ=60 isize=255 C=33 | GGTGGCGCAAGCATTGGAGTCCTCCAGTCCCCACACCTACAAGGCCAGCAGATCCCTAATTCTCTCTCCAATCAAGTGCGCTCTCCCCAGCCTGTCCCTTCTCCACGGCCACAGTCCCAG\|C\|CCCCCCaCtCCAgTCCTTCCCCAaggatgc |
| 1:1108:5323:39155#2#R | \| \| |
| MQ=60 isize=-131 C=37 | CCAGTCCCCACACCTACAAGGCCAGCAGATCCCTAAtTCTCTCTCCAATCAAGTGCGCTCTCCCCAGCCTGTCCCTTCtCCACGGCCACAGTCCCAG\|C\|CCCCCCACTCCAGTCCTTCCCCAAGGATGCanGACTCCAATCACTTATATCCA |
| 1:1116:11704:67221#2#R | \| \| |
| MQ=60 isize=-132 C=33 | cCAGTCCCCaCACCTACAAGGCCAGCaGATCCCTAATTCTCtCtCCAAtCAAGTGCGCTCTCCCCAGCCTGTCCCTTCTCCACGGCCACAGTCCCAG\|C\|CCCCCCACTCCAGTCCTTCCCCAAGGATGCAGGACTCCAATTATCGACCTTAG |
| 1:1202:19280:38367#2#R | \| \| |
| MQ=60 isize=-132 C=22 | CCAGTCCcaccaCCTaCAAGGCCAGCAGATCCCTAAttCTCTCTCCAATcAAGTGCGCTCTCCcCAGCCTGTcCCTTCTCCACGGCCACAGTCCCAG\|C\|CCCCCCACTCCAGTCCTTCCCCAAGGATGCAGGACTCCAATAATAAATAGTCT |
| 1:2205:17551:31546#2#R | \| \| |
| MQ=60 isize=-129 C=15 | ccagtccccacacctgcaaggccagcagatccctaattccccctCCAaTcAAGtgCgCtctCCCCAgCcTgtCCCtTCcccACGGCCaCAGTCCcag\|c\|ccCCCCACtCCAGTCCTTCCCCAAGGATGAanGACTCCAATAGCCCaGACCGC |
| 1:2215:16320:32740#2#R | \| \| |
| MQ=60 isize=-132 C=33 | CCAGTCCCcaccCCtacAAGGCCaGCaGATCCCTAATTCtCtCTCCAATCAAGTgCGCTCtCCCCAGCCTGTCCCTTCTCCACGGCCACAGTCCCAG\|C\|CCCCCCACTCCAGTCCTTCCCCAAGGATGCAgnACTCCAATGCTGGCTGTCGG |
| 2:1207:14713:93378#2#R | \| \| |
| MQ=60 isize=-131 C=37 | CCAGTCCCCACACCTACAAGGCCAGCAGaTCCCTAATTCTCTCTCCAATCAAGTGCGCTCTCCCCAGCCTGTCCCTTCTCCACGGCCACAGTCCCAG\|C\|CCCCCCACTCCAGTCCTTCCCCAAGGATGCanGACTCCAATACCGGACGGCAG |
| 2:2110:11998:84536#2#R | \| \| |
| MQ=60 isize=-132 C=15 | ccagtcccCACcCcTacAaGGCCAGcagatCCCTAattctctCtCCAAtCAagtgCGCtCtCCcCAGCCTGTCCCTtcTCCaCgGCCACAGTCCCAG\|c\|CCCCCcaCTCCAGTCCTTCCCCAAGGATGCAGGACTCCAATATTAACAACAGG |
| 2:2113:17927:79254#2#R | \| \| |
| MQ=60 isize=-132 C=33 | CCAGtCCccaCACCTACaAGGCcAGCAGATCCCTAATtCTCtCTCCAATCAAgTGCGCTCTCCCCAGCCTGTCCCTtCTCCACGGCCACAGTCCCAG\|C\|CCCCCCACTCCAGTCCTTCCCCAAGGATGCAgnACTCCAAtCCGAGACGTTGT |
| 2:2206:2269:21344#2#R | \| \| |
| MQ=60 isize=-131 C=33 | ccagtccccccacctacaaGGCCAGCAGATCCCTAATTCTCtCTCCAATCAAGtGCGCtCTCCCCAGCCTGTCCCTTCTCCACGGCCACAGTCCCAG\|C\|CCCCCCACTCCAGTCCTTCCCCAAGGATGCanGACTCCAATGACGTTTAGCCT |
| 2:2209:7559:91865#2#R | \| \| |
| MQ=60 isize=-120 C=27 | gcgttttggccagtccccacacctacaaggCCAGCAgatCCCTAATTCTctCTCCaaTCAaGTGCgCtCTCCcCAgCCTGTCCCTTCtCcaCGGCcaCAGTCCCAG\|C\|CCCCCcaCTCCAGTCCTTCCCanGACTCCAAttCaCTGTCCGCA |
| 2:2216:10851:27216#2#R | \| \| |
| MQ=60 isize=-132 C=22 | ccagtccccaccCCTaCAAGGCCAGCAGATCCcTAAttCtctCTcCAATCAAGtGCgctCtCCCCAGCCTGTCCCTTCTCCACGGCCACAGTCCCAG\|C\|CCCCCCaCTCCAGtCCTTCCCCAAGGATGCAGGACTCCAATCAACCGCTGCGC |
| 2:2201:4897:29685#2#F | \| \| |
| MQ=60 isize=254 C=27 | GTTTCGATCTTAATTGGAGTCntCAGTCCCCACACCTACAAGGCCAGCAGATCCCTAATTCTCTCTCCAATCAAGTGCGCtCTCCCCAGCCTgtCCCTTCtcCACGcCCACAgtCCCAg\|C\|CCCCCCaCtCCAGtccTtCCCCaaggatgca |
| 1:2113:7687:36316#2#R | \| \| |
| MQ=60 isize=-131 C=2 | cagtccccacacctacaaggccagcagatccctaattctctctccaatcaagtgcgctctccccagcctgtcccttctccacggccacagtcccag\|c\|cCCCCCACTCCAGTCCTTCCCCAAGGATGCaannACTCCAATACCTCCCTGCCC |
| 1:2201:4495:54018#2#F | \| \| |
| MQ=60 isize=387 C=33 | CTGGGAAAGAATATTGGAGTCCTCACACCTACAAGGCCAGCAGATCCCTAATTCTCTCTCCAATCAAGTGcGCTCTCCCCAGcctgTcCCTTCTCCACGGCCAcAgTcCCAG\|C\|ccCCCCACTCCAGTCCTTCcCCAaGgATGcagcctcag |
| 1:1111:11890:77129#2#R | \| \| |
| MQ=60 isize=-132 C=22 | AgTCCCcacACCTACAAGGCCAGCagaTCccTAaTTctcTCTCCAaTCAAGTGCGCtCTCCCCAGCCTGTcCcttCTCcaCGGCCaCaGtCCCAg\|C\|CCCCCCaCTCCAGTCCTTCCCCAAGGATGCAGAGGACTCCAATCACCCCTTGCAC |
| 1:1213:13721:35644#2#R | \| \| |
| MQ=60 isize=-132 C=33 | AGTCCCCACaCCtACAAGGCCAGCAGATCCCTAATTCTCtCTCCAAtCAAgtGCGCTCTCCCCaGCCTGtCCCTTCtCCACGGCCACAGTCCCAG\|C\|CCCCCCACTCCAGTCCTTCCCCAAGGATGCAGagnACTCCAATGAACTTCACATA |
| 1:2102:1815:93976#2#R | \| \| |
| MQ=60 isize=-132 C=37 | aGTCCCCACaCCTACAAGGCCAGCAGATCCCTAATTCTCTCTCCAATCAAGTGCGCTCTCCCCAGCCTGTCCCTTCTCCACGGCCACAGTCCCAG\|C\|CCCCCCACTCCAGTCCTTCCCCAAGGATGCAGAGGACTCCAATCACAACCCCCTC |
| 1:2209:7631:22456#2#R | \| \| |
| MQ=60 isize=-132 C=33 | cgtCCCCAcacCtaCAAGGCCAGCAgATCCCTAATTCTCTCTCCAATCAAGTGCGCTCTCCCCAGCCTGTCCCTTCTCCACGGCCACAGTCCCAG\|C\|CCCCCCACTCCAGTCCTTCCCCAAGGATGCAGanGACTCCAATAACTGCGCTGTT |
| 2:1108:10004:9367#2#R | \| \| |
| MQ=60 isize=-132 C=37 | cGTCCCCACACCTAcaAGGCCAGCaGAtCCcTaATTCTCTCTCCAATCAAGTGCgCTCTCCCCAGCCTGCCCCTtCTCCACGGCCACAGTCCCAG\|C\|CCCCCCACTCCAGTCCTTCCCCAAGGATGCAGanGACTCCAATCATTACAACTAA |
| 2:1203:15277:77102#2#R | \| \| |
| MQ=60 isize=-132 C=37 | aGTCCCCACACCTACAAGGCCAGCAGATCCCTAATTCTCTCTCCAATCAAGTGCGCTCTCCCCAGCCTGTCCCTTCTCCACGGCCACAGTCCCAG\|C\|CCCCCCACTCCAGTCCTTCCCCAAGGATGCAGAGGACTCCAATAGGAGACTGTTT |
| 2:2109:8822:4014#2#R | \| \| |
| MQ=60 isize=-132 C=37 | cgtccccacccctacaAGGCCAGCagatCCCTAAtTccCtctCCAATCaAGTgcgCTcTCCCCAGCCTGTCCCTTCcCCACGGCCACAGtCCCAG\|C\|CCCCCCACTCCAGTCCTTCCCCAAGGATGCAGAGGACTCCAATCGCGTTATTTCA |
| 2:2112:15865:47750#2#R | \| \| |
| MQ=60 isize=-132 C=37 | cgtcccCACACCTACAAGGCCAGCAgaTCCcTAATtCtCTCTCcAATCAAgtgCGCTCTCCCCAGCCTGTCCCTTCtCCACGGCCACAGTCCCAG\|C\|CCCCCCACTCCAGTCCTTCCCCAAGGATGCAGAgnACTCCAATTAGACCTTCGAG |
| 2:2204:14595:76151#2#R | \| \| |
| MQ=60 isize=-132 C=33 | aGTCCaCACACCTACaAGGCCAGCAGATCCCTAATtCTCTCTCCAATCAAGTGCGCTCTCCCCAGCCTGTCCCTTCTCCACGGCCACAGTCCCTG\|C\|CCCCCCACTCCAGTCCTTCCCCAAGGATGCAGanGACTCCAATCGCGGTGGCCGG |
| 2:2211:16910:82807#2#R | \| \| |
| MQ=60 isize=-132 C=33 | aGTCCCCACaCCTACAAGGCCAGCAGATCCCTAATtcgCTCTCCAATCAAGTGCGCTCTCCCCAGCCTGTCCCTTCTCCACGGCCACAGTCCCAG\|C\|CCCCCCACTCCAGTCCTTCCCCAAGGATGCAGAGGACTCCAATAACTGCGCTGTT |
| 2:2207:20992:94096#2#F | \| \| |
| MQ=60 isize=252 C=33 | GAGGCTGCAACTATTGGAGTCntGTCCCCACACCTACAAGGCCAGCAGATCCCTAATTCTCTCTCCAATCAAGTGCGCTCTCCCCAGCCTGTCCCTTCTCCACGGCCACAGTCCCAG\|C\|CCCCCCACTCCAGTCCTTCCCCAagGatGCAGC |
| 1:1204:16999:62470#2#R | \|C\| |
| MQ=60 isize=-132 C=33 | gtccccacacctacaaggCCAGCAGATCCCTAattCtCTCtCCAATCAAgtgcGCTccCCCCAGCCTGTCCCTTCTCCACGGCCACAGTCCCAI\|C\|CCCCCCaCTCCAGTCCTTCCCCAAGGATGCAganGACTCCAatCCCCGCCCCCAT |
| 1:2102:2609:57038#2#R | \| \| |
| MQ=60 isize=-133 C=37 | gTCCCCaCaCCTaCAAGGCCAGCAGATCTCTAATtCTCTCTCCAATCAAgTGCGCTCTCCCCAGCCTGTcCCTTCTCCACGGCCACAGTCCCAG\|C\|CCCCCCACTCCAGTCCTTCCCCAAGGATGCAGCAGGACTCCAATGAACTTCAAACG |
| 2:1208:11976:67031#2#R | \| \| |
| MQ=60 isize=-133 C=33 | gtCCCCACACCTACAAGGCCAGCAGATCCCTAATtCTCTCTCCAATCAAGTGCGCTCTCCCCAGCCTGTCCCTTCTCCACGGCCACAGTCCCAG\|C\|CCCCCCACTCCAGTCCTTCCCCAAGGATGCAGCanGACTCCAATGCGCACATCACC |
| 1:1206:19196:22594#2#F | \| \| |
| MQ=60 isize=251 C=33 | TTGGCGGATTAAATTGGAGTCCTCCCCACACCTACAAGGCCAGCAGATCCCTAATTCTCTCTCCAATCAAGTGCGCTCTCCCCAGCCTGTCCCTTCTCCACGGCCACAGTCCCAG\|C\|CCCCCCACTCCAGTCCTTCCCCAaggATGCAGCCc |
| 2:2111:2694:73440#2#F | \| \| |
| MQ=60 isize=251 C=2 | TATGTCAACAGTATTGGAGTCCTCCCCACACCTACAAGGCCAGCAGATCCCTAATTCTCTCTCCAATCAAGtgCGCtctCCCCAccctgccccttctccacggccccattcccag\|c\|cccccccccccacccctcccccaaggaggccgccc |
| 1:1214:18559:41342#2#R | \| \| |
| MQ=60 isize=-134 C=33 | tCCCCACaCCtacaAGGCCaGCAGATCCCTAATTCTCTCTCCAATCAAGTGCGCTCTCCCCAGCCTGTCCCTTCTCCACGGCCACAGTCCCAG\|C\|CCCCCCACTCCAGTCCCTCCCCAAGGATGCAGCCAgnACTCCAATGGCGCGGAGTGG |
| 1:2202:5638:86785#2#R | \| \| |
| MQ=60 isize=-134 C=37 | cccccACACCTACAAGGCCAGCAGAtCCCTAATTCTCTCTCCAATCAAGTGCGCtCTCCCCAGCCTGTCCCTTCTCCACGGCCACAGTCCCAG\|C\|CCCCCCACTCCAGCCCTTCCCCAAGGATGCAGCcanGACTCCAATCATGTCTGGGTG |
| 2:2106:11992:27454#2#R | \| \| |
| MQ=60 isize=-123 C=22 | cttttggccacatccccacacctacaaggccagcagatccctaattctcgccccaaccaagcgccctcgcccCAGCCTGTCcctTCTCCACGGCCACAGTCCCAg\|C\|CCCCCcgCTCCAGTCCTTCCCcanGACTCCAATCGCCCCaCCCCC |
| 2:2108:6453:51342#2#R | \| \| |
| MQ=60 isize=-134 C=33 | cccccccacccacaaggccagcagaTCCCTaattctCTCTCCAAtCAAGTGCgctctccCCAGCCTGTCCCTTCTCCACGGCCACaGTCCCAG\|C\|CCCCCccCTCCAGTCCTTCCCCAAGGATGCAGCCAGGACTCCAATTGCGCCGGGGAG |
| 1:2201:9009:6076#2#R | \| \| |
| MQ=60 isize=-132 C=22 | cgcccccacacctacaaggccagcagATCCCTaattctCTCtCCAAtCaaGTgCGCtCTCCCCAGCCTGTCCCTTCTCCaCgGCCACAGTCCCAG\|C\|cCCCCCACTCCAGTCCTTCCCCAAgGATGCAGAGGACTCCAATAGGAGACTGTTT |
| 1:1103:11926:9660#2#F | \| \| |
| MQ=60 isize=248 C=33 | CGTACCTTGGCGATTGGAGTCCTCCCACACCTACAAGGCCAGCAGATCCCTAATTCTCTCTCCAATCAAGTGCGCTCTCCCCAGCCTGTCCCTTCTCCACGgCCACAGTCCCAG\|C\|CCCCCCaCTCCAGTCCTtCCCCAAGGATgCagCCTC |
| 1:2103:9435:51977#2#F | \| \| |
| MQ=60 isize=249 C=33 | CGGGTGTTCGAAATTGGAGTCCTCCCACACCTACAAGGCCAGCAGATCCCTAATTCTCTCTCCAATCAAGTGCGCTCTCCCCAGCCTGTCCCTTCTCCACGGCCACAGTCCCAG\|C\|CCCCCCACTCCAGTCCTTCCCCAAGGATgCAGCCTC |
| 1:2202:10482:57743#2#F | \| \| |
| MQ=60 isize=249 C=33 | CGCCCGGGTACAATTGGAGTCntCCCACACCTACAAGGCCAGCAGATCCCTAATTCTCTCTCCAATCAAGTGCGCTCTCCCCAGCCTGTCCCTTCTCCACGGCCACAGTCCCAG\|C\|CCCCCCaCTCCagtCCTTCCCCAaGGATgCAGCCcc |
| 2:1101:13225:79545#2#F | \| \| |
| MQ=60 isize=249 C=2 | GCGGTTGAGACAAtTgGagTCCTCCCACAcCTaCaAGgCCaGCAGaTCCCTAATTCTCTCTCCAATCaAgTGCgctctccccagcctgtcccttctccaccgccccagtcccag\|c\|cccccccctcccatccttccccacggctgcgacccc |
| 1:1112:6995:34695#2#R | \| \| |
| MQ=60 isize=-138 C=37 | CCCACaCCTaCAAGGCCAGCAGATCCCTAAttCtCtCtCCAATCAAGTGCGCTCTCCCCAGCCTGTCCCTTCTCCACGGCCACAGTCCCAG\|C\|CCCCCCaCTCCAGTCCTTCCCCAAGGATGCAGCCTCAgnACTCCAATATCGCCCTCCGA |
| 1:2108:4505:87736#2#R | \| \| |
| MQ=60 isize=-137 C=33 | CCCACACCTACAAGGCCAGCAGATCCCTAATTCTCTCTCCAATCAAGTGCGCTCTCCCCAGCTTGTCCCTTCTCCACGGCCACAGTCCCAG\|C\|CCCCCCACTCCAGTCCTTCCCCAAGGATGCAGCCTcanGACTCCAATATGCCGAACAAG |
| 2:1101:7586:52466#2#R | \| \| |
| MQ=60 isize=-138 C=33 | cccacccctccaagcccagctgatccccaattccctccccaatcaaggtccctcccccCAGCCTgtCCCTtctCcaCgGCCaCAGtCCCAG\|C\|CCCCCCcCtCCAGTCCTTCCCCAAGGATGCAGCCTCAGGACTCCAATCGCCACCCTTTA |
| 1:2112:16576:22446#2#R | \| \| |
| MQ=60 isize=-137 C=27 | CCACaCCTaCAAGGCCAGCAGATCCCTAAttCtCtCTCCAATCAAGtGCgCtCTCCCCaGCCtGtCCCTTCTCCACGGCCACAGTCCCAG\|C\|CCCCcCaCTCCAGTCCTTCCCCAAGGATGCAGCCTCAAGGACTCCAAtACGCCGAGGACA |
| 2:2202:12041:22341#2#R | \| \| |
| MQ=60 isize=-137 C=37 | CCACACCTACAAGGCCAGCAGATCCCTAATtCTCTCTCCAATCAAGTGCGCTCTCCCCAGCCTGTCCCTTCTCCACGGCCACAGTCCCAG\|C\|CCCCCCACTCCAGTCCTTCCCCAAGGATGCAGCCTCAAGGACTCCAATACGCCGAGGACA |
| 1:1102:19324:88779#2#R | \| \| |
| MQ=60 isize=-138 C=37 | CACaCCTACAAGGCCAGCAGATCCCTAATtCTCTctCCAATCAAGTgCgCtCTCCCCAGCCTGTCCCtTCTCCACGGCcaCAGTCCCAG\|C\|CCCCCCACTCCAGTCCTTCCCCAAGGATGCAGCCTCAGanGACTCCAATAAAATTTTATTC |
| 1:2202:10673:68770#2#R | \| \| |
| MQ=60 isize=-138 C=33 | cacccctacaaGGCCAGCAGATCCCTAAttCtCtctCCAATCAaGtgCgctCTCCCCAGcCTGTCCCTTCTCCACGGCCaCAGTCCCAG\|C\|CCCCCCaCTCCAGTCCTTCCCCAAGGATGCAGCCTCAGanGACTCCAATCCTCATTGCACT |
| 2:1201:4699:69379#2#R | \| \| |
| MQ=60 isize=-139 C=37 | CACaCCTACAAGGCccGCAGaTCCCCAATtccCtCTCCAATCAAGtGCGctCtCCCCAGCCTGTCCCTtCTCCACGGCCACAGTCCCAG\|C\|CCCCCCACTCCAGTCC\_TCCCCAAGGATGCAGCCTCAGCAGGACTCCAATAGGACGGACATA |
| 2:1211:2252:63209#2#R | \| \| |
| MQ=60 isize=-138 C=33 | CACACCTAcaAgGCCAGCcGaTCCCtaattctCcCTCCAATCAAGtgcgCGCTCCCCAGCCtgtCCCTTCTCCACGGCCACAGTCCCAG\|C\|CCCCCCACTCCAGTCCTTCCCCAAGGATGCAGCCTCAGAGGACTCCAATACATCTTCAAGA |
| 2:2110:8179:2699#2#R | \| \| |
| MQ=60 isize=-139 C=33 | cac\_cctaCAAGGCCAGcaGATCCcTAAttCtCTCtCCAATCAAGTGCgCtCtCCCCAGCCTGTCCCTTCTCCACGGCCACAGTCCCAG\|C\|CCCCCCACTCCAGTCCTTCCCCAAGGATGCAGCCTCAGCAGGACTCCAATTTCCTACCTATT |
| 2:2213:4036:4627#2#R | \| \| |
| MQ=60 isize=-138 C=37 | caccCCtaCAAGgCCAGCAGATCCCTAATTCtCtCTCCAATCAAGTGCgCtCTCCCCAGCCTGTCCCTTCTCCACGGCCACAGTCCCAG\|C\|CCCCCCACTCCAGTCCTTCCCCAAGGATGCAGCCTCAGAGGACTCCAATCACCCGATCCCC |
| 1:1101:1457:16610#2#R | \| \| |
| MQ=60 isize=-139 C=33 | cCaCCTACAaGGCCAGCAGATCCCTAATTCTCTCtCCAATCAAGTGCGCTCTCCCCAGCCTGTCCCTTCTCCACGGCCACAGTCCCAG\|C\|CCCCCCACTCCAGTCCTTCCCCAAGGATGCAGCCTCAGCanGACTCCAATGTGGCCAGCCCC |
| 1:1201:4501:21809#2#R | \| \| |
| MQ=60 isize=-139 C=37 | ccacctacAAGGCcaGCAGatCCCTAATtCTCTCcCCAAtCAAgtgCGCTCtCCCCAGCCTgtcccTtCTCCACGGCCACAGTCCCAG\|C\|CCCCCCACTCCAGTCCTTCCCCAAGGATGCAGCCTCAGCanGACTCCAATCTAAAAGACCAG |
| 1:1206:18404:58961#2#R | \| \| |
| MQ=60 isize=-139 C=37 | cCACCtACAAGGCCAgcAGATCCCTAATTCTCtCTCCAATCaaGTGCgCtCtCCCCAGCCTGTCCCTTCTCCaCGGCCACAGTCCCAG\|C\|CCCCCCACTCCAGTCCTTCCCCAAGGATGCAGCCTCAGCAGGACTCCAATATAGCCCCTACC |
| 1:1215:12143:60200#2#R | \| \| |
| MQ=60 isize=-139 C=27 | ccacctacaaggccagcagatccCTAATtctCtCTCCaatCAAgtGCgCtCtCCCCAGcctgTCCCTTCTCCACGGCCACAGtCcCAG\|C\|CCCCCCAcTCCAGTCCTtCCCCAAGGATGCAGcCTCAGCAGGACTCCAATAGGACGgACATA |
| 1:2110:19393:48120#2#R | \| \| |
| MQ=60 isize=-139 C=22 | aCaCCTACAAGGCCAGCAGATCCCTAATTCTCTCTCCAATCAAGTGCGCtCTCCCCAGCCTGTCCCTtCTCCACGGCCaCAGTCCCAG\|C\|CCCCCCACTCCAGTCCTTCCCCAAGGATGCAGCCTCAGCAGGACTCCAATCAAGGACCAGTT |
| 1:2206:7485:84245#2#R | \| \| |
| MQ=60 isize=-139 C=37 | ccaCCtAcaAGGCCAgtcGATCCCTAATTCtCTCTCCAATCaAGTGCGCtCTCCCCAGCCTGTCCCTTCTCCaCGGCCACAGTCCCAG\|C\|CCCCCCACTCCAGTCCTTCCCCAAGGATGCAGCCTCAGcanGACTCCAATATAGCCCCTACC |
| 2:1110:17283:62010#2#R | \| \| |
| MQ=60 isize=-139 C=37 | acaCCTACAAGgcCAGCAgatCCCTAattCtctCTCCAATCAAGtGCGCtCTCCCCAGCCtGtCCCTTCtCCACGGCCACAGTCCCAG\|C\|CCCCCCACTCCAGTCCTTCCCCAAGGATGCAGCCTCAGCAGGACTCCAATCGATACTCCTTC |
| 2:1201:2619:65873#2#R | \| \| |
| MQ=60 isize=-139 C=27 | ccacctccaaggccagcagatCCCTaATTCtCTCtccaATCaagtGCGccCTcCCCAGcctGtCccttctaCaCggCcaCAGTCCCAG\|C\|CccCccgCTCCAGTCCTTCCCCAaGGATGCAGCCTCAGCAGGACTCCAATCCGGGGGTCTGT |
| 2:2214:15686:26053#2#R | \| \| |
| MQ=60 isize=-139 C=33 | ccaCCTACAAGGCCAGCAGAtCCCTAATtctCTCTcCAATCAAGTGCgctCtCCCCAGCCTGTCCCTTCTCCACGGCCACAGTCCCAG\|C\|CCCCCCaCTCCAGTCCTTCCCCAAGGATGCAGCCTCAGCAGGACTCCAATGGTGGCCACTAA |
| 1:2209:9365:53418#2#F | \| \| |
| MQ=60 isize=245 C=27 | ATGCGGCTGCGCATTGGAGTCntCACCTACAAGGCCAGCAGATCCCTAATTCTCTCTCCAATCAAGTGCGCTCTCCCCAGCCtgtCCCTTCTCCaCAGCCacAGtCCCAG\|C\|CCCCCCaCtCCaGtcCTTCCCCAAGGATGCAGCCTCAGCC |
| 2:2110:7226:91087#2#F | \| \| |
| MQ=60 isize=245 C=33 | TGAGTAGGGGTGATTGGAGTCCTCACCTACAAGGCCAGCAGATCCCTAATTCTCTCTCCAATCAAGTGCGCTCTCCCCAGCCTGTCCCTTCTCCaCGGCCACAGTCCCAG\|C\|CCCCCCactCCAGTCCTTCcccaaggatgcagcctcaccc |
| 1:1205:10704:50798#2#R | \| \| |
| MQ=60 isize=-140 C=33 | CaCCTacaaggCCAGCAGATCCCTAatcctCTCtccaATCAAGtGCgctctCCCCAGCCtgTCCcTTCTCCACGGCCACAGTCCCAG\|C\|CCCCCCACTCCAGTCCTTCCCCAAGGATGCAGCCTCAGCCanGACTCCAATTACCGTCAGACC |
| 1:1206:19084:65287#2#R | \| \| |
| MQ=60 isize=-140 C=27 | CAcctaCAAGgCCAGCcGAtCCCTAATTCTCTctCCAATCAaGtgCgctctcCCCAGCCTGTCCCTtCTCCACGGCCACAGTCCCAG\|C\|CCCCCCACTCCAGTCCTTCCCCAAGGATGCAGcCtCAGCCAGGACTCCAATAGCACGTCGTAG |
| 1:2109:18423:39446#2#R | \| \| |
| MQ=60 isize=-140 C=2 | cacctccaaggccagcagatccctaattctcccgccaatcaagtgcgcgcaccccagactgccccttctccacggccacagtcccgg\|c\|ccccccactccagtcctcccccaaggatgcagcctcagccanGACTCCAATTACCgTCAgaCC |
| 1:2112:4752:52767#2#R | \| \| |
| MQ=60 isize=-140 C=37 | CACCTACAAGGCCAGCAGATCCCTAATTCTCTCTCCAATCAAGTGCGCtCTCCCCAGCCTGTCCCTTCTCCACGGCCACAGTCCCAG\|C\|CCCCCCACTCCAGTCCTTCCCCAAGGATGCAGCCTCAGCCAGGACTCCAATCCCGGTCCGTAC |
| 2:1104:21059:95595#2#R | \| \| |
| MQ=60 isize=-140 C=33 | cccctacacggccagcagatccctacttctctCTCCAATCAagTGCcCTCtCCCCAGCCTgcCCCTTCtCCACGgCCACAGTCCCAG\|C\|CCCCCCACTCCAGTCCTTCCCCAAGGATGCAGCCTCAGCCAGGACTCCAATTGCGGGGTAGCC |
| 2:1115:4222:40659#2#R | \| \| |
| MQ=60 isize=-140 C=37 | cacccacaAgGCcagcaGatCCCTAATTctCTCtCCAAtCAAGtgcgctCtCCCCAGCCtGTCCCTTCtCCACGGCCACAGtCCCAG\|C\|CCCCCCACTCCAGTCCTTCCCCAAGGATGCAGCCTCAGCCAgnACTCCAATACCCAGAAATCC |
| 2:2205:18951:10568#2#R | \| \| |
| MQ=60 isize=-140 C=37 | caCCTACAGGGCCAGCAGaTCCCTAATtCTCtCTCCAATCAAGTGCGCTCTCCCCAGCCTGTCCCTTCTCCACGGCCACAGTCCCAG\|C\|CCCCCCACTCCAGTCCTTCCCCAAGGATGCAGCCTCAGCCanGACTCCAATTGTGCCCATCTA |
| 2:1107:8591:28730#2#R | \| \| |
| MQ=60 isize=-141 C=33 | ccctacAAGGCCAGCagatcCCtaattCtCtccCCAATCAAgtgCGCtCtCCCCAGCCtGtCcCTTCtCCACGGCCACAGTCCCAG\|C\|CCCCCCACTCCAGTCCTTCCCCAAGGATGCAGCCTCAGCCTanGACTCCAATGCTAGGGGGATT |
| 1:2114:4080:55959#2#F | \| \| |
| MQ=60 isize=243 C=33 | TTCGTAAGCTCTATTGGAGTCCTACAAGGCCAGCAGATCCCTAATTCTCTCTCCAATCAAGTGCGCTCTCCCCAGCCTGTCCCTTCTCCACGGCCACAGTCCCAG\|C\|CCCCCCACTCCAGTCCTTCCCCAAGGATGCAGCCTCAGCCTTCTC |
| 1:1104:15138:76312#2#R | \| \| |
| MQ=60 isize=-139 C=37 | cccCCTACAAGGCCAGCAGATCCCTAAttCTCTCTCCAATCAAGTgCGCcCCCCCCAGCCTGTCCCTTCTCCACGGCCACAGTCCCAG\|C\|CCCCCCACTCCAGTCCTTCCCCAAGGATGCAGCCTCAGCanGACTCCAATCTGGGGGCGGCG |
| 1:2202:19940:87181#2#R | \| \| |
| MQ=60 isize=-133 C=33 | gtcccccgtcctacaaggccagcaggtccCTAatcCTCTCTCCAATCAagtgCgctCTCCCCAGCCTGTCCCTtctccaCGGCCaCAGTCCCaG\|C\|CCCCCCaCTCCAGTCCTTCCCCAAGGATGCAGCanGACTCCAATCTCTGCTTAGCG |
| 2:1101:19266:25045#2#R | \| \| |
| MQ=60 isize=-137 C=22 | caacccctacaaggccagcagatccctaattctcgctCCAATCAAGtgCgCTCTCCCCAGCCTgtcccttCtCCaCGGCCACAGTCCCAG\|C\|CCCCCCaCTCCAGTCCTTCCCCAAGGATGCAGCCTCAAGGACTCCAATCCTGCGCCTCAA |
| 1:1109:7199:2973#2#F | \| \| |
| MQ=60 isize=242 C=33 | GTTGTGATTTGTATTGGAGTCCTCTACAAGGCCAGCAGATCCCTAATTCTCTCTCCAATCAAGTGCGCTCTCCCCAGCCTGTCCCTTCTCCACGGCCACAGTCCCAG\|C\|CCCCCCACTCCAGTCCTTCCCCAAGGATGCAGCCTCAGCCTTC |
| 1:2103:3877:15779#2#F | \| \| |
| MQ=60 isize=242 C=33 | TGAATTATGTCCATTGGAGTCCTCTACAAGGCCAGCAGATCCCTAATTCTCTCTCCAATCAAGTGCGCTCTCCCCAGCCTGTCCCTTCTCCACGGCCACAGTCCCAG\|C\|CCCCCCACTCCAGTCCTTCCCCAAGGatGCaGCCTCAGCCTTC |
| 1:1114:19667:9240#2#R | \| \| |
| MQ=60 isize=-144 C=27 | CtaCAaggCCAGCagaTCCCTAATTCtCTCcCCAATCAAGTGCGCTctcCCCAGCCTGTcCCTTCtCCACGGCCACAGTCCCAG\|C\|CCCCCCACTCCAGTCCTTCCCCAAGGATGCAGCCTCAGCCTtCTAGGAcTCCATTAGTGAGCCGGC |
| 1:1215:5872:31777#2#R | \| \| |
| MQ=60 isize=-143 C=33 | CTACAAGGCCAGCAGATCCCTAATTCTCtCTCCAATCAAGtGCGCTCTCCCCAGCCTGTCCCTTCTCCACGGCCACAGTCCCAG\|C\|CCCCCCACTCCAGTCCTTCCCCAAGGATGCAGCCTCAGCCTTCAGAACTCCAATCCCCAACTGGCC |
| 2:1112:12335:83569#2#R | \| \| |
| MQ=60 isize=-143 C=37 | CtACAAGGCCAGCAGATCCCTAAttCtCtCtCCAATCAAGTGCGCtCTCCCCAGCCtGTCCCttCTCCACGGCCACAGTCCCAG\|C\|CCCCCCACTCCAGTCCTTCCCCAAGGATGCAGCCTCAGCCTTCAGGACTCCAATACCGGCAGCTCA |
| 2:2204:6229:29060#2#R | \| \| |
| MQ=60 isize=-143 C=33 | CTACaagGCCAGCaGATCCCTAATtctcTccCCAatCAAGTGcgctccCCCCAGCCTGtCCCTTCtCCaCGGCCACAGTCCCAG\|C\|CCaCaCaCTCCAGTCCTTCCCCAAGGATGCAGCCTCAGCCTTCanGACTCCAATTGACAGGACCCC |
| 2:2107:19381:49645#2#F | \| \| |
| MQ=60 isize=241 C=22 | GTCATTCGGGAAATTGGAGTCCTTACAAGGCCAGCAGATCCCTAATTCTCTCTCCAATCAAGTGCGCTCTCCCCAGCCTGTCCCTTCTCCACGGCCaCAGTcCCAg\|C\|CCCCCCACTCCAGTCCTTCCCCaaggaTgCAGCCTCAgCCTTCc |
| 2:2212:20376:44659#2#F | \| \| |
| MQ=60 isize=241 C=37 | GACACTACCTGGATTGGAGTCCTTACAAGGCCAGCAGATCCCTAATTCTCTCTCCAATCAAGTGCGCTCTCCCCAGCCTGTCCCTTCTCCACGGCCACAGTCCCAG\|C\|CCCCCCACtCCAGTCCCTCCCCAAGGaTGCAGCCTCAGCCttct |
| 1:1106:12819:16097#2#R | \| \| |
| MQ=60 isize=-144 C=37 | taCAAGGCCAGCAGATCCCTaATTCtCTCTCCAATCAAGTGCGCTCTCCCCAGCCTGTCCCTTCTCCACGGCCACAGTCCCAG\|C\|CCCCCCACTCCAGTCCTTCCCCAAGGATGCAGCCTCAGCCTTCTanGACTCCAATACATTACCTGGT |
| 1:1201:1982:35691#2#R | \| \| |
| MQ=60 isize=-144 C=15 | tACAAGGCCAgCAgaTCCCTAATTCtctCTCCAATCAagtGCgccctCccCAGCCTGTCccTtCTCCACGGCCACAGTCCCAG\|c\|CCCCCCaCTCCAGTCCTTCCCCAAGGATGCAGCCTCAGCCTTCTanGACTCCAATCAAGGTATTTCT |
| 1:2110:9428:10907#2#R | \| \| |
| MQ=60 isize=-144 C=33 | TACAAGGCCGGCAGATCCCTAATTCTCTCTCCAATCAAgtGCGCtCTCCCCAGCCTGtCCCTTCTCCACGGCCACAGTCCCAG\|C\|CCCCCCACTCCAGTCCTTCCCCAAGGATGCAGCCTCAGCCTTCTAGGACTCCAATCCTTGCCCCTGC |
| 2:1110:15318:20030#2#R | \| \| |
| MQ=60 isize=-144 C=33 | tACAAGGCCAGCAGATCCCTAaTTCTCTCTCCAATCAAGTGCGCTCTCCCCAGCCTGTCCCTTCTCCACGGCCACAGTCCCAg\|C\|CCCCCCACTCCAGTCCTTCCCCAAGGATGCAGCCTCAGCCTTCTAGGACTCCAATTATTTAACGTCG |
| 2:1205:8478:45470#2#R | \| \| |
| MQ=60 isize=-144 C=33 | taCAaGGCCAGCAGAtCcCtAATTCTCTCTCCAATCAAGTGCGCtCTCCCCAGCCTGTCCCTTCTCCACGGCCACAGTCCCAG\|C\|CCCCCCACTCCAGTCCTTCCCCAAGGATGCAGCCTCAGCCTTCTanGACTCCAATCGACGCCGCCGC |
| 2:2107:11976:28503#2#R | \| \| |
| MQ=60 isize=-144 C=27 | cacaaggccagcagatCCCTAAttccctctCCAATCaAGtgcgctCtCCCCAGCCTGtCCCTTCTCCACGGCCACAGTCCCAG\|C\|CCCCCCACTCCAGTCCTTCCCCAAGGATGCAGCCTCAGCCTTCTAGGACTCCAATTAGTTTTCCAGC |
| 2:2208:13727:10451#2#R | \| \| |
| MQ=60 isize=-144 C=37 | taCAAgGCCAGCAGATCCCTAATTCTCtCTCCAATCAAGTGCGCtCTCCCCAGCCTGTCCCTTCTCCATGGCCACAGTCCCAG\|C\|CCCCCCACTCCAGTCCTTCCCCAAGGATGCAGCCTCAGCCTTCTanGACTCCAATCCAAACCTAGAC |
| 1:1105:11184:49264#2#R | \| \| |
| MQ=60 isize=-145 C=37 | aCAAGGCCAGCAGATCCCTAATtCTCTCTCCAATCAAGTGCGCTCTCCCCAGCCTGTCCCTTCTCCACGGCCACAGTCCCAG\|C\|CCCCCCACTCCAGTCCTTCCCCAAGGATGCAGCCTCAGCCTTCTCanGACTCCAATCGCAAAGATGTA |
| 1:1116:13315:10469#2#R | \| \| |
| MQ=60 isize=-145 C=27 | aCAAggCCAGCAGATCCCTAATTCtCTCTCCAATCAAgTGCGCTCTCCCCAGCCTGTCCCTTCTCCACGGCCACAGTCCCAG\|C\|CCCCCCACTCCAGTCCTTCCCCAAGGATGCAGCCTCAGCCTTCTCAGGACTCCAATGGTTCATAACGT |
| 1:2104:16016:49868#2#R | \| \| |
| MQ=60 isize=-145 C=33 | cCaaggCCAgCagatCCCTaattctctCtCcaAtCAAGTgCgctCtCCCCAGCCTGTCCCTtCTCCACGGCCACAGTCCCAG\|C\|CCCCCCACTCCAGTCCTTCCCCAAGGATGCAGCCTCAGCCTTCTcanGACTCCAATTCACGTATCGAA |
| 2:1209:20691:20719#2#R | \| \| |
| MQ=60 isize=-145 C=33 | acaaggCCAGCAGATCccTaatTCtCtCTCcAatCAagtgCgctCtCCCCAGCctgTCCCTTCtCCACGGCCACAGTCCCAg\|C\|CCCCCCACccCAGTCCTTCCCCAAGGATGCAGCCTCAGCCTTCTCAGGACTCCAATGGTTCATAACGT |
| 2:2109:8660:82689#2#R | \| \| |
| MQ=60 isize=-145 C=33 | ccaaGGCCAGcaGatCCcTAAttccctCTCCAatCaaGtgCgCTCtCCccAgCccgTCCCTTCTCCaCGGCCACAGTCCCaG\|C\|CCCCCCACTCCAGTCCTTCCCCAAGGATGCAGCCTCAGCCTTCTCAGGACTCCAATTAGACATCGGCT |
| 2:2203:7779:33474#2#R | \| \| |
| MQ=60 isize=-145 C=37 | ACAaGGCCAGCAGATCCCTAAttCcCTCTCCAATCAAgTgCGCcCTCCCCAGCCTGTcCCtTCTCCaCGGCCACAGTCCCAG\|C\|CCCCCCACTCCAGTCCTTCCCCAAGGATGCAGCCTCAGCCTTCTCanGACTCCAATCTCGACTCATAC |
| 2:2215:18972:53510#2#R | \| \| |
| MQ=60 isize=-145 C=33 | ccaaggccagcagATCCCtAattctCTCTCCAATCAAgtgCgCTctCcccAGCCTGcCCCttCtCCACGGCCAcaGtCCCAG\|C\|CCCCCCaCTCCAGTCCTTCCCCAAGGATGCAGCCTCAGCCTTCTCAGGACTCCAATGGACCCTGGCGC |
| 1:1102:2276:41225#2#R | \| \| |
| MQ=60 isize=-147 C=37 | CAAGGCcAGCAGAtCCCTAAttcTCTCTCCAATCAAGTGCGCtCTCCCCaGCCTGtCCCTTCtCCACgGCCACAGTCCCAG\|C\|CCCCCCACTCCAGTCCTTCCCCAAGGATGCAGCCTCAGCCTTCTCCanGACTCCAATTAGCCCAGACAC |
| 1:1204:18949:2088#2#R | \| \| |
| MQ=60 isize=-147 C=33 | cAAGGCCaGCAGATCCCTAATTCtCTCTCCAatCAAGTGCGCTCTCCCCAGCCTGTCCCTTCTCCaCGGCCACAGTCCCAG\|C\|CCCCCCACTCCAGTCCTTCCCCAAGGATGCAGCCTCAGCCTTCTCCanGACTCCAATCTCTCAGCCCCA |
| 1:1207:5173:42104#2#R | \| \| |
| MQ=60 isize=-147 C=33 | caagGCCAgCAGaTCCCTaaTtCTCtCTCcAATCAAGTgCgCtCTCCCCAgCCTGTCCCTTcTCCACGGCCACAGTCCCAG\|C\|CCCCCCACTCCAGTCCTTCCCCAAGGATGCAGCCTCAGCCTTCTCcanGACTCCAATGAATAAAAAACT |
| 1:1213:7418:63630#2#R | \| \| |
| MQ=60 isize=-147 C=37 | CAAGGCCAGCAGaTCCCTAATTCTCTCTCCAATCAAGTGCGCTCTCCCCAGCCTGTCCCTTCTCCACGGCCACAGTCCCAG\|C\|CCCCCCACTCCAGTCCTTCCCCAAGGATGCAGCCTCAGCCTTCTCCAgnACTCCAATCTTCCACCGTCC |
| 1:2106:15392:66430#2#R | \| \| |
| MQ=60 isize=-147 C=37 | CAAGGCCAGCAGATCCCTAATTCTCTCTCCAATCAAGTGCGCTCTCCCCAGCCTGTCCCTTCTCCACGGCCACAGTCCCAG\|C\|CCCCCCACTCCAGTCCTTCCCCAAGGATGCAGCCTCAGCCTTCTCCanGACTCCAATCAGGTCTGCTAG |
| 1:2205:11507:8395#2#R | \| \| |
| MQ=60 isize=-147 C=2 | caaggcccgcagatccctaattctctctccaatcaagtgcgctctccccagcctgtcccttctccacggccacagtcccag\|c\|ccccccactcCAGTCCTTCCCCAAGGATGCAGCCTCAGCCTTCTccanGACTCCAATAGGCGGGCTGCG |
| 1:2215:13177:64134#2#R | \| \| |
| MQ=60 isize=-147 C=27 | caaGGCCAgcaGATCCCTAAttctCTCtCCAATCAAGTGCGCTCTCCCCAgCctGTCCcTTCTCcaCGGCCACAGTCCCAG\|C\|CCCCCccCTCCAGTCCTTCCCCAAGGATgCAGCCTCAGCCTTCTCCAgnACTCCAATGAGAGGCGCTTT |
| 1:2216:15934:12095#2#R | \| \| |
| MQ=60 isize=-147 C=33 | CAAggCCAGCAGATCCCTAATTCtCtCTCCAATCAAGtGCgctCTCCCCAGCCTGTCCCTTCTCCACGGCCACAGTCCCAG\|C\|CCCCCCACTCCAGTCCTTCCCCAAGGATGCAGCCTCAGCCTTCTCCAgnACTCCAATGACGCTCCACCA |
| 2:1107:3981:71789#2#R | \| \| |
| MQ=60 isize=-127 C=33 | tggcccaggccccaccccctcaaggccaacagatccctaattccccctccaatcaagtgcgccctcCCCaGCCTGTCCCTTctcCACGGCCaCAGTCCCAG\|C\|CCCCCCACTCCAGTCCTTCCCCAAGganGACTCCAATGCGACCAATCCC |
| 2:1109:14393:45764#2#R | \| \| |
| MQ=60 isize=-147 C=27 | caaggccagcagatccctaattcccgccccaatCAAGTGCgctCTCCCCaGCCTGTCCCTTCgCCACGGCCACAGTCCCAG\|C\|CCCCCCACtCCAGTCCTtCCCCAAGGATGCAGCCTCAGCCTTCTCCanGACTCCAAtCCAACGTCCGCT |
| 2:1116:21213:11262#2#R | \| \| |
| MQ=60 isize=-147 C=33 | caaggccagcagacccctaatcctccctccaatcaagtcccctctccccagcctcccccttccccacggCCACAgcCCcag\|C\|CCCCCCaCtCCagcCCttCCCCAAgGAtGCAgCCtCAgCCTtCTCCAGGACTCCAATTGCAtCgaCCGC |
| 2:2106:14935:5406#2#R | \| \| |
| MQ=60 isize=-147 C=37 | caaggcccgcagatccctaattctctctccaatcAAGTGCgCtCtCCCCAGCCTGTCCCTTCTCCACGGCCACAGTCCCAG\|C\|CCCCCCACTCCAGTCCTTCCCCAAGGATGCAGCCTCAGCCTTCTCCanGACTCCAATGACGCTCCACCA |
| 2:2115:11701:68588#2#R | \| \| |
| MQ=60 isize=-147 C=37 | CAAGGCCAGCAGATCCCTAATtCTCTCTCCAATCAAGtGCGCtCTCCCCAGCCTGTCCCTTCTCCACGGCCACAGTCCCAG\|C\|CCCCCCACTCCAGTCCTTCCCCAAGGATGCAGCCTCAGCCTTCTCCAgnACTCCAATCGGTGGCATTCA |
| 2:1104:12641:87637#2#R | \| \| |
| MQ=60 isize=-132 C=33 | cgtcccccaccctaaaaggccagcaGATCCCTaattctccCtCCAAtCAagtGccCtCTCCCCAGCCTgtCCCTTCTCCACGGCCACAGTCCCAG\|C\|CCCCCCACTCCAGTCCTTCCCCAAGGATGCAGAGGACTCCAATAGCGTGTCCACT |
| 1:1107:15065:12644#2#R | \| \| |
| MQ=60 isize=-149 C=27 | AGGCCAGCAGATCCCTAATtCTCTCTCCAAtCAaGtGCGCtCTCCCCaGCCTGTCCCttCTCCAcGGCCACAGTCCCAG\|C\|CCCCCCACTCCAGTCCTTCCCCAAGGATGCAGCCTCAGCCTTCTCCACAGGACTCCAATAGGGTCGGTTCG |
| 1:1115:10303:100545#2#R | \| \| |
| MQ=60 isize=-149 C=37 | AGGCCAGCAGatCCCTAATTCTCTCTCCAATCAAGtGCGCTCTCCCCAGCCTGTCCCTTCTCCACGGCCACAGTCCCAG\|C\|CCCCCCACTCCAGTCCTTCCCCAAGGATGCAGCCTCAGCCTTCTCCACAGGACTCCAATCTGGGCCACGCT |
| 1:2212:8739:4624#2#R | \| \| |
| MQ=60 isize=-149 C=37 | AGGCCAGCAGATCCCTAATtCTctCTCCAAtCAAGTgCgCtCTCCCCAGCCTGTCCCTTCTCCACGGCCACAGTCCCAG\|C\|CCCCCCaCTCCAGTCCTTCCCCAAGGATGCAGCCTCAGCCTTCTCCACAGGACTCCAATGGAGCACACTTG |
| 2:1113:12446:17821#2#R | \| \| |
| MQ=60 isize=-149 C=2 | aggcctgcacatccctaattctatctccaatcaagtccgctctccccagcctggcccttcaccgcggccacagtcccag\|c\|ccccccactccagtccttccccaaggatgcagcctcagccttCTCCACAgnACTCCAATcgCCACcgcTtc |
| 2:1205:17586:69841#2#R | \| \| |
| MQ=60 isize=-149 C=37 | AgGCCAGCAGAtCCCTAaTTCTCTCtCCAATCAAGtGCGCtCTCCcCAGCCTGTCCCTTCTCCACGGCCACAGTCCCAG\|C\|CCCCCCACTCCAGTCCTTCCCCAAGGATGCAGCCTCAGCCTTCTCCACanGACTCCAATCTGGGCCACGCT |
| 2:1214:9857:81920#2#R | \| \| |
| MQ=60 isize=-149 C=33 | aggccaGCAGAtcCCTAATtCtCtccCCAATCAAGTgCgCTctCCCCAGCCTGTCCCTTCTCCACGGCCACAGTCCCAG\|C\|CCCCCCaCTCCAGTCCTTCCCCAAGGATGCAGCCTCAGCCTTCTCCACAGGACTCCAATTCAGAGAACATA |
| 2:2106:12485:25740#2#R | \| \| |
| MQ=60 isize=-144 C=33 | caccaggccagcagatccctaattctccccccaatcaagtgcgctctcCCCAGCccGTCCCttctCCACGGCcaCAGTCCCAG\|C\|CCCCCCACTCCAGTCCTTCCCCAAGGATGCAGCCTCAGCCTTCtanGACTCCAATTATTTAACGtCG |
| 2:2116:15977:51515#2#R | \| \| |
| MQ=60 isize=-149 C=37 | aggCCAgCaGaTCCCTAATTCTCtCTCCAATCAagtgCGCtCTCCCCAGCCTGTCCCTTCTCCACGGCCACAGTCCCAG\|C\|CCCCCCACTCCAGTCCTTCCCCAAGGATGCAGCCTCAGCCTTCTCCACAgnACTCCAATATAATAGCGCAG |
| 2:2208:17586:4661#2#R | \| \| |
| MQ=60 isize=-149 C=37 | aGGCCaGcAGatCcCTAattCTCTCTCCAATCAAGTGCGCTCTCCCCAGCCTGTCCCTTCTCCACGGCCACAGTCCCAG\|C\|CCCCCCACTCCAGTCCTTCCCCAAGGATGCAGCCTCAGCCTTCTCCACanGACTCCAATCACACCACCGGG |
| 1:2112:8846:78970#2#F | \| \| |
| MQ=60 isize=236 C=27 | AGGTGTAGCGGGATTGGAGTCCTGGCCAGCAGATCCCTAATTCTCTCTCCAATCAAGTGCGCTCTCCCCAGCCTGTCCCTTCTCCACGGCCACAGTCCCAG\|C\|CCCCCCACTCCAGTCCTTCCCCAAGGaTGCAGCCTCAGCCTtCTCCacc |
| 2:1107:3468:65692#2#F | \| \| |
| MQ=60 isize=236 C=2 | GTTCTTTTATGTATTGGAGTCntGGCCAGCAGATCCCTAATTCTCTCTCCAATCAAgtgCGCTCTCCCCAgCCTGtCCCTTCcCCAcGgcCACAGtcccac\|c\|ccccccactccagtccttccccaagactgcaccctcacccttccccccc |
| 1:1204:11014:8931#2#R | \| \| |
| MQ=60 isize=-149 C=2 | cgccaccagatccctaattctctctccaaccaagtgcgctctccccagcctggcccttctccccggccacagtcccag\|c\|ccccccactccagtccttccccaaggatgcagcctcagccttctccacaangactccaattgccctcccccc |
| 2:1109:7041:47650#2#R | \| \| |
| MQ=60 isize=-149 C=37 | agccagcagatccctaattctccctccaatcaagtgcgctctCCCCAGCctgtCCCTTCtCCAcGgCCACaGtCcCAG\|C\|CCCCCCACTCCAGTCCTTCCCCAAGGATGCAGCCTCAGCCTTCTCCACAanGACTCCAATGGCCACACACGT |
| 2:1111:3652:35097#2#R | \| \| |
| MQ=60 isize=-134 C=33 | gccccccacctacggggccagcagcTCCCTAATTCtCtCTCCAATCAAgtgcgctCtCCCCAGCCTGTCCCTTCTCCACGGCCACAGTCCCAG\|C\|CCCCCCACTCCAGTCCTTCCCCAAGGATGCAGCCAGGACTCCAATGGCGCGGAGTGG |
| 1:1112:4027:2261#2#F | \| \| |
| MQ=60 isize=235 C=33 | GTTACATTTGCCATTGGAGTncTGCCAGCAGAACCCTAATTCTCTCTCCAATCAAGTGCGCTCTCCCCAGCCTGTCCCTTCTCCACGGCCACAGTCCCAG\|C\|CCCCCCACTCCAGTCCTTCCCCAAGGATGCAGCCTCAgCCTTCCCCaCAC |
| 1:1212:11955:56351#2#F | \| \| |
| MQ=60 isize=235 C=37 | CCGACGGCCCTGATTGGAGTCCTGCCAGCAGATCCCTAATTCTCTCTCCAATCAAGTGCGCTCTCCCCAGCCTGTCCCTTCTCCACGGCCACAGTCCCAG\|C\|CCCCCCACTCCAGTCCTTCCCCAAGGATGCAGCCTCAGCCTTCTCCACAC |
| 1:1210:14837:52317#2#R | \| \| |
| MQ=60 isize=-150 C=27 | gCCGGCaGATCCCTAAttctctCTCCaatCAAGTGCgCTCTCCCCAGCCtGTCCCTtCtCCACGGCCACAGTCCCAG\|C\|CCCCCCACTCCAGTCCTTCCCCAAGGATGCAGCCTCAGCCTTCTCCACACAGGACTCCAATGGCAACTGCGCA |
| 1:2212:21014:3744#2#R | \| \| |
| MQ=60 isize=-149 C=33 | gCCAGCAGATCCCTAATtCTCTCTCCAATCAAGTGCGCTCtCCCCAGCCTgTCCCTTCTCCaCGGCCACAGTCCCAG\|C\|CCCCCCACTCCAGTCCTTCCCCAAGGATGCAGCCTCAGCCTTCTCCACACAGGACTCCAATTCACACGACGCG |
| 2:2103:13128:53412#2#R | \| \| |
| MQ=60 isize=-150 C=37 | gccagcagatccctaattccccccccaatcaagggccctccCCCCAGCCtgtCCCTTCtCCACGGCCACAGTCCCAG\|C\|CCCCCCACTCCAGTCCTTCCCCAAGGATGCAGCCTCAGCCTTCTCCACACAGGACTCCAATTCACACGACGCG |
| 1:2201:1890:16792#2#F | \| \| |
| MQ=60 isize=234 C=33 | AGTGGTCTAGAGATTGGAGTCCTCCAGCAGATCCCTAATTCTCTCTCCAATCAAGTGCGCTCTCCCCAGCCTGTCCCTTCTCCACGGCCACAGTCCCAG\|C\|CCCCCCACTCCAGTCCTTCCCCAAGGATGCAGCCTCAGCCTTCTCCACACC |
| 2:2107:2698:95155#2#F | \| \| |
| MQ=60 isize=234 | GATCCGCAGATGATTGGAGTCCTCCAGCAGATCCCTAATtCTCTCTCcAATCAAgtgCGcTCTCCccagcctgtcctttctccacggctaaagcccaac\|c\|cccccccccccactgctccccacagaagccagccccggacttccccccacc |
| 1:1208:18571:34893#2#F | \| \| |
| MQ=60 isize=233 C=33 | CATATCACTGCAATTGGAGTCntCAGCaGATCCCCAATTCTCTCTCCAATCAAGTGCGCTCTCCCCAGCCTGTCCCTTCTCCACGGCCACAGTCCCAG\|C\|CCCCCCACTCCAGTCCTTCCCCAAGGATGCAGCCTCAGCCTTCTCCACACCC |
| 2:1116:15870:96100#2#F | \| \| |
| MQ=60 isize=233 C=27 | CATATCACTGCAATTGGAGTCCTCAGCAGATCCCTAATTCTCTCTCCAATCAAGTGCGCTCTCCCCAGCCTGTCCCTTCTCCACGGCCACAGTCCCAg\|C\|cCCCCCACTCCAGTCCTTCCCCAAGGATGCAGCCTCAGCCTTCTCCACACCc |
| 2:2104:19379:52829#2#F | \| \| |
| MQ=60 isize=233 C=27 | CTACCGGGGATAATTGGAGTCCTCAGCAGATCCCTAATTCTCTCTCCAATCAAGTGCGCTCTCCCCAGCCTGTCCCTTCTCCACGGCCACAGTCCCAg\|C\|CccCCCACTCCAGTCCTTCCCCAAgGATgCAGCCTCAGCCTTCTCCacaccc |
| 2:2215:7742:78420#2#F | \| \| |
| MQ=60 isize=233 C=22 | CGTGTTTTAGTGATTGGAGTCCTCAGCAGATCCCTAATTCTCTCTCCAATCAAGTGCGCTCTCCCCAGCCTGTCCCTTCTCCACGGCCACAGTCCCAG\|C\|CCCCCCACTCCAGTCCTTCCCCaaGGATGCAGCCTCAgCCTTCtCCaCACCc |
| 2:2210:16019:79624#2#F | \| \| |
| MQ=60 isize=232 C=33 | CGTATGCAAGCGATTGGAGTCCTAGCAGATCCCTAATTCTCTCTCCAATCAAGTGCGCTCTCCCCAGCCTGTCCCTTCTCCACGGCCACAGTCCCAG\|C\|CCCCCCACTCCAGTCCTTCCCCAAGGATGCAGCCTCAGCCTTCTCCACACCAC |
| 2:1206:16493:50641#2#R | \| \| |
| MQ=60 isize=-153 C=33 | cgCAGATCCCTAAtTCtCTCTCCAATCAAGTGCGCTCTCCCCAGCCTGTCCCTTCTCCACGGCCACAGTCCCAG\|C\|CCCCCCACTCCAGTCCTTCCCCAAGGATGCAGCCTCAGCCTTCTCCACACCACanGACTCCAATACCCTTCGCAAC |
| 1:2103:14464:10058#2#R | \| \| |
| MQ=60 isize=-154 C=2 | gcagatccctaattctccctccaatcaagtgccctctccccagcctgtcccttccccacggccacagtcccag\|c\|cccccccctccagtccttccCCAAGGATGCAGCCtCAGCCTTCTCccCaCCACGgGgaccCcAaTGTtCCCGCCGAG |
| 1:2208:14767:29670#2#F | \| \| |
| MQ=60 isize=229 C=33 | GTGTTATTGCTCATTGGAGTCntAGATCCCTAATTCTCTCTCCAATCAAGTGCGCTCTCCCCAGCCTGTCCCTTCTCCACggCCACAGTCCCAG\|C\|CCCCCCACtCCAGTCCTTCCCCAAGGatgCaGCCTCAgCCTtCTCCACACCacgtt |
| 1:2212:7911:41090#2#R | \| \| |
| MQ=60 isize=-156 C=33 | aGATCCCTAATtctCtCtCCAATCaAgtcCgCTCtCCCCAGCCtgTCCCTtCtCCACGGCCaCAGTCCCAG\|C\|CCCCCCACTCCAGTCCTTCCCCAAGGATGCAGCCTCAGCCTTCTCCACaCCACGTTAGGACTCCAATTCGAACGGGCAC |
| 1:2208:18759:86008#2#R | \| \| |
| MQ=60 isize=-157 C=33 | gatcCCTAATTCtCTctCCAAtCAAGTgCGCtCtCCCCAGCCTGTCCCTTCTCCACGGCCaCAGTCCCAg\|C\|CCCCCCACTCCAGTCCTTCCCCAAGGATGCAGCCTCAGCCTTCTCCACACCACGTTTanGACTCCAATGGCTGGTACGTT |
| 1:1207:8274:81728#2#F | \| \| |
| MQ=60 isize=227 C=33 | GATGCGACGCTTATTGGAGTCntaTCCCTAATTCTCTCTCCAATCAAGTGCGCTCTCCCCAGCCTGTCCCTTCTCCACGGCCACAGTCCCAG\|C\|CCCCCCACTCCAGTCCTTCCCCAAGGATGCAGCCTCAGCCTTCTCCACACCACGTTTC |
| 1:1211:16401:9721#2#R | \| \| |
| MQ=60 isize=-158 C=33 | ctCCCtAattCtCTCTCCAatccAGTGCgCtCTCCCCAGCCTgTCCCttctCccCggCCACaGTCCCAG\|C\|CCCCCCaCTCCAGTCCTTCCCCAAGGATGCAGCCTCAGCCTTCTCCACACCACGTTTCAGGACTCCAAtCCCAGTGCCCCC |
| 1:1211:5557:43701#2#F | \| \| |
| MQ=60 isize=226 C=37 | CCCGTCGGGCTGATTGGAGTCCTTCCCTAATTCTCTCTCCAATCAAGTGCGCTCTCCCCAGCCTGTCTCTTCTCCACGGCCACAGTCCCAG\|C\|CCCCCCACTCCagTCCTTCCCCAAGGATGCAGCCTCAGCCTTCTCCaCACCACGTTTCC |
| 1:2106:2662:47488#2#F | \| \| |
| MQ=60 isize=226 C=33 | GGCACCTAGGGGATTGGAGTCntCCCTAATTCTCTCTCCAATCAAGTGCGCTCTCCCCAGCCTGTCCCTTCTCCACGGCCACAGTCCCAG\|C\|CCCCCCACTCCAGTCCTTCCCCAaGGATGCAGCCTCAGCCTTCTCCACACCaCGtTTCCC |
| 1:1101:15566:100007#2#R | \| \| |
| MQ=60 isize=-160 C=33 | tCCCTAattCtCTCTCcAATCAAGtgCgCtCtCCCCAgCCTGTCCCtTCTCCACGGCCACAGTCCCaG\|C\|CCCCCCACTCCAGTCCTTCCCCAAGGATGCAGCCTCAGCCTTCTCCACACCACGTTTCCcnAGGACtCCAATGTCCACCGTC |
| 1:1116:20239:84118#2#R | \| \| |
| MQ=60 isize=-159 C=37 | tccctaattctctCTCCAAtCaAGTgCgcgCTCCCCAGCCTGTCCCTTCTCCACGGCCaCAGtcCCaG\|C\|CCCCCCACTCCAGTCCTTCCCCAAGGATGCAGCCTCAGCCTTCTCCACACCACGTTTCCAGGACTCCAATATAGGACGTAAA |
| 1:2207:15870:100697#2#R | \| \| |
| MQ=60 isize=-160 C=33 | ccCCTAAttctCtctCCAATCAAgtgcgctCtCCCCAGCCtGtCCCTTCTCCACGGCCACAGTCCCAG\|C\|CCCCCCaCTCCAGTCCTTCCCCAAGGATGCAGCCTCAGCCTTCTCCACACCACGTTTCCcnGGACTCCAATGTGAATACCTC |
| 2:1202:7620:100113#2#R | \| \| |
| MQ=60 isize=-159 C=37 | tCCCTAATtCtCcCTCCAATCAAGtGCgCTCTCCCCAGCCTGTCCCTTCTCCACGGCCACAGTCCCAG\|C\|CCCCCCACTCCAGTCCTTCCCCAAGGATGCAGCCTCAGCCTTCTACACACCACGTTTCCanGACTCCAATTTCCAGCGTGCT |
| 2:2202:13507:84017#2#R | \| \| |
| MQ=60 isize=-159 C=37 | cCCCTAAtcCTCTCTCCAATCAAGTGCGCcCTCCCCAGCCTGTCCCTtCTCCACGGCCACAGTCCCAG\|C\|CCCCCCaCTCCAGTCCTTCCCCAAGGATGCAGCCTCAGCCTTCTCCACACCACGTTTCCAGGACTCCAATTTCCAGCGTGCT |
| 1:1206:12198:32583#2#R | \| \| |
| MQ=60 isize=-160 C=37 | CCCTAAttCtCtCTCCAATCAAGTGCgCtCTCCCCAGCCTGTCCCTTCTCCaCGGCCaCAGTCCCAG\|C\|CCCCCCACTCCAGTCCTTCCCCAAGGATGCAGCCTCAGCCTTCTCCACACCACGTTTCCCAGGACTCCAATTCTCCCCAATCC |
| 1:1216:16882:24981#2#R | \| \| |
| MQ=60 isize=-160 C=27 | cCCTaATTCTCTCTCCaAtcaAGTGCGCTCTCCCCaGCCTGTCCCttCtCCACgGCCACAGTCCCAG\|C\|CCCCCCACTCCAGTCCTTCCCCAAGGATGCAGCCTCAGCCTTCTCCACACCACGTTTCCCAgnACTCCAATAACTTCCGTAAG |
| 1:2202:3728:40609#2#R | \| \| |
| MQ=60 isize=-160 C=33 | ccctaatcctctctcacatcaagtgcgctctcccCAGCCTGTcCCTtCtccacGGCCACAGTCCCAG\|C\|CCCCCCACTCCAGTCCTTCCCCAAGGATGCAGCCTCAGCCTTCTCCACACCACGTTTCCCanGACTCCAATCGCTAAATGTCG |
| 2:1101:6885:49107#2#R | \| \| |
| MQ=60 isize=-140 C=27 | cccctcacacgcccggcgacccctaagtctctctccaatccagggccctccccccaGCctgaCCCttCtcccCGGccaCaGtCCCAG\|C\|CCCCCCACtCCAGTCCTTCCCCAAGGATGCAGCCTCAGCCAGGACTCCgATTGTGCCCATCTA |
| 2:2112:20711:16329#2#R | \| \| |
| MQ=60 isize=-160 C=37 | CCCTAattCtCtctCCaatCAaGtGCGCTCTCCCCAGCCTGTCCCTTCTcCACGGCCACAGTCCCAg\|C\|CCCCCCACTCCAGTCCTTCCCCAAGGATGCAGCCTCAGCCTTCTCCACACCACGTTTCCCAgnACTCCAATGATTCGGTATAC |
| 1:1110:2342:54526#2#F | \| \| |
| MQ=60 isize=224 C=33 | GGTGGGCgGCGCATTGGAGTCCTCCTAATTCTCTCTCCAATCAAGTGCGCTCTCCCCAGCCTGTCCCTTCTCCACGGCCACAGTCCCAG\|C\|CCCCCCACTCCAGtCCTTCCCCAAGGATGCAGcCTCAGCCTTCTCCaCACCACGcTTCCCC |
| 1:2102:2875:49965#2#F | \| \| |
| MQ=60 isize=224 C=33 | GGGCGGAAAAAGATTGGAGTCCTCCTAATTCTCTCTCCAATCAAGTGCGCTCTCCCCAGCCTGTCCCTTCTCCACGGCCACAGTCCCAG\|C\|CCCCCCACTCCAGtCCTTCCCCAagGaTgCAGCCTCAGCCtTCTCCACACCaCGTttCCCC |
| 2:1206:12150:81741#2#F | \| \| |
| MQ=60 isize=224 C=33 | GGGAGGGGTAGGATTGGAGTCntCCTAATTCTCTCTCCAATCAAGTGCGCTCTCCCCAGCCTGTCCCTTCTCCACGGCCACAGTCCCAg\|C\|CCCCCCACTCCAGTCCTTCCCCAAGGATGCAGCCTCAGCCTTCTCCACACCACGTTTCCCC |
| 2:2109:6701:17712#2#F | \| \| |
| MQ=60 isize=224 C=22 | GTTCCGTTATACATTGGAGTCCTCCTAATTCTCTCTCCAATCAAGTGCGCTCTCCCCAGCCTGTCCCTTCTCCAcGGCCACAGTCCCag\|C\|CCCCCCACtccagtCctTCCCcaagtttccaccctcagccttccccccccccggtttcccc |
| 2:2202:12761:76135#2#F | \| \| |
| MQ=60 isize=224 C=37 | GGGCGGTCGGTCATTGGAGTCCTCCTAATTCTCTCTCCAATCAAGTGCGCTCTCCCCAGCCTGTCCCTTCTCCACGGCCACAGTCCCAG\|C\|CCCCCCACTCCAGTCCTTCCCCAAGGATGCAGCCTCAGCCTTCTCCACACCACGtTTCCCC |
| 2:2216:17831:20305#2#F | \| \| |
| MQ=60 isize=224 C=37 | GTTCCGTTATACATTGGAGTCCTCCTAATTCTCTCTCCAATCAAGTGCGCTCTCCCCAGCCTGTCCCTTCTCCACGGCCACAGTCCCAG\|C\|CCCCCCACTCCAGTCCTTCCCCAAGGATGCAGCCTCAGCCTTCTCCACACCACGTTTCCCC |
| 1:1103:5282:34261#2#R | \| \| |
| MQ=60 isize=-162 C=2 | cctaattctccccccaatcaagtgcgctctccccagcctgtcccttctccacggccacagccccag\|c\|cccccccctccAGTCCTTCCCCAAGGATGCAGCCTCAGCCTTCTCCaCaCCACGTTtCCCCAGGACtCCAAtCtGtCCGACCGC |
| 1:1214:17316:5733#2#R | \| \| |
| MQ=60 isize=-162 C=22 | cctaattctccctccaatcaagtgcgctctccccagcctgtcccctccccccggcCACAGtCCCcG\|C\|CCCCCCaCtCCAGTCCtTCCCCAAGGATGCAGCCTCAGCCTTCTCCACACCACGTTTCCCCAgnACTCCAATCTCCCTTGCCCt |
| 1:2106:7807:66453#2#R | \| \| |
| MQ=60 isize=-162 C=2 | cctaatcctctctccaatcaagtgcgctctccccagcctgtcccttctccacggccacagtcccag\|c\|ccccccactccagtccttcccCAAGGATGCAGcCtCAGcCTTCTCCACaCCACGTttCCCCanGACTCCAATAGTACGTAtGTT |
| 1:2215:9276:22849#2#R | \| \| |
| MQ=60 isize=-162 C=33 | CCTAAttCtCTCTCCAAtCAAGTGCgCtCTCCCCAGCCTGTCCCTtCtCcACGGCCACAGTCCCAG\|C\|CCCCCCACTCCAGTCCTTCCCCAAGGATGCAGCCTCAGCCTTCTCCACACCACGTTTCCCCAgnACTCCAATCTCGGCGCTCCG |
| 2:1106:20663:50494#2#R | \| \| |
| MQ=60 isize=-162 C=33 | cctaattctctctccaatcaagtgcgctctcCCCAGCCTGtCCCTtctCCACGGCCAcAGTCCCAG\|C\|CCCCCCACTCCAGTCCTTCCCCAAGGATGCAGCCTCAGCCTTCTCCACACCaCGTTTCCCCanGACTCCAATCTGAGCTACCGA |
| 2:1116:19127:35970#2#R | \| \| |
| MQ=60 isize=-162 C=37 | cctaattctctccccAATCAAGtgCGCtCTCCCCAGCCTGTCcCttctCCACGGCCACAGTCCCAG\|C\|CCCCCCACTCCAGTCCTTCCCCAAGGATGCAGCCTCAGCCTTCTCCACACCACGTTTCCCCAGGACTCCAAtCCCCCGCACACC |
| 2:1209:17562:52008#2#R | \| \| |
| MQ=60 isize=-162 C=37 | cctaattctctctccaatCAAGTGCGctCTCCCCAGCCTGTCCCttcTCcaCGGCCACAGTCCCAG\|C\|CCCCCCACtCCAGTCCTTCCCCAAGGATGCAGCCTCAGCCTTCTCCACACCACGTTTCCCCAGGACTCCAATCCCCCGCACACC |
| 2:2208:10450:77939#2#R | \| \| |
| MQ=60 isize=-162 C=37 | CctAaTTCTCtCtCCAATCaaGtGCGCtCcCCCCAGCCTGTCCCTTCTCCaCGGCCaCAGTCCCaG\|C\|CCCCCCaCTCCAGTCCTTCCCCAAGGATGCAGCCTCAGCCTTCTCCACACCACGTTTCCCcanGACTCCaacCCCCCGCACACC |
| 1:1115:7537:55633#2#F | \| \| |
| MQ=60 isize=328 C=37 | ACTGTTCTTCTGATTGGAGTCCTCTAATTCTCTCTCCAATCAAGTGCGCTCTCCCCAGCCTGTCCCTTCTCCACGGCCACAGTCCCAG\|C\|CCCCCCACTCCAGtCCTTCCCCAAGGATGCAGCCTCAGCCTTCTCCACACCACGTTTCCCCa |
| 1:2103:15575:41522#2#F | \| \| |
| MQ=60 isize=357 C=33 | TCATGTCCAGTAATTGGAGTCCTCTAATTCTCTCTCCAATCAAGTGCGCTCTCCCCAGCCTGTCCCTTCTCCACGGCCACAGTCCCAG\|C\|CCCCCCaCTCCAGTCCtTCCCCaagGAtgcAGCCtCAGCCTTCTCCACaCCACgTTtcCCCc |
| 2:1111:16597:77151#2#F | \| \| |
| MQ=60 isize=223 C=37 | GGtGGTGGGAGTATTGGAGTCCTCTAATTCTCTCTCCAATCAAGTGCGCTCTCCCCAGCCTGTCCCTTCTCCACGGCCACAGTCCCAG\|C\|CCCCCCAcTCCaGTCCTTCCCCAAGGATGCAGCCTCAGCCTTCTCCaCACCACGTTTCCCCc |
| 2:1215:7513:77457#2#F | \| \| |
| MQ=60 isize=223 C=33 | GGCGTGCCGGCTATTGGAGTncTCTAATTCTCTCTCCAATCAAGtgCGCTCTCCCCAGCCTGTCCCTTCTCCACGGCCACAGTCCCAg\|C\|CCCCCCACtCCAGTCCTTCCCCAAgGATGCAGCCTCAGCCTTCTCCACACCACGTTTCCCCA |
| 2:2116:4570:36930#2#F | \| \| |
| MQ=60 isize=328 C=2 | TATACCGCGCTGATTGGAGTncTCtAATTcTcTCTCCAATCAAgtgCGcTcTCCCCAGCCTgTCCCTTctccacggccacagtcccag\|c\|ccccccactccggtccttccccaaggagccagcctcagccgtctccacaccacgttcccccc |
| 1:1206:13562:90799#2#F | \| \| |
| MQ=60 isize=222 C=33 | GTTATAGTGGGTATTGGAGTCCTTAATTCTCTCTCCAATCAAGTGCGCTCTCCCCAGCCTGTCCCTTCTCCACGGCCACAGTCCCAG\|C\|CCCCCCACTCCAgtCCTTCCCCAAGGATGCAGCCTCAgCCTTCtCCaCACCACgtttCCCCAC |
| 2:1109:9491:37962#2#R | \| \| |
| MQ=60 isize=-164 C=33 | taattcgctctccaatcaagtgcgctctCCCCAGCCtgtCCCTTCtcCacGGCCACAGtCCCAG\|C\|CCCCCCACTCCAGTCCTTCCCCAAGGATGCAGCCTCAGCCTTCTCCACACCACGTTTCCCCACanGACTCCAAtCACTCCCCACTG |
| 1:2202:9715:34020#2#R | \| \| |
| MQ=60 isize=-162 C=2 | ctaaattctccctccaatcaagtgcgccctccccagcctgtcccttctccacggccccagccccag\|c\|ccccccactccagtccttCCCCAAGGATGCAGCCTCAGCCTTCTCCACACCACGTTTCCCCanGACTCCAATCTGTCCGACCGC |
| 2:2210:9676:61987#2#R | \| \| |
| MQ=60 isize=-165 C=33 | AATTCtCtCTCCAATCAAGTGCGCtCtCCCCAGCCtgTCCCTtCTCCACGGCCACAGTCCCAG\|C\|CCCCCCaCTCCAGTCCTTCCCCAAGGATGCAGCCTCAGCCTTCTCCACACCACGTTTCCCCACAGGGACTCCAATAAACACCGTCCA |
| 2:2213:10050:77399#2#R | \| \| |
| MQ=60 isize=-164 C=33 | AATTCTCTCTCCAATCAAGTGCGCtCTCCCCAGCCTGTCCCTtCTCCACGGCCACAGTCCCAG\|C\|CCCCCCACtCCAGTCCTTCCCCAAGGATGCAGCCTCAGCCTTCTCCACACCACGTTTCCCCACAAGGACTCCAAtCACAACACTTCA |
| 1:2101:1376:39289#2#R | \| \| |
| MQ=60 isize=-166 C=37 | attCTCTCTCCAaTCAaGtGCGCtCTCCCCAGCCTGTCCCTTCTCCACGGCCACAGTCCCAG\|C\|CCCCCCACTCCAGTCCTTCCCCAAGGATGCAGCCTCAGCCTTCTCCACACCACGTTTCCCCACAGAGGACTCCAATTCCAACGCCGAC |
| 1:2213:20315:22573#2#R | \| \| |
| MQ=60 isize=-166 C=37 | ATTCtCtctCCAATCAAGTGCGCTCTCCCCAGCCTGTCCCTTCTCCACGGCCaCAGTCCCAG\|C\|CCCCCCACTCCAGTCCTTCCCCAAGGATGCAGCCTCAGCCTTCTCCACACCACGTTTCCCCACAGAgnACTCCAATCCCTTTGCCTAA |
| 2:1205:4235:62397#2#R | \| \| |
| MQ=60 isize=-166 C=27 | attctctctccaATCAAGTgccctCtccCCagccTgtCCCtTCTCcaCGGCCACAGTCCCag\|C\|CCCCCCACTCCAGTCctTCCCCAAGGATGCAGCCTCAGCCTTCtCCACACCACGTTTCCCCACAGanGACTCCAATTTGCCTAATACG |
| 2:2216:15071:14938#2#R | \| \| |
| MQ=60 isize=-166 C=33 | attctctCTCCAATCAAGTGCGCtCTCCCCAGCCTGTTCCTCCTCCACGGCCACAGTCCCAG\|C\|CCCCCCaCTCCAGTCCTTCCCCAAGGATGCAGCCTCAGCCTTCTCCACaCCACGTTTCCCCACAGAGGACTCCAATTGTACACAATAA |
| 1:1210:18842:30579#2#F | \| \| |
| MQ=60 isize=219 C=33 | GGCCTGAGGCGAATTGGAGTCCTTCTCTCTCCAATCAAGTGCGCTCTCCCCAGCCTGTCCCTTCTCCACGGCCACAGTCCCAG\|C\|CCCCCCACTCCAGTCCTTCCCCAaGGAtGCAGCCTCAgCCTTCTCCACACCACGTTTCCCCACAGAC |
| 1:2110:5998:88890#2#F | \| \| |
| MQ=60 isize=219 C=33 | ATTTTTTATCTAATTGGAGTCCTTCTCTCTCCAATCAAGTGCGCTCTCCCCAGCCTGTCCCTTCTCCACGGCCACAGTCCCAG\|C\|CCCCCCACTCCAGTCCTTCCCCAAGGATGCAGCCTCAGCCTTCTCCaCACCACGTTTCCCCACAGAC |
| 1:2205:19179:38662#2#F | \| \| |
| MQ=60 isize=219 C=33 | ggCCTGAGGCGAATTGGAGTCnttCTCTCTCCAATCAAGTGCGCTCTCCCCAGCCTGTCCCTTCTCCACGGCCACAGTCCCAG\|C\|CCCCCCACtcCAgtCCTTCCCCAAGgatgCAGCCtCAGCCTTCTCCacACCACgttTCCCCACAgac |
| 2:2102:14908:92694#2#F | \| \| |
| MQ=60 isize=353 C=33 | GGCGCGGGGCGGATTGGAGTCCTTCTCTCTCCAATCAAGTGCGCTCTCCCCAGCCTGTCCCTTCTCCACGGcCACAGTCCCAG\|C\|CCCCCCACtCCAGTcCTTCCCCaaggatgcagcctcagccttctcccccccccgtttcccccccgac |
| 1:1203:19671:77294#2#F | \| \| |
| MQ=60 isize=218 C=27 | TGCGGTTTAATCATTGGAGTCntCTCTCTCCaatCAAGTGCGCTCTCCCCAgCCTGTCcCtTCTCCACGGCCACAgTCCCAG\|C\|CCCCCCACTCCAGTCCTtCccCAAGGatgCagcCTCaGCCTTCtccacaccacccttccccacagacc |
| 2:2208:10099:23360#2#F | \| \| |
| MQ=60 isize=218 C=33 | AAATGCCGAGCGATTGGAGTCnctCTCCAATCAAGTGCGCTCTCCCCaGCCTGTCCCTTCTCCACGGCCaCAGTCCCAG\|C\|CCCCCCACtCCAGtCCTTCCCCAAGGAtgCAGCCTCAGCCTTCTCCACACCACGTTTCCcCacagACAAgt |
| 2:2214:10054:57806#2#F | \| \| |
| MQ=60 isize=217 C=33 | GGCATGGGTCAATTGGAGTCcTCTCTCCAATCAAGTGCGCTCTCCCCAGCCTGTCCCTTCTCCACGGCCACAGTCCCAG\|C\|CCCCCCACTCCAGTCCTTCCCCAAGGATGCAGCCtCAGCCTTCTCCACACCACGTTTCCCCACAGACAAGT |
| 2:1105:17784:85550#2#R | \| \| |
| MQ=60 isize=-169 C=22 | ctctCTCCAAtCAAgtgCGCtctCCCCAGCCtgtCCCTTctCCaCGGCCACAGtCCCAG\|C\|CCCcCcaCTCCAGTCCttCCCCAAGGATGCAGCCTCAGCCTTCTCCACACCACGTTTCCCCACAGACAanGACTCCAATATATGTGCACCC |
| 2:2202:20699:88407#2#R | \| \| |
| MQ=60 isize=-170 C=33 | CtCTCTCCAATCAAGtGCGCtCTCCCCAGCCTGTCCCTTCTCCACGGCCACAGTCCCAG\|C\|CCCCCCACTCCAGTCCTTCCCCAAGGATGCAGCCTCAGCCTTCTCCACACCACGTTTCCCCACAGACAAGGACTCCAATACTCCCACCCTT |
| 2:2112:6964:3580#2#F | \| \| |
| MQ=60 isize=216 C=27 | TGGTCGGAGACAATTGGAGTncTTCTCTCCAATCAAGTGCGCTCTCCCCAGCCTGTCCCTTCTCCACGGCCACAGTCCCAG\|C\|CCCCCCACtCCAGcCCTTCCCCAAGgAtGcaGCCtcagCCTtCtCCcccccacgTTTCCCCACagacaa |
| 1:2203:5630:9546#2#R | \| \| |
| MQ=60 isize=-169 C=33 | cctctcccATCAaGTGCGCtCTCCCCAGCCTGtCCCTTCTCCACGGCCACAGTCCCaG\|C\|CCCCCCACTCCAGTCCTTCCCCAAGGATGCAGCCTCAGCCTTCTCCACACCACGTTTCCCCACAGACAAAGGACTCCAATGATTCGGGGCAA |
| 2:1114:14890:80836#2#R | \| \| |
| MQ=60 isize=-169 C=33 | cctCtCcAATCAAGTGCGCTCTCCCCAGCcTGTCCCTTCtCCACGGCCACAGTCCCAG\|C\|CCCCCCACTCCAGTCCTTCCCCAAGGATGCAGCCTCAGCCTTCTCCACACCACGTTTCCCCACAGACAAAgnACTCCAATGTACCATATATA |
| 1:1113:15062:36581#2#F | \| \| |
| MQ=60 isize=320 C=27 | GAAACTTACTGCATTGGAGTCCTCTCCAATCAAGTGCGCTCTCCCCAGCCTGTCCCTTCTCCACGGCCACAGTCCCAG\|C\|CCCCCCACTCCAGTCCTTCCCCAAGGATGCAGCCTCAGCCTTCTCCACACCACGTTTCCCCACAGACAAGTT |
| 1:2204:15463:13221#2#F | \| \| |
| MQ=60 isize=215 C=33 | GCCGGGGGCGGTATTGGAGTCCTCTCCAATCAAGTGCGCTCTCCCCAGCCTGTCCCTTCTCCACGGCCACAGTCCCAG\|C\|CCCCCCACTCCAGTCCTTCCCCAAGGATGCAGCCTCAGCCTTCTCCACACCaCGTTTCCCCACagACAagtc |
| 2:1116:4356:68902#2#F | \| \| |
| MQ=60 isize=215 C=33 | GGTTCGGCCCCGATTGGAGTCCTCTCCAATCAAGTGCGCTCTCCCCAGCCTGTCCCTTCTCCACGGCCACAGTCCCAG\|C\|CCCCCCaCcCCAgccCTTCCCCaaGGatgcacccccagcctcctccccacccctttcccccccacacaagtc |
| 2:1215:10896:17350#2#F | \| \| |
| MQ=60 isize=215 C=37 | GCGTAAAGGCGGATTGGAGTncTCCTCTCCAATCAAGTGCGCTCTCCCCAGCCTGTCCCTTCTCCACGGCCACAGTCCCAG\|C\|CCCCCCACTCCAGtCCTTCCCCAAGGATGCAGCCtCAGCCTTCTCCACACCACGTTTCCCCaCAgaCAa |
| 2:2102:15217:70550#2#F | \| \| |
| MQ=60 isize=215 C=27 | TTTGTTTCCTGCATTGGAGTCCTCTCCAATCAAGTGCGCTCTCCCCAGCCTGTCCCTTCTCCACGGCCACAGTCCCAG\|C\|CCCCCCACTCCAGTCCTTCCCCaaggatGCAGCctCAgCCTtctCCaccCcccgtttCCCCcccgacaagtc |
| 2:2211:17358:60221#2#F | \| \| |
| MQ=60 isize=215 C=37 | CTGGCGCGCGGCATTGGAGTCCTCTCCAATCAAGTGCGCTCTCCCCAGCCTGTCCCTTCTCCACGGCCACAGTCCCAG\|C\|CCCCCCACTCCAGTCCTTCCCCAAGGaTGCAGCCTCAGCCTTCTCCACACCaCGTTTCcCCACagACAAGTT |
| 1:1107:21155:65033#2#R | \| \| |
| MQ=60 isize=-170 C=33 | ccctccaatccagtgcgctctccccagcctgtcccttctccacggCCacagTCCCAg\|C\|CCCCCCACTCCAGTCCttCCCCAAGGATGCAGCCTCAGCCTTCTCCACaCCaCGTTTCCCCACAGACAAGAGGACTCCAATCCTTTTCGAGGG |
| 1:1116:5589:47430#2#R | \| \| |
| MQ=60 isize=-170 C=33 | ccctCCAATCAAGTGCGCTCTCCCCAGCCTGTCCCTTCTCCACGGCCaCAGtCCCAG\|C\|CCCCCCACTCCAGTCCTTCCCCAAGGATGCAGCCTCAGCCTTCTCCACACCACGTTTCCCCACAGACAAGAGGACTCCAAtCGTCCCCTGCCC |
| 1:1214:9941:77663#2#R | \| \| |
| MQ=60 isize=-170 C=27 | CtcTcCAAtCAAGTGCGCTCTCCCCAGCCTGTCCCTTCTCCACGGCCACAGTCCCAG\|C\|CCCCCCACTCCAGTCCTTCCCCAAGGATGCAGCCTCAGCCTTCTCCACACCACGTTTCCCCACAGACAAGAgnACtCCAATCCTTTTCGAGGG |
| 1:2105:10584:22573#2#R | \| \| |
| MQ=60 isize=-170 C=37 | ccctCCAATCAAgtgCGCtCtCCCCAGCCTGTCCCTTCTCCACGGCCACAGTCCCAG\|C\|CCCCCCACTCCAGTCCTTCCCCAAGGATGCAGCCTCAGCCTTCTCCACACCACGTTTCCCCACAGACAAGanGACTCCAATGTAGGACTGATA |
| 1:2113:5939:87051#2#R | \| \| |
| MQ=60 isize=-170 C=2 | ctctccaatcaagtgcgctctccccagcctgtcccttctccacggccacagtcccag\|c\|ccccccactccagtccttccCCAAGGATGCAGCCTCAGCCTTCTCCACACCACGTTTCCCCACAGACAAgannACTCCAATGTAGGACTGATA |
| 2:1107:7563:31730#2#R | \| \| |
| MQ=60 isize=-170 C=22 | ctctccaatcaagcgcgctctcccCAGCCtggCcCTtCtCCACGGcCAcaGtCCCaG\|C\|CACCCCACTCCAGTCCTTCCCCAAGGATGCAGCCTCAGCCTTCTCCACACCACGTTTCCCCACAGACAAGanGACTCCAATTGATGCCCAGTT |
| 2:1216:2406:65558#2#R | \| \| |
| MQ=60 isize=-170 C=15 | CcCtCCAatCAagggCgcTCtcCCCAGCCTGtCCCTtCtCCACGGCCACAGTCccag\|c\|CCCCCCactCCGGTCCTtCCCCAAGGATGCAGCCTCAGCCTTCTCCACACCACGTTTCCCCACAGACAAGAgnACTCCAATCGTCCCCTGCCC |
| 1:1102:20709:7783#2#F | \| \| |
| MQ=60 isize=214 C=37 | CGTGTGAGTGGGATTGGAGTCntcTCCAATCAAGTGCGCTCTCCCCAGCCTGTCCCTTCTCCACGGCCACAGTCCCAG\|C\|CCCCCCACTCCAgtCCTTCCCCAAGGATGCAGCCTCAGCCTTCTCCACACCACGTTTCCCCACAGACAAGTT |
| 1:1201:7005:41359#2#F | \| \| |
| MQ=60 isize=214 C=33 | TGTGACCTGCTTATTGGAGTCntcTCCAATCAAGTGCGCTCTCCCCAGCCTGTCCCTTCTCCACGGCCACAGTCCCAG\|C\|CCCCCCACTCCAGTCCTTCCCCAAGGATGCAGCCTCAGCCTTCTCCACACCaCGtTTCCCCaCAGACAAGtc |
| 1:2106:3073:77471#2#F | \| \| |
| MQ=60 isize=214 C=33 | GGTTCGGCCCCGATTGGAGTCntcTCCAATCAAGTGCGCTCTCCCCAGCCTGTCCCTTCTCCACGGCCACAGTCCCAG\|C\|CCCCCCaCTCCAGTCCtTCCCCAAGGATGCAGCCTCAGCCTTCTCCACACCcCGtttCCCCaCAGACAAGtt |
| 2:1106:2610:27859#2#F | \| \| |
| MQ=60 isize=214 C=2 | ACGGTGTGgTGGATTGGAGTCntcTCCAATCAAGTGCGCTCTCCCCAGCCtGtCCCTTCTCCACGgccacagtcccac\|c\|cccccccctccagccctcccccacggatcaacccccaccctctcccccccaccgctcccccccgacaaagtc |
| 2:1109:14993:71484#2#F | \| \| |
| MQ=60 isize=214 C=33 | AGATGGCACGGGATTGGAGTCntTCTCCAATCAAGTGCGCTCTCCCCAGCCTGTCCCTTCTCCACGGCCACAGTCCCAG\|C\|CCCCCCACTCCAGTCCTTCCCcAaggATGCAGCctCaGCCTTCTCCaCACCacgttTCCCCacacacaagt |
| 2:1216:19895:21634#2#F | \| \| |
| MQ=60 isize=214 C=33 | AGATGGCACGGGATTGGAGTncTTCTCCAATCAAGTGCGCTCTCCCCAGCCTGTCCCTTCTCCACGGCCACAGTCCCAG\|C\|CCCCCCACTCCAGTCCTTCCCCAAGGATGCAGCCTCAGCCTTCTCCACACCACGTTTCCCCACAGACaagt |
| 1:1215:19930:43615#2#R | \| \| |
| MQ=60 isize=-172 C=2 | tcttcaatcaagtgcgctctccccagcctgtcccttatccacggcaacagtcccag\|c\|ccccccactccagtcCtTcccCAAgGATgtAgCCtcAGcCTTcTCCAcacctCGTTTCccCaCAGaCaAgtTAGGacTCCAATGtTGGtACCtA |
| 1:2111:7623:27228#2#F | \| \| |
| MQ=60 isize=213 C=27 | ACGCGCCCATCGATTGGAGTCCTCCAATCAAGTGCGCTCTCCCCAGCCTGTCCCTTCTCcACGGCCACaGTCcCAG\|C\|CCCCCCACTCcaGTCCTtCCCCAagGATGCAGCCtCAGCCTTCTCCACACCACGtttCCCCAcagacaagttcc |
| 1:1215:8643:21495#2#R | \| \| |
| MQ=60 isize=-172 C=37 | CTCCAATCAAGtGCGCtctCCCCAGCCTGtCCCTTCTCCACGGCCACAGTCCCAG\|C\|CCCCCCACTCCAGTCCTTCCCCAAGGATGCAGCCTCAGCCTTCTCCACACCACGTTTCCCCACAGACAAGTTAGGACTCCAATACGGGTTGTCAT |
| 1:2102:12144:80222#2#F | \| \| |
| MQ=60 isize=212 C=22 | TCTCTGGGTACGATTGGAGTCCTTCCAATCAAGTGCGCTCTCCCCAGCCTGTCCCTTCTCCACGGCCACAGTCCCAG\|C\|CCCCCCACTCCAGTCCTTCCCCAAGGATGCAGCCTCAGCCTTCTCCACACCACGTTTCCCCACAGACAAGTTC |
| 1:1101:3471:34116#2#R | \| \| |
| MQ=60 isize=-173 C=37 | tCCAATCAAGTGCGCTCTCCCCAGCCTGTCCCTTCTCCACGGCCACAGTCCCAG\|C\|CCCCCCACTCCAGTCCTTCCCCAAGGATGCAGCCTCAGCCTTCTCCACACCACGTTTCCCCACAGACAAGTTCanGACTCCAATTCCCGGTAACAA |
| 1:2215:11623:40497#2#R | \| \| |
| MQ=60 isize=-173 C=27 | cccaatcaagtgcgctctCCCCAGCCtgtCCCTTCTCCaCGGCCaCAGTCCCAG\|C\|CCCCCCACTCCAGTCCTTCCCCAAGGATGCAGCCTCAGCCTTCTCCACACCACGTTTCCCCACAGACAAGTTCAgnACTCCAATCGCGCTCCCCCT |
| 2:2204:16025:4148#2#R | \| \| |
| MQ=60 isize=-173 C=37 | cCCAAtCaAgtGCGCtCtCCCCAGCCTGTCCCTTCTCCACGGCCACAGTCCCAG\|C\|CCCCCCACTCCAGTCCTTCCCCAAGGATGCAGaCTCAGCCTTCTCCACaCCACGTTTCCCCACAGACAAGttCanGACTCCAATTCCCGGTAACAA |
| 1:2115:12805:47003#2#R | \| \| |
| MQ=60 isize=-174 C=37 | ccaatcaagtgcgccctCCCCAGCCTGTCCCttCTCCACGGCCACAGtCCCAG\|C\|CCCCCCACtCCAGTCCTTCCCCAAGGATGCAGCCTCAGCCTTCTCCACACCACGTTTCCCCACAGACAAGTTCCAGGACTCCAATCGGCCAGCCCCC |
| 2:1108:8231:99252#2#R | \| \| |
| MQ=60 isize=-169 C=2 | ccccccccaatcaagtccgctccccccagcctgtcccttctccacggccacagtcccag\|c\|ccccccactccagtccttccccaaggatgcagcctcagCCTTCTCCACaCCaCGTTtcCCCACAGAcaanGACTCCAATCCaCCACGCCTA |
| 2:1116:20247:68540#2#R | \| \| |
| MQ=60 isize=-174 C=33 | CCAATCAAgtgcgctCTCCccAGCCTGTCCCttCTCCACGGCCACAGTCCCAG\|C\|CCCCCCaCTCCAGTCCTTCCCCAAGGATGCGGCCTCAGCCTTCTCCACACCACGTTTCCCCACAGACAAGTTCCAGGACTCCAATGCTCCGCCTGGC |
| 2:1215:2768:59259#2#R | \| \| |
| MQ=60 isize=-174 C=33 | ccaaccaagtgcgctctCCCCAGCCtGTCCCttCTCCaCGGCCACAGTCCCaG\|C\|CCCCCCaCTCCAGTCCTTCCCCAAGGATGCAGCCTCAGCCTTCTCCACACCACGTTTCCCCACAGACAAGTTCCAgnACTCCAaTCGGCCAGCCCCC |
| 2:2103:17149:89905#2#R | \| \| |
| MQ=60 isize=-174 C=33 | ccaatcaagtgcgctcccCCCAGCctgtcaCTTCTCCACGGCcaCAGTCCCAG\|C\|CCCCCCACTCCAGTCCTTCCCCAAGGATGCGGCCTCAGCCTTCTCCACACCACGTTTCCCCACAGACAAGTTCCAGGACTCCAATGCTCCGCCTGGC |
| 1:1111:5284:77785#2#R | \| \| |
| MQ=60 isize=-175 C=37 | caatcaagcgcgctCTCCCCcGCCtgtCcCttctCcACGgCCcCAGtCCCaG\|C\|CCCCCCACtCCAGTCCtTCCCCAAGGATGCAGCCTCAGCCTTCTCCaCACCACGTTTCCCCACAGACAAGTTCCCAGGACTCCAATCGTGCCTCAAAC |
| 1:1216:10075:62212#2#R | \| \| |
| MQ=60 isize=-173 C=37 | CAATCAAGTgCgctCTCCCCAGCCTGTCCCTTCTCCACGGCCACAGTCCCaG\|C\|CCCCCCACTCCAGTCCTTCCCCAAGGATGCAGCCTCAGCCTTCTCCACACCACGTTTCCCCACAGACAAGTTCATAgnACTCCAATCCCAAACTGAGC |
| 1:2116:1516:95388#2#R | \| \| |
| MQ=60 isize=-175 C=37 | CAATCAagtGCgCtCtCCCCAGCCTGTCCCttCtCCcCGGCCACAGTCCCAG\|C\|CCCCCCaCTCCAGTCCTTCCCCAAGGATGCAGCCTCAGCCTTCTCCACACCACGTTTCCCCACAGACAAGTTCCCanGACTTCAATTGTTTCAACAAC |
| 2:1101:12486:82246#2#R | \| \| |
| MQ=60 isize=-177 C=22 | caatcaagcgcgctctcccccggctgtcCCTtcccCACGgCCACAGtCCcag\|C\|CCcccCaCTCCAGTCCtTCCCCAAGGATGCAGCCTCAGCCTTCTCCACACCACGTTTCCCCACAGACAAGTTCCCCAGGACTCCAATTGCGGGTTGGA |
| 2:1107:19518:74391#2#R | \| \| |
| MQ=60 isize=-175 C=2 | caatcaagtgtgctcgccccagcctgcccctgctccacggccacagtcccag\|c\|ccccccactccagtccttccccaaGGATGCAgCCTCaGCCtTCTCCacaCcaCGTTTCCCCACAGACAAGTtCCcanGACTCCAagCCCCCGCCCCCG |
| 2:1203:17008:89798#2#R | \| \| |
| MQ=60 isize=-175 C=33 | CAATcAAgTGCGCTCTCCCCAGCCTGtCCCTTCTCCaCGGCCACAGTCCCAG\|C\|CCCCCCACTCCAGTCCTTCCCCAAGGATGCAGCCTCAGCCTTCTCCACACCACGTTTCCCCACAGACAAGTTCCCAGGACTCCAATTGGAAGGTAACC |
| 2:1212:19522:70817#2#R | \| \| |
| MQ=60 isize=-175 C=33 | CAATCAAGTGCGCTCTCCCCAGCCTGTCCCTTCTCCACGGCCACAGTCCCAG\|C\|CCCCCCACTCCAGTCCTTCCCCAAGGATGCAGCCTCAGCCTTCTCCACACCACGTTTCCCCACAGACAAGTTCCCAGGACTCCAATTCCAAAGAACGA |
| 2:2115:6585:71576#2#R | \| \| |
| MQ=60 isize=-175 C=33 | caatcaagtgcgctcccCCCAgCctgtCCCTTccCCAcGGCCACAGTCCCAG\|C\|CCCCCcactCCAGTCCTTCCCCAAGGATGCAGCCTCAGCCTTCTCCACaCCACGTTTCCCCACAGACAAGTTCCCAgnACTCCAATCATTCCCTTCGT |
| 2:2207:3147:91112#2#R | \| \| |
| MQ=60 isize=-175 C=37 | CAATCaAGtgCgctCtCCCCAGCCTGTCCCtTCtCCaCGGCcaCAGTCCCAG\|C\|CCCCCCaCTCCAGTCCTTCCCCAAGGATGCAGCCTCAGCCTTCTCCACACCACGTTTCCCCACAGACAAGTTCCCanGACTCCAATTGTCACACAACC |
| 1:1209:15796:95282#2#F | \| \| |
| MQ=60 isize=209 C=33 | CAAGACCACCGAATTGGAGTCntAATCAAGTGCGCTCTCCCCAGCCTGTCCCTTCTCCACGGCCACAGTCCCAG\|C\|CCCCCCACTCCAGTCCTTCCCCAAGGATGCAGCCTCAGCCTTCtcCaCACCACGtTTCCCCACAGACAAGTTCCCC |
| 1:2113:18054:81195#2#F | \| \| |
| MQ=60 isize=209 C=27 | AGCCTGCGGCTTATTGGAGTnntaATCAAGTGCGCTCTCCCCAGCCTGTCCCTTCTCCACGGCCACAGTCCCAG\|C\|CCCCCCACTCCAGTCCTTCCCCAAGGATGCAGCCTCAGCCTTCTCCACACCACGTTTCCCCACAGACAAGTTCCCC |
| 1:2204:5869:79351#2#F | \| \| |
| MQ=60 isize=209 C=27 | ACTGTCCTGCCTATTGGAGTCCTAATCAAGTGCGCTCTCCCCAGCCTGTCCCTTCTCCACGGCCACAGTCCCAG\|C\|CCCCCCACTCCAGTCCTTCCCCAAGGATGCAgCCTCAGCCttCTCCaCACCaCGtTTCCCCACAGACAcgtTCCCC |
| 2:2115:18301:20763#2#F | \| \| |
| MQ=60 isize=209 C=33 | TTATTATCAACGATTGGAGTncTAATCAAGTGCGCTCTCCCCAGCCTGTCCCTTCTCCACGGCCACAGTCCCAG\|C\|CcCCCCACTCCAGTCCTTCCCCAAGGATGcAGCCtCaGCCTTCTCCACACCacgtTTCCCCacagacaagttcccc |
| 1:1107:11338:20494#2#R | \| \| |
| MQ=60 isize=-177 C=33 | AATCAAgTgcgCtCTCCCCAGCCTGTCCCTTCtCCACGGCCACaGTCCCag\|C\|CCCCCCACTCCAGTCCTTCCCCAAGGATGCAGCCTCAGCCTTCTCCACACCACGTTTCCCCACAGACAAGTTCCCCAGGACTCCAATATTAGCAGCGGT |
| 1:1114:18853:77429#2#R | \| \| |
| MQ=60 isize=-177 C=27 | aaccaagtgcgctctCCCCAGcCtGTCCCtTCtCCaCgGCcaCacTCcCAG\|C\|cCCCCCaCTCCAGTCCTtccCCAAGGaTGCAGCCTCAGCCTTCTCCaCACCaCGTTTCCCCACAGACAAGTTCCCCAGGACTCCAATCCCAGCCTCCTC |
| 1:1213:3459:60372#2#R | \| \| |
| MQ=60 isize=-177 C=33 | aATCAAgTGCGCTCTCCCCAgCCTGTCCCTTCTCCaCGGCCACAGTCCCAG\|C\|CCCCCCACTCCAGTCCTTCCCCAAGGATGCAGCCTCAGCCTTCTCCACACCACGTTTCCCCACAGACAAGTTCCCCAgnACTCCAATACTCTAGCCTGG |
| 1:2105:8098:49255#2#R | \| \| |
| MQ=60 isize=-177 C=33 | aatcaagtgcGCTCTcCCCAGCCtGTCCCTTCTCCACGGCCACAGtCCCAG\|C\|CCCCCCactCCAGTCCTTCCCCAAGGATGCAGCCTCAGCCTTCTCCACACCaCGTTTCCCCGCAGACAAGTTCCCCanGACTCCAATAACCTGGCACGC |
| 1:2113:4574:16227#2#R | \| \| |
| MQ=60 isize=-177 C=2 | aatcaagtgcgctctccccagcctgtcccttctccacggccacagtcccag\|c\|ccccccactccagtccttccccaaggatgcagcctcagccttctccacaccacgttTCCCCACAGACTAGTTCCCcannACTCCAATGGACGCCGCGTA |
| 2:1105:6866:88820#2#R | \| \| |
| MQ=60 isize=-177 C=2 | aatcaagtcccctccccccagcctgtcccttctccacggccacagtcccag\|c\|cccccccctccagtccttccccaaggatgcagcctcagccttctccacaccacgtttccccacagacaagttccccangaCTCCAAtCGCCGCCGGTAT |
| 2:1201:6578:94068#2#R | \| \| |
| MQ=60 isize=-177 C=33 | aATCaAGTGCcCtCtCCCCAGCCTgtCCCTTCtCCACGGCCaCAGtCCCAG\|C\|CCCCCcactCCAGTCCTTCCCCAAGGATGCAGCCTCAGCCTTCTCCACACCACGTTTCCCCACAGACAAGTTCCCCAGGACTCCAATGACCCGGCCACG |
| 2:1215:20640:81841#2#R | \| \| |
| MQ=60 isize=-177 C=37 | AATCAAGTGCGCtCTCCCCAGCCTgTCCCTTCTCCACGGCCACAGTCCCAG\|C\|CCCCCCACTCCAGTCCTTCCCCAAGGaTGCAGCCTCAGCCTTCTCCACACCACGTTTCCCCACAGACAAGTTCCCCAgnACTCCAAtCTGCTCGCGGGC |
| 2:2115:5934:80299#2#R | \| \| |
| MQ=60 isize=-177 C=33 | aatcaaggtccctctCCCCAgCCtgtcCCTTCtcCacgGCCAcAGTCCcaG\|C\|CCCCCCaCTCCAGTCCTTCCCCAAGGATGCAGCCTCAGCCTTCTCCACACCACGTTTCCCCACAGACAAGTTCCCCAgnACTCCAATTATCTCAAGCAC |
| 1:1203:17375:10437#2#F | \| \| |
| MQ=60 isize=208 C=33 | GATTGGCGCCCTATTGGAGTCntATCAAGTGCGCTCTCCCCAGCCTGTCCCTTCTCCACGGCCACAGTCCCAG\|C\|CCCCCCACtCCAGtCCTTCCCCaAgGAtGCAgCCtCAGCCTTCTCCacaCCACgtTTCCCCaCAGacaAGtTCCCca |
| 1:2211:15869:34889#2#F | \| \| |
| MQ=60 isize=208 C=33 | GGGTGGGGGGCGATTGGAGTCCTATCAAGtGCGCTCTCCCCAGCCTGTCCCTTCTCCACGGCCACAGTCCCaG\|C\|CCCCCCACTCCAGtcCTTCCCCAAGGatgCAGCCTCAGCCTTCtCCaCaCcccgtTTCCCCACAgacAAgtcccccc |
| 2:2104:17768:41519#2#F | \| \| |
| MQ=60 isize=208 C=37 | GTTTCTTACTGTATTGGAGTCCTATCAAGTGCGCTCTCCCCAGCCTGTCCCTTCTCCACGGCCACAGTCCCAG\|C\|CCCCCCACTCCAGtcCTTCCcCAAgGATGCAGCCTCAGCCTTCTCCACACCACGTtTCCCCacAgacaagtTCCCCc |
| 2:2215:18001:38369#2#F | \| \| |
| MQ=60 isize=208 C=37 | TATGCGGGCATGATTGGAGTCCTATCAAGTGCGCTCTCCCCAGCCTGTCCCTTCTCCACGGCCACAGTCCCAG\|C\|CCCCCCACTCCAGTCCTTCCCCAAGGATGCAGCCTCAGCCTTCTCCACACCACGtTTCCCCaCaGACAAGtTCCCCc |
| 2:1115:11699:40986#2#R | \|C\| |
| MQ=60 isize=-177 C=37 | atcaagtgcgctctccCCAGCCTGTCCcTTCtCCACGGCCACAGTCCCAI\|C\|CCCCCCaCTCCAGTCCTTCCCCAAGGATGCAGCCTCAGCCTTCTCCACACCACGTTTCCCCACAGACAAGTTCCCCAgnACTCCAAtCGTGACGACCAT |
| 2:1201:13171:44691#2#F | \| \| |
| MQ=60 isize=207 C=37 | TGGGGGGGCGGAATTGGAGTCCTCAAGTGCGCTCTCCCCAGCCTGTCCCTTCTCCACGGCCACAGTCCCAG\|C\|CCCCCCACTCCAGTCCTTCCCCAAGGAtGcAGCCTCAGCCTTCTCCACaCCACgtttCCCCACAgACAAGTTCCCCACa |
| 1:1213:15885:76075#2#R | \| \| |
| MQ=60 isize=-179 C=33 | tcaagtgcgctctCCCCAGCCTGTCCCTTCtCCACGGCCaCAGTCCCAG\|C\|CCCCCCaCtCCAGTCCTTCCCCAAGGATGCAGCCTCAGCCTTCTCCACACCACGTTTCCCCACAGACAAGTTCCCCACAgnACTCCAATGCCCGTACCCCC |
| 2:1108:6838:2186#2#R | \| \| |
| MQ=60 isize=-179 C=33 | ccaagtgccctctcccccGCctgtCCCTtCcCCaCGgcCaCAGtCCCaG\|C\|CCCCCcactCCAGtCCTTCCCCAAGGATGCAGCCTCAGCCTTCTCCACACCACGTTTCCCCACAGACAAGTTCCCCACanGACTCCAATTGCGTTGCCCGA |
| 2:1203:1523:84945#2#R | \| \| |
| MQ=60 isize=-179 C=15 | tcaagtgcgctctccCCaGCCTGTCCCTTCTCCACGGCCaCaGtCCCAG\|c\|CCTCCCaCtCCAGTCCTTCCCCAAGGATGCAGCCTCAGCCTTCTCCACACCACGTTTCCCCACAGACAAGTTCCCCACAGGACTCCAATATCAAAGACCGT |
| 2:2215:16656:39702#2#R | \| \| |
| MQ=60 isize=-179 C=37 | tCAAGTGCGCTCTCCCCAGCCTGTCCCTTCTCCACGGCCACAGTCCCAG\|C\|CCCCCCACTCCAGTCCTTCCCCAAGGATGCAGCCTCAGCCTTCTCCACACCACGTTTCCCCACAGACAAGTTCCCCACAGGACTCCAAtCAAACCCACGGG |
| 2:2102:4505:56596#2#R | \| \| |
| MQ=60 isize=-139 C=33 | ccacctccaaggcccgggaatcccctattccccctcccaacaagtgccctccccccagccgctcccctctccacggccacagtCCCaG\|C\|CCCCCCcCtCCAGTCCtTCCCCAAGGATGCaGCCTCAGCAGGACTCCAATGGCCATTAGAAA |
| 1:1104:2163:18113#2#F | \| \| |
| MQ=60 isize=310 C=33 | GGCGGGCGCGCGATTGGAGTCntAAGTGCGCTCTCCCCAGCCTGTCCCTTCTCCACGGCCACAGTCCCAG\|C\|CCCCCCACTCCAGTCCTTCCCCAAGGATGCAGCCTCAGCCTTCTCCACACCACGTTTCCCCACAGaCAAGTTCCCCACAT |
| 1:1202:9974:38036#2#R | \| \| |
| MQ=60 isize=-123 C=2 | ccgattctgcccggaccccacacccccaaggccagccgacccctaattccccccccaacaaagtgccctctccccagcctgtcccttctccacggcaacaggccctg\|c\|ccccccactccagtccttccccaccgaagggaaagaaattgtt |
| 2:1110:9196:90991#2#F | \| \| |
| MQ=60 isize=204 C=37 | CTTAAAGTCAATATTGGAGTCCTAGTGCGCTCTCCCCAGCCTGTCCCTTCTCCACGGCCACAGTCCCAG\|C\|CCCCCCACTCCAGTCCTTCCCCAAGGATgCAgCCTCAGCCTTCTCCAcACCaCGTTTCCCCacagacaagttCCCCACAtc |
| 1:1107:20312:39819#2#F | \| \| |
| MQ=60 isize=308 C=33 | AaTATCGAAATTATTGGAGTCCTGTGCgCTCTCCCCAGcCTGTCcCTTCTCCACGgCCACAGTCCCAG\|C\|CcCCCCACTCCAgtccttcCCCAAGgatgcAGcCTCAgCcttCTCCACACCaCGtttCCCCaCAgaCAAGctcCCCACAtCC |
| 1:2203:9327:33236#2#F | \| \| |
| MQ=60 isize=203 C=37 | GTTATATGTTTTATTGGAGTCCTGTGCGCTCTCCCCAGCCTGTCCCTTCTCCACGgCCACAGTCCCAG\|C\|CCCCCCACTCCAGTCCTTCCCCAAGGAtGCAGccTCAGCCTTCTCCaCaCCACGtTTCCCCaCAGACAagtTCCCCACAtCC |
| 1:2211:10405:30424#2#F | \| \| |
| MQ=60 isize=203 C=37 | CATGTAGTGCCCATTGGAGTCCTGTGCGCTCTCCCCAGCCTGTCCCTTCTCCACGGCCACAGT\_\_CAG\|C\|CCCCCCACTCCAGTCCTTCCCCAAGGATGCAGCCTCAGCCTTCTCCACACCACGTTTCCCCACAGACAAGTTCCCCACATCCTG |
| 2:1205:12574:98284#2#F | \| \| |
| MQ=60 isize=203 C=37 | TCGAGACAAATCATTGGAGTCntGTGCGCTCTCCCCAGCCTGTCCCTTCTCCACGGCCACAGTCCCAG\|C\|CCCCCCACTCCAGTCCTTCCCCAAGGATGCAGCCTCAGCCTTCTCCACACCACGTTTCCCCACAGACAAGtTCCCCACATCC |
| 2:1208:8538:98481#2#F | \| \| |
| MQ=60 isize=203 C=33 | TCGAGACAAATCATTGGAGTCntgtGCGCTCTCCCCAGCCTGTCCCTTCTCCACGGCCACAGTCCCAG\|C\|CCCCCCACTCCAGtCCtTCCCCAAGGATGCAGCCTCAGCCTTCTCCACACCACGTTTCCCCACAGACAAGTTCcCCACAtcC |
| 1:1201:3017:21022#2#R | \| \| |
| MQ=60 isize=-182 C=27 | gcgcgctcccCCCAGCCtgcCCCttcTCCaCGGCCaCAgTCCCAG\|C\|CCCCCCaCtCCAGTCCTTCCCCAAGGATGCAGCCTCAGCCTTCTCCACACCACGTTtCCCCACAGACAAGTTCCCCACATCCanGACTCCAATGAGGCGTGCTAT |
| 1:1210:1859:13788#2#F | \| \| |
| MQ=60 isize=202 C=37 | ATTTCACGACCGATTGGAGTCCTGCGCTCTCCCCAGCCTGTCCCTTCTCCACGGCCACAGTCCCAG\|C\|CCCCCCACTCCAGTCCTTCCCCAAGGATGCAGCCTCAGCCTTCTCCACACCACGtTTCCCCACAGACAAGtTCCCCACATCCTg |
| 1:2207:10496:83549#2#F | \| \| |
| MQ=60 isize=202 C=37 | CGCTGAGCCAATATTGGAGTCntGCGCTCTCCCCAGCCTGTCCCTTCTCCACGGCCACAGTCCCAG\|C\|CCCCCCACTCCAGTCCTTCCCCAAGGATGCAGCCTCAGCCTTCTCCACACCaCGTTTCCCCACAGaCAAGTTCCCCaCATCCTG |
| 2:2109:14853:32067#2#F | \| \| |
| MQ=60 isize=202 C=37 | TAGGTTGATTTAATTGGAGTCCTGCGCTCTCCCCAGCCTGTCCCTTCTCCACGGCCACAGTCCCAG\|C\|CCCCCCACTCCAGTCCTTCCCCaagGATGCAGCCTCAGCCTTCTCCACACCACgttTCCCCACAgccaagttccccacatcctg |
| 2:2213:12243:57470#2#F | \| \| |
| MQ=60 isize=202 C=37 | TACCGCAGCCTCATTGGAGTCCTGCGCTCTCCCCAGCCTGTCCCTTCTCCACGGCCACAGTCCCAG\|C\|CCCCCCACTCCAgTCCTTCCCCAAGGATGCAGCCTCAGCCTTCTCCACACCACGTTTCCCCACAGACAAGTTCCCCACATCCTG |
| 1:1203:19837:75495#2#R | \| \| |
| MQ=60 isize=-184 C=22 | gcgctctCCCCAGCctGtCCCTTCTCCACGGCCaCaGTCCCAg\|C\|CCCCcccCTCCAGTCCTTCCCCAAGGATGCAGCCTCAGCCTTCTCCACaCCACGTTTCCCCACAGACAAGTTCCCCACATCCTGanGACTCCAATACCCGCAACTTA |
| 2:2201:7784:31421#2#R | \| \| |
| MQ=60 isize=-169 C=37 | cccccccaaccaagcgcgctccCCCCAGCCtgtcCCTTCTCcacgGCCaCAGTCCCaG\|C\|CCCCCccCtCCAGTCCTTCCCCAAGGATGCAGCCTCAGCCTTCTCCACACCACGTTTCCCCACAGACAAanGACTCCAATACACCACACGAC |
| 2:2209:18890:94846#2#R | \| \| |
| MQ=60 isize=-184 C=33 | gCGCTCTCCCCAGCCTGTCCCTTCTCCACGGCCACAGTCCCAG\|C\|CCCCCCACTCCAGTCCTtccCCAAGGATGCAGCCTCAGCCTTCTCCACACCACGTTTCCCCACAGACAAGTTCCCCACATCCTGanGACTCCAATATTAGTCCGGTG |
| 2:2108:8834:25245#2#F | \| \| |
| MQ=60 isize=200 C=2 | tAtCATCCATTCagggtcctcCgctCTCCCCAGcctgtcccttctccacggccacagtcccac\|c\|ccccccactccagtccttccccaacgatgccgccccagccttcccccccccccgcttccccacgacaaagttcccccccccccgccc |
| 1:1106:19795:22811#2#R | \| \| |
| MQ=60 isize=-186 C=22 | CgCTCTCCCCAGCCTGtCcCTtCTCCaCGGCCaCAGTCCCAg\|C\|CCCCCCACTCCAGTCCTtCCCCAAGGATGCAGCCTCAGCCTTCTCCaCACCACGTTTCCCCACAGACAAGTTCCCCAcATCCTGGanGACTCCAATGCACATATCCTG |
| 1:2210:17336:66493#2#R | \| \| |
| MQ=60 isize=-186 C=33 | cgctctCCCCAGCCTGTCCCTTcTCCACGGCCACAGTCCCAG\|C\|CCCCCCACtCCAGTCCTTCCCCAAGGATGCaGCCTCAGCCTTCTCCACaCCACgTTTCCCCACAGACAAGTTCCCCACATCCTGGAGGACTCCAATCCTCCGACGCCC |
| 2:1202:12029:88500#2#R | \| \| |
| MQ=60 isize=-123 C=2 | tttggcctccgcccccaccccaaaagggccagcagccccataattctccccccatccaaggccgctccccccggcctttcccttctccacgcccacagtcccag\|c\|cccccccctccagtccttccccagngaccccaatcctCCcCCGCCG |
| 2:2213:7126:47743#2#R | \| \| |
| MQ=60 isize=-186 C=27 | cgctctccccagcctgtcccctctccacgGCCACAgTCCCaG\|C\|CCCCCCActCCAGTCCTTCCCCAAGGATGCAGCCTCAGCCTTCTCCACACCACGTTTCCCCACAGACAAGTTCCCCACATCCTGGAGGACTCCAATACGTAACGGACG |
| 1:1211:17313:40255#2#F | \| \| |
| MQ=60 isize=199 C=33 | GCTGGACGTGGGATTGGAGTCCTGCTCTCCCCAGCCTGTCCCTTCTCCACGGCCACAGTCCCAG\|C\|CCCCCCACTCCAGTCCTTCCCCAAGGATGCAGCCTCAGCCTTCTCCACACCACGTTTCCCCACAGACAAGTTCCCCACATCCtgga |
| 1:2216:3587:79850#2#R | \| \| |
| MQ=60 isize=-186 C=33 | cCTCTCCCCAGCCTGTCCCTTCTCCACGGCCACAGTCCCAG\|C\|CCCCCCaCTCCAGTCCTTCCCCAAGGATGCAGCCTCAGCCTTCTCCACACCACGTTTCCCCACAGACAAGTTCCCCACATCCTGGAAgnACTCCAAtGCCCCCAACCCA |
| 1:1112:5782:90133#2#F | \| \| |
| MQ=60 isize=198 C=33 | GTGCCGGAGGACATTGGAGTncTCTCTCCCCAGCCTGTCCCTTCTCCACGGCCACAGTCCCAG\|C\|CCCCCCACTCCAGTCCTTCCCCAAGGATGCAGCCTCAGCCTTCTCCACACCACGTTTCCCCACAGACAAGTTCCCCACATCCTGGAC |
| 1:1215:6547:34372#2#F | \| \| |
| MQ=60 isize=198 C=37 | GTATGTTTTTTCATTGGAGTCCTCTCTCCCCAGCCTGTCCCTTCTCCACGGCCACAGTCCCAG\|C\|CCCCCCACTCCAGTCCTTCCCCAAGGATGCAGCCTCAGCCTTCTCCACACCACGTTTCCCCACAGACAAGTTCCCCACATCCTGGAC |
| 1:2206:4737:45907#2#F | \| \| |
| MQ=60 isize=198 C=37 | CTCACACGTTTGATTGGAGTCntCTCTCCCCAGCCTGTCCCTTCTCCACGGCCACAATCCCAG\|C\|CCCCCCACTCCAGTCCTTCCCCAAGGATGCAGCCTCAGCCTTCTCCACACcaCGTTTCCCCACAGACAAGTTCCCCACATCCTGGAC |
| 1:2213:4398:36603#2#F | \| \| |
| MQ=60 isize=198 C=37 | CTCACACGTTTGATTGGAGTncTCTCTCCCCAGCCTGTCCCTTCTCCACGGCCACAGTCCCAG\|C\|CCCCCCACTCCAGTCCTTCCCCAAGGATGCAGCCTCAGCCTTCTCCACACCACGTTTCCCCACAGACAAGTTCCCCACATCCTGgAC |
| 2:1111:9696:68795#2#F | \| \| |
| MQ=60 isize=198 C=33 | GCCACCGATGGCATTGGAGTCCTCTCTCCCCAGCCTGTCCCTTCTCCACGGCCACAGTCCCAG\|C\|CCCCCCAcTCCAgtCCTtCCCCaAGGaTGCAGCCtcagCCTTCTCCaCACCaCgtttCCCCACagACAAGtTCCCCacaTCctggac |
| 2:1216:2383:59805#2#F | \| \| |
| MQ=60 isize=198 C=37 | GACGGTGTCGTCATTGGAGTncTCTCTCCCCAGCCTGTCCCTTCTCCACGGCCACAGTCCCAG\|C\|CCCCCCACTCCAGTCCTTCCCCAAGGATGCAGCCTCAGCCTTCTCCACACCACGTTTCCCCaCAGACAaGTTCCCCACATCCTGGac |
| 2:2114:11558:89728#2#F | \| \| |
| MQ=60 isize=198 C=33 | AACGCGGCCCCGATTGGAGTncTCTCTCCCCAGCCTGTCCCTTCTCCACGGCCACAGTCCCAG\|C\|CCCCCCACTCCAGTCCTTCCCCAAGGATGCAGCCTCAGCCTTCTCCACACCACGTTTCCCCACAGACAAGTTCCCCACATCCtGGAC |
| 2:2211:12065:5298#2#F | \| \| |
| MQ=60 isize=198 C=37 | CTGGGTTACGGCATTGGAGTCCTCTCTCCCCAGCCTGTCCCTTCTCCACGGCCACAGTCCCAG\|C\|CCCCCCACTCCAGTCCTTCCCCAAGGATGCAGCCTCAGCCTTCTCCACACCACGTTTCCCCACAGACAAGTTCCCCACATCCTGGAC |
| 1:1112:14834:82560#2#R | \| \| |
| MQ=60 isize=-187 C=27 | CTCTCCCCAGCCTGTCCCTTCTCCACGGCCACAGTCCCAG\|C\|CCCCCCACTCCAGTCCTTCCCCAAGGATGCAGCCTCAGCCTTCTCCACACCACGTTTCCCCACAGACAAGTTCCCCACATCCTGGACAgnACTCCAATTGGGGAATCGAC |
| 1:1211:12484:35217#2#R | \| \| |
| MQ=60 isize=-187 C=22 | CTCTCCCCAGCCTGTCCCTtCTCCACggCCACaGTCccag\|C\|CCCCCCACTCCAGTCCTTCCCCAAGGATGCAGCCTCAGCCtTCTCCACACCACGTTTCCCCACAGACAAGTTCCCCACATCCTGGACAGGACTCCAATCCGCCCCGGCCG |
| 1:2104:12765:86674#2#R | \| \| |
| MQ=60 isize=-187 C=37 | ctctCCCCAGCCtGTCCCTTCTCCaCGGCCACAGTCCCAG\|C\|CCCCCCACTCCAGTCCTTCCCCAAGGGTGCAGCCTCAGCCTTCTCCACACCACGTTTCCCCACAGACAAGTTCCCCACATCCTGGACanGACTCCAATTCGTGCCACTTG |
| 1:2211:14114:12115#2#R | \| \| |
| MQ=60 isize=-187 C=27 | ctCtCCCCaGCCTGTCCCtTCcCCaCGGCCaCaGTCCCaG\|C\|CCCCCCACTCCAGTCCTTCCCCAAGGATGCaGCCTCAGCCTTCTCCACACCACGTTTCCCCACAGACAAGTTCCCCACATCCTGGACAGGACTCCAATCAAAAAATGCAG |
| 2:1210:13260:79747#2#R | \| \| |
| MQ=60 isize=-187 C=33 | CtCTCCCCAGCCTGTCCCTTCTCCACGGCCACAGTCCCAG\|C\|CCCCCCACTCCAGTCCTTCCCCAAGGATGCAGCCTCAGCCTTCTCCACACCACGTTTCCCCACAGACAAGTTCCCCACATCCTGGACAGGACTCCAATGTACATGGTTCG |
| 2:1215:12948:29055#2#R | \| \| |
| MQ=60 isize=-186 C=37 | CTCTCCCCAGCCTGTCCCTTCTCCACGGCCACAGTCCCAG\|C\|CCCCCCACTCCAGTCCTTCCCCAAGGATGCAGCCTCAGCCTTCTCCACACCACGTTTCCCCACAGACAAGTTCCCCACATCCTGGACAgnACTCCAATTCGTGCCACTTG |
| 2:2112:17420:17090#2#R | \| \| |
| MQ=60 isize=-187 C=33 | ccctCCCCAGCCTgtCCCTTCtCCACGGCCACAGtCCCAG\|C\|CCCCCCACtCCAGTCCTTCCCCAAGGATGCAGCCTCAGCCTTCTCCACACCACGTTTCCCCACAGACAAGTTCCCCACATCCTGGACAgnACTCCAATTTACACGCACGC |
| 2:2206:18445:8540#2#R | \| \| |
| MQ=60 isize=-187 C=27 | ccctccccagcctgtccctTCTCcACGGCCACAGTCCCaG\|C\|CCCCCCaCTCCAGTCCTTCCCCAAGGATGCAGCCtCaGCCTtCTcCACACCACGTTTCCCCACAGACAAGTTCCCCACATCCTGGAcanGaCTCCaagCCTAGCCCCGCT |
| 1:2209:19903:28594#2#F | \| \| |
| MQ=60 isize=197 C=37 | TAATTAGGTGCAATTGGAGTCntTCTCCCCAGCCTGTCCCTTCTCCACGGCCACAGTCCCAG\|C\|CCCCCCACTCCAGTCCTTCCCCAAGGATGCAGCCTCAGCCTTCTCCACACCACGTTtCCCCACAGACAAGTTCCCCACATCCTGGACc |
| 1:1111:13024:55549#2#F | \| \| |
| MQ=60 isize=196 C=27 | GTACAGCAGGGAATTGGAGTCCCTCCCCAGCCTGTCCCTTCTCCACGGCCACAGTCCCAG\|C\|CCCCCCACTCCAGtCCTTCCCCAAGGATGCAGCCTCAGCCTTCTCCACACCACGTTTCCCCACAgACAAgtTCCCCACaTCCTggactgg |
| 2:1103:8874:15261#2#F | \| \| |
| MQ=60 isize=196 C=33 | GGTCGCGGTTGTATTGGAGTCCTCCCCAGCCTGTCCCTTCTCCACGGCCACAGTCCCAG\|C\|CCCCCCACtCCAGtCCTTCCCCaAGGAtgCagCCtCaccCTtCtCCCccccaCgtTTCCCcccagacaagttccccacaccctggactgga |
| 1:2111:2936:19602#2#R | \| \| |
| MQ=60 isize=-189 C=33 | CTcCCCaGCCTGTCCCttctcCACGGCcacaGtCcCAG\|C\|CCCCcCACTCCAGTCCTcCCCCAAGGATGCAGCCTCAGCCTTCTCCACACCACGTtTCCCCACAGACAAGTTCCCCACAtCCTGGACTgAGGACTCCAATGCGACCCGAGAA |
| 2:1209:4889:7972#2#R | \| \| |
| MQ=60 isize=-189 C=37 | ctCCCCAGCctgTCCCTTCTCCACggcCaCAGTCCCaG\|C\|CCCCCCaCTCcAgTCCtTCCCcAAGGATGCAGCCTCAGCCTTCTCCACACCACGTTTCCCCACAGACAAGTTCCCCACATCCTGGACTGAGGACTCCAATCCACCCAGGCAC |
| 1:1106:21025:19470#2#F | \| \| |
| MQ=60 isize=195 C=33 | GGTCGCGGTTGTATTGGAGTCntcCCCAGCCTGTCCCTTCTCCACGGCCACAGTCCCAG\|C\|CCCCCCACTCCAGTCCTTCCCCAAGGATGCAGCCtCaGCcttCTCCACACCACGtTTCCCCACAGACAAGTTCCCCACATCctGGaCtGGG |
| 1:2106:3964:41871#2#F | \| \| |
| MQ=60 isize=195 C=37 | TGTCATATGTGGATTGGAGTCntcCCCAGCCTGTCCCTTCTCCACGGCCACAGTCCCAG\|C\|CCCCCCACTCCagtCCTTCCCCAAGGATGCAGCCTCAGCCTTCTCCACACCACGtTTCCCCACAGACAAGTTCCCCACATCCTGGaCTGGt |
| 2:1104:14439:55584#2#R | \| \| |
| MQ=60 isize=-190 C=2 | cccccagcctgtcacttctccacggccgcagtcccag\|c\|ccccccactccagTCCttCCCcAAgGATGCaGcctCAGCCTTCTCcacacCACGtttccCCACAGACAAGTTCCCCACATCCtGgACTGGAGGACTCcAATCGCGCCGCGAtG |
| 1:2103:13691:71100#2#F | \| \| |
| MQ=60 isize=194 C=15 | TAGTAGtCTACCATTGGAGTCCCCCAgCCTGTCCCCtCcCcacGGCcAcAGtcCCAc\|c\|CCCCCccccccagtcctTCCCCAaggctgcagcctcagccttcccccccccccgcttccccacaaccaagttccccacatcctgacctggaac |
| 1:1215:19829:37592#2#R | \| \| |
| MQ=60 isize=-193 C=33 | CCCCagCCtgtcCCtTCtCCaCgGCccCaGTCCCaG\|C\|CCCCCCaCtCCAGtCCTTCCCCAAGGATGCAGCCTCAgCCTTCTCCACACCACGTTTCCCCACAGACAAGTTCCCCACATCCTGGACTGGTAGGACTCCAATTAAACCACGGGG |
| 1:2210:19124:17117#2#R | \| \| |
| MQ=60 isize=-162 C=33 | cctacctctccccccaaacaggggcccccccCCCAGCCtGtcccttCtccaCGgCCAcagTCCCAG\|C\|CCCCCCacTCCAGtCCTTCCCCAAGGATGCagcCTCAGCCTTCtCCaCACCACGTTTCCCCAGGACTCCAATAtCTTCTAGAAC |
| 1:2215:16027:49099#2#R | \| \| |
| MQ=60 isize=-193 C=2 | cccgagcctgtccctaccccaagcccactgtcccag\|c\|ccccccactccagtccttccccaaggatgcagcctcagccttctgcacaccccgtttccCCAcAGAcAAgtTcCccaCaTccTgGACTGGTAgnACTCCAATTAAACcaCgggg |
| 2:1102:8705:95791#2#R | \| \| |
| MQ=60 isize=-173 C=2 | cccaacgcagtccccctcccccagcccgtcccttccccgcggccaccgtcccag\|c\|ccccccactccagtCCTTCCCCAAGGATGCaGCcTCaGcCTTCTCcCCaCCACGTTTCCCCacAGaCAAGTTCAGGACTCCAATCGCGctCcCCCT |
| 2:1107:2783:79531#2#R | \| \| |
| MQ=60 isize=-162 C=33 | cccaattccccccccaattaagtgccccccccccagcccgtcCCTTCtCcaCGGCCACAGTCCCaG\|C\|CCCCCCACTCCAGTCCTTCCCCAAGGATGCAGCCTCAGCCTTCTCCACACCACGTTTCCCcanGACTCCAATTGATCGTAATTG |
| 2:2111:3260:70447#2#R | \| \| |
| MQ=60 isize=-140 C=33 | cccctgtaaggccaggaccacccttattccccctccaacccaggccgccccccccagcctgtgccttctccgcggccacagtCCCAg\|C\|CCCCCCCctcCaGTCCtTCCCcAAGGATGCAGCCTCAGCCAGGACTCCAATCCCTCGAGCGGC |
| 1:1109:11724:48632#2#F | \| \| |
| MQ=60 isize=193 C=37 | CACGGGAGACAAATTGGAGTCCTCCCAGCCTGTCCCTTCTCCACGGCCACAGTCCCAG\|C\|CCCCCCACTCCAGtCCTTCCCCAAGGATGCAGCCTCAGCCTTCTCCACACCACGTTT\_CCCACAgaCAAGtTCCCCACATCCTGGaCTGGtag |
| 1:1204:11489:78501#2#F | \| \| |
| MQ=60 isize=193 C=37 | TCAAGCGTGGAGATTGGAGTCntCCCAGCCTGTCCCTTCTCCACGGCCACAGTCCCAG\|C\|CCCCCCaCtCCAGTCCTTCCCCAAGGATGCAGCCTCAGCCTTCtCCACACCACGTtTCCCCaCagACAAGTTCCCCaCATCCTGGaCtGGta |
| 1:2103:1705:78936#2#F | \| \| |
| MQ=60 isize=298 C=37 | GTCACGCTGCGCATTGGAGTCCTCCCAGCCTGTCCCTTCTCCACGGCCACAGTCCCAG\|C\|CCCCCCACTCCAGTCCTTCCCCAAGGATGCAGCCTCAGCCTTCTCCACACCACGTTTCCCCACAGACAAGTTCCCCACATCtggacttgtaa |
| 2:1111:9391:29105#2#F | \| \| |
| MQ=60 isize=298 C=37 | CTAGGAGCCCGCATTGGAGTCCTCCCAGCCTGTCCCTTCTCCACGGCCACAGTCCCAG\|C\|CCCCCCACTCCAGTCCTTCCCCAAGgatgCAGCCTCAGCCTtcTCCcCacCACGttTCCCCACAgacaAGTTCCCCACATCCtggactcgta |
| 1:1108:6180:35901#2#R | \| \| |
| MQ=60 isize=-192 C=33 | CCCAGCCTGTCCCtTCTCCACGGCCACAGTCCCAG\|C\|CCCCCCACTCCAGTCCtTCCCCAAGGATGCAGCCTCAGCCTTCTCCACACCACGTTTCCCCACAGACAAGTTCCCCACATCCTGGACTGGTAanGACTCCAAtCGACTCACTAGC |
| 1:2214:13290:70750#2#R | \| \| |
| MQ=60 isize=-192 C=33 | CCCAGCCTGTCCCTTCTCCACGGCCACAGTCCCaG\|C\|CCCCCCACTCCAGTCCTTCCCCAAGGATGCAGCCTCAGCCTTCTCCACACCACGTTTCCCCACAGACAAGTTCCCCACATCCTGGACTGGTAAgnACTCCAaTCCCGAGCGGGTC |
| 2:2204:20398:95911#2#R | \| \| |
| MQ=60 isize=-192 C=33 | CCCagcCTGTcCCTTCTCcaCGGcCACAGTCcCAG\|C\|CcCCCCaCtCcAGTCCTTCCCCAAGGATGCAGCCTCAGCCTTCTCCACAcctCGTTTCCCCACAGACAAGTTCCCCACATCCTGGAcTGGTAanGACTCCAATGAACCTCCTGAG |
| 1:1111:2562:56429#2#F | \| \| |
| MQ=60 isize=192 C=33 | CGCCGACGCTAAATTGGAGTCCTCCAgCCTGTCCCTTCTCCACGGCCACAGTCtCAG\|C\|CCCCCCACTCCaGtcCTTCCCCAAGGaTGCAgcctCAGcCtTCtcCAcaCCacGtTtcCCCACAGACaAgttcCCCACatCCcgGAcTGgTAG |
| 1:1212:17656:88159#2#F | \| \| |
| MQ=60 isize=192 C=33 | GCCTCTTCGACCATTGGAGTCCTCCAGCCTGTCCCTTCTCCACGGCCACAGTCCCAG\|C\|CCCCCCACTCCAGTCCTTCCCCAAGGATGCAGCCTCAGCCTTCtCCACACCaCGtTTCCCCACAGacAAGTTCCCCACATCCtGGaCTGgtag |
| 1:2106:13516:96836#2#F | \| \| |
| MQ=60 isize=192 C=27 | ACACGTGTAGTGATTGGAGTCntCCAGCCTGTCCCTTCTCCACGGCCACgGTCCCAG\|C\|CCCCCCACTCCAGTCCTTCCCCAAGGATGCAGTCTCAGCCTTCTCCACACCACGTTTCCCCACAGACAAGTTCCCCACATCCTGGACTGGTAG |
| 1:2206:11758:93372#2#F | \| \| |
| MQ=60 isize=192 C=33 | TAGCCTGACGCGATTGGAGTCntCCAGCCTGTCCCTTCTCCACGGCCACAGTCCCAG\|C\|CCCCCCACTCCAGTCCTTCCCCAAGGATGCAGCCTCAGCCTTCTCCACaCCaCGCTTCCCCacagaCAAGTTCCCCacatCCTGGACTGgaag |
| 2:1201:2856:86560#2#F | \| \| |
| MQ=60 isize=192 C=40 | ACACGTGTAGTGATTGGAGTCCTCCAgCCTGTCCCTTCTCCACGGCCACAGTCCCAG\|C\|CCCCCCACTCCAGTCCTTCCCCAAGGATGCAGCCTCAGCCTtCTCCAcaCCACGTTTCCCCaCAGACAagttCCCCACAgcCtggactggtag |
| 2:2201:17879:94328#2#F | \| \| |
| MQ=60 isize=192 C=33 | CCGAGGAGGGGCATTGGAGTCntCCAGCCTGTCCCTTCTCCACGGCCACAGTCCCAG\|C\|CCCCCCACTCCAGTCCTTCCCCAAGGATGCAGCCTCAGCCTTCTCCACACCACGtTTCCCCACAGACAAGTTCCCCACATCCTGGACTGGTAg |
| 2:2208:9587:80595#2#F | \| \| |
| MQ=60 isize=192 C=33 | GTGGGGCGGGCCATTGGAGTCntcCAGCCTGTCCCTTCTCCACGGCCACAGTCCCAG\|C\|CCCCCCACTCCAGTCCTTCCCCAAGGATGCAGCCTCAGCCTTCTCCACaCCACGTTTCCCCACAGACAAGTTCCCCACATCCTGGACTGGtag |
| 2:1208:5942:36365#2#R | \| \| |
| MQ=60 isize=-194 C=33 | CAGCCtGTCCCtTCTCCACGGCCACAGTCCCcG\|C\|CCCCCCACTCCAGtCcTTCCCCAAGGATGCAGCCTCAGCCTTCTCCaCACCACGTTTCCCCACAGACAAGTTCCCCACATCCTGGACTGGTAGCanGACTCCAATACCCGAGGTGAT |
| 1:1111:15364:56053#2#F | \| \| |
| MQ=60 isize=190 C=37 | TCTCTCTCTCCGaTTGGAGTCCTAGCCTGTCCCTTCTCCACGGCCACAGTCCCAG\|C\|CCCCCCACTCCAGTCCTTCCCCAAgGATGCAGCCTCAGCCTTCTCCaCACCACGTTtCCCCACAGACAAGTTCCCCACATCCTggaCTGGtaGCt |
| 1:1108:5778:92966#2#R | \| \| |
| MQ=60 isize=-195 C=33 | agcctgtcCCTTCTCCACGGCCaCAGTCCCAG\|C\|CCCCCCACTCCAGTCCTTCCCCAAGGATGCAGCCTCAGCCTTCTCCACACCACGTTTCCCCACAGACAAGTTCCCCACATCCTGGACTGGTAGCTanGACTCCAATCGCTGACGTATA |
| 2:2216:9546:84760#2#R | \| \| |
| MQ=60 isize=-195 C=37 | aGCCTGTCCCTTCTCCACGGCCACAGTCCCAG\|C\|CCCCCCACTCCAGTCCTTCCCCAAGGATGCAGCCTCAGCCTTCTCCACACCACGTTTCCCCACAGACAAGTTCCCCACATCCTGGACTGGTAGCTAGGACTCCAATTGAGGACGGCCT |
| 1:2106:5987:63988#2#F | \| \| |
| MQ=60 isize=189 C=37 | TTCCGCTGTTCCATTGGAGTCntGCCTGTCCCTTCTCCACGGCCACAGTCCCAG\|C\|CCCCCCACTCCAGTCCTTCCCCAAGGATGCAGCCTCAGCCTTCTCCACACCACGTTTCCCCACAGACAAGTTCCCCACATCCTGGACTGGTAgCTg |
| 1:1104:14047:74709#2#R | \| \| |
| MQ=60 isize=-187 C=33 | cccccccccgcctgtcccccctCCaCggCcaCagtCCcaG\|C\|CCCCCcactCCAGTCCttCCCCAAggATGCAGCCTCAGCCTTCTCCACACCACGTTTCCCCACAGACAAGTTCCCCACATCCTGGAcanGACTCCAAtCTATATCTCTGA |
| 2:1116:13097:54138#2#R | \| \| |
| MQ=60 isize=-196 C=27 | cCCTgtCCCttCtCCACGGCCACAGtCCCag\|C\|CCCCCCaCTCCAGTCCTTCCCCAAGGATGCAGCCTCAGCCTTCTCCACACCACGTTTCCCCACAGACAAGTTCCCCACATCCTGGACTGGcAGCTGAGGACTCCAATGACCGACAGCTA |
| 1:1201:8957:83333#2#F | \| \| |
| MQ=60 isize=188 C=33 | AGTGTTTGTCGAATTGGAGTCntCCTGTCCCTTCTCCACGGCCACAGTCCCAG\|C\|CCCCCCACTCCAGTCCTTCCCCAAGGATGCAGCCTCAGCCTTCTCCacACCacGTTTCCCCACGGACAAGttCCCCACATCCTgGACTGGtagctgc |
| 1:2214:13335:96316#2#F | \| \| |
| MQ=60 isize=188 C=37 | AGTGTTTGTCGAATTGGAGTncTCCTGTCCCTTCTCCACGGCCACAGTCCCAG\|C\|CCCCCCACTCCAGTCCTTCCCCAAGgATGCAGCCTCAGCCTTCTCCACACCACGTTtCCCCaCAGACAAGTTCcCCACAtCCTGGACTGGTAgCTGC |
| 2:1212:10437:34321#2#F | \| \| |
| MQ=60 isize=188 C=27 | TGCTTACATGCTATTGGAGTCCTGTCCCTTCTCCACGGCCACAGTCCCAG\|C\|CCCCCCACTCCAGTCCTTCCCCAaGGATGCAGCCTCAGCCTTCTCCACACCACGTTTCCCCACaGACAAGTTCCCCACATCCTGGaCTGGtAGCTGCCCa |
| 1:1212:12207:7679#2#R | \| \| |
| MQ=60 isize=-195 C=22 | cCtGgCCCTTcTccaCGGCCACaGTCCCAg\|C\|cCCCCCaCtCCAgtccttcCCCAaggaTgCAGCctcAGCCTTCTCCaCaCCAcgcTTCccCACAGACAaGtTCCCcACATCCTGGACTGGTAgCTGCAGGACTCCAATTGGGCCTAAACC |
| 2:1113:13449:46523#2#R | \| \| |
| MQ=60 isize=-197 C=2 | cctgccacttcaccacggccacagtcccag\|c\|ctcaccactccagtccttcccCAAGGATGCAGcgtCAgCcttctccAcACCACGTttccCCaCAGacAAGTtCCCCACATCCtGGACTGGtAGCTGCAgnACTCCAATAACGTATCaCac |
| 2:2101:19159:18132#2#R | \| \| |
| MQ=60 isize=-195 C=2 | cctgccccttctccacggccacagtcccag\|c\|ccccccaCTCCAgtCCTtccccgAGGATGCAgCcgCAGCCTTCTCCACaCCACGTTtCCCCaCAGACAAGTTCCCcACatCCtGGACTGGTaGCTGCAGGACTCCAATTGGGCCTAAACC |
| 2:2215:3724:88560#2#R | \| \| |
| MQ=60 isize=-195 C=37 | CCTGTCCCTtCTCCACGGCCACAGTCCCAG\|C\|CCCCCCaCTCCAGTCCTTCCCCAAGGATGCAGCCTCAGCCTTCTCCACACCACGTTTCCCCACAGACAAGTTCCCCACATCCTGGACTGGTAGCTGCAGGACTCCAATTGGGCCTAAACC |
| 1:2204:7982:43748#2#F | \| \| |
| MQ=60 isize=186 C=40 | TCGGGAATAGAAATTGGAGTCCTCTGTCCCTTCTCCACGGCCACAGTCCCAG\|C\|CCCCCCACTCCAGTCCTTCCCCAAGGATGCAGCCTCAGCCTTCTCCACACCACGtTTCCCCACAGACAAGtTCCCCACATCCtGGACTGGtAGCTgCC |
| 2:1105:1413:12068#2#F | \| \| |
| MQ=60 isize=187 C=37 | ACGGCCTGACGTATTGGAGTCntCTGTCCCTTCTCCACGGCCACAGTCCCAG\|C\|CCCCCCcCTCCagtccTTCCCcaaggatgcagcctcagccttctcccacccacgttcccccccaaccagattccccccacccgggcctggagccgccc |
| 2:2115:15255:64375#2#F | \| \| |
| MQ=60 isize=292 C=37 | CAATGTTTACTAATTGGAGTncTCTGTCCCTTCTCCACGGCCACAGTCCCAG\|C\|CCCCCCACTCCAGTCCTTCCCCAAGGATGCAGCCTCAGCCTTCTCCACACCACGTTTCCCCaCAGACAAGTTCCCCACATCCTGGACTGGTAGCTGCC |
| 1:1116:7439:62487#2#R | \| \| |
| MQ=60 isize=-198 C=33 | CTGTCCCTtCtCCaCGGCCACAGTCCCAG\|C\|CCCCCCaCTCCAGTCCTTCCCCAAGGATGCAGCCTCAGCCTTCTCCaCACCACGTTTCCCCACAGACAAGTTCCCCACATCCTGGACTGGTAGCTGCCAGGACTCCAATTCGCGAGCCTCG |
| 1:1205:16542:17926#2#R | \| \| |
| MQ=60 isize=-197 C=37 | CTgtcCCTTCtCCACGGCCaCAGTCCCAG\|C\|CCCCCCaCTCCAGTCCTTCCCCAAGGATGCAGCCTCAGCCTTCTCCACACCACGTTTCCCCACAGACAAGTTCCCCACATCCTGGACTGGTAGCTGCCanGACTCCAATCGCCTAGGCTAT |
| 2:1103:14677:6090#2#R | \| \| |
| MQ=60 isize=-198 C=2 | ccgtcccctcgccacggccacagtcacag\|c\|ccccccactccagtccttccccaaggatGCAGCCTCAGCcttCtccaCACCacGtttcCcCACAGACAAGTTCCgCaCAtCCtGGACTGGTAGCTGCCAGGACTCCAATCACCCTCGAGAG |
| 2:2103:20418:9721#2#R | \| \| |
| MQ=60 isize=-198 C=37 | ctgtcccttctccccggccccAGtCCCAG\|C\|CCCCcCactCCAGtCCttCCCCAAGGATGCAGCCTCAGCCTTCTCCACACCACGTTTCCCCACAGACAAGTTCCCCaCATCCTGGACTGGTAGCTGCCAGGACTCCAATAGAAATAGAGAG |
| 2:2115:11350:13659#2#R | \| \| |
| MQ=60 isize=-198 C=33 | ctgtcccttccccacgGCCaCAGTCCCAG\|C\|CCCCCCaCTCCAGTCCTTCCCCAAGGATGCAGCCTCAGCCTTCTCCACACCACGTTTCCCCACAGACAAGTtCCCCACATCCTGGACTGGTAGCTGCcAgnACTCCAATAAACCCCCCGCG |
| 2:1111:19892:11927#2#F | \| \| |
| MQ=60 isize=186 C=37 | GTAGTGGCACCGATTGGAGTCCTTGTCCCTTCTCCACGGCCACAGTCCCAG\|C\|CCCCCCACTCCAGTCCTTCCCCAagGATGCAGCCTCAGCCTTCTCCACACCACGtTTCCCCACAGACAAGttCCCCACATCCtggactggtagctgccc |
| 2:2208:7302:95501#2#F | \| \| |
| MQ=60 isize=320 C=2 | tgctcgagccacattggagccntacaggatgtcccttctccacagccagagccccac\|c\|ccccccactccagtccttccccaggggtgcagtctaatccttctccccaccacgcttcccatcagcctagttctctccatccttgcctgattc |
| 1:1202:19260:40197#2#R | \| \| |
| MQ=60 isize=-198 C=22 | tgtctcttctcCACGgCCaCAGtCCCAg\|C\|CCCCCCaCtccgGTCCTTCccCAAgGATGCAGCCTCAGCCTTCTCcaCAcCaCGTTTcCCCaCAGACAAGTTCCCCACATCCTGGACTGGTAGCTGCCAGGactcCAAtGCCTTTCAGAGCC |
| 1:1215:4260:93146#2#R | \| \| |
| MQ=60 isize=-198 C=37 | tgtCCCTtCTCCACGGCCACAGTCCCAG\|C\|CCCCCCaCTCCAGTCCTTCCCCAAGGATGCAGCCTCAGCCTTCTCCACACCACGTTTCCCCACAGACAAGTTCCCCACATCCTGGACTGGTAGCTGCCAGGACTCCAATGCCTTTCAGAGCC |
| 1:1110:7072:94762#2#R | \| \| |
| MQ=60 isize=-202 C=33 | gtcccttctccaCGGCCACAgtCCCAG\|C\|CCCCCcaCTCCAGTCCTTCCCCAAGGATGCAGCCTCAGCCTTCTCCaCACCACGTTTCCCCACAGACAAGTTCCCCACATCCTGGACTGGTAGCTGCCCAGGGACTCCAATCCCCGATCACCG |
| 2:2204:5367:82689#2#R | \| \| |
| MQ=60 isize=-200 C=15 | ctcccttctccccggccaccgtCCCag\|c\|CCCCCcaCtCCAGTCCTTCCCcAAGGAtGCaGCCTCAGCCTTCTCcaCACCACGTTTCCACACAGACAAGTTCCCCACATCCTGGACTGGTAGCTGCCCAanGACTCCAATTGCACCATTGGC |
| 1:1107:16377:42363#2#F | \| \| |
| MQ=60 isize=184 C=40 | GACCGATGGATGATTGGAGTCCTCCCTTCTTCACGGCCACAGTCCCAG\|C\|CCCCCCACTCCAGTCCTTCCCCAAGGATGCaGCCTCAGCCTTCTCCACACCACGTTTCCCCACAGACAAGTTCCCCACaTCCTGGaCTGGTAGCTGCCCAGg |
| 1:1206:7145:45268#2#F | \| \| |
| MQ=60 isize=184 C=37 | GATATCAGCTCTATTGGAGTCCTCCCTTCTCCACGGCCACAGTCCCAG\|C\|CCCCCCACTCCAGTCCTTCCCCAAGGATGCAGCCTCAGCCTTCTCCACACCACGtTTCCCCACAGACAAGTTCCCCACATCCTGGACTGGTAGCTGCCCAGg |
| 1:2102:18348:71703#2#F | \| \| |
| MQ=60 isize=289 C=40 | GTTGTCATACATATTGGAGTCCTCCCTTCTCCACGGCCACAGTCCCAG\|C\|CCCCCCACTCCAGCCCTTCCCCAAGGATGCAGCCTCAGCCTTCTCCACACCacGTTTCCCCACAGACAaGtTCCCCACAtCCTGGaCTGGTAGCTGCCCAGg |
| 2:1116:7110:44871#2#F | \| \| |
| MQ=60 isize=184 C=40 | GTAAACATACCCATTGGAGTCCTCCCTTCTCCACGGCCACAGTCCCAG\|C\|CCCCCCACTCCAGTCCTTCCCCAAGGATGCAGCCTCAGCCTTCTCCACACCACGTTTCCCCACAGACAAgTTCCCCACATCCTGGaCTGGtagcTgcccagc |
| 2:1207:9316:96125#2#F | \| \| |
| MQ=60 isize=184 C=37 | TGCAAGACCCAGATTGGAGTCntCCCTTCTCCACGGCCACAGTCCCAG\|C\|CCCCCCACTCCAGTCCTTCCCCAAGGATGCAGCCTCAGCCTTCTCCACACCACGTTTCCCCACAGaCAAGTTCCCCACATCCTGgACTGGtAGCTGCCCAgg |
| 2:2206:8336:9460#2#F | \| \| |
| MQ=60 isize=183 C=40 | GTCCATCTGGAGATTGGAGTCntCCCTTCTCCACGGCCACAGTCCCAG\|C\|CCCCCCACTCCAGTCCTTCCCCAAGGATGCAGCCTCAGCCTTCTCCACACCACGTTTCCCCACAgACAAGTTCCCCaCAtCCTGGACTGGTAGCTGCCCAGg |
| 1:2103:16765:96910#2#R | \| \| |
| MQ=60 isize=-201 C=33 | tcCCttCTCCACGGCCcCAgTCCCAG\|C\|CCCCCCaCTCCAGTCCTTCCCCaAGGATGCAGCCTCAGCCTTCCCCACACCACGTTTCCCCACAGACAAGTTCCCCACATCCTGGACTGGTAGCTGCCCAGAGGACTCCAATCCACGGGAGCCC |
| 2:1102:10710:50587#2#R | \| \| |
| MQ=60 isize=-198 C=2 | cctcccttcaccaccgccccagccccag\|c\|cccccccctCCagtCCttCCCCAAgGatgCaGCCTcAGCCttctcCACACcacGgTTCCCCaCAGACAAGtTCCCCACATCCTGGaCTGGTAGCTGCCaGGACTCCAAATAGCGGACGCGGC |
| 2:2113:7125:10840#2#R | \| \| |
| MQ=60 isize=-201 C=2 | tcccttctccacggccacagtcccag\|c\|ccccccacgccagtCCTTCCCCAAGGATgCAGCCTCAGCCTTCTCCACaCCACGTTTCCCCACAGACAAGTTCCCCACATCCTGGACTGGTAGCTGCCCAGAgnACTCCAATTCGCGTTGGTGG |
| 1:2104:20324:89268#2#F | \| \| |
| MQ=60 isize=183 C=37 | TCGCCCTTCCTCATTGGAGTCncCCTTCTCCACGGCCACAGTCCCAG\|C\|CCCCCCACTCCAGTCCTTCCCCAAGGATGCAGCCTCAGCCTTCTCCACACCaCGTTTCCCCACAGACAAGTTCCCCACATCCTGGACTGGTAgCTGCCCAGGC |
| 2:1111:17428:12043#2#R | \| \| |
| MQ=60 isize=-160 C=2 | ccctaactctctctccacacaagttccccccccccagccgtccccttctccgcggccacagtcccag\|c\|cccccccccccagtccttccccAAGGAtGCAGCcTcAGCCTTCTCcaCcCCacGtTtCCcaagactccaatTAGTTGgtcTCA |
| 1:2102:15229:90780#2#F | \| \| |
| MQ=60 isize=182 C=37 | GGGTAATAACTTATTGGAGTCCTCCTTCTCCACGGCCACAGTCCCAG\|C\|CCCCCCACTCCAGTCCTTCCCCAAGGATGCAGCCTCAGCCTTCTCCACACCACGTTTCCCCACAGACAAGtTCCCCACATCCTGGACTGGTAGCTGCCCAGGC |
| 2:1103:4602:88594#2#F | \| \| |
| MQ=60 isize=182 C=33 | TGcGAATCGCATATTGGAGTCCTTCtCCACGgCCACAGcCcCAG\|C\|CCCCcCACTCCAGtCCTtCCCCAaggatccaccctcagcctcctccacaccacgttcccccacagaaaagttccccacatcctgaacgggaagcgccccagaccac |
| 1:1101:7409:11057#2#R | \| \| |
| MQ=60 isize=-203 C=37 | CCTTCTCCACGGCCACAGTCCCAG\|C\|CCCCCCACTCCAGTCCTTCCCCAAGGATGCAGCCTCAGCCTTCTCCACACCACGTTTCCCCACAGACAAGTTCCCCACATCCTGGACTGGTAGCTGCCCAGGCanGACTCCAATAGCGCACACCCA |
| 1:1202:16246:61915#2#R | \| \| |
| MQ=60 isize=-203 C=27 | ccttctCCaCgGCCaCAGTCCCAG\|C\|CCCCCCACTCCAGTCCTTCCCCAAGGATGCaGCCTCAGCCTTCTCCACACCACGTTTCCCCACAGACAAGTTCCCCACATCCTGGACTGGTAGCTGCCCAGGCAGGACTCCAATCACGGCGCCCAC |
| 1:2208:5656:73326#2#R | \| \| |
| MQ=60 isize=-203 C=33 | CCTTCtCcaCGGCCacagtCCCAG\|C\|CCCCCCACTCCAGTCCTTCCCCAAggatGCAGCCTCAGCCTgCTCCacaCCACGTTTCCCCACAGACAAGTTCCCCACATCCTGGACTGGTAGCTGCCCAGGCanGACTCCAATACCACCCGTACA |
| 1:2212:9125:31349#2#R | \| \| |
| MQ=60 isize=-199 C=2 | ccttctccacggccacagtcccag\|c\|ccCCCCaCTCCAGTCCTTCCCCAAGGATGCAGCCTCAGCCTTCTCCACACCACGTTTCCCCACAGACAAGTTCCCCACATCCTGGACTGGTAGCTGCCCGGGCAGGACTCCAATCCCCGTCTACCG |
| 2:1105:17963:14165#2#R | \| \| |
| MQ=60 isize=-203 C=22 | ccttccccacggcCACAGTCCCag\|C\|CCCCccaCTCCAGTCCtTCCCCAAGGATGCAGCCTCAGCCTTctCCACACCaCGTttCCCCACAGACAAGTTCCCCACATCCTGGACTGGTAGCTGCCCAGGcanGACTCCAATTATTTTCTTGAA |
| 2:1206:12170:8282#2#R | \| \| |
| MQ=60 isize=-203 C=22 | CCTTCTCCACgGCCACAGtCCCAG\|C\|tCCCCCaCTCCAGTCCtTCCCCAAGGATGCAGCCTCAGCCTtCCCCACACCACGTTTCCCCACAGACAAGTTCCCCACATCCTGGACTGGTAGCTGCCCAGGcanGACTCCAATAATTGACGGTGA |
| 2:2215:6421:60462#2#R | \| \| |
| MQ=60 isize=-203 C=33 | CCcTCTCCaCGGCcacagtCCccG\|C\|CCCCCcaCTCCAGTCCTTCCCCAAGGATGCAGCCTCAGCCTTCTCCACACCACGTTTCCCCACAGACaAGTTCCCCACATCCTGGACTGGTAGCTGCCCAGGCAGGACTCCAATCAAGAAGACGAC |
| 2:1214:13108:46080#2#F | \| \| |
| MQ=60 isize=315 C=37 | TATCCTCCTCGTATTGGAGTCCTCTTCTCCACGGCCACAGTCCCAG\|C\|CCCCCCACTCCAGTCCTTCCCCAAGGATGCAGCCTCAGCCTTCTCCACACCACGTTTCCCCACAGACAAGTTCCCCACATCCTGGaCTGGTAGCTGCCCAgGCC |
| 1:1102:14902:35380#2#R | \| \| |
| MQ=60 isize=-205 C=37 | CTTCTCCACGGCCACaGTCCCaG\|C\|CCCCCCACTCCAGTCCTTCCCCAAGGATGCAGCCTCAGCCTTCTCCACACCACGTTTCCCCACAGACAAGTTCCCCACATCCTGGACTGGTAGCTGCCCAGGCcanGACTCCAATCAGTGAATACTA |
| 1:1214:17411:33343#2#R | \| \| |
| MQ=60 isize=-205 C=33 | CTTCtCcaCGGCCACAGTCCCAG\|C\|CCCCCCaCTCCAGTCCTTCCCCAAGGATGCAGCCTCAGCCTTCTCCACACCACGTTTCCCCACAGACAAGTTCCCCACATCCTGGACTGGTAGCTGCCCAGGCCAgnACTCCAATAAATGCTCAACA |
| 2:1209:11658:41425#2#R | \| \| |
| MQ=60 isize=-205 C=37 | CTTCTCCACGGCCACAGTCCCAG\|C\|CCCCCCACTCCAGTCCTTCCCCAAGGATGCAGCCTCAGCCTTCTCCACACCACGTTTCCCCACAGACAAGTTCCCCACATCCTGGACTGGTAGCTGCCCAGGCCAGGACTCCAATTATCACTTTTCC |
| 2:2202:4973:44987#2#R | \| \| |
| MQ=60 isize=-205 C=27 | cttctccccggccccagtccCAG\|C\|CCCCCCaCtCCAGTCCtTCCCCAAGGATGCAGCCTCAGCCTTCTCCACACCACGTTTCCCCACAGACAAGTTCCCCACATCCTGGACTGGTAGCTGCCCAGGCCAGGACTCCAATGTTCCTAGGTCA |
| 2:2215:20981:90327#2#R | \| \| |
| MQ=60 isize=-205 C=2 | cttctccacggccacagtcccag\|c\|ccccccactccagtccttcccCAAGGATGCAGCCTCAGcctTCTCCaCACCACGTtcCCCCACAGACAAGTTCCCCACATCCTGGACTGGTAGCTgCCCaggcCAGGACTCCAATGCtAtTTTTGAT |
| 2:1108:12925:23562#2#F | \| \| |
| MQ=60 isize=180 C=40 | GGACCAGACGCCATTGGAGTCntTTCTCCACGGCCACAGTCCCAG\|C\|CCCCCCACTCCAGTCCTTCCCCAAGGATGCAGCCTCAGCCTTCTCCACACCACGttTCCCCAcaGaCAAGTTCCCCacAtCCTGGACtGGtAGCtgCCCAGGCCA |
| 2:2215:13002:30980#2#F | \| \| |
| MQ=60 isize=180 C=37 | GGGCCCCAGCTTATTGGAGTCCTTTCTCCACGGCCACAGTCCCAG\|C\|CCCCCCACTCCAGTCCTTCCCCAAGGATGCAGCCTCAGCCTTCTCCACACCACGTTTCCCCACAGACAAGTTCCCCACATCCTGGACTGGtAGCTGCCCAGGCca |
| 2:2215:15607:87127#2#R | \| \| |
| MQ=60 isize=157 C=33 | ttctccaCGGCCACAGTCCCaG\|C\|CCCCCCACTCcAGTCCTTCCCCAAGGATGCAGCCTCAGcCTTCTCCACaCCaCGTTTCCCCACAGACAAGTTCCCCACATCCTGGACTGGTAGCTGCCCAGGCCAAGGACTCCAATCACCTCACCGCC |
| 2:2204:20366:14567#2#F | \| \| |
| MQ=60 isize=179 C=33 | TCCTGGGGaGCCATTGGAGTCntCTCCACGGCCACAGTCCCAG\|C\|CCCCCCACTaCAGTCCTtCacCAAGgATgcAGCCtcaGcCttctcCACACCACGtttccCCACAGACAAGTTcccCACATCCtggactggtagctgccccgtccaac |
| 1:1216:18755:83682#2#R | \| \| |
| MQ=60 isize=-207 C=33 | TCtCCACgGCCaCaGtCCCAG\|C\|CCCCCc\_CTCCAGTCCTTCCCCAAGGATGCAGCCTCAGCCTTCTCCACACCACGTTTCCCCACAGACAAGTTCCCCACATCCTGGACTGGTAGCTGCCCAGGCCAACAgnACTCCAAtCgCACTCCACGC |
| 2:1104:11531:71184#2#R | \| \| |
| MQ=60 isize=-207 C=2 | tccccaccgccacagtcccag\|c\|cccccc\_ctccagtccttccccaaggatgcagcctcagccttctCCACACCACgtTtCCCCACAGACAAGTTCCCCACATCCTGGACTGGTAGCTGCCCAGGCCAACAGGACTCCAATCGCACTCCACGC |
| 2:2207:6337:48800#2#R | \| \| |
| MQ=60 isize=-185 C=27 | tctccacgGCCACAGtCCCaG\|C\|CCCCCCaCTCCAGTCCTTCCCCAAGGATGCAGCCTCAGCCTTCTCCaCACCACGTTTCCCCACAGACAAGTTCCCCACATCCTGGCCTGGGCAGCTACCAGTCCAganGACTCCAAtCAGGTCGGGCAC |
| 1:2214:6443:16508#2#F | \| \| |
| MQ=60 isize=312 C=6 | ATGTCgtTCCGTATTGGAGTncTCCACGGCCACAGTCCCAG\|c\|CCCCCcactccagtccttccccaaggatgcagcctcagccttctccacaccacgtttccccacagacaagtttcccacatcctggacttgtagctgcccacgccaacac |
| 1:1106:17389:43121#2#R | \| \| |
| MQ=60 isize=-207 C=2 | ctccacggccccagtcccag\|c\|ccccccactccagtccttccccaaggatgcagcctcAGCCTTcTCCACACCACGTTTCCCCACAGACAAGTTCCCCaCATCCTGGACTGGTAGCTGCCCAGGCCAAcanGGCTccaatcCCCCCTCGATG |
| 1:1204:14520:7143#2#R | \| \| |
| MQ=60 isize=-207 C=33 | ctccacggccacagtCCCAG\|C\|CCcCccacTCCAGTCCttCcCcAAGGATGCAGCCTCAGCCTTCTCCaCaCCaCGTTTCCCCACAGACAAGTTCCCCACATCCTGGACTGGTAGCTGCCCAGGCCAAcanGACTCCAATGCTTCCCCCGCC |
| 1:2204:8853:56634#2#R | \| \| |
| MQ=60 isize=-205 C=2 | ctccacggccacagtcccgg\|c\|CCCcCtcCtCCAGTCCTTCCCcAAGGATgCAGCCTCAGCCTTCTCCACACCACGTTTCCCCACAGACAAGTTCCCCaCATCCTGGACTGGTAGCTGCCCAGGCCAGgaaTCCACCAAtCaCCGCCCACCT |
| 1:2209:5410:34185#2#R | \| \| |
| MQ=60 isize=-207 C=33 | CtCCACGGCCACAGTCCCAG\|C\|CCCCCcaCTCCAGTCCTTCCCCAAGGATGCAGCCTCAGCCTTCTCCACaCCACGTTTCCCCACAGACAAGTTCCCCGCATCCTGGGCTGGTAGCTGCCCAGGCCAacanGACTCCAaTGCTTCCCCCGCC |
| 2:1104:18779:16808#2#R | \| \| |
| MQ=60 isize=-207 C=22 | ccccacgGCcaCAGTCCCac\|C\|CCCCCCaCtCCAGTCCTTCCCCAAGGATGCAGCCTCAGCCTtCTCCACACCACGTttCCCCACAGACAAGTTCCCCACATCCTGGACTGGTAGCTGCCCAGGCCAACAGGACTCCAATTGACACCGGAGC |
| 2:2107:9475:77122#2#R | \| \| |
| MQ=60 isize=-207 C=27 | ccccacggccccagtcCCAG\|C\|CCCCCCCCtCcagtcCtTCCCCAAGGATGCAGCCTCAGCCTTCTCCACACCaCGTTTCCCCACAGACAAGTTCCCCACATCCTGGACTGGTAGCTGCCCAGGCCAACAGGACTCCAATGATTTTCCACAT |
| 2:2114:11527:96658#2#R | \| \| |
| MQ=60 isize=-207 C=2 | ccccacggccacagtcccag\|c\|ccccccactccggtccttccccaaggatgcagcctcagccttctccacaccacgtttccCCACAGACAAGTTCCCCACATCCTGGACTGGTAGCTGCCCAGGCCAACagnaCTCCAagCCCACAGCGCCT |
| 1:1203:19640:47326#2#F | \| \| |
| MQ=60 isize=177 C=37 | CCTTTCAGACAAATTGGAGTCntCCACGGCCACAGTCCCAG\|C\|CCCCCCACTCCAGTCCTTCCCCAAGGATGCAGCCTCAGCCTTCTCCACACCACGTTTCCCCACAGACAAGTTTCCCaCATcCTGGACTGGTAgCtgCCCAGgCCAACCC |
| 2:2212:5776:56597#2#F | \| \| |
| MQ=60 isize=177 C=40 | AAACGTCGGCGAATTGGAGTCCTTCCACGGCCACAGTCCCAG\|C\|CCCCCCACTCCAGTCCTTCCCCAAGGATGCAGCCTCAGCCTTCTCCACACCACGTTTCCCCaCAGACAAGTTCCCCACATCCTGGACTGGTAGCTGCCCAGGCCAccc |
| 1:1211:11399:22017#2#R | \| \| |
| MQ=60 isize=-208 C=22 | tCCACGGCCACAGTCCCAG\|C\|CCCCCCACTCCAGTCCTTCCCCAAGGATGCAGCCTCagCCTTCTCCACACCaCGTTTCCCCACAGACAAGTTCCCCACATCCTGGACTGGTAGCTGCCCAGGCCAACCAGGACTCCAAtCTGGCGAGCCGG |
| 1:2207:17557:70177#2#R | \| \| |
| MQ=60 isize=-208 C=33 | cCCACGgCccCAgtCCCAG\|C\|CCCCCcaCTCCAGTCCTTCCCCAAGGATgcaGCCTCAGCCTTctCCaCaCCACGTTTCCCCACAGACAAGTTCCCCACATCCTTGACTGGTAGCTGCCCAGGCCAACCanGACTCCAATGCTATCCACCGG |
| 2:1211:17102:19160#2#R | \| \| |
| MQ=60 isize=-208 C=22 | tccaCGGCtaCAGTCCcAg\|C\|CCCCccaCTCCAgtCcttCCCCAAGGatgcagccTCAGCCtTCtCCACaCCaCGTtTCCCCACAGcCAAGTTCCCCACATCCTGgACtGGTaGCTGCCCAGgaCaaCCAGGACTCCAAtCCCCTTACCCCA |
| 1:1206:11196:68965#2#F | \| \| |
| MQ=60 isize=176 C=40 | TGTATTAATACTATTGGAGTCCCACGGCCACAGTCCCAG\|C\|CCCCCCACTCCAGTCCTTCCCCAAGGATGCAGCCTCAGCCTTCTCCACACCACGTTTCCCCACAGACAAGTTCCCCACATCCTGGACTGGTAGCTGCCCaGgcCAACCCCa |
| 1:1201:11894:31574#2#R | \|C\| |
| MQ=60 isize=-208 C=27 | ccacggccaCAGtCCcaI\|C\|CCCCCCACtcCAGTCCTTCCCCAAGGATGcaGcCTCAGCCTTctCCACACCACGTTTCCCCACAGACAAGTtCCCCACATCCTGGACTGGTAGCTGCCCAGGCCAACcanGACTCCaatCCCAGTGGTACG |
| 1:2113:14783:21753#2#R | \| \| |
| MQ=60 isize=-209 C=2 | ccacggccacagtcccag\|c\|ccccccactccagtccttccccaaggatgcagcctcagccttctccacaccacgtttcccCACAGACAAGTTCCCCACATCCTGGACTGGTAGCTGCCCAGGCCAACCCannACTCCAATCGCACCAGACGC |
| 2:1216:11076:68329#2#R | \| \| |
| MQ=60 isize=-192 C=2 | ccacgcctgaccctaccccacggccacagtcccag\|c\|ccccccactccagtccttccccaaggatgcagcctcagccttctccactccacgtttccccacagacaagttccccacatcctgaACTGGTAAgnAcTCCAATACgACCagcGAa |
| 2:2115:14383:30663#2#R | \| \| |
| MQ=60 isize=-209 C=2 | ccacggccacagtcccac\|c\|ccctccactccagtccttcccCAAGGATGCaGCCTCAGCCTTctccacaCCACGTtTCcCCACAGACaAgTTCcccaCAtCCTGGACTGGTAGCTGCCCAGGCCAAcccagnaCTCCAATTCTAGCAGtGCG |
| 1:1211:14575:8764#2#F | \| \| |
| MQ=60 isize=175 C=40 | AATTTGAGCGCCATTGGAGTCCTCACGGCCACAGTCCCAG\|C\|CCCCCCACTCCAGTCCTTCCCCAAGGATGCAGCCTCAGCCTTCTCCACACCACGTTTCCCCACAGACAAGTTCCCCACATCCTgGaCTGGTAGCTGCCCAGGCCAACCCC |
| 1:2111:11679:2038#2#F | \| \| |
| MQ=60 isize=175 C=37 | AATTTGAGCGCCATTGGAGTCCTCACGGCCACAGTCCCAG\|C\|CCCCCCACTCCAGTCCTTCCCCAAGGATGCAGCCTCAGCCTTCTCCACACCACGtTTCCCCACAGACAAGTTCCCCACATCCTGGACTGGTAGCTGCCCAGGCCAACCCC |
| 1:2203:6405:78342#2#F | \| \| |
| MQ=60 isize=175 C=40 | GCCGGTACCTAGATTGGAGTCCTCACGGCCACAGTCCCAG\|C\|CCCCCCACTCCAGTCCTTCCCCAAGGATGCAGCCTCAGCCTTCTCCACACCACGtTTCCCCACAGACAAGTTCCCCACATCCTGGACTGGTAGCTGCCCAGgcCAACCCC |
| 1:2216:17355:49707#2#F | \| \| |
| MQ=60 isize=175 C=40 | AAGGCCGATTGCATTGGAGTncTCACGGCCACAGTCCCAG\|C\|CCCCCCACTCCAGTCCTTCCCCAAGGATGCAGCCTCAGCCTTCTCCACACCacGTTTCCCCACAGaCAAGTTCCCCaCATCCTGGgCTGGtAGctgCCCAGgcCAACCCC |
| 1:2216:4806:64179#2#F | \| \| |
| MQ=60 isize=175 C=37 | TAGTAGACCATGATTGGAGTncTCACGGCCACAGTCCCAG\|C\|CCCCCCACTCCAGTCCTTCCCCAAGGATGCAGCCTCAGCCTTCTCCACACCACGTTTCCCCACAGACAAGtTCCCCACATCCTGGACTGGtAGCTGCCCAGGCCAACCCC |
| 2:1211:2478:74297#2#F | \| \| |
| MQ=60 isize=280 C=37 | GCGTTTGGGGCTATTGGAGTCCTCACGGCCACAGTCCCAG\|C\|CCCCCCACTCCAGTCCTTCCCCAAGGATGCAGCCTCAGCCTTCTCCACACCACGTTTCCCCACAGACAaGTTCCCCACATCCTGGACtGGTAGCTGCCCAGgCCGACCCC |
| 2:2215:18831:13445#2#F | \| \| |
| MQ=60 isize=309 C=27 | GTATCATCTGCCATTGGAGTCCTCACGGCCACAGTCCCAG\|C\|CCCCCCACTCCAGTCCtTCCCCAAGGATGCAGCCTCAGCCTTCTCCaCACCacGTtTCCCCACAGAcAAGTTCCCCACAtCCTGGACTgGTAGCTGCCCAGGCCAACCCC |
| 1:1108:16275:95424#2#R | \| \| |
| MQ=60 isize=-211 C=33 | cacggCCACAGTCCCAG\|C\|CCCCCCACTCCAGTCCTTCCCCAAGGATGCAGCCTCAGCCTTCTCCaCACCACGTTTCCCCACAGACAAGTTCCCCACATCCTGGACTGGTAGCTGCCCAGGCCAACCCCanGACTCCAAtCCCGTTCACTAC |
| 1:2107:11958:63898#2#R | \| \| |
| MQ=60 isize=-211 C=33 | CaCGGCCaCagtCcCAG\|C\|CCCCCCaCTCCAGTCCTTCCCCAAGGATGCAGCCTCAGCCTTCTCCacaCCACGTTTCCCCACAGACAAGTTCCCCACATCCTGGACTGGTAGCTGCCCAGGCCAACCCCAGGACTCCAATCCGATGGTTCAA |
| 2:1109:3518:84498#2#R | \| \| |
| MQ=60 isize=-211 C=2 | cccggccccagtccccg\|c\|CCCCCcactccaGtCCcTCCCCaAGggtgcagCCtCAGcCtTctCcacaccaCGtTTCCCCACAGACAAGTTCCCCACATCCTGGACTGGTAGCTGCCCAGGCCAACCCCanGACTCCAATTGATTTTATGGA |
| 1:1113:2853:59196#2#F | \| \| |
| MQ=60 isize=174 C=37 | GTCCACCAGTTGATTGGAGTCCTACGGCCACAGTCCCAG\|C\|CCCCCCACTCCAGTCCTTCCCCAAGGATGCAGCCTCAGCCTTCTCCACACCACGTTTCCCCACAGACAAGTTCCCCACATCCTGGACtggtAGCTGCCCAGGCCAACCCCa |
| 1:2110:10904:37611#2#F | \| \| |
| MQ=60 isize=174 C=37 | GGAGACGGGCGTATTGGAGTCCTACGGCCACAGTCCCAG\|C\|CCCCCCACTCCAGTCCTTCCCCAAGGATGCAGCCTCAGCCTTCTCCACACCACGTTTCCCCACAGACAAGTTCCCCACATCCTGGACTGGTAGCTGCCCAGGCCAACCCCA |
| 2:2110:21264:39125#2#F | \| \| |
| MQ=60 isize=174 C=33 | ACTATAGTCAACATTGGAGTCCTACGGCCACAGTCCCAG\|C\|CCCCCCACTCCAGTCCTTCCCCAAGGATGCAGcCTCAGCCTTCTCCACACCACGtttccCCaCAGACAAgttCCcCACATCCTGGACTGGTAGCTGCccaGGCCaacccca |
| 1:1206:13124:24637#2#R | \| \| |
| MQ=60 isize=-211 C=27 | cCGGCcACAgtCCCAG\|C\|CCCCCCACtCCAGtCCTTCCCCAAGGATGCAGCCTCAGCCTTCTCCaCACCACGTTTCCCCACAGACAAGTTCCCCACATCCTGGACTGGTAGCTGCCCAGGCCAACCCCAAGGACTCCAATACACGCCAACAC |
| 1:1105:15580:76952#2#F | \| \| |
| MQ=60 isize=307 C=2 | GCGGGGGCCCggggtgggggcncggccacaggcccag\|c\|cccccccctcccgtccttccccaaagatgcagccccagccttctccccaccacgtttccccccagacaagttccccccatcctggattgggagctgcccccgcccaccccatg |
| 1:1111:10114:59039#2#F | \| \| |
| MQ=60 isize=173 C=40 | TATTATTGTTACATTGGAGTCCTCGGCCACAGTCCCAG\|C\|CCCCCCACTCCAGTCCTTCCCCAAGGATGCAGCCTCAGCCTTCTCCACACCACGTTTCCCCACAgaCAAGTTCCCCACATCCTGGACTGGTaGCTGCCCAGGCCaaCCCCAT |
| 1:1113:13546:45344#2#F | \| \| |
| MQ=60 isize=173 C=27 | GTACCGTTTCAAATTGGAGTCCCGGCCACAGTCCCAG\|C\|CCCCCCACTCCAGTCCTTCCCCAAGGATGCAGCCTCAGCCTTCTCCACACCACGTTTCCCCACAGACAAGTTCCCCACATCCTGgACTGGTAGCTGCCCAGGCCAACCCCATG |
| 1:2208:8949:92528#2#F | \| \| |
| MQ=60 isize=173 C=40 | GGTTACCGCTAAATTGGAGTCntCGGCCACAGTCCCAG\|C\|CCCCCCACTCCAGTCCTTCCCCAAGGATGCAGCCTCAGCCTTCTCCACACCACGTTTCCCCAcAGACAAGTTCCCCACATCCTGGACTGGTAGCTGCCCAGGCCAACCCCAt |
| 2:2104:7169:8085#2#F | \| \| |
| MQ=60 isize=173 C=40 | GTACCGTTTCAAATTGGAGtCCCGGCCACAGTCCCAG\|C\|CCCCCCACTCCAGTCCTTCCCCAAGGATGCAGCCTCAGCCTTctccaccccacgTTtCCCCacAGcaaaGTtCCCcccatcctggcctggaacctcccca\_gccaaacccatgg |
| 1:2201:1438:21125#2#R | \| \| |
| MQ=60 isize=-212 C=2 | cggccacagtcccag\|c\|ccccccgctccagtccttccccaaggatgcagcctcagccctctccgcaccgcgttaccccacagacaagttccccgcaccctgcactggtagCTGCCCAGGCCAaCCCCATAGGACTCCAAttCAgATCCCCga |
| 2:1212:7745:93565#2#R | \| \| |
| MQ=60 isize=-212 C=37 | CGGCCACAgTCCCaG\|C\|CCCCCCaCTCCAGTCCTTCCCCAAGGATGCaGCCTCAGCCTTCTCCACaCCACGTTTCCCCACAGACAAGTTCCCCACATCCTGGACTGGTAGCTGCCCAGGCCAACCCCATAGGACTCCAATAAGCAGGGACTG |
| 2:2115:4265:2103#2#R | \| \| |
| MQ=60 isize=-212 C=33 | cggcccccgtcccag\|C\|CCCCccactccagtccttcCCCAAgGaTgCagCctCAGCCTTCTCCacaCCaCgTTTCCCCACAGACAAGTTCCCCaCATCCTGGACTGGTAGCTGCCCAGGCCAACCCCATAgnACTCCAAtCCCTTTGCCCCA |
| 1:1211:6038:72056#2#F | \| \| |
| MQ=60 isize=306 C=40 | AGCTTGAGTTGCATTGGAGTCCTGGCCACAGTCCCAG\|C\|CCCCCCACTCCAGTCCTTCCCCAAGGATGCAGCCTCAGCCTTCTCCACACCACGtTTCCCCACAGACAAGTTCCCCACATCCTGGACtgGTAGCTGCCCAgGCCAACCCCATG |
| 1:1214:16532:55501#2#F | \| \| |
| MQ=60 isize=172 C=37 | GGCGTGGGGGAGATTGGAGTncTGGCCACAGTCCCAG\|C\|CCCCCCaCTCCAGTCCTTCCCCAAGGaTGCAGCCTCagcCTTCTCCAcaCCACGtTTCCCCACaGACAAgTtCCCCACATCCTGGACTAGtAGCtGCCCAgtccAACCCCATG |
| 2:1111:12895:40343#2#F | \| \| |
| MQ=60 isize=172 C=37 | GCGACTTTGAAAATTGGAGTCCTGGCCACAGTCCCAG\|C\|CCCCCCACTCCAGTCCTTCCCCAAGGATGCAGCCTCAGCCTTCTCCACACCaCGTTTCCCCACAGACAAGTTCCCCACATCCTGGACTGGTagCtGCCCAGGCCAACCCCATg |
| 2:2112:1909:100597#2#F | \| \| |
| MQ=60 isize=306 C=37 | GACATTGGTCGAATTGGAGTncTGGCCACAGTCCCAG\|C\|CCCCCCACTCCAGTCCTTCCCCAAGGATGCAGCCTCAGCCTtCtCCacaCcccGTTtCCCCacaGacaagttCCCCccatcctggacgggtagctccccgggccacccccaga |
| 2:2214:2758:45843#2#F | \| \| |
| MQ=60 isize=172 C=40 | GCGCGTCGGCCCATTGGAGTCCTGGCCACAGTCCCAG\|C\|CCCCCCACTCCAGTCCTTCCCCAAGGATGCAGCCTCAGCCTTCTCCACACCACGTTTCCCCACAGACAAGTTCCCCacATCCTGGaCTGGTAGCTGCCCAGgCCAACCCCATG |
| 1:1108:16828:35161#2#F | \| \| |
| MQ=60 isize=171 C=27 | TAAATACTTCAAatTGGAGtcntgCCACAGTCCCAG\|C\|CCCCCCACTCCAGTCCTTCCCCAAGgATGCAGCCTCAGCCTTCTCCACACCaCGTtTCCCCACAGACAAGTTCCCCACATCCTggaCtGgtagCTGCCCaggccaaccccatgg |
| 1:1209:12181:37048#2#F | \| \| |
| MQ=60 isize=276 C=37 | CCGCATCGCACCATTGGAGTCntGCCACAGTCCCAG\|C\|CCCCCCACTCCAGTCCTcCCCCAAGGATGCAGCCTCAGCCTTCTCCACACCACGTTTCCCCaCAGACAAGTTCCCCACATCCTGGACTgGtAGCTGCCCAGGCCAACCCCATGg |
| 1:2208:17071:53410#2#F | \| \| |
| MQ=60 isize=171 C=37 | AGGCAATTTAGAATTGGAGTCntGCCACAGTCCCAG\|C\|CCCCCCACTCCAGTCCTTCCCCAAGGATGCAGCCTCAGCCTTCTCCACACCACGTTTCCCCaCaGACAAGTTCCCCACATCCTGGACTGGtAgCTGCCCaggcCAACCCCATGg |
| 2:1106:7985:51260#2#F | \| \| |
| MQ=60 isize=171 C=40 | TCCGCCAAAGCGATTGGAGTCntGCCACAGTCCCAG\|C\|CCCCCCACTCCAGTCCTtCCCCAAGGATGCAGCCtCAgCCTTctccaCACcaCGTTtCCCCccagacaagttccccccatcctggcctgtcacctgcccaggccaaccccatca |
| 2:2114:9483:48128#2#F | \| \| |
| MQ=60 isize=276 C=37 | GCAGCTCGTGCCATTGGAGTncTGCCACAGTCCCAG\|C\|CCCCCCACTCCAGTCCTTCCCCAAGGATGCAGCCTCAGCCTTCTCCaCACCACGTTTCCCCATAGACAAGtTCCCCACATCCTGGACTGGTAGctGcCCAggCcaACCCCAtgg |
| 2:2216:14593:5396#2#F | \| \| |
| MQ=60 isize=305 C=37 | TAAATACTTCAAATTGGAGTCCTGCCACAGTCCCAG\|C\|CCCCCCACTCCAGTCCTTCCCCAAGGATgCAGCCTCAGCCTTCTCCACACCACGTTTCCCCACAGACAAGTTCCCCACATCCTgGACTGGtagCTGCCCAGGCCaACCCCATGG |
| 2:1109:12888:33611#2#R | \| \| |
| MQ=60 isize=-215 C=33 | GCcACAgtCCCAG\|C\|CCCCCCACtCCAGTCCTTCCCCAAGGATGCAGCCTCAGCCTTCTCCACaCCACGTTTCCCCACAGACAAGTTCCCCACATCCTGGACTGGTAGCTGCCCAGGCCAACCCCATGGanGACTCCAATTCCGACCCTCGC |
| 1:1113:20652:92677#2#F | \| \| |
| MQ=60 isize=170 C=40 | TCGAATGACTATATTGGAGTCCTCCACAGTCCCAA\|C\|CCCCCCACTCCAGTCCTTCCCCAAGGATGCAGCCTCAGCCTTCTCCACACCACGTTTCCCCACAGACAAGTTCCCCACATCCTGGACTGGTAGCTGCCCAGGCCAACCCCATGGa |
| 1:2104:14856:45030#2#F | \| \| |
| MQ=60 isize=170 C=40 | GTGTACAGGCTGATTGGAGTCntCCACAGTCCCAG\|C\|CCCCCCACTCCAGTCCTTCCCCAAGGATGCAGCCTCAGCCTTCTCCACACCACGTTTCCCCACAGACAAGTTCCCCACATCCtGGACTGGTAGCTGCCCAGGCCAACCCCATGGa |
| 2:1103:18538:31189#2#F | \| \| |
| MQ=60 isize=170 C=37 | GGTGGAGGCCAAATTGGAGTCCTCCACAGTCCCAG\|C\|CCCCCCACTCCAGTCCTTCCCCAAGGATGCAGCCTCAGCCTTCTCCACACCACGTTTCCCCACAGACAAGTTCCCCACATCCTGGACTGGTAGCTGCCCAGGcCAAcCCCAtgGa |
| 2:2205:3959:90754#2#F | \| \| |
| MQ=60 isize=170 C=40 | GGGGGAGAGTAGATTGGAGTCntCCACAGTCCCAG\|C\|CCCCCCACTCCAGTCCTTCCCCAAGGATGCAGCCTCAGCCTTCTCgaCACCACGTTTCCCCACAGACAAGTTCCCCACATCCTGGACTGGTAGCTGCCCAGgCCAACCCCATGGA |
| 2:1109:10666:44299#2#R | \| \| |
| MQ=60 isize=-208 C=2 | ccccgcgtccacagtcccag\|c\|ccccccactCCAGTCCTTCcccAagGatgCAGCCtgAGcCTTCtCCACACCaCGTttCCCCACAGACAAGTTcCCcaCATCCTGGACTGGTAGCTGCCCAGGCCAACcnGGACTCCAATCCCCAGGAGCG |
| 1:1114:18318:71911#2#F | \| \| |
| MQ=60 isize=169 C=33 | AGACGGAACGGAATTGGAGTCCtCACAGTCCCAG\|C\|CCCCCCACTCCAGtCCTTCCCCAAGGATGCAGCCTCAGCCTTCTCCaCACCACGTTTCCCCACAGACaAGTtCCCCACATCCTGGACTGGtagCTGCCCAGgCCAACCCCATGgaa |
| 1:2112:16240:12007#2#F | \| \| |
| MQ=60 isize=169 C=37 | GTGCGGGGTTGCATTGGAGTCCTCACAGTCCCAG\|C\|CCCCCCACTCCAGTCCTTCCCCAAGGATGCAGCCTCAGCCTTCTCCACACCACGTTTCCCCACAGACAAGTTCCCCACATCCTGGACTGGTAGCTGCCCAGGCCaaCCCCATGGAA |
| 2:1110:10143:52224#2#F | \| \| |
| MQ=60 isize=274 C=33 | TTAAGCTAGCAAATTGGAGTCCTCACAGTCCCAG\|C\|CCCCCCACTCCAGTCCTTCCCCAAGGATGCAGCCTcaGCCTTCTCCACACCACGTTTCCCCACAGACAAGTTCCCCACAtCCTGGACTGGTagcTGCCcAGgCCAACCCCATGGAa |
| 2:1215:10508:12473#2#F | \| \| |
| MQ=60 isize=169 C=37 | GCCGGACGGTGAATTGGAGTncTCACAGTCCCAG\|C\|CCCCCCACTCCAGTCCTTCCCCAAGGATGCAGCCTCAGCCTTCTCCACACCACGTTTCCCCACAGACAAGTTCCCCACATCCTGGACTGGTAGCTGCCCAGGCCaACCCCATGGAA |
| 2:2203:2659:23302#2#F | \| \| |
| MQ=60 isize=169 C=37 | GGGATAGCACCGATTGGAGTCntCACAGTCCCAG\|C\|CCCCCCACTCCAGTCCTTCCCCAAGGATGCAGCCTCAGCCTTCTCCACGCCaCGTTTCCCCaCAgacAAGTTCCCCaCatCCTGgaCgggtaGCTGCCCaggCCAaCCCCATGgaa |
| 2:2102:5645:4052#2#R | \| \| |
| MQ=60 isize=-179 C=2 | cccacgtgcgctccccccgcccggcccctccccccgggcacagtcccag\|c\|ccccccaccccagtcCctCCCCAAGgatgCcGCCtcAGcCTTCtCCacaCCaCGTTTCCCCACAGACAAGTTCCCCACAGGACTCCAATGGTTACACCCCT |
| 1:1111:17398:73213#2#R | \| \| |
| MQ=60 isize=-218 C=37 | aCAGTCCCAG\|C\|CCCCCCACTCCAGTCCTtCCCCAAGGATGCAGCCTCAGCCTTCTCCACACCACGTTTCCCCACAGACAAGTTCCCCACATCCTGGACTGGTAGCTGCCCAGGCCAACCCCATGGAACAGGACTCCAATATCGGGCAAAAC |
| 2:1104:19742:49071#2#R | \| \| |
| MQ=60 isize=-218 C=22 | ccagtCCCAG\|C\|CCccCcACtCCAGTCCTtCCccaaGGATGCAGCCTCAGcCTtcTCCAcaCCaCGTTtCCCCACAGACAAGTTCCCCACATCCTGGACTGGaAGCTGCCCAGGCCAACCCCATGGAACAGGACTCCAATCCCCGTGCCACT |
| 1:1108:10562:52948#2#F | \| \| |
| MQ=60 isize=167 C=40 | AATGCCGATTCCATTGGAGTCntCAGTCCCAG\|C\|CCCCCCACTCCAGTCCTTCCCCAAGGATGCAGCCTCAGCCTTCTCCACACCACGTTTCCCCACAGACAAGTTCCCCACATCCTGGACTGGTAGCTGCCCAGgCCAACCCCATGGAACA |
| 2:1109:15464:68347#2#F | \| \| |
| MQ=60 isize=167 C=40 | CGGGGATGGGGGATTGGAGTCntCAGTCCCAG\|C\|CCCCCCACTCCAGTCCTTCCCCAAGGATGCAGCCTCAGCCTTCTCCACACCACGtTTCCCCACAGACAAGtTCCCCaCATCCTGGACTGGtAGCTGCCCAGGCCAACCCCATGGaaca |
| 2:2110:2774:42642#2#F | \| \| |
| MQ=60 isize=301 C=37 | ACCATGGGACGCATTGGAGTCCTCAGTCCCAG\|C\|CCCCCCACTCCAgTCCTTCCCCAAGGATgcaccctcagccttctccacaccacttttccccccagacaatttccccccaccctggcgtggtagcgccccgggcaaccccccggaaaca |
| 2:2206:19779:15055#2#R | \| \| |
| MQ=60 isize=-209 C=2 | ccaggcccccagtcccag\|c\|cccccccctccactcctcccccaaggatgcaccctcagccttctccaccccacgttcccccccagacaagttccccacatccgggcctggtagctgcccaggccaacccgngccccaaacctccccccCCGC |
| 1:1212:4749:62749#2#F | \| \| |
| MQ=60 isize=166 C=40 | AACCCCATGATAATTGGAGTCCTAG\|C\|CCCCCCACTCCAGTCCTTCCCCAAGGATGCAGCCTCAGCCTTCTCCACACCACGTTTCCCCACAGACAAGTTCCCCACATCCTGGACTGGTAGCTGCCCAGGCCAACCCCATGGAACAAGGGCAT |
| 1:2103:3831:86763#2#F | \| \| |
| MQ=60 isize=166 C=40 | GGAGGAGAGAGCATTGGAGTCCTAG\|C\|CCCCCCACTCCAGTCCTTCCCCAAGGATGCAGCCTCAGCCTTCTCCACACCACGTTTCCCCACAGACAAGTTCCCCACATCCTGGACTGGTAGCTGCCCaGGCCAACCCCATGGAACAAggGCAT |
| 2:2104:15460:23477#2#F | \| \| |
| MQ=60 isize=166 C=37 | TCGTCGAGCCGAATTGGAGTCCTAG\|C\|CCCCCCACTCCAGTCCTTCCCCAAGGATGCAGCCTCAGCCTTCTCCACACCACGttTCCCCACAGacAAgtTCCCcacAtCCtgGcctgGtAgcggCCCAGGCCAAcCCCAtGGAacaagggcat |
| 2:2212:14988:74500#2#F | \| \| |
| MQ=60 isize=166 C=37 | ACGATACGAACGATTGGAGTCCTAG\|C\|CCCCCCACTCCAGTCCTTCCCCAAGGATGCAGCCTCAGCCTTCTCCACACCACGTTTCCCCACAGACAAGTTCCCCACATCCTGGACTGGTAGCTGCCCAGGCCAACCCCATGGAACAAGGGCAT |
| 1:2102:18123:64008#2#R | \| \| |
| MQ=60 isize=-219 C=27 | agtCCCaG\|C\|CCCCCcaCTCCAGTCCTTCCCCAAGGATGCAGCCTCAGCCTTctCCACaCCACGTTTCCCCACAGACAAGTtCCCCACATCCTGGACTGGTAGCTGCCCAGGCcAaCCCCATGGAACAAAGGACTCCAATCCAGCCCCCCCG |
| 1:1111:7969:41102#2#F | \| \| |
| MQ=60 isize=270 C=37 | TTACGAGATTTTATTGGAGTCCTGTCCCAG\|C\|CCCCCCACTCCAGTCCTTCCCCAAGGaTGCAGCCTCAGCCTTCTCCACACCACGTTTCCCCACAGACAAGTTCCCCACATCCTGGACTGGtGgCTGCCCAcgcCAACCCCATGGaACaag |
| 1:1115:5774:82894#2#F | \| \| |
| MQ=60 isize=165 C=37 | TTACGAGATTTTATTGGAGTCCTGTCCCAG\|C\|CCCCCCACTCCAGTCCTTCCCCAAGGATGCAGCCTCAGCCTTCTCCACACCACGTTTCCCCACAGACAAGTTCCCCACATCCTGGACTGGTAGCtgCCCAGGCCAACCCCATggAaCAAG |
| 1:1216:15197:57606#2#F | \| \| |
| MQ=60 isize=165 C=40 | TGAGTTGCAATAATTGGAGTncTGTCCCAG\|C\|CCCCCCACTCCAGTCCTTCCCCAAGGATGCAGCCTCAGCCTTCTCCACACCACGTTTCCCCACAGACAAGTTCCCCACATCCTGGACTGGTAGCTGCCCAGgCCAACCCCATGGAACAAg |
| 1:2202:12267:78045#2#F | \| \| |
| MQ=60 isize=165 C=40 | ACGCAGCATTGTATTGGAGTCntGTCCCAG\|C\|CCCCCCACTCCAGTCCTTCCCCAAGGATGCAGCCTCAGCCTTCTCCACACCACGTTTCCCCACAGACAAGTTCCCCACATCCTGGaCTGGtAGCTGCCCAGgccAACCCCaTGGAacaAg |
| 2:1112:7014:92729#2#F | \| \| |
| MQ=60 isize=165 C=33 | GGTACGTGAGTGATTGGAGTCCTGTCCCAg\|C\|CCCCCCACTCCAGTCCTTCCCCAAGGATGCAGCCTCAgCcttCTCCCCACCACGTTtccCCACAgACAAgttCCCCaCATCCTggaCTGgtagcggcCCAGgCCAACCCCAtGGAacaag |
| 2:1214:15570:96036#2#F | \| \| |
| MQ=60 isize=165 C=37 | ACGCTTGCATGTATTGGAGTCCTGTCCCAG\|C\|CCCCCCACTCCAGTCCTTCCCCAAGGATGCAGCCTCAGCCTTCTCCACACCACGTTTCCCCACAGACAAGTTCCCCACATCCTGGACTGGTAGCTGCCCAGGCCAACCCCATGGAACAAG |
| 1:1205:14932:49229#2#R | \| \| |
| MQ=60 isize=-220 C=33 | gcCCCag\|C\|CCCCCcactcCAGTCCTTCCCCAAGGATGCaGCCTCaGCCTTCTCCaCaCCACGTTTCCCCACAGACAAGTTCCCCACATCCTGGACTGGTAGCTGCCCAGGCCAACCCCATGGAACAAGanGACTCCAATACAAGCTCCACG |
| 1:2115:15013:73320#2#R | \| \| |
| MQ=60 isize=-220 C=37 | GTCCCaG\|C\|CCCCCCaCTCCAGTCCTTCCCCAAGGATGCAGCCTCAGCCTTCTCCACACCACGTTTCCCCACAGACAAGTTCCCCaCATCCTGGACTGGTAGCTGCCCAGGCCAACCCCATGGAACAAGAGGACTCCAATACAAGCTCCACG |
| 2:2111:3220:27929#2#R | \| \| |
| MQ=60 isize=-187 C=2 | cccccccccggccggcccctctcccccgcccccgtcccag\|c\|ccccccaccccaggccctcccccaggaagcaggctcagccttctcccccccaCGTTTCCCCaCAGAcAAGTTCCCCACATCCTGgACAGGACTCCAATCAAAAAATGcAG |
| 1:1214:8181:80070#2#F | \| \| |
| MQ=60 isize=164 C=37 | GGGAGTGTAACCATTGGAGTncTCCCAG\|C\|CCCCCCACTCCAGTCCTTCCCCAAGGATGCAGCCTCAGCCTTCTCCACACCACGTTTCCCCACAGACAAGtTCCCCACATCCTGGACTGGTAGCTGCCCAGGCCAACCCCATGGAACAAGGG |
| 1:2213:8603:91238#2#F | \| \| |
| MQ=60 isize=164 C=40 | AGTGTGGCAGGCATTGGAGTncTCCCAG\|C\|CCCCTCACTCCAGTCCTTCCCCAAGGATGCAGCCTCAGCCTTCTCCACACCACGTTTCCCCACAGACAAGTTCCCCACATCCTGGACTGGTAGCTGCCCAGGCCAACCCCATGGaaCAAGGg |
| 2:1113:14443:39823#2#F | \| \| |
| MQ=60 isize=164 C=37 | CTGCTGTTGCACACTGGAGTncTCCCAG\|C\|CCCCCCACTCCAGTCCTaCCCCaAGGATGCAGCCTCAGCCTTCTCCACaCCaCGTTTCCCCACAGaCAAGTTCCCCaCATCCTGGACTGGTaGCTGCCCaGGCCaacCCCatGgAacaaggg |
| 2:1216:10374:39901#2#F | \| \| |
| MQ=60 isize=367 C=33 | GGAGAATGCGCCATTGGAGTncTCCCAG\|C\|CCCCCCACTCCAGTCCTTCCCCAAGGATGCAGcCTCAGCCTTCTCCACACCACGttTCCCCACAGACAAGTTCCCCACATCCTGGaCTgGtAGCTGCCCagGCCaaCccCATGGAACAAGGG |
| 2:2104:5140:7521#2#F | \| \| |
| MQ=60 isize=164 C=37 | ATTATACAGGCTATTGGAGTCCTCCCAG\|C\|CCCCCCACTCCAGtCCTTCCCCAAGGATGCAGCCTCAgCCTtcccccccccccgtttccccccagcaaagtccccccactccgggccggtagcccgcccagcccacccccctgaaacaaggc |
| 1:1113:9491:76232#2#R | \| \| |
| MQ=60 isize=-221 C=27 | tCCCAG\|C\|CCCCCCaCTCCAGTCCtTCCCCAAGGATGCAGCCTCAGCCTTCTCCACaCCACGTTTCCCCACAGACAAGTTCCCCACATCCTGGACTGGTAGCTGCCCAGGCCAACCCCATGGAACAAGGAGGACTCCAATTTAGCGTATCCC |
| 1:2113:12936:4942#2#R | \| \| |
| MQ=60 isize=-221 C=27 | tCCcaG\|C\|CCCCCCACTCCAGTCcTTCCCCAAGGATGCAGCCTCAGCCTtctCCACaCcaCGTTTCCCCACAGACAAGTTCCCCACATCCTGGACTGGTAGCTGCCCAGGCCAACCCCATGGAACAAGgannACTCCAATCCTTCCGGGGTT |
| 1:2213:12211:38407#2#R | \| \| |
| MQ=60 isize=-221 C=2 | tcccac\|c\|ccccccacTCCAGTCCTTcCCCAAGGATGCAGcCTCAGCCtTcTCCACACCACGtTtCCCCACAGACAAGtTcCCCACAtCCTGGACTGGTAGCTGCCCAGGCCAACCCCATGGAACAAGGAgnACTCCAATCCTTCCggGGTT |
| 2:2108:3256:29144#2#R | \| \| |
| MQ=60 isize=-181 C=2 | cctgcgcccgccccaacctggtccttttcccccgcccccatcccag\|c\|cccccccctccagtccttccccaaagatgcagcctcagccttctccaccccacgttTCCCCACAGaCaAGttcCcCACAtCAGGACTCCAATCGGGTGcCCGCc |
| 1:1103:11093:11985#2#R | \| \| |
| MQ=40 isize=-139 C=2 | actcccccacggccggcccatccctactcctctcccccatcccctgcccccccaccagcctctccccgcgccaaggccaccgccccag\|c\|cccccccccccagtacttccccaaggactcagcctcagcgtgggcgtggaacccacaggaga |
| 1:2211:6706:82014#2#R | \| \| |
| MQ=60 isize=-222 C=37 | CCCAG\|C\|CCCCCCACTCCAGTCCTTCCCCAAGGATGCAGCCTCAGCCTTCTCCaCACCACGTTTCCCCACAGACAAGTTCCCCACATCCTGGACTGGTAGCTGCCCAGGCCAACCCCATGGAACAAGGGAGGACTCCAATAGTGTTGAACCA |
| 2:1109:7180:97624#2#R | \| \| |
| MQ=60 isize=-207 C=2 | ctccccggccccacgcccag\|c\|cccccccctccagtccttcCCCAAGGaTgCAGCCTCAGCCTTCTCCACACCaCGTTTCCCCACAGACAAGTTCCCCACATCCTGGACTGGTAGCTGCCCAGGCCAACanGACTCCAAtTGCGCACGGCAC |
| 1:1102:14430:94301#2#F | \| \| |
| MQ=60 isize=162 C=40 | GCATCTTTCCTCATTGGAGTCntCCAG\|C\|CCCCCCACTCCAGTCCTTCCCCAAGGATGCAGCCTCAGCCTTCTCCACACCACGtTTCCCCACAGACAAGTTCCCCACATCCTGGACTGGTAGCTGCCCAGGCCAaCCCCATGGAACAAGGGC |
| 1:2203:3650:81034#2#F | \| \| |
| MQ=60 isize=161 C=37 | CGTTCGGGGGTTATTGGAGTCCTCCAG\|C\|CCCCCCACTCCAGTCCTTCCCCAAGGATGCAGCCTCAGCCTTCTCCACACCACGTTTCCCCACAGACAAGTTCCCCACATCCTGGACTGGTAGCTGCCCAGGCCAACCCCATGGaACAAggGC |
| 1:1205:20042:93382#2#R | \| \| |
| MQ=60 isize=-224 C=2 | ccag\|c\|ccccccactccAGTCCttCCCcAAgGATGCAGCCtCAgCcttctCCacaCccCGTttCCCCACAGACAAGTTCCCCACATCCTGGACTGGTAGCTGCCCAGGCCAaCcCCATGGAACAAGGGCanGACTCCAATAATAGTAATCCA |
| 1:2214:2264:68699#2#F | \| \| |
| MQ=60 isize=266 C=40 | TACATCACAATAATTGGAGTncTCAG\|C\|CCCCCCACTCCAGTCCTTCCCCAAGGATGCAGCCTCAGCCTTCTCCACACCACGtTTCCCCACAgACAAGTTCCCCACATCCTGGACTGGTAGCTGCCCAgGCCAACCCCATgGAACAAgGGCa |
| 2:2116:7886:34206#2#F | \| \| |
| MQ=60 isize=161 C=37 | GCGACAACGATTATTGGAGTncTCAG\|C\|CCCCCCACTCCAGTCCTTCCCCAAGGATGCAGCCTCAGCCTCCTCCACACCACGTTTCCCCACAGACAAGTTCCCCACATCCTGGaCTGGTAGCTGCCCAGGCCAACCCcaTGgAAcaAGGgct |
| 1:1204:16380:11870#2#F | \| \| |
| MQ=60 isize=160 C=37 | GTCTACGGGAAGATTGGAGTCntAG\|C\|CCCCCCACTCCAGTCCTTCCCCAAGGATGCAGCCTCAGCCTTCTCCACACCACGTTTCCCCACAGACAAGTTCCCCACATCCTGGACTGGTAGCTGCCCAGGCCAACCCCATGGAAcAAgGGCAT |
| 1:1213:5708:10633#2#F | \| \| |
| MQ=60 isize=160 C=33 | GTCTACGGGAAGATTGGAGTncTAG\|C\|CCCCCCACTCCAGTCCTTCCCCAAGGATGCAGCCTCAGCCTTCTCCACACCACGtTTCCCCACAGcCAAGTTCCCcaCATCCTGGACTgGTAGCTgcCCAGgCCAACcCCatggaacaagggcat |
| 1:1202:20780:14654#2#R | \| \| |
| MQ=60 isize=-220 C=33 | cG\|C\|CCCCCCaCTCCAGTCCTTCCCCAAGGATGCAGCCTCAGCCCTCTCCACACCACGTTTCCCCACAGACAAGTTCCCCACATCCTGGACTGGTAGCTGCCCAGGCCAACCCCATGGAACAAGAGCATAGGACTCCAATTAGTCGGCGCCA |
| 1:2111:3217:44256#2#R | \| \| |
| MQ=60 isize=-176 C=37 | AG\|C\|CCCCCCCCTCCAGTCCtTCCCCAAGGATGCAGCCTCAGCCTTCTCCACACCACGTTTCCCCaCAGACAAGTtCCCCCCATATTAGGACACCTATGATCTCTATTTTACACATGAGGAAATTGAAGaGGACTCCAATATACCTAACCTC |
| 1:1112:13895:68041#2#F | \| \| |
| MQ=60 isize=293 C=33 | TCTAGTTTATGCATTGGAGTncTG\|C\|CCCCCCACTCCAGTCCTTCCCCAAGGAtGCAGCCTCAGCCTTCTCCACACCACGTTTCCCCACAGACAAGTTCCCCACATCCTGGACTGGtAGCTGCCCAGGCCAACcCCAtGGAACAAGgGCaTT |
| 1:2203:17767:70623#2#F | \| \| |
| MQ=60 isize=293 C=40 | TGCCGAAAGCGCATTGGAGTCCTG\|C\|CCCCCCACTCCAGTCCTTCCCCAAGGATGCAGCCTCACCCTtCTCCACACCACGTTTCCCCACAGaCAAGTTCCCCACATCCTGGACTGGTAGCtGCCCAGGCCAACCCCctgGAACaAGggCaTT |
| 2:1209:19991:64473#2#F | \| \| |
| MQ=60 isize=395 C=33 | TAATCGGGCTTAATTGGAGTCCTG\|C\|CCCCCCACTCCAGTCCTTCCCCAAgGATGCAGCCtCAGCCTTCTCCACACCACgttTCCCCACAGACAaGTTCCCCACATCCTGgACTGGTAGCTGCCCAggCCAACCCCATggAACAAgggcatt |
| 2:1107:8031:71310#2#R | \| \| |
| MQ=60 isize=-186 C=2 | gccccccccccaaccgttcccttctcccaggccccaggccccg\|c\|ccccccgccccagtccttccccaaagatgcaacctcagccttctccacaccacgtttccccacagacaagttccccacatcctggngactccagcccccccctGTGC |
| 2:2106:4737:38612#2#R | \| \| |
| MQ=60 isize=-175 C=2 | caatcccagtcccccccccccggcctgcccttccccccgggccccggccccg\|c\|ccccccacccccggcCttCCCCAAGGATGCAGCCTCAGCCTTCTCCaCACCaCGTTTCCCCACAGACAAGTTCCCanGACTCCAATAGACAATCTCAT |
| 2:2206:6054:73469#2#R | \| \| |
| MQ=60 isize=-226 C=37 | c\|C\|CCCCccaCtCCaGTCCTTCCCCAAGGATGCAGCCTCAGCCTTCTCCAcaCCACGTTTCCCCACAGACAAGTTCCCCACATCCTGGACTGGTAGCTGCCCAGGCCAACCCCATGGAACAAGGGCATtanGACTCCAATTCTCTGCCAGCC |
| 2:1210:20868:42207#2#F | \| \| |
| MQ=60 isize=158 C=33 | GTAcTCGTGCGCATTGGAGTCCT\|C\|CCCCCCACTCCAGTCCTTCCCCAAGGATGCAGCCTCAGCCTTCTCCACACCACGTTTCCCCACAGACAAGTTCCCCACATCCTGGACTGGTaGCTGCCCAGGCCaACCCCATGGAACAAGGGCATTT |
| 1:2216:11366:71888#2#R | \| \| |
| MQ=60 isize=-229 C=27 | \|C\|CCCCCcaCTCCAGtCCTTCCCCAAGGATGCAGCCTCAGCCTTCtCCACACCACGTTTCCCCACAGACAAGTtCCCCACATCCTGGACTGGTAGCTGCCCAGGCCAACCCCATGGAACAAGGGCATTTtgnACTCCAATTACTCAACGTA |

| 22:41574679 | all:2838 A:0 a:1 T:1 t:4 C:2557 c:263 G:1 g:4 Ins:0 ins:0 del:7 n:0 match:2827 mismatch:18 softclip:9 |
| --- | --- |
| Normal\_Reference | CCTGTTCAGCCCAACCCCATGAGCCCCCAGCAGCATATGCTCCCAAATCAGGCCCAGTCCCCACACCTACAAGGCCAGCAGATCCCTAATTCTCTCTCCAATCAAGTGCGCTCTCCCCAGCCTGTCCCTTCTCCACGGCCACAGTCCCAG\|C\|CCCCCCACTCCAGTCCTTCCCCAAGGATGCAGCCTCAGCCTTCTCCACACCACGTTTCCCCACAGACAAGTTCCCCACATCCTGGACTGGTAGCTGCCCAGGCCAACCCCATGGAACAAGGGCATTTTGCCAGCCCGGACCAGAATTCAA |
| 1:2215:14945:33270#2#F | \| \| |
| MQ=60 isize=262 | GGAAAAGTAGGAATTGGAGTncTaTCAGGCCCAGTCCCCACAccTACAAGGCCAGCAGATCcCTAATTCTCTCTCCAATCAAGtgcgctcTCcCCAgcCTGTCCCTTCTCCACggCCaCAGTCccag\|\_\|cccCCCACTCCagtcCTTCccCAa |
| 1:1211:11907:36720#2#R | \| \| |
| MQ=60 isize=-125 | TTTgGCCcagtCCCCACcCCTACAAgGCcAGCAGATCCCTAATTCTcTCTCCAATCAAGTGCgCtCTcCCCAGCCTgTCCCtTCTCCACGGCCACAGTCCCAG\|\_\|CCCCCCACTCCAGTCCTTCCCCAAGAGGACTCCAATCAAAACACAAAC |
| 1:1105:10480:58712#1#F | \| \| |
| MQ=60 isize=149 | GCCCAGTCCCCACACCTACAAGGCCAGCAGATCCCTAATTCTCTCTCCAATCAAGTGCGCTCTCCCCAGCCTGTCCCTTCTCCACGGCCACAGTCCCAG\|\_\|CCCCCCACTCCAGTCCTTCcCCAAGGATGCAGCCTCAGCCTTCTCCAcAGga |
| 1:1211:11907:36720#1#F | \| \| |
| MQ=60 isize=125 | GCCCAGTCCCCACACCTACAAGGCCAGCAGATCCCTAATTCTCTCTCCAATCAAGTGCGCTCTCCCCAGCCTGTCCCTTCTCCACGGCCACAGTCCCAG\|\_\|CCCCCCACTCCAGTCCTTCCCCAAGAGGACTCCAATCAAAACACAAACCTGt |
| 1:2109:2810:35508#1#F | \| \| |
| MQ=60 isize=177 | GCCCAGTCCCCACACCTACAAGGCCAGCAGATCCCTAATTCTCTCTCCAATCAAGTGCGCTCTCCCCAGCCTGTCCCTTCTCCACGGCCACAGTCCCAg\|\_\|CCCCCCactccaatccttccccaaagatgcagactcagccttctccacacca |
| 1:2208:2757:64975#1#F | \| \| |
| MQ=60 isize=179 G=33 | GCCCAGTCCCCACACCTACAAGGCCAGCAGATCCCTAATTCTCTCTCCAATCAAGTGCGCTCTCCCCAGCCTGTCCCTTCTCCACGGCCACAGTCCCAg\|G\|CCCCCCACTCCAGTCCTTCCCCAAGGATGCAGCCTCAGCCTTCTCCACACC |
| 2:1207:14028:55477#1#F | \| \| |
| MQ=60 isize=131 A=2 | GCCCAGTCCCCACACCTACAAGGCCAGCAGATCCCTAATTCTCTCTCCAATCAAGTGCGCTCTCCCCAGCCTGTCCCTTCTCCACGGCCACAGtcccaa\|a\|ccccccactccagtccttcccccaggatgcaaggactccaatcccccccgc |
| 2:2103:15508:15977#1#F | \| \| |
| MQ=60 isize=125 G=2 | GCCCAGTCCCCACACCTACAAGGCCAGCAGATCCCTAATTCTCTCTCCAATCAAGTGCGCTCTCCCCAGCCTGTCCCTTCTCCACGGCCACAGTCCCaa\|g\|ccccccactccagcccttccccaagatccctccaattcccctccgaacccg |
| 2:2204:6383:57817#1#F | \| \| |
| MQ=60 isize=131 T=6 | GCCCAGTCCCCACACCTACAAGGCCAGCAGATCCCTAATTCTCTCTCCAATCAAGTGCGCTCTCCCCAGCCTGTCCCTTCTCCACGGCCACAGTCCCAG\|t\|cCCCCCACTCCAGTCCTTCCCCAAGGATGCAGGACTCCAATTATACACAAg |
| 2:2204:9866:63894#1#F | \| \| |
| MQ=60 isize=234 G=15 | GCCCAGTCCCCACACCTACAAGGCCAGCAGATCCCTAATTCTCTCTCCAATCAAGTGCGCTCTCCCCAGCCTGTCCCCTCTCCACGGCCaCAGTCCCAg\|g\|CCCCCcaCtCCAGTCCTTCCCCaaggatgcagcctcagccttctccacacc |
| 2:2204:4518:82045#1#F | \| \| |
| MQ=60 isize=0 T=2 | GCCCAGTCCCcACAcctacAAGgcCaGcAGATccttaAtTCacTctCcAATcAAgTgCGCTcTcCCcAaccTGTcCCTTCaCCACagctacagtcccag\|t\|ccccccactccagtccttccccaagtatgcagcctcggcctcctccacacc |
| 2:2204:2062:87757#1#F | \| \| |
| MQ=60 isize=208 T=6 | GCCCAGTCCCCACACCTACAAGGCCAGCAGATCCCTAATTCTCTCTCCAATCAAGTGCGCTCTCCCCAGCCTGTCCCTTCTCCACGGCtaCAGTCCCAG\|t\|cccCCCACTCCaGTCCTTCCCCAAGGATGCAGCCTCAGCCTTCTCCACACC |
| 1:1105:10480:58712#2#R | \| \| |
| MQ=60 isize=-149 | aaggccagcagatccctaattctctctCCAATCAAgtGCgctaTCCCCAGCCtGTCCCTTctCCACGGCCACAGTCCCAG\|\_\|CCCCCCACtCCAGTCCTTCCCCAAGGATGCAGcCtCAGCCTTCTcCACanGACTCCAATCGTAAGTCCGTT |
| 1:2209:4553:43530#2#F | \| \| |
| MQ=60 isize=177 T=6 | CGATATATGCACATTGGAGTCntCCACGGCCACAGTCCCAG\|t\|cCCCCCACTCCAGTCCTTTCCCAAGGATGCAGCCTCAGCCTTCTCCACAtcACGtTTCCCCACAGACAAGTTCCCCACATCCTgGACTGGTAGCTGCCCAGGCCAACCC |
| 1:1216:20191:11933#2#F | \| \| |
| MQ=60 isize=173 T=37 | AAGGCTGTGGGTATTGGAGTncTCGGCCACAGTCCCAG\|T\|CCCCCCACTCCAGTCCtTCCCCAAGGATGCAGCCTcAGccTtCTCCacacCacGTTTCCCCacAGacAAGTTCCCcacatcctggacgggtacctgcccaggccaacccctg |
| 2:1214:7687:9162#2#F | \| \| |
| MQ=60 isize=303 | CACTGCGGTGCCATTGGAGTCCTCACAGTCCCAG\|\_\|CCCCCCACTCCAGTCCTTCCCCAAGGATGCAGCCTCAGCCTTCTCCACACCACGTTTCCCCACAGACAAGTTCCCCACATCCTGGACtGgTagCTGCCCAGGCCAACCCCATGgAAC |
| 2:1102:4360:98765#2#R | \| \| |
| MQ=60 isize=-200 G=2 | ctgcccttcccccggccacacgcccag\|g\|cccccccctcccgtccctccccaaagattcagcctccgccttctccccaccacgtTTCCCCACAGaCaAGTTCCCCACATCCTGGACTGGTAGCTGCCCAAGGACTCCAATGCACCCACACGT |
| 2:2101:8024:49122#2#R | \| \| |
| MQ=60 isize=-220 G=2 | cgcccag\|g\|ccccccactcccgtccttccccaagggtgccgcctcagccttctccacaccacgtttCCCCacagaCaAGtTCCCCACAtCCTGGACTGGTAGCTGCCCAGGCCAACCCCATGGAACAAGAGGACTCCAATTAAGTCACGGCG |
| 1:2112:20262:27901#2#R | \| \| |
| MQ=60 isize=-435 | cctgttcagcccaaccccatgagcccccaccagcatatgctcccaaatcaggccccgtccccacacctacaaggccagcagatccctaATtCTcTCTCCAaTcAAGTgcgCTcTCCcCAGCCTGTCCCaGGACTCCAataGCGTCTTTTC\|G\| |
| 2:1216:5765:6140#2#R | \| \| |
| MQ=60 isize=-443 | GCCCAACCCCATGAGCCCCCAGCAGCATATGCtcCCAAATCAGGCCCAGTCCCCaCACCTACAAGGCCAGCAGATCCCTAATTCTCTCTCCAATCAAGTGCGCTCTCCCCAGCCTGTCCCTTCTCCACAgnACTCCAATGTT\|G\|GGACGAAT |
| 1:1205:5406:98835#2#R | \| \| |
| MQ=60 isize=177 | AGGACtGgAGTGGGGGGGCTggGACTGTggCCGTGGAGAAGGGACAGGCCCAGtCCCCACACCTACAAggCCAGCAgATCCCTAATTCTCtCTCCAATCAAGTGTGCTCTCCCCAGCCTGTCCCTTCTanGACTCCAATACTCTCA\|T\|CCCt |
| 1:2215:16251:39833#2#R | \| \| |
| MQ=60 isize=177 | aggactggagtgggggggctGGGACTGTGGcCGTGGAGAAGGGACAGGCCCAGTCCCCACaCCTATAAGGCCAGCAGATCCCTAATTCTCTCTCCAATCAAGTGCGCTCTCCCCAGCCTGTCCCTTCTAgnActCCAATACTCTCA\|T\|CCCT |
| 1:1109:15607:89837#1#F | \| \| |
| MQ=60 isize=0 | GCCCAGTCCCCACACCTACAAGGCCAGCAGATCCCTAATTCTCTCTCCAATCAAGTGCGCTCTCCCCAGGAAAAAGATAATCCACCATGATCAAGTGTG\|T\|TTCATACCAGGGATGCAAGGATGGCTTAACATGGGAGGACTCCAATAATCc |
| 1:2105:1210:9016#1#F | \| \| |
| MQ=60 isize=0 | GCCCAGTCCCCACACCTACAAGGCCAGCAGATCCCTAATTCTCTCTCCAATCAAGTGCGCTCTCCCCAGCCTGTCCCTTCTCCACGGCCACAGTCCCAG\|A\|GGACTCCAATGGTCAGTGGGCC |
| 1:2109:20539:81692#1#F | \| \| |
| MQ=60 isize=0 | GCCCAGTCCCCACACCTACAAGGCCAGCAGATCCCTAATTCTCTCTCCAATCAAGTGCGCTCTCCCCAGCCTGTCAGGtGGATGAGGGGCTCCTTCCTT\|T\|TGTCCTCAGATGACCCAAGCCAAGtGCAGAaGGCtaGCCTCCTCCCAAAGC |
| 1:2207:15546:18931#1#F | \| \| |
| MQ=60 isize=0 | GCCCAGTCCCCACACCTACAAGGCCAGCAGATCCCTAATTCTCTCTCCAATCAAGTGCGCTCTCCCCAGCCTGTCCCTTCTCCACGGCCACAGTCCCAT\|A\|GAAAGTGGAAGGGCAAGCCAGCCTAAGACCCTCTTTATGCACGGTCTTAAt |
| 2:1104:9196:77176#1#F | \| \| |
| MQ=60 isize=0 | GCCCAGTCCCCACACCTACAAGGcCAgCaGatcccTAATgcTCTctcCAATCAAGtGagCTCTCCCCAGCCTGtCCCTTCTCctcACATACCTCcccct\|g\|AttAtGtgTAAGgcACTgTTCtaaACCttagtcTAaATtactgcatgaagt |
| 1:1113:16066:13342#2#F | \| \| |
| MQ=60 isize=285 C=33 | TTGTGAgCTGTAttgtagtcctcCCCCAGCAGCATATGCtCCCAAATCAGGcccAGTcccCACACCTACAaGgCcAgCAGaTCCCtaATTctcTCTcCaATCaagagCGCtctCCCCAGcCtGTCCCttcTCCACGgCCaCAgTCCCAG\|C\|C |
| 1:2108:19074:36910#2#F | \| \| |
| MQ=60 isize=388 C=33 | TCGCTTTCCGAGATTGGAGTCntCCCAGCAGCATATGCTCCCAAATCAGGCCCAGTCCCCACACCTACAAGGCCAGCAGATCCCTAATTCTCTCtCCAATCAagagCGCTCTCCCCAGCCTGTCCCTTCTCCACGgCCACAGTCCCAG\|C\|CC |
| 1:1107:7542:9750#2#F | \| \| |
| MQ=60 isize=281 C=33 | GTGTCGAGGAGAATTGGAGTCCTCAGCAGCATATGCTCCCAAATCAGGCCCAGTCCCCACACCTACAAGGCCAGCAGATCCCTAATTCTCTCTCCAATCAAGTGCGCTCTCCCCAGCCTGTCCCTTCTCCACGGCCACAGTCCCAG\|C\|CCCC |
| 1:2209:4269:96925#2#F | \| \| |
| MQ=60 isize=281 C=33 | GCCCTGGTCTCGATTGGAGTCntcAGCAGCATATGCTCCCAAATCAGGCCCAGTCCCCACACCTACAAGGCCAGCAGATCCCTAATTCTCTCTCCAaTCAAGTGCGCTCTCCCCAGCCTGTCCCTTCTCCACGGCCACAGTCCCAG\|C\|CCCC |
| 1:2107:3960:40701#2#F | \| \| |
| MQ=60 isize=280 C=37 | CGATGTACTAATATTGGAGTCCTAGCAGCATATGCTCCCAAATCAGGCCCAGTCCCCACACCTACAAGGCCAGCAGATCCCTAATTCTCTCTCCAATCAAGTGCGCTCTCCCCAGCCTGTCCCTTCTCCACGGCCaCAGTCCCAG\|C\|CCCCC |
| 1:1216:12141:52897#2#F | \| \| |
| MQ=60 isize=279 C=33 | CCGGGTACGTGCATTGGAGTncTGCAGCATATGCTCCCAAATCAGGCCCAGTCCCCACACCTACAAGGCCAGCAGATCCCTAATTCTCTCTCCAATCAAGTGCGCTCTCCCCAGCCTGTCCCTTCTCCACGGCCACAGTCCCAG\|C\|CCCCCC |
| 1:2214:18852:6974#2#F | \| \| |
| MQ=60 isize=277 C=33 | ACAGCTGTAAGAATTGGAGTncTAGCATATGCTCCCAAATCAGGCCCAGTCCCCACACCTACAAGGCCAGCAGATCCCTAATTCTCTCTCCAATCAAGTGCGCTCTCCCCAGCCTGTCCCTTCTCcaCGgCCaCAGtCCCAG\|C\|CCCCCCac |
| 1:2101:4010:100313#2#F | \| \| |
| MQ=60 isize=408 C=33 | GTAGGTAGGCGAATTGGAGTCCTATATGCTCCCAAATCAGGCCCAGTCCCCACACCTACAAGGCCAGCAGATCCCTAATTCTCTCTCCAATCAAGTGCGCTCTCCCCAGCCTGTCCCTTCTCCACGGCCACAGTCCCAG\|C\|CCCCCCaCTCC |
| 1:1213:3053:86051#2#F | \| \| |
| MQ=60 isize=271 C=2 | TGTACCCTAAAGATTGGAGTnctgctcCCAaATCAGgcccAGTCCCCACACCTACAagGCCagCAGATcCCtAaTTCTCTCTCcaatcaagtgcgctctccccagcctgtcccttctccacggccacagtcccag\|c\|ccccccactcgagtc |
| 1:2115:10928:96763#2#F | \| \| |
| MQ=60 isize=269 C=33 | AAACGTCTAGGCATTGGAGTCCTCCCAAATCAGGCCCAGTCCCCACACCTACAAGGCCAGCAGATCCCTAATTCTCTCTCCAATCAAGTGCGCTCTCCCCAGCCTGTCCCTTCTCCACGGCCACAGTCCCAG\|C\|CCCCCCACTCCAGTCCTt |
| 2:2112:15728:46756#2#F | \| \| |
| MQ=60 isize=269 C=2 | TGCCGAGGtGTCATTGGAGTncTCCCAAATCAGGCCCAGTCCCCACACCTACAAGGCCAGCAGATCCCTAATTCTCTCTCCAATCAAGtgcGCTCTCCCCAGCCTGTCCCTTCTCCACGGCCACAGTCCCag\|c\|ccccccactccagtcctc |
| 1:2107:17437:55497#2#F | \| \| |
| MQ=60 isize=268 C=33 | AAGGTAGTTGTGATTGGAGTCCTTCCCAAATCAGGCCCAGTCCCCACACCTACAAGGCCAGCAGATCCCTAATTCTCTCTCCAATCAAGTGCGCTCTCCCCAGCCTGTCCCTTCTCCACGGCCACAGTCCCAG\|C\|CCCCCCaCTCCaGtCCT |
| 1:2210:8422:91087#2#F | \| \| |
| MQ=60 isize=268 C=33 | tCTCaTGACTATATTGGAGTCCTTCCCAAATCAGGCCCAGtCCCCACaCCTACAaGGCCaGCAGATCCCTAATTCTCTCTCCAATCAAgagCGctCtcCCCAgcCTGTCCCTtCTCCACGGCCACAgTCCCAG\|C\|CCcccccctccagccct |
| 2:1207:3498:77010#2#F | \| \| |
| MQ=60 isize=373 C=33 | CGCGGTGCGAGTATTGGAGTCntCCCAAATCAGGCCCAGTCCCCACACCTACAAGGCCAGCAGATCCCTAATTCTCTCTCCAATCAAGTGCGCTCTCCCCAGCCTGTCCCTTCTCCACGgcCaCAgTCCCAG\|C\|CCCCCCaCtCCagtcctt |
| 1:2210:2182:50124#2#F | \| \| |
| MQ=60 isize=265 C=33 | CTACCGGTGCGCATTGGAGTCCTCAAATCAGGCCCAGTCCCCACACCTACAAGGCCAGCAGATCCCTAATTCTCTCTCCAATCAAGTGCGCTCTCCCCAGCCTGTCCCTTCTCCACGGCCACAGtCCCAG\|C\|CCCCCCACTCCAgTCCTTCC |
| 2:2203:5785:7425#2#F | \| \| |
| MQ=60 isize=501 C=33 | ATACCGATGGCCATTGGAGTCntTCTCAAATCAGGCCCAGTCCCCACACCTACAAGGCCAGCAGATCCCTAATTCTCCCTCCAATCAAGtgCGCTCTCCCCAGCCTGTCCCTTCTCCACGgCCACAGTCCCAG\|C\|CCCCCCACTCCAgtCCc |
| 2:2101:6352:30588#2#F | \| \| |
| MQ=60 isize=263 | GGCCAGTCAGGTATTGGAGTCCTAATCAGGCCCAGTCCCCACACCTACAAGGCCAGCAGATCCCTAATTCTCTCtCCAAtCAAGTgcgctctccccacccggccccttctccacgccccagtcccaag\|c\|cccccccccccgcccttccccc |
| 1:2104:16747:35033#2#F | \| \| |
| MQ=60 isize=253 C=33 | GCTTGCGTGGTGATTGGAGTCntATCAGGCCCAGTCCCCACACCTACAAGGCCAGCAGATCCCTAATTCTCTCTCCAATCAAGTGCGCTCTCCCCAGCCTGTCCCTTCTCCACGGCCACAGTCCCAG\|C\|CCCCCCaCtCCAGTCCTTCCCCa |
| 2:1202:10258:99588#2#F | \| \| |
| MQ=60 isize=262 C=33 | GAGTGGGATAATATTGGAGTCntATCAGGCCCAGTCCCCACACCTACAAGGCCAGCAGATCCCTAATTCTCTCTCCAATCAAGTGCGCTCTCCCCAGCCTGTCCCTTCTCCACGGCCACAGTCCCAG\|C\|CCCCCCACTCCAGTCCTTCCCCA |
| 1:1201:9424:79646#2#F | \| \| |
| MQ=60 isize=261 C=15 | CTGCTCGTGACGATTGGAGTCntCAGGCCCAGTCCCCACACCTACAAGGCCAGCAGATCCCTAATTCTCTCTCCAATCAAGTGCGCTCTCCCCAGCCTGTCCCTTCTCCACGGCCACAGtCCCAg\|c\|CcCCCCcCtCCagtCCTtCCCCaag |
| 2:2107:19371:58240#2#F | \| \| |
| MQ=60 isize=259 C=15 | CACGGGCAGGGTATTGGAGTCCTAGGCCCAGTCCCCACACCTACAAGGCCAGCAGATCCCTAATTCTCTCTCCAATCAAGTGCGCTCTCCCCAGCCtgtCCCTTCTCCACGGCCACAgtcCCAg\|c\|CCCCCCACTCCAGtCCTTCCCCAagg |
| 1:2107:14048:59785#2#R | \| \| |
| MQ=60 isize=-485 C=33 | cggcccagtCCCCACaCCTACAAGGCCAGCAGATCCCTAATTCTCTCtCcAATCAAGTGCGCtCTCCCCAGCCTGTCCCTTCTCCACGGCCaCAGTCCCAG\|C\|CCCCCCACTCCAGTCCTTCCCCAAGGAGGACTCCAATGACGCCCTCCTC |
| 2:1116:12229:97011#2#R | \| \| |
| MQ=60 isize=-127 C=33 | tggCCCAGtCcccaCcCCTACAAGGCCAGCAGATCCCTAATTCTCTCTCCAATCAAGTGCGCTCTCCCCAGCCTGTCCCTTCTCCACGGCCACAGTCCCAG\|C\|CCCCCCACTCCAGTCCTTCCCCAAGGAGGACTCCAATAACGTATCTCGC |
| 2:1210:1534:45884#2#R | \| \| |
| MQ=60 isize=-127 C=33 | TGGCCcactCCCCACcCCtACaagGCCcGCAgatCCCTAATTCTCtCtCCaAtCaAGtGCgCtCTCCCCAGCCTGTCCCTTCTCCACGGCCACAGTCCCAG\|C\|CCCCCCACTCCAGTCCTTCCCCAAGGAGGACTCCAATGGACAGAATCAG |
| 1:2115:15389:82105#2#F | \| \| |
| MQ=60 isize=258 C=33 | ACCAGAGCGGGAATTGGAGTCCTGGCCCAGTCCCCACACCTACAAGGCCAGCAGATCCCTAATTCTCTCTCCAATCAAGTGCGCTCTCCCCAGCCTGTCCCTTCTCCACGGCCACAGTCCCAG\|C\|CCCCCCACTCCAGTCCTTCCCCAAGGA |
| 1:1107:15969:55331#2#R | \| \| |
| MQ=60 isize=-121 C=37 | CGTTTTGGCCCAgTCCCCaCaCCTACAAGGCCAGCAGATCCCTAaTTCTCTCTCCAATCAAGTGCGCTCtCCCCAGCCTGTCCCTTCTCCACGGCCACAGTCCCAG\|C\|CCCCCCACTCCAGTCCTTCCCAGGACTCCAATCACTCCGACCCG |
| 1:1114:7852:4888#2#R | \| \| |
| MQ=60 isize=-128 C=37 | GTTTTGGCCCAGTCCCCACaCCTACAAGGCCAGCAGATCCCTAATTCTCTCTCCAATCAAGTGCGCTCTCCCCAGCCTGTCCCTTCTCCACGGCCACAGTCCCAG\|C\|CCCCCCACTCCAGTCCTTCCCCAAGGATCCAATCGACACAGCTAA |
| 1:1214:16634:96231#2#R | \| \| |
| MQ=60 isize=-125 C=2 | ttggcccagtccccacacctacaaggccagcagatccctaattctctctccaatcaagtgcgctctccccagcctgtcccttctccacggccacagtcccag\|c\|cCCCCCaCtCCAGTCCTTCCCCAAGAgnACTCCAATtCTCAggTGTCG |
| 1:2101:17723:43263#2#R | \| \| |
| MQ=60 isize=-123 C=37 | GTTTTGGCCCAgTCCCCacaCCTACAAGgCCaGCAGATCCCTAATTCTCtCTCCAATCAAGTGCGCTCTCCCCAGCCTGTCCCTTCTCCACGGCCaCAGTCCCAG\|C\|CCCCCCACTCCAGTCCTTCCCCAGGACTCCAATCTGCCGAGACaT |
| 1:2102:19958:27039#2#R | \| \| |
| MQ=60 isize=-121 C=37 | CGTTTTGGCCCAGTCCCCACaCCTACAAGGCCAGCAGATCCCTAATTCTCTCTCCAATCAAGTGCGCTCTCCCCAGCCTGTCCCTTCTCCACGGCCACAGTCCCAG\|C\|CCCCCCACTCCAGTCCTTCCCAGGACTCCAATGTGAGTCTGATA |
| 1:2105:7102:40577#2#R | \| \| |
| MQ=60 isize=-125 C=37 | TTGGCCCAGTCCCCaCACCTACAaGGCCAGCAGATCCCTAATTCTCTCTCCAATCAAGtGCGCtCtCCCCAGCCTGTCCCTTCTCCACGGCCACAGTCCCAG\|C\|CCCCCCACTCCAGTCCTTCCCCAAGanGACTCCAATTCTCAGGTGTCG |
| 1:2111:7952:14112#2#R | \| \| |
| MQ=60 isize=-125 C=33 | TTGGCCCAGtCCCCACaCCtACAAGGCCAGCAGATCCCTAATTCTCTCTCCAAtCAAGTGCGCTCTCCCCAGCCTGTCCCTTCTCCACGGCCaCAGTCCCAG\|C\|CCCCCCACTCCAGTCCTTCCCCAAGAGGACTCCAATAGATGGCGATGA |
| 1:2202:17756:27179#2#R | \| \| |
| MQ=60 isize=-125 C=33 | ttggcccagtccccacccctacAAGGCCAGCAGATCCCTAattCTCgCtCCAATCAAGTGCGCtCtCCCCAGCCTGtCCCTtCtCCACGGCCaCAGTCCCAG\|C\|CCCCCCACTCCAGTCCTTCCCCAAGanGACTCCAAtGCCCCCAGTCCG |
| 1:2205:1800:24401#2#R | \| \| |
| MQ=60 isize=-127 C=33 | ttttggcccagtCCCCAccCCTaCAAGgCCAGCAGaTCCCTAattCtCtctCCAatCAAGTGCGCtCtCCCCAGCCTGTCCCTtCTCCACGGCCACAGTCCCAG\|C\|CCCCCCACTCCAGTCCTTCCCCaanGACTCCAATTCTGACCACGCC |
| 1:2207:4222:16741#2#R | \| \| |
| MQ=60 isize=-125 C=37 | TTGGCCCAGTCCCCACACCTACAAGGCCAGCAGATCCCTAAtTCTCTCTCCAATCAAGTGCGCTCTCCCCAGCCTGTCCCTTCTCCACGGCCACAGTCCCAG\|C\|CCCCCCACTCCAGTCCTTCCCCAAGanGACTCCAATCGCTGCCTCCGC |
| 1:2216:4016:30670#2#R | \| \| |
| MQ=60 isize=-125 C=37 | TTTTGGCCCAGTCCCCACACCTAcaAGGCCAGCAGATCCCTAAttCTCTCTCCAATCAAGTGCGCTCTCCCCAGCCTGTCCCTTCtCCACGGCCACAGTCCCAG\|C\|CCCCCCaCTCCAGTCCTTCCCCAAgnACTCCAatCCCCTCCCCCAA |
| 2:1106:13999:42952#2#R | \| \| |
| MQ=60 isize=-124 C=37 | tttggccCAGTCCCCaCaCCTACAAGGCCAGCAgaTCCCTAATtCCCTCTCCAATCAAGTGCGCtCTCCCCAGCCTGTCCCTTCTCCACGGCCACAGTCCCAG\|C\|CCCCCCACTCCAGTCCTTCCCCAAanGACTCCAATCaCCGAATCCCA |
| 2:1112:13497:6706#2#R | \| \| |
| MQ=60 isize=-127 C=33 | tTTTGGCCCAGtCCCCACACCTACAAGGCCAGCAGaTCCCTAATTCTCTCTCCAATCAAGTGCGCTCTCCCCAGCCTGTCCCTTCTCCACGGCCACAGTCCCAG\|C\|CCCCCCACTCCAGTCCTTCCCCAAGGACTCCAATCAAAACCGTAAG |
| 2:1116:19644:3150#2#R | \| \| |
| MQ=60 isize=-121 C=33 | cgTTTTGGCCcagtCCCCAccCCTACAAGGCCAGCAgaTCCcTaatTCtctctCCAATCAAGTGCGCTCTCCCcAGCCTGTCCCTTCTCCACGGCCACAGTCCCAG\|C\|CCCCCCACTCCAGTCCTTCCCAGGACTCCAATTCCGCCTGTCTG |
| 2:1204:14406:8597#2#R | \| \| |
| MQ=60 isize=-127 C=33 | TTTTgGCCcAGTCCCCACaCCTACAAGGCCAGCAGATCCCTAATTCTCTCTCCAATCAAGTGCGCTCTCCCCAGCCTGTCCCTTCTCCACGGCCACAGTCCCAG\|C\|CCCCCCACTCCAGTCCTTCCCCAanGACTCCAATGCCTTCCCCGGC |
| 2:1215:12337:60139#2#R | \| \| |
| MQ=60 isize=-121 C=37 | CGTTTTGGCCCAGTCCCCACACCTACAAGGCCAGCAGATCCCTAATTCTCTCTCCAATCAAGTGCGCTCTCCCCAGCCTGTCCCTTCTCCACGGCCACAGTCCCAG\|C\|CCCCCCACTCCAGTCCTTCCCAgnACTCCAATGCAATCCCACGG |
| 2:2103:15508:15977#2#R | \| \| |
| MQ=60 isize=-125 C=37 | ttggcccagcccccacacctacaaggccagcagatcactaatcctccctcCAAtCAAgtgcGctctCCCCAGCCTGTCCCTTCtcCACGGCcaCAGTCCCAG\|C\|CCCCCCACTCCAGTCCTTCCCCAAGAGGACTCCAATTCCCCCCCGAAC |
| 2:2108:19838:27641#2#R | \| \| |
| MQ=60 isize=-120 C=33 | gcgtTTTGGCCCAGTCCCcacccCTacagGGCCAGCAggtCccTAAttCtCtCtcCAATCaagTGCGCTcTCCCCAGCCtGTCCcTgCTCCaCGGCCACAgTCCCAG\|C\|CCCCCCACTCcAGTCCTTCCAGGACTCCAATAACACGCTAGGA |
| 2:2203:9067:34326#2#R | \| \| |
| MQ=60 isize=-127 C=37 | TTTTGGCCCaGtCCCCACACCTACAAGGCCAGCAGATCCCTAATTCtCtCTCCAATCAAGTGCGCtCTCCCCAGCCTGTCCCTTCTCCACGGCCACAGTCCCAG\|C\|CCCCCCACTCCAGTCCTTCCCCAanGACTCCAATTGACGGCAGGTC |
| 2:2209:16670:45509#2#R | \| \| |
| MQ=60 isize=-120 C=33 | GCGTTTTGGCCCAGTCCCCACACCTACAAgGCCAGCAgatCCCTAATTCTCTCTCCAATCAAGTGCGCTCTCCCCAGCCTGTCCCTTCTCCACGGCCACAGTCCCAG\|C\|CCCCCCACTCCAGTCCTTCCanGACTCCAATCGACCATTTTGC |
| 2:2212:14081:79401#2#R | \| \| |
| MQ=60 isize=-120 C=33 | gCGTTTTGGCCCAGTCCCCaCACCTACAAGGCCAGCAGATCCCTAATTCTCTCTCCAATCAAGTGCGCTCTCCCCAGCCTGTCCCTTCTCCaCGGCCACAGTCCCAG\|C\|CCCCCCACTCCAGTCCTTCCAGGACTCCAATGGCCACCCTTCG |
| 2:2214:8249:43957#2#R | \| \| |
| MQ=60 isize=-127 C=33 | GGCCCAGTCCCCACaCCTACAAGGCCAGCAGATCCCTAATTCTCTCTCCAATCAAGTGCGCTCTCCCCAGCCTGTCCCTTCTCCACGGCCACAGTCCCAG\|C\|CCCCCCACTCCAGTCCTTCCCCAAGGAAGGACTCCAATAACGCCTGTCAC |
| 1:1101:1249:61375#1#F | \| \| |
| MQ=60 isize=175 C=37 | GCCCAGTCCCCACACCTACAAGGCCAGCAGATCCCTAATTCTCTCTCCAATCAAGTGCGCTCTCCCCAGCCTGTCCCTTCTCCACGGCCACAGTCCCAG\|C\|CCCCCCACTCCAGTCCTTCCCCAAGGATGCAGCCTCAGCCTTCTCCACACC |
| 1:1102:18948:8131#1#F | \| \| |
| MQ=60 isize=143 C=37 | GCCCAGTCCCCACACCTACAAGGCCAGCAGATCCCTAATTCTCTCTCCAATCAAGTGCGCTCTCCCCAGCCTGTCCCTTCTCCACGGCCACAGTCCCAG\|C\|CCCCCCACTCCAGTCCTTCCCCAAGGATGCAGCCTCAGCCTTCAGGACTCC |
| 1:1102:10780:41113#1#F | \| \| |
| MQ=60 isize=168 C=37 | GCCCAGTCCCCACACCTACAAGGCCAGCAGATCCCTAATTCTCTCTCCAATCAAGTGCGCTCTCCCCAGCCTGTCCCTTCTCCACGGCCACAGTCCCAG\|C\|CCCCCCACTCCAGTCCTTCCCCAAGGATGCAGCCTCAGCCTTCTCCaCACC |
| 1:1102:12162:70923#1#F | \| \| |
| MQ=60 isize=143 C=37 | GCCCAGTCCCCACACCTACAAGGCCAGCAGATCCCTAATTCTCTCTCCAATCAAGTGCGCTCTCCCCAGCCTGTCCCtTCTCCACGGCCACAGtCCCAG\|C\|CCCCCCACTCCAGtcCTTCCCCAAGGAtgCaGCCTCAGCCTTCAGGACTCC |
| 1:1102:19514:91277#1#F | \| \| |
| MQ=60 isize=147 C=37 | GCCCAGT\_CCCACACCTACAAGGCCAGCAGATCCCTAATTCTCTCTCCAATCAAGTGCGCTCTCCCCAGCCTGTCCCTTCTCCACGGCCACAGTCCCAG\|C\|CCCCCCACTCCAGTCCTTCCCCAAGGATGCAGCCTCAGCCTTCTCCAGGACt |
| 1:1103:17951:22979#1#F | \| \| |
| MQ=60 isize=125 C=15 | GCCCAGTCCCCATACCTACAAGGCCAGCAGATCCCTAATTCTCCCTCCAATCAAGTGCGCTCTCCCCAGCCTGTCCCTTCTCCACGGCCACAGTCCCag\|c\|cccCCCaCtCCagcccttcCCCaagaggaCtCCacttCCCCCCccaacccc |
| 1:1103:11122:33041#1#F | \| \| |
| MQ=60 isize=149 C=37 | GCCCAGTCCCCACACCTACAAGGCCAGCAGATCCCTAATTCTCTCTCCAATCAAGTGCGCTCTCCCCAGCCTGTCCCTTCTCCACGGCCACAGTCCCAG\|C\|CCCCCCACTCCAGTCCTTCCCCAAGGATGCAGCCTCAGCCTTCTCCACAGG |
| 1:1103:12939:69974#1#F | \| \| |
| MQ=60 isize=255 C=33 | GCCCAGTCCCCACACCTACAAGGCCAGCAGATCCCTAATTCTCTCTCCAATCAAGTGCGCTCTCCCCAGCCTGTCCCTTCTCCACGGCCACAGTCCCAG\|C\|CCCCCCACTCCAGTCCTTCCCCaAgGATGCAGCCTCAGCCTTCTCCACACC |
| 1:1104:17635:16813#1#F | \| \| |
| MQ=60 isize=242 C=33 | GCCCAGTCCCCACACCTACAAGGCCAGCAGATCCCTAATTCTCTCTCCAATCAAGTGCGCTCTCCCCAGCCTGTCCCTTCTCCACGGCCACAGTCCCAG\|C\|CCCCCCACTCCAGtCCTTCCCCAAGGATGCAGCCTCAGCCTTCTCCACACC |
| 1:1104:9410:72943#1#F | \| \| |
| MQ=60 isize=220 C=33 | GCCCAGTCCCCACACCTACAAGGCCAGCAGATCCCTAATTCTCTCTCCAATCAAGTGCGCTCTCCCCAGCCTGTCCCTTCTCCACGGCCACAGTCCCAG\|C\|CCCCCCACTCCAGTCCTTCCCCAAGGAtGCAGCCTCAGCCTTCTCCACACC |
| 1:1104:11752:100886#1#F | \| \| |
| MQ=60 isize=197 C=37 | GCCCAGTCCCCACACCTACAAGGCCAGCAGATCCCTAATTCTCTCTCCAATCAAGTGCGCTCTCCCCAGCCTGTCCCTTCTCCACGGCCACAGTCCCAG\|C\|CCCCCCACTCCAGTCCTTCCCCAAGGATGCAGCCTCAGCCTTCTCCACACC |
| 1:1105:14410:12027#1#F | \| \| |
| MQ=60 isize=169 C=33 | GCCCAGTCCCCACACCTACAAGGCCAGCAGATCCCTAATTCTCTCTCCAATCAAGTGCGCTCTCCCCAGCCTGTCCCTTCTCCACGGCCACAGTCCCAG\|C\|CCCCCCACTCCAGTCCTTCCCCAAGGATgCAgCCtCAGCCTTCTCCacacc |
| 1:1105:15787:30488#1#F | \| \| |
| MQ=60 isize=162 C=33 | GCCCAGTCCCCACACCTACAAGGCCAGCAGATCCCTAATTCTCTCTCCAATCAAGTGCGCTCTCCCCAGCCTGTCCCTTCTCCACGGCCACAGTCCCAG\|C\|CCCCCCACTCCaGTCCTTCCCCAAGGATGCaGCCTCAGCCTTCTCCACACC |
| 1:1105:2976:89067#1#F | \| \| |
| MQ=60 isize=147 C=37 | GCCCAGTCCCCACACCTACAAGGCCAGCAGATCCCTAATTCTCTCTCCAATCAAGTGCGCTCTCCCCAGCCTGTCCCTTCTCCGCGGCCACAGTCCCAG\|C\|CCCCCCACTCCAGTCCTTCCCCAAGGATGCAGCCTCAGCCTTCTCCAGGAC |
| 1:1106:18835:3196#1#F | \| \| |
| MQ=60 isize=135 C=33 | GCCCAGTCCCCACACCTACAAGGCCAGCAGATCCCTAATTCTCTCTCCAATCAAGTGCGCTCTCCCCAGCCTGTCCCTTCTCCACGGCCACAGTCCCAG\|C\|CCCCCCACTCCAGTCCTTCCCCAAGGATGCAGCCTAGGACTCCAATCGATA |
| 1:1106:6944:32050#2#F | \| \| |
| MQ=60 isize=257 C=33 | TCGTCGATTAAAATTGGAGCCngCCCAGTCCCCACACCTACAAGGCCAGCAGATCCCTAATTCTCTCTCCAATCAAGTGCGCTCTCCCCAGCCTGTCCCTTCTCCACGGCCACAGTCCCAG\|C\|CCCCCCACTCCAGTCCTTCCCCAAGGATG |
| 1:1106:3504:66811#1#F | \| \| |
| MQ=60 isize=139 C=33 | GCCCAGTCCCCACACCTACAAGGCCAGCAGATCCCTAATTCTCTCTCCAATCAAGTGCGCTCTCCCCAGCCTGTCCCTTCTCCACGGCCACAGTCCCAG\|C\|CCCCCcactccagtcctcccccaaggatgcagcctcagcgggaccccaatc |
| 1:1106:16189:91235#1#F | \| \| |
| MQ=60 isize=266 C=22 | GCCCAGcCCCcACACCTACAAGGCCAGCAGATCCCTAATTCTCTCTCCAATCAaGtGCGCTCTCCCCAGCCTGTCCCTTCTCCACGGCCACAgtCCCAG\|C\|CCCCCCACtCCAGtccTTCCCCAAGgaTGcAGCCTCAGCCTTCTCCACACC |
| 1:1107:14173:13249#1#F | \| \| |
| MQ=60 isize=307 C=33 | GCCCAGTCCCCACACCTACAAGGCCGGCAGATCCCTAATTCTCTCTCCAATCAAGTGCGCTCTCCCCAGCCTGTCCCTTCTCCACGGCCACAGTCCCAG\|C\|CCCCCCACTCCAGTCCTTCCCCAaGGATGCAGCCTCAGCCTTCTCCACACC |
| 1:1107:12410:31751#1#F | \| \| |
| MQ=60 isize=201 C=33 | GCCCAGTCCCCACACCTACATGGCCAGCAGATCCCTAATTCTCTCTCCAATCAAGTGCGCTCTCCCCAGCCTGTCCCTTCTCCACGGCCACAGTCCCAG\|C\|CCCCCCACTCCAGTCCTTCCCCAAGGATGCAGCCTCAGCCTTCTCCACACC |
| 1:1107:19432:34430#1#F | \| \| |
| MQ=60 isize=138 C=33 | GCCCAGTCCCCACACCTACAAGgCCAGCAGATCCCTAATTcTCTCTCCAATCAAGtGCGCTCTCCCCAgCCTGTCCCTTCTCCACGGCCAcaGTcCCAG\|C\|CCCCCCACtCCAGtCCTTCCCCAAGGAtGCAGCCtcAGAGGACtCCaATCC |
| 1:1107:8454:44999#1#F | \| \| |
| MQ=60 isize=126 C=37 | GCCCAGTCCCCACACCTACAAGGCCAGCAGATCCCTAATTCTCTCTCCAATCAAGTGCGCTCTCCCCAGCCTGTCCCTTCTCCACGGCCACAGTCCCAG\|C\|CCCCCCACTCCAGTCCTTCCCCAAGGCTGAGGCTGCATCCTTGGGGAAGAG |
| 1:1107:19687:87027#1#F | \| \| |
| MQ=60 isize=120 C=33 | GCCCAGTCCCCACACCTACAGGGCCAGCAGATCCCTAATTCTCTCTCCAATCAAGTGCGCTCTCCCCAGCCTGTCCCTTCTCCACGGCCACAGTCCCAG\|C\|CCCCCCACTCCAGTCCTTCCAGGACTCCAATAGATGTACCTCCCTGTCTCT |
| 1:1108:5684:27522#1#F | \| \| |
| MQ=60 isize=149 C=22 | GCCCAGTCCCCACACCTACAAGGCCAGCAGATCCCTAATTCTCTCTCCAATCAAGTGCGCTCTCCCCAGCCTGTCCCTTCTCCACGGCCACAGtCCCag\|C\|CCCCCCACTCCAGTCCTTCCCCAAGGATGCAGCCTCAgCCTTCTCCACaGG |
| 1:1108:4360:57933#1#F | \| \|      T |
| MQ=60 isize=139 C=33 | GCCCAGTCCCCACACCTACAAGGCCAGCAGATCCCTAATTCTCTCTCCAATCAAGTGCGCTCTCCCCAGCCTGTCCCTTCTCCACGGCCACAGTCCCAG\|C\|CCCCCIACTCCAGTCCTTCCCCAAGGATGCAGCCTCAGCAGGACTCCAAt |
| 1:1108:12360:70423#1#F | \| \| |
| MQ=60 isize=210 C=33 | GCCCAGTCCCCACACCTACAAGGCCAGCAGATCCCTAATTCTCTCTCCAATCAAGTGCGCTCTCCCCAGCCTGTCCCTTCTCCACGGCCACAGTCCCAG\|C\|CCCCCCACTCCAGTCCTTCCCCAAGGATGCAGCCTCAGCCTTCTCCACACC |
| 1:1109:8900:29951#1#F | \| \| |
| MQ=60 isize=174 C=37 | GCCCAGTCCCCACACCTACAAGGCCAGCAGATCCCTAATTCTCTCTCCAATCAAGTGCGCTCTCCCCAGCCTGTCCCTTCTCCACGGCCACAGTCCCAG\|C\|CCCCCCACTCCAGTCCTTCCCCAAGGATGCAGCCTCAGCCTTCTCCACACC |
| 1:1109:2639:56775#1#F | \| \| |
| MQ=60 isize=0 C=33 | GCCCAGTCCCCACACCTACAAGGCCAGCAGATCCCTAATTCTCTCTCCAATCAAGTGCGCTCTCCCCAGCCTGTCCCTTCTCCACGGCCACAGTCCCAG\|C\|CCCCCCACTCCAGTCCTTCCCCcAGGATGCAGCCTCAGCCTTCTCCCCGGA |
| 1:1110:7685:22675#1#F | \| \| |
| MQ=60 isize=150 C=37 | GCCCAGTCCCCACACCTACAAGGCCAGCAGATCCCTAATTCTCTCTCCAATCAAGTGCGCTCTCCCCAGCCTGTCCCTTCTCCACGGCCACAGTCCCAG\|C\|CCCCCCACTCCAGTCCTTCCCCAAGGATGCAGCCTCAGCCTTCTCCACACa |
| 1:1110:15467:59848#1#F | \| \| |
| MQ=60 isize=147 C=33 | GCCCAGTCCCCACACCTACAAGGCCAGCAGATCCCTAATTCTCTCTCCAATCAAGTGCGCTCTCCCCAGCCTGTCCCTTCTCCACGGCCACAGTCCCAG\|C\|CCCCCCACTCCAGTCCTTCCCCAAGGATGCAGCCTCAGCCTTCTCCAGGAC |
| 1:1110:15052:86338#1#F | \| \| |
| MQ=60 isize=0 C=37 | GCCCAGTCCCCACACCTACAAGGCCAGCAGATCCCTAATTCTCTCTCCAATCAAGTGCGCTCTCCCCAGCCTGTCCCTTCTCCACGGCCACAGtCCCAG\|C\|CCCCCCACTCCAGTCCTTCCCCAAGGATGCAGCCTCAGCCTTCTCCACaCC |
| 1:1111:19882:3660#1#F | \| \| |
| MQ=60 isize=187 C=33 | GCCCAGTCCCCACACCTACAAGGCCAGCAGATCCCTAATTCTCTCTCCAATCAAGTGCGCTCTCCCCAGCCTGTCCCTTCTCCACGGCCACAGTCCCAG\|C\|CCCCCCACTCCAGTCCTTCCCCAAGGATGCAGCCTCAGCCTTCTCCACACC |
| 1:1111:14140:61596#1#F | \| \| |
| MQ=60 isize=160 C=33 | GCCCAGTCCCCACACCTACAAGGCCAGCAGATCCCTAATTCTCTCTCCAaTCAaGTGCGCTCTCCCCAGCCTGTCCCTTCTCCACGGCCACAGTCCCAG\|C\|CCCCCCACTCCAGTCCTTCCCCAAGGATGCAGCCTCAGCCTTCTCCACaCC |
| 1:1111:16255:74385#1#F | \| \| |
| MQ=60 isize=230 C=37 | GCCCAGTCCCCACACCTACAAGGCCAGCAGATCCCTAATTCTCTCTCCAATCAAGTGCGCTCTCCCCAGCCTGTCCCTTCTCCACGGCCACAGTCCCAG\|C\|CCCCCCaCTCCAGTCCTTCCCCAAGGATGCAGCCtCAGCCTTCTCCACACC |
| 1:1111:7534:85552#1#F | \| \| |
| MQ=60 isize=240 C=33 | GCCCAGTCCCCACACCTACAGGGCCAGCAGATCCCTAATTCTCTCTCCAATCAAGTGCGCTCTCCCCAGCCTGTCCCTTCTCCACGGCCACAGTCCCAG\|C\|CCCCCCACTCCAGTCCTTCCCCAAGGATGCAGCCTCAGCCTTCTCCACACC |
| 1:1111:14675:93972#1#F | \| \| |
| MQ=60 isize=187 C=33 | GCCCAGTCCCCACACCTACAAGGCCAGCAGATCCCTAATTCTCTCTCCAATCAAGTGCGCTCTCCCCAGCCTGTCCCTTCTCCACGGCCACAGTCCCAG\|C\|CCCCCCACTCCAGtCCTTCCCCAAGGATGCAGCCTCAGCCTTCTCCaCACC |
| 1:1112:5839:11355#1#F | \| \| |
| MQ=60 isize=132 C=33 | GCCCAGTCCCCACACCTACAAGGCCAGCAGATCCCTAATTCTCTCTCCAATCAAGTGCGCTCTCCCCAgCCTGTCCCTTCTCCACGGCCaCagTCCCAG\|C\|CCCCCCACTCCagtcCtTCCCCaAGGATGCAGAGGaCTCCaATTCGAATta |
| 1:1112:12843:40179#1#F | \| \| |
| MQ=60 isize=187 C=22 | GCCCAGTCCCCACACCTACAAGGCCAGCAGATCCCTAATTCTCTCTCCAATCAAGTGCGCTCCCCCCAGCCTGTCCCtTCTCCACGgCCACAgtCCCAG\|C\|CCCCCCaCTCCAGTCCttCCCCAaggccgccGCCTtagcCttCTCCACccc |
| 1:1112:6609:54692#1#F | \| \| |
| MQ=60 isize=203 C=33 | GCCCAGTCCCCACACCTACAAGGCCAGCAGATCCCTAATTCTCTCTCCAATCAAGTGCGCTCTCCCCGGCCTGTCCCTTCTCCACGGCCACAGTCCCAG\|C\|CCCCCCACTCCAGTCCTTCCCCAAGGATGCAGCCTCAGCCTTCTCCACACC |
| 1:1113:1974:8827#1#F | \| \| |
| MQ=60 isize=189 C=15 | GCCCAGTCCCCACACCTACAAGGCCAGCAGATCCCTAATTCTCTCTCCAATCAAGTGCGCTCTCCCCAGCCTGTCCCTTCTCCACGGCCACaGTcCCag\|c\|ccCCCCccccccggccttccccacaggcgccacctccgccctccccccccc |
| 1:1113:19689:33319#1#F | \| \| |
| MQ=60 isize=139 C=33 | GCCCAGTCCCCACACCTACAAGGCCAGCAGATCCCTAATTCTCTCTCCAATCAAGTGCGCTCTCCCCAGCCTGTCCCTTCTCCACGGCCACAGtCCCAG\|C\|CCCCCCACTCCAGTCCTTCCCCAAGGATGCAGCCTCAGCAGGACTCCAATT |
| 1:1113:20270:73654#1#F | \| \| |
| MQ=60 isize=261 C=37 | GCCCAGTCCCCACACCTACAAGGCCAGCAGATCCCTAATTCTCTCTCCAACCAAGTGCGCTCTCCCCAGCCTGTCCCTTCTCCACGGCCACAGTCCCAG\|C\|CCCCCCACTCCAGTCCTTCCCCAAGGATGCAGCCTCAGCCTTCTCCACACC |
| 1:1113:13542:89001#1#F | \| \| |
| MQ=60 isize=147 C=37 | GCCCAGTCCCCACACCTACAAGGCCAGCAGATCCCTAATTCTCTCTCCAATCAAGTGCGCTCTCCCCAGCCTGTCCCTTCTCCACGGCCACAGTCCCAG\|C\|CCCCCCACTCCAGTCCTTCCCCAAGGATGCAGCCTCAGCCTTCTCCAGGAC |
| 1:1114:3568:19305#1#F | \| \| |
| MQ=60 isize=144 C=33 | GCCCAgTCCCCACACCTACAAGGCCAGCAGATCCCTAATTCTCTCTCCAATCAAGTGCGCTCTCCCCAGCCTGTCCCTTCTCCACGGCCACAATCCCAG\|C\|CCCCCCACTCCAGTCCTTCCTCAAGGATGCAGCCTCAGCCTTCTAGGACTC |
| 1:1114:5282:71270#1#F | \| \| |
| MQ=60 isize=300 C=33 | GCCCAGTCCCCACACCTACAAGGCCAGCAGATCCCTAATTCTCTCTCCAATCAAGTGCGCTCTCCCCAGCCTGTCCCTTCTCCACGGCCACAGTCCCAG\|C\|CCCCCCACTCCAGTCCTTCCCCAAGGATGCAGCCTCAGCCTTCTCCACACC |
| 1:1114:13814:86269#1#F | \| \| |
| MQ=60 isize=235 C=37 | GCCCAGTCCCCACACCTACAAGGCCAGCAGATCCCTAATTCTCTCTCCAATCAAGTGCGCTCTCCCCAGCCTGTCCCTTCTCCACGGCCACAGTCCCAG\|C\|CCCCCCACTCCAGTCCTTCCCCAAGGATGCAGCCTCAGCCTTCtCCACACC |
| 1:1115:4532:40840#1#F | \| \| |
| MQ=60 isize=160 C=27 | GCCCAGTCCCCACACCTACAAGGCCAGCAGATCCCTAATTCTCTCTCCAATCAAGTGCGCTCTCCCCAGCCTGTCCCTTCTCCACGGCCACAGTCCCAG\|C\|CCCCCCACTCCAGTCCTTCCCCAAGGATGCAGCCTCAGCCTTCTCCACACC |
| 1:1115:19376:89263#1#F | \| \| |
| MQ=60 isize=351 C=33 | GCCAAGTCCCCACACCTACAAGGCCAGCAGATCCCTAATTCTCTCTCCAATCAAGtGCGCTCTCCCCAGCCTGTCCCTTCTCCACGGCCACAGTCCCAG\|C\|CCCCCCACTCCAGtCCTTCCCCAAGGATGCAGCCTCAGCCTTCTCCACACC |
| 1:1116:13944:35949#1#F | \| \| |
| MQ=60 isize=170 C=33 | GCCCAGTCCCCACACCTACAAGGCCAGCAGATCCCTAATTCTCTCTCCAATCAAGTGCGCTCTCCCCAGCCTGTCCCTTCTCCACGGCCACAGTCCCAG\|C\|CCCCCCACTCCAGTCCTTCCCCAAGGATGCAGCCTCAGCCTTCTCCACACC |
| 1:1116:4085:84989#1#F | \| \| |
| MQ=60 isize=212 C=33 | GCCCAGTCCCCACACCTACAAGGCCAGCAGATCCCTAATTCTCTCTCCAATCAAGTGCGCTCTCCCCAGCCTGTCCCTTCTCCACGGCCACAGTCCCAG\|C\|CCCCCCACTCCaGTCCTTCCCCAAGGATGCAGCCTCAGCCTTCTCCACACC |
| 1:1201:8153:4334#1#F | \| \| |
| MQ=60 isize=160 C=37 | GCCCAGTCCCCACACCTACAAGGCCAGCAGATCCCTAATTCTCTCTCCAATCAAGTGCGCTCTCCCCAGCCTGTCCCTTCTCCACGGCCACAGT\_CCAG\|C\|CCCCCCACTCCAGTCCTTCCCCAAGGATGCAGCCTCAGCCTTCTCCaCACcc |
| 1:1201:14897:38639#1#F | \| \| |
| MQ=60 isize=133 C=33 | GCCCAGTCCCCACACCTACAAGGCCAGCAGATCCCTAATTCTCTCTCCAATCAAGTgCGCTCTCCCCAGCCTGTCCCTTCTCCACGGCCACAGtcCCAG\|C\|CcCCCCACtcCggtCCTTCCCCAAGGATGCAgcaggactccaatctaaatc |
| 1:1201:15824:63826#1#F | \| \| |
| MQ=60 isize=158 C=22 | GCCCAGTCCCCACACCTACAAGGCCAGCAGATCCCTAATTCTCTcTCCAATCAAGTGCGCTCTCCCCAgcCTGTCCCtTCTCCACGGCCACAGtCCcAG\|C\|CCCCCcAcTccaGTCCTTCCCCAAgGAtGcagcCtccgccttccccacacc |
| 1:1201:13091:75531#1#F | \| \| |
| MQ=60 isize=205 C=37 | GCCCAGTCCCCACACCTACAAGGCCAGCAGATCCCTAATTCTCTCTCCAATCAAGTGCGCTCTCCCCAGCCTGTCCCTTCTCCACGGCCACAGTCCCAG\|C\|CCCCCCACTCCAGTCCTTCCCCAAGGATGCAGCCTCagCCTTCTCCaCACC |
| 1:1202:6181:8644#1#F | \| \| |
| MQ=60 isize=205 C=37 | GCCCAGTCCCCACACCTACAAGGCCAGCAGATCCCTAATTCTCTCTCCAATCAAGTGCGCTCTCCCCAGCCTGTCCCTTCTCCACGGCCACAGTCCCAG\|C\|CCCCCCACTCCAGTCCTTCCCCAAGGATGCAGCCTCAGCCtTCTCCACACC |
| 1:1202:12528:18956#1#F | \| \| |
| MQ=60 isize=177 C=33 | GCCCAGTCCCCACACCTACAAGGCCAGCAGATCCCTAATTCTCTCTCCAATCAAGTGCGCTCTCCCCAGCCTGTCCCTTCTCCACGGCCACAGTCCCAG\|C\|CCCCCCacTCCAgtCCTTCCCCAAGGatGCaGCctcAGCcTtCtCCACACc |
| 1:1202:13206:45960#1#F | \| \| |
| MQ=60 isize=197 C=37 | GCCCAGTCCCCACACCT\_CAAGGCCAGCAGATCCCTAATTCTCTCTCCAATCAAGTGCGCTCTCCCCAGCCTGTCCCTTCTCCACGGCCACAGTCCCAG\|C\|CCCCCCACTCCAGtCCTTCCCCAAGGATGCAGCCTCAGCCTTCTCcacaccc |
| 1:1203:16186:17864#1#F | \| \| |
| MQ=60 isize=271 C=33 | GCCCAGTCCCCACACCTACAAGGCCAGCAGATCCCTAATTCTCTCTCCAATCAAGTGCGCTCTCCCCAGCCTGTCCCTTCTCCACGGCCACAGTCCCAG\|C\|CCCCCCACTCCAGTCCTTCCCCAAGGATGCAGCCTCAGCCTTCTCCACACC |
| 1:1204:10286:10347#1#F | \| \| |
| MQ=60 isize=232 C=33 | GCCCAGTCCCCACACCTACAAGGCCAGCAGATCCCTAATTCTCTCTCCAATCAAGTGCGCTCTCCCCAGCCTGTCCCTTCTCCACGGCCACAGTCCCAG\|C\|CCCCCCACTCCAGTCCTTCCCCAAGGATGcAgCCTCAGCCTTctcCACACC |
| 1:1204:14912:17011#1#F | \| \| |
| MQ=60 isize=144 C=37 | GCCCAGTCCCCACACCTACAAGGCCAGCAGATCCCTAATTCTCTCTCCAATCAAGTGCGCTCTCCCCAGCCTGTCCCTTCTCCACGGCCACAGTCCCAG\|C\|CCCCCCACTCCAGTCCTTCCCCAAGGATGCAGCCTCAGCCTTCTAGGACTC |
| 1:1204:6769:23442#1#F | \| \| |
| MQ=60 isize=212 C=37 | GCCCAGTCCCCACACCTACAAGGCCAGCAGATCCCTAATTCTCTCTCCAATCAAGTGCGCTCTCCCCAGCCTGTCCCTTCTCCACGGCCACAGTCCCAG\|C\|CCCCCCACTCCAGTCCTTCCCCAAGGATGCAGCCTCAGCCTTCTCCACACC |
| 1:1204:2723:66789#1#F | \| \| |
| MQ=60 isize=207 C=37 | GCCCAGTCCCCACACCTACAAGGCCAGCAGATCCCTAATTCTCTCTCCAATCAAGTGCGCTCTCCCCAGCCTGTCCCTTCTCCACGGCCACAGTCCCAG\|C\|CCCCCCACTCCAGTCCTTCCCCAAGGATGCAGCCTCAGCCTTCTCCACACC |
| 1:1204:21244:89461#1#F | \| \| |
| MQ=60 isize=131 C=27 | GCCCAGTCCCCACACCTACAAGGCCAGCAGATCCCTAATTCTCTCTCCAATCAAGTGCGCTCTCCCCAGCCTGTCCCTTCTCCACGGCCACAGTCCCAG\|C\|ccCCCCACTCCAGTCCTTCCCCAAGGATGCAGGACTcCAATATGCCGACCG |
| 1:1205:1566:29610#1#F | \| \| |
| MQ=60 isize=0 C=37 | GCCCAGTCCCCACACCTACAAGGCCAGCAGATCCCTAATTCTCTCTCCAATCAAGTGCGCTCTCCCCAGCCTGTCCCTTCTCCACGGCCACAGTCCCAG\|C\|CCCCCCACTCCAGTCCTTCCCCAAGGATGCAGCCTCAGCCTTCTCCACACC |
| 1:1205:10478:70361#1#F | \| \| |
| MQ=60 isize=301 C=37 | GCCCAGTCCCCACACCTACAAGGCCAGCAGATCCCTAATTCTCTCTCCAATCAAGTGCGCTCTCCCCAGCCTGTCCCTTCTCCACGGCCACAGTCCCAG\|C\|CCCCCCACTCCAGTCCTTCCCCAAGGATGCAGCCTCAGCCTTCtCCACACC |
| 1:1205:20547:95230#1#F | \| \| |
| MQ=60 isize=293 C=37 | GCCCAGTCCCCACACCTACAAGGCCAGCAGATCCCTAATTCTCTCTCCAATCAAGTGCGCTCTCCCCAGCCTGTCCCTTCTCCACGGCCACAGTCCCAG\|C\|CCCCCCACTCCAGTCCTTCCCCAAGGATGCAGCCTCAGCCTTCTCCACACC |
| 1:1206:3736:17198#1#F | \| \| |
| MQ=60 isize=200 C=33 | GCCCAGTCCCCACACCTACAAGGCCAGCAGATCCCTAATTCTCTCTCCAATCAAGTGCGCTCTCCCCAGCCTGTCCCTTCTCCACGGCCACAGTCCCAG\|C\|CCCCCCACTCCAGTCCTTCCCCAAGGatGCAgcCtCAGCCTTCTCCAcacc |
| 1:1206:18816:32508#1#F | \| \| |
| MQ=60 isize=237 C=33 | GCCCAGTCCCCACACCTACAAGGCCAGCAGATCCCTAATTCTCTCTCCAATCAAGTGCGCTCTCCCCAGCCTGTCCCTTCTCCACGGCCACAGTCCCAG\|C\|CCCCCCACTCCAGTCCTTCCCCAAGGATGCAGCCTCAGCCTTCTCCACACC |
| 1:1206:16166:82486#1#F | \| \| |
| MQ=60 isize=140 C=37 | GCCCAGTCCCCACACCTACAAGGCCAGCAGATCCCTAATTCTCTCTCCAATCAAGTGCGCTCTCCCCAGCCTGTCCCTTCTCCACGGCCACAGtCCCAG\|C\|CCCCCCACTCCAGTCCTTCCCCAAGGATGCAGCCTCAGCCAGGACTCCAAa |
| 1:1207:7126:21008#1#F | \| \| |
| MQ=60 isize=175 C=22 | GCCCAGTCCCCACACCTACAAGGCCAGCAGATCCCTAATTCTCTCTCCAATCAAGTGCGCTCTCCCCAGCCTGTCCCTTCTCCACGGCCACAGTCCCAG\|C\|CCCCCCACTCCAGTCCTTCCCCAAgGATGCAGCCTCacccTTCTCCacacc |
| 1:1207:16923:67258#1#F | \| \| |
| MQ=60 isize=175 C=37 | GCCCAGTCCCCACACCTACAAGGCCAGCAGATCCCTAATTCTCTCTCCAATCAAGTGCGCTCTCCCCAGCCTGTCCCTTCTCCACGGCCACAGTCCCAG\|C\|CCCCCCACTCCAGTCCTTCCCCAAGGATGCAGCCtCAGCCTTCTCCACACC |
| 1:1208:12614:29909#1#F | \| \| |
| MQ=60 isize=232 C=15 | GCCCAGTCCCCACACCTACAAGGCCAGCAGATCCCTAATTCTCTCTCCAATCAAGTGCGCTCTCCCCAGCCTGTCCCTTCTCCACGGCCACAGTCCCAG\|c\|CCCCCCACTCCAGTCCTTCCCCAAgGATGCAgCCTCAgCCTTctCCccacc |
| 1:1208:17039:51193#1#F | \| \| |
| MQ=60 isize=162 C=33 | GCCCAGTCCCCACACCTACAAGGCCAGCAGATCCCTAATTCTCTCTCCAATCAAGTGCGCTCTCCCCAgCCTGTCCCTTCTCCACGGCCACAGTCCCAG\|C\|CCCCCCACTCCagtCCTTCCCCAAGgaTGCAGCCtCAGCCTTctccacacc |
| 1:1208:2789:73373#1#F | \| \| |
| MQ=60 isize=177 C=33 | GCCCAGTCCCCACACCTACAAGGCCAGCAGATCCCTAATTCTCTCTCCAATCAAGTGCGCTCTCCCCAGCCTGTCCCTTCTCCACGGCCACAGTCCCAG\|C\|CCCCCCACTCCAGTCCTTCCCCaAGGATGCAGCCTCAGCCTTCtCCACACC |
| 1:1209:3890:6590#1#F | \| \| |
| MQ=60 isize=265 C=33 | GCCCAGTCCCCACACCTACAAGGCCAGCAGATCCCTAATTCTCTCTCCAATCAAGTGCGCTCTCCCCAGCCTGTCCCTTCTCCACGGCCACAGTCCCAG\|C\|CCCCCCACTCCAGTCCTTCCCCAAGGATGCAGCCTCAGCCTTCTCCACACC |
| 1:1209:12891:40925#1#F | \| \| |
| MQ=60 isize=213 C=37 | GCCCAGTCCCCACACCTACAAGGCCAGCAGATCCCTAATTCTCTCTCCAATCAAGTGCGCTCTCCCCAGCCTGTCCCTTCTCCACGGCCACAGTCCCAG\|C\|CCCCCCACTCCAGTCCTTCCCCAAGGATGCAGCCtCAGCCTTCTCCacacc |
| 1:1209:4459:56572#1#F | \| \| |
| MQ=60 isize=168 C=33 | GCCCAGTCCCCACACCTACAAGGCCAGCAGATCCCTAATTCTCTCTCCAATCAAGTGCGCTCTCCCCAGCCTGTCCCTTCTCCACGGCCACAGTCCCAG\|C\|CCCCCCACTCCAGTCCTTCCCCAAGGATGCAGCCTCAGCCTTCTCCACACC |
| 1:1210:3414:7863#1#F | \| \| |
| MQ=60 isize=186 C=33 | GCCCAGTCCCCACACCTACAAGGCCAGCAGATCCCTAATTCTCTCTCCAATCAAGTGCGCTCTCCCCAGCCTGTCCCTTCTCCACGGCCACAGTCCCAG\|C\|CCCCCCACTCCAGTCCTTCCCCAaGGATGCAGCCTCAGCCTTCTCCACACC |
| 1:1210:5125:39611#1#F | \| \| |
| MQ=60 isize=192 C=37 | GCCCAGTCCCCACACCTACAAGGCCAGCAGATCCCTAATTCTCTCTCCAATCAAGTGCGCTCTCCCCAGCCTGTCCCTTCTCCACGGCCACAGTCCCAG\|C\|CCCCCCACTCCAGTCCTTCCCCAAGGATGCAGCCTCAGCCTTCTCCACGCC |
| 1:1210:3449:74135#1#F | \| \| |
| MQ=60 isize=165 C=27 | GCCCAGTCCCCACACCTACAAGGCCAGCAGATCCCTAATTCTCTCTCCAATCAAGTGCGCTCTCCCCAGCCTGTCCCTTCTCCACGGCCACAGTcccAG\|C\|CCCCCCACTCCaGTCCtTCCCCAAGGATGCaGCCTcAGcCTTCTcCACaCC |
| 1:1210:14809:92261#1#F | \| \| |
| MQ=60 isize=125 C=27 | GCCCAGTCCCCACACCTACAAGGCCAGCAGATCCCTAATTCTCTCTCCAATCAAGTGCGCTCTCCCCAGCCTGTCCCTTCTCCACGGCCACAGTCCCAG\|C\|CcCCCCACTCCAGTCCTTCCCCAAgacTCCAATCAtAGCCCTTTTctgtct |
| 1:1211:10917:16029#1#F | \| \| |
| MQ=60 isize=149 C=15 | ccCCAgTCCcCACACCTACAAGGCCAGCAGAtCCCTAAtTCTCTCTCCAATCAAGTGCGCTCTCCCCAGCCTGTcCCTTCTCCACGGcCaCagtCCCcg\|c\|CCCCCCACTCCAgtCCTTCCCCaaggatgcagcctcagccttctccccagg |
| 1:1211:9249:36802#1#F | \| \| |
| MQ=60 isize=150 C=33 | GCCCAGTCCCCACACCTACAAGGCCAGCAGATCCCTAATTCTCTCTCCAATCAAGTGCGCTCTCCCCAGCCTGTCCCTTCTCCACGGCCACAGTCCCAG\|C\|CCCCCCACTCCAgTCCTTCCCCAAggATGCAGCCtCAGCCTTCTCCaCACA |
| 1:1211:7340:63199#1#F | \| \| |
| MQ=60 isize=147 C=33 | GCCCAGTCCCCACACCTACAAGGCCAGCAGATCCCTAATTCTCTCTCCAATCAAGTGCGCTCTCCCCAGCCTGTCCCTTCTCCACGGCCACAGTCCCAG\|C\|CCCCCCACTCCAGTCCTTCCCCAAGGATGCAGCCTCAGCCTTCTCCAGGAC |
| 1:1211:19212:72638#1#F | \| \| |
| MQ=60 isize=133 C=33 | GCCCAGTCTCCACACCTACAAGGCCAGCAGATCCCTAATTCTCTCTCCAATCAAGTGCGCTCTCCCCAGCCTGTCCCTTCTCCACGGCCACAGTCCCAG\|C\|CCCCCCACTCCAGTCCTTCCCCAAGGATGCAGCAGGAcTCCAATAATCaCa |
| 1:1212:2413:9331#1#F | \| \| |
| MQ=60 isize=209 C=33 | GCCCAGTCCCCACACCTACAAGGCCAGCAGATCCCTAATTCTCTCTCCAATCAAGTGCGCTCTCCCCAGCCTGTCCCTTCTCCACGGCCACAGTCCCAG\|C\|CCCCCCACTCCAGTCCTTCCCCAAGGATGCAGCCTCAGCCTTCTCCaCACC |
| 1:1212:5369:72433#1#F | \| \| |
| MQ=60 isize=205 C=33 | GCCCAGTCCCCACACCTACAAGGCCAGCAGATCCCTAATTCTCTCTCCAATCAAGTGCgCTCTCCCCAGcCTGTCCCTTCTCCACGGCCACAGTCCCAG\|C\|CCCCCCACTCCAGTCCTTCCCCAAGGATGCAGCCTCAGCCtTCTCCACACC |
| 1:1212:10743:96725#1#F | \| \| |
| MQ=60 isize=184 C=33 | GCCCAGTCCCCACACCTACAAGGCCAGCAGATCCCTAATTCTCTCTCCAATCAAGTGCGCTCTCCCCAGCCTGTCCCTTCTCCACGGCCACAGTCCCAG\|C\|CCCCCCACTCCAGTCCTTCCCCAAGGAtGCAGCCTCAGCCTTCTCcacacc |
| 1:1213:17709:29605#1#F | \| \| |
| MQ=60 isize=144 C=33 | GCCCAGTCCCCACACCTACAAGGCCAGCAGATCCCTAATTCTCTCTCCAATCAAGTGCGCTCTCCCCAGCCTGTCCCTTCTCCACGGCCACAGTCCCAG\|C\|CCCCCCACTCCAGTCCTTCCCCAAGGATGCAGCCTCAgCCTTCTAGGcctC |
| 1:1213:8499:91112#1#F | \| \| |
| MQ=60 isize=271 C=33 | gCCCAGTCCCCACACCTACAAGGCCAGCAGATCCCTAATTcTCTCTCCAATCAAGTGCGCTCTCCCCAGCCTGTCcCTTCTCCACgGCCACAGTCCCAG\|C\|CCCcCCACTCCaGTCCTTCCCCAAGGATgCAGCCTCAGCCTTCTCcaCACC |
| 1:1214:12173:39460#1#F | \| \| |
| MQ=60 isize=162 C=37 | GCCCAGTCCCCACACCTACAAGGCCAGCAGATCCCTAATTCTCTCTCCAATCAAGTGCGCTCTCCCCAGCCTGTCCCTTCTCCACGGCCACAGTCCCAG\|C\|CCCCCCACTCCAGTCCTTCCCCAAGGATGCAGCCTCAGCCTTCTCCACACC |
| 1:1214:2931:58360#1#F | \| \| |
| MQ=60 isize=187 C=33 | GCCCAGTCCCCACACCTACAAGGCCAGCAGATCCCTAATTCTCTCTCCAATCAAGTGCGCTCTCCCCAGCCTGTCCCTTCTCCACGGCCACAGTCCCAG\|C\|CCCCCCACTCCAGTCCTTCCCCAAGGATGCAGCCTCAGCCTTCTCCACACC |
| 1:1214:16634:96231#1#F | \| \| |
| MQ=60 isize=125 C=33 | GCCCAGTCCCCACACCTACAAGGCCAGCAGATcCCTAaTTCTCTCTCCAATCAAGTGCGCTCTCCCCAgCCTGTCCCTTCTCCACGGCCACAGTCCCAg\|C\|CCCCCCaCtCCAGtCCTTCCCCAagaggactccaattctcaggtgtcgctg |
| 1:1215:14185:13877#1#F | \| \| |
| MQ=60 isize=149 C=6 | gCCCAGTCCCCACACCTACAAGGCCAGCaGaTCCCTAATTCTCTCTCCAaTCccGtGCgCtCtCCCCagCCTGTCCCTTctCCACGgCCACAGTCCCAg\|c\|CCCCCCACtCCagtcctTCCCCAaggAtgCagCCTCAgCCtTctcCACAgg |
| 1:1215:1874:38297#1#F | \| \| |
| MQ=60 isize=120 C=33 | GCCCAGTCCCCACACCTACAAGGCCAGCAGATCCCTAATTCTCTCTCCAATCAAGTGCGCTCTCCCCAGCCTGTCCCTTCTCCACGGCCACAGTCCCAG\|C\|CCCCCCACTCCAGTCCTTCCAGGGCTCCAATCTTATCGCCCTCCTGTCTCT |
| 1:1215:9271:82839#1#F | \| \| |
| MQ=60 isize=261 C=33 | GCCCAGTCCCCACACCTACAAGGCCAGCAGATCCCTAATTCTCTCTCCAATCAAGTGCGCTCTCCCCAGCCTGTCCCTTCTCCACGGCCACAGTCCCAG\|C\|CCCCCCACTCCAGTCCTTCCCCAAGGATGCAGCCTCAGCCTTCTCCACACC |
| 1:1216:14921:28727#1#F | \| \| |
| MQ=60 isize=251 C=33 | GCCCAGTCCCCACACCTACAAGGCCAGCAGATCCCTAATTCTCTCTCCAATCAAGTGCGCTCTCCCCAGCCTGTCCCTTCTCCACGGCCACAGTCCCAG\|C\|CCCCCCACTCCAGTCCTTCCCCAAGGATGCAGCCTCAGCCTTCTCCACACC |
| 1:1216:13164:38101#1#F | \| \| |
| MQ=60 isize=245 C=33 | GCCCAGTCCCCACACCTACAAGGCCAGCAGATCCCTAATTCTCTCTCCAATCAAGtGCGCTCTCCCCAGCCTGTCCCTTCTCCACGGCCACAGTCCCAG\|C\|CCCCCCACTCCAGTCCTTCCCCAAGGATGCAGCCTCAGCCTTCTCCACACC |
| 1:1216:15476:49577#1#F | \| \| |
| MQ=60 isize=0 C=33 | GCCCAGTCCCCACACCTACAAGGCCAGCAGATCCCTAATTCTCTCTCCAATCAAGTGCGCTCTCCCCAGCCTGTCCCTTCTCCACGGCCACAGTCCCAG\|C\|CCCCCCACTCCAGtCCTTCCCCAAGGATGCAGCCTCagCCTTCTCCACACC |
| 1:1216:18265:69190#1#F | \| \| |
| MQ=60 isize=140 C=33 | GCCCAGTCCCCACACCTACAAGGCCAGCAGATCCCTAATTCTCTCTCCAATCAAGTGCGCTCTCCCCAGCCTGTCCCTTCTCCACGGCCACAGTCCCAG\|C\|CCCCCCaCtCCaGtCCTTCCCCAAGGATGCAgCCtcAccCAggactCCAAc |
| 1:1216:5632:91031#1#F | \| \| |
| MQ=60 isize=201 C=22 | GCCCAGTCCCCACACCTACAAGGCCAGCAGATcccTAATTCTCTCtCCAATCAAGTGCGCTCTCCCCAGCCTGTCCCTTCTCCACGGCCACAGTCCCAg\|C\|CcccCCactCCagTCCTTccCCaAggaTgCcGCCTCAgccttctccccacc |
| 1:2101:5981:41311#1#F | \| \| |
| MQ=60 isize=0 C=37 | GCCCAGTCCCCACACCTACAAGGCCAGCAGATCCCTAATTCTCTCTCCAATCAAGTGCGCTCTCCCCAGCCTGTCCCTTCTCCACGGCCACAGTCCCAG\|C\|CCCCCCACTCCAGTCCTTCCCCAAGGATGCATATTTAATAGAGAATTCAGt |
| 1:2101:10390:54073#1#F | \| \| |
| MQ=60 isize=187 C=37 | GCCCAGTCCCCACACCTACAAGGCCAGCAGATCCCTAATTCTCTCTCCAATCAAGTGCGCTCTCCCCAGCCTGTCCCTTCTCCACGGCCACAGTCCCAG\|C\|CCCCCCACTCCAGTCCTTCCCCAAGGATGCAGCCTCAGCCTTCTCCACACC |
| 1:2101:13358:57553#1#F | \| \| |
| MQ=60 isize=233 C=37 | GCCCAGTCCCCACACCTACAAGGCCAGCAGATCCCTAATTCTCTCTCCAATCAAGTGCGCTCTCCCCAGCCTGTCCCTTCTCCACGGCCACAGTCCCAG\|C\|CCCCCCACTCCAGTCCTTCCCCAAGGATGCAGCCTCAGCCTTCTCCACACC |
| 1:2101:12799:60988#1#F | \| \| |
| MQ=60 isize=138 C=33 | GCCCAGTCCCCACACCTACAAGGCCAGCAGATCCCTAATTCTCTCTCCAATCAAGTGCGCTCTCCCCAgCCTGTCCCTTCTCCACGGCCACAGTCCCAG\|C\|CCCCCCACTCCAGTCCTTCCCCAAGGATGCAGCCTCAGAGGACTCCAATAT |
| 1:2101:4845:71047#1#F | \| \| |
| MQ=60 isize=144 C=33 | GCCCAGTCCCCACACCTACAAGGCCAGCAGATCCCTAATTCTCTCTCCAATCAAGTGCGCTCTCCCCAGCCTGTCCCTTCTCCACGGCCACAGtCCCAG\|C\|CCCCCCaCTCCAGTCCTTCCCCAAGGATGCAGCCTCAGCCTTCTAGGACTC |
| 1:2102:13135:3007#1#F | \| \| |
| MQ=60 isize=138 C=37 | GCCCAGTCCCCACACCTACAAGGCCAGCAGATCCCTAATTCTCTCTCCAATCAAGTGCGCTCTCCCCAGCCTGTCCCTTCTCCACGGCCACAGTCCCAG\|C\|CCCCCCACTCCAGTCCTTCCCCAAGGATGCAGCCTCAGAGGACTCCAATGT |
| 1:2102:14230:13420#1#F | \| \| |
| MQ=60 isize=149 C=37 | GCCCAGTCCCCACACCTACAAGGCCAGCAGATCCCTAATTCTCTCTCCAATCAAGTGCGCTCTCCCCAGCCTGTCCCTTCTCCACGGCCACAGTCCCAG\|C\|CCCCCCACTCCAGTCCTTCCCCAAGGATGCAGCCTCAGCCTTCTCCACAGG |
| 1:2102:4818:38451#1#F | \| \| |
| MQ=60 isize=332 C=33 | GCCCAGTCCCCACACCTACAAGGCCAGCAGATCCCTAATTCTCTCTCCAATCAAGTGCGCTCTCCCCAGCCTGTCCCTTCTCCACGGCCACAGTCCCAG\|C\|CCCCCCACTCCAGTCCTTCCCCaAGGATGCAGCCTCAGCCTTCCCCACACC |
| 1:2102:19464:55676#1#F | \| \| |
| MQ=60 isize=127 C=37 | GCCCAGTCCCCACACCTACAAGGCCAGCGGATCCCTAATCCTCTCTCCAATCAAGTGCGCTCTCCCCAGCCTGTCCCTTCTCCACGGCCACAGTCCCAG\|C\|CCCCCCACTCCAGTCCTTCCCCAAGGACTCCAATGCCTTCCCCGGCCTGTC |
| 1:2102:5443:73863#1#F | \| \| |
| MQ=60 isize=175 C=37 | GCCCAGTCCCCACACCTACAAGGCCAGCAGATCCCTAATTCTCTCTCCAATCAAGTGCGCTCTCCCCAGCCTGTCCCTTCTCCACGGCCACAGTCCCAG\|C\|CCCCCCACTCCAGTCCTTCCCCAAGGATGCAGCCTCAGCCTTCTCCACACC |
| 1:2102:15692:94492#1#F | \| \| |
| MQ=60 isize=124 C=37 | GCCCAGTCCCCACACCTACAAGGCCAGCAGATCCCTAATTCTCTCTCCAATCAAGTGCGCTCTCCCCAGCCTGTCCCTTCTCCACGGCCACAGTCCCAG\|C\|CCCCCCACTCCAGTCCTTCCCCAAAGGACTCCAATACAGCGCACCTCCTGT |
| 1:2103:12820:16109#1#F | \| \| |
| MQ=60 isize=269 C=37 | GCCCAGTCCCCACACCTACAAGGCCAGCAGATCCCTAATTCTCTCTCCAATCAAGTGCGCTCTCCCCAGCCTGTCCCTTCTCCACGGCCACAGTCCCAG\|C\|CCCCCCACTCCAGTCCTTCCCCAAGGATGCAGCCTCAGCCTTCTCCACACC |
| 1:2103:3124:29303#1#F | \| \| |
| MQ=60 isize=147 C=33 | GCCCAGTCCCCACACCTACAAGGCCAGCAGATCCCTAATTCTCTCTCCAATCAAGTGCGCTCTCCCCAGCCTGTCCCTTCTCCACGGCCACAGTCCCAG\|C\|CCCCCCACTCCAGTCCTTCCCCAAGGATGCAGCCTCAGCCTTCTCCAGGAC |
| 1:2103:17384:56711#1#F | \| \| |
| MQ=60 isize=262 C=33 | GCCCAGTCCCCACACCTACAAGGCCAGCAGATCCCTAATTCTCTCTCCAATCAAGTGCGCTCTCCCCAGCCTGTCCCTTCTCCACGGCCACAGTCCCAG\|C\|CCCCCCACTCCAGTCCTTCCCCAAGGATGCAGCCTCAGCCTTCTCCACACC |
| 1:2103:20374:89239#1#F | \| \| |
| MQ=60 isize=0 C=33 | GCCCAGTCCCCACACCTACAAGGCCAGCAGATCCCTAATTCTCTCTCCAATCAAGTGCGCTCTCCCCAGCCTGTCCCTTCTCCACggccACAGTCCCAG\|C\|CCCCCCACTCCAgtCCTTCCCCAAggAtGCAGCCTCAGCCTTCTCCACAcC |
| 1:2104:21036:19617#1#F | \| \| |
| MQ=60 isize=177 C=37 | GCCCAGTCCCCACACCTACAAGGCCAGCAGATCCCTAATTCTCTCTCCAATCAAGTGCGCTCTCCCCAGCCTGTCCCTTCTCCACGGCCACAGTCCCAG\|C\|CCCCCCACTCCAGTCCTTCCCCAAGGATGCAGCCTCAGCCTTCTCCACACC |
| 1:2104:7285:72809#1#F | \| \| |
| MQ=60 isize=188 C=33 | GCCCAGTCCCCACACCTACAAGGCCAGCAGATCCCTAATTCTCTCTCCAATCAAGTGCGCTCTCCCCAGCCTGTCCCTTCTCCACGGCCACAGTCCCAG\|C\|CCCCCCACTCCAGTCCTTCCCCAAGGATGCAGCCTCAGCCTTCTCCACACC |
| 1:2105:5993:31720#1#F | \| \| |
| MQ=60 isize=302 C=33 | GCCCAGTCCCCACACCTACAAGGCCAGCAGATCCCTAATTCTCTCTCCAATCAAGTGCGCTCTCCCCAGCCTGTCCCTTCTCCACGGCCACAGTCCCAG\|C\|CCCCCCACTCCAGTCCTTCCCCAAGGATGCAGCCTCAGCCTTCTCCACACC |
| 1:2106:8805:2848#1#F | \| \| |
| MQ=60 isize=169 C=33 | GCCCAGTCCCCACACCTACAAGGCCAGCAGATCCCTgaTTCTCTCTCCAATCAAGTGCGCTCTCCCCAGCCTGTCCCTTCTCCACGGCCACAGTCCCAG\|C\|CCCCCCACTCCAGTCCTTCCCCAAGGATGCAGCCTCAGCCTTCTCCACACC |
| 1:2106:18569:25723#1#F | \| \| |
| MQ=60 isize=186 C=33 | GCCCAGTCCCCACACCTACAAGGCCAGCAGATCCCTAATTCTCTCTCCAATCAGGTGCGCTCTCCCCAGCCTGTCCCTTCTCCACGGCCACAGTCCCAG\|C\|CCCCCCaCTCCAGTCCTTCCCCAAGGATGCAGCCTCAgcctTcTCCACACC |
| 1:2106:16704:40375#1#F | \| \| |
| MQ=60 isize=159 C=37 | GCCCAGTCCCCACACCTACAAGGCCAGCAGATCCCTgaTTCTCTCTCCAATCAAGTGCGCTCTCCCCAGCCTGTCCCTTCTCCACGGCCACAGTCCCAG\|C\|CCCCCCACTCCAGTCCTTCCCCAAGGATGCAGCCTCAGCCTTCTCCACACC |
| 1:2106:18685:66837#1#F | \| \| |
| MQ=60 isize=271 C=37 | GCCCAGTCCCCACACCTACAAGGCCAGCAGATCCCTAATTCTCTCTCCAATCAAGTGCGCTCTCCCCAGCCTGTCCCTTCTCCACGGCCACAGTCCCAG\|C\|CCCCCCACTCCAGTCCTTCCCCAAGGATGCAGCCTCAGCCTTCTCCACACC |
| 1:2106:3939:97521#1#F | \| \| |
| MQ=60 isize=0 C=37 | GCCCAGTCCCCACACCTACAAGGCCAGCAGATCCCTAATTCTCTCTCCAATCAAGTGCGCTCTCCCCAGCCTGTCCCTTCTCCACGGCCACAGTCCCAG\|C\|CCCCCCACTCCAGTCCTTCCCCAAGGATGCAGCCTCAGCCTTCTCCTTTTC |
| 1:2107:2273:16254#1#F | \| \| |
| MQ=60 isize=225 C=37 | GCCCAGTCCCCACACCTACAAGGCCAGCAGATCCCTAATTCTCTCTCCAATCAAGTGCGCTCTCCCCAGCCTGTcCCTTCTCCACGGCCACAGTCCCAG\|C\|CCCCCCACTCCAGTCCTTCCCCAAGGATGCAGCCTCAGCCTCCTCCACACC |
| 1:2107:2014:22522#1#F | \| \| |
| MQ=60 isize=209 C=37 | GCCCAGTCCCCACACCTACAAGGCCAGCAGATCCCTGATTCTCTCTCCAATCAAGTGCGCTCTCCCCAGCCTGTCCCTTCTCCACGGCCACAGTCCCAG\|C\|CCCCCCACTCCAGTCCTTCCCCAAGGATGCAGCCTCAGCCTTCTCCACACC |
| 1:2107:3050:54372#1#F | \| \| |
| MQ=60 isize=150 C=33 | GCCCAGTCCCCACACCTACAAGGCCAGCAGATCCCTAATTCTCTCTCCAATCAAGTGCGCTCTCCCCAGCCTGTCCCTTCTCCACGGCCACAGTCCCAG\|C\|CCCCCCACTCCcgtCCTTCCCCaaGgatGCaGCCTCAGCCTtcTCCACACc |
| 1:2107:8164:64341#1#F | \| \| |
| MQ=60 isize=132 C=37 | GCCCAGTCCCCACACCTACAAGGCCAGCAGATCCCTAATTCTCTCTCCAATCAAGTGCGCTCTCCCCAGCCTGTCCCTTCTCCACGGCCACAGTCCCAG\|C\|CCCCCCACTCCAGTCCTTCCCCAAGGATGCAGGACTCCAATCCTTCACCGT |
| 1:2107:16901:78282#1#F | \| \| |
| MQ=60 isize=175 C=33 | GCCCAGTCCCCACACCTACAAGGCCAGCAGATCCCTAATTCTCTCTCCAATCAAGTGCGCTCTCCCCAGCCTGTCCCTTCTCCACGGCCACAGTCCCAG\|C\|CCCCCCACTCCAGTCCTTCCCCAAGGATGCAGCCTCaGCCTTCTCCACACC |
| 1:2108:9782:3529#1#F | \| \| |
| MQ=60 isize=287 C=33 | GCCCAGTCCCCACACCTACAAGGCCAGCAGATCCCTAATTCTCTCTCCAATCAAGtGCGCTCtCCCCAGCCTGTCCCTTCTCCACGGCCACAGTCCCAG\|C\|CCCCCCACTCCAGTCCTTCCCCAAGGATGCAGCCTCAGCCTTCTCCACACC |
| 1:2108:20211:38299#1#F | \| \| |
| MQ=60 isize=0 C=33 | GCCCAGTCCCCACACCTACAAGGCCAGCAGATCCCTAATTCTCTCTCCAATCAAGTGCGCTCTCCCCAGCCTGTCCCTTCTCCACGGCCACAGTCCCAG\|C\|CCCCCCACTCCAGTCCTTCCCCAAGGATGCAGCCTCAGGATAACTAGAAGA |
| 1:2108:8927:80065#1#F | \| \| |
| MQ=60 isize=165 C=37 | GCCCAGTCCCCACACCTACAAGGCCAGCAGATCCCTAATTCTCTCTCCAATCAAGTGCGCTCTCCCCAGCCTGTCCCTTCTCCACGGCCACAGTCCCAG\|C\|CCCCCCACTCCAGTCCTTCCCCAAGGATGCAGCCTCAGCCTTCTCCACACC |
| 1:2108:7286:98637#1#F | \| \| |
| MQ=60 isize=195 C=37 | GCCCAGTCCCCACACCTACAAGGCCAGCAGATCCCTAATTCTCTCTCCAATCAAGTGCGCTCTCCCCAGCCTGTCCCTTCTCCACGGCCACAGTCCCAG\|C\|CCCCCCACTCCAGTCCTTCCCCAAGGATGCAGCCTCAGCCTTCTCCACACC |
| 1:2108:14011:100407#1#F | \| \| |
| MQ=60 isize=186 C=27 | GCCCAGTCCCCACACCTACAAGGCCAGCAGATCCCTAATTCTCTCTCCAATCAAGTGCGCTCTCCCCAGCCTGTCCCTTCTCCACGGCCACAGTCCCAg\|C\|CCCCCCactCCaGtCCTTCCCCAAGGATGCAGCCTCAGCCTTCTCCACACC |
| 1:2109:2042:15172#1#F | \| \| |
| MQ=60 isize=185 C=37 | GCCCAGTCCCCACACCTACAAGGCCAGCAGATCCCTAATTCTCTCTCCAATCAAGTGCGCTCTCCCCAGCCTGTCCCTTCTCCACGGCCACAGTCCCAG\|C\|CCCCCCACTCCAGTCCTTCCCCAAGGATGCAGCCTCAGCCTTCTCCACACC |
| 1:2109:9717:23748#1#F | \| \| |
| MQ=60 isize=164 C=33 | GCCCAGTCCCCACACCTACAAGGCCAGCAGATCCCTAATTCTCTCTCCAATCAAGTGCGCTCTCCCCAGCCTGTCCCTTCTCCACGGCCACAGTCCCAG\|C\|CCCCCCACTCCAGTCCTTCCCCAAGGATGCAGCCTCAGCCTTCTCCACACC |
| 1:2109:13352:58370#1#F | \| \| |
| MQ=60 isize=208 C=33 | GCCCAGTCCCCACACCTACAAGGCCAGCAGATCCCTAATTCTCTCTCCAATCAAGTGCGCTCTCCCCAGCCTGTCCCTTCTCCACGGCCACAGTCCCAG\|C\|CCCCCCACTCCAGTCCTTCCCCAAGGATGCAGCCTCAGCCTTCTCCACACC |
| 1:2109:20196:67243#1#F | \| \| |
| MQ=60 isize=176 C=33 | GCCCAGTCCCCACACCTACAAGGCCAGCAGATCCCTAATTCTCTCTCCAATCAAGTGCGCTCTCCCCAGCCTGTCCCTTCTCCACGGCCACAGTCCCAG\|C\|CCCCCCACTCCAGTCCTTCCCCAAGGATGCAGCCTCAGCCTTCTCCACACC |
| 1:2109:15750:80470#1#F | \| \| |
| MQ=60 isize=187 C=37 | GCCCAGTCCCCACACCTACAAGGCCAGCAGATCCCTAATTCTCTCTCCAATCAAGTGCGCTCTCCCCAGCCTGTCCCTTCTCCACGGCCACAGTCCCAG\|C\|CCCCCCACTCCAGTCCTTCCCCAAGGATGCAGCCTCAGCCTTCTCCACACC |
| 1:2109:10990:89878#1#F | \| \| |
| MQ=60 isize=140 C=37 | GCCCAGTCCCCACACCTACAAGGCCAGCAGATCCCTAATTCTCTCTCCAATCAAGTGCGCTCTCCCCAGCCTGTCCCTTCTCCACGGCCACAGTCCCAG\|C\|CCCCCCACTCCAGTCCTTCCCCAAGGATGCAGCCTCAGCCAGGACTCCAAT |
| 1:2110:4609:63254#1#F | \| \| |
| MQ=60 isize=210 C=33 | GcCCAGTCCCCACACCTACAAGGCCAGCAGATCCCTAATTCTCTCTCCAATCAAGCGCGCTCTCCCCAGCCTGTCCCTTCTCCACgGCCACAGTCCCAG\|C\|CCCCCCACTCCAGTCCTTCCCCAAGGATGCAgcCtcagcctTCTCCACACC |
| 1:2110:4075:85946#1#F | \| \| |
| MQ=60 isize=125 C=33 | GCCCAGTCCCCACACCTACAAGGCCAGCAGATCCCTAATTCTCTCTCCAATCAAGTGCGCTCTCCCCAGCCTGTCCCTTCTCCACGGCCACAGTCCCAG\|C\|CCCCCCACTCCAGTCCTTCCCCAAGAGGACTCCAATTGTGACATGAGCCTg |
| 1:2111:4201:17202#1#F | \| \| |
| MQ=60 isize=208 C=33 | GCCCAGTCCCCACACCTACAAGGCCAGCAGATCCCTAATTCTCTCTCCAATCAAGTGCGCTCTCCCCAGCCTGTCCCTTCTCCACGGCCACAGTCCCAG\|C\|CCCCCCACTCCAGTCCTTCCCCAAGGATGCAGCCTCAGCCTTCTCCACACC |
| 1:2111:19199:58613#1#F | \| \| |
| MQ=60 isize=187 C=37 | GCCCAGTCCCCACACCTACAAGGCCAGCAGATCCCTAATTCTCTCTCCAATCAAGTGCGCTCTCCCCAGCCTGTCCCTTCTCCACGGCCACAGTCCCAG\|C\|CCCCCCACTCCAGTCCTTCCCCAAGGATGCAGCCTCAGCCTTCTCCACACC |
| 1:2112:20171:16565#1#F | \| \| |
| MQ=60 isize=199 C=33 | GCCCAGTCCCCACACCTACAAGGCCAGCAGaTCCCTAATTCTCTCTCCAAtCAAGTGCGCTCTCCCCAGCCTGtCCCTTCTCCACGGCCaCAGTCCCAG\|C\|CCcCCCaCtCCAGTCCTTCCCCAAggATGCagCCtCAGCCTTCTCCACACC |
| 1:2112:16490:35491#1#F | \| \| |
| MQ=60 isize=236 C=33 | GCCCAGTCCCCACACCTACAAGGCCAGCAGATCCCTAATTCTCTCTCCAATCAAGTGCGCTCTCCCCAGCCTGTCCCTTCTCCACGGCCACAGTCCCAG\|C\|CCCCCCACTCCAGTCCTTCCCCAAGGATGCAGCCTCAGCCTTCTCCACACC |
| 1:2112:6053:79145#1#F | \| \| |
| MQ=60 isize=269 C=33 | GCCCAGTCCCCACACCTACAAGGCCAGCAGATCCCTAATTCTCTCTCCAATCAAGTGCGCTCTCCCCAGCCTGTCCCTTCTCCACGGCCACAGTCCCAG\|C\|CCCCCCACTCCAGTCCTTCCCCAAGGATGCAGCCTCAGCCTTCTCCACACC |
| 1:2112:6270:96168#1#F | \| \| |
| MQ=60 isize=125 C=37 | GCCCAGTCCCCACACCTACAAGGCCAGCAGATCCCTAATTCTCTCTCCAATCAAGTGCGCTCTCCCCAGCCTGTCCCTTCTCCACGGCCACAGTCCCAG\|C\|CCCCCCACTCCAGTCCTTCCCCAAGAGGaCTCCAATAGACGGCCTACTCTG |
| 1:2113:6008:16947#1#F | \| \| |
| MQ=60 isize=132 C=33 | GCCaaGTCCCCACACCTACAAGGCCAGCAGATCCCTAATTCTCTCTCCAATCAAGTGCGCTCTCCCCAGCCTGTCCCTTCTCCACGGCCACAGtCCCAG\|C\|CCCCCCaCTCCAGTCCTTCCCCAAGGATGCAGAGGACTCCAATTAGTCAGG |
| 1:2113:3891:57270#1#F | \| \| |
| MQ=60 isize=147 C=22 | gCccAGTCCCCACACCTACAAGGCCaGCAGATCCCTaATTCTCTCTCCAATCAAGTGCGCTCTCCCCAGCCTGTCCCTTCTCCACGGCCACaGtCCCAG\|C\|CcCCCCACTCCaGtCCTTCCCCAAGGATGCAGCCTCAGCCTTCTCcaGGAC |
| 1:2113:17528:93294#1#F | \| \| |
| MQ=60 isize=134 C=33 | GCCCAGTCCCCACACCTACAAGGCCAGCAGATCCCTAATTCTCTCTCCAATCAAGTGCGCTCTCCCCAGCCTGTCCCTTCTCCACGGCCACAGTCCCAG\|C\|CCCCCCACTCCAgTCCTTCCCCAAGGATGCAGCCAGGACTCCAATGGACAG |
| 1:2114:6806:7320#1#F | \| \| |
| MQ=60 isize=159 C=37 | GCCCAGTCCCCACACCTACAAGGCCAGCAGATCCCTAATTCTCTCTCCAATCAAGTGCGCTCTCCCCAGCCTGTCCCTTCTCCACGGCCACAGTCCCAG\|C\|CCCCCCACTCCAGTCCTTCCCCAAGGATGCAGCCTCAGCCTTCTCCACACC |
| 1:2114:3364:10915#1#F | \| \| |
| MQ=60 isize=115 | GCCCAGTCCCCACACCTACAAGGCCAGCAGATCCCTAATTCTCTCTCCAATCAAGTGCGCTCTCCcccccctgccccctctacaccgccacagccaccc\|c\|ccccccacccctcacagctccccaatacccccccgaca |
| 1:2114:10881:57734#1#F | \| \| |
| MQ=60 isize=177 C=33 | GCCCAGTCCCCACACCTACAAGGCCAGCAGATCCCTAATTCTCTCTCCAATCAAGTGCGCTCTCCCCAGCCTGTCCCTTCTCCACGGCCACAGTCCCAG\|C\|CCCCCCACTCCAGTCCTTCCCCAAGGATGCAGCCTCAGCCTTCTCCACACC |
| 1:2114:14497:78941#1#F | \| \| |
| MQ=60 isize=212 C=37 | GCCCAGTCCCCACACCTACAAGGCCAGCAGATCCCTAATTCTCTCTCCAATCAAGTGCGCTCTCCCCAGCCTGTCCCTTCTCCACGGCCACAGTCCCAG\|C\|CCCCCCACTCCAGTCCTTCCCCAAGGATGCaGCCTCAGCCTTCTCCAcAcC |
| 1:2115:10474:12324#1#F | \| \| |
| MQ=60 isize=203 C=37 | GCCCAGTCCCCACACCTACAAGGCCAGCAGATCCCTAATCCTCTCTCCAATCAAGTGCGCCCTCCCCAGCCTGTCCCTTCTCCACGGCCACAGTCCCAG\|C\|CCCCCCACTCCAGTCCTTCCCCAAGGATGCAGCCTCAGCCTTCTCCACACC |
| 1:2115:4779:37669#1#F | \| \| |
| MQ=60 isize=282 C=33 | GCCCAGTCCCCACACCTACAAGGCCAGCAGATCCCTAATTCTCTCTCCAATCAAGTGCGCTCTCCCCAGCCTGTCCCTTCTCCACGGCCACAgtCCCAG\|C\|CCCCCCACTCCAGTCCTTCCCCAAGGATGCAGCCTCAGCCTTCTCCACACC |
| 1:2115:19606:84434#1#F | \| \| |
| MQ=60 isize=280 C=33 | GCCCAGTCCCCACACCTACAAGGCCAGCAGATCCCTAATTCTCTCTCCAATCAAGTGCGCTCTCCCCAGCCTGTCCCTTCTCCACGGCCACAGTCCCAG\|C\|CCCCCCACTCCAGTCCTTCCCCAAGGAtGCAGCCTCAGCCTTCTCCACACC |
| 1:2116:8814:26318#1#F | \| \| |
| MQ=60 isize=133 C=33 | GCCCAGTCCCCACACCTACAAGGCCAGCAGATCCCTAATTCTCTCTCCAATCAAGTGCGCTCTCCCCAGCCTGTCCCTTCTCCACGGCCACAGtCCCAG\|C\|CCCCCCACTCCAGTCCTTCCCCAAGGATGCAGCAGGACTCCAATAAACGGt |
| 1:2116:12881:31382#1#F | \| \| |
| MQ=60 isize=166 C=33 | GCCCAGTCCCCACACCTACAAGGCCAGCAGATCCCTAATTCTCTCTCCAATCAAGTGCGCTCTCCCCAGCCTGTCCCTTCTCCACGGCCACAGTCCCAG\|C\|CCCCCCACTCCAGTCCTTCCCCAAGGATGCAGCCTCAGCCTTCTCCACACC |
| 1:2116:13362:52751#1#F | \| \| |
| MQ=60 isize=187 C=33 | GCCCAGTCCCCACACCTACAAGGCCAGCAGATCCCTAATTCTCTCTCCAATCAAGTGCGCTCTCCCCAGCCTGTCCCTTCTCCACGGCCACAGTCCCAG\|C\|CCCCCCACTCCAGTCCTTCCCCAAGGATGCAGCCTCAGCCTTCTCCACACC |
| 1:2116:18955:64912#1#F | \| \| |
| MQ=60 isize=230 C=33 | GCCCAGTCCCCACACCTACAAGGCCAGCAGATCCCTAATTCTCTCTCCAATCAAGTGCGCTCTCCCCAGCCTGTCCCTTCTCCACGGCCACAGTCCCAG\|C\|CCCCCCACTCCAGTCCTTCCCCAAGGATGCAGCCTCAGCCTTCTCCACACC |
| 1:2116:13284:75555#1#F | \| \| |
| MQ=60 isize=250 C=33 | GCCCAGTCCCCACACCTACAAGGCCAGCAGATCCCTAATTCTCTCTCCAATCAAGTGCGCTCTCCCCAGCCTGTCCCTTCTCCACGGCCACAGTCCCAG\|C\|CCCCCCACTCCAGTCCTTCCCCAAGGATGCAGCCTCAGCCTTCTCCACACC |
| 1:2116:6236:97323#1#F | \| \| |
| MQ=60 isize=240 C=33 | GCCCAGTCCCCACACCTACAAGGCCAGCAGATCCCTAATTCTCTCTCCAATCAAGTGCGCTCTCCCCAGCCTGTCCCTTCTCCACGGCCACAGTCCCAG\|C\|CCCCCCACTCCAGTCCTTCCCCAAGGATGCAGCCTCAGcCtTCTCCACACC |
| 1:2201:5972:50063#1#F | \| \| |
| MQ=60 isize=160 C=33 | GCCCAGTCCCCACACCTACAAGGCCAGCAGATCCCTAATTCTCTCTCCAATCAAGTGCGCTCTCCCCAGCCTGTCCCTTCTCCACGGCCACAGTCCCAG\|C\|CCCCCCACTCCAGTCCTTCCCCAAGGATGCAGCCtCAGCCTTCTCCACACC |
| 1:2201:2256:85458#1#F | \| \| |
| MQ=60 isize=147 C=33 | GCCCAGTCCCCACACCTACAAGGCCAGCAGATCCCTAATTCTCTCTCCAATCAAGTGCGCTCTCCCCAGCCTGTCCCTTCTCCACGGCCACAGTCCCAG\|C\|CCCCCCACTCCAGTCCTTCCCCAAGGATGCAGCCTCAGCCTTCTCCAGGAC |
| 1:2202:5552:23134#1#F | \| \| |
| MQ=60 isize=162 C=33 | GCCCAGTCCCCACACCTACAAGGCCAGCAGATCCCTAATTCTCTCTCCAATCAAGTGCGCTCTCCCCAGCCTGTCCCTTCTCCACGGCCACAGTCCCAG\|C\|CCCCCCACTCCAGTCCTTCCCCAAGGATGCAGCCTCAGCCTTCTCCACACC |
| 1:2202:17756:27179#1#F | \| \| |
| MQ=60 isize=125 C=27 | GCCCAGTCCCCACACCTACAAGGCCAGCAGATCCCTAATTCTCTCTCCAATCAAGTGCGCTCTCCCCAgCCTGTCCCTTCtCCcCGGCCACAgTCCCAG\|C\|CCCCCCaCtCCcgtccttccccaagaggactcccacgccccccgtcccccc |
| 1:2202:11478:37472#1#F | \| \| |
| MQ=60 isize=160 C=15 | GCCCAGTCCCCACACCTACAAGGCCAGCAGATCCCTAATTCTCTCTCCAATCAAGTGCGCTCTCCCCAGCCTGTCCCTTCTCCaCGGCCACAGTCCCAG\|c\|CCCCCCACTCCaGccCTTCCCCAAGGATGCAgCCtcagccttctccacacc |
| 1:2202:14108:64497#1#F | \| \| |
| MQ=60 isize=169 C=33 | GCCCAGTCCCCACACCTACAAGGCCAGCAGATCCCTaaTTCTCTCTCCaATCAAGTGCGCTCTCCCCAGCCTGTCCCTTCTCCaCGGCcccaGTCCCaG\|C\|CcCCCCccTCCAGtCCTTCCCCAagGATGCagCCTCcgCCTtctccacacc |
| 1:2202:1856:69930#1#F | \| \| |
| MQ=60 isize=121 C=33 | GCCCAGTCCCCACACCTACAAGGCCAGCAGATCCCTAATTCTCTCTCCAATCAAGCGCGCTCTCCCCAGCCTGTCCCTTCTCCACGGCCACAGTCCCAG\|C\|CCCCCCACTCCAGTCCTTCCCAGGACTCCAATAGCAAGGAACTACTGTCTc |
| 1:2202:16673:93147#1#F | \| \| |
| MQ=60 isize=181 C=37 | GCCCAGTCCCCACACCTACAAGGCCAGCAGATCCCTAATTCTCTCTCCAATCAAGTGCGCTCTCCCCAGCCTGTCCCTTCTCCACGGCGACAGtCCCAG\|C\|CCCCCCACTCCAGtCCTTCCCCAAGGAtGCAGCctCAgCCTtctCCccacc |
| 1:2203:19305:26523#1#F | \| \| |
| MQ=60 isize=198 C=33 | GCCCAGTCCCCACACCTACAAGGCCAGCAGATCCCTAATTCTCTCTCCAATCAAGTGCGCTCTCCCCAGCCTGTCCCTTCTCCACGGCgaCAGTCCCAG\|C\|CCCCCCACTCCAGTCCTTCCCCAAGGATGCAGCCTCAGCCTTCTCCacacc |
| 1:2203:13700:57432#1#F | \| \| |
| MQ=60 isize=162 C=15 | GCCCAGTCCCCACACCTACAAGGCCAGCAGATCCCTAaTTCTCTCTCCAATCAAGTGCGCTCTCCCCAGCCTGTCCCTTCTCCaCGgCCACAGtCCCAg\|c\|cccCCCACTCcagTCCTTCcccAgGgATGCAgcctaagCCTTCTCCacacc |
| 1:2204:20533:7454#1#F | \| \| |
| MQ=60 isize=175 C=33 | GCCCAGTCCCCACACCTACAAGGCCAGCAGATCCCTAATTCTCTCTCCAATCAAGTGCGCTCTCCCCAGCCTGTCCCTTCTCCACGGCaaCAGTCCCAG\|C\|CCCCCCACTCCAGTCCTTCCCCAAGGATGCAGCCTCAGCCTTCTCCACACC |
| 1:2204:7128:62766#1#F | \| \| |
| MQ=60 isize=147 C=33 | GCCCAGTCCCCACACCTACAAGGCCAGCAGATCCCTAATTCTCTCTCCAATCAAGTGCGCTCTCCCCAGCCTGTCCCTTCTCCACGGCCACAGTCCCAG\|C\|CCCCCCaCTCCAGTCCTTCCCCAaGGaTGCAGCCTCAgCCTTCTCCaggac |
| 1:2204:2435:86546#1#F | \| \| |
| MQ=60 isize=133 C=33 | GCCCAGTCCCCACACCTACAAGGCCAGCAGATCCCTAATTCTCTCTCCAATCAAGTGCGCTCTCCCCAGCCTGTCCCTTCTCCACGGCCACAGTCCCAG\|C\|CCCCCCACTCCAGTCCTTCCCCaaggATGCAGCAGGACTCCAATGTCGGCC |
| 1:2204:11375:91497#1#F | \| \| |
| MQ=60 isize=232 C=37 | GCCCAGTCCCCACACCTACAAGGCCAGCAGATCCCTAATTCTCTCTCCAATCAAGTGCGCTCTCCCCAGCCTGTCCCTTCTCCACGGCCACAGTCCCAG\|C\|CCCCCCACTCCAGTCCTTCCCCAAGGAtGCAGCCTCAGCCTTCTCCACaCC |
| 1:2205:11421:13114#1#F | \| \| |
| MQ=60 isize=305 C=37 | GCCCAGTCCCCACACCTACAAGGCCAGCAGATCCCTAATTCTCTCTCCAATCAAGTGCGCTCTCCCCAGCCTGTCCCTTCTCCACGGCCACAGTCCCAG\|C\|CCCCCCaCTCCAGTCCTTCCCCAaGgATGCAGCCtcAGCCTtctccccacc |
| 1:2205:15776:22028#1#F | \| \| |
| MQ=60 isize=224 C=15 | GCCCAGTCCCCACACCTACAaGGCCAGCAGATcCCTAATTCTCTCTCCAATCAAGTGCGCTCTCCCCAGCCTGTCCCTTCTCCACGGCCACAGTCCCAG\|c\|CcCCCcaCTCCAGtCCTtCCCCAaGGATGcAGCCTCAGCCTTcTCcacACc |
| 1:2205:19728:51791#1#F | \| \| |
| MQ=60 isize=186 C=37 | GCCCAGTCCCCACACCTACAAGGCCAGCAGATCCCTAATTCTCTCTCCAATCAAGTGCGCTCTCCCCAGCCTGTCCCTTCTCCAcggCCACAGTCCCAG\|C\|CCCCCCACTCCAGTCCTTCCCCAAGGATGCAGCCTCAGCCTTCTCCaCACC |
| 1:2205:8921:73651#1#F | \| \| |
| MQ=60 isize=156 C=27 | GCCCAGtcCCCACACCTACAaGGCCAGCAGATCcCTAATtcTCTCTCCAATCAAGTgcGCTcTCCCCAGCCTGTCCCTTcTCCACGgccaCAGtccCaG\|C\|CCCCCcACtccAGTCCTTCCccaaggatgcagcctcagccttccccacgcc |
| 1:2205:5898:91632#1#F | \| \| |
| MQ=60 isize=127 C=37 | GCCCAGTCCCCACACCTACAAGGCCAGCAGATCCCTAATTCTCTCTCCAATCAAGTGCGCTCTCCCCAGCCTGTCCCTTCTCCACGGCCACAGTCCCAG\|C\|CCCCCCACTCCAGTCCTTCCCCAAGGACTCCAATTGACGGCAGGTCCTGTc |
| 1:2206:4164:8382#1#F | \| \| |
| MQ=60 isize=139 C=33 | GCCCAGTCCCCACACCTACAAGGCCAGCAGATCCCTAATTCTCTCTCCAATCAAGTGCGCTCTCCCCAGCCTGTCCCTTCTCCACGGCCACAGTCCCAG\|C\|CCCCCCACTCCAGTCCTTCCCCAAGGATGCAGCCTCAGCAGGACTCCAATC |
| 1:2206:14677:43087#1#F | \| \| |
| MQ=60 isize=162 C=33 | GCCCAGTCCCCACACCTACAAGGCCAGCAGATCCCTAATTCTCTCTCCAATCAAGTGCGCTCTCCCCAGCCTGTCCCTTCTCCACGGCCACAGTCCCAG\|C\|CCCCCcactCCAGTCCTTCCCCAAGgaTGCagCCtCagCcttctccccacc |
| 1:2206:4696:54895#1#F | \| \| |
| MQ=60 isize=144 C=33 | GCCCAGTCCCCACACCTACAAGGCCAGCAGATCCCTAATTCTCTCTCCAATCAAGTGCGCTcTCCCCGGCCTGTCCCTTCTCCACGGCCACAGTCCCAG\|C\|CCCCCCACTCCAGTCCTtCccCAAGGatGCAGCCTCAgCCTTCTAGGACTC |
| 1:2206:8834:72032#1#F | \| \| |
| MQ=60 isize=177 C=37 | GCCCAGTCCCCACACCTACAAGGCCAGCAGATCCCTAATTCTCTCTCCAATCAAGTGCGCTCTCCCCAGCCTGTCCCTTCTCCACGGCCACAGTCCCAG\|C\|CCCCCCACTCCAGTCCTTCCCCAAGGATGCAGCCTCAGCCTTCTCCACACC |
| 1:2206:3082:90242#1#F | \| \| |
| MQ=60 isize=162 C=33 | GCCCAGTCCCCACACCTACAAGGCCAGCAGATCCCTAATTCTCTCTCCAATCAAGTGCGCTCTCCCCAGCCTGTCCCTTCTCCACGGCCACAGTCCCAG\|C\|CCCCCCACTCCAGTCCTTCCCCAAGGATGCAGCCtcagCCTTCTCCACACC |
| 1:2207:6864:26267#1#F | \| \| |
| MQ=60 isize=162 C=37 | GCCCAGTCCCCACACCTACAAGGCCAGCAGATCCCTAATTCTCTCTCCAATCAAGTGCGCTCTCCCCAGCCTGTCCCTTCTCCACGGCCACAGTCCCAG\|C\|CCCCCCACTCCAGTCCTTCCCCAAGGATGCAGCCTCAGCCTTCTCCACACC |
| 1:2207:4629:51801#1#F | \| \| |
| MQ=60 isize=211 C=33 | GCCCAGTCCCCACACCTACAAGGCCAGCAGATCCCTAATTCTCTCTCCAATCAAGTGCGCTCTCCCCAGCCTGTCCCTTCTCCaCGGCcACAGTCCCAG\|C\|CCCCCCACTCCAGTCCTTcCCCAAGGATGCAGCCtCAGCCTTCTCCACACc |
| 1:2207:8259:80185#1#F | \| \| |
| MQ=60 isize=224 C=33 | GCCCAGTCCCCACACCTACAAGGCCAGCAGATCCCTAATTCTCTCTCCAATCAAGTGCGCTCTCCCCAGCCTGTCCCTTCTCCACGGCCACAGTCCCAG\|C\|CCCCCCACTCCAGTCCTTCCCCAAGGATGCAGCCTCAGCCTTCTCCACACC |
| 1:2207:14274:96247#1#F | \| \| |
| MQ=60 isize=175 C=33 | GCCCAGTCCCCACACCTACAAGGCCAGCAGATCCCTAATTCTCTCTCCAATCAAGTGCGCTCTCCCCAGCCTGTCCCTTCTCCACGGCCACAGTCCCAG\|C\|CCCCCCaCTCCAGTCCTTCCCCAAGGATGCAGCCTCAgCCTTCtCCaCACC |
| 1:2208:10791:52716#1#F | \| \| |
| MQ=60 isize=143 C=2 | GCCCAGTCCCCACACCTaCAAGgCcAGcagaTccCtAAtTcTCTCtCcAaTcAagtgCGttctccctagcctgacccttctccacggccacaggcccat\|c\|ccccccactccagtccttccccaaggatgcagcattagcctgcaggactca |
| 1:2208:10848:76326#1#F | \| \| |
| MQ=60 isize=293 C=33 | GCCCAGTCCCCACACCTACAAGGCCAGCAGATCCCTAATTCTCTCTCCAATCAAGTGCGCTCTCCCCAGCCTGTCCCTTCTCCACGGCCACAGTCCCAG\|C\|CCCCCCACTCCAGTCCTTCCCCAAGGATGCAGCCTCAGCCTTCTCCACACC |
| 1:2209:3225:43225#1#F | \| \| |
| MQ=60 isize=205 C=33 | GCCCAGTCCCCACACCTACAAGGCCAGCAGATCCCTAATTCTCTCTCCAATCAAGTGCGCTCTCCCCAGCCTGTCCCTTCTCCACGGCCACAGTCCCAG\|C\|CCCCCCACTCCAGTCCTTCCCCAAGGATGCAGCCTCAGCCTTCTCCACACC |
| 1:2209:6454:81943#1#F | \| \| |
| MQ=60 isize=137 C=37 | GCCCAGTCCCCACACCTACAAGGCCAGCAGATCCCTAATTCTCTCTCCAATCAAGTGCGCTCTCCCCAGCCTGTCCCTTCTCCACGGCCACAGTCCCAG\|C\|CCCCCCACTCCAGTCCTTCCCCAAGGATGCAGCCgcagGACTCCAATAGCC |
| 1:2209:8235:99339#1#F | \| \| |
| MQ=60 isize=174 C=37 | GCCCAGTCCTCACACCTACAAGGCCAGCAGATCCCTAATTCTCTCTCCAATCAAGTGCGCTCTCCCCAGCCTGTCCCTTCTCCACGGCCACAGTCCCAG\|C\|CCCCCCACTCCAGTCCTTCCCCAAGGaTGCAGCCTCAGCCTTCTCCACACC |
| 1:2210:15656:13841#1#F | \| \| |
| MQ=60 isize=194 C=33 | GCCCAGTCCCCACACCTACAAGGCCAGCAGATCCCTAATTCTCTCTCCAATCAAGTGCGCTCTCCCCAGCCTGTCCCTTCTCCACGGCCACAgtCCCaG\|C\|CCCCCCACtCCAGtCCTTCCCCaAGgATGCAGCCtCAgCCTTCTCCaCACc |
| 1:2210:13356:41372#1#F | \| \| |
| MQ=60 isize=187 C=33 | GCCCAGTCCCCACACCTACAAGGCCAGCAGATCCCTAATTCTCTCTCCAATCAAGTGCGCTCTCCCCAGCCTGTCCCTTCTCCACGGCCACAGTCCCAG\|C\|CCCCCCACTCCAGTCCTTCCCCAAGGATGCAGCCTCAgCCTTCTCCACACC |
| 1:2210:8201:81020#1#F | \| \| |
| MQ=60 isize=254 C=2 | GCCcAGTCccCACACcTACAAGGCCAgCAgaTccCTAATTcTCTCTCCAATCAAGTgCGCTCTCCCCAgcctgtcccttctccacggccacagtcccag\|c\|ccccccactccagtccttccccaaggatgcagcctcagccttctccacacc |
| 1:2211:11934:3390#1#F | \| \| |
| MQ=60 isize=168 C=33 | GCCCAGTCCCCACACCTACAAGGCCAGCAGATCCCTAATTCTCTCTCCAATCAAGTGCGCTCTCCCCAGCCTGTCCCTTCTCCACGGCCACAGTCCCAG\|C\|CCCCCCACTCCAgtCCTTCCCCAAGGATGCAGCCtCAGCCTTctCCACaCC |
| 1:2211:17727:68660#1#F | \| \| |
| MQ=60 isize=140 C=33 | GCCCAGTCCCCACACCTACAAGGCCAGCAGATCCCTaATTCTCTCTCCAATCAAGTGCGCTCTCCCCAGCCTGTCCCTTCtCCACGGCCACAGTCCCAG\|C\|CCCCCCcctCCagtCCTTCCCCtaGGaTGCAGCCtCagCCcggactccaaa |
| 1:2211:17941:72101#1#F | \| \| |
| MQ=60 isize=186 C=33 | GCCCAGTCCCCACACCTACAAGGCCAGCAGATCCCTAATTCTCTCTCCAATCAAGTGCgCTcTCCCCAGCCTGTCCCTTCTCCACGGCCACAGtcCCAG\|C\|CcCCCCACTCCAgtCcTtcCCCAAGGatgcagcctaagccttctccccacc |
| 1:2211:15342:81737#1#F | \| \| |
| MQ=60 isize=221 C=37 | GCCCAGTCCCCACACCTACAAGGCCAGCAGATCCCTAATTCTCTCTCCAATCAAGTGCGCTCTCCCCAGCCTGTCCCTTCTCCACGGCCACAGTCCCAG\|C\|CCCCCCACTCCAGTCCTTCCCCAAGGATGCAGCCTcAgCCTTCtCCaCACC |
| 1:2211:20744:95116#1#F | \| \| |
| MQ=60 isize=127 C=37 | GCCCAGTCCCCACACCTACAAGGCCAGCAGATCCCTAATTCTCTCTCCAATCAAGTGCGCTCTCCCCAGCCTGTCCCTTCTCCACGGCCACAGTCCCAG\|C\|CCCCCCACTCCAGTCCTTCCCCAAGGAGGGCTCCAATTAAACGAAGTCACt |
| 1:2212:20909:49556#1#F | \| \| |
| MQ=60 isize=230 C=33 | GCCCAGTCCCCACACCTACAAGGCCAGCAGATCCCTAATTCTCTCTCCAATCAAGTGCGCTCTCCCCAGCCTGTCCCTTCTCCACGGCCACAGTCCCAG\|C\|CCCCCCaCTCCAGTCCTTCCCCAAGGATGCAGCCTCAGCCTTCTCCACACc |
| 1:2212:9931:90426#1#F | \| \| |
| MQ=60 isize=158 C=33 | GCCCAGTCCCCACACCTACAAGGCCAGCAGATCCCTAATTCTCTCTCCAATCAAGTGCGCTCTCCCCAGCCTGTCCCTTCTCCACGGCCACAGTCCCAG\|C\|CCCCCCACTCCAGTCCTTCCCCAAGGATGCAGCCTCAGCCTTCTCCACACC |
| 1:2213:14236:5174#1#F | \| \| |
| MQ=60 isize=287 C=33 | GCCCAGTCCCCACACCTACAAGGCCAGCAGATCCCTAATTCTCTCTCCAATCAAGTgCGCTCTCCCCAGCCTGTCCCTtCTCCACGGCCaCAGTCCCAg\|C\|CCCCCCcCtCCcGTCCTtCCCCaaGGaggcagCCTCagCCTTcTCCaCacC |
| 1:2213:15175:21355#1#F | \| \| |
| MQ=60 isize=137 C=33 | GCCCAGTCCCCACACCTACAAGGCCAGCAGATCCCTAATTCTCTCTCCAATCAAGTGCGCTCTCCCCAGCCTGTCCCTTCTCCACGGCCaCAGTCCCAG\|C\|CCCCCCaCtCCAGTCCTTCCCCAAGGAtGCAGCCtCAAGGactcCAATATt |
| 1:2213:11909:44715#1#F | \| \| |
| MQ=60 isize=185 C=33 | GCCCAGTCCCCACACCTACAAGGCCAGCAGATCCCTAATTCTCTCTCCAATCGAGTGCGCTCTCCCCAGCCTGTCCCTTCTCCACGGCCACAGTCCCAG\|C\|CCCCCCaCTCCAGTCCTTCCCCAAGGATGCAGCCTCAGCCTTCtCCACACC |
| 1:2213:8333:60976#1#F | \| \| |
| MQ=60 isize=168 C=33 | GCCCAGTCCCCACACCTACAAGGCCAGCAGATCCCTAATTCTCTCTCCAATCAAGTGCGCTCTCCCCAGCCTGTCCCTTCTCCACGGCCACAGtCCCAG\|C\|CCCCCCACTCCAGtCCTTCCCCAAGGATGCAGCCTCAGCCTTCtcCACACC |
| 1:2213:7158:78447#1#F | \| \| |
| MQ=60 isize=140 C=37 | GCCCAGTCCCCACACCTACAAGGCCAGCAGGTCCCTAATTCTCTCTCCAATCAAGTGCGCTCTCCCCAGCCTGTCCCTTCTCCACGGCCACAGTCCCAG\|C\|CCCCCCACTCCAGTCCTTCCCCAAGGATGCAGCCTCAGCCAGGACTCCAAt |
| 1:2214:14842:42358#1#F | \| \| |
| MQ=60 isize=203 C=33 | GCCCAGTCCCCACACCTACAAGGCCAGCAGATCCCTAATTCTCTCTCCAATCAAGTGCGCTCTCCCCAGCCTGTCCCTTCTCCACGGCCACAGTCCCAG\|C\|CCCCCCACTCCAGtCCTTCCCCAaGGATGcagCCtCaGCcTTCTCcacacc |
| 1:2214:3510:98660#1#F | \| \| |
| MQ=60 isize=186 C=33 | GCCCAGTCCCCACACCTACAAGGCCAGCAGATCCCTAATTCTCTCTCCAATCAAGTGCGCTCTCCCCAGCCTGTCCCTTCTCCACGGCCACAGTCCCAG\|C\|CCCCCCACTCCAGTCCTTCCCCAAGGATGCAGCCTCAGCCTTCTCCACACC |
| 1:2215:19379:5929#1#F | \| \| |
| MQ=60 isize=160 C=33 | GCCCAGTCCCCACACCTACAAGGCCAGCAGATCCCTAATTCTCTCTCCAATCAAGTGCGCTCTCCCCAGCCTGTCCCTTCTCCACGGCCACAGTCCCAG\|C\|CCCCCCACtCCAGtCCTTCCCCAAGgAtgcagCCTCAgCcTtctCCacaCC |
| 1:2215:19651:69827#1#F | \| \| |
| MQ=60 isize=287 C=27 | GCCCAGTCCCCACACCTACAAGGCCAGCAGATCCCTAATTCTCTCTCCAATCAAGTGCGCTCTCCCCAGCCTGTCCCTTCTCCaCGGcCaCAGTCCCAg\|C\|CCCCCCaCTCCAGcCCTTCCCCAaGGATGCaGCCtCAGCcttctccacccc |
| 1:2215:15206:81361#1#F | \| \| |
| MQ=60 isize=162 C=33 | GCCCAGTCCCCACACCTACAAGGCCAGCAGATCCCTAATTCTCTCTCCAATCAAGTGCGCTCTCCCCAGCCTGTCCCTTCtCCACGGCCACAGTCCCAG\|C\|CCCCCCACtCCAGTCCTTCCCCAAGGAtGCagCCtCAGCCTTCTCCcCACC |
| 1:2216:13715:5291#1#F | \| \| |
| MQ=60 isize=149 C=37 | GCCCAGTCCCCACACCTACAAGGCCAGCAGATCCCTAATTCTCTCTCCAATCAAGTGCGCTCTCCCCAGCCTGTCCCTTCTCCACGGCCACAGtCCCAG\|C\|CCCCCCACTCCAGtCCTTCCCCAAGGaTGCAGCCTCAGCCTTCTCCaCAAG |
| 1:2216:8593:31830#1#F | \| \| |
| MQ=60 isize=160 C=37 | GCCCAGTCCCCACACCTACAAGGCCAGCAGATCCCTAATTCTCTCTCCAATCAAGTGCGCTCTCCCCAGCCTGTCCCTTCTCCACGGCCACAGTCCCAG\|C\|CCCCCCACTCCAGTCCTTCCCCAAGGATGCAGCCTCAGCCTTCTCCACACC |
| 1:2216:7065:54176#1#F | \| \| |
| MQ=60 isize=289 C=33 | GCCCAGTCCCCACACCTACAAGGCCAGCAGATCCCTAATTCTCTCTCCAATCAAGTGCGCTCTCCCCAGCCTGTCCCTTCTCCACGGCCACAGTCCCAG\|C\|CCCCCCACTCCAGTCCTTCCCCAAGGATGCAGCCTCAGCCTTCTCCACACC |
| 1:2216:10224:84607#1#F | \| \| |
| MQ=60 isize=230 C=33 | GCCCAGTCCCCACACCTACAAGGCCAGCAGATCCCTAATTCTCTCTCCAATCAAGTGCGCTCTCCCCAGCCTGTCCCTTCTCCACGGCCACAGTCCCAG\|C\|CCCCCCACTCCAGTCCTTCCCCAAGGATGCAGCCTCAGCCTTCTCCACACC |
| 2:1101:7146:15041#1#F | \| \| |
| MQ=60 isize=156 C=33 | GCCCAGTCCCCACACCTACAAGGCCAGCAGATCCCTAATTCTCTCTCCAATCAAGTGCGCTCTCCCCAGCCTGTCCCTTCTCCACGGCCACAGTCCCAG\|C\|CCCCCCACTCCaGtcCTTCCCCAAggAtgcagCCtcagCCTTctCCACACC |
| 2:1101:15649:50204#1#F | \| \| |
| MQ=60 isize=0 C=27 | GCCCAGTCCCCACACCTACAAGGCCAGCAGATCCCTAATTCTCTCTCCAATCAAGTGCGCTCTCCCCAGCCTGTCCCTTCTCCACGGCCACAGTCCCAG\|C\|ACCTAAATCCAAATGCATGATcCTTCCTCAAGAGCATTTCCTCAAcCCAGG |
| 2:1102:7156:20931#1#F | \| \| |
| MQ=60 isize=154 C=33 | GCCCAGTCCCCACACCTACAAGGCCAGCAGATCCCTAATTCTCTCTCCAATCAAGTGCGCTCTCCCCAGCCTGTCCCTTCTCCACGGCCACAGTCCCAG\|C\|CCCCCCaCtCCAGtCCTTCCCCAAGGAtgcAGCCtcagCCTTCTCCaCACc |
| 2:1102:2925:49394#1#F | \| \| |
| MQ=60 isize=186 C=33 | GCCCAGTCCCCACACCTACAAGGCCAGCAGATCCCTAATTCTCTcTCCAATCAAgTGCGCtCTCCCCAGCCTGgCCCTTCtCcACGgcCACAGccCCAg\|C\|CCCCCcActCCAGacCTcCCCcaaggatccagcctccgccttctcacaacc |
| 2:1102:4360:98765#1#F | \| \| |
| MQ=60 isize=200 C=33 | GCCCAGTCCCCACACCTACAAGGCCAGCAGATCCCTAATTCTCTCTCCAATCAAGTGCGCTCTCCCCAgCCTGTCCCTTCTCCaCGGCCACAgtCCCAG\|C\|CCCCCCACtCCAGcCcTtCCCCAAGGATGCagcctCAGCCTtcccCACaCc |
| 2:1103:18627:28578#1#F | \| \| |
| MQ=60 isize=150 C=33 | GCCCAGTCCCCACACCTACAAGGCCAGCAGATCCCTAATTCTCTCTCCAATCAAGTGCGCTCTCCCCAGCCTGTCCCTTCTCCACGGCCACAGTCCCAG\|C\|CCCCCCACTCCAGTCCTTCCCCAAGGATGCAGCCTCAGCCTTCTCCACACa |
| 2:1103:9186:51031#1#F | \| \| |
| MQ=60 isize=268 C=33 | GCCCAGTCCCCACACCTACAAGGCCAGCAGATCCCTAATTCTCTCTCCAATCAAGTGCGCTCTCCCCAGCCTGTCCCTTCTCCACGGCCACAGTCCCAG\|C\|CCCCCCACTCCAGTCCTTCCCCAAGGATGCAGCCTCAgCCTTCTCCACACC |
| 2:1103:2741:84031#1#F | \| \| |
| MQ=60 isize=138 C=33 | GCCCAGTCCCCACACCTACAAGGCCAGCAGATCCCTAATTCTCTCTCCAATCAAGTGCGCTCTCCCCAGCCTGtCCCTTCTCCACGGCCAcAgtCCCAG\|C\|CCCCCCaCtCCAGtCCTtCCCCAaggatgcagcctcaggactccaatctgg |
| 2:1104:19234:45960#1#F | \| \| |
| MQ=60 isize=149 C=37 | GCCCAGTCCCCACACCTACAAGGCCAGCAGATCCCTAATTCTCTCTCCAATCAAGTGCGCTCTCCCCAGCCTGTCCCTTCTCCACGGCCACAGTCCCAG\|C\|CCCCCCACTCCAGTCCTTCCCCAAGGATGCAGCCTCAGCCTTCTCCACAGG |
| 2:1105:8813:6596#1#F | \| \| |
| MQ=60 isize=175 C=37 | GCCCAGTCCCCACACCTACAAGGCCAGCAGATCCCTAATTCTCTCTCCAATCAAGTGCGCTCTCCCCAGCCTGTCCCTTCTCCACGGCCACAGTCCCAG\|C\|CCCCCCaCTCCAGtCCTTCCCCAAGGATGCAGCCTCAGCCTTCTCCACACC |
| 2:1105:19038:39909#1#F | \| \| |
| MQ=60 isize=186 C=27 | GCCCAGTCCCCACACCTACAAGGCCAGCAGATCCCTAATTCTCTCTCCAATCAAGTGCGCTCTCCCCAgCCTGTCCCTTCTCCACGGCCACAGTCCCaG\|C\|CCCCCCACtCCaGTCCtTCCCCAAGGATGCAGCCTCAGCCTTCTCCACACC |
| 2:1105:6996:100836#1#F | \| \| |
| MQ=60 isize=147 C=22 | GCCCAGTCCCCACACCTACAAGGCCAGCAGATCCCTAATTCTCTCTCCAATCAAGtGCGCTCCCCCCAGCCTGTCCCTTCTCCACGGCcacAGTCCCag\|C\|CCCCCCACtCCAGTCCTtCCCCaaggatgcagcctcacccttctccaggac |
| 2:1106:19299:18158#1#F | \| \| |
| MQ=60 isize=164 C=33 | GCCCAGTCCCCACACCTACAAGGCCAGCAGATCCCTAATTCTCTCTCCAATCAAGTGCGCTCTCCCCAGCCTGTCCCtTCTCCACggCCACAgtcCCag\|C\|acccCCACtccAgtcCTTcCCCAaggaggcAgccTCAGCCttctccacacc |
| 2:1106:20523:43674#1#F | \| \| |
| MQ=60 isize=233 C=15 | GCCCAGTCCCCACACCTACAAGGCCAGCAGATCCCTAATTCTCTCTCCAATCAAGTGCGCTCTCCCCAGCCTGTCCCTTCTCCACGGCCACAGTCCCAG\|c\|CCCCCCACTCCAGTCCTTCCCCAAGGATGCAGcCTCAGCCTTCTCCACACC |
| 2:1106:8861:96841#1#F | \| \| |
| MQ=60 isize=0 C=33 | GcCCAGTCCCCACACCTACAAGGCCAGCAGATCCCTAATTCTCTCTCCAATCAAGTGCGCTCTCCCCAGCCTGTCCCTTCTCCACGGCCACAgTCCCAG\|C\|CcCCCCACTCCAgtCCTtCCCCAAAGgacTCCAAtaatgtactaaccctgt |
| 2:1107:4620:19186#1#F | \| \| |
| MQ=60 isize=125 C=37 | GCCCAGTCCCCACACCTACAAGGCCAGCAGATCCCTAATTCTCTCTCCAATCAAGTGCGCTCTCCCCAGCCTGTCCCTTCTCCACGGCCACAGTCCCAG\|C\|CCCCCCACtCCaGTCCTTCCCCAAGAGGACTCCAATAGtaACAAgATCctg |
| 2:1107:13636:54687#1#F | \| \| |
| MQ=60 isize=242 C=33 | GCCCAGTCCCCACACCTACAAGGCCAGCAGATCCCTAATTCTCTCTCCAATCAAGTGCGCTCTCCCCAGCCTGTCCCTTCTCCACGGCCACAGTCCCAG\|C\|CCCCTCACTCCAGTCCTTCCCCAAGGaTGCAGCCTCAGCCTTCTCCACACC |
| 2:1107:13110:90950#1#F | \| \| |
| MQ=60 isize=237 C=33 | GCCCAGTCCCCACACCTACAAGGCCAGCAGATCCCTAATTCTCTCTCCAATCAAGTGCGCTCTCCCCAGCCTGTCCCTTCTCCACGGCCACAGTCCCAG\|C\|CCCCCCACTCCAGTCCTTCCCCAAGGATGCAGCCTCAGCCTTCTCCACACC |
| 2:1108:18677:46916#1#F | \| \| |
| MQ=60 isize=273 C=33 | GCCCAGTCCCCACACCTACAAGGCCAGCAGATCCCTAatTCTCTCTCCAATCAAGTGCGCTCTCCCCAGCCTGTCCCTTCTCCACGGCCACAGTCCCAG\|C\|CCCCCCACTCCAGTCCTTCCCCAAGGATGCAGCCTCAGCCTTCTCCaCACC |
| 2:1108:9106:70978#1#F | \| \| |
| MQ=60 isize=201 C=33 | GcCCAGTCCCCACACCTACAAGGCCAGCAGATCCCTAATTCTCTCTCCAATCAAGTGCGCTCTCCCCAGCCTGTCCCTTCTCCACGgCCaCAgTCCCAG\|C\|CCCCCCactCCagtccTtcCCCAaGGATgcAgCCtCAgCCTTCTCCacacc |
| 2:1108:13413:90089#1#F | \| \| |
| MQ=60 isize=205 C=33 | GCCCAGTCCCCACACCTACAAGGCCAGCAGATCCCTAATTCTCTCTCCAATCAAGTGCGCTCTCCCCAgCCTGTCCCTTCTCCACGGCCACAGTCCCAG\|C\|CCCCCCACTCCAGTCCTTCCCCAAGGATGCAGCCTCAGCCTTCTCCACACC |
| 2:1109:13603:3802#1#F | \| \| |
| MQ=60 isize=131 C=37 | GCCCAGTCCCCACACCTACAAGGCCAGCAGATCCCTAATTCTCTCTCCAATCAAGTGCGCTCTCCCCAGCCTGTCCCTTCTCCACGGCCACAGTCCCAG\|C\|CCCCCCACTCCAGTCCTTCCCCAAGGATGCAAGGACTCCAATTCTAGACAt |
| 2:1109:20674:39061#1#F | \| \| |
| MQ=60 isize=137 C=37 | GCCCAGTCCCCACACCTACAAGGCCAGCAGATCCCTAATTCTCTCTCCAATCAAGTGCGCTCTCCCCAGCCTGTCCCTTCTCCACGGCCACAGTCCCAG\|C\|CCCCCCACTCCAGTCCTTCCCCAAGGATGCAGCCTCAGGACTCCAATAGGG |
| 2:1109:7237:69035#1#F | \| \| |
| MQ=60 isize=192 C=33 | GCCCAGTCCCCACACCTACAAGGCCAGCAGATCCCTAATTCTCTCTCCAATCAAGTGCGCTCTCCCCAGCCTGTCCCTTCTCCACGGCCACAGTCCCAG\|C\|CCCCCCACTCCAGTCCTTCCCCAAGGaTGCAGCCTCAgCCTTCTCCACACC |
| 2:1110:3923:6541#1#F | \| \| |
| MQ=60 isize=121 C=27 | GCCCAGTCCCCACACCTACAAGGCCAGCAGATCCCTAATTCTCTCTCCAATCAAGTGCGCTCTCCCCAGCCTGtCCCTTCTCCACGgCCACaGtcccAG\|C\|CCCCCCACTCCAGtCcTtCCcaggactccaataacaagacggccctgtctc |
| 2:1110:9111:25920#1#F | \| \| |
| MQ=60 isize=229 C=33 | GCCCAGTCCCCACACCTACAAGGCCAGCAGATCCCTAATTCTCTCTCCAATCAAGTGCGCTCTCCCCAGCCTGTCCCTTCTCCACGGCCACAGTCCCAG\|C\|CCCCCCACTCCAGTCCTTCCCCAAGGATGCAGCCTCAGCCTTCTCCACACC |
| 2:1110:1408:42620#1#F | \| \| |
| MQ=60 isize=121 C=33 | GCCCAGTCCCCACACCTACAAGGCCAGCAGATCCCTAATTCTCTCTCCAATCAAGtGCGCTCTCCCCAGCCTGTCCCTTCTCCaCGGCCaCAGtCCCAG\|C\|CCCCCCcctCCAGtCcTTCCCaGGacccCAaTTTTTcatGGAAactgtctc |
| 2:1110:8701:58647#1#F | \| \| |
| MQ=60 isize=140 C=33 | GCCCAGTCCCCACACCTACAAGGCCAGCAGATCCCTAATTCTCTCTCCAATCAAGTGCGCTCTCCCCAGCCTGTCCCTTCTCCACGGCCACAGTCCCAG\|C\|CCCCCCACTCCAGTCCTTCCCCAAGGATGCAGCCTCAGCCAGGACTCCAAt |
| 2:1110:1675:78099#1#F | \| \| |
| MQ=60 isize=0 C=27 | GCCCAGTCCCCACACCTACAAGGCCAGCAGATCCCTAATTCTCTCTCCAATCAAGTGCGCTCTCCCCAGCCTGTCCCTTCTCCACGGCCACAGTCCCAg\|C\|CCCCCCaCtCCAGtCCTTCCCCAaggatgcagccccagccttccccccccc |
| 2:1110:5508:87715#1#F | \| \| |
| MQ=60 isize=162 C=2 | gcCCAGTCCCCACACCtACAAGGCCAGcagAtccCTAATTcTcTCTCCAAtcaagtgcgctttccccagcctgtcccttctccacgcccacagccccag\|c\|ccccccactccggtccttccccaacgatgcagcctcacccttctccccacc |
| 2:1111:5163:44641#1#F | \| \| |
| MQ=60 isize=162 C=33 | GCCCAGTCCCCACACCTACAAGGCCAGCAGATCCCTAATTCTCTCTCCAATCAAGTGCGCtCTCCCCAGCCTGtCCCTTCTCCACGGCCaCAGTCCcaG\|C\|CCCCCCACtccagccCTtCCCCAAggatgcaccctcagccttctccacccc |
| 2:1111:14541:74142#1#F | \| \| |
| MQ=60 isize=147 C=37 | GCCCAGTCCCCACACCTACAAGGCCAGCAGATCCCTAATTCTCTCTCCAATCAAGTGCGCTCTCCCCAGCCTGTCCCTTCTCCACGGCCACAGTCCCAG\|C\|CCCCCCACTCCAGTCCTTCCCCAAGGATGCAGCCTCAGCCTTCTCCAGGAC |
| 2:1112:13497:6706#1#F | \| \| |
| MQ=60 isize=127 C=37 | GCCCAGTCCCCACACCTACAAGGCCAGCAGATCCCTAATTCTCTCTCCAATCAAGTGCGCTCTCCCCAGCCTGTCCCTTCTCCACGGCCACAGTCCCAG\|C\|CCCCCCACTCCAGTCCTTCCCCAAGGACTCCAATCAAAACCGTAAGCTGTC |
| 2:1112:16959:10357#1#F | \| \| |
| MQ=60 isize=159 C=33 | GCCCAGTCCCCACACCTACAAGGCCAGCAGATCCCTAATTCTCTCTCCAATCAAGTGCGCTCTCCCCAGCCTGTCCCTTCTCCACGGCCACAGTCCCAG\|C\|CCCCCCACTCCAGTCCTTCCCCAAGGATGCAGCCTCAGCCTTCTCCACACC |
| 2:1112:10082:20674#1#F | \| \| |
| MQ=60 isize=170 C=33 | GCCCAGTCCCCACACCTACAAGGCCAGCAGATCCCTAATTCTCTCTCCAATCAAGTGTGCTCTCCCCAGCCTGTCCCTTCTCCACGGCCACAGTCCCAG\|C\|CCCCCCACTCCAGTCCTTCCCCAAGGATGCAGCCTCAGCCTTCTCCACACC |
| 2:1112:20958:54069#1#F | \| \| |
| MQ=60 isize=186 C=6 | GCCCAGTCCCCACACCTACAAGGCCAGCAGATCCCTAATTCTcTCTCCAATCAAGTGCGCTCTCCCCAGCCTGTCCCTTCTCCACGGtCACAGTccCAG\|c\|ccccCCacttcagtccttcccccaggacggagcctcagtcttctccacacc |
| 2:1112:5163:67573#1#F | \| \| |
| MQ=60 isize=121 C=37 | GCCCAGTCCCCACACCTACAAGGCCAGCAGATCCCTAATTCTCTCTCCAATCAAGTGCGCTCTCCCCAGCCTGTCCCTTCTCCACGGCCACAGTCCCAG\|C\|CCCCCCACTCCAGtCCTTCCCAGGACTCCAATCTCTGCCAgcCTCTGTCTc |
| 2:1112:5472:98144#1#F | \| \| |
| MQ=60 isize=143 C=27 | GCCCAGTCCCCACACCTACAAGGCCAGCAGATCCCTAATTCTCTCTCCAATCAAGTGCGCTCTCCCCAGCCTGTCCCTTCTCCACGGCCACAGTCCCAg\|C\|CCCCCCACTCCAGTCCTTCCCCAAGGATGCAGCctcAGCCTTCAGGACTCC |
| 2:1113:21239:50704#1#F | \| \| |
| MQ=60 isize=209 C=27 | GCCCAGTCCCCACACCTACAAGGCCAGCAGATCCCTAATTCTCTCTCCAATCAAGTGCGCTCTCCCCAGCCTGTCCCTTCTCCACGGCCACAGTCCCAG\|C\|CCCCCCACTCCAGTCCTTCCCCAAGGATGCaGCCTCAGCCTTCTCCACACC |
| 2:1113:3948:69868#1#F | \| \| |
| MQ=60 isize=143 C=37 | GCCCAGTCCCCACACCTACAAGGCCAGCAGATCCCTAATTCTCTCTCCAATCAAGTGCGCTCTCCCCAGCCTGTCCCTTCTCCACGGCCACAGTCCCAG\|C\|CCCCCCACTCCAGTCCTTCCCCAAGGATGCAGCCTCAGCCTTCAGGACTCC |
| 2:1113:16145:76094#1#F | \| \| |
| MQ=60 isize=149 C=22 | GCCCAGTCCCCACACCTACAAGGCCAGCAGATCCCTAATTCTCTCTCCAATCAAGTGCGCTCTCCCCAGCCTGTCCCTTCTCCACGGCCACAGTCCCAg\|C\|CCCCCCACTCCAgtCCTTCCCCaAgGaTGCagCCTCAgCCttCtcCcccgg |
| 2:1114:3637:13809#1#F | \| \| |
| MQ=60 isize=183 C=33 | GCCCAGTCCCCACACCTACAAGGCCAGCAGATCCCTAATTCTCTCTCCAATCAAGTGCGCTCTCCCCAGCCTGTCCCTTCTCCACGGCCACAGTCCCAG\|C\|CCCCCCACTCCAGtCCTTCCCCAAGGATGCAGCCTCAGCCTTCTCCACACC |
| 2:1114:12799:35798#1#F | \| \| |
| MQ=60 isize=307 C=33 | GCCCAGTCCCCACACCTACAAGGCCAGCAGATCCCTAATTCTCTCTCCAATCAAGTGCGCTCTCCCCAGCCTGTCCCTTCTCCACGGCCACAGTCCCAG\|C\|CCCCCCACTCCAGTCCTTCCCCAAGGATGCAGCCTCAGCCTTCTCCACACC |
| 2:1114:17920:79284#1#F | \| \| |
| MQ=60 isize=133 C=37 | GCCCAGTCCCCACACCTACAAGGCCAGCAGATCCCTAATTCTCTCTCCAATCAAGTGCGCTCTCCCCAGCCTGTCCCTTCTCCACGGCCACAGTCCCAG\|C\|CCCCCCACTCCAGTCCTTCCCCAAGGATGCAGCAGGACTCCAATtCTGCTG |
| 2:1115:12859:2979#1#F | \| \| |
| MQ=60 isize=126 C=37 | GCCCAGTCCCCACACCTACAAGGCCAGCAGATCCCTAATTCTCTCTCCAATCAAGTGCGCTCTCCCCAGCCTGTCCCTTCTCCACGGCCACAGTCCCAG\|C\|CCCCCCACTCCAGTCCTTCCCCAAGGACTCCAATATTTTTATGCCGACTGT |
| 2:1115:14816:33142#1#F | \| \| |
| MQ=60 isize=123 C=33 | GCCCAGTCCCCACACCTACAAGGCCAGCAGATCCCTAATTCTCTCTCCAATCAAGTGCGCTCTCCCCAGCCTGTCCCTTCTCCACGGCCACAGTCCCAG\|C\|CCCCCCACTCCAGTCCTTCCCCAGGACTCCAATCTGGGTATAGGACTGTCT |
| 2:1115:13850:50386#1#F | \| \| |
| MQ=60 isize=175 C=37 | GCCCAGTCCCCACACCTACAAGGCCAGCAGATCCCTAATTCTCTCTCCAATCAAGTGCGCTCTCCCCAGCCTGTCCCTTCTCCACGGCCACAGTCCCAG\|C\|CCCCCCACTCCAGTCCTTCCCCAAGGATGCAGCCTCAGCCTTCTCCACACC |
| 2:1116:19644:3150#1#F | \| \| |
| MQ=60 isize=121 C=37 | GCCCAGTCCCCACACCTACAAGGCCAGCAGATCCCTAATTCTCTCTCCAATCAAGTGCGCTCTCCCCAGCCTGTCCCTTCTCCACGGCCACAGTCCCAG\|C\|CCCCCCACTCCAGTCCTTCCCAGGACTCCAATTCCGCCTGTCTGCTGTCTC |
| 2:1116:14465:36742#1#F | \| \| |
| MQ=60 isize=127 C=33 | GCCCAGTCCCCACACCTACAAGGCCAGCAGATCCCTAATTCTCTCTCCAATCAAGTGCGCTCTCCCCAGCCTGTCCCTTCTCCACGGCCACAGTCCCAG\|C\|CCCCCCACTCCAGTCCTTCCCCAaGGACTCCaATTTGCCAGAACCTCTGTC |
| 2:1116:19816:59304#1#F | \| \| |
| MQ=60 isize=156 C=22 | GCCCAGTCCCCACACCTACAAGGCCAGCAGATCCCTAATTCTCTCTCCAATCAAGtGCGCTCTCCCCAGCCTGTCCCTTCTCCACGGCCACAGtCCCAg\|C\|CCCCCCACTCCAGTCCTTCCCCAaGGATGCAGCCTCAGCCTTCTCCACaCC |
| 2:1116:7153:78034#1#F | \| \| |
| MQ=60 isize=160 C=37 | GCCCAGTCCCCACACCTACAAGGCCAGCAGATCCCTAATTCTCTCTCCAATCAAGTGCGCTCTCCCCAGCCTGTCCCTTCTCCACGGCCACAGTCCCAG\|C\|CCCCCCACtCCAGTCCTTCCCCAAgGaTGCAgCCtCAGCCTTCTCCACACC |
| 2:1116:5018:97401#1#F | \| \| |
| MQ=60 isize=139 C=33 | GCCCAGTCCCCACACCTACAAGGCCAGCAGATCCCTAATTCTCTCTCCAATCAAGTGCGCTCTCCCCAGCCTGTCCCTTCTCCACGGCCACAGTCCCAG\|C\|CCCCCCACTCCaGtCCTTCCCCAaGGATGCagCCccagCAGGactCCAATC |
| 2:1201:9622:5864#1#F | \| \| |
| MQ=60 isize=186 C=33 | GCCCAGTCCCCACACCTACAAGGCCAGCAGATCCCTAATTCTCTCTCCAATCAAGTGCGCTCTCCCCAGCCTGTCCCTTCTCCaCGGCCACAGtCCCAG\|C\|CCCCCCACtCCagtccTTCCCCAAgGAtgCAGCCTCAcCCTTttccccacc |
| 2:1201:8705:27110#1#F | \| \| |
| MQ=60 isize=0 C=37 | GCCCAGTCCCCACACCTACAAGGCCAGCAGATCCCTAATTCTCTCTCCAATCAAGTGCGCTCTCCCCAGCCTGTCCCTTCTCCACGGCCACAGTCCCAG\|C\|CCCCCCACTCCAGTCCTTCCCCAAGGATGCAGCCTCAGCCTTCTCCACACC |
| 2:1201:11452:52066#1#F | \| \| |
| MQ=60 isize=162 C=33 | GCCCAGTCCCCACACCTACAAGGCCAGCAGATCCCTAATTCTCTCTCCAATCAAGTGCGCTCTCCCCAGCCTGTCCCTTCTCCACGGCCACAGTCCCAG\|C\|CCCCCCACTCCAGtCCTTCCCCAAGGATGCAGCCTCAGCCTTCTCCACACC |
| 2:1201:8059:75241#1#F | \| \| |
| MQ=60 isize=147 C=37 | GCCCAGTCCCCACACCTACAAGGCCAGCAGATCCCTAATTCTCTCTCCAATCAAGTGCGCTCTCCCCAGCCTGTCCCTTCTCCACGGCCACAGTCCCAG\|C\|CCCCCCACTCCAGTCCTTCCCCAAGGATGCAGCCTCAGCCTTCTCCAGGAC |
| 2:1201:7970:89612#1#F | \| \| |
| MQ=60 isize=138 C=33 | GCCCAGTCCCCACACCTACAAGGCCAGCAGATCCCTAATTCTCTCTCCAATCAAGTGCGCTCTCCCCAGCCTGTCCCTTCTCCACGGCCACAGTCCCAG\|C\|CCCCCCACTCCAGTCCTTCCCCAAGGATGCAGCCTCAGGACTCCAATCTGA |
| 2:1202:15176:20236#1#F | \| \| |
| MQ=60 isize=152 C=37 | GCCCAGTCCCCACACCTACAAGGCCAGCAGATCCCTAATTCTCTCTCCAATCAAGTGCGCTCTCCCCAGCCTGTCCCTTCTCCACGGCCACAGTCCCAG\|C\|CCCCCCACTCCAGTCCTTCCCCAAGGATGCAGCCTCAGCCTTCTCCACACC |
| 2:1202:19396:32312#1#F | \| \| |
| MQ=60 isize=280 C=27 | GCCCAGTCCCCACACCTACAAGGCCAGCAGATCCCTAATTCTCTCTCCAATCAAGTGCGCTCTCCCCAGCCTGTCCCTTCTCCACGGCCACAGTCCCAg\|C\|CCCCCCACTCCAGTCCTTCCCCAAGGATGCAGCCTCAGCCTTCTCCACACC |
| 2:1202:15083:59086#1#F | \| \| |
| MQ=60 isize=127 C=33 | GCCCAGTCCCCACACCTACAAGGCCAGCAGATCCCTAATTCTCTCTCCAATCAAGTGCGCTCTCCCCAGCCTGTCCCTTCTCCACGGCCACAGTCCCAG\|C\|CCCCCCACTCCAGTCCTTCCCCAAGGACTCCAATAGGCTTACCAACCTGTc |
| 2:1202:17767:75362#1#F | \| \| |
| MQ=60 isize=232 C=33 | GCCCAGTCCCCACACCTACAAGGCCAGCAGATCCCTAATTCTCTCTCCAATCAAGTGCGCTCTCCCCAGCCTGTCCCTTCTCCACGGCCACAGTCCCAG\|C\|CCCCCCACTCCAGTCCTTCCCCAAGGATGCAGCCTCAGCCTTCTCCACACC |
| 2:1203:14365:10510#1#F | \| \| |
| MQ=60 isize=131 C=37 | GCCCAGTCCCCACACCTACAAGGCCAGCAGATCCCTAATTCTCTCTCCAATCAAGTGCGCTCTCCCCAGCCTGTCCCTTCTCCACGGCCACAGTCCCAG\|C\|CCCCCCACTCCAGTCCTTCCCCAAGGATGCAAGGACTCCAATGACACCTGG |
| 2:1203:11568:44965#1#F | \| \| |
| MQ=60 isize=236 C=33 | GCCCAGTCCCCACACCTACAAGGCCAGCAGATCCCTAATTCTCTCTCCAATCAAGTGCGCTCTCCCCAGCCTGTCCCTTCTCCACGGCCACAGTCCCAG\|C\|CCCCCCACTCCAGTCCTTCCCCAAGGATGCAGCCTCAGCCTTCTCCACACC |
| 2:1203:7737:70048#1#F | \| \| |
| MQ=60 isize=162 C=37 | GCCCAGTCCCCACACCTACAAGGCCAGCAGATCCCTAATTCTCTCTCCAATCAAGTGCGCTCTCCCCAGCCTGTCCCTTCTCCACGGCCACAGTCCCAG\|C\|CCCCCCACTCCAGTCCTTCCCCAAGGATGCAGCCTCAGCCTTCTCCACACC |
| 2:1203:17499:95292#1#F | \| \| |
| MQ=60 isize=233 C=37 | GCCCAGTCCCCACACCTACAAGGCCAGCAGATCCCTAATTCTCTCTCCAATCAAGTGCGCTCTCCCCAGCCTGTCCCTTCTCCACGGCCACAGTCCCAG\|C\|CCCCCCACTCCAGTCCTTCCCCAAGGATGCAGCCTCAGCCTTCTCCACACC |
| 2:1204:13557:37828#1#F | \| \| |
| MQ=60 isize=177 C=33 | GCCCAGTCCCCACACCTACAAGGCCAGCAGATCCCTAATTCTCTCTCCAATCAAGTGCGCTCTCCCCAGCCTGTCCCTTCTCCACGGCCACAGTCCCAG\|C\|CCCCCCACTCCAGtCCTTCCCCAAGGATGCAGCCTCAGCCTtCTCCACACC |
| 2:1204:12096:67699#1#F | \| \| |
| MQ=60 isize=137 C=37 | GCCCAGTCCCCACACCTACAAGGCCAGCAGATCCCTAATTCTCTCTCCAATCAAGTGCGCTCTCCCCAGCCTGTCCCTTCTCCACGGCCACAGtCCCAG\|C\|CCCCCCACTCCAGTCCTTCCCCAAGGATGCAGCCTCAAGGACTCCAATTAA |
| 2:1204:1910:75260#1#F | \| \| |
| MQ=60 isize=127 C=33 | GCCCAGTCCCCACACCTACAAGGCCAGCAGATCCCTAATTCTCTCTCCAATCAAGTGCGCTCTCCCCAGCCTGTCCCTTCTCCACGGCCACAGTCCCAG\|C\|CCCCCCACTCCAGTCCTTCCCCAAGGAGGACTCCAATGTAACTCGTGCCCT |
| 2:1204:6046:87893#1#F | \| \| |
| MQ=60 isize=132 C=33 | GCCCAGTCCCCACACCTACAAGGCCAGCAGATCCCTAATTCTCTCTCCAATCAAGTGCGCTCTCCCCAGCCTGTCCCTTCTCCACGGCCACAGTCCCAg\|C\|CCCCCCACTCCAGTCCTTCCCCAAGGatgCAGAGGACtCCaATCCACGCGC |
| 2:1205:8322:37182#1#F | \| \| |
| MQ=60 isize=184 C=37 | GCCCAGTCCCCACACCTACAAGGCCAGCAGATCCCTAATTCTCTCTCCAATCAAGTGCGCTCTCCCCAGCCTGTCCCTTCTCCACGGCCACAGTCCCAG\|C\|CCCCCCACTCCAGTCCTTCCCCAAGGATGCAGCCTCAGCCTTCTCCACACC |
| 2:1205:6434:73240#1#F | \| \| |
| MQ=60 isize=159 C=33 | GCCCAGTCCCCACACCTACAAGGCCAGCAGATCCCTAATTCTCTCTCCAATCAAGTGCGCTCTCCCCAGCCTGTCCCTTCTCCACGGCCACAGTCCCAG\|C\|CCCCCCaCTCCAGTCCTTCCCCAAGGAtGCagCCTCAGCCTTCTCCGCACC |
| 2:1205:19756:100305#1#F | \| \| |
| MQ=60 isize=203 C=33 | GCCCAGTCCCCACACCTACAAGGCCAGCAGATCCCTAATTCTCTCTCCAATCAAGTGCGCTCTCCCCAGCCTGTCCCTTCTCCACGgCCACAGtCCCAg\|C\|cCCCCCACTCCAgTCCTTCCCCAAGGAtGCAGCCTCagCCTTCTCCACACC |
| 2:1206:17664:5652#1#F | \| \| |
| MQ=60 isize=162 C=37 | GCCCAGTCCCCACACCTACAAGGCCAGCAGATCCCTAATTCTCTCTCCAATCAAGTGCGCTCTCCCCAGCCTGTCCCTTCTCCACGGCCACAGTCCCAG\|C\|CCCCCCACTCCAGTCCTTCCCCAAGGATGCAGCCTCAGCCTTCTCCACACC |
| 2:1206:16537:46445#1#F | \| \| |
| MQ=60 isize=0 C=15 | GCCCAGacCCCACACCTACAAggCCAGCAgATCCCTAATTCTCTCTCCAATCAAGTGCGCTaTCCCCAGCCtGTCCCtTCTCCACGgccaCAgtCCCag\|c\|CCCCCCACTCcAGTCCTtCCCCaagGAtgcAGCCtCAgCCTtCtccACACC |
| 2:1206:17344:59281#1#F | \| \| |
| MQ=60 isize=177 C=33 | GCCCAGTCCCCACACCTACAAGGCCAGCAGATCCCTAATTCTCTCTCCAATCAAGTGCGCTCTCCCCAGCCTGTCCCtTCTCCACGGCCACAGTCCCAG\|C\|CCCCCCACTCCAGTCCTTCCCCAAGGATGCAGCCTCAGCCTTCTCCACACC |
| 2:1206:19838:94162#1#F | \| \| |
| MQ=60 isize=257 C=33 | GCCCAGTCCCCACACCTACAAGGCCAGCAGATCCCTAATTCTCTCTCCAATCAAGTGCGCTCTCCCCAGCCTGTCCCTTCTCCACGGCCACAGTCCCAG\|C\|CCCCCCACTCCAGTCCTTCCCCAAGGATGCAGCCTCAGCCTTCTCCACACC |
| 2:1207:12240:34426#1#F | \| \| |
| MQ=60 isize=285 C=33 | GCCCAGTCCCCACACCTACAAGGCCAGCAGATCCCTAATTCTCTCTCCAATCAAGTGCGCTCTCCCCAGCCTGTCCCTTCTCCACGGCCACAGTCCCAG\|C\|CCCCCCACTCCAGTCCTTCCCCAAGGATGCAGCCTCAGCCTTCTCCACACC |
| 2:1207:5412:62742#1#F | \| \| |
| MQ=60 isize=231 C=33 | GCCCAGTCCCCACACCTACAAGGCCAGCAGATCCCTAATTCTCTCTCCAATCAAGTGCGCTCTCCCCAGCCTGTCCCTTCTCCACGGCCACAGTCCCAG\|C\|CCCCCCACTCCAGTCCTTCCCCAAGGATGCAGCCTCAGCCTTCTCCACACC |
| 2:1207:4462:78684#1#F | \| \| |
| MQ=60 isize=135 C=33 | GCCCAGTCCCCACACCTACAAGGCCAGCAGATCCCTAATTCTCTCTCCAATCAAGTGCGCTCTCCCCAGCCTGTCCCTTCTCCACGGCCACAGTCCCAG\|C\|CCCCCCACTCCAGtCCTTCCCCAAGGATGCAGCCTAGGACTCCaAtCTGCC |
| 2:1207:7739:88952#1#F | \| \| |
| MQ=60 isize=160 C=33 | GCCCAGTCCCCACACCTACAAGGCCAGCAGATCCCTAATTCTCTCTCCAATCAAGTGCGCTCTCCCCAGCCTGTCCCTTCTCCACGGCCACAGTCCCAG\|C\|CCCCCCACTCCAGTCCTTCCCcaaggatgcagctccagccttctccccacc |
| 2:1208:2845:3577#1#F | \| \| |
| MQ=60 isize=307 C=33 | GCCCAGTCCCCACACCTACAAGGCCAGCAGATCCCTAATTCTCTCTCCAATCAAGTGCGCTCTCCCCAGCCTGTCCCTTCTCCACGGCCACAGTCCCAG\|C\|CCCCCCACTCCAGTCCTTCCCCAAGGATGCAGCCTCAGCCTTCTCCACACC |
| 2:1208:18484:48987#1#F | \| \| |
| MQ=60 isize=170 C=33 | GCCCAGTCCCCACACCTACAAGGCCAGCAGATCCCTAATTCTCTCTCCAATCAAGTGCGCTCTCCCCAGCCTGTCCCTTCTCCACGGCCACAGTCCCAG\|C\|CCCCCCACTCCAGTCCTTCCCCAAGGATGCAGCCTCAGCCTTCTCCAcacc |
| 2:1209:5080:27582#1#F | \| \| |
| MQ=60 isize=144 C=33 | GCCCAGTCCCCACACCTACAAGGCCAGCAGATCCCTAATTCTCTCTCCAATCAAGCGCGCTCTCCCCAGCCTGTCCCTTCTcCACGgCcACAGtCCCAG\|C\|CCCCCCACTCCAgtCCTTCCCCAaggAtgCAgCCTcagCCttctaggactc |
| 2:1209:13761:90965#1#F | \| \| |
| MQ=60 isize=158 C=37 | GCCCAGTCCCCACACCTACAAGGCCAGCAGATCCCTAATTCTCTCTCCAATCAAGTGCGCTCTCCCCAGCCTGTCCCTTCTCCACGGCCACAGTCCCAG\|C\|CCCCCCACTCCAGTCCTTCCCCAAGGATGCAGCCTCAGCCTTCTCCACACC |
| 2:1209:17171:96786#1#F | \| \| |
| MQ=60 isize=236 C=33 | GCCCAGT\_CCCACACCTACAAGGCCAGCAGATCCCTAATTCTCTCTCCAATCAAGTGCGCTCTCCCCAGCCTGTCCCTTCTCCACGGCCACAGTCCCAG\|C\|CCCCCCACTCCaGTCCTTCCCCAAGGATGCAGCCTCAGCCTTCTCCACACCa |
| 2:1210:1807:31827#1#F | \| \| |
| MQ=60 isize=165 C=37 | GCCCAGTCCCCACACCTACAAGGCCAGCAGATCCCTAATTCTCTCTCCAATCAAGTGCGCTCTCCCCAGCCTGTCCCTTCTCCACGGCCACAGTCCCAG\|C\|CCCCCCACTCCAGtCCTTCCCCAAGGATGCAGCCTCAGCCTTCtCCACacc |
| 2:1210:18317:46026#1#F | \| \| |
| MQ=60 isize=144 C=37 | GCCCAGTCCCCACACCTACAAGGCCAGCAGATCCCTAATTCTCTCTCCAATCAAGTGCGCTCTCCCCAGCCTGTCCCTTCTCCACGGCCACAGTCCCAG\|C\|CCCCCCACTCCAGTCCTTCCCCAAGGATGCAGCCTCAGCCTTCTAGGACTC |
| 2:1210:9556:92949#1#F | \| \| |
| MQ=60 isize=153 C=37 | GCCCAGTCCCCACACCTACAAGGCCAGCAGATCCCTAATTCTCTCTCCAATCAAGTGCGCTCTCCCCAGCCTGTCCCTTCTCCACGGCCACAGTCCCAG\|C\|CCCCCCACTCCAGTCCTTCCCCAAGGATGCAGCCTCAGCCTTCTCCACACC |
| 2:1211:6488:18971#1#F | \| \| |
| MQ=60 isize=149 C=33 | GCCCAGTCCCCACACCTACAAGGCCAGCAGATCCCTAATTCTCTCTCCAATCAAGTGCGCTCTCCCCAGCCTGTCCCTTCTCCACGGCCACAGTCCCAG\|C\|CCCCCCACTCCAGTCCTTCCCCAAGGATGCAGCCTCAGCCTTCTCCACAGG |
| 2:1211:5840:37640#1#F | \| \| |
| MQ=60 isize=251 C=33 | GCCCAGTCCCCACACCTACAAGGCCAGCAGATCCCTAATTCTCTCTCCAATCAAGTGCGCTCTCCCCAGCCTGTCCCTTCTCCACGGCCACAGTCCCAG\|C\|CCCCCCACTCCAGtCCTTCCCCAAGGATGCAGCCTCAGCCTTCTCCACACC |
| 2:1211:16364:61899#1#F | \| \| |
| MQ=60 isize=0 C=33 | GCCCAGTCCCCACACCTACAAGGCCAGCAGATCCCTAATTCTCTCTCCAATCAAGTGCGCTCTCCCCAGCCTGTCCCTTCTCCACGGCCACAGTCCCAG\|C\|CCCCCCACTCCAGTCCTTCCCCAAGGATGCAGCCTCAGCCTAAcCCTGTTG |
| 2:1211:15881:92983#1#F | \| \| |
| MQ=60 isize=138 C=33 | GCCCAGTCCCCACACCTACAAGGCCAGCAGATCCCTAATTCTCTCTCCAATCAAGTGCGCTCTCCCCAGCCTGTCCCTTCTCCACGGCCACAGTCCCAG\|C\|CCCCCCACTCCaGTCCTTCCCCAAGGATGCAGCCTCAGGACTCCAATCAAt |
| 2:1212:9303:38424#1#F | \| \| |
| MQ=60 isize=0 C=33 | GCCCAGTCCCCACACCTACAAGGCCAGCAGATCCCTAATTCTCTCTCCAATCAAGTGCGCTCTCCCCAGCCTGTCCCTTCTCCACGGCCACAGTCCCAG\|C\|CCCCCCACTCCAGTCCTTCCCCAAGGATGCAGCCTCAGCCTTCTCCACACC |
| 2:1212:20148:63065#1#F | \| \| |
| MQ=60 isize=123 C=37 | GCCCAGTCCCCACACCTACAAGGCCAGCAGATCCCTAATTCTCTCTCCAATCAAGTGCGCTCTCCCCAGCCTGTCCCTTCTCCACGGCCACAGTCCCAG\|C\|CCCCCCACTCCAGTCCTTCCCCAGGACTCCAATCATTGTAGGGGGCTGTCT |
| 2:1212:14284:90325#1#F | \| \| |
| MQ=60 isize=168 C=33 | GCCCAGTCCCCACACCTACAAGGCCAGCAGATCCCTAATTCTCTCTCCAATCAAGTGCGCTCTCCCCAGCCTGTCCCTTCTCCACGGCCACAGTCCCAG\|C\|CCCCCCACTCCAGTCCTTCCCCAAGGATGCAGCCTCAGCCTTCTCCACaCC |
| 2:1213:16301:9686#1#F | \| \| |
| MQ=60 isize=144 C=37 | GCCCAGTCCCCACACCTACAAGGCCAGCAGATCCCTAATTCTCTCTCCAATCAAGTGCGCTCTCCCCAGCCTGTCCCTTCTCCACGGCCACAGTCCCAG\|C\|CCCCCCACTCCAGTCCTTCCCCAAGGATGCAGCCTCAGCCTTCTAGGACTC |
| 2:1213:8305:26437#1#F | \| \| |
| MQ=60 isize=269 C=33 | GCCCAGTCCCCACACCTACAAGGCCAGCAGATCCCTAATTCTCTCTCCAATCAAGTGCGCTCTCCCCAGCCTGTCCCTTCTCCACGGCCACAGTCCCAG\|C\|CCCCCCACTCCAGTCCTTCCCCAAGGATGCAGCCTCAGCCTTCTCcacaCC |
| 2:1213:11869:55823#1#F | \| \| |
| MQ=60 isize=121 C=33 | GCCCAGTCCCCACACCTACAAGGCCAGCAGATCCCTAATTCTCTCTCCAATCAAGTGCGCTCTCCCCAgCCTGTCCCTTCTCCACGGCCACAGTCCCAG\|C\|CCCCCCACTCCAGTCCTTCCCAGGACTCCAATGCCTAATGGACCCTGTCTC |
| 2:1213:1785:62075#1#F | \| \| |
| MQ=60 isize=222 C=37 | GCCCAGTCCCCACACCTACAAGGCCAGCAGATCCCTAATTCTCTCTCCAATCAAGTGCGCTCTCCCCAGCCTGTCCCTTCTCCACGGCCACAGTCCCAG\|C\|CCCCCCACTCCAGTCCTTCCCCAAGGATGCAGCCTCAGCCTCCTCCACACC |
| 2:1213:1680:87418#1#F | \| \| |
| MQ=60 isize=127 C=33 | GCCCAGTCCCCACACCTACAAGGCCAGCAGATCCCTAATTCTCTCTCCAATCAAGTGCGCTCTCCCCAGCCTGTCCCTTCTCCACGGCCACAGTCCCAG\|C\|CCCCCCACTCCAGTCCTTCCCCaAGGaGGACTCCAATAACGTATCTCGCCt |
| 2:1214:1604:32669#1#F | \| \| |
| MQ=60 isize=170 C=37 | GCCCAGTCCCCACACCTACAAGGCCAGCAGATCCCTAATTCTCTCTCCAATCAAGTGCGCTCTCCCCAGCCTGTCCCTTCTCCACGGCCACAGTCCCAG\|C\|CCCCCCACTCCAGTCCTTCCCCAAGGATGCAGCCTCAGCCTTCTCCACACC |
| 2:1214:17312:61468#1#F | \| \| |
| MQ=60 isize=186 C=33 | GCCCAGTCCCCACACCTACAAGGCCAGCAGATCCCTAATTCTCTCTCCAATCAAGTGCGCTCTCCCCAGCCTGTCCCTTCTCCACGGCCACAGtCCCAG\|C\|CCCCCCACTCCAGTCCTTCCCCAAGGATGCAGCCTCAGCCTTCTCCACACC |
| 2:1214:9843:75556#1#F | \| \| |
| MQ=60 isize=160 C=37 | GCCCAGTCCCCACACCTACAAGGCCAGCAGATCCCTAATTCTCTCTCCAATCAAGTGCGCTCTCCCCAGCCTGTCCCTTCTCCACGGCCACAGTCCCAG\|C\|CCCCCCACTCCAGTCCTTCCCCAAGGATGCAGCCTCAGCCTTCTCCACACC |
| 2:1215:1551:38837#1#F | \| \| |
| MQ=60 isize=170 C=37 | GCCCAGTCCCCACACCTACAAGGCCAGCAGATCCCTAATTCTCTCTCCAATCAAGTGCGCTCTCCCCAGCCTGTCCCTTCTCCACGGCCACAGTCCCAG\|C\|CCCCCCACTCCAGTCCTTCCCCAAGGATGCAGCCTCAGCCTTCTCCACACC |
| 2:1215:12337:60139#1#F | \| \| |
| MQ=60 isize=121 C=33 | GCCCAGTCCCCACACCTACAAGGCCAGCAGATCCCTAATTCTCTCTCCAATCAAGTGCGCTCTCCCCAGCCTGTCCCTTCTCCACGGCCACAGTCCCAG\|C\|CCCCCCACTCCAGTCCTTCCCAGGACTCCAATGCAATCCCACGGCTGTCTC |
| 2:1215:16948:69598#1#F | \| \| |
| MQ=60 isize=208 C=27 | GCCcAGTcCCCACACCtACAAGGCCAGCAGaTcCCtaaTTCTTTCtcCAATCAAGTGCGCTCTCCCcAgCCTGTCCCTTCtCCACGGCCACAGTCcCAG\|C\|cCCCCCActcCAGtCCTtCCCCAAGGATGCAGCCTCAGCctTCTCCACACC |
| 2:1215:4261:78870#1#F | \| \| |
| MQ=60 isize=140 C=37 | GCCCAGTCCCCACACCTACAAGGCCAGCAGATCCCTAATTCTCTCTCCAATCAAGTGCGCTCTCCCCAGCCTGTCCCTTCTCCACGGCCACAGTCCCAG\|C\|CCCCCCACTCCAGTCCTTCCCCAAGGATGCAGCCTCAGCCAGGACTCCAAT |
| 2:1216:11149:3073#1#F | \| \| |
| MQ=60 isize=266 C=33 | GCCCAGTCCCCACACCTACAAGGCCAGCAGATCCCTAATTCTCTCTCCAATCAAGTGCGCTCTCCCCAGCCTGTCCCTTCTCCACGGCCACAGTCCCAG\|C\|CCCCCCACTCCAGTCCTTCCCCAAGGACGCAGCCTCAGCCTTCTCCACACC |
| 2:1216:16217:18028#1#F | \| \| |
| MQ=60 isize=132 C=33 | GCCCAGTCCCCACACCTACAaGGCCAGCAGATCCCTAATTCtCTCTCCAATCAAGTgCGCTCTCCCCAGCCTGTCCCTTCTCCACGGCcACAGtCCCAG\|C\|CCCCCCACTCCAGTCCTTCCCCAAGGATGCAGGACTCCAATGGGGCCTAGA |
| 2:1216:14916:45624#1#F | \| \| |
| MQ=60 isize=113 C=33 | GCCcAGTCCCCACACCTACAAGGTCAGCAGATCCCTAATTCTCTCTCCAATCAAGTGCgcTCTCCCCaGCCTGTCCCTTCTCCACGgcCACAgtcCCAG\|C\|Ccccccactccgcgactccaacaacgccaccccgcgttttcttatcaaaat |
| 2:1216:18750:95942#1#F | \| \| |
| MQ=60 isize=162 C=33 | GCCCAGTCCCCACACCTACAAGGCCAGCAGATCCCTAATTCTCTCTCCAATCAAGTGCGCTCTCCCCAGCCTGTCCCTTCTCCACGGCCACAGtCCCAG\|C\|CCCCCCACTCCAgtCCttcCCCAAGGATGCAgCCTCAgCCTTCTCCAcaCC |
| 2:2101:19393:42041#1#F | \| \| |
| MQ=60 isize=360 C=33 | GCCCAGTCCCCACACCTACAAGGCCAGCAGATCCCTAATTCTCTCTCCAATCAAGTGCGCTcTCCCCAGCCTGTCcCTTCTCCACGGCCACAGTCCCAG\|C\|CCCCCCACTCCagtcCTTCCCCAaGgATGCAGCCTCAgCCTTCTCCACACC |
| 2:2101:14365:69034#1#F | \| \| |
| MQ=60 isize=121 C=37 | GCCCAGTCCCCACACCTACAAGGCCAGCAGATCCCTAATTCTCTCTCCAATCAAGTGCGCTCTCCCCAGCCTGTCCCTTCTCCACGGCCACAGTCCCAG\|C\|CCCCCCACTCCAGTCCTTCCCAGGACTCCAATGTTCCTAGATGACTGTCTC |
| 2:2102:14168:17702#1#F | \| \| |
| MQ=60 isize=0 C=37 | GCCCAGTCCCCACACCTACAAGGCCAGCAGATCCCTAATTCTCTCTCCAATCAAGTGCGCTCTCCCCAGCCTGTCCCTTCTCCACGGCCACAGTCCCAG\|C\|CCCCCCaCTCCAGTCCTTCCCCaaGGATGCAGCCTCaGCCTTCTCCaCACC |
| 2:2102:3451:50724#1#F | \| \| |
| MQ=60 isize=162 C=2 | GCCCAGTCCCCACACCTACAAGGCcAgCaGATCCCTAATTctCTCtCcAATCAAGtGCGCTCtcCCCAgCctGtcCCTtCtCCacggccacagccccac\|c\|ccccccaccccattccttccccaaggatgcagcctcacccttccccacacc |
| 2:2103:2674:17704#1#F | \| \| |
| MQ=60 isize=312 C=15 | GCCCAGTCCCCACACCTACAAGGCCAGCAGATCCCTAATTCTCTCTCCAATCAAGTGCGCTCTCCCCAGCCTGTCCCTTCTCCaCGGCcccagtcCcac\|c\|CCCCCCactccagccctcccccagggctgcccccccaccttcccccacccc |
| 2:2103:2029:28035#1#F | \| \| |
| MQ=60 isize=133 C=2 | GCCCAGTCCCCACACCTACAAGGCCAGCAGATCCCTAATTCTCTCTCCAATCAAGTGCGCTCTCCCCAgCCTGcCCCTtctccacggccacagtcccag\|c\|cccccccctccagcccttcccaaagaatgcgccaggcccccaaggcgggcc |
| 2:2103:12167:78886#1#F | \| \| |
| MQ=60 isize=278 C=37 | GCCCAGTCCCCACACCTACAAGGCCAGCAGATCCCTAATTCTCTCTCCAATCAAGTGCGCTCTCCCCAGCCTGTCCCTTCTCCACGGCCACAGTCCCAG\|C\|CCCCCCACTCCAGTCCTTCCCCAAGGATGCAGCCTCAGCCTTCTCCacaCC |
| 2:2103:5373:87649#1#F | \| \| |
| MQ=60 isize=233 C=27 | GCCCAGTCCCCACACCTACAAGGCCAGCAGATCCCTAATTCTCTCTCCAATCAAGTGCGCTCTCCCCAGCCTGTCCCTTCTCCACGGCCACAGTCCCAg\|C\|CCCCCCACTCCAgtCCTTCCCCAaGGAtgcggcctcagccttctccccacc |
| 2:2104:2926:11166#1#F | \| \| |
| MQ=60 isize=261 C=15 | GCCCAGTCCCCACACCTACAAGGCCAGCAGATCCCTgaTTaTCTCTcCAATCAAGTGCGCtCTCCCCaGCCTGtCCCTTcTCcACGGCcacagcCCcac\|c\|CCCCCCcctcaagcccttcccaagggttcaacctccacccttctcaacccc |
| 2:2104:11820:76172#1#F | \| \| |
| MQ=60 isize=125 C=15 | GCCCAGTCCCCACACCTACAAGGCCAGCAGATCCCTAATTCTCTCTCCAATCAAGTGCGCTCTCCCCAGCCTGTCCCTTCTCCACGGCCACAGTCCCAG\|c\|CCCCCCACTCCAGTCCTTCCCCAAGAGGACTCCAATAGTaaCAAGATCCTg |
| 2:2104:18723:100452#1#F | \| \| |
| MQ=60 isize=162 C=33 | GCCCAGTCCCCACACCTACAAGGCCAGCAGATCCCTAATTCTCTCTCCAATCAAGTGCGCTCTCCCCAGCCTGTCCCTTCTCCACGGCCACAGTCCCAG\|C\|CCCCCCACTCCAGTCCTTCCCCAaGGATGCAGCCTCAGCCTTCTCCACACc |
| 2:2105:12458:20784#1#F | \| \| |
| MQ=60 isize=481 C=33 | GCCCAGTCCCCACACCTACAAGGCCAGCAGATCCCTAATTCTCTCTCCAATCAAGTGCGCTCTCCCCAGCCTGTCCCTTCTCCACGGCCACAGTCCCAG\|C\|CCCCCCACTCCAGTCCTTCCCCAAGGATGCAGCCTCAGcCTTCTCCAcACC |
| 2:2105:12591:58791#1#F | \| \| |
| MQ=60 isize=162 C=33 | GCCCAGTCCCCACACCTACAAGGCCAGCAGATCCCTAATTCTCTCTCCAATCAAGTGCGCTCTCCCCAGCCTGTCCCTTCTCCACGGCCACAGTCCCAG\|C\|CCCCCCACTCCAGTCCTTCCCCAAGGATGCAGCCTCAGCCTTctcCACACC |
| 2:2105:14589:83682#1#F | \| \| |
| MQ=60 isize=144 C=33 | GCCCAGTCCCCACACCTACAAGGCCAGCAGATCCCTAATTCTCTCTCCAATCAAGTGCGCTCTCCCCAGCCTGTCCCCTCTCCACGGCCACAGTCCCAG\|C\|CCCCCCACTCCAGTCCTTCCCCAAGGATGCAGCCTCAGCCTTCTAGGACTC |
| 2:2106:12370:5820#1#F | \| \| |
| MQ=60 isize=229 C=33 | GCCCAGTCCCCACACCTACAAGGCCAGCAGATCCCTAATTCTCTCTCCAATCAAGTGCGCTCTCCCCAGCCTGTCCCTTCTCCACGGCCACAGTCCCAG\|C\|CCCCCCACTCCAGTCCTTCCCCAAGGATGCAGCCTCAGCCTTCTCCaCACC |
| 2:2106:9548:18193#1#F | \| \| |
| MQ=60 isize=207 C=22 | gCCCAGTCCCCaCACCTACAAGGCCAGCAGATCCCTAATTCTCTCTCCAATCAAGTGCGCTCTCCCCAGCCTGTCcCTTCTCCACGGCCACAgtCCCag\|C\|CcCCCCACTCCagTCCTTCCCCAagGatGCAgCCTCAGCCTtccccacccc |
| 2:2106:17220:23273#1#F | \| \| |
| MQ=60 isize=293 C=37 | GCCCAGTCCCCACACCTACAAGGCCAGCAGATCCCTAATTCTCTCTCCAATCAAGTGCGCTCTCCCCAGCCTGTCCCTTCTCCACGGCCACAGTCCCAG\|C\|CCCCCCACTCCAGtCCTTCCCCAAGGATGCAGCCTCAGCCTTCTCCACACC |
| 2:2106:11347:67319#1#F | \| \| |
| MQ=60 isize=131 C=33 | GCCCAGTCCCCACACCTACAAGGCCAGCAGATCCCTAATTCTCTCTCCAATCAAGTGCGCTCTCCCCAGCCTGTCCCTTCTCCACGGCCACAGTCCCAG\|C\|CCCCCCACTCCAGTCCTTCCCCAAGGATGCAGGACTCCAATCAACTCCGCC |
| 2:2106:18012:85175#1#F | \| \| |
| MQ=60 isize=133 C=33 | GCCCAGTCCCCACACCTACAAGGCCAGCAGATCCCTAATTCTCTCTCCAATCAAGTGCGCTCTCCCCAgCCTGTCCCTTCTCCACGGCCACAGTCCCAG\|C\|CCCCCCACTCCAGTCCTTCCCCAAGGATGCAGCAGGACTCCAATAATCACA |
| 2:2107:1705:2465#1#F | \| \| |
| MQ=60 isize=166 C=27 | GCCCAGTCCCCACACCTACAAGGCCAGCAGATCCCTAATTCTCTCTCCAATCAAGTGCGCTCTCCCCAgCCTGcCCCTTCTCCACGgCCaCAGtCCCAg\|C\|CCCCCCcCtCcaGtccTtCCCCAaggatgcagcctccgccttctccacccc |
| 2:2107:16049:32941#1#F | \| \| |
| MQ=60 isize=160 C=27 | GCCCAGTCCCCACACCTACAAGGCCAGCAGATCCCTAATTCTCTCTCCAATCAAGTGCGCTCTCCCCAGCCTGTCCCTTCTCCACGGCCACAGTCCCAg\|C\|CCCCCCACTCCAGTCCTTCCCCAAGGATGCAGCCtCaGCCTTCTCCACACC |
| 2:2107:5350:36715#1#F | \| \| |
| MQ=60 isize=177 C=27 | GCCCAGTCCCCACACCTACAAGGCCAGCAGATCCCTAATTCTCTCTCCAATCAAGTGCGCTCTCCCCAGCCTGTCCCTTCTCCACGGCCACAGtCCCAG\|C\|CCCCCCACcCCagccCTTCCCCAaGGATgCAcCCtcagccttctccacacc |
| 2:2107:4942:54138#1#F | \| \| |
| MQ=60 isize=207 C=22 | GCCCAGTCCCCACACCTACAAGGCCAGCAGATCCCTAATTCTCTCTCCAaTCAAGTGCGCTcTCCCCAGCCTGTCCCTTCTCCACGgCCaCAGtcCCAg\|C\|CcCCCcaCtCCagtCCTtCCCCAaggatccggcctcagccttctccacacc |
| 2:2107:1590:78807#1#F | \| \| |
| MQ=60 isize=143 C=27 | GCCCAGTCCCCACACCTACAAGGCCAGCAGATCCCTAATTCTCTCTCCAATCAAGTGCGCTCTCCCCAGCCTGTCCcTTcTCcACGGCCAcaGtCCCag\|C\|CCCCCCACcCCagcCcTtCCCCAgggtggcagcctcacccttcagaccccc |
| 2:2108:8717:20178#1#F | \| \| |
| MQ=60 isize=192 C=33 | GCCCAGTCCCCACACCTACAAGGCCAGCAGATCCCTAATTCTCTCTCCAATCAAGTGCGCTCTCCCCAGCCTGTCCCTTCTCCACGGCCACAgTCCCAG\|C\|CCCCCCACTCCAGTCCTTCCCCAAGgATGCAGCCTCAGCCTtcTCCaCaCC |
| 2:2108:2728:31544#1#F | \| \| |
| MQ=60 isize=120 C=33 | GCCCAGTCCCCACACCTACAAGGCCAGCAGATCCCTAATTCTCTCTCCAATCAAGTGCGCTCTCCCCAGCCTGTCCCTTCTCCACGGCCaCAGtCCCAG\|C\|CCCCCCACTCCAGccCTTCCAGGACtCCAatCCagactatgtcctgtctct |
| 2:2108:15620:59816#1#F | \| \| |
| MQ=60 isize=123 C=33 | GCCCAGTCCCCACACCTACAAGGCCAGCAGATCCCTAATTCTCTCTCCAATCAAGTGCGCTCTCCCCAGCCTGTCCCTTCTCCACGGCCACAGTCCCAG\|C\|CCCCCCACTCCAGTCCTTCCCCaGGGCTCCAATCGTGCCCGaATCCTGTCc |
| 2:2108:2516:97703#1#F | \| \| |
| MQ=60 isize=250 C=27 | GCCCAGTCCCCACACCTACAAGGCCAGCAGATCCCTAATTCTCTCTCCAATCAAGTGCGCTCTCCCCAGCCTGTCCCTTCTCCACGGCCACAGtCCCag\|C\|CCCCCCACTCCagtCcTTCCCCAaGgatgaagcctcagccttctccaaccc |
| 2:2109:14690:5107#1#F | \| \| |
| MQ=60 isize=325 C=37 | GCCCAGTCCCCACACCTACAAGGCCAGCAGATCCCTAATTCTCTCTCCAATCAAGTGCGCTCTCCCCAGCCTGTCCCTTCTCCACGGCCACAGTCCCAG\|C\|CCCCCCaCTCCAGTCCTTCCCCAAGGATGCAGCCTCAGCCTTCTCCACACC |
| 2:2109:2602:38808#1#F | \| \| |
| MQ=60 isize=187 C=27 | GCCCAGTCCCCACACCTACAAGGCCAGCAGATCCCTAATTCTCTCTCCAATCAAGTGCGCTCTCCCCAGCCTGTCCCTTCTCCaCGGCCACagtCCCAg\|C\|CCCCCCaCtCCAgtcCTtCCCCAAGgatgcagcctcagccttctccacacc |
| 2:2109:2379:86524#1#F | \| \| |
| MQ=60 isize=125 C=15 | GcCcAGTCCCCACACCTACAAGGCCAGCAGATCCCTAATTCTCtCTCCAATcAAGTGCGCTCTCCCCAGCcTGTCCCTTCTCCACGGCCACAggcCcAA\|c\|ccCCCcccccCAGtCCTtCCCcaagaggccccaattcaccccgccccctat |
| 2:2109:20508:94608#1#F | \| \| |
| MQ=60 isize=156 C=33 | GCCCAGTCCCCACACCTACAAGGCCAGCAGATCCCTAATTCTCTCTCCAATCAAGTGCGCTCTCCCCAGCCTGTCCCTTCTCCACGGCCACAGTCCCAG\|C\|cCCCCCACTCCAGTCCTTCCCCAAgGATGCAGCCTCAgCCTTCTCCACACC |
| 2:2110:4514:8025#1#F | \| \| |
| MQ=60 isize=147 C=33 | GCCCAGTCCCCACACCTACAAGGCCAGCAGATCCCTAATTCTCTCTCCAATCAGGTGCGCTCTCCCCAGCCTGTCCCTTCTCCACGGCCACAGTCCCAG\|C\|CCCCCCACtCCAGTCCTTCCCCAAGGAtgCAgCCtCagCCTTCtCCAGgAC |
| 2:2110:11389:10083#1#F | \| \| |
| MQ=60 isize=139 C=37 | GCCCAGTCCCCACACCTACAAGGCCAGCAGATCCCTAATTCTCTCTCCAATCAAGTGCGCTCTCCCCAGCCTGTCCCTTCTCCACGGCCACAGTCCCAG\|C\|CcCCCCACTCCAGTCCTTCCCCAAGGATGCAGCCTCAGCAGGaCTCCAATC |
| 2:2110:4219:27722#1#F | \| \| |
| MQ=60 isize=234 C=33 | GCCCAGTCCCCACACCTACAAGGCCAGCAGATCCCTAATTCTCTCTCCAATCAAGTGCGCTCTCCCCAGCCTGTCCCTTCTCCACGGCCACAGTCCCAg\|C\|CCCCCCacTCCAGTCCTtCCCCAaGGATGCAGCCtcagCCTTCtCcacacc |
| 2:2110:9945:60189#1#F | \| \| |
| MQ=60 isize=262 C=37 | GCCCAGTCCCCACACCTACAAGGCCAGCAGATCCCTAATTCTCTCTCCAATCAAGTGCGCTCTCCCCAGCCTGTCCCTTCTCCACGGCCACAGTCCCAG\|C\|CCCCCCACTCCAGTCCTTCCCCAAGGATGCAGCCTCAGCCTTCTCCACACC |
| 2:2110:20671:86331#1#F | \| \| |
| MQ=60 isize=207 C=33 | GCCCAGTCCCCACACCTACAAGGCCAGCagGtcCCTAAtTCTttcTCCAAtCAAGTGCGCTCTCCCCaGCCTGTcCCTTCTCCACGgccACAGtCcCaG\|C\|ccCCCCACtCCagtCCTTCCCCAAGgAcgcAGCCtCAGCCTTcTCCACACC |
| 2:2110:16836:89793#1#F | \| \| |
| MQ=60 isize=239 C=33 | GCCCAGTCCCCACACCTACAAGGCCAGCAGATCCCTAATTCTCTCTCCAATCAAGTGCGCTCTCCCCAGCCTGTCCCTTCTCCACGgCCaCAGTCCCAG\|C\|ccCCCCActCCAGtCCTTCCCCAAGGATgCagCCTCAGCCTtCTCCACACC |
| 2:2111:4006:9313#1#F | \| \| |
| MQ=60 isize=127 C=22 | GCCCAGTCCCCACACCTACAAGGCCAGCAGATCCCTAATTCTCTCTCCAATCAAGTGCGCTCTCCCCAGCCTGTCCCTTCTCCaCGGCCaCAGtCCCAg\|C\|CCCCCCactCcagtccTtCCCCAAGGAGGaCTCCAATCCCTACGGAgcgct |
| 2:2111:2541:18600#1#F | \| \| |
| MQ=60 isize=169 C=27 | GCCCAGTCCCCACACCTACAAGGCCAGCAGATCCCTAATTCTCTCTCCAATCAAGTGCGCTCTCCCCAGCCTGTCCCTTCTCCACGGCCACaGtCCCAG\|C\|CCCCCCactCCAGtCCTTCCCCAAGgatgcagcctcagccttctccacacc |
| 2:2111:11510:33808#1#F | \| \| |
| MQ=60 isize=201 C=37 | GCCCAGTCCCCACACCTACAAGGCCAGCAGATCCCTAATTCTCTCTCCAATCAAGTGCGCTCTCCCCAGCCTGTCCCTTCTCCACGGCCACAGTCCCAG\|C\|CCCCCCACTCCAGTCCTTCCCCAAGGATGCAGCCTCAGCCTTCTCCACACC |
| 2:2111:9735:58039#1#F | \| \| |
| MQ=60 isize=0 C=37 | GCCCAGTCCCCACACCTACAAGGCCAGCAGATCCCTAATTCTCTCTCCAATCAAGTGCGCTCTCCCCAGCCTGTCCCTTCTCCACGGCCACAGTCCCAG\|C\|CCCCCCACTCCAGTCCTTCCCCAAGGATGCCGTCAATGGGTTCCATGGCCT |
| 2:2111:5278:78657#1#F | \| \| |
| MQ=60 isize=132 C=33 | GCCCAGTCCCCACACCTACAAGGCCAGCAGATCCCTAATTCTCTCTCCAATCAAGTGCGCTCTCCCCAGCTTGTCCCTTCTCCACGGCCACAGtcCCag\|C\|CCCCCCACtCCcgtcCTTCCCCaAGGATGCagAGGaCtCCAATTATtatcc |
| 2:2111:16253:97619#1#F | \| \| |
| MQ=60 isize=235 C=37 | GCCCAGTCCCCACACCTACAAGGCCAGCAGATCCCTAATTCTCTCTCCAATCAAGTGCGCTCTCCCCAGCCTGTCCCTTCTCCACGGCCACAGTCCCAG\|C\|CCCCCCACTCCaGTCCTTCCCCAAGGATGCAGCCTCAgCCTTCTCcacacc |
| 2:2112:18056:34058#1#F | \| \| |
| MQ=60 isize=350 C=22 | GCCCAGTCCCCACACCTACAAGGCCAGCAGATCCCTAATTCTCTCTCCAATCAAGTGCGCTCTCCCCAGCCTGTCCCTTCTCCACGGCCACAGTCcCAG\|C\|CcCCCCACTCCAGTCCTTCCCccaggatgCAgCCTCAGCcTtCTCCaCaCC |
| 2:2113:1637:4526#1#F | \| \| |
| MQ=60 isize=127 C=22 | GCCCAGTCCCCACACCTACAAGGCCAGCAGATCCCTAATTCTCTCTCCAATCAAGTGCGCTCTCCCCAGCCTGTCCCTTCTCCACGgCCACAGtcCCAg\|C\|CCCCCCcctcCAGtCCTtCCCCAAGgaggactccatcatctgtcccccctt |
| 2:2113:3033:19175#1#F | \| \| |
| MQ=60 isize=120 C=33 | GCCCAGTCCCCACACCTACAAGGCCAGCAGATCCCTAATTCTCTCTCCAATCAAGTGCGCTCTCCCCAGCCTGTCCCTTCTCCACGGCCACAGTCCCAG\|C\|CCCCCCaCtCCAGtCCTTCCAGGACTCCAATGAAAAAATTGgCCtgtctcT |
| 2:2113:12676:59368#1#F | \| \| |
| MQ=60 isize=216 C=33 | GCCCAGTCCCCACACCTACAAGGCCAGCAGATCCCTAATTCTCTCTCCAATCAAGTGCGCTCTCCCCAGCCTGTCCCTTCTCCACGGCCACAGTCCCAG\|C\|CCCCCCACTCCAGTCCTTCCCCAAGGATGCAGCCTCAGCCTTCTCCACACC |
| 2:2113:7290:80799#1#F | \| \| |
| MQ=60 isize=187 C=33 | GCCCAGTCCCCACACCTACAAGGCCAGCAGATCCCTAATTCTCTCTCCAATCAAGTGCGCTCTCCCCAGCCTGTCCCTTCTCCACGGCCACAGTCCCAG\|C\|CCCCCCACTCCAGTCCTtCCCCAAGGATGCagcCtCAGCCTTcTCCACACC |
| 2:2113:5337:89936#1#F | \| \| |
| MQ=60 isize=131 C=37 | GCCCAGTCCCCACACCTACAAGGCCAGCAGATCCCTAATTCTCTCTCCAATCAAGTGCGCTCTCCCCAGCCTGTCCCTTCTCCACGGCCACAGTCCCAG\|C\|CCCCCCACTCCAGTCCTTCCCCAAGGATGCAAGGACTCCAATCGTTCGGtc |
| 2:2114:14156:15603#1#F | \| \| |
| MQ=60 isize=138 C=33 | GCCCAGTCCCCACACCTACAAGGCCAGCAGATCCCTAATTCTCTCTCCAATCAAGTGCGCTCTCCCCAGCCTGTCCCTTCTCCACGGCCACAGTCCCAG\|C\|CCCCCCACTCCAGTCCTTCCCCAAGGATGCAGCCTCAGAGGACTCCAATAA |
| 2:2114:1589:48560#1#F | \| \| |
| MQ=60 isize=289 C=33 | GCCCAGTCCCCACACCTACAAGGCCAGCAGATCCCTAATTCTCTCTCCAATCAAGTGCGCTCTCCCCAGCCTGTCCCTTCTCCACGGCCACAGTCCCAG\|C\|CCCCCCaCtCCAGTCcTtCCCcaaggatgcagcctcagccttctccacacc |
| 2:2114:1401:77164#1#F | \| \| |
| MQ=60 isize=0 C=33 | GCCCAGTCCCCACACCTACAAGGCCAGCAGATCCCTAATTCTCTCTCCAATCAAGTGCGCTCTCCCCAGCCTGTCCCTTCTCCaCGGCCacAgtCCCag\|C\|CCCCCCaCtCCaGtCCTtCCCCAAGgatgcagcctcacccttctccaaccc |
| 2:2114:17225:95446#1#F | \| \| |
| MQ=60 isize=185 C=33 | GCCCAGTCCCCACACCTACAAGGCCAGCAGATCCCTAATTCTCTCTCCAATCAAGTGCGCTCTCCCCAGCCTGTCCCTTCTCCACGGCCACAGTCCCAG\|C\|CCCCCCACTCCAGTCCTTCCCCAAGGATGCAGCCTCAGCCTTCTCCACACC |
| 2:2115:4873:66275#1#F | \| \| |
| MQ=60 isize=134 C=33 | GCcCAGTCCCCACACCTACAAGGCCAGCAGATCCCTAATTCTCTCTCCAATCAAGTGCGCTCTCCCCAGCCTGTCCCTTCTCCACGGCCACAGtCCcAG\|C\|CCCCCCACTCCAgtCCTTCCCCAAGGATGcagCCAGGACtcCAATCGgata |
| 2:2115:17696:77687#1#F | \| \| |
| MQ=60 isize=212 C=33 | GCCCAGTCCCCACACCTACAAGGCCAGCAGATCCCTAATTCTCTCTCCAATCAAGTGCGCTCTCCCCAGCCTGTCCCTTCTCCACGGCCACAGTCCCAG\|C\|CCCCCCACTCCAGTCCTTCCCCaAGGATGCAGCCTCAGCCTTCTCCACACC |
| 2:2116:8918:7201#1#F | \| \| |
| MQ=60 isize=205 C=15 | GCCCAGTCCCCACACCTACAAGGCCAGCAGATCCCTAATTCTCTCTCCAATCAAGTGCGCTCTCCCCAGCCTGTCCCTTCTCCACGGCCACAGTCCCAG\|c\|CCCCCCACTCCAGTCCTTCCCCAAGGATGCAGCCTCAGCCTTCTCCACACC |
| 2:2116:15257:48826#1#F | \| \| |
| MQ=60 isize=0 C=33 | GCCCAGTCCCCACACCTACAAGGCCAGCAGATCCCTAATTCTCTCTCCAATCAAGTGCGCTCTCCCCAGCCTGTCCCTTCTCCACGGCCACAGTCCCAG\|C\|CCCCCCACTCCAGTCCTTCCCCAAGGATGCAGCCTCAGCCTTCTCCACACC |
| 2:2116:12778:74983#1#F | \| \| |
| MQ=60 isize=147 C=33 | GCCCAGTCCCCACACCTACAAGGCCAGCAGATCCCTAATTCTCTCTCCAATCAAGTGCGCTCTCCCCAGCCTGTCCCTTCTCCACGGCCACAGTCCCAG\|C\|CCCCCCACTCCAGTCCTTCCCCAAGGATGCAGCCTCAGCCTTCTCCAGGAC |
| 2:2201:3250:10552#1#F | \| \| |
| MQ=60 isize=160 C=33 | GCCCAGTCCCCACACCTACAAGGCCAGCAGATCCCTAATTCTCTCTCCAATCAAGTGCGCTCTCCCCAGCCTGTCCCTTCTCCACGGCCACAGTCCCAG\|C\|CCCCCCACTCCAGTCCTTCCCCAAGgATGCAgCCTCAgCCTTCtCCacaCC |
| 2:2201:10899:41382#1#F | \| \| |
| MQ=60 isize=150 C=37 | GCCCAGTCCCCACACCTACAAGGCCAGCAGATCCCTAATTCTCTCTCCAATCAAGTGCGCTCTCCCCAGCCTGTCCCTTCTCCACGGCCACAGTCCCAG\|C\|CCCCCCACTCCAGTCCTTCCCCAAGGATGCAGCCTCAGCCTTCTCCACACA |
| 2:2201:3203:62616#1#F | \| \| |
| MQ=60 isize=140 C=37 | GCCCAGTCCCCACACCTACAAGGCCAGCAGATCCCTAATTCTCTCTCCAATCAAGTGCGCTCTCCCCAGCCTGTCCCTTCTCCACGGCCACAGTCCCAG\|C\|CCCCCCTCTCCAGTCCTTCCCCAAGGATGCAGCCTCAGCCAGGACTCCAAT |
| 2:2201:6504:89323#1#F | \| \| |
| MQ=60 isize=240 C=37 | GCCCAGTCTCCACACCTACAAGGCCAGCAGATCCCTAATTCTCTCTCCAATCAAGTGCGCTCTCCCCAGCCTGTCCCTTCTCCACGGCCACAGTCCCAG\|C\|CCCCCCACTCCAGTCCTTCCCCAAGGATGCAGCCTCAGCCTTCTCCACACC |
| 2:2202:4153:24731#1#F | \| \| |
| MQ=60 isize=350 C=37 | GCCCAGTCCCCACACCTACAAGGCCAGCAGATCCCTAATTCTCTCTCCAATCAAGTGCGCTCTCCCCAGCCTGTCCCTTCTCCACGGCCACAGTCCCAG\|C\|CCCCCCACTCCAGTCCTtCCCCAAGGAtgcaGCCTCAGCCTtCTCcacACC |
| 2:2202:17538:79285#1#F | \| \| |
| MQ=60 isize=137 C=33 | GCCCAGTCCCCACACCTACAAGGCCAGCAGATCCCTAATTCTCTCTCCAATCAAGTGCGCTCTCCCCAGCCTGTCCCTTCTCCACGGCCACAGTCCCAG\|C\|CCCCCCACTCCAGTCCTTCCCCAAGGATGCAGCCTCAAGGACtCCaaTGCC |
| 2:2202:13555:83007#1#F | \| \| |
| MQ=60 isize=159 C=37 | GCCCAGTCCCCACACCTACAAGGCCAGCAGATCCCTAATTCTCTCTCCAATCAAGTGCGCTCTCCCCAGCCTGTCCCTTCTCCACGGCCACAGTCCCAG\|C\|CCCCCCACTCCAGTCCTTCCCCAAGGATGCAGCCTCAGCCTTCTCCACACC |
| 2:2203:6609:13653#1#F | \| \| |
| MQ=60 isize=135 C=33 | GCCCAGTCCCCACACCTACAAGGCCAGCAGATCCCTAATTCTCTCTCCAATCAAGTGCGCTCTCCCCAGCCTGTCCCTTCTCCACGGCcACAGTCCCAG\|C\|CCCCCCACTCCAGTCCTTCCCCAAGGATGCAGCCTAGGACTCCAATACAGC |
| 2:2203:9067:34326#1#F | \| \| |
| MQ=60 isize=127 C=37 | GCCCAGTCCCCACACCTACAAGGCCAGCAGATCCCTAATTCTCTCTCCAATCAAGTGCGCTCTCCCCAGCCTGTCCCTTCTCCACGGCgaCAGTCCCAG\|C\|CCCCCCACTCCAGTCCTTCCCCAAGGACTCCAATTGACGGCAGGgCCTGTC |
| 2:2203:5660:54104#1#F | \| \| |
| MQ=60 isize=194 C=33 | GCCCAGTCCCCACACCTACAAGGCCAGCAGATCCCTAATTCTCTCTCCAATCAAGTGCGCTCTCCCCAGCCTGTCCCTTCTCCACGGCcACAGTCCCAG\|C\|CCCCCCACTCCAGTCCTTCCCCAAGGATGCAGCCtCAGCCTTCTCCACACC |
| 2:2203:10239:67940#1#F | \| \| |
| MQ=60 isize=160 C=33 | GCCCAGTCCCCACACCTACAAGGCCAGCAGATCCCTAATTCTCTCTCCAATCAAGTGCGCTCTCCCCAGCCTGTCCCTTCTCCACGGCCACAGTCCCAG\|C\|CCCCCCACTCCAGTCCTTCCCCAAGGATGCAGCCTCAGCCTTtTCCACACC |
| 2:2203:10877:82913#1#F | \| \| |
| MQ=60 isize=251 C=27 | GCCCAGTCCCCACACCTACAAGGCCAGCAGATCCCTAATTCTCTCTCCAATCAAGTGCGCTCTCCCCAGCCTGTCCCTTCTCCACGGCcACAGTCCCAG\|C\|CCCCCCACTCCAGTCCTTCCCCAAGGATGCAGCCTCAGCCTTCTCCACACC |
| 2:2204:3897:26454#1#F | \| \| |
| MQ=60 isize=0 C=2 | GCCCAGTCCCCACACCTaCAAGGCCAGCAGATCCCTaATtCtCTCTCCAATCaAgTGCGCtcTCcccagcctgtcccttctccacggccacagcccccg\|c\|ccccccgcacccaaccttgcccaaagatggggcccctgccctctccccccc |
| 2:2204:1974:46175#1#F | \| \| |
| MQ=60 isize=127 C=33 | GCCCAGTCCCCACACCTACAAGGCCAGCAGATCCCTAATTCTCTCTCCAATCAAGTGCGCTCTCCCCAGCCTGTCCCTTCTCCACGGCCACAGTCCCAG\|C\|CCCCCCACTCCAGTCCTTCCCCAAGGAGGaCTCCAATCACTAGCCCTCACt |
| 2:2204:9341:70189#1#F | \| \| |
| MQ=60 isize=205 C=15 | GCCCAGTCCCCACACCTACAAGGCCAGCAGATCCCTAATTCTCTCTCCAATCAAGTGCGCTCTCCCCAGCCTGTCCCTTCTCCACGGCCACAGTCCCAG\|c\|CCCCCCACTCCAGTCCTTCCCCAAGGATGCAGCCTCAGCCTTCTCCACACC |
| 2:2205:11712:37312#1#F | \| \| |
| MQ=60 isize=189 C=33 | GCCCAGTCCCCACACCTACAAGGCCAGCAGATCCCTAATTCTCTCTCCAATCAAGTGCGCTCTCCCCAGCCTGTCCCTTCTCCACGGCCACAGTCCCAG\|C\|CCCCCCACTCCAGTCCTTCCCCAAGgATGCAGCCTCAGCCTTCTCCACACC |
| 2:2205:4713:46156#1#F | \| \| |
| MQ=60 isize=160 C=37 | GCCCAGTCCCCACACCTACAAGGCCAGCAGATCCCTAATTCTCTCTCCAATCAAGTGCGCTCTCCCCAGCCTGTCCCTTCTCCACGGCCACAGTCCAAG\|C\|CCCCCCACTCCAGTCCTTCCCCAAGGATGCAGCCTCAGCCTTCTCCACACC |
| 2:2205:7348:69271#1#F | \| \| |
| MQ=60 isize=175 C=33 | GCCCAGTCCCCACACCTACAAGGCCAGCAGATCCCTAGTTCTCTCTCCAATCAAGTGCGCTCTCCCCAGCCTGTCCCTTCTCCACGGCCACAGTCCCtg\|C\|CCCCCCACTCCAGTCCTTCCCCAAGGATGCAGCCTCAGTCTTCTCCACACC |
| 2:2205:15696:95697#1#F | \| \| |
| MQ=60 isize=133 C=2 | GCCCAGTCCCCACACCTACAAGGCCAGCAGATCCCTAATTCTCTCTCCAATCAAGTGCGCTCTCCCCAGCCTGTCCCTTCTCCACGGCCACAGTCCCag\|c\|ccccccactccaatccttcccccaagatgccgcaagactcccacccccccc |
| 2:2206:10589:11859#1#F | \| \| |
| MQ=60 isize=207 C=33 | GCCCAGTCCCCACACCTACAAGGCCAGCAGATCCCTAATTCTCTCTCCAATCAAGTGCGCTCTCCCCAGCCTGTCCCTTCTCCACGGCCACAGTCCCAG\|C\|CCCCCCACTCCAGTCCTTCCCCAAGGATGCAgcCTCAGCCTTCTCCACACC |
| 2:2206:17168:22278#1#F | \| \| |
| MQ=60 isize=251 C=37 | GCCCAGTCCCCACACCTACAAGGCCAGCAGATCCCTAATTCTCTCTCCAATCAAGTGCGCTCTCCCCAGCCTGTCCCTTCTCCACGGCCACAGTCCCAG\|C\|CCCCCCACTCCAGTCCTTCCCCAAGGATGCAgcctCAGCCTTCTCCACACC |
| 2:2206:17693:53445#1#F | \| \| |
| MQ=60 isize=307 C=33 | GCCCAGTCCCCACACCTACAAGGCCAGCAGATCCCTAATTCTCTCTCCAATCAAGTGCGCTCTCCCCAGCCTGTCCCTTCTCCACGGCCACAGTCCCAG\|C\|CCCCCCACTCCAGTCCTTCCCCAAGGATGCAGCCTCAGCCTTCTCCACACC |
| 2:2206:15850:84480#1#F | \| \| |
| MQ=60 isize=175 C=37 | GCCCAGTCCCCACACCTACAAGGCCAGCAGATCCCTAATTCTCTCTCCAACCAAGTGCGCTCTCCCCAGCCTGTCCCTTCTCCACGGCCACAGTCCCAG\|C\|CCCCCCACTCCAGTCCTTCCCCAAGGATGCAGCCgCAGCCTTCTCCACACC |
| 2:2207:4209:6170#1#F | \| \| |
| MQ=60 isize=187 C=33 | GcCCAGTCCCCACACCTACAAGGCCAGCAGATCCCTAATTCTCTCTCCAATCAAGTGCGCTCTCCCCAGCCTGTCCCTTCTCCACGGCCACaGTCCCAG\|C\|CCCCCCaCTCCAGtcCTtcCCCAaGGatGCAGcCTcAgCCTTCtccacacc |
| 2:2207:19132:24476#1#F | \| \| |
| MQ=60 isize=134 C=37 | GCCCAGTCCCCACACCTACAAGGCCAGCAGATCCCTAATTCTCTCTCCAATCAAGTGCGCTCTCCCCAGCCTGTCCCTTCTCCACGGCCACAGTCCCAG\|C\|CCCCCCACTCCAGTCCTTCCCCAAGGATGCAGCCAGGACTCCAATTCCGCt |
| 2:2207:10087:43817#1#F | \| \| |
| MQ=60 isize=319 C=15 | GCCCAGTCCCCACACCTACAAGGCCAGCAGATCCCTAATTCTCTCTCCAATCAAGTGCGCTCTCCCCAGCCTGTCCCTTCTCCACGGCCACAGTCCCAg\|c\|ccCCCCACTCCAGTCCTTCCCCAAGGATGCAGCCTCAGCCTTCTCCACACC |
| 2:2207:5493:72228#1#F | \| \| |
| MQ=60 isize=198 C=33 | GCCCAGTCCCCACACCTACAAGGCCAGCAGATCCCTAATTCTCTCTCCAATCAAGTGCGCTCTCCCCAGCCTGTCCCTTCTCCACGGCCACAGTCCCAG\|C\|CCCCCCACTCCAGTCCTTCCCCAAGGATGCAGCCTCAGCCTTCTCCACACC |
| 2:2208:9445:13153#1#F | \| \| |
| MQ=60 isize=262 C=33 | GCCCAGTCCCCACACCTACAAGGCCAGCAGATCCCTAATTCTCTCTCCAATCAAGtGCGCTCTCcCCAGCCTGTCCCTTCTCCACGGCCACAGtCCCAG\|C\|CCCCCCACTCCAGtcCTTCCCCaaggATcCAGcCtCAGCCttctccacacc |
| 2:2208:7601:42914#1#F | \| \| |
| MQ=60 isize=236 C=33 | GCCCAGTCCCCACACCTACAAGGCCAGCAGATCCCTAATTCTCTCTCCAATCAAGTGCGCTCTCCCCAGCCTGTCCCTTCTCCACGGCCACAGTCCCAG\|C\|CCCCCCaCTCCAGTcCTTCCCCAAGGATGCAGCCTCAGCCTtCTCCACACC |
| 2:2208:9802:59047#1#F | \| \| |
| MQ=60 isize=185 C=27 | GCCCAATCCCCACACCTACAAGGCCAGCAGATCCCTAATTCTCTCTCCAATCAAGTGCGCTCTCCCCAGCCTGTCCCTTCTCCACGGCCACAGtCCCAg\|C\|CCCCCCACTCCAGTCCTTCCCCAAGGATGCAGCCTCAGCCTTCTCCACaCC |
| 2:2209:20565:10805#1#F | \| \| |
| MQ=60 isize=139 C=33 | GCCCAGTCCCCACACCTACAAGGCCAGCAGATCCCTAATTCTCTCTCCAATCAAGTGCGCTCTCCCCAGCCTGTCCCTTCTCCACGGCCACAGTCCCAG\|C\|CCCCCCACTCCAGTCCTTCCCCAAGGATGCAGCCTCAGCAGGACTCCAATT |
| 2:2209:16670:45509#1#F | \| \| |
| MQ=60 isize=120 C=37 | GCCCAGTCCCCACACCTACAAGGCCAGCAGATCCCTAATTCTCTCTCCAATCAAGTGCGCTCTCCCCAGCCTGTCCCTTCTCCACGGCCACAGTCCCAG\|C\|CCCCCCACTCCAGTCCTTCCAGGACTCCAATCGACCATTTTGCCTGTCTCT |
| 2:2209:9042:69858#1#F | \| \| |
| MQ=60 isize=139 C=27 | GCcCAGTCCCCACACCTACAAgGcCAGCAGATCCCTAATTCTCTCTcCAATCAAGtGCGCTCTCCCCAGCCtGTCCCTTCTCCACggcCACAGtCCCAg\|C\|CcCCCCACTCCAGtcCTTCCCCAAGgatgcAgcCtCAgCAGgACtCCAATA |
| 2:2210:7146:47459#1#F | \| \| |
| MQ=60 isize=124 C=33 | GCCCAGTCCCCACACCTACAAGGCCAGCAGATCCCTAATTCTCTCTCCAATCAAGTGCGCTCTCCCCAGCCTGTCCCTTCTCCACGGCCACAGTCCCAG\|C\|CCCCCCACTCCAGTCCTTCCCCAAAGGACTCCAATCGACgCCCCACCCTGt |
| 2:2210:2397:55353#1#F | \| \| |
| MQ=60 isize=194 C=37 | GCCCAGTCCCCACACCTACAAGGCCAGCAGATCCCTAATTCTCTCTCCAATCAAGTGCGCTCTCCCCAGCCTGTCCCTTCTCCACGGCCACAGTCCCAG\|C\|CCCCCCACTCCAGTCCTTCCCCAAGGATGCAGCCTCAGCCTTCTCCACACC |
| 2:2210:6507:90433#1#F | \| \| |
| MQ=60 isize=234 C=37 | GCCCAGTCCCCACACCTACAAGGCCAGCAGATCCCTAATTCTCTCTCCAATCAAGTGCGCTCTCCCCAGCCTGTCCCTTCTCCACGGCCACAGTCCCAG\|C\|CCCCCCACTCCAGTCCTTCCCCAAGGATGCAGCCTCAGCCTTCTCCaCACC |
| 2:2211:12709:2995#1#F | \| \| |
| MQ=60 isize=233 C=22 | GCCCAGTCCCCACACCTACAAGGCCAGCAGATCCCTAATTCTCTCTCCAATCAAGTGCGCTCTCCCCAGCCTGTCCCTTCTCCACGGCCACAGTCCCAG\|C\|CCCCCCACTCCAGTCCTTCCCCAAGGAtGCAGCCtcAgCCTTCTCCACACC |
| 2:2211:6611:50793#1#F | \| \| |
| MQ=60 isize=402 C=33 | GCCCAGTCCCCACACCTACAAGGCCAGCAGATCCCTAATTCTCTCTCCAATCAAGTGCGCTCTCCCCAGCCTGTCCCTTCTCCACGGCCACAGTCCCAG\|C\|CCCCCCaCTCCAGTCCTTCCCCAAGGATGCAGCCTCAGCCTTCTCCACACc |
| 2:2211:15500:69750#1#F | \| \| |
| MQ=60 isize=162 C=33 | GCCCAGTCCCCACACCTACAAGGCCAGCAGATCCCTAATTCTCTCTCCAATCAAGTGCGCTCTCCCCAGCCTGTCCCTTCTCCACGGCCACAGtCCCAG\|C\|CCCCCCACTCCAGTCCTTCCCCAAGGATGCAGCCTCaGCCTTCTCCACACC |
| 2:2212:14749:5605#1#F | \| \| |
| MQ=60 isize=140 C=27 | GCCCAGTCCCCACACCTACAAGGCCAGCAGATCCCTAATTCTCTCTCCAATCAAGTGCGCTCTCCCCAGCCTGTCCCTTCTCCACGGCCACAGTCCCAg\|C\|CcCCCCACTCCAGTCCTTCCCCAAGGATGCAGCCTCAGCCAGGACTCCAAT |
| 2:2212:9702:11895#1#F | \| \| |
| MQ=60 isize=266 C=37 | GCCCAGTCCCCACACCTACAAGGCCAGCAGATCCCTAATTCTCTCTCCAATCAAGTGCGCTCTCCCCAGCCTGTCCCTTCTCCACGGCCACAGTCCCAG\|C\|CCCCCCACTCCAGTCCTTCCCCAAGGATGCAGCCTCAGCCTTCTCCACACC |
| 2:2212:12654:32202#1#F | \| \| |
| MQ=60 isize=177 C=37 | GCCCAGTCCCCACACCTACAAGGCCAGCAGATCCCTAATTCTCTCTCCAATCAAGTGCGCTCTCCCCAGCCTGTCCCTTCTCCACGGCCACAGTCCCAG\|C\|CCCCCCACTCCAGTCCTTCCCCAAGGATGCAGCCTCAGCCTTCTCCACACC |
| 2:2212:12771:40898#1#F | \| \| |
| MQ=60 isize=132 C=37 | GCCCAGTCCCCACACCTACAAGGCCAGCAGATCCCTAATTCTCTCTCCTATCAAGTGCGCTCTCCCCAGCCTGTCCCTTCTCCACGGCCACAGTCCCAG\|C\|CCCCCCACTCCAGTCCTTCCCCAAGGATGCAGAGGACTCCAATAAAATTCC |
| 2:2212:10442:66530#1#F | \| \| |
| MQ=60 isize=175 C=33 | GCCCAGTCCCCACACCTACAAGGCCAGCAGATCCCTAATTCTCTCTCCAATCAAGtGCGCTCTCCCCAGCCTGTCCCTTCTCCACGGCCGCAGTCCCAG\|C\|CCCCCCaCTCCAGtCCTTCCCCAAGGATGCAgCCTCagCCtTCTCCACACC |
| 2:2212:17349:82042#1#F | \| \| |
| MQ=60 isize=458 C=33 | GCCCAGTCCCCACACCTACAAGGCCAGCAGATCCCTAATTCTCTCTCCAATCAAGTGCGCTCTCCCCAGCCTGTCCCTTCTCCACGGCCACAGTCCCAG\|C\|CCCCCCACTCCAGTCCTTCCCCAAGGATGCAGCCTCAGCCTTCTCCACACC |
| 2:2213:16697:12473#1#F | \| \| |
| MQ=60 isize=233 C=33 | GCCCAGTCCCCACACCTACAAGGCCAGCAGATCCCTAATTCTCTCTCCAATCAAGTGCGCTCTCCCCAGCCTGTCCCTTCTCCACGGCCACAGTCCCAG\|C\|CCCCCCACTCCAGTCCTTCCCCAAGGATGCAGCCTCAGCCTTCTCCACACC |
| 2:2213:3847:22717#1#F | \| \| |
| MQ=60 isize=0 C=15 | GCCCAGTCCCCACACCTACAAGGCcAGCAGaTcccTaaTTCTctCTCCAATCAAGTGCGCTCTCCCCAGCCTGTCCCTTCTCCACGGCCAcAGTCCCAG\|c\|cCcCCCACTCCAGTCCTtCCCCAAGGATgCagCcTcAGCCttCtccACACC |
| 2:2213:1942:47183#1#F | \| \| |
| MQ=60 isize=120 C=37 | GCCCAGTCCCCACACCTACAAGGCCAGCAGATCCCTAATTCTCTCTCCAATCAAGTGCGCTCTCCCCAGCCTGTCCCTTCTCCACGGCCACAGTCCCAG\|C\|CCCCCCACTCCAGTCCTTCCAGGACTCCAATAGATGTACCTCCCTGTCTCT |
| 2:2213:9058:50039#1#F | \| \| |
| MQ=60 isize=209 C=15 | GCCCAGtCCCCACACCTACAAGGCCAGCAGATCcCTAATTCTCTCTCCAATCAAGTGCGCTCTCCCCAGCCtGTCcTTTCTCCACGGCCACAGTcCCag\|c\|CCCCCCacTCCAgtCctcCCCCAAgGatgcAGCCTcAgcCttCTCCaCaCc |
| 2:2213:7723:76135#1#F | \| \| |
| MQ=60 isize=212 C=37 | GCCCAGTCCCCACACCTACAAGGCCAGCAGATCCCTAATTCTCTCTCCAATCAAGTGCGCTCTCCCCAGCCTGTCCCTTCTCCACGGCCACAGTCCCAG\|C\|CCCCCCACTCCAGTCCTTCCCCAAGGATGCAGCCTCAGCCTTCTCCACACC |
| 2:2213:11685:89232#1#F | \| \| |
| MQ=60 isize=177 C=33 | GCCCAGTCCCCACACCTACAAGGCCAGCAGATCCCTAATTCTCTCTCCAATCAAGTGCGCTCTCCCCAGCCTGTCCCTTCTCCACGGCCACAGTCCCAG\|C\|CCCCCCACTCCAGTCCTTCCCCAAGGATGCAGCCTCAGCCTTCTCCACACC |
| 2:2214:15081:2598#1#F | \| \| |
| MQ=60 isize=0 C=33 | GCCCAGTCCCCACACCTACAAGGCCAGCAGATCCCTAATTCTCTCTCCAATCAAGTGCGCTCTCCCCAGCCTGTCCCTTCTCCACGGCCACAGTCCCAG\|C\|CCCCCCACTCCAGTCCTTCCCCAAGGATGCAGCCTCAGCCTTCTCCACACC |
| 2:2214:18455:36418#1#F | \| \| |
| MQ=60 isize=218 C=37 | GCCCAGTCCCCACACCTACAAGGCCAGCAGATCCCTAATTCTCTCTCCAATCAAGTGCGCTCTCCCCAGCCTGTCCCTTCTCCACGGCCACAGTCCCAG\|C\|CCCCCCACTCCAGTCCTTCCCCAAGGATGCAGCCTCAGCCTTCTCCACACC |
| 2:2214:8249:43957#1#F | \| \| |
| MQ=60 isize=127 C=37 | GCCCAGTCCCCACACCTACAAGGCCAGCAGATCCCTAATTCTCTCTCCAATCAAGTGCGCTCTCCCCAGCCTGTCCCTTCTCCACGGCCACAGTCCCAG\|C\|CCCCCCACTCCAGTCCTTCCCCAAGGAAGGACTCCAATAACGCCTGTCACC |
| 2:2214:15385:53909#1#F | \| \| |
| MQ=60 isize=153 C=33 | GCCCAGTCCCCACACCTACAAGGCCAGCAGATCCCTAATTCTCTCTCCAATCAAGTGCGCTCTCCCCAGCCTGtCCCTTCTCCACGGCCACAGTCCCAG\|C\|CCCCCCACTCCagTCCTTCCCCAaGGATGCAGCCTCagCCTTCTCCACAcc |
| 2:2214:11351:84393#1#F | \| \| |
| MQ=60 isize=158 C=37 | GCCCAGTCCCCACACCTACAAGGCCAGCAGATCCCTAATTCTCTCTCCAATCAAGTGCGCTCTCCCCAGCCTGTCCCTTCTCCACGGCCACAGTCCCAG\|C\|CCCCCCACTCCAGTCCTTCCCCAAGGATGCAGCCTCAGCCTTCTCCACACC |
| 2:2215:2798:21656#1#F | \| \| |
| MQ=60 isize=187 | GCCCAGTCCCCACACCTACAAGGCCAGCAGATCCCTAATTCTCTCTCCAATCAAGTGCGCTCTCCCCAgCCTGTCCCTTCTCCACGGCCACAGTCCCca\|c\|ccCCCcccccccacccctccccaaggcggccgcctcacccctctccacccc |
| 2:2215:3877:42399#1#F | \| \| |
| MQ=60 isize=132 C=22 | GCCCAGTCcCCAcACCTACAAGGcCAGCagaTCacTaatTctCTCtCCAATCAAGCGCGCTcTCCCCAGcCTGTCCCTTCTCCACGgCCACagtCCcag\|C\|CcCCCCACTCCAgtCCTTccCcaaggatgCAGAGGaCTCCAATTGGGCCGG |
| 2:2215:12875:89136#1#F | \| \| |
| MQ=60 isize=147 C=33 | GCCCAGTCCCCACACCTACAAGGCCAGCAGATCCCTAATTCTCTCTCCAATCAAGTGCGCTCTCCCCAGCCTGTCCCTTCTCAACGGCCACAGTCCCAG\|C\|CCCCCCaCTCCaGTCCTTCCCCAAGGATGCAGCCTCAGCCTTCTCCAGGAC |
| 2:2216:16474:10049#1#F | \| \| |
| MQ=60 isize=293 C=37 | GCCCAGTCCCCACACCTACAAGGCCAGCAGATCCCTAATTCTCTCTCCAATCAAGTGCGCTCTCCCCAGCCTGTCCCTTCTCCACGGCCACAGTCCCAG\|C\|CCCCCCACTCCAGTCCTTCCCCAAGGATGCAGCCTCAGCCTTCTCCACACC |
| 2:2216:3455:32016#1#F | \| \| |
| MQ=60 isize=166 C=33 | GCCCAGTCCCCACACCTACAAGGCCAGCAGATCCCTAATTCTCTCTCCAATCAAGTGCGCTCTCCCCAGCCTGTCCCTTCTCCACGGCCACAGTCCCAG\|C\|CCCCCCACTCCAGTCCTTCCCCAAGGATGCAGCCTCAGCCTTCTCCACACC |
| 2:2216:18274:48594#1#F | \| \| |
| MQ=60 isize=232 C=37 | GCCCAGTCCCCACACCTACAAGGCCAGCAGATCCCTAATTCTCTCTCCAATCAAGTGCGCTCTCCCCAGCCTGTCCCTTCTCCACGGCCACAGTCCCAG\|C\|CCCCCCACTCCAGTCCTTCCCCAAGGATGCAGCCTCAGCCTTCTCCACACC |
| 2:2216:14527:74393#1#F | \| \| |
| MQ=60 isize=186 C=33 | GCCCAGTCCCCACACCTACAAGGCCAGCAGATCCCTAATTCTCTCTCCAATCAAGTGCGCTCTCCCCAGCCTGTCCCTTCTCCACGGCCACAGTCCCAG\|C\|CCCCCCACTCCagTCCTTCCCCAAGGATGCAGCCTCAGCCTTCTCCACACC |
| 2:2216:4453:80982#1#F | \| \| |
| MQ=60 isize=132 C=33 | GCCCAGTCCCtACACCTACAAGGCCAGCAGATCCCTAATTCTCTCTCCAATCAAGTGCGCTCTCCCCAGCCTGTCCCTTCTCCACGGCCACAGTCCCAG\|C\|CCCCCCACTCCAGTCCTTCCCCAaGGATGCAGGACTCCAATTTGaACGCCC |
| 2:1107:13947:59242#2#R | \| \| |
| MQ=60 isize=-128 C=37 | GCCCAGtCCCCACaCCTACAAGGCCAGCAGAtCCCTAATTCTCTCTCCAATCAAGTGCGCTCTCCCCAGCCTGTCCCTTCTCCACGGCCACAGTCCCAG\|C\|CCCCCCACTCCAGTCCTTCCCCAAGGATanGACTCCAATGAACTTATACTC |
| 2:1201:7301:16656#2#R | \|C\| |
| MQ=60 isize=-127 C=22 | gcccagtccccacacctacaaggccagcagatccctaatgctccccccaatcaagtgcgctcccCCCAGCcTGtCCCtTCTCCACGGCCacagTcCCAI\|C\|CCCCCCaCTCCAGTCCTtCCCCAAGGAAGGACTCCAATtCTCGCTCCCCC |
| 1:1109:1347:23606#1#F | \| \| |
| MQ=60 isize=248 C=37 | CCCAGTCCCCACACCTACAAGGCCAGCAGATCCCTAATTCTCTCTCCAATCAAGTGCGCTCTCCCCAGCCTGTCCCTTCTCCACGGCCACAGTCCCAG\|C\|CCCCCCACTCCAGTCCTTCCCCAAGGATGCAGCCTCAGCCTTCTCCACACCA |
| 1:1216:1627:42737#1#F | \| \| |
| MQ=60 isize=165 C=33 | CCCAGTCCCCACACCTACAAGGCCAGCAGATCCCTAATTCTCTCTCCAATCAAGTGCGCTCtcCCCAGcCTGTCCCTTCTCCACgGCCACAGtcCCAG\|C\|CCCCCCactcccgtccttccccaaggatgcagcctcagccttctccacaccc |
| 1:2112:11709:100832#1#F | \| \| |
| MQ=60 isize=131 C=33 | CCCAGTCCCCACACCTACAAGGCCAGCAGATCCCTAATTCTCTCTCCAATCAAGTGCGCTCTCCCCAGCCTGTCCCTTCTCCACGGCCACAGTCCCAG\|C\|CCCCCCACTCCAGTCCTTCCCCAAGGATGCAGAGGACTCCAATCAATCGTTA |
| 1:2211:13470:93251#1#F | \| \| |
| MQ=60 isize=328 C=33 | CCCAGTCCCCACACCTACAAGGCCAGCAGATCCCTAATTCTCTCTCCAATCAAGTGCGCTCTCCCCAGCCTGTCCCTTCTCCACGGCCACAGTCCCAG\|C\|CCCCCCACTCCAGTCCTTCCCCAAGGATGCAGCCTCAGCCTTCTCCACACCc |
| 2:2108:20993:88501#1#F | \| \| |
| MQ=60 isize=318 C=33 | CCCAGTCCCCACACCTACAAGGCCAGCAGATCCCTAATTCTCTCTCCAATCAAGTGCGCTCTCCCCAGCCTGTCCCTTCTCCACGGCCACAGTCCCAG\|C\|CCCCCTACTCCAGTCCTTCCCCAAGGATGCAGCCTCAGCCTTCTCCACACCa |
| 2:2208:6674:43239#1#F | \| \| |
| MQ=60 isize=124 C=37 | CCCAGTCCCCACACCTACAAGGCCAGCAGATCCCTAATTCTCTCTCCAATCAAGTGCGCTCTCCCCAGCCTGTCCCTTCTCCACGGCCACAGTCCCAG\|C\|CCCCCCaCTCCAGTCCTTCCCCAAGAGGATTCCAATAGTAACAAGATCCTGT |
| 1:2203:9856:66549#2#R | \| \| |
| MQ=60 isize=-132 C=22 | CCCAgtCCCccCCCCtACAAGGCCAGCAGATCCCTAAttctCtctCCAAtCaaGtGccCcCTCCCCaGCCTgtCCCttctCCaCGGCCaCAGTCCCAG\|C\|CCCCCCACTCCAGTCCTTCCCCAAGGATGCAGACTCCAATTGAAGGCCGCTA |
| 2:2108:15608:88698#2#R | \| \| |
| MQ=60 isize=-129 C=33 | cccagtccccacccctacaaggccagcagatccctaattctCTCTCCaATCAAGTGCGctctcCCcaGCCTGTCCCTTCTCCACGGCCACAGTCCcAG\|C\|CCCCCcaCTCCAGTCCTTCCCCAAGGATGAGGACTCCAATATCGAGCTCCAA |
| 2:1214:14055:8294#2#F | \| \| |
| MQ=60 isize=255 C=33 | GGCCGTCGGGGCATTGGAGTCCTCCAGTCCCCACACCTACAAGGCCAGCAGATCCCTAATTCTCTCTCCAATCAAGTGCGCTCTCCCCAGCCTGTCCCTTCTCCACGGCCACAGTCCCAG\|C\|CCCCCCACTCCAGTCCTTCCCCAAGGATGC |
| 1:1201:15655:65013#2#R | \| \| |
| MQ=60 isize=-131 C=22 | ccagtccccacaCCtaCAaGgCCAGCAGATCCcTAATTCtctctccAatcAagtgcgctCtCCCCAGCCTgTcCcTtCTCCaCGGCCacaGTCCCaG\|C\|CCCCCCACTCCAGtCCTtCCCCAAGGATGCanGACTCCAATgTGCGACATCCA |
| 1:2107:8164:64341#2#R | \| \| |
| MQ=60 isize=-132 C=37 | CCAGTCCCCACACCTACAAGGCCAGCAGAtcCCTAATTCTCTCTCCAATCAAGTGCGCTCTCCCCAGCCTGTCCCTTCTCCACGGCCACAGTCCCAG\|C\|CCCCCCACTCCAGTCCTTCCCCAAGGATGCAGGACTCCAATCCTTCACCGTTG |
| 2:1110:2305:82606#2#R | \| \| |
| MQ=60 isize=-132 C=33 | ccagcccccacacctacaaggccagcagatccctaattctctctccaatcaagtgccctctcCCCAAcctgTCCctTctCcaCGGCCaCAGTCCCAG\|C\|CCCCCCACTCCAGTCCTTCCCCAAGGATGCAGGACTCCAATTTGAACGCCCAC |
| 2:1204:1726:70743#2#R | \| \| |
| MQ=60 isize=-131 C=33 | ccagcccccacacctccaaggccaggagatccCTAAttCTCtCTCCaatcaagtGcgctctCCCCAGCCTgccCCTtctcaacgGcccCAGtCCCaG\|C\|CCCCcCaCTCCAGTCCTTCCCCAAGGATGCanGACTCCAATAGCACACAGCTG |
| 2:2106:11347:67319#2#R | \| \| |
| MQ=60 isize=-131 C=37 | CCAGtCCCCacaCCtacAAGGCCAGCAGATCCCTAATTCtCTCTCCAATcAAGTGCGCTCTCCCCAGCCTGTCCCTTCTCCACGGCCACAGTCCCAG\|C\|CCCCCCACTCCAGTCCTTCCCCAAGGATGCanGACTCCAATCAACTCCGCCAG |
| 2:2216:4453:80982#2#R | \| \| |
| MQ=60 isize=-132 C=15 | CCaGTCCCTaCacctACAAGGCCAGCAGatCCCTAATtctctCTCCAATcAAGTgCGCTCTCCCCAGCctGTCCCTTCtCCaCGGCcaCAGTCCCAG\|c\|CCCCCCaCTCCAGTCCTTCCCCAAGGATGCAGGACTCCAATTTGAACGCCCaC |
| 2:1113:3674:95463#2#R | \| \| |
| MQ=60 isize=-489 C=22 | cagtccccacacctacaaggacagcagatccctaattctccctccaatcaagtgcgctctccccagcctgtcccttctccacggcCACAGTCCCaG\|C\|CCCCCCaCTCCAGTcCTTCCCCAAGGATGCAAgnacaCcaagcCCCCAATGCAA |
| 2:2110:16568:49429#2#R | \| \| |
| MQ=60 isize=-121 C=37 | ggttttGGCaCAGtCCcaACACCTACAAGGCCAGCAGATCCCTAATTCtcTCTCCAATCAAGTGCGCTCTCCCCAGCCTGTCCCTTCTCCACGGCCACAGTCCCAG\|C\|CCCCCCACTCCAGTCCTTCCCAGGACTCCAATCCAAATATCACC |
| 1:1111:8260:62322#2#F | \| \| |
| MQ=60 isize=253 C=33 | GCGGGGGTAAGTatTTGaGTCCTCACACCTACAAGGCCAGCAGATCCCtAATTCTCTCTCCAATCAAgtgCGCtCTCCCCAgCCTgTCCCtTCtCCcCGGCCaCAGtCCCAG\|C\|CCCCCCaCTCCAgtCCtTCCCCCAggATGCggcctcag |
| 2:1214:12022:38432#2#F | \| \| |
| MQ=60 isize=253 C=33 | GAATATTAGCTTATTGGAGTCCTAGTCCCCACACCTACAAGGCCAGCAGATCCCTAATTCTCTCTCCAATCAAGTGCGCTCTCCCCAGCCTGTCCCTTCTCCACGGCCACAGTCCCAg\|C\|CCCCCCaCCCCagtCCtTCCCCaaggatgcag |
| 1:1114:11530:74529#2#R | \| \| |
| MQ=60 isize=-132 C=27 | agtCCCCACACCTACAAGGCCAGCAGATCCCTAATTCTCTCTCCAATCAAGTGCGCTCTCCCCAGCCTGTCCCTTCTCCACGGCCACAGTCCCAG\|C\|CCCCCCACTCCAGTCCTTCCCCAAGGATGCAGAGGACTCCAATAATTCACACCCT |
| 1:1216:16335:62419#2#R | \| \| |
| MQ=60 isize=-132 C=15 | cgtccccacccctacaaggcccgcagatCCCTaATTctctCTCCaAtcaagtgCGCtCtCCCCAGCCTGtCCcTtCtCCACGGCCACAGTCCCaG\|c\|CCCCCCACTCCAGTCCTTCCCCAAGGATGCAGAgnACTCCAAtCAATCGTTAAAT |
| 1:2112:11709:100832#2#R | \| \| |
| MQ=60 isize=-131 C=37 | AGTCCCCaCaCCTACAAGGCCAGCAGATCCCTAATTCTCTCTCCAATCAAGTGCGCTCTCCCCAGCCTGTCCCTTCTCCACGGCCACAGTCCCAG\|C\|CCCCCCACTCCAGTCCTTCCCCAAGGATGCAGAGGACTCCAATCAATCGTTAAAT |
| 2:1113:18577:75939#2#R | \| \| |
| MQ=60 isize=-132 C=37 | cgtCCCcACaCCTACAAGGCCAGCAGatCCCTAATTCtCTCtCcAATCAaGtGCGCTCTCCCCAGCCTGTCCCTTCTCCACGGCCACAGTCCCAG\|C\|CCCCCCACTCCAGTCCTTCCCCAAGGATGCAGAgnACTCCAATGCAGCAGGTGTA |
| 2:1206:18996:35336#2#R | \| \| |
| MQ=60 isize=-132 C=37 | agTCCCCACaCCTACAAGGCCAGCAGATCCCTAATTCTCTCTCCAATCAAGTGCGCTCTCCCCAGCCTGTCCCTTCTCCACGGCCACAGTCCCAG\|C\|CCCCCCACTCCAGTCCTTCCCCAAGGATGCAGanGACTCCAATAATTCACACCCT |
| 2:2112:4699:13431#2#R | \| \| |
| MQ=60 isize=-132 C=33 | cgTCCCCACACctacaAggcCaGCAGatCCCTaattctCtCcCCaAtcaagtgCgCtctCCCCAGCCTGtCCCTTCtCCACGGCCACAGTCCCAG\|C\|CCCCCCaCTCCAGTCCTTCCCCAAGGATGCAGAgnACTCCAATTATTATCTCCTA |
| 2:2215:3877:42399#2#R | \| \| |
| MQ=60 isize=-132 C=22 | agcccccccaCCTaCaaGGCCagcagatCCCTAaTTCTCTCTccAatCAaGCGCGctctCCCCAGcCTgTCCcTtCtccaCggccgCAGTCCCAg\|C\|cCcccCaCTCCaGTCCTTCCCCAAGGATGCAGAGGACTCCAATTGGGCCGGAGTG |
| 1:1201:14897:38639#2#R | \| \| |
| MQ=60 isize=-133 C=33 | gtccccacccctacaaggccaGCaGAtCCCTAATTCTCTCTCCaatCAagtgCgCTctCCcCAgcCtGtcccTTCTCCaCGgCcaCAGTCCCAG\|C\|CCCCcCACTCCAGTCCTTCCCCAAGGATGCAGCanGACTCCAATCTAAATCTAAGT |
| 1:2101:15328:42253#2#R | \| \| |
| MQ=60 isize=-133 C=33 | GTCCCCACACCTACAAGGCCAGCAGATCCCTAAtTCTCTCTCCAAtCAAGTGCGCTCTCCCCAGCCTGTCCCTTCTCCACGGCcACAGTCCCAG\|C\|CCCCCCACTCCAGTCCTTCCCCAAGGATGCAGCAGGACTCCAATGTCTCAAATTCA |
| 1:2204:2435:86546#2#R | \| \| |
| MQ=60 isize=-133 C=37 | gtccccacacctacaaGGCCAgCAGATCcCTAATtCTCTCTCCAAtcaAGtGCgctCTCCCCAGCCTGtCCCTtCTCCACGGCCACAGTCcCAG\|C\|CCCCCCACTCCAGTCCTTCCCCAAGGATGCAGCAGGACTCCAATGTCGGCCCGACC |
| 2:2106:18012:85175#2#R | \| \| |
| MQ=60 isize=-133 C=33 | gtCCCCaCcCCTACAAGGCCAGCAGatCCCTAATtCtCtCtCCAATCAagTGCGCTCTCcCCAGCCTGTCCCTTCTCCACGGCCACAGTCCCAG\|C\|CCCCCCACTCCAGTCCTTCCCCAAGGATGCAGCanGACTCCAATAATCACAACCaC |
| 2:2205:15696:95697#2#R | \| \| |
| MQ=60 isize=-133 C=37 | GTCCCCACACCTACAAGGCCAGCAGAtCCCTAAtTCTCTCTCCAATCAAGTGCGCtCtCCCCAGCCTGTCcCTTCTCCACGGCCACAGTCCCAG\|C\|CCCCCCACTCCAGTCCTTCCCCAAGGATGCAGcanGACTCCAAtCCCCCCCTCTAC |
| 1:2113:17528:93294#2#R | \| \| |
| MQ=60 isize=-134 C=2 | cccccacacctacaaggccagcagatccctaattctctctccaatcaagtgcgctctccccagcctgtcccttctccacggccacagtcccag\|c\|ccCCCCaCTCCAGTCCTTCCCCAAGGATGCAGCcannACTCCAATGGACAGCCCCGT |
| 2:2115:4873:66275#2#R | \| \| |
| MQ=60 isize=-134 C=22 | gccccacccctaaaaggccagcagttccctaattctctctccaatcaagttccccctccCCagCCTGTCCCTTCTCCACGGCCACAgtcCCAg\|C\|CCCCCcaCTCCAGTCCTTCCCCAAGGATGCAGcCAgnACTCCAATCGGATAGCTTTG |
| 1:1106:18835:3196#2#R | \| \| |
| MQ=60 isize=-135 C=27 | CCCCACaCCTACAAGgCCAGCAgaTCCCTAATTCtCtCtCCAATCAAGTgCgctCTCCCCAGCCTGTCCCTTCTCCACGGCCACAGTCCCAG\|C\|CCCCCCACTCCAGTCCTTCCCCAAGGATGCAGCCTanGACTCCAATCGATACTAAAAG |
| 2:1107:4620:19186#2#R | \| \| |
| MQ=60 isize=-125 C=37 | ttggcccaccccccacccctacaaggccagcagaTCCCTAATTCcCtccCCAAtCAAGTGCgCtccCCCCAGCCTGTCCCtTCtCCaCGGCCACAGTCCCAG\|C\|CCCCCCACTCCAGTCCTTCCCCAAGanGACTCCAATAGTAACAAGATC |
| 1:1113:16641:34316#2#F | \| \| |
| MQ=60 isize=249 C=2 | TAAaGACGGccTATTGGAGTCCTCcCACACCTACAAGgCCAGCAGATCCcTAATTctCTctccaatcaagtgcgctctccccagcctgtcccttctccacggccacagtcccag\|c\|ccccccactccagtccttccccaaggatgccgcctc |
| 2:1105:6602:56249#2#F | \| \| |
| MQ=60 isize=249 C=15 | GGCGGTTTGCCTATTGGAGTCntCCCACACCTACAAGGCCAGCAGATCCCTAATTCTCTCTCCAAtCAAGtGCGCtCtCCCCAGCCTgtCCCTTCtCCccGgcCccAgcccCAg\|c\|CCCCCcaccccagccctcccccaaggactcaccccc |
| 2:2111:2730:87007#2#F | \| \| |
| MQ=60 isize=249 C=2 | CTGAAATACCCAATTGGAGTCCTCCCACACCTACAAGGCCAGCAGATCCCTAATTCTCTCTCCAATcAAGTgcgCTCtCCCcacCCTGcCCCTTCtCCccggccccattcccac\|c\|ccccccccccccgcccttccccaaggacgccccctc |
| 1:2209:6454:81943#2#R | \| \| |
| MQ=60 isize=-137 C=37 | CCCACaCCTACAAGGCCAGCAGATCCCTAATTCTCTCTCCAATCAAGTGCGCTCTCCCCAGCCTGTCCCTTCTCCACGGCCACAGTCCCAG\|C\|CCCCCCACTCCAGTCCTTCCCCAAGGATGCAGCCTCanGACTCCAATAGCCCCGGGGTC |
| 2:1109:20674:39061#2#R | \| \| |
| MQ=60 isize=-137 C=27 | CcCACaCCTAcAAgGCCAGcAgatccCTAaTtctCTCTcCAATCAAGTGcGCTCTCCCCAGCCTGTCCCTTCTCCACGGCCACAGTCCCAG\|C\|CCCCCCACTCCAGTCCTTCCCCAAGGATGCAGCCTCanGACTCCAATAGGGAACCTAAT |
| 2:1211:15881:92983#2#R | \| \| |
| MQ=60 isize=-138 C=37 | CCCaCaCCTaCAAGgCCAGCAGaTCcCTaAttctCTCTCCAatCAAGTGcgCTCtcCCCAGCCTGTCCCTTCTCCACGGCCACAGTCCCAG\|C\|CCCCCCACTCCAGTCCTTCCCCAAGGATGCAGCCTCAGGACTCCAATCAATGTGACCCC |
| 1:2213:15175:21355#2#R | \| \| |
| MQ=60 isize=-137 C=37 | ccaccCCTACAAGGCCAGCAGaTCCCTAATtCTCTCTCCAATCAAGTGCgctCTCCCCAGCCTGTCCCTTCTCCACGGCCACAGTCCCAG\|C\|CCCCCCACTCCAGTCCTTCCCCAAGGATGCAGCCTCAAgnACTCCAATATTGACCGCGCA |
| 2:1212:5500:8570#2#R | \| \| |
| MQ=60 isize=-137 C=37 | CCACACCTacaaggtCAGcagaTCCCTAATTCtCtCTCCAAtCAAGTGcgCTCTCCCCAGCCTGTCCCTtCTCCACGGCCACAGTCCCAG\|C\|CCCCCCACTCCAGTCCTTCCCCAAGGATGCAGCCTCAAGGACTCCAATTGAACCAACTTA |
| 1:2206:13077:67278#2#F | \| \| |
| MQ=60 isize=247 C=33 | GCGGACCTACAGATTGGAGTCntCACACCTACAAGGCCAGCAGATGCCTAATTCTCTCTCCAATCAAGTGCGCTCTCCCCAGCCTGTCCCTTCTCCACGGCCaCAGtCCCAG\|C\|CCCCCCACTCCAGTCCTTCCCCAAggATGCAGCCTCac |
| 1:1108:6199:47556#2#R | \| \| |
| MQ=60 isize=-138 C=33 | caccCCTaCAAGGCCAGCAGATCCCTAAtTCTCTCtCCaAtCaAGTGCgctCTCCCCAGCCTGTCCCTTCTCCACGGCCACAGTCCCAG\|C\|CCCCCCACTCCAGTCCTTCCCCAAGGATGCAGCCTCAGanGACTCCAATAGATGAATTCCG |
| 1:2101:12799:60988#2#R | \| \| |
| MQ=60 isize=-138 C=33 | CACACCTACAAGGCCAGCAGATCCCTAATTCTCTCTCCAATCAAgTGCGCTCTCCCCAGCCTGTCCCTTCTCCACGGCCACAGTCCCAG\|C\|CCCCCCACTCCAGTCCTTCCCCAAGGATGCAGCCTCAGAGGACTCCAATATTACACCGACA |
| 1:2115:19246:25394#2#R | \| \| |
| MQ=60 isize=-138 C=37 | CaCACCTACAAGGCCAGCAGATCCCTAATTCTCtCTCCAATCAAGTGCGCtCTCCCCAGCCTGTCCCTTCTCCACGGCCACAGTCCCAG\|C\|CCCCCCACTCCAGTCCTTCCCCAAGGATGCAGCCTCAGAGGACTCCAATAAAAAAATAGTG |
| 2:1201:6700:65111#2#R | \| \| |
| MQ=60 isize=-138 C=15 | cacacccacaaggccagcaGATCCCtAAttCtctCtccaatCAAgtgCgctcTCCCCAGCCTGTCCCTTCTCCaCGGCcACAGTCCCAG\|c\|CCCCCCACTCCAGTCCTTCCCCAAgGATGCAGCCTCAGAGGACTCCAATGCGGTCAGTCGA |
| 2:2210:12480:6870#2#R | \| \| |
| MQ=60 isize=-138 C=27 | CACACCTACAAGGCCAGCAGAtCCCTGATtCTCTCTCCAATCAAGTGCgctCTCCCCAGCCTGTCCCTTCTCCACGGCCACAGTCCCAG\|C\|CCCCCcACtCCAGTCCTTCCCCAAGGATGCAGCCTCAGaGGACTCCAATAAATGAAAAAGG |
| 1:1116:20637:62128#2#R | \| \| |
| MQ=60 isize=-139 C=37 | acaccaacaaggccagcagatccctaattctctccCCAatCAaGTGccctctCCCCAGCCTGTCCCTtCTCCaCGGCCACagtCCCAG\|C\|CCCCCCACtCCAGTCCTTCCCCAAGGATGCAGCCTCAGCAGGACTCCAaTCGCATAATTCGA |
| 1:2112:13456:84657#2#R | \| \| |
| MQ=60 isize=-139 C=33 | aCaCCTaCAAGGCCAGCAGATCCCTAATTCTCTCTCCAATCAAGTgCgctCtCCCCAGCCTGTCCCTTCTCCACGGCCACAGTCCCAG\|C\|CCCCCCACTCCAGTCCTTCCCCAAGGATGCAGCCTCAGCAGGACTCCAATGCATACGCGTAA |
| 1:2215:4061:76083#2#R | \| \| |
| MQ=60 isize=-139 C=37 | cCACCTACAAGGCCAGCAGATCCCTAAttCTCTCTCCAATCAAGTGCGCTCTCCCCAGCCTGTCCCTTCTCCACGGCCACAGTCCCAG\|C\|CCCCCCACTCCAGTCCTTCCCCAAGGATGCAGCCTCAGCAgnACTCCAATGCCCCTCCCGCC |
| 1:1210:1946:63065#2#F | \| \| |
| MQ=60 isize=245 C=33 | ATCTGGATGGTCATTGGAGTCCTCACCTACAAGGCCAGCAGATCCCTAATTCTCTCTCCAATCAAGTGCGCTCTCCCCAGCCTGTCCCTTCTCCACGGCCACAGTCCCAG\|C\|CCCCCCACTCCAGTCCTTCCCCAAGGATGCAGCCTCaGCC |
| 2:1206:17800:93422#2#F | \| \| |
| MQ=60 isize=245 C=37 | CAGACCCGGAGAATTGGAGTCntCACCTACAAGGCCAGCAGATCCCTAATTCTCTCTCCAATCAAGTGCGCTCTCCCCAGCCTGTCCCTTCTCCACGGCCACAGTCCCAG\|C\|CCCCCCACTCCAGTCCTTCCCCAaGGaTGCAGCCTCAGCC |
| 2:2214:18420:61063#2#F | \| \| |
| MQ=60 isize=245 C=37 | ATCTGGATGGTCATTGGAGTCCTCACCTACAAGGCCAGCAGGTCCCTAATTCTCTCTCCAATCAAGTGCGCTCTCCCCAGCCTGTCCCTTCTCCACGGCCACAGTCCCAG\|C\|CCCCCCACTCCAGTCCTTCCCCAAGGATGCAGCcTCAGCC |
| 1:1206:1374:9587#2#R | \| \| |
| MQ=60 isize=-140 C=37 | CaCCTACAaGgCCAGCagATCCCTaattCtCTCtCCAAtCAaGTGCGCtCTCCCCAGCCTGTCCCTTCTCCACGGCCACAGTCCCAG\|C\|CCCCCCACTCCAGTCCTTCCCCAAGGATGCAGCCTCAGCCAGGACTCCAATGCACGCGTCAAC |
| 1:1216:20790:55864#2#R | \| \| |
| MQ=60 isize=-140 C=33 | CACCtaCaAGGCCAGCAGATCCCTAATTCtCtCtCCAATCAAGTGCGCtCtCCCCAGCCTGTCCCTTCTCCACGGCCACAGTCCCAG\|C\|CCCCCCACTCCAGTCCTTCCCCAAGGATGCAGCCTCAGCCAgnACTCCAATAGAGAGCGGAGT |
| 1:2102:12711:39437#2#R | \| \| |
| MQ=60 isize=-140 C=33 | cacctaccaggccagcagatccctaattctctctccaatcaagtgcgctctCCCCaGCCTgTCCCttCtCCacGGCCACAGTCCCAG\|C\|CCCCCCACtCCAGTCCTTCCCCAAGGATGCAGCCTCAGCCAGGACTCCAatCCTGCGGTTCAC |
| 1:2112:1642:94661#2#R | \| \| |
| MQ=60 isize=-140 C=33 | CACCTaCAAGGCCAGCaGATCCCTAATTCTCCCTCCAATCAAGTGCGCTCTCCCCAGCCTGTCCCTTCTCCACGGCCACAGTCCCAG\|C\|CCCCCCACtCCAGTCCTTCCCCAAGGATGCAGCCTCAGCCAGGACTCCAATGGGGACGAGGGG |
| 1:2210:18638:11792#2#R | \| \| |
| MQ=60 isize=-140 C=37 | CcCCTACAAGGCCAGCaGatCCCTAATtctctCtCCAATCAAGtgCGCtCTCCCCAGCCTGTCCCTTCTCCACGGCCACAGTCCCAG\|C\|CCCCCCACTCCAGTCCTTCCCCAAGGATGCAGCCTCAGCCAGGACTCCAATGAAACGTTTTGT |
| 2:1108:18428:51456#2#R | \| \| |
| MQ=60 isize=-140 C=37 | cccctACAAGGCcAGCaGATCCCTAATTCtCtCtCCAatcAAGTGcgCTCTCCCCAGCCTGTCCCTTCTCCACGGCCACAGTCCCAG\|C\|CCCCCCACTCCAGTCCTTCCCCAAGGATGCAGCCCCAGCCanGACTCCAATTCCTCAGCACCC |
| 2:1215:4261:78870#2#R | \| \| |
| MQ=60 isize=-140 C=37 | CACCTACAAGGCCAGCAGATCCCTAATTCtCTCTCCAATCAAGTGCgCtCTCCCCAGCCTGTCCCTTCTCCACGGCCACAGTCCCAG\|C\|CCCCCCACTCCAGTCCTTCCCCAAGGATGCAGCCTCAGCCAgnACTCCAATATGTATCAAAAA |
| 2:2108:7102:97973#2#R | \| \| |
| MQ=60 isize=-140 C=37 | cacctacaaggcccgaaaatccctaattccctCTCCAATCAAGtgcgctctCCCCaGCCTGTCCCTTCtCCACGGCCACAGTCCCAG\|C\|CCCCCCACTCCAGTCCTTCCCCAAGGATGCAGCCTCAGCCAGGACTCCAATAGGTTTGCAGAT |
| 2:2201:3203:62616#2#R | \| \| |
| MQ=60 isize=-140 C=33 | CcCCtacaAGGCCAGCAGATCCCTAATTCtCtCtCCAATCAAGtgCgCtCTCCCCAGCCTGTCCCtTCtCCACGGCCACAGTCCCAG\|C\|CCCCCCTCTCCAGTCCTTCCCCAAGGATGCAGCCTCAGCCanGACTCCAATATTTATTGCCCG |
| 1:1103:17951:22979#2#R | \| \| |
| MQ=60 isize=-125 C=27 | ttGGCCCAGcCCCCAtcCCtACAagGCCAGCAGcTCCCTaAttCtCCCtCCAATCAAGTGCgCTCTCCCCAGCCTGTCCCTTCTCCaCGGCCACAGTCCCAG\|C\|CCCCCCACTCCAGTCCTTCCCCAAGAGGACTCCAAtTCCCCCCCGAAC |
| 1:2206:4164:8382#2#R | \| \| |
| MQ=60 isize=-139 C=33 | cccCCTACAAGGCCAGCAGATCCCTAATTCtCTCTCCAATCAAGTGCGCTCTCCCCAGCCTGTCCCTTCTCCACGGCCaCAGTCCCAG\|C\|CCCCCCACTCCAGTCCTTCCCCAAGGATGCgGCCTCAGCanGACTCCAATCGTGGTTGGCCG |
| 2:1112:5163:67573#2#R | \| \| |
| MQ=60 isize=-121 C=37 | cgtTTTGgcCcaGtCCcacCCCCTaCAAGgCCaGCaGATCCCTAATtCtctccCCAATCAAGTGCGctCTCCCCAGCCTGTCCCTTCTCCACGGCCACAGTCCCAG\|C\|CCCCCCaCTCCAGTCCTTCCCAGGACTCCAATCTCTGCCAGCCT |
| 2:2110:11389:10083#2#R | \| \| |
| MQ=60 isize=-139 C=37 | ccgcctacaAGGCCagcagaTCCCTAATTCtctcccCAAtCAAGTgCgCTCTCccCAGCCTGTCCCTTCTCCACGGCCACAGTCCCAG\|C\|CCCCCCACTCCAGTCCTTCCCCAAGGATGCAGCCTCAGCAGGACTCCAATCGCGGCCCCAAA |
| 1:1102:18948:8131#2#R | \| \| |
| MQ=60 isize=-143 C=33 | CTACAAGGCCAGCAGATCCCTAattCtCtCtCCAATCAAGTGCGCTCTCCCCAGCCTGTCCCTTCTCCACGGCCACAGTCCCAG\|C\|CCCCCCACTCCAGTCCTTCCCCAAGGATGCAGCCTCAGCCTTCanGACTCCAATCTACTCAAACAT |
| 1:2208:10791:52716#2#R | \| \| |
| MQ=60 isize=-143 C=2 | cttcaaggccagcagatccataattcggtctccaatccagtgcgccctccccagcctgtcccatctctacggccacagtcccag\|c\|ccccccactgcagtccttccccaaggatgcagcctaagccttcanGACTCCAATTgtgcTAGCcta |
| 2:1113:3948:69868#2#R | \| \| |
| MQ=60 isize=-143 C=37 | ctaccAGGCCAGCagatCCCTAATtctccCcCCAAtCaaGtGcgctctCCCCAGCCTGTCCCTTCTCCACGGCCACAGTCCCAG\|C\|CCCCCCACTCCAGTCCTTCCCCAAGGATGCAGCCTCAGCCTTCAgnACTCCAATGCCCATGACACC |
| 2:2106:9771:87218#2#R | \| \| |
| MQ=60 isize=-143 C=37 | ctaccaggccagcaggtccctaattctctctccaatcaagtgcgctctccCCAGCCTGTCCCTTCTCCaCGGCCACAGTCCCAG\|C\|CCCCCCACTCCAGTCCTTCCCCAAGGATGCAGCCTCAGCCTTCanGACTCCAATGGTTATGACGTA |
| 2:2212:20945:37447#2#R | \| \| |
| MQ=60 isize=-143 C=33 | CTACAAGgCCAGCAGATCCCTAAttCTCTCTCCAATCAAGTgCgCTCTCCCCAGCCTgtCCCTTCTCCACGGCCACAGTCCCAG\|C\|CCCCCCACTCCAGTCCTTCCCCAAGGATGCAGCCTCAGCCTTCAGGACTCCAATGGTGATATATTA |
| 1:1116:18930:55009#2#F | \| \| |
| MQ=60 isize=241 C=22 | TGCCGTGGGTGGATTGGAGTCCTTACAAGGCCAGCAGATCCCTAATTCTCTCTCCAATCAAGTGCGCTCTCCCCAGCCTGTCCCTTCTCCACGGCCACAGTCCCAg\|C\|CCCCCCACTCCAGtCCTTCCCCAAGGATGCAGCCTCAgcCCTCt |
| 1:1114:3568:19305#2#R | \| \| |
| MQ=60 isize=-144 C=33 | taCAAGGCCAGCAGATCCCTAATTCTCTCTCCAATCAAGtgCgctCTCCCCAGCCTGTCCCTTCTCCACGGCCACAATCCCAG\|C\|CCCCCCACTCCAGTCCTTCCTCAAGGATGCAGCCTCAGCCTTCTAGGACTCCAATTGCCTCATAGGA |
| 1:1206:5156:63554#2#R | \| \| |
| MQ=60 isize=-144 C=2 | cacaaggccagcagatccctaatccgctctccaatcaagtgcgctctccccagcctgtcccttctccacggccacagtcccag\|c\|ccccccactccaGTCCTTCCCCAAGGATGCAGCCtCAGCCTTCTAGGACTCCAATAAGATTGGTACa |
| 1:2101:4845:71047#2#R | \| \| |
| MQ=60 isize=-144 C=37 | taCAAGGCCAGCAGATCCCTAaTTCTCTCTCCAATCAAGTGCGCTCTCCCCAGCCTGTCCCTTCTCCACGGCCACAGTCCCAG\|C\|CCCCCCACTCCAGTCCTTCCCCAAGGATGCAGCCTCAGCCTTCTAGGACTCCAATCATAGCATTGTG |
| 1:2213:14765:2762#2#R | \| \| |
| MQ=60 isize=-144 C=33 | caCAAGGCCAGCAGATCCCTAAttctccCtCCAAtCAAGTgcgctctcCCCAGCCTGtCCCTTCTCcaCGGCCACAGTCCCAG\|C\|CCCCCCACTCCAGTCCTTCCCCAAGGATGCAGCCTCAGCCTTCTAgnACTCCAATCCGCCCCGCGGG |
| 2:1210:18317:46026#2#R | \| \| |
| MQ=60 isize=-144 C=37 | tACAAGGCCAGCAGATCCCTAATTCTCTCTCCAATCAAGTGCGCTCTCCCCAGCCTGTCCCTTCTCCACGGCCACAGTCCCAG\|C\|CCCCCCACTCCAGTCCTTCCCCAAGGATGCAGCCTCAGCCTTCTAGGACTCCAATCTACCAGCTGCG |
| 2:2203:20972:31641#2#R | \| \| |
| MQ=60 isize=-144 C=33 | taCaaggCCaGCAGATCCCTAAtTCtCtCTCCAATCAAgtGCgCTCTCCCCAGCCTGTCCCTtCTCCACGGCCACAGtCCCAG\|C\|CCCCCCACTCCAGTCCTTCCCCAAGGATGCAGCCTCAGCCTTCTanGACTCCAATAAGATTGGTACA |
| 2:2210:6949:51710#2#R | \| \| |
| MQ=60 isize=-144 C=33 | tacaaggccagcAGaTCCCTAATtccCTCtCCaATCaaGTgCGCtctCCCCagCctGTCCCttcTCccCGGCCAcaGTCCcag\|C\|CCCCcCaCtCCAGTCCTTCCCCAAGGatGCAGCCTCAGCCTTCTAGGACTCCAATtGTTCATtCTAA |
| 1:1114:10715:71942#2#R | \| \| |
| MQ=60 isize=-145 C=37 | aCAAGGCCAGCAGATCCCTAATTCTCTCTCCAATCAAGTGCGCTCTCCCCAGCCTGTCCCTTCTCCACGGCCACAGTCCCAG\|C\|CCCCCCACTCCAGTCCTTCCCCAAGGATGCAGCCTCAGCCTTCTCAGGACTCCAATGCGCCTGACCTG |
| 2:1116:4571:17541#2#R | \| \| |
| MQ=60 isize=-145 C=33 | ccaaggccagcagatCCCtAATTCtCtctCCAaTCAAgtgCgctCTCCCCaGCCTgtCCCTTCTCCaCGGCCACaGTCCCAG\|C\|CCCCCCACTCCAGTCCTTCCCCAAGGATGCAGCCTCAGCCTTCTCAGGACTCCAATATCGCTCACCCG |
| 2:2111:5617:15868#2#R | \| \| |
| MQ=60 isize=-127 C=37 | cccggccccacccccacaaggccagcagatccCTAATtCtCcCTCCAATCaAGTGcgCtCtCCCCagcctgTCCCtTCtCCaCGGCCaCAGTCCCAG\|C\|CCCCCCACTCCAGTCCTTCCCCAAGGACGCAGGACTCCAATTACCTACTACAT |
| 1:1209:15887:24102#2#F | \| \| |
| MQ=60 isize=239 C=2 | GAATGGTGTACAAtTgGggTcntcaAGGCCAGCAGATCCCTAAtTcTctcTCCAATCAagtgcgctctccccagcctgtcccttctccacggccacagtcccag\|c\|cccccccctcccgtccttccccaaggctgcagcctcatccttctcc |
| 1:1102:15126:49425#2#R | \| \| |
| MQ=60 isize=-147 C=37 | CAAGGCCAGCAGATCCCTAATTCtCtCTCCAATCAAGTgcgCtCTCCCCAGCCTGtCCCTTCtCcaCGGCCACAGTCCCAG\|C\|CCCCCCACTCCAGTCCTTCCCCAAGGATGCAGCCTCAGCCTTCTCCanGACTCCAATTGGTTAATCTGC |
| 1:1103:9528:2899#2#R | \| \| |
| MQ=60 isize=-147 C=37 | CAAGGCCAGCAGATCCCTAaTTCTCTCTCCAATCAAgTgCGCtCTCCCCAGCCTGTCCCTTCTCCaCGGCCACAGTCCCAG\|C\|CCCCCCACTCCAGTCCTTCCCCAAGGATGCAGCCTCAGCCTTCTCCAGGACTCCAATCTTCCCATACGA |
| 1:1110:15467:59848#2#R | \| \| |
| MQ=60 isize=-147 C=22 | cAAGGCCAGCagaTCCCTaAttCTCtCtCcAATCAAGTGCgCtCTCCCCAGCCTGTCCCTTCTCCACgGCCaCAGTCCCAG\|C\|CCCCCCACTCCAGTCCTTCCCCAAGGATGCAGCCTCAGCCTTCTCCAGGACTCCAATTATAGCCGCGAC |
| 1:1207:10225:16676#2#R | \| \| |
| MQ=60 isize=-147 C=33 | CAAGGCCAGCAGATCCCTAATTCTCTCTCCAATCAAGTGCGCTCTCCCCAGCCTGTCCCTTCTCCACGGCCACAGTCCCAG\|C\|CCCCCCACTCCAGTCCTTCCCCAAGGATGCAGCCTCAGCCTTCTCCanGACTCCAATCATGACCTAAGT |
| 1:2113:3891:57270#2#R | \| \| |
| MQ=60 isize=-147 C=2 | caaggccagcagacccctaattccctctccaatcaagtgcgctctccccagcctgtcccttctccacggccacagtcccag\|c\|ccccccactcCAGTCCttCCCcAAGGaTGCAGCCTCAGCCTTCTCcannACTCCAATCtTcCCATaCGA |
| 1:2201:2256:85458#2#R | \| \| |
| MQ=60 isize=-147 C=33 | CAAGcCCAGCAGATCCcTAattCtCTCTCCAATCAAGTGCGCTCTCCCCAGCCTGTCCCTTCTCCACGGCCACAGTCCCAG\|C\|CCCCCCACTCCAGTCCTTCCCCAAGGATGCAGCCTCAGCCTTCTCCAGGACTCCAATAGCCTGTCGTGT |
| 1:2206:13697:54546#2#R | \| \| |
| MQ=60 isize=-147 C=37 | CAAGGCCAGCAGATCCCTAAttctctccCCAAtCaaatgCgCTCTCCCCAGCCTGTCCCTTCtCCACGGCCACAGTCCCAG\|C\|CCCCCCACTCCAGTCCTTCCCCAAGGATGCAGCCTCAGCCTTCTCCanGACTCCAATCTTCCCATACGA |
| 2:1111:14541:74142#2#R | \| \| |
| MQ=60 isize=-147 C=33 | caagGCCAGCAGATCCcTAATTCTCTCTCCAATCAagTGCGCtCTCCCCAGCCTGTCCCTTCTCCACGGCCACAGTCCCAG\|C\|CCCCCCACTCCAGTCCTTCCCCAAGGATGCAGCCTCAGCCTTCTCCAGGACTCCAATCCTAGGCCTGTT |
| 2:1204:13192:69775#2#R | \| \| |
| MQ=60 isize=-147 C=33 | CAAGGCCAGCAGATCCCTAATTCTCTCTCCAatCAAGtGCGCTCTCCCCAGCCTGTCCCTTCTCCACGGCCACAGTCCCAG\|C\|CCCCCCaCTCCAGTCCTTCCCCAAGGATGCAGCCTCAGCCTTCTCCanGACTCCAATACCGAGTGGGGT |
| 2:2110:10451:91394#2#R | \| \| |
| MQ=60 isize=-147 C=22 | caaggcccgcatatccctaattcccgctccaatccagtgcgctctccccagcctggcccttctccacgcCCACAGTCCCAG\|C\|CCcCCcACTCCAGTCCTTCCCCAAGGATGCAGCCTCAGcCTTCTCCAGGACTCCAAtGGCTAACCCCAA |
| 2:2116:12778:74983#2#R | \| \| |
| MQ=60 isize=-147 C=37 | CAAGGCCAGcaGaTCCcTAATTCtCTCTCCAAtCAAgTGCGCTCTCCCCagCCTGTCCCTTCTCCACGGCCACAGTCCCAG\|C\|CCCCCCACTCCAGTCCTTCCCCAAGGATGCAGCCTCAGCCTTCTCCAgnACTCCAATGTACGTTAAAAA |
| 2:2207:9759:65184#2#R | \| \| |
| MQ=60 isize=-147 C=2 | caaggccagcagctccctaattctctccccaatcaagtgcgctctccccagcctgtccctcccccacggccccagtcccag\|c\|ccccccactccagtccttccCCAAGGATGCAGCCTCAGCCTTCTCcangtctctaatAACGcgcCCCCC |
| 2:2215:12875:89136#2#R | \| \| |
| MQ=60 isize=-147 C=37 | CAAGGCCAGCAGatCCCTAAttctCtCTCCAaTCAAGTGCGCtCtCCCCaGCCTGTCCCTTCTCAACGGCCACAGTCCCAG\|C\|CCCCCCACTCCAGTCCTTCCCCAAGGATGCAGCCTCAGCCTTCTCCAGGACTCCAATCTGGCCCGCGCG |
| 1:2204:16414:82876#2#R | \| \| |
| MQ=60 isize=-144 C=37 | cagaaggccagcatatccctaattccccctCCAAtCaAGtgCGCtCTCCaCAGCCtGTCCCTTCtCCACGGCCaCAGTCCCaG\|C\|CCCCCCACTCCAGTCCTTCCCCAAGGATGCAGCCTCAGCCTTCTAGGACTCCAATCTACCAGCTGCG |
| 1:1103:11122:33041#2#R | \| \| |
| MQ=60 isize=-149 C=27 | aggCCAGCAGatCCCTAatcCTCTCTCCAATCAAgtGCgCtCTCCCCAGCCTGTCCCTTCtCCACGGCCACAGTCCCAG\|C\|CCCCCCACTCCAGTCCTTCCCCAAGGATGCAGCCTCAGCCTTCTCCACAGGACTCCAATATATATGCTCTC |
| 1:1211:10917:16029#2#R | \| \| |
| MQ=60 isize=-149 C=22 | aggCCagcAgaTCCCTAAttctCTctCCAAtcaaGTGcgctctCCCCAGCCTgccCcTtCtCCaCgGCCACAGTcCcac\|C\|CCCCCCACtCCAgTCCTTCCCCAAGGATGCAgCCtCAGCCTTCTCCACAGGACCCCAATGATATACAAACG |
| 1:2106:19865:53118#2#R | \| \| |
| MQ=60 isize=-149 C=33 | AGGCcaGCAgATCCCTAATtCTCTccCCAATCAAGTGCgctCtCCcCAGCCTGTCCCTTCTCCACGGCcacAGTCCCAG\|C\|CCCCCCaCTCCAGTCCTTCCCCAAGGATGCAGCCTCAGCCTTCTCCACanGACTCCAATTGATATTCCAAC |
| 1:2211:10073:80702#2#R | \| \| |
| MQ=60 isize=-149 C=27 | aggccagcagatccctaattctctctcCAAtCaAGTGCGCtcTCCcCAGCCTGTCCCtTCtCCACGGCCaCAGTCCCAG\|C\|CCCCCCACtCCAGTCCTTCCCCAAGGAtGCAGCCTCAGCCTTCTCCACAGGACTCCAATCCCATCCGGAAT |
| 2:1104:19234:45960#2#R | \| \| |
| MQ=60 isize=-149 C=37 | aggccagccgttCCCtAAttCcCtctCCAAtCAAGTgCGcTCTCCCCAGcCTGTCCcTtCTCCACGGCCACAGTCCCAG\|C\|CCCCCCACTCCAGTCCTTCCCCAAGGATGCAGCCTCAGCCTTCTCCACAGGACTCCAATGGCTAGATGAAT |
| 2:1113:16145:76094#2#R | \| \| |
| MQ=60 isize=-149 C=33 | aGGCCAGCagacCcCTaattCtctCtCCAAtCaaGTgCgctctCCCCaGCCTGtCCCTtCTCCaCGGCCaCAGTCCCAG\|C\|CCCCCCaCTCCAGTCCTTCCCCAAGGATGCAGCCTCAGCCTTCTCCACAgnACTCCAAtCGACCTCCCCCT |
| 2:1211:6488:18971#2#R | \| \| |
| MQ=60 isize=-149 C=33 | AGGCCAGCaGaTCCCTAATtcTCtCTCCAATCAAGTGcgCtCTCCCCAGCcTGTCCCTTCTCCACGGCCaCAGTCCCAG\|C\|CCCCCCaCTCCAGTCCTTCCCCAAGGATGCAGCCTCAGCCTTCTCCACAGGACTCCAATCCGCTGCGCGGC |
| 2:2213:9344:97571#2#R | \| \| |
| MQ=60 isize=-149 C=27 | aggccagCAGATCCCtaattcTCtCTCccaTCAAGTgcgctcTCCCCAGCCTGtcCCTTCtCCaCGGcctCagtccCaG\|C\|CCCCcCACtCCaGTCCTtCcCCAAGGATGCAGCCTCAGCCtTCTccACAGGACTCCAATTGAtATTCCAAC |
| 2:2215:13708:23861#2#R | \| \| |
| MQ=60 isize=-149 C=37 | aggCCAGCAGAtCCCTAAttccCTCTCCAaTCAAGtGCGCtCTCCCCAGCCTGTCCCTtCTCCACGGCCACAGTCCCAG\|C\|CCCCCCACTCCAGTCCTTCCCCAAGGATGCAGCCTCAGCCTTCTCCACAGGACTCCAAtCCGCTGCGCGGC |
| 1:1113:15049:15944#2#F | \| \| |
| MQ=60 isize=236 C=22 | GCATAGGTCGTGATTGGAGTCCTGGCCAGCAGATCCCTAATTCTCTCTCCAATCAAGTGCGCTCTCCCCAGCCTGTCCCTTCTCCACGGCCACAGTCCCAG\|C\|CCCCCCACTCCAGTCCTTCCCCAAGGATGCAGCCTCAGCCTTCTCCACa |
| 1:2104:7117:90583#2#F | \| \| |
| MQ=60 isize=236 C=2 | CtAACCCGCcACATTGGAGTCntGGCCagcAGAtcccTAaTTCTCTCTCCAATCAAgtgcgctctccccagccagtcccttctccacggccacagtcccag\|c\|ccccccactccagtccttccccaaggatgcagcctcagccttctccaca |
| 1:1107:8454:44999#2#R | \| \| |
| MQ=60 isize=-126 C=37 | GGCCAGCAGATCCCTAATTCTCTCTCCAaTCAAgTGCgCtCTCCCCAGCCTGTCCCTTCTCCACGGCCACAGTCCCAG\|C\|CCCCCCACTCCAGTCCTTCCCCAAGGCTGAGGCTGCATCCTTGGGGAAGAGGACTCCAATGACAAATTAAAC |
| 1:2213:12084:42702#2#R | \| \| |
| MQ=60 isize=-149 C=37 | GGCCAGCAGATCCCTAATTCTCtCtCCaATCAAGTGCGCTCtCCCCAGCCTGTCCCTTCTCCACGGCCACAGTCCCAG\|C\|CCCCCCACTCCAGTCCTTCCCCAAGGATGCAGCCTCAGCCTTCTCCACAAgnACTCCAATCCGCTTAAGCGC |
| 2:2108:16879:73298#2#R | \| \| |
| MQ=60 isize=-149 C=27 | gcccagcagatccctaattctctctccaatcaagtgcgctctgcccagcctgtcccttctccacggccacagtCccAg\|C\|CCCccCACTCCAGTCCTTCCCCAAGGATGCAGCCTCAGCCTTCTCCACAAGGACTCCAATCACTCTCATACC |
| 1:1112:8483:76532#2#F | \| \| |
| MQ=60 isize=235 C=33 | CCGGAAGCGGGTATTGGAGTncTGcCAGCAGATCCCTAATTCTCTCTCCAATCAAGTGCGCTCTCCCCAGCCTGTCCCTTCTCCACGGCCACAGtCCCAG\|C\|CCCCCCACTCCaGTCCTTCCCCAAGGATGCAGCCTCAGCCTTCTCCACAC |
| 2:1202:2499:78795#2#F | \| \| |
| MQ=60 isize=235 C=22 | TCTCTGGGGTTATTGGAGTCCngCCAGCAGATCCCTAATTCTCTCTCCAATCAAGTGCGCTCTCCCCAGCcTGTCCCTTCTCCACGgCCACAGTCCCAg\|C\|CCCCCCaCtCCAGtCCTTCCCCAAGGAtGCAGCCTCagCCTTCTCCacccc |
| 1:1107:2255:25876#2#R | \| \| |
| MQ=60 isize=-150 C=33 | gCCAGCAGATCCCTAATtCTCtCcCCAATCAAGTGCGCTCTCCCCAGCCtGTCCCtTCtCCACGGCCACAGTCCCAG\|C\|CCCCCCACTCCAGTCCTTCCCCAAGGATGCAGCCTCAGCCTTCTCCACACAGGACTCCAATCTTTACATTTCG |
| 1:2101:10323:57114#2#R | \| \| |
| MQ=60 isize=-150 C=33 | GCCAGCAGaTCCCTAATTCTCTCTCCAATCAAGTGCGctCTCCCCAGCCTGTCCCTTCTCCACGGCCACAGTCCCAG\|C\|CCCCCCaCTCCAGTCCTTCCCCAAGGATGCAGCCTCAGCCTTCTCCACACAGGACTCCAATGATGATCCAGCT |
| 1:2213:1789:62094#2#R | \| \| |
| MQ=60 isize=-150 C=33 | gccagcagatccctaattctCTCTCCAATCAAGTgCgCTCtCCCCAGCCTGTCCCTTCTCCACGGCCACAGTCCCAG\|C\|CCCCCCACTCCAGTCCTTCCCCAAGGATGCAGCCTCAGCCTTCTCCACACAgnACTCCAATGGTCGTCGCGCG |
| 2:1208:14335:65264#2#R | \| \| |
| MQ=60 isize=-508 C=2 | gccagcagatccctaatacgccccccaatcaagtgcgctctccccagcctgtcccttctccacggctacaggcccag\|c\|ccccccacaccAGTCCTtCCCCAAGGaTGcAGCCTCAGCCTTctcCAcacanGACTCCAATAtCAaGTGtCGG |
| 2:2201:10899:41382#2#R | \| \| |
| MQ=60 isize=-150 C=37 | GCCAGCAGATCCCTAAttCTCTCTCCAATCAAGTGCgctCtCCCCAGCCTGTCCCTTCTCCACGGCCACAGTCCCAG\|C\|CCCCCCACTCCAGTCCTTCCCCAAGGATGCAGCCTCAGCCTTCTCCACACanGACTCCAATCTCTGTATCGTG |
| 1:1113:18011:19293#2#F | \| \| |
| MQ=60 isize=234 C=33 | CTTCCTACTGGTATTGGAGTCCTCCAGCAGATCCCTAATTCTCTCTCCAATCAAGTGCGCTCTCCCCAGCCTGTCCCTTCTCCACGGCCACAGTCCCAG\|C\|CCCCCCACTCCAGTCCTTCCCCAAGgaTGCAGCCTCAGCCTTCTCCACACC |
| 1:1208:5962:39042#2#R | \| \| |
| MQ=60 isize=-152 C=37 | CCAGCAGaTCCCTAATTCTCtCtCCAATCAAGTGCGCTCTCCCCAGCCTGTCCCTTCTCCACGGCCACAGTCCCAG\|C\|CCCCCCACTCCAGTCCTTCCCCAAGGATGCAGCCTCAGCCTTCTCCACACcanGACTCCAATCAACCAAGGCCG |
| 2:2210:16382:100159#2#R | \| \| |
| MQ=60 isize=-152 C=37 | CCAGCAGaTCCCTAAttCTCtCTCCAATCAAGTGCGCtctCCCCAGCCTGTCCCTTCTCCACGGCCACAGTCCCAG\|C\|CCCCCCACTCCAGTCCTTCCCCAAGGATGCAGCCTCAGCCTTCTCCACACCAGGACTCCAATACCTCGGCAGTC |
| 1:1204:7492:66628#2#F | \| \| |
| MQ=60 isize=367 C=22 | TTAGTTTCGGCTAtAGGAgTCntcAGCAGATCCCTAATTCTCTCTCCAATCAAGTGCGCTCTCCCCAGCCTGTCCCTTCTCCACGGCCaCAGTCCCAg\|C\|CCCCCCACTCCAGTCCTTCCCCAAGGATGCAGCCtCAGCCTTCtCCACaccc |
| 2:2102:11873:98105#2#F | \| \| |
| MQ=60 isize=233 C=2 | CAGACTCACTATaTTGGAGTCCTCAGCAGATCCCTAATTCTCTCTCCAATCAAGTGCGCTCTCCcCAGCcTGtCCCTTCTCCAcgGCCacagtcccag\|c\|ccccccactccagcccttccccaggatgctacccccacccttcccccccccc |
| 2:1105:16795:30411#2#F | \| \| |
| MQ=60 isize=337 C=33 | GTGGGGCTGGTGATTGGAGTCntaGCAGATCCCTAATTCTCTCTCCAATCAAGTGCGCTCTCCCCAGCCTGTCCCTTCTCCACGGCCACAGTCCCAG\|C\|cCCCCCACTCCAGTCCTTCCCCAAggATGCAGCCTCAGCCttCTCCaCACCAC |
| 2:2109:20118:52610#2#F | \| \| |
| MQ=60 isize=232 C=33 | GATGCAATTGGGATTGGAGTCCTAGCAGATCCCTAATTCTCTCTCCAATCAAGTGCGCTCTCCCCAGCCTGTCCCTTCTCCACGGCCACAGTCCCAG\|C\|CCCcCCACTCCAGTCCTTCCCCaAggatgCAGCCTCAGCCTTCTCCACACCAC |
| 2:2201:18192:27408#2#F | \| \| |
| MQ=60 isize=337 C=33 | GTGGGGCTGGTGATTGGAGTCntAGCAGATCCCTAATTCTCTCTCCAATCAAGTGCGCTCTCCCCAGCCTGTCCCTTCTCCACGGCCACAGTCCCAG\|C\|CCCCCCACTCCAGTCCTTCCCCAAGGaTGCAGCCTCAGCCTTCTCCccaccac |
| 2:1210:9556:92949#2#R | \| \| |
| MQ=60 isize=-153 C=37 | aGCAgaTCCctaATTCtCTCtCCAATCAagtgCgCTCTCCCCAGCCTGTCCCTTCTCCACGGCCACAGTCCCAG\|C\|CCCCCCACTCCAGTCCTTCCCCAAGGATGCAGCCTCAGCCTTCTCCACACCACAGGACTCCAATGCGGCGCGGTGC |
| 1:2109:7233:85385#2#F | \| \| |
| MQ=60 isize=231 C=33 | AACTGTCACGCGATTGGAGTCntgCAGATCCCTAATTCTCTCTCCAATCAAGTGCGCTCTCCCCAGCCTGTCCCTTCTCCACGGCCACAGTCCCag\|C\|CCCCCCACTCCagTCCTTCCCCAAgGAtGCaGCCTCAGCCTTCTCCACACCacg |
| 2:2210:7659:19061#2#F | \| \| |
| MQ=60 isize=231 C=33 | AACTGTCACGCGATTGGAGTCCTGCAGATCCCTAATTCTCTCTCCAATCAAGTGCGCTCTCCCCAGCCTGTCCCTTCTCCACGgCCACAGTCCCAG\|C\|CCCCCCACTCCAGTCCTTCCCCAAgGAtGCAGCCTCAGCCTtcTCCacaCCACg |
| 1:2112:14136:50109#2#F | \| \| |
| MQ=60 isize=230 C=33 | CCTGTCGAGCAAATTGGAGTCCTCAGATCCCTAATTCTCTCTCCAATCAAGTGCGCTCTCCCCAGCCTGTCCCTTCTCCACGGCCACAGTCCCAG\|C\|CCCCCCACTCCAGtCCTTCCCCAAGGATGCAGCCTCAGCCTtCTCCACACCACgt |
| 2:1213:18755:28967#2#F | \| \| |
| MQ=60 isize=230 C=27 | GTGGaTGTATGCATTGGAGTCCTCAgATCCCTAATTCTCTCTCCaATCAAGTGcgCTCTCCCCAtCCTGTcCCTTCTcCACggCCACagTCcCAG\|C\|cCCCCCaCTCcagtCCttCCCCAaggatgCagCCtcagCcTTCTCCACACCACgt |
| 1:1210:5445:82505#2#R | \| \| |
| MQ=60 isize=-156 C=33 | agatCCCTAATtCTCtCtCCAagCaAGTGCgctCtCCCCagCCTGtCCCTtCtCCACGgCcaCAGTCCCAG\|C\|CCCCCCACTCCAGTCCTTCCCCAAGGATGCAGCCTCAGCCTTCTCCACACCACGTTAGGACTCCAATCCAATCTGCGAC |
| 2:1116:19816:59304#2#R | \| \| |
| MQ=60 isize=-156 C=22 | cgatccctAAttCTctcTCCAATCaagTGCgCtctCCCCAGCCTGTCcCtTCTCctcGGCCACAGTCCCAG\|C\|CCCCCCACTCCAGTCCTTCCCcAAGGATGCAGCCTCAGCCTTCTCCACACCACGTTAGGACTCCAATCTTtTTGACaCC |
| 1:1211:11562:14360#2#F | \| \| |
| MQ=60 isize=228 C=22 | GGTCGCCCCGCCATTGGAGTCCTGATCCCTAATTCTCTCTCCAATCAAGTGCGCTCTCCCCAGCCTGTCCCTTCTCTACGGCCACAGTCCCAg\|C\|CCCCCCACtCCaGtCCTTCCCCAAGGATgCAGCCTCAGCCTTCTCCACACCaCGTTt |
| 2:1207:1942:70376#2#R | \| \| |
| MQ=60 isize=-155 C=27 | cccgatccctaattctctctccaatccagtgcgctctccccagcctgtcccttctccACGGCcacAGTCCCAG\|C\|CCCCCCACTCCAGTCCTTCCCCAaGGATGCAGCCTCAGCCTTCTCCACaCCaCgtnAGGACTCCATCAAATTCACAT |
| 1:2106:7874:58853#2#R | \| \| |
| MQ=60 isize=-158 C=37 | aTCCCTAattCtCTCTCCAATCAAGTgcGCTCTCCCCAGCCTGTCCCTTCTCCACGGCCACAGTCCCAG\|C\|CCCCCCACTCCAGTCCTTCCCCAAGGATGCAGCCTCAGCCTTCTCCACACCACGTTTCanGACTCCAATGTAATCCTCTTT |
| 1:2211:17686:70656#2#R | \| \| |
| MQ=60 isize=-158 C=33 | ATCCCTAAttCTCTCtCCAATCAAGtGCGctCcCCCCAGCCTGTCCCTTctCCACGGCCACAGTCCCAG\|C\|CCCCCCACTCCAGTCCTTCCCCAAGGATGCAGCCTCAGCCTTCTCCACACCACGTTTCAGGACTCCAATGGTTGCCGGAAC |
| 2:1209:13761:90965#2#R | \| \| |
| MQ=60 isize=-158 C=37 | atCCcTAATtCTCtCTCCAATCAAGtgCGCTCTCCCCAGCCTGTCCCTtctCCACGGCCACAGTCCCAG\|C\|CCCCCCACTCCAGTCCTTCCCCAAGGATGCAGCCTCAGCCTTCTCCACACCACGTTTCAGGACTCCAATGACCCAGTGCGC |
| 1:1201:8153:4334#2#R | \| \| |
| MQ=60 isize=-160 C=33 | cccctaattccctctccaatcaagtgcgCTCtCCCCAgCCTGTCCCTTCtCCACGgCCACAGT\_CCAG\|C\|CCCCCCaCtCCAGTCCTTCCCCAAGGATGCAGCCTCAGCCTTCTCCACACCACGTTTCCCanGACTCCAATGCGTACAGTGGC |
| 1:2101:19338:3433#2#R | \| \| |
| MQ=60 isize=-159 C=37 | tCCCTAATTCtCTCTCCAATCAAGTGCGCTCTCTCCAGCCTGTCCCTTCTCCACGGCCACAGTCCCAG\|C\|CCCCCCACTCCAGTCCTTCCCCAAGGATGCAGCCTCAGCCTTCTCCACACCACGTTTCCAGGACTCCAATACGCGGGGAGGG |
| 1:2109:2901:88984#2#R | \| \| |
| MQ=60 isize=-159 C=2 | cccctaattctccctccaatcaagtgcgctctccccagcctgtcccttctccacggccacagtcccag\|c\|ccccccactccagtcCTTCCCCAAGGATGCAGCCTCAGCCTTCTCCACACCACGTTTCCanGACTCCAATCaCTCATTCCCA |
| 1:2210:10150:79767#2#R | \| \| |
| MQ=60 isize=-159 C=27 | tCCCTAAttCCCTCTCCAATCAAGtGCGCtCtCCCCAGCCTGtcCCTTCTCCACGGCCACAGTCCCAG\|C\|CCCCCCaCTCCAGTCCTTCCCCAAGGATGCAGCCTCAGCCTTCTCCACACCACGTTTCCAGGACTCCAATTGTGCGACTTAC |
| 2:1102:17463:26261#2#R | \| \| |
| MQ=60 isize=-159 C=33 | tccctaattccctctcCAATCATGTgCGCtctcCccAGCCTGTCcCTTCtCcACGGCcACAGTCCCaG\|C\|CCcCcCACTCCAGTCCTTCCCCAAGGATGCAGCCTCAGCCTTCTCCACACCACGTTTCCAGGACTCCAATACTATACAAAtC |
| 2:1202:18768:93131#2#R | \| \| |
| MQ=60 isize=-159 C=33 | tCCCTAATtCTCTCTCCAATCAAGTGCGCTCTCCCCAGCCTGTCCCTTCTCCACGGCCACAGTCCCAG\|C\|CCCCCCACTCCAGTCCTTCCCCAAGGATGCAGCCTCAGCCTTCTCCACACCACGTTTCCanGACTCCAATCACTCATTCCCA |
| 2:1208:19006:17283#2#R | \| \| |
| MQ=60 isize=-159 C=27 | tccctaattctCTCTCCAATCAAGtgcgCtCTCCCCAGCCTGTCCCTTCTCCACGGCCacagTCCcAG\|C\|CCCCCCACTCCAGTCCTTCCCCAAGGATGCAGCCTCAGCCTTCTCCACACCACGTTTCCanGACTCCAATCACTCATTCCCA |
| 2:1203:7695:35101#2#F | \| \| |
| MQ=55 isize=225 C=2 | TaaACGtTcggCCttgccctCccCTaAcTCtCtCTCCAaccaagcacgctctccccagcctcacccttctcctctcccacaggccagg\|c\|ccccccactccagtccttccccccggctgccgcctcagccgtccccccaccccggtcccccc |
| 1:1111:11826:90992#2#R | \| \| |
| MQ=60 isize=-160 C=37 | CCCTAATtCTCtCTCCAATCAAGTGCGCtCTCCCCAGCCTGTCCCTtCTCCACGGCCACAGTCCCAG\|C\|CCCCCCACTCCAGTCCTTCCCCAAGGATGCAGCCTCAGCCTTCTCCACACCACGTTTCCCAGGACTCCAATACCGGTTAAGCC |
| 1:1115:4532:40840#2#R | \| \| |
| MQ=60 isize=-160 C=33 | CCCTAattCTCTCtCCAATCaAGTGCGCTCTCCCCAGCCTGTCCCTtCTcCACGGCCACAGTCCCAG\|C\|CCCCCCACTCCAGTCCTTCCCCAAGGATGCAGCCTCAGCCTtCTCCACACCACGTTTCCCAGGACtCCAATACGCGCCCGGAG |
| 1:2109:20616:84559#2#R | \| \| |
| MQ=60 isize=-160 C=33 | CCCTAATtctccCtCcAATCAAGCgCgCTCTCCcCAGCCTGTCCCTtCTccaCGGCCACAGTCCCAG\|C\|CCCCCCACTCCAGTCCTTCCCCAAGGATGCAGCCTCAGCCTTCTCCACACCACGTTTCCCanGACTCCAATTGTACTTTTATC |
| 1:2114:17885:75764#2#R | \| \| |
| MQ=60 isize=-160 C=2 | ccccaattctccctccaatcaagtgcgctctccccagcctgtcccctctccacggccacagtcccag\|c\|ccccccactcCAGtCCtTCCCCAAGGATGCAGCCTCaGCCTTCTCCACACCACGTTtCCCAGGACtcCAAtAACAGTTTAAAT |
| 1:2202:8908:63222#2#R | \| \| |
| MQ=60 isize=-160 C=27 | ccctaattctctctCCAATCaAgTgcgctcccCCCAGCCTGTCCCTTCtCCACGgCCACAGTCCCAG\|C\|CCCCCCACTCCAGTCCTTCCCCAAGGATGCAGCCTCAGCCTTCTCCACACCACGTTTCCCanGACTCCAATGCAGAATCAAAG |
| 1:2216:8593:31830#2#R | \| \| |
| MQ=60 isize=-160 C=33 | CCCTAAttCtCTCtCCAATCAAGTGCGCTCTCCCCAGCCTGTCCCTTCTCCaCGGCCACAGTCCCAG\|C\|CCCCCCACTCCAGTCCTTCCCCAAGGATGCAGCCTCAGCCTTCTCCACACCACGTTTCCCAgnACTCCAATGTGCGGGCAAAT |
| 2:1116:7153:78034#2#R | \| \| |
| MQ=60 isize=-160 C=27 | ccctaattctccccccaatcaagtccgctctccccagcctgtcccttctccccggccacagcCCCaG\|C\|CCCCcCaCtCCagcCCTTCCCCAaGGATGCAGCCtCAGCCTTCTCCACACCACGTTTCCCaGGaCtCCAATGAAACCCCGACT |
| 2:1211:4548:43595#2#R | \| \| |
| MQ=60 isize=-160 C=22 | ccCTAATTctctCTCCaatCaAGTGCGCTCtcCCCAGCCTGTCCCtTCtCCAcGGCCaCAGTCCCAG\|C\|CCCCCCACTCCAGTCCTTCCCCAAGGATGCAGCCTCAGCCTTCTCCACACCACGTTTCCCAGGACTCCAATCACGCTTAGTAG |
| 2:2106:20187:20994#2#R | \| \| |
| MQ=60 isize=-160 C=27 | CCCtAATtCtCtctccAAtcaAGTGCGCTctCCCCAGCCTGTCccTTCTCCACGGCCACAGTCccAG\|C\|CCCCCCACTCCAGTCCTTCCCCAAGGATGCAGCCTCAGCCTTCTCCACACCACGTTTCCCanGACTCCAATGCGTTTCCTATG |
| 2:2111:4920:79148#2#R | \| \| |
| MQ=60 isize=-160 C=33 | ccctaaatccctctcaaatcacgtgcgccctccccagcCTGTCCCcTcTccaCGGCCacAGtCccag\|C\|CCCCCcActcCAGtCCTTcCCCAAGGATGcAgCCTCAgCCtTCTCCACACCACgTTTCcCAGGACTCCAATAACAGTTTAAAT |
| 2:2205:4713:46156#2#R | \| \| |
| MQ=60 isize=-160 C=37 | CCCTAATtctccCtCCAAtCatgtgCGCtCtCCCCAGCCTGTCCCTTCTCCaCGGCCACAGTCCAAG\|C\|CCCCCCACTCCAGTCCTTCCCCAAGGATGCAGCCTCAGCCTTCTCCACACCACGTTTCCCanGACTCCAATCACCATCTTGCA |
| 1:1115:17232:33267#2#F | \| \| |
| MQ=60 isize=224 C=33 | GGTGGACACGAGATTGGAGTCCTCCTAATTCTCTCTCCAATCAAGTGCGCTCTCCCCAGCCTGTCCCTTCTCCACGGCCACAGTCCCAG\|C\|CCCCCCACTCCAGTCCTTCCCCAAGGatgcaGCCTCAGCCTTCTCCACACCACGTTTCCCC |
| 2:1107:12803:36109#2#F | \| \| |
| MQ=60 isize=329 C=27 | GTAGTTTCTTTGaTTGGAGTCntcCTAATTCTCTCTCCAATCAAGTGCGCTCTCCCCAGCCTGTCCCTTCTCCACGgCCACAGTCCCAG\|C\|CCCCCCACTCCAGTCCTTCCCCAaggctgCaGCctcagccttctccacaccccgtttcccc |
| 2:2107:19141:81858#2#F | \| \| |
| MQ=60 isize=224 C=33 | TTACCCTAGGTCATTGGAGTCCTCCTAATTCTCTCTCCAATCAAGTGCGCTCTCCCCAGCCTGTCCCTTCTCCACGGCCACAGTCCCAG\|C\|CCCCCCACTCCAGTCCTTCCCCAaGGATGCAGCCTCAGCCtTCTCCACACCACGtTTCCCC |
| 1:1112:4378:49557#2#R | \| \| |
| MQ=60 isize=-162 C=37 | CCTAATtCtCTCTCCaATCAAGtGCGCTCTCCCCAGCCTGTCCCTtCTCCACGGcCaCAGTCcCaG\|C\|CCCCCCACTCCAGTCCTTCCCCAAGGATGCAGCCTCAGCCTTCTCCACACCACGTTTCCCCAgnACTCCAATAAGCCTTTATAA |
| 1:1208:17039:51193#2#R | \| \| |
| MQ=60 isize=-162 C=22 | cctaattctctctccgatcaagggcgctctccCCAGCCTGTCCCTTCtCcacggCCaCAGTCcCag\|C\|CCCCCCaCtCCAGtCCTTCCCCAAGGAtGCAGCCTCAGCCtTCtCCAcaCCACGTTTCCCCanGACTCCAATACTCAAACATGC |
| 1:2113:3377:94396#2#R | \| \| |
| MQ=60 isize=-162 C=27 | CCTAATtCtCTCTCCAATCAAGTGCGCtCTCCCCAGCCTGTCCCTTCTCCACGGCCACAGTCCCAG\|C\|CCCCCCACTCCAGTCCTTCCCCAAGGATGCAGCCTCAGCCTTCTCCACACCACGTTTCCCcannACTCCAATTACAAGCCAACC |
| 1:2206:14677:43087#2#R | \| \| |
| MQ=60 isize=-162 C=37 | cgtaattccccctccaatcaagtgcgccctCCCCagaCTGTCCCtTctCcaCGGCCaCAGTCCCaG\|C\|CCCCCCaCtCCAGTCCTTCCCCAAGGAgGCAGCCTCAGCCTTCTCCACACCAcGTTTCCCCanGACTCCAATCTAGCCGCCGGC |
| 1:2207:7719:64058#2#R | \| \| |
| MQ=60 isize=-162 C=37 | CCTAAtTCTCTCtCCaatcaAgtgcgCtCTCCCCAGCCTGTCCCTTCTCCACGGCCACAGTCCCAG\|C\|CCCCCCACTCCAGTCCTTCCCCAAGGATGCAGCCTCAGCCTTCTCCACACCACGTTTCCCCanGACTCCAATTGCAAAGCCCAT |
| 1:2215:15206:81361#2#R | \| \| |
| MQ=60 isize=-162 C=33 | cctaattctctctccaatcaagtgcgctctCCcCaGCCTGtCCCTTCTCcaCGGcCACaGTCCCAG\|C\|CCCCCCACTCCAGTCCTTCCCCAAGGATGCAGCCTCAGCCTTCTCCACaCCACGTTTCCCCAgnACTCCAATTGGATTTAATTT |
| 2:1111:5163:44641#2#R | \| \| |
| MQ=60 isize=-162 C=2 | cctaaatccctctccaaccaagtgcggcccccccagcctgtcccctctccccggccacagtcccag\|c\|CCCCCCaCtCCAGTCCTTCCCCAAGGATGCAGCCTCAGCCTTCTCCACACCACGTTTCCCCAGGACTCCAATTGTCCTGACGTC |
| 2:1203:4883:70742#2#R | \| \| |
| MQ=60 isize=-162 C=33 | ccttattctctctCCAATCAAGtgCGCtCtCCcCaGcctgTCCCtTCTCCACGGcCaCAGTCcccg\|C\|CCCCCCaCTCCAGTcCTTCCCCAAGGATGCAGCCTCAGCCTTCTCCACACCACGTTTCCCCAGGACTCCAATCCCACACCCCTC |
| 2:1206:12123:41760#2#R | \| \| |
| MQ=60 isize=-162 C=37 | CCTAATTCTCTCTCCAATCAAGTGCGCTCTCCCCAGCCTGTCCCTTCtCcaCGGCCACAGTCCCAG\|C\|CCCCCCACTCCGGTCCTTCCCCAAGGATGCAGCCTCAGCCTTCTCCACACCACGTTTCCCCanGACTCCAATAGCTGGAGAGGC |
| 2:1216:18750:95942#2#R | \| \| |
| MQ=60 isize=-162 C=33 | CCTAATTctccccCCAAtCAaGtGCGCtctCCCcAgCCTGtCCCTTCtCcaCGGCCACAGTCCCag\|C\|CCCCCCaCTCCAGTCCTTCCCCAAGGATGCAGCCTCAGCCTTCTCCACACCACGTTTCCCCAgnACTCCAATCTAGCCGCCGGC |
| 2:2105:12591:58791#2#R | \| \| |
| MQ=60 isize=-162 C=33 | cctaattccccctccaatcaagtgcgctctCCCCAGCCTGTCCCTTCtccacGGCCACAGTCCCAG\|C\|CCCCCCaCTCCAGTCCTTCCCCAAGGATGCAGCCTCAGCCTTCTCCACACCACGTTTCCCCanGACTCCAATGGTTCGGGGACA |
| 2:2109:11829:91779#2#R | \| \| |
| MQ=60 isize=-162 C=33 | cctaattccctctccaatcaagtgcgctctcccCAGCCTGTCCCTtCtCcACGGCCACAGtCCCAg\|C\|CCCCCCACTCCAGTCCTTCCCCAAGGATGCAGCCTCAGCCTTCTCCACACCACGTTTCCCCAGGACTCCAATCTTCATCTCCTA |
| 2:2201:12490:100077#2#R | \| \| |
| MQ=60 isize=-162 C=33 | CCTAAttCtCtCtCCAATCAAgtgCgCtCTCCCCAGCCTGTCCCTtCTCCACGGCCACAGTCCCAG\|C\|CCCCCCaCTCCAGTCCTTCCCCAAGGATGCAGCCTCGGCCTTCTCCACACCACGTTTCCCCanGACTCCAATCCGCATGACCTC |
| 2:2211:15500:69750#2#R | \| \| |
| MQ=60 isize=-162 C=33 | CCTAATTCTCTCTCCAATCAaGTGCGCTCTCCCCAGCCTGTCCCTTCTCCACGGCCACAGTCCCAG\|C\|CCCCCCACTCCAGTCCTTCCCCAAGGATGCAGCCTCAGCCTTCTCCACACCACGTTTCCCCAGGACTCCAATATGTCATAATTA |
| 1:1206:16163:47251#2#F | \| \| |
| MQ=60 isize=223 C=33 | AGGTGTAGGACAATTGGAGTCCTCTAATTCTCTCTCCAATCAAGTGCGCTCT\_\_CCAGCCTGTCCCTTCTCCACGGCCACAGTCCCAG\|C\|CCCCCCACTCCAGtcCTTCCCCAAGGATGCAGCCTCAGCCTTCTCCacACCACGTTTCCCCAca |
| 1:1214:6877:67560#2#F | \| \| |
| MQ=60 isize=223 C=37 | CGTCACAAGATGATTGGAGTncTCTAATTCTCTCTCCAATCAAGTGCGCTCTCCCCAGCCTGTCCCTTCTCCACGGCCACAGTCCCAG\|C\|CCCCCCACTCCAGTCCTTCCCCAagGATGCAGCCTCAGCCTTCTCCACACCACGTTTCCCCa |
| 1:2113:14118:68217#2#F | \| \| |
| MQ=60 isize=223 C=2 | TATTTAAGTATGATTGGAGTnntcTAATTCTCTCTCCAATCAAGTGCGCTCTCCCCAGCCTGTCCCTTCTCCACGGCCACAgtcccag\|c\|ccccccactccagtccttccccaaggatgcagcctcagccttctccacaccacgtttcccca |
| 2:1103:15724:39485#2#F | \| \| |
| MQ=60 isize=223 C=27 | ATGTGAAAGAGAATTGGAGTCCTCTAATTCTCTCTCCAATCAAGTGCGCTCTCCCCAGCCTGTCCCTTCTCCACGGCCACAGTCCCAG\|C\|CCCCCCACTCCAgTCCTTCCCCAAGGaTGCAGCCtCAgCCTtCTCCCCACCaCgtTTCCCCC |
| 2:1212:15446:35870#2#F | \| \| |
| MQ=60 isize=223 C=33 | GGGAGCGTCGCCATTGGAGTCCTCTAATTCTCTCTCCAATCAAGTGCGCTCTCCCCAGCCTGTCCCTTCTCCACGGCCACAGTCCCAG\|C\|CCCCCCACtCCAGTCCTTCCCCAAGGATGCAGCCTCAGCCTTCTCCACACCaCGTTTCCCCc |
| 2:2112:5801:29561#2#F | \| \| |
| MQ=60 isize=223 C=33 | TGTCTGCGCATCATTGGAGTncTCTAATTCTCTCTCCAATCAAGTGCGCTCTCCCCAGCCTGTCCCTTCTCCACGGCCACAGTCCCAG\|C\|CCCCCCACTCCagtCCTtCCCCAAGgaTGCAGCCtcaccCTtccccacaCccCGTTtCCCcC |
| 1:1102:20654:86170#2#R | \| \| |
| MQ=60 isize=-164 C=37 | taattctctctCCAATCAaGTGCGCtCtCCCCAGCCTGTCCCTtCTCCaCGGCCACAGTCCCAG\|C\|CCCCCCACtCCAGTCCTTCCCCAAGGATGCAGCCTCAGCCTTCTCCACACCACGTTTCCCCACanGACTCCAATTCATTACGAGTA |
| 1:2104:13738:70450#2#R | \| \| |
| MQ=60 isize=-164 C=37 | TAATtCTCtCTCCAATCAAGTGCGCTCTCCCCAGCCTGTCCCTtCTCCACGGCCACAGTCCCAG\|C\|CCCCCCACTCCAGTCCTTCCCCAAGGATGCAGCCTCAGCCTTCTCCACACCACGTTTCCCCAcanGACTCCAATCAATATAAGCCC |
| 2:1210:1807:31827#2#R | \| \| |
| MQ=60 isize=-165 C=37 | taattctctccccaatcaagtgcgctccCCCCaGCCtGTCCCtTCTCCaCgGCCACAGtCCCAG\|C\|CCCCCCaCtCCAGTCCTTCCCCAAGGATGCAGCCTCAGCCTTCTCCACACCACGTTTCCCCACAGGACTCCAATTAAGCTAAATAA |
| 2:1215:7800:71762#2#R | \| \| |
| MQ=60 isize=-165 C=37 | taATTCTCaCTCCAATCAAGTGCgctCTCCCCaGCCTGTCCCTTCTCCaCGGCCACAGTCCCAG\|C\|CCCCCCACTCCAGTCCTTCCCCAAGGATGCAGCCTCAGCCTTCTCCACaCCACGTTTCCCCACAgnACTCCAATTTCGACTCTCCA |
| 2:2215:15502:48494#2#R | \| \| |
| MQ=60 isize=-161 C=2 | taattctctctccaatcaagtgcgctctccccagcctgtcccttctccacggccacagtcccag\|c\|ccccccactccagtccttccccaaggaTGCAGCCTCAGCCTTCTCCACACCACGtttcCCCgCAGGACTCCAATttCCCTCaCCTC |
| 1:2109:14651:11072#2#R | \| \| |
| MQ=60 isize=-165 C=37 | AATtCTCcCTCCAATCAAGTGCGCtCTCCCCAGCCTGTCCCTTCTCCACGGCCACAGTCCCAG\|C\|CCCCCCACTCCAGTCCTTCCCCAAGGATGCAGCCTCAGCCTTCTCCACACCACGTTTCCCCACAgnGGACTCCAATACCTCTGGACG |
| 1:1211:3443:74994#2#F | \| \| |
| MQ=60 isize=220 C=37 | CCGTCAGTAATTATTGGAGTCCTATTCTCTCTCCAATCAAGTGCGCTCTCCCCAGCCTGTCCCTTCTCCACGGCCACAGTCCCAG\|C\|CCCCCCACTCCAGTCCTTCCCCAAgGATGCAGCCTCAGCCTTCTCCACACCACGTTTCCCCACAg |
| 1:1216:16879:34733#2#R | \| \| |
| MQ=60 isize=-166 C=37 | ATTCtCTCTCCAATCAAGtgCGctctCCCCAGCCTGTCCCTTCTCCaCGGCCACAGTCCCAG\|C\|CCCCCCACTCCAGTCCTTCCCCAAGGATGCAGCCTCAGCCTTCTCCACACCACGTTTCCCCACAGAgnACTCCAATGCCGGCGCTTCC |
| 1:2107:15865:14205#2#R | \| \| |
| MQ=60 isize=-166 C=33 | atcCtCTCTCCAATCAAGTGCGCTCTCCCCAGCCTGTCCCTTCTCCACGGCCaCAGTCCCAG\|C\|CCCCCCaCTCCAGTCCTTCCCCAAGGATGCAGCCTCAGCCTTCTCCACACCACGTTTCCCCACAGAGGACTCCAATATTGCGTTAAAG |
| 2:1112:6322:15130#2#R | \| \| |
| MQ=60 isize=-166 C=33 | attCTCtCTCcaAtCaAGTGcgCtCTCCCCAGCCTGTCCCttcTCCACGGCCaCAGTCCCAG\|C\|CCCCCCACTCCAGTCCTTCCCCAAGGATGCAGCCTCAGCCTTCTCCACACCACGTTTCCCCACAGAGGACTCCAATGAGGATTGATGA |
| 2:1201:19233:86728#2#R | \| \| |
| MQ=60 isize=-166 C=33 | attCTCtctccaATCAAGTGCgCtCTCCCCAGCCTGTCCCTtCTCCACGGCCACAgTCcCAG\|C\|CcCCCCACTCCAGTCCTTCCCCAAGGATGCAGCCTCAGCCTTCTCCACACCACGTTTCCCCACAGAGGACTCCAATAGGATAATTCGT |
| 1:1106:4028:89671#2#F | \| \| |
| MQ=60 isize=219 C=22 | CGGATTCACCCAATTGGAGTCntTCTCTCTCCAATCAAGTGCGCtCTCCCCAGCCTGTCCCtTCTCCACGGCCACAGTCCCAG\|C\|CCCCCCACTCCAgtcCTtCCCCAAGGATGCAGCCTCAGCCTTctccACaCcacgttTCCCCACAGAC |
| 1:1205:4388:99896#2#F | \| \| |
| MQ=60 isize=219 C=33 | ATAAAGAGTCACATTGGAGTCntTCTCTCTCCAATCAAGTGCGCTCTCCCCAGCCTGTCCCTTCTCCACGGCCACAGTCCCAG\|C\|CCCCCCACTCCAGTCCTTCCCCAAGGATGCAGCCTCAGCCTTCTCCaCACCACGtTTcCCCACAGAc |
| 1:2105:5527:61083#2#F | \| \| |
| MQ=60 isize=219 C=33 | GTTAATCCTCGGATTGGAGTCntTCTCTCTCCAATCAAGTGCGCTCTCCCCAGCCTGTCCCTTCTCCACGGCCACAGTCCCAG\|C\|CCCCCCACTCCAGTCCTTCCCCAAGGATGCAGCCTCAGCCTTCTCCACACCACGTTTCCCCACAGAC |
| 1:2109:3296:17287#2#F | \| \| |
| MQ=60 isize=219 C=33 | TCATAACTATCTATTGGAGTCntTCTCTCTCCAATCAAGTGCGCTCTCCCCAGCCTGTCCCTTCTCCACGGCCACAGTCCCAG\|C\|CCCCCCACTCCAGTCCTTCCCCAAGGATGCAGCCTCAGCCTTCTCCACACCACGTTtCCCCACAGAC |
| 2:2103:9952:26946#2#F | \| \| |
| MQ=60 isize=353 C=33 | CGTGTTGTCCTCATTGGAGTCCTTCTCTCTCCAATCAAGTGCGCTCTCCCCAGCCTGTCCCTTCTCCACGGCCACAgtCCCAG\|C\|CCCCCCcctccaGTCcTccCCCAaggatgcagcctcagccttccccccaccccgtttccccccagac |
| 1:1210:18667:83807#2#R | \| \| |
| MQ=60 isize=-166 C=37 | ttctCTCtCCAATCAAGtgCgCtCTCCCCAGCCTGTCCCTTCTCCACGGCCACAGTCCCAG\|C\|CCCCCCaCTCCAGTCCTTCCCCAAGGATGCAGCCTCAGCCTTCTCCACACCACGTTTCCCCACAGAAGGACTCCAATCCCCTCTACCTC |
| 2:1204:15965:32134#2#F | \| \| |
| MQ=60 isize=218 C=33 | GGAGCAAGCTGCATTGGAGTCntCTCTCTCCAATCAAGTGCGCTCTCCCCAGCCTGTCCCTTCTCCACGGCCACAGTCCCAG\|C\|CCCCCCACTCCAGTCCTTcCCCAAGGATGCAGCCTCaGCCTTCTCCACACCACGTTTCCCCACAGACA |
| 1:1102:10780:41113#2#R | \| \| |
| MQ=60 isize=-168 C=27 | cctctctccaaccaagtgcgctctccccagCCTGTCCCTTCTCCaCggccACAgTCCCAG\|C\|CCCCCCACTCCAGTCCTTCCCCAAGGATGCAGCCtCAGCCTTCTCCACACCACGTTTCCCCACAGAcanGACTCCAATATAGATAATGAG |
| 1:2213:8333:60976#2#R | \| \| |
| MQ=60 isize=-168 C=37 | tctCTCTCCAATCAaGtGCGCtccCCCCAGCCTGTCCCTTCtCCaCGGCcaCAGTCCCAG\|C\|CCCCCCACTCCAGTCCTTCCCCAAGGATGCAGCCTCAGCCTTCTCCACACCACGTTTCCCCACAGACAgnACTCCAATAGAGCACCCTAT |
| 2:2107:17073:34343#2#R | \| \| |
| MQ=60 isize=-168 C=33 | tccctctccaatcaagtgcgctcccCCCAGCCTGTCCCTTCTCCACGGCCACAgtCCCAG\|C\|CCCCCCACtCCAGTCCTtCCCCAAGGAtGcAGCCTCAGCCTTCTCCACACCACGTTTCCCCACAGACAGGACTCCAATAGAGCACCCTAT |
| 1:1102:9376:67348#2#R | \| \| |
| MQ=60 isize=-164 C=33 | caatcctctctccaatcaagtgccctctCCCCAgcCTgtcCCTtCtcCacGGCCACAGTCCCAG\|C\|CCCCCCACTCCAGTCCTTCCCCAAGGATGCAGCCTCAGCCTTCTCCACACCACGTTTCCCCAcanGACTCCAATCAGGGGTTCCAC |
| 1:2212:5201:9277#2#R | \| \| |
| MQ=60 isize=-170 C=2 | ctctctccaaccaagtgcgctctccccagcctgtcccttctccacggccacagtcccag\|c\|ccccccactccagtccttcccCAAGGATGCAGCCTCAGCCTTCTCCACACCACGTTTCcCCACAGACaaGGaccCCAatCAtCcCCCCCCA |
| 1:1112:1470:37121#2#F | C    \| \| |
| MQ=60 isize=216 C=33 | GCGCTGCTGGTGATTGGAGTncTTCTCTCCAATCAAGTgCGCcCTCCCCAGCCTGTCCCTTCTCCaCGGCCACAGICCCAG\|C\|ccCCCCACTCCaGtcCTTCCCCAAgGatGcagcCTCagCCttCTCCaCACCaCGTTTCCCCACAGACA |
| 1:2116:17657:57274#2#R | \| \| |
| MQ=60 isize=-170 C=37 | cctCTCCAatCAaGTgCGCTCTCCCCAGCCTGTCCCTTCTCCACGGCCACAGTCCCAG\|C\|CCCCCCACTCCAGTCCTTCCCCAAGGATGCAGCCTCAGCCTTCTCCACACCACGTTTCCCCACAGACAAgnGGAATCCAATAACCCAGCCCG |
| 2:2211:20853:34611#2#R | \| \| |
| MQ=60 isize=-169 C=33 | cCTCTCCAATCAAGTGCGCtCTCCCCAGCCTgTCCCTtCtCCaCGGCCACAGTCCTAG\|C\|CCCCCCaCTCCAGTCCTTCCCCAAGGATGCAGCCTCAGCCTTCTCCACACCACGTTTCCCCACAGACAAAGGACTCCAATCCTTCCCCGAGA |
| 2:1101:4815:49838#2#F | \| \| |
| MQ=60 isize=320 C=27 | ATTCGCTCTGTAATTGGAGTCCTCTCCAATCAAGTGCGCTCTCCCCAGCCTGtCCcTTCTCCACGgCCACAGTCCcaG\|C\|CCCCCccccccagccctcccccacgtaggcgccctcgcccttctccccccccctttcccccccaaccaattc |
| 2:1216:5624:27653#2#F | \| \| |
| MQ=60 isize=215 C=37 | GGTACTTCTGCGATTGGAGTncTCTCCAATCAGGTGCGCTCTCCCCAGCCTGTCCCTTCTCCACGGCCACAGTCCCAG\|C\|CCCCCCACTCCAGTCCTTCCCCAAGGATGCAGCCTCAGCCTTCTCCACACCACGTTTCCCCACAGACAAGTT |
| 2:2116:5862:97116#2#F | \| \| |
| MQ=60 isize=215 C=22 | TTTCTTATCATGATTGGAGTncTCTCCAATcAAgtgCGCTCTCCCCAGCCTgTCCCTTCTCcccggccAcagcCCCac\|C\|ctccCCactccagtcCTtCCCCAAGgatgcggcctcggcctcttccacaccccgcttcacccccgcaaagtc |
| 1:1205:4117:87913#2#R | \| \| |
| MQ=60 isize=-170 C=33 | ctctCCAATCAAGTGCGCTCTCCCCAGCCTGTCCCTTCTCCACGGCCACAGTCCCAG\|C\|CCCCCCACTCCAGTCCTTCCCCAAGGATGCAGCCTCAGCCTTCTCCACACCACGTTTCCCCACAGACAAGanGACTCCAATTCCCATGTGGTG |
| 2:1112:10082:20674#2#R | \| \| |
| MQ=60 isize=-170 C=27 | ccctccaatcaAgTGtGCtcTCCCCAGCCTGtCCCTTCtCCACGGcCACAGtCCCAG\|C\|CCCCCCACTCCAGTCCTtCCCCAAGGATGCAGCCTCAGCCTTCTCCACACCACGTTTCCCCACAGACAAGAGGACTCCAATATCGGCGCAAAA |
| 2:1203:17372:98215#2#R | \| \| |
| MQ=60 isize=-170 C=27 | ctctccaatcaagtgcgctctcCCCAGcCtgtcCCTTctCCaCGgCCACAGTCCCaG\|C\|CCcCCCACTCCAGTCCTTCCCcAAGGATGCAGCCTCAGCCTTCTCCACACCACGTTTCCCCACAGACAAGAGGACTCCAATAGCGTCCAGACT |
| 2:1214:1604:32669#2#R | \| \| |
| MQ=60 isize=-170 C=37 | ctctCCAAtCaAGtgCgCCCtCCCCAGCCtgTCCCTTCTCCACGGCCaCAgtCCCaG\|C\|CCCCCcaCTCCAGTCCTTCCCCAAGGATGCAGCCTCAGCCTTCTCCACACCACGTTTCCCCACAGACAAGAGGACTCCAATGCAGACCGGGAA |
| 2:2110:10219:88444#2#R | \| \| |
| MQ=60 isize=-170 C=37 | CtcTCCAATCaAgtgCgCTCtCCCCAGCCTGtcccTTCTCCACGGCCACAGTCCCAG\|C\|CCCCCCACTCCAGTCCTTCCCCAAGGATGCAGCCTCAGCCTTCTCCACACCACGTTTCCCCACAGACAAGAGGACTCCAATCATCCACTCACG |
| 1:1201:6231:89873#2#F | \| \| |
| MQ=60 isize=213 C=2 | agagggatgaaacccccggttngctccaatcaagagtgctctcccaagcctgtcccttctccacggcccaagtcccag\|c\|ccccccactccagtccttccccaagcatgcagcctcagcctccccccccctccgtttccccccccacactcc |
| 2:2110:10252:6308#2#R | \| \| |
| MQ=60 isize=-169 C=33 | cccctccaatcaagtgcgccctcccCAGCCTGTCCCTTctCCACGGCCACAGtCCCAG\|C\|CCCCCCACTCCAGTCCTTCCCCAAGGATGCAGCCTCAGCCTTCTCCACACCACGTTTCCCCACAGACAAAGGACTCCAAtCCTTCCCCGAGA |
| 1:1109:8900:29951#2#R | \| \| |
| MQ=60 isize=-174 C=33 | CCAATCAAGTGCGCTCTCCCCAGCCTGTCCCTTCTCCACGGCCACAGTCCCAG\|C\|CCCCCCaCTCCAGTCCTTCCCCAAGGATGCAGCCTCAGCCTTCTCCACACCACGTTTCCCCACAGACAAGTTCCAGGACTCCAATGATGGGTATAGA |
| 1:1212:19755:55736#2#R | \| \| |
| MQ=60 isize=-174 C=33 | ccaatcaagtgcgctCTCCCcaGcCTGtCCCTTCtCCaCGGCCaCAGtCCCAG\|C\|CCCCCCaCTCCAGTCCTTCCCCAAGGATGCAGCCTCAgcCTTCTCCACACCaCGTTTCCCCACAGACAAGTTCCAGGACTCCAATGTAtCCCACCCT |
| 1:1216:12771:29324#2#R | \| \| |
| MQ=60 isize=-174 C=37 | CCAatCAAgtcCgCTCTCCCCAGCCtgtcCCTTCTCCACGGCCACAgtCCCAG\|C\|CCCCCCaCtCCAGTCCtTCCCCAAGGATGCAGCCTCAGCCTTCTCCACACCACGTTTCCCCACAGACAAGTTCCAgnACTCCAATACAATCACGGTA |
| 1:2112:8572:45541#2#R | \| \| |
| MQ=60 isize=-174 C=33 | CCAATCAAGTGCgCtCTCCCCAGCCTGTCCCttCTCCACGGCCACAGTCCCAG\|C\|CCCCCCaCTCCAGTCCTTCCCCAAGGATGCAGCCTCAGCCTTCTCCACACCACGTTTCCCCACAGACAAGTTCCAGGACTCCAATAACCAGCTAAAA |
| 1:2211:9674:60503#2#R | \| \| |
| MQ=60 isize=-174 C=22 | CCAATCAAGTGCgCTCTCCCCAGCCTGTCCCTtCTCCACGGCCACAGTCCCAG\|C\|CCCCCCaCTCCAGTCCTTCCCCAAGGATGCAGCCTCAGCCTTCTCCACaCCACGTTTCCCCACAGACAAGTTCCAGGACTCCAATCTCTCCCTACTT |
| 2:1106:1410:56954#2#R | \| \| |
| MQ=60 isize=-174 C=33 | ccaaacaagcgcgccccCCCCAGCCTggCCCtTCcCCaCGGCCcCagtccCcg\|C\|CCCCCCaccCCagTCCtTCCCCAAGGAtgCAGCCTCAGCCTTCTCCACACCACGTTTCCCCACAGACAAGTTCCanGACTCCAATGCCGTTGTTCGG |
| 2:1201:7428:58289#2#R | \| \| |
| MQ=60 isize=-174 C=27 | ccaatcaagtgcgccctCCCCAGCCTGTCCCTTCtCCACGGCCACAGTCCCAg\|C\|CCCCCCaCTCCAGtCCTTCCCCAAgGATGCAGCCTCAGCCTTCTCCACACCACGTTTCCCCACAGACAAGTTCCAGGACTCCAAtCCGTACGTCCCA |
| 2:2209:13878:30589#2#R | \| \| |
| MQ=60 isize=-174 C=37 | CcAAtCAAGtgCGCTCTCCCCAGCCTGTCCCTTCTCCACGGCCACAGTCCCAG\|C\|CCCCCCACTCCAGTCCTTCCCCAAGGATGCAGCCTCAGCCTTCTCCACACCACGTTTCCCCACAGACAAGTTCCanGACTCCAATACGATATAAGTG |
| 2:1103:6127:33874#2#F | \| \| |
| MQ=60 isize=210 C=33 | GCGCTAGGCACCAtTGGAGTCCTCAATCAAGTGCGCTCTCCCCAGCCTGCCCCTACTCCACGGCCACAgcCCCAG\|C\|CCCCCCACcCCAGtCCTccCCCAaggatgcaccctcacccttcccccccccccgtttcccacccgacaagttccc |
| 1:1104:7798:86535#2#R | \| \| |
| MQ=60 isize=-175 C=33 | caatcaagtgcgctccCCCCaGCCTGTCCCTTCTCCACGGCCaCAGTCCCaG\|C\|CCCCCCaCTCCAGTCCTTCCCCAAGGATGCAGCCTCAGCCTTCTCCACACCACGTTTCCCCACAGACAAGTTCCCanGACTCCAATCCCCCTCTACTC |
| 1:1113:4715:79837#2#R | \| \| |
| MQ=60 isize=-175 C=27 | CAATCAAGtGCGCTCTCCCCAGCCTGTCCCTTCTCCACGGCCACAGTCCCAG\|C\|CCCCCCaCTCCAGTCCTTCCCCAAGGATGCAGCCTCAGCCTTCTCCACACCACGTTTCCCCACAGACAAGTTCCCAGGACTCCAAtCCCCCTCTACTC |
| 1:1207:7126:21008#2#R | \| \| |
| MQ=60 isize=-175 C=33 | caatcaagtgcgctcTCCCCaGCCTGTCcCTTCTCcACGGCcACAgTCCcAG\|C\|CCCCCcaCtCCAGTCCTTCCCCAAGGATGCAGCCTCAGCCTTCTCCACACCaCGTTTCCCCACAGACAAGTTCCCanGACTCCAATCCCCCTCTACTC |
| 1:2102:5443:73863#2#R | \| \| |
| MQ=60 isize=-175 C=37 | caatcaagtgcgcccTCCCCAGCCTGTCCCTTCTCCACGGCCACAGTCCCAG\|C\|CCCCCCACTCCAGTCCTTCCCCAAGGATGCAGCCTCAGCCTTCTCCACACCACGTTTCCCCACAGACAAGTTCCCAGGACTCCAATTAACCCGCAATC |
| 1:2107:16901:78282#2#R | \| \| |
| MQ=60 isize=-175 C=33 | CAATCaAGtgCgCtCtCCCCAGCCTGTCCCTTCTCCACGGCCACAGTCCCAG\|C\|CCCCCCaCTCCAGTCCTTCCCCAAGGATGCAGCCTCAGCCTTCTCCACACCACGTTTCCCCACAGACAAGTTCCCAGGACTCCAATCCCCCTCTACTC |
| 1:2202:20365:8689#2#R | \| \| |
| MQ=60 isize=-175 C=37 | CAATCAAGTGCgCtCTCCCCAGCCTGtCCCTTCTCCACGGCCACAGTCCCAG\|C\|CCCCCCACTCCAGTCCTTCCCCAAGGATGCAGCCTCAGCCTTCTCCACACCACGTTTCCCCACAGACAAGTTCCCanGACTCCAATCTAGCGCCCCAC |
| 1:2207:14274:96247#2#R | \| \| |
| MQ=60 isize=-175 C=33 | CAATCAAGTgCgCTCTCCCCAGCCTGtCCCTTCTCCACGGCCACAGTCCCAG\|C\|CCCCCCACTCCAGTCCTTCCCCAAGGATGCAGCCTCAGCCTTCTCCACACCACGTTTCCCCACAGACAAGTTCCCanGACTCCAATCAAGCGCGCCTG |
| 2:1107:8715:5437#2#R | \| \| |
| MQ=60 isize=-175 C=33 | CcATCAAgtgcgccctCCCcagcctggcCcttctCcccgGCCACagtcCcaG\|C\|CCCCccaCTCCAGTCCTTCCCCAAGGATGCAGCCTCAGCCTTCTCCACACCACGTTTCCCCACAGACAAGTTCCCanGACTCCAATCAAATTCGAAGC |
| 2:2202:10759:84997#2#R | \| \| |
| MQ=60 isize=-175 C=37 | CAATCAAGTgCGCTCTCCCCAGCCTGTCCCTTCTCCACGGCCACAGTCCCAG\|C\|CCCCCCaCTCCAGTCCTTCCCCAAGGATGCAGCCTCAGCCTTCTCCACACCACGTTTCCCCACAGACAAGTTCCCAGGACTCCAATTAATACCGGTTG |
| 2:2210:2083:59153#2#R | \| \| |
| MQ=60 isize=-175 C=27 | caaacaagtgcgctctcccccgccTGtCaCtTCTCcacGGCCACagtCcCag\|C\|CCccCccccCCAGTCCTTCCCCAAGGAtgCAGaCTCaGCCtTCTCCACaCCACGTTTCCCCACAGACAAgTTCCCAGGACTCCAATGtTgcCACCATA |
| 2:2215:14036:14493#2#R | \| \| |
| MQ=60 isize=-175 C=2 | caatcaagtgcgccccccccagcctgtcccttctccacggccacagtcccag\|c\|ccccccactccagtccttccccaaggatgCAGCCTCAGCCTTCTCCACACCACGTTTCCCCACAGACAAGTTCCcagGACTCCaaTTTGTGATCAAGT |
| 1:1201:14952:11422#2#F | \| \| |
| MQ=60 isize=314 C=27 | CGGTCATTGCCCATTGGAGTCntaaTCAAGTGCGCTCTCCCCAgCCTGTCCCTTCTCCACGGCCACAGTCCCAG\|C\|CCCCCCaCtCCAGtCCTTCCCCAAGgatgCAGCCTCagCCttCtcCaCacCaCGTTtcCCCacagacaagttcccc |
| 1:2211:14201:34199#2#F | \| \| |
| MQ=60 isize=209 C=37 | TCCGAGAAAAATATTGGAGTCCTAATCAAGTGCGCTCTCCCCAGCCTGTCCCTTCtCCACGGCCAcAGTCCCAG\|C\|CCCCCCacTCCAGtCCTtCCCCAaGgatgCAGCCtcAGCCTTCTCCACACCACgtTTCCccaCAgACAAGTTCCCC |
| 2:2108:6407:96771#2#F | \| \| |
| MQ=60 isize=209 C=2 | TAAATTGTCGTCATTGGAGTCCTAATCAAGTGCgcTcTCCCCAGCCTGTCCCTTCTCCACGGCaACAGtCCcag\|c\|ccccccaccccagtccttcccccaggatgcaccctcacctttctccaacccatcttccccaccaagaaagttcccc |
| 1:1107:10118:42959#2#R | \| \| |
| MQ=60 isize=-177 C=37 | aATCAAGTGCGCtCtCCcCAGCCTGTCCCTTCtCCcCGGCCACAGTCCCAG\|C\|CCCCCCACTCCAGTCCTTCCCCAAGGATGCAGCCTCAGCCTTCTCCACACCACGTTTCCCCACAGACAAGTTCCCCAGGACTCCAATAGATGAAGGGGA |
| 1:1112:10759:2709#2#R | \| \| |
| MQ=60 isize=-177 C=33 | aatcaagtgcgctctccccagcctgtccctTCTCCAcggCcacagtCCCAG\|C\|CCCCcCactCCAGTcCttCCCCAAgGaTGCAGCCTCAgcCTTCTcCACACCACGTtaCCCCACAGACAAGTtCCCCAgnACTCCAATCGCTGGACctCA |
| 1:1202:2920:51564#2#R | \| \| |
| MQ=60 isize=-177 C=22 | aatcaagtgcgctctCCCCAGCCTGtCCCTtCtCCACGGCcACaGtCCCAG\|C\|CCCCCCaCTCCAGTCCTTCCCCAAGGATGCAGCCTCAGCCTTCTCCACACCACGTTTCCCCACAGACAAGTTCCCCAGGACTCCAATAAACCGGCCGAC |
| 1:2107:11320:70182#2#R | \| \| |
| MQ=60 isize=-165 C=37 | aatcaagtgcgctctCCCCAGCCTGTCCCttCTCCACGgCaaCAGTCCCAG\|C\|CCCCCCACTCCAGTCCTTCCCCAAGGATGCAGCCTCAGCCTTCTCCACACCACGTTTCCCCACAGGCTGGAGTGCAAGGACTCCAATACAAGACGCCAG |
| 1:2204:6320:17488#2#R | \| \| |
| MQ=60 isize=-177 C=33 | aaTCaagtgCGctCTCCCCAGCCTGTCCCttctcCACGGCCACAGTCCCAG\|C\|CCCCCCACTCCAGTCCTTCCCCAAGGATGCAGCCTCAGCCTTCTCCACaCCACGTTTCCCCACAGACAAGTTCCCCAGGACTCCAATGATAGGTGGCAA |
| 2:1103:16507:69684#2#R | \| \| |
| MQ=60 isize=-177 C=22 | aatcaagtccgctctccccagcctggcccttccccacGGCCACAGTCCCAG\|C\|CCCCCCACTCCAGTCCTTCCCcAAGGATGCAGCCtCAGCCTTCtCCACACCACGTTTCCCCACAGACAAGTTCCCCAGGACTCCAATGATAGGTGGCAA |
| 2:1203:14499:43994#2#R | \| \| |
| MQ=60 isize=-177 C=27 | AATCAAGTgccCtCTCCCCAgCCTgtcCCtTCTCCACGGCCACAGTCCCaG\|C\|CCCCCCACTCCAGTCCTTCCCCAAGGATGCAGCCTCAGCCTTCTCCACACCACGTTTCCCCACAGACAAGTTCCCCAGGACTCCAATTACAATACCGCT |
| 2:1206:17344:59281#2#R | \| \| |
| MQ=60 isize=-177 C=37 | AATCAAGTGCGCTCTCCCCAGCCTGTCCCTTCTCCACGGCCACAGTCCCAG\|C\|CCCCCCACTCCAGTCCTTCCCCAAGGATGCAGCCTCAGCCTTCTCCACACCACGTTTCCCCACAGACAAGTTCCCCanGACTCCAATCAAAGGCCTCCC |
| 2:2107:19809:27711#2#R | \| \| |
| MQ=60 isize=-177 C=33 | aATCAAGtGCgCTCTCCCCAGCCTGTCCCTTCTCCACGGCCACAgcCCCAG\|C\|CCcCCCACTCCAGTCCTTCCCCAAGGATGCAGCCTCAGCCTTCTCCACACCACGTTTCCCCACAGACAAGTTCCCCAGGACTCCAATTCCGACCTCCCG |
| 2:2204:7112:49728#2#R | \| \| |
| MQ=60 isize=-165 C=37 | aATCAAGTGCGCtcTCCCCAgCCTGTCCCTTCTCCACGGCCACAGTCCCAG\|C\|CCCCCcaCTCCAGTCCTTCCCCAAGGATGCAGCCTaAGCCTTCTcCacgCCACGTTTCCCCACAGGCTGGAGTGCAanGACTCCAATACAAGATGCCAG |
| 2:2212:12654:32202#2#R | \| \| |
| MQ=60 isize=-177 C=33 | AATCAAGTGCgCtCTCCCCAGCCTGTCCCTTCTCCACGGCCACAGTCCCAG\|C\|CCCCCCACTCCAGTCCTTCCCCAAGGATGCAGCCTCAGCCTTCTCCACACCACGTTTCCCCACAGACAAGTTCCCCAGGACTCCAATTAGGGACACTAC |
| 1:1110:10429:19059#2#F | \| \| |
| MQ=60 isize=208 C=33 | GGCTATACCGGTATTGGAGTCCTATCAAGTGCGCTCTCCCCAGCCTGTCCCTTCTCCACGGCCACAGTCCCAG\|C\|CCCCCCACTCCAGTCCTTCCCCAAGGATGCAGCCTCAGCCTTCTCCACACCACGTTTCCCCaCAGACAAGtTCCCCA |
| 1:2207:11726:13711#2#F | \| \| |
| MQ=60 isize=208 C=33 | GCTGGTCGGTGGATTGGAGTCntATCAAGTGCGCTCTCCCCAGCCTGTCCCTTCTCCACGGCCACAGTCCCAG\|C\|CCCCCCaCTCCAGtCCtTCCCCaAgGaTGCAGCCtCAGCCTTCtCCACACCaCGttTCCCCACAgACAAGTTCCCCc |
| 2:1202:11466:24041#2#F | \| \| |
| MQ=60 isize=208 C=33 | TAGGCGTCTGGGATTGGAGTCntATCAAGTGCGCTCTCCCCAGCCTGTCCCTTCTCCACGGCCACAGTCCCAG\|C\|CCCCCCACTCCAGTCCTTCCCCAAGGATGCAGCCTCAGCCTTCTCCACACCACGTTTCCCCaCAGACAAGTTCCCCc |
| 1:2202:14108:64497#2#R | \| \| |
| MQ=60 isize=-169 C=37 | ccccccacatcaagtgccctctcCCCAGCCTGTCCCTTCTCCACGGCCaCAGTCCCAG\|C\|CCCCCCACtCCAGTCCTTCCCCAAGGATGCAGCCTCAGCCTTCTCCACACCACGTTTCCCCACAGACAAanGACTCCAATCCGCCACCTGTC |
| 2:2116:10850:66391#2#R | \| \| |
| MQ=60 isize=-177 C=27 | atcAAGTgCgCTCtCCCCAGCCTGTCCCTTCtCCACGGCCaCAGTCCCAG\|C\|CCCCCCACTCCAGTCCTTCCCCAAGGATGCAGCCTCAGCCTTCTCCACACCACGTTTCCCCACAGACAAGTTCCCCAAgnACTCCAATACAAATCGTCTC |
| 2:2214:10468:4873#2#F | \| \| |
| MQ=60 isize=207 C=33 | GAGGGTGTTCGGATTGGAGTCCTCAAGTGCGCTCTCCCCAGCCTGTCCCTTCTCCACGGCCACAGTCCCAG\|C\|CCCCCCACTCCAGTCCTTCCCCAAGGATGCAGCCTCAGCCTTCTCCACACCACGTTTCCCCaCAGACAAGTTCCCCACa |
| 1:2208:2757:64975#2#R | \| \| |
| MQ=60 isize=-179 C=33 | tCAAGTGCGCTCTCCCCAGCCTGTCCCTTCTCCACGGCCACAGTCCCAG\|C\|CCCCCCACTCCAGTCCTTCCCCAAGGATGCAGCCTCAGCCTTCTCCACACCACGTTTCCCCACAGACAAGTTCCCCACanGACTCCAATTGTATGCAAGCT |
| 2:1103:7226:94252#2#F | \| \| |
| MQ=60 isize=205 C=33 | GCCGAGGGTAGCATTGGAGTCCTAAGTGCGCTCTCCCCAGCCTGTCCCTTCTCCACGGCCACAGTCCCAG\|C\|CCCCCCACTCCAGtCCTTCCCCAAGgaggCAgCCtCAgCCTTCTCCacccCcCgtTTCCCcccacacaggttccccacac |
| 1:1102:6432:85915#2#R | \| \| |
| MQ=60 isize=-181 C=33 | agtgcgctctCCCCAGCCTGTCCCtTCTCCACGGCCACAGTCCCAG\|C\|CCCCCCACtCCAGTCCAtCCCCAAGGATGCAGCCTCAGCCTTCTCCACaCCACGTTTCCCCACAGACAAGTTCCCCACATCanGACTCCAAtCCTGCCGTGTCG |
| 1:2202:16673:93147#2#R | \| \| |
| MQ=60 isize=-181 C=22 | agggcgctctcCCCAGCctgtcCCTTCTCCaCGGCCaCAGTCCCAG\|C\|CCCCCCACcCCAGTCCTTCCCCAAGGATGCAGCCTCAGCCTTCTCCACACCACGTTtCCCCACAGACAAGTTCCCCACATCanGACTCCAAtCCTGCCGTGTCG |
| 2:1201:18805:42574#2#R | \| \| |
| MQ=60 isize=-181 C=37 | AGtGCGCTCTCCCCAGCCTGTCCCTTCTCCACGGCCACAGTCCCAG\|C\|CCCCCCACTCCAGTCCTTCCCCAAGGATGCAGCCTCAGCCTTCTCCACACCACGTTTCCCCACAGACAAGTTCCCCACATCAGGACTCCAATCAAAAAGGGAAT |
| 1:1116:11539:77364#2#F | \| \| |
| MQ=60 isize=308 C=37 | CAAGGAAATAGCATTGGAGTCCTGTGCGCTCTCCCCAGCCTGTCCCTTCTCCACGGCCACAGTCCCAG\|C\|CCCCCCACTCCAGTCCTTCCCCAAGGATGCAGCCTCAGCCTCCTCCACACCACGTTTCCCCGCAGACAAGTTCCCCACATCC |
| 1:2207:1402:36373#2#F | \| \| |
| MQ=60 isize=203 C=22 | CGCGGGGTGCCGATTGGAGTCntgTGcGCTCTCCCCAtccTGTCCCTTCTcCACGGccacAgtcccAg\|C\|CCCCCCACTCCAGTcCTtCcCCaaggaTGCAGCCTCAgcctTCTccacAccaCggttccccacagacaatgtccccacatcc |
| 1:2209:3800:71936#2#F | \| \| |
| MQ=60 isize=203 C=37 | AAGATTATAGTTATTGGAGTCntGTGCGCTCTCCCCAGCCTGTCCCTTCTCCACGGCCACAGTCCCAG\|C\|CCCCCCACTCCAGTCCTTCCCCAAGGATGCAGCCTCAGCCTTCTCCACACCACGtTTCCCCACAGACAAGTTCCCCACATCC |
| 2:1109:15642:63283#2#F | \| \| |
| MQ=60 isize=203 C=37 | AACCTGTCGTGCATTGGAGTCntGTGCGCTCTCCCCAGCCTGTCCCTTCTCCACGGCCACAGTCCCAG\|C\|CCCCCCACTCCAGTCCTTCCCCaAGGatGCAGcCTCAGCCTTCTCCACACCACGtTTCCCCACAGaCaAGTTCCCCACATCC |
| 2:1214:10345:86232#2#F | \| \| |
| MQ=60 isize=203 C=37 | TACGGTAACAACATTGGAGTCCTGTGCGCTCTCCCCAGCCTGTCCCTTCTCCACGGCCACAGTCCCAG\|C\|CCCCCCACTCCAGTCCTTCCCCAAGGATGCAGCCTCAGCCTTCTCCACACCACGTTTCCCCACAGACAAGTTCCCCACATCC |
| 2:2208:11447:84347#2#F | \| \| |
| MQ=60 isize=203 C=37 | TCCGGTTCTAATATTGGAGTCntGTGCGCTCTCCCCAGCCTGTCCCTTCTCCACGGCCACAGTCCCAG\|C\|CCCCCCACTCCAGTCCTTCCCCAAGGATGCAGCCTCAGCCTTCTCCACACCACGTTTCCCCACAGACAAGTTCCCCACATCC |
| 1:1109:9510:14469#2#F | \| \| |
| MQ=60 isize=202 C=33 | GGGCCCGAGCGGATTGGAGTCCTGCGCTCTCCCCAGCCTGTCCCTTCTCCACGGCCACAGTCCCAG\|C\|CCCCCCACTCCAGTCCTTCCCCAAGGATGCGGcCTCAGCCTTCTCCACACCACGTTTCCCCACAGACAAGTTCCCCACaTCCTg |
| 1:2203:11796:23714#2#F | \| \| |
| MQ=60 isize=202 C=33 | GTGTGCAGGATTATTGGAGTCCTGCGCTCTCCCCAGCCTGTCCCTTCTCCACGGCCACAGTCCCAG\|C\|CCCCCCACTCCAGtCCTTCCCCAaGGATGCagCCtcagCctTCTCCaCaCCaCgttTCCCCACagaCAagttCCCCacatcctg |
| 1:2213:13101:12800#2#F | \| \| |
| MQ=60 isize=202 C=37 | CGCTGGGGGTGGATTGGAGTncTGCGCTCTCCCCAGCCTGTCCCTTCTCCACGGCCACAGTCCCAG\|C\|CCCCCCACTCCAGTCCTTCCCCAAGGATGCAGCCTCAGCCTTCTCCACACCACGTTTCCCCACAGACAAGTTCCCCACATCCTG |
| 2:1206:8664:22963#2#F | \| \| |
| MQ=60 isize=202 C=37 | CCGAGACTGTCGATTGGAGTCntGCGCTCTCCCCAGCCTGTCCCTTCTCCACGGCCACAGTCCCAG\|C\|CCCCCCACTCCAGTCCTTCCCCAAGGATGCAGCCTCAGCCTTCtCCACACCACGTTTCCCCACagACAAGTTCCCCACATCCtg |
| 2:2109:9785:66475#2#F | \| \| |
| MQ=60 isize=307 C=37 | GAAATTCTTGGTATTGGAGTCCTGCGCTCTCCCCAGCCTGTCCCTTCTCCACGGCCACAGTCCCAG\|C\|CCCCCCACTCCAGtCCTTCCCCAcgGATGCAGCCTCAGCCTTCtCCACaCCaCGtTTCCCCACagaCAagTtccccccatcctg |
| 1:2101:10190:65948#2#R | \| \| |
| MQ=60 isize=-183 C=33 | ggcgctctcCCCAGCCTGTCCCtTCTCCACGGCCACAGtCCCAG\|C\|CCCCcCACTCCAGtCCttCCCCAAGGATGCAGCCTCAGCCTTCTCCACaCCACGTTTCCCCACAGACAAGTTCCCCACATCCTAGGACTCCAATATCGCTTACCCC |
| 1:2109:14261:15214#2#R | \| \| |
| MQ=60 isize=-183 C=33 | ggcgctctCCCCAGCCTGtCCCTtCTCCACGGCCACAGTCCCAG\|C\|CCCCCCaCTCCAGTCCTTCCCCAAGGATGCAGCCTCAGCCTTCTCCACACCACGTTTCCCCACAGACAAGTTCCCCACATCCTanGACTCCAATCTCCAACGGACC |
| 1:2109:2042:15172#2#R | \| \| |
| MQ=60 isize=-185 C=37 | gcgctctCCCCAGCCTGTCCCTTCTCCACGGCCACAGTCCCAG\|C\|CCCCCCACTCCAgTCCTTCCCCAAGGATGCAGCCTCAGCCTTCTCCACACCACGTTTCCCCACAGACAAGTTCCCCACATCCTGgnGACTCCAATAGGTTCAACCAG |
| 2:1205:8322:37182#2#R | \| \| |
| MQ=60 isize=-184 C=37 | gCGCTCtCCCCAGCCtgTCCCTtCTCCaCGGCCACAGTCCCAG\|C\|CCCCCCACTCCAGTCCTTCCCCAAGGATGCAGCCTCAGCCTTCTCCACACCACGTTTCCCCACAGACAAGTTCCCCACATCCTGanGACTCCAATGGGGTCTTCTCG |
| 2:2105:7495:20568#2#R | \| \| |
| MQ=60 isize=-164 C=27 | caaatccctctcccactcaaggcgctccccccagcctgtcccctacccacggccacaggCCCAG\|C\|CCCccCaCTCCAgTCCttCCCCaAGGAtgcagCCTCAGCCTTCTCCACACCACGTTTCCCCAcanGACTCCAATTCATTACGAGTA |
| 2:2205:11700:52921#2#R | \| \| |
| MQ=60 isize=-184 C=37 | gcGCTCTCCCCAGCCTGTCCCTTCTCCACGGCCACAGTCCCAG\|C\|CCCCCCACTCCAGTCCTTCCCCAAGGATGCAGCCTCAGCCTTCTCCACACCACGTTTCCCCACAGACAAGTTCCCCACATCCTGanGACTCCAATACGACTCGGTAC |
| 1:1211:15108:34926#2#R | \| \| |
| MQ=60 isize=-186 C=33 | CGCTCtCCcCaGCCTGTCCCTTCTCCACGGCCaCAGTCCCAG\|C\|CCCCCCACTCCAGTCCTTCCCCAAGGATGCAGCCTCAGCCTTCTCCACACCACGTTTCCCCACAGACAAGTTCCCCACATCCTGGAGGACTCCAATCAACATTTAAAC |
| 1:2108:14011:100407#2#R | \| \| |
| MQ=60 isize=-186 C=37 | CGCTCTCCCCAGCCTGTCCCTtCtCCaCGGCCACagTCCCAG\|C\|CCCCCCACTCCAGTCCTTCCCCAAGGATGCAGCCTCAGCCTTCTCCACACCaCGTtTCCCCACAGACAAGTTCCCCACATCCTGGanGACTCCAAtCCCCGCGCCCCC |
| 1:2214:3510:98660#2#R | \| \| |
| MQ=60 isize=-186 C=37 | CGCTCTCCCCAGCCTGTCCCTtCTCCaCGGCCACAGTCCCAG\|C\|CCCCCCACTCCAGTCCTTCCCCAAGGATGCAGCCTCAGCCTTCTCCACACCACGTTTCCCCACAGACAAGTTCCCCACATCCTGGAgnACTCCAATCTAGTTTATACA |
| 2:1115:19492:84523#2#R | \| \| |
| MQ=60 isize=-186 C=2 | cgctctccccagcctgtcccttctccacggccacagtcccag\|c\|ccccccactccagtccttccccaaGGATGCAGCCTCAGCCTTCTCCaCACCaCGTTTCCCcACAGACAAGtTCCCCACATCCTGgagnACTCCAagCCCCGCGCCCCC |
| 2:1204:13180:87822#2#R | \| \| |
| MQ=60 isize=-186 C=33 | cgctCTCCCCAGCCTgTCCCTTCTCCACGGCCACAGTCCCAG\|C\|CCCCCCACTCCAGTCCTTCCCCAAGGATGCAGCCTCAGCCTTCTCCACACCACGTTTCCCCACAGACAAGTTCCCCACATCCTGGanGACTCCAATCGGCGGCAAGCC |
| 2:2204:8608:14279#2#R | \| \| |
| MQ=60 isize=-186 C=37 | cgctctCCCCAGCCTGTCCCTTCTCCACGGCCACAGTCCCAG\|C\|CCCCCCACTCCAGTCCTTCCCCAAGGATGCAGCCTCAGCCTTCTACACACCACGTTTCCCCACAGACAAGTTCCCCACATCCTGGanGACTCCAATAATGAAGGTTGA |
| 2:2216:14527:74393#2#R | \| \| |
| MQ=60 isize=-186 C=22 | cgctctccCCAGCCtGTcCCTTCTCCaCGgccACAGTCCCaG\|C\|CCCCCCACTCCAGTCCTTCCCCAAGGATGCagCCTCAGCCTTCTCCACACcaCGTTTCCCCACAGACAAGTTCCCCACATCCTGGAGGACTCCAAtTGAGCTCACCTC |
| 2:1105:2885:13436#2#F | \| \| |
| MQ=60 isize=199 C=33 | AAGCGACTGCTGATTGGAGTCntGCTCTCCCCAGCCTGTCCCTTCTCCACGGCCACAGTCCCAG\|C\|CCCCCCACcCCAGtCCTTCCCCAaGgatccagccccaccctcctccaccccccgttcccccccacccaagtcccccccaccccgtc |
| 2:2116:4204:90859#2#F | \| \| |
| MQ=60 isize=199 C=33 | AAGCGACTGCTGATTGGAGTncTGCTCTCCCCAGCCTGTCCCTTCTCCACGGCCACAGTCCCAG\|C\|CCCCCCACTCCaGTCCTTCCCCAaGGAtgcagCCtCAGCCTTCTCCACaCCacGTTTCCCCacagacAAGTTCCCcccatcctggc |
| 1:1115:2225:95075#2#F | \| \| |
| MQ=60 isize=303 C=37 | GGAGGGGTGTTAATTGGAGTCCTCTCTCCCCAGCCTGTCCCTTCTCCACGGCCACAGTCCCAG\|C\|CCCCCCACTCCAGTCCTTCCCCAAGGATGCAGCCTCAGCCTTCTCCaCACCACGTTTCCCCACAGACAAGTTCCCCACATCCTGGAC |
| 1:2107:6346:31993#2#F | \| \| |
| MQ=60 isize=198 C=37 | TTAATCTAAAAAATTGGAGTCCTCTcTCCCCAGCCTGTCCCTTCTCCACGGCCACAGTCCCAG\|C\|CCCCCCACTCCAGTCCTTCCCCAAGGaTGCAGCCTCAgCGTTCTCCACACCACGTTTCCCCACAGaCaAGCTCCCCACATCCTGGaC |
| 2:1101:5532:85154#2#F | \| \| |
| MQ=60 isize=198 C=33 | GTTACACGGGCTATTGGAGTCCTCTCTCCCCAGCCTGTCCCTTCTCCACGGCCACaGTCCCAG\|C\|CCCCCcactccAGtCCTTCCCCaaggaggcagcccaggccttctccaccccacgttcccccacagccaagttccccaatcccgggcc |
| 2:2110:7367:17294#2#F | \| \| |
| MQ=60 isize=198 C=33 | TAaCTGGTCCGGATTGGAGTCCTCTCCCCAGCcTGTCCCTTCTCCACGGCCACAGTCCCAG\|C\|CCCCccactccagtccttccccaaggatgcagcctcagcctcccccacaccacgtttccccacagacaagttcctcaaaccccggacgg |
| 1:1110:18858:34181#2#R | \| \| |
| MQ=60 isize=-187 C=37 | CTCTCCCCAGCCTGTCCCTTCTCCACGGCCACAGTCCCAG\|C\|CCCCCCaCTCCAGTCCTTCCCCAAGGATGCAGCCTCAGCCTTCTCCACACCACGTTTCCCCACAGACAAGTTCCCCACATCCTGGACAGGACTCCAAtCGCCTCAGCCAG |
| 1:1111:14675:93972#2#R | \| \| |
| MQ=60 isize=-187 C=33 | ctctcCCCAGCCTGTCCCTtCtCCaCGGCCACAGTCCCaG\|C\|CCCCCCaCTCCAGTCCTTCCCCAAGGATGCAGCCTCAGCCTTCTCCaCaCCACGTTTCCCCaCAGACAAGTTCCCCACATCCTGGACAGGACTCCAATGGACCTCCACTC |
| 1:1214:2931:58360#2#R | \| \| |
| MQ=60 isize=-187 C=2 | ctctccccagcatgccccttctccacggccacagcccccg\|c\|cccccccctccagtccttccccaaggatgcagcctcagccttctccacaccacGTTTCCCCACAGATAAGTtCCCCACATCCTGGacagnACTCaAAtGGCCGGCACCGC |
| 1:2109:15750:80470#2#R | \| \| |
| MQ=60 isize=-187 C=37 | CTCTCCCCAGCCTGTCCCTTCTCCACGGCCaCAGTCCCAG\|C\|CCCCCCACTCCAGTCCTTCCCCAAGGATGCAGCCTCAGCCTTCTCCACACCACGTTTCCCCACAGACAAGTTCCCCACATCCTGGACanGACTCCAATCCAAAGGTCACC |
| 1:2116:15925:91247#2#R | \| \| |
| MQ=60 isize=-187 C=33 | ctctCCCCAGCCTGTCCCttCTCCaCGGCCACAGTCCCAG\|C\|CCCCCCaCTCCAGTCCTTCCCCAAGGATGCAGCCTCAGCCTTCTCCACACCACGTTTCCCCACAGACAAGTTCCCCACATCCTGGACanGACTCCAATGAAGCATGCAGA |
| 1:2210:13356:41372#2#R | \| \| |
| MQ=60 isize=-187 C=33 | ctctCCCCAGCCTGTCCCtTCtCCACGGCcACAGTCCCAG\|C\|CCCCCCaCTCCAGTCCtTCCCCAAGGATGCAGCCTCAGCCTTCTCCACACCACGTTTCCCCACAGACAAGTTCCCCACATCCTGGACAGGACTCCAATGAGACTTAAAGC |
| 2:2207:4209:6170#2#R | \| \| |
| MQ=60 isize=-187 C=15 | ccCTCCCCAGCctGTCccttCTCCACGGCCACAGTCCCag\|c\|CCCcCCACTCCagTCCTtCCCCAAGGATGCAGCCTCAGCCTTCTCCACACCACGTTTCCCCACAGACAAGTTCCCCACATCCTGGACanGACTCCAATACGCCGAACTGC |
| 1:1106:15200:76600#2#F | \| \| |
| MQ=60 isize=197 C=33 | CATTTTTTGTGAATTGGAGTCntTTCTCCCCAGCCTGTCCCTTCTCCACGGCCACAGTCCCAG\|C\|CCCCCCACTCCAGTCCTTCCCCAAGGATGCAGCCTCAGCCTTCTCCACACCACGTTTCCCCACAGACAAGTTCCCCACATCCTGGAC |
| 1:2105:21081:68888#2#F | \| \| |
| MQ=60 isize=197 C=37 | ATCGGCATATATATTGGAGTCntCTCCCCAGCCTGTCCCTTCTCCACGGCCACAGTCCCAG\|C\|CCCCCCACTCCAGTCCTTCCCCAAGGATGCAGCCTCAGCCTTCTCCACACCACGTTTCCCCACAGACAAGTTCCCCaCATCCTGGaCTG |
| 2:1204:5985:14037#2#F | \| \| |
| MQ=60 isize=197 C=37 | TAACTGGTCCGGATTGGAGTCntCTCCCCAGCCTGTCCCTTCTCCACGGCCACAGTCCCAG\|C\|CCCCCCACTCCAGTCCTTCCCCAAgGATGCAGCCTCAGCCTTCTCCACACCACGTTTCCCCACAGaCAAGTTCCCCACATCCtGGACTG |
| 1:2104:7285:72809#2#R | \| \| |
| MQ=60 isize=-188 C=37 | cctcccCAGCCtGtCCCTTCtCCACGGCCACAGTCCCaG\|C\|CCCCCCaCTCCAGTCCTTCCCCAAGGATGCAGCCTCAGCCTTCTCCACACCACGTTTCCCCACAGACAAGTTCCCCACATCCTGGACTanGACTCCAATAGTCTCCTACGA |
| 1:2210:16169:46914#2#F | \| \| |
| MQ=60 isize=196 C=33 | ACGTGTGACGGGATTGGAGTCCTCCCCAGCCTGTCCCTTCTCCACGGCCACAGTCCCAG\|C\|CCCCCCACTCCAGTCCTTCCCCAAGGATGCagCCTCAgCCTTCTCCACACCACGttTCCCCACAgACAAgttcCCcacatcctggactggt |
| 2:1212:8065:96528#2#F | \| \| |
| MQ=60 isize=196 C=37 | TGTGAGGTAGGGATTGGAGTCCTCCCCAGCCTGTCCCTTCTCCACGGCCACAGTCCCAG\|C\|CCCCCCACTCCAGTCCTTCCCCAAGGATGCAGCCTCAGCCTTCTCCACACCACGTTTCCCCACAGAcAAGtTCCCCACATCCTGGACTGGt |
| 1:2109:9717:23748#2#R | \| \| |
| MQ=60 isize=-164 C=2 | tacttccctctccaatccagccccccctccccagcctgtcccgtctccacgcccactgtcccag\|c\|cccccccctccagtccttccccaaggatgcagcctcagccttctccccaacacggttccccacangactccaattaccctcacctc |
| 2:2109:2379:86524#2#R | \| \| |
| MQ=60 isize=-125 C=2 | ttgcacccgtccccacccccaacaggccaggagatccctaaatacccccccaaacaagttccccctccccagcctgtcccttctccacggccacagtcccaa\|c\|CCCCCCaCtCCAGTCCTTCCCCAAGAGGACTCCAATCCaCCCgACCCA |
| 1:1208:11313:46332#2#F | \| \| |
| MQ=60 isize=195 C=33 | GCACGGATGGCCATTGGAGTCntcCCCAGCCTGTCCCTTCTCCACGGCCACAGTCCCAG\|C\|CCCCCCACTCCAGTCCTTCCCCAAGGATGCAGCCTCAGCCTTCTCCACACCACGtttcCCCccaGACAAGTTCCCCaCATCCtggactgga |
| 2:2206:5440:99033#2#F | \| \| |
| MQ=60 isize=300 C=22 | GCAGACTGACCTATTGGAGTCntcCCCAGCCTGTCCCTTCTCCACGGCCACAGTCCCAg\|C\|CCCCCCcCtCCagTCCTtcCCCAAgGatgCAGCctcagccttctccacaccacgtttccccacagccaagttccccacctcctggactgga |
| 1:1203:10624:32133#2#F | \| \| |
| MQ=60 isize=194 C=22 | CGTCGCGTACATATTGGAGTCntCTCCCCCCAGCCTGTCCCTTCTCCACGGCCACAGTCCCAg\|C\|CCCCCCACtCCagtcCttCCCCAagGaTGCagCCtCAccCtTCTCCACACCACGtTtCCCCACAgacaagttcCCCacatcctggcc |
| 1:1201:15824:63826#2#R | \| \| |
| MQ=60 isize=-158 C=2 | cccaatttcgccttccaatgaagtcggctcaccccagcctgtcccctctccacggccccagtcccag\|c\|ccccccactccagtccttccccaAGGatgCAGCCTcAgCcTTCtCCaCaCcAcGttTctcanGACTCCAATaAgGGAtGtaTC |
| 1:2102:1332:10707#2#R | \| \| |
| MQ=60 isize=-193 C=37 | CCCCAgCCTGtCCCTTCTCCACGGCCACAGTCCCaG\|C\|CCCCCCACTCCAGTCCttCCCCAAGGATGCAGCCTCAGCCTTCTCCACACCACGTTTCCCCACAGACAAGTTCCCCaCATCCTGGACTGGTAGGACTCCAATGTGTTCGGTGAC |
| 2:1101:7051:7123#2#R | \| \| |
| MQ=60 isize=-162 C=33 | ccttactccccctccaaaccagtgcgccccccccagcccgtcccttctccccggccccagtcCCAG\|C\|CCCCCcaccCCAGTCCTTCCCCAAGGATGCAGCCTCAGCCTTCTCCACACCaCGTTTCCCCAGGACTCCAATCGTTCTAAGTGG |
| 2:2103:2029:28035#2#R | \| \| |
| MQ=60 isize=-133 C=22 | ccgcccccccccccaagggccagagatccccaattccccctccaaacaagtgccccccccccagcCTGTCCCttccCCcCGGccaCAGtCCCAG\|C\|CCCCCCaCtCCAGTCCTTCCCCAAGGATGCAGCAGGACTCCAATGTCGGCCCGACC |
| 2:2115:5888:71005#2#R | \| \| |
| MQ=60 isize=-116 C=37 | ttttgccttttggccccgtccccccaccctcaaagccaggcaaccccttattccccccccaatcaaaggcccccccCCCAGCCTGTCCCTTCTCCACGGCCACAGTCCCAG\|C\|CCCCCCACTCCAGTCCAgnACTCCAATCGGGCCTACAGA |
| 1:1210:19351:28266#2#F | \| \| |
| MQ=60 isize=193 C=33 | GGGAGTGGGGGGatTGGAgTCCTCCCAGCCTGTCCCTTCTCCACGGCCACAGTCCCAG\|C\|CCCCCCACTCCAGTCCTTCCCCAAGGATGCAGCCTCAGCCTTCTCCACACCACGTTTCCCCACAGACaAGTTCCCCACATCCTGGACTGgta |
| 1:1216:15328:3973#2#F | \| \| |
| MQ=60 isize=193 C=37 | GGGAGTGGGGGGATTGGAGTncTCCCAGCCTGTCCCTTCTCCACGGCCACAGTCCCAG\|C\|CCCCCCACTCCAGtCCTTCCCCAAGGATGCAGCCTCAGCCTTCTCCACACCACGTTtCCCCACAGACGAgTTCCCCaCaTCCTGGaCTGGta |
| 2:1101:21092:59924#2#F | \| \| |
| MQ=60 isize=180 C=37 | CCCTCCGCCAAGATTGGAGTCCTCCCAGCCTGTCCCTTCTCCACGGCCACAGTCCCAG\|C\|CCCCCCACTCCAGTCCTTCccCaAGGAtgCagccTCAgcctTCTCCACACCACGttTCccCACAGacaagtttcccacatcctggactggta |
| 2:1206:19118:84073#2#F | \| \| |
| MQ=60 isize=193 C=2 | GGATTATGGCGGaTTggaGTCntcCCAGCCTGTCCCTTCTccacggccacagtcccag\|c\|ccccccactccagtccttccccaaggatgcagcctcagccttctccccaccacgtttccccccagacaagttccccccatcctggactggtc |
| 2:2212:10492:72214#2#F | \| \| |
| MQ=60 isize=193 C=40 | CATCATATCGGTATTGGAGTCCTCCCAGCCTGTCCCTTCTCCACGGCCACAGTCCCAG\|C\|CCCCCCACTCCAGTCCTTCCCCAAGGATGCAGCCTCAGCCTTCTCCAtACCACGTTTCCCCACAGACAAGTTCCCCACATCCTGGACTGGtA |
| 1:1207:19170:36349#2#R | \| \| |
| MQ=60 isize=-192 C=33 | CCCAGCCTGTCCCTTCTCCaCGGCCacAGTCCCAG\|C\|CCCCCCACTCCAGTCCTTCCCCAAGGATGCAGCCTCAGCCTTCTCCACACCACGTTtCCCCACAGACAAGTTCCCCACATCCTGGACTGGTAanGACTCCAATCCAATCTCCCCA |
| 2:1116:2945:96438#2#R | \| \| |
| MQ=60 isize=-192 C=2 | ccccgcctgtcccttccccccggccaccgtccccg\|c\|cccccccctccagtCCcCCCCCaAGgatgCaGCCtcaGCCttCtCCACACCaCGTTTCCCCACAGaCAAGTTCCCCaCATCCTGGACTGGTAAGGACTCCAATCGCACCCCTGCC |
| 2:2108:8717:20178#2#R | \| \| |
| MQ=60 isize=-192 C=33 | ccccgcctgtcccttctccacggccacagtccccg\|C\|CCCCCCACTCCAGTCCTTCCCCAAGGATGCAGCCTCAGCCTTCtCCACaCCACGTTTCCCCACAGACAAGTTCCCCACATCCTGGACTGGTAAGGACTCCAATGCCGGCCCCaCC |
| 1:1111:6883:14606#2#F | \| \| |
| MQ=60 isize=297 C=37 | CAAGAGTTATTGATTGGAGTCCTCCAGCCTGTCCCTTCTCCACGGCCACAGTCCCAG\|C\|CCTCCCACTCCAGTCCTTCCCCAAGGATGCAGCCTCAGCCTTCTCCACACCACGTTTCCCCACAGACAAGTTCCCCaCATCCTgGACTGGtag |
| 1:1213:9035:84295#2#F | \| \| |
| MQ=60 isize=192 C=22 | CGGCGGACCGGAATTGGAGTncTCCAGCCTGTCCCTTCTCCACGGCCACAGTCCCAg\|C\|CCCCCCACTCCAGTCCTTCCCCAAGGATGCAGCCTCAGCCTTCTCCACACCACGTTTCCCCACAgACAAGTTCCCCACATCCTGGACTGGtag |
| 1:2211:6015:41021#2#F | \| \| |
| MQ=60 isize=326 C=37 | ATCCAATTGTCCATTGGAGTCCTCCAGCCTGTCCCTTCTCCACGGCCACAGTCCCAG\|C\|CCCCCCACTCCAGTCCTTCCCCAAGGATGCAGCCTCAGCCTTCTCCACACCACGtTTCCCCACAGACAAGtTCCCCACATCCTGGACtggtag |
| 2:1110:16198:82071#2#F | \|C\| |
| MQ=60 isize=192 C=33 | GAGCGGGTGGAGATTGGAGTCCTCCAGCCTGTCCCTTCTCCACGGCCACAGTCCCAI\|C\|CCCCCCACTCCAGTCCTTCCCCAAGGATGCAGCCTCaGCCTTCTCCACACCACGTTTCCCCAcagaCAAgttccccacatcctggactggtg |
| 2:1116:12936:54959#2#F | \| \| |
| MQ=60 isize=297 C=37 | CAAGAGTTATTGATTGGAGTCCTCCAGCCTGTCCCTTCTCCACGGCCACAGTCCCAG\|C\|CCTCCCACTCCAGTCCTTCCCCAAGGATGCAGCCTCAGCCTTCTCCACACCACGTTTCCCCACAGACAAGTTCCCCACATCCTGGACTGGTAG |
| 2:1208:7538:75772#2#F | \| \| |
| MQ=60 isize=192 C=2 | gGggCCcGaACCATTGGAGTCntccagcctgtcccttctccacggccacagtcccag\|c\|ccccccactccagtccttccccaaggatgcagcctcatccttctccacaccacgtttccccacagacaagttccccacagcctggactggtaa |
| 2:2107:1801:49996#2#F | \| \| |
| MQ=60 isize=192 C=33 | CAGACGAACCTAATTGGAGTCCTCCAGCCTGTCCCTTCTCCACGGCCACAGtCCCAG\|C\|CCCCCCACtCCAGTCCTtcccccaggaggcagccccagcctttcccaccccccgttccccccccaccaagtccccacctcccggcccgggacc |
| 2:2211:9601:11084#2#F | \| \| |
| MQ=60 isize=192 C=37 | CGTTGACGATAGATTGGAGTCCTCCAGCCTGTCCCTTCTCCACGGCCACAGTCCCAG\|C\|CCCCCCACTCCAGTCCTTCCCCAAGGATGCAGCCTCAGCCTTCTCCACACCACGTTTCCCCACagACAAGTTCCCCACATCCTGGACTGgtag |
| 2:2105:13526:57575#2#R | \| \| |
| MQ=60 isize=-116 C=27 | CCaGcCTGTCCCTTCTCCACGgccaCAGtCCCAG\|C\|CCCCCcacTcCAgtcC |
| 1:2107:15588:14032#2#R | \| \| |
| MQ=60 isize=-194 C=37 | CAGCCTGTCCCTTCTCCaCGGCCACAGTCCCAG\|C\|CCCCCCACTCCAGTCcTTCCCCAAGgATGCAGCCTCAGCCTTCTCCACACCACGTTTCCCCACAGACAAGTTCCCCACATCCTGGACTGGTAGCAGGACTCCAAtCGCCTACACAAT |
| 2:2203:5660:54104#2#R | \| \| |
| MQ=60 isize=-194 C=37 | cagcctgtcccttctccacgGCCACaGTCCCaG\|C\|CCCCCCACtCCAGTCCTTCCCCAAGGATGCaGCCtCAGCCTTCTCCACACCACGTTTCCCCACAGACAAGTTCCCCACATCCTGGACTGGTAGCanGACTCCAATCGCCTACACAAT |
| 2:2216:5353:42300#2#R | \| \| |
| MQ=60 isize=-194 C=33 | CAGCCTGtCCCtTCtCCACGGCCACAGTCCCaG\|C\|CCCCCCACTCCAGTCCTTCCCCAAGGATGCAGCCTCAGCCTTCTCCACACCACGTTTCCCCACAGACAAGTTCCCCACATCCTGGACTGGTAGCAGGACTCCAaTCGCCTACACAAT |
| 1:2103:2651:75585#2#F | \| \| |
| MQ=60 isize=190 C=40 | GTCCTATTCAGGATTGGAGTCCTAGCCTGTCCCTTCTCCACGGCCACAGTCCCAG\|C\|CCCCCCACTCCAGTCCTTCCCCAAGGATGCAGCCTCAGCCTTCTCCACACCACGTTTCCCCACAGACAAGTTCCCCACATCCTGGACTGGTAGCT |
| 2:1215:10752:80672#2#F | \| \| |
| MQ=60 isize=190 C=37 | CGAagGGGAcCaATTGGAGTncTAGCCTGTCCcTTCTCCACGGCCACAGtcCCAG\|C\|CCCCCCACTCCAgtccttCCCCAAGgATGCAGacacAGCcttctCCACACCACGTttcCCCaCAGACAAGTTCCccacatcctggactggtagcc |
| 2:1210:13344:9946#2#R | \| \| |
| MQ=60 isize=-195 C=33 | agCCTGTCCCTTCTCCACGGCCACAGTCCCAG\|C\|CCCCCCACTCCAGTCCTTCCCCAAGGATGCAGCCTCAGCCTTCTCCACACCACGTTTCCCCACAGACAAGTTCCCCACATCCTGGACTGGTAGCTAGGACTCCAATGCTCCAGCACAA |
| 1:1214:9151:95345#2#F | \| \| |
| MQ=60 isize=189 C=27 | CCCGCGTGTCTAATTGGAGTnctgccTGTCCCTTCTCCACGGcCACaGtCCCAG\|C\|CCCCCCACTCCagtCCTTCCCCaaggaTgCagCCTCAGCCTtCtcCACaCCACgcttCCCCaCagAcAagctCCCCAcatcctggactggtagctc |
| 2:1202:6372:50963#1#F | \| \| |
| MQ=60 isize=76 C=37 | GAAAGCATTGAATTCTGGTCCGGGCTGGCAAAATGCCCTTGTTCCATGGGGTTGGCCTGTCCCTTCTCCACGGCCACAGTCCCAG\|C\|CCCCCCACTCCAGTCCTTCCCCAAGGATGCAGCCTCAGCCTTCTAGGACTCCAATGTCGCAGTAG |
| 1:2111:18590:83432#2#R | \| \| |
| MQ=60 isize=-196 C=33 | GCCTGtCCCtTCTCCaCGGCCaCAGTCCCaG\|C\|CCCCCCACTCCAGTCCTTCCCCAAGGATGCAGCCTCAGCCTTCTCCACACCACGTTTCCCCACAGACAAGTTCCCCACATCCTGGACTGGTAGCTGAGGACTCCAATGTGATGTGAACC |
| 2:1202:6372:50963#2#R | \| \| |
| MQ=60 isize=-76 C=37 | AAGCATTGAATtCTGGTCcGGGCTGGCAAAATGCCCTTGTTCCATGGGGTTGGCCTGTCCCTTCTCCACGGCCACAGTCCCAG\|C\|CCCCCCACTCCAGTCCTTCCCCAAGGATGCAGCCTCAGCCTTCTanGACTCCAATGTCGCAGTAGCT |
| 2:2105:21003:11398#2#R | \| \| |
| MQ=60 isize=-196 C=37 | cCCTgtCCCttctCcaCGgccACAgtCCCAG\|C\|CCCCCCACTCCAGTCCttCCCCAAGGATGCAGCCTCAGCCTTCTCCACACCACGTTTCCCCACAGACAAGTtCCCCACATCCTGGACTGGTAGCTGanGACTCCAATTGTTTTCACTCG |
| 2:1111:11948:90305#2#F | \| \| |
| MQ=60 isize=188 C=37 | TATCAGCGCGCTATTGGAGTCCTGTCCCTTCTCCACGGCCACAGTCCCAG\|C\|CCCCCCACTCCAGTCCTTCCCCAAGGATGCAGcCTCAGCCTTCTCCACACCACGTTTCCCCACaGACAAGTTCCCCACATCCTGGACtgGTAGCTGCCCa |
| 2:2103:14483:28641#2#F | \| \| |
| MQ=60 isize=322 C=33 | GAAGGCGCGGGGATTGGaGTCCTGTCCCTTCTCCACGGCCACAgTCCCAG\|C\|CCCCCCACTCCAGTCCTTCCCCAAGGATGCAGCCTCAGCCTTCTCCACACCaCgtTTCCCCaCagacAAGTTCCCCccatcccggactggtagctgcccc |
| 1:1104:11752:100886#2#R | \| \| |
| MQ=60 isize=-197 C=33 | cctGTCCCTtCtCCACGGCCaCAGTCCCaG\|C\|CCCCCCaCTCCAGTCCTTCCCCAAGGATGCAGCCTCAGCCTTCTCCACaCCACGTTTCCCCACAGACAAGTTCCCCACATCCTGGACTGGTAGCTGCanGACTCCAATTGTTCAGGCtCG |
| 2:1212:10750:92815#2#R | \| \| |
| MQ=60 isize=-197 C=33 | CCTGTCcCtTCTCCACGGCCACAGtCCCAG\|C\|CCCCCCaCTCCAGTCCTTCCCCAAGGATGCAGCCTCAGCCTTCTCCACACCACGTTTCCCCACAGACAAGTTCCCCACATCCTGGACTGGTAGCTGCAGGACTCCAATAGCAAACCAGTC |
| 1:1105:13575:59867#2#F | \| \| |
| MQ=60 isize=187 C=37 | GGCAACGCGAAGATTGGAGTCntCTGTCCCTTCTCCACGGCCACAGTCCCAG\|C\|CCCCCCACTCCAGTCCTTCCCCAAGGATGCAGCCTCAGCCTTCTCCACACCACGTTTCCCCACAGACAAGTTCCCCACATCCTGGACTGGTAGCTGCC |
| 2:1215:5621:62106#2#F | \| \| |
| MQ=60 isize=423 C=27 | AGCCGGCCCGTGATTGGAGTnctGTCCCTTCTCCACGGCCACAGTCCCAG\|C\|CCCCCCACtCCAGtCCTTCCCCAAGGATGCAGCcTCAGCCTTCTCCACACCaCGttTCCCCACAGACAAgtTCCCCaCatCCTggacTGgTacctGCCCa |
| 1:1206:20847:43744#2#R | \| \| |
| MQ=60 isize=-198 C=37 | CtGTCCCTTCTCCACGgCCACaGtCCCAG\|C\|CcCCCCaCTCCAGTCCTtCCCCAAGGAtGCAGCCTCAGCCTTCTCCACACCACGttTCCCCACAGACAAGTTCCCCaCATCCTGGACTGGTAGCTGCCAGGACTCCAATCGAGGTCCCCCA |
| 2:1207:5207:60264#2#R | \| \| |
| MQ=60 isize=-198 C=37 | CTGtCCCTTCTCCACGGCCACAGTCCCAG\|C\|CCCCCccCTCCAGTCCTTCCCCAAGGATGCAGCCTCAGCCTTCTCCACaCCACGTTTCCCCACAGACAAGTTCCCCACATCCTGGACTGGTAGCTGCCanGACTCCAATACAACGCCAGCA |
| 2:2109:14575:79130#2#R | \| \| |
| MQ=60 isize=-198 C=33 | CTGTCCcTTCTCcACGGCcAcAGtcCCAG\|C\|CCCcccaCTCCAGTCCTTCCCCAAGGATGCAGCCTCAGCCTTCTCCACACCACGTTTCCCCACAGACAAGTTCCCCACATCCTGGACTGGTAGCTGCCAGGACTCCAATACAACGCCAGCA |
| 2:2212:13291:61112#2#R | \| \| |
| MQ=60 isize=-198 C=33 | CTGTCCCTTCTCCaCGGCCAcaGTCCCAG\|C\|CCCCCCaCTCCAGTCCTTCCCCAAGGATGCAGCCtCAGCCTtCTCCACACCACGTTTCCCCACAGACAcGTTCCCCaCATCCTGGACTGGTAGCTGCCAGGACTCCAATACAACGCCAGCA |
| 1:2106:9382:5521#2#R | \| \| |
| MQ=60 isize=-200 C=37 | tGTCCCTTCTCCACGGCCACAGTCCCAG\|C\|CCCCCCaCTCCAGTCCTTCCCCAAGGATGCAGCCTCAGCCTTCTCCACACCACGTTTCCCCACAGACAAGTTCCCCACATCCTGGACTGGTAGCTGCCCanGACTCCAATTTTTTGTCCCGG |
| 2:1216:5423:37592#2#F | \| \| |
| MQ=60 isize=421 C=40 | GCAAAAGTCCTGATTGGAGTnccTTCTCCACGGCCACAGTCCCAG\|C\|CCCCCCACTCCAGTCCTTCCCCAAGGATGCAGCCTCAGCCTTCTCCACACCACGTTTCCCCACAGACAAGTTCCCCACATCCTGGACTcgTAGCTGCCCAGGCCa |
| 2:1215:12978:71515#2#R | \| \| |
| MQ=60 isize=-200 C=33 | GTCCCTTCtCCACGGCCACaGTCCCAG\|C\|CCCCCCACtCCAGTCCTTCCCCAAGGATGCAGCCTCAGCCTTCTCCaCACCaCGTTTCCCCACAGACAAGTTCCCCACATCCTGGACTGGTAGCTGCCCAAgnACTCCAATGCACCCACACGT |
| 2:2202:8131:97726#2#R | \| \| |
| MQ=60 isize=-200 C=33 | gTCCCTTCTCCACGGCCACAGTCCCAG\|C\|CCCCCCACTCCAGTCCTTCCCCAAGGATGCAGCCTCAGCCTTCTCCACACCACGTTTCCCCACAGACAAGTTCCCCACATCCTGGACTGGTAGCTGCCCAAGGACTCCAATCAACCTCATATT |
| 1:1115:14228:21179#2#F | \| \| |
| MQ=60 isize=184 C=37 | AACGATGAATGTATTGGAGTCCTCCCTTCTCCACGGCCACAGTCCCAG\|C\|CCCCCCACTCCAGTCCTTCCCCAAGGATGCAGCCTCAGCCTTCTCCACACCACGTTTCCCCACAGACAAGTTCCCCACATCCTGGACtgGTAGCTGCCCAGG |
| 1:2106:6238:78525#2#F | \| \| |
| MQ=60 isize=184 C=37 | GTCTCGGGTGCTATTGGAGTCntCCCTTCTCCACGGCCACAGTCCCAG\|C\|CCCCCCACTCCAGTCCTTCCCCAAGGATGCAGCCTCAGCCTTCTCCACACCACGTTTCCCCACAGACAAGTTCCCCACATCCTGGACTGGTAGCTGCCCAGg |
| 1:2206:16725:27598#2#F | \| \| |
| MQ=60 isize=184 C=37 | ACGGGACCCTGGATTGGAGTCntCCCTTCTCCACGGCCACAGTCCCAG\|C\|CCCCCCACTCCAGTCCTTCCCCAAGGATGCAgCCTCAGCCTTCTCCACACCACGTTTCCCCACAGACAAGTTCCCCACATCCTGGACTGGTAGCTGCCCAGc |
| 2:1115:16046:32459#2#F | \| \| |
| MQ=60 isize=184 C=37 | AGGCCGGGCGATATTGGAGTncTCCCTTCTCCACGGCCACAGTCCCAG\|C\|CCCCCCACTCCAGTCCTTCCCCAAGGATGCAGCCTCAGCCTTCTCCACACCACGtTTCCCCACAGACAAGTTCCCCACATCCTGGACTGGtaGCTGCCCAGg |
| 2:2111:17968:70614#2#F | \| \| |
| MQ=60 isize=184 C=37 | CGGCGCCCGGGCATTGGAGTCCTCCCTTCTCCACGGCCACAGTCCCAG\|C\|CCCCCCACTCCAGTCCTTCCCCAAGGATGCAGCCTCAGCCTTCTCCACACCACGtTTCCCCACAGACAagtTCCCCACATCCtgGACtGGTAGCTGcCCAGc |
| 2:2115:20385:38217#2#F | \| \| |
| MQ=60 isize=184 C=40 | GTCTGTGCAGTGATTGGAGTncTCCCTTCTCCACGGCCACAGTCCCAG\|C\|CCCCCCACTCCAGTCCTTCCCCAAgGATGCAGCCTCAGCCTTCTCCACACCACGTTTCCCCACAgACAAgtTCCCCACAtCCTGGACTcGTagcTGcCCAgc |
| 2:2206:9089:36047#2#F | \| \| |
| MQ=60 isize=184 C=37 | TTCGGAAATGTTATTGGAGTCntCCCTTCTCCACGGCCACAGTCCCAG\|C\|CCCCCCACTCCAGTCCTTCCCCAAGGATGCAGCCTCAGCCTTCTCGACACCACGTTTCCCCACAGACAAGTTCCCCACATCCTGGACTGGTAGCTgCCCAGc |
| 1:2113:5950:28529#2#R | \| \| |
| MQ=60 isize=-201 C=2 | ccccttctccacggccacagtcccag\|c\|ccccccactccagtccttccccaaggatgcagcctcagccttctccacaccacgtttcCCCACAGACAAGTTCcCCACATCCTGGACTGGTAGCTGCCCAgannACTCCAATCGCATTCACACA |
| 2:1110:18875:57966#2#R | \| \| |
| MQ=60 isize=-201 C=33 | tcCCtTCTCCACGGCCACAGTCCCAG\|C\|CCCCCCACTCCAGTCCTTCCCCaAGGATGCAGCCTCAGCCTTCTCCACACCACGTTTCCCCACAGACAAGTTCCCCACATCCTGGACTGGTAGCTGCCCAGAGGACTCCAATTCTACGGCTCTC |
| 2:2111:11510:33808#2#R | \| \| |
| MQ=60 isize=-201 C=33 | cCCCTTCtCCACGGCcaccGTCCCAG\|C\|CCCCCCACTCCAGTCCTtCCCCAAGGATGCAGCCTCAGCCTTCTCCACACCACGTTTCCCCACAGACAAGTTCCCCACATCCTGGACTGGTAGCTGCCCAGAGGACTCCAATTTTCCGGCTCAA |
| 1:1201:11452:89854#2#R | \| \| |
| MQ=60 isize=-198 C=33 | ccgccccttctccacggCCaCAGTCCCAG\|C\|CCCCCcaCtCCAGTCCTtCCCCAAGGATGCAGCCTCAGCCTTCTCCACaCCACGTTTCCCCACAGACAAGTTCCCCACATCCTGGACTGGTAGCTGCCanGACTCCAATCGTTTCACTTAG |
| 1:2115:10474:12324#2#R | \| \| |
| MQ=60 isize=-203 C=37 | CCCTTCTCCACGGCCACAGTCCCAG\|C\|CCCCCCaccCCAGTCCTTCCCCAAGGATGCAGCCTCAGCCTTCTCCACACCACGTTTCCCCACAGACAAGTT\_CCCACATCCTGGACTGGTAGCTGCCCAGGCAGGACTCCAATCGCGCCGCCCCG |
| 1:2108:15813:5210#2#F | \| \| |
| MQ=60 isize=182 C=37 | ATAGGGGCGGACATTGGAGTCntCCTTCTCCACGGCCACAGTCCCAG\|C\|CCCCCCACTCCAGTCCTTCCTCAAGGATGCAGCCTCAGCCTTCTCCACACCACGTTTCCCCACAGACAAGTTCCCCACATCCTGGACTGGTAGCTGCCCAGGC |
| 2:1216:2623:96002#2#F | \| \| |
| MQ=60 isize=182 C=40 | CTTACTGTAGTGATTGGAGTncTCCTTCTCCACGGCCACAGTCCCAG\|C\|CCCCCCACTCCAGTTCTTCCCCAAGGATGCAGCCTCAGCCTTCTCCACACCACGTTTCCCCACAGACAAGTTCCCCACATCCtGGACTGGTAGCtGCCCAGGC |
| 1:1102:19746:29630#2#R | \| \| |
| MQ=60 isize=-203 C=33 | CCtTCTcCACGGCCaCaGTCCCAG\|C\|CCCCCCACTCCAGTCCTTCCCCAAGGATGCAGCCTCAGCCTTCTCCTCACCACGTTTCCCCACAGACAAGTTCCCCACATCCTGGACTGGTAGCTGCCCAGGCanGACTCCAATACACACCTCaCc |
| 1:2214:14842:42358#2#R | \| \| |
| MQ=60 isize=-203 C=2 | ccctctccacggccacagtcccag\|c\|cccccccctccAGTCCTTCCCCAAGGATGCAgCCTCAGCCTTCtCCACACCACGTTTCCCCACAGACAAGTTCCCCACATCCTGGACTGGTAGCTGCCCAGGCAgnACTCCAATGCGGACAGAGAC |
| 2:2203:20478:73129#2#R | \| \| |
| MQ=60 isize=-203 C=37 | CCTTCtCCACGGCCACAGTCCCAG\|C\|CCCCCCaCTCCAGTCCTTCCCCAAGGATGCAGCCTCAGCCTTCTCCACACCACGTTTCCCCGCAGACAAGTTCCCCACATCCTGGACTGGTAGCTGCCCAGGCanGACTCCAATAGCACCCGAACT |
| 1:1106:6423:2658#2#F | \| \| |
| MQ=60 isize=315 C=40 | TGCGCTCGGGGTATTGGAGTCntCTTCTCCACGGCCACAGTCCCAG\|C\|CCCCCCACTCCAGTCCTTCCCCAAGGATGCAGCCTCAGCCTTCTCCACACCACGTTTCCCCACAGACAAGTTCCCCACATCCTGGACTGgTAGCTGCCCAGGCC |
| 1:2112:4339:16203#2#F | \| \| |
| MQ=60 isize=286 C=27 | AGAGtGACAGCCATTGGAGTCCTCTTCTCCACGGCTACAGTCCCAG\|C\|CCCCcCACTCCAGTCCTTCCCCAAGGaTGCAgccTcagCCTTCTCCACACCacGtTtcCCCAcagaCAAGttCcccaCAtccTGGACTgGtAGCTGCCCAGGCC |
| 1:1201:13091:75531#2#R | \| \| |
| MQ=60 isize=-205 C=33 | cttctccccgGCCaCaGTCCCag\|C\|CCCCCCACTCCaGTCCTTCCCCAAGGATGCAGCCTCAGCCTTCTCCACACCACGTTTCCCCACAGACAAGTTCCCCACATCCTGGACTGGTAGCTGCCCAGGCCanGACTCCAATGTTTACAGCCGC |
| 1:1211:5780:82298#2#R | \| \| |
| MQ=60 isize=-198 C=27 | CTTCTCCACggCCacAGTCCCAG\|C\|CCCCCCaCTCCAGTCCTTCCCCAAGGATGCAGCCTCAGCCTTCTCCACACCACGTTTcCCCACAGACaAGTTCCCCACATCCTGGACTGGTAGCTGcCAAGgctAGGACTCCAATATCAAGCTAGGA |
| 2:1108:13413:90089#2#R | \| \| |
| MQ=60 isize=-205 C=27 | cttctccaCGGCCACAGtCCCAG\|C\|CCCCCcaCTCCAGTCCTTCCcCAAGGATGCAGCCTCAGCCTTCTCCACACCACGTTTCCCCACAGACAAGTTCCCCACATCCTGGACTGGTAGCTGCCCAGGCCanGACTCCAATTTTGAGCGAATA |
| 2:2116:8918:7201#2#R | \| \| |
| MQ=60 isize=-205 C=22 | cttCTCCACgGCCACAGTCCCAG\|C\|CCCCCCAcTCCAgtCcttCCCCAagGATGCAGCCTCAGCCTTCTCCACACCACGTTTCCcCACAGACAAGTTCCCCACATCCTGGACTGGTAGCTGCCCAGGCCAgnACTCCAATACCTAGGCGTAA |
| 2:1114:18539:16328#2#F | \| \| |
| MQ=60 isize=180 C=37 | GAGCGAGCGGACATTGGAGTncTTTCTCCACGGCCACAGTCCCAG\|C\|CCCCCCACTCCAGTCCTTCCCCAAGGATGCAGCCTCAGCCTTCTCCACACCACGTTTCCCCACAGACAAGTTCCCCACATCCTGGACTGGTAGCTGCCCAGGCCa |
| 2:2210:10402:7531#2#F | \| \| |
| MQ=60 isize=314 C=37 | TGGAGCCGCGTGATTGGAGTCCTTTCTCCACGGCCACAGTCCCAG\|C\|CCCCCCACTCCAGTCCTTCCCCAAGGATGCAGCCTCAGCCTTCTCCACACCACGTTTCCCCACAGACAAGTTCCCCACATCCTGGACTGGtaGCTGCCCAGGCCa |
| 2:2215:7558:47401#2#R | \| \| |
| MQ=60 isize=-185 C=2 | ttctccacggccacagtcccag\|c\|ccccCCACTCCaGtCctTCCCCAAGGatGCAGCCTcAGCCTtCTCCaccCCACGTTTCCCCACAGACAAGTTCCCCACATCCTGGCCTGGGCAGCTACCAGTCCAAGGACTCCAATCATTTAACTGGc |
| 2:2204:6692:5915#2#F | \| \| |
| MQ=60 isize=313 C=37 | GCCCGTGTGAGCATTGGAGTCntcTCCACGGCCACAGTCCCAG\|C\|CCCCCCACTCCAGTCCTTCgcCAAGGATGCAGCCtCAGCCTTCTCCaCACCACGTtTcCCCACAGACaaGtTCCCCaCATCCTGGaCtggtagCTGCCCagcccaac |
| 2:2113:2115:91101#2#R | \| \| |
| MQ=60 isize=-154 C=2 | cccgacccctaaattccccccccatcaagggggcgcccccccgcctcttccctctccaccgccacaggccccg\|c\|CCCCCcaCtCCaGTCCTTCCCCAAGGATGCAGCCTCAGCCTTCTCCACACCACGAgnACTCCAATCCCCAACGCCCG |
| 2:2113:4117:92064#2#F | \| \| |
| MQ=60 isize=178 C=37 | TTATTGAATGAGATTGGAGTncTCCACGGCCACAGTCCCAG\|C\|CCCCCCACTCCAGTCCTTCCCCAAGGATGCaGCCtcaGCCTTCTCCacaCCACGTTTCcccacagacaagttccccccaccctggcctggaagctgcccaggccacccc |
| 1:1201:16484:54742#2#R | \| \| |
| MQ=60 isize=-207 C=22 | CTCCACGGCCACagtCCCaG\|C\|CCCCCCaCTCCAGTCCTTCCCCAAGGATGCAGCCTCAGCCTTCTCCACACCACGTTTCCCCaCAGACAAGTTCCCCACATCCTGGACTGGTAGCTGCCCAGGCCAACanGACTCCAATAAGCGCCGTCGC |
| 1:2106:12010:14606#2#R | \| \| |
| MQ=60 isize=-207 C=37 | ctccaCGGCCaCAGTCCCAG\|C\|CCCCCCACTCCAGTCCTTCCCCAAGGATGCAGCCTCAGCCTTCTCCACACCACGTTTCCCCACAGGCAAGTTCCCCACATCCTGGACTGGTAGCTGCCCAGGCCAACanGACTCCAATATTAGAGCATAC |
| 2:1102:17803:5144#2#R | \| \| |
| MQ=60 isize=-207 C=33 | ccccacggccacAGtCCCaG\|C\|CCCCCCaCtCCAgTCCTTcCCCAAGGATGCAGCCTCAGCCTTCtCCACaCCACGTTTCCCCACAGACAAGTTCCCCACATCCTGGACTGGTAGCTGCCCAGGCCAACAGGACTCCAATGCCTTCGACCAT |
| 2:2206:10589:11859#2#R | \| \| |
| MQ=60 isize=-207 C=22 | ctCCaCGGCCACAGTCCCaG\|C\|CCCCCcaCtCCAGTCCTTCCCCAAGGATGCAGCCTCAGCCTTCTCCACACCACGTTTCCCCACAGACAAGTTCCCCACATCCTGGACTGGTAGCTGCCCAGGCCAACanGACTCCAATTTAAGCCTTGTA |
| 1:1209:5365:57197#2#F | \| \| |
| MQ=60 isize=177 C=37 | CACCCCATGGACATTGGAGTCntTCCACGGCCACAGTCCCAG\|C\|CCCCCCACTCCAGTCCTTCCCCAAGGATGCAGCCTCAGCCTTCTCCACACCACGTTTCCCCACAGACAAGTTCCCCACATCCTGGACTGGTAGCTGCCCAGGCCAACC |
| 2:2102:20589:64494#2#F | \| \| |
| MQ=60 isize=177 C=37 | GCACAAAATCTAATTGGAGTCCTTCCACGGCCACAGTCCCAG\|C\|CCCCCCACTCCAGTCCTTCCCCAAGGATGCAGCCTCAGCCTTCTCCACACCACGtTTCCCCaCAGACAAgTTcCCCACATCCTGGactGGtagcTGCCCAggcCaACC |
| 1:2109:13352:58370#2#R | \| \| |
| MQ=60 isize=-208 C=37 | cCCACGGCCACAGTCCCAG\|C\|CCCCCCaCTCCAGTCCTTCCCCAAGGATGCAGCCTCAGCCTTCTCCACACCACGTTTCCCCACAGACAAGTTCCCCACATCCTGGACTGGTAGCTGCCCAGGCCAACCanGACTCCAATACAGCTCCGTAT |
| 2:1211:9850:11903#2#R | \| \| |
| MQ=60 isize=-208 C=33 | cccacggccaCAGTCCCAG\|C\|CCCCCCACtCCAGTCCtTCCCCAAGGATGCAGCCTCAGCCTTCTCCACACCACGTTTCCCCACAGACAAGTTCCCCACATCCTGGACTGGTAGCTGCCCAGGCCAACCAGGACTCCAATATCAGAACCCTG |
| 2:2114:3018:7849#2#R | \| \| |
| MQ=60 isize=-185 C=2 | cccacggccacaggcccag\|c\|cccccccctccagtccttccccaaggatgcagccccaaccttccccaccccacgTTTCCCcacagacAAaTTCCCCACATCCtgGgCagCTaCCAGTCCAGGATGTGGAgnACTCCAATTTCACCGGGCCA |
| 1:1105:20559:94956#2#R | \| \| |
| MQ=60 isize=-184 C=2 | cccccccccccaccctgcccctcccccacggccacagtccccg\|c\|ccccccactccagtccttccccaaggatgcagcctcagccttctCCACaCCACgTTTCCCCACAGACAAGTTCCCCACAtCCTganGACTCCAAgGTATACCTGTGC |
| 1:1210:6185:23415#2#R | \| \| |
| MQ=60 isize=-209 C=37 | ccacGGCCacaGtCcCaG\|C\|CCCCCctCTCCAGTCCTTCCCCAAGGATGCAGCCTCAGCCTTCTCCACACCaCGTTTCCCCACAGACAAGTTCCCCACATCCTGGACTGGTAGCTGCCCAGGCCAACCCAGGACTCCAATATGTAGTATTTG |
| 1:2107:2014:22522#2#R | \| \| |
| MQ=60 isize=-209 C=37 | CCACGGCCACAGTCCCAG\|C\|CCCCCCACTCCAGTCCTTCCCCAAGGAttCAGCCTCAGCCTTCTCCACACCACGTTTCCCCACAGACAAGTTCCCCACATCCTGGACTGGTAGCTGCCCAGGCCAACCCAGGACTCCAATACCAACAGTCCA |
| 2:2107:5350:36715#2#R | \| \| |
| MQ=60 isize=-177 C=2 | caaccagggccccctcccccgcctggcccctccccacggccacagtccccg\|c\|CCCCCCACtCCAGTCCTTCCCCAAGGATGCAGCCTCAGCCTTCTCCACACCACGtcTCCCCACAGACAAGTTCCCCAGGACTCCAATATCAAAGCAGGT |
| 2:2213:8334:19829#2#R | \| \| |
| MQ=60 isize=-209 C=2 | ccacggccacagtcccag\|c\|ccccccactccagtccttccCCAAGGAtgctGCCTCAGCcttcTccActCccCGTTtccCCACAGACAAGTTCCccacaTCCTgGacTGGTAGCTGCCCAGGCCAACcCAGGACTCCAATTGCACCCGCCCA |
| 1:1207:10812:82595#2#F | \| \| |
| MQ=60 isize=175 C=33 | GTCTAAGACGAAATTGGAGTCntCACGGCCACAGTCCCAG\|C\|CCCCCCACTCCAGTCCTTCCCCAAGGATGCAGCCTCAGCCTtCTCCACACCACGTTTCCCCACAgAcAAGtTCCCCaCATCCTGGACTGGtAGCTGCCCAGgcCAACCCc |
| 1:1213:16121:62923#2#F | \| \| |
| MQ=60 isize=175 C=40 | GCCGTGATCAACATTGGAGTncTCACGGCCACAGTCCCAG\|C\|CCCCCCACTCCAGTCCTTCCCCAAGGATGCAGCCtCAGCCTTCTCCACACCACGTttCCCCACAGaCAAGTTCCCCACATCCTGgactgGtAGCTGCCCAGgCCAACCCC |
| 2:1103:14708:8403#2#F | \| \| |
| MQ=60 isize=175 C=22 | CTGATAACGGgAATTGGAGTCCTCACGGCCACAGTCcCaG\|C\|CCCCCCACTCCAGTCCTtCCcCAAGGATGcAgccTaAGCctTctcCaCaccaCGcTTCcCCACAGacaagTTCcCCACaTCCtGgaCtGGtAGCTGCCCAGgcCAACgcc |
| 2:2103:12749:3813#2#F | \| \| |
| MQ=60 isize=309 C=37 | ATGGTCGACTGGATTGGAGTCCTCACGGCCACAGTCCCAG\|C\|CCCCCCACTCCAGTCCTTCCCCAAGGATGCAGCCTCAGCCTTCTCCACACcacgTTTCCCCccagacAAGTTCCCcccatcctggaccggtacctccccagcccaccccc |
| 2:2206:14099:11274#2#F | \| \| |
| MQ=60 isize=175 C=33 | GGACAGCGGATCATTGGAGTCntcACGGCCACAGTCCCAG\|C\|CCCCCCACTCCAGTCCTTCCCCAAGGAtGCAGCCtCAgcCTTCTCCACACCACGtTTCCCcACAGACAAGtTCCCCACATCCtgGaCTGgtAGCTGCCCaGGCCAAcCCC |
| 1:2207:4629:51801#2#R | \| \| |
| MQ=60 isize=-211 C=27 | CAcGGCCACaGTCCCaG\|C\|CCCCCCaCTCCAGTCCTTCCCCAAGGATGCAGCCTCAGCCTTCTCCACACCACGTTTCCCCACAGACAAGTTCcCCACATCCTGGACTGGTAGCTGCCCAGGCCAACCCCanGACTCCAATAACAGAGTACTC |
| 2:1110:10717:32740#2#R | \| \| |
| MQ=60 isize=-211 C=33 | cacggccacagtCCCAG\|C\|CCCCCCactCCAGTCCTTCCCCAAGGatGCAGCCTCAGCCTTCtCCacaCCACGTTTCCCCACAGACAAGTTCCCCACATCCTGGACTGGTAGCTGCCCAGGCCAACCCCAGGACTCCAatAAAGAGACGCCT |
| 1:1213:10321:55192#2#F | \| \| |
| MQ=60 isize=174 C=37 | GTATTCCGACACATTGGAGTncTACGGCCACAGTCCCAG\|C\|CCCCCCACTCCAGTCCTTCCCCAAGGATGCAGCCTCAGCCTTCTCCACACCACGTTTCCCCACAGACAAGTTCCCCACATCCTGGACTGGTAGCTGCCCAGGCCAACCCCa |
| 2:1111:3799:38719#2#F | \| \| |
| MQ=60 isize=174 C=37 | TGCATCCTCGGAATTGGAGTCCTACGGCCACAGTCCCAG\|C\|CCCCCCACTCCAGTCCTTCCCCAAGGATGCAGCCTTAGCCTTCTCCacACCaCGTTtcCCCacagaCAAgTTCCCCcCAtCCtggcctGGTAgctgCCCaggCCAacCCcc |
| 2:1214:9541:7058#2#F | \| \| |
| MQ=60 isize=174 C=37 | TGCATCCTCGGAATTGGAGTCCTACGGCCACAGTCCCAG\|C\|CCCCCCACTCCAGTCCTTCCCCAAGGATGCAGCCTTAGCCTTCTCCACACCACGTTTCCCCaCAGACAAGTTCCCCACATCCTGGACTGGTAGCtGCCCAGGCCAACCCCA |
| 2:1211:16364:61899#2#R | \| \| |
| MQ=60 isize=-141 C=33 | cCGGCCaCAGtCCCAG\|C\|CCCCCCACTCCAGTCCTtCCCCAAGGATGCAGCCTCAGCCT |
| 1:1115:8335:95771#2#F | \| \| |
| MQ=60 isize=173 C=40 | GGAGGGGCAACGATTGGAGTCCTCGGCCACAGTCCCAG\|C\|CCCCCCACTCCAGTCCTTCCCCAAGGATGCAGCCTCAGCCTTCTCCACACCACGTTTCCCCACAGACAAGTTCCCCACATCCTGGACTGGTAGCTGCCCAGGCCAACCCCAT |
| 1:1214:5423:39941#2#F | \| \| |
| MQ=60 isize=173 C=40 | CAGGTTCATGGTATTGGAGTncTCGGCCACAGTCCCAG\|C\|CCCCCCACTCCAGTCCTTCCCCAAGGATGCAGCCTCAGCCTTCTCCACACCACGTTTCCCCACAGACAAGTTCCCCACATCCTgGACTGGTAGCTGCCCAgGCCAACCCCAT |
| 1:2206:15750:32949#2#F | \| \| |
| MQ=60 isize=307 C=15 | TCTGGcGCGATTATTGGAGgCntcGgCCACAtTCcCAg\|c\|CCCCCcactccagtccttccccaaggatgcagcctcagccttctccacaccacgtttccccacagccaagttccccacatcctggactggcagccgcccagcccaaccccat |
| 2:1206:13001:48141#2#F | \| \| |
| MQ=60 isize=173 C=37 | CGCGGGGCAGGAATTGGAGTCntCGGCCACAGTCCCAG\|C\|CCCCCCACTCCAGTCCTTCCCCAAGGATGCAGCCTCAGCCTTCTCCACACCACGTTTCCCCACAGACAAGTTCCCCACATCCTGGACTGGTAGCTGCCCAGGCCAACCCCat |
| 2:1211:1884:2474#2#F | \| \| |
| MQ=60 isize=307 C=37 | TCTGGCGCGATTATTGGAGTCCTCGGCCACAGTCCCAG\|C\|CCCCCCACTCCAGTcCTTCCCCAAGGAtGCaGCCTCAGCCTTCTCCaCgccacGTTTCCCCAcacacaatttccccacatcctggactggtagctgcccaggccaaccccat |
| 2:2202:6376:4936#2#F | \| \| |
| MQ=60 isize=173 C=40 | TCACGGATTAGAATTGGAGTCCTCGGCCACAGTCCCAG\|C\|CCCCCCACTCCAGTCCTTCCCCAAGGATGCAGCCTCAGCCTTCTCCACACCACGTTTCCCCACAGACAAGTTCCCCACATCCTGGactGGtaGCTGCCCAGGCCAACCCCAt |
| 2:2210:15272:34008#2#F | \| \| |
| MQ=60 isize=278 C=22 | GAGGCGGAGGATATTGGAGTCCTCGGCCACAGTCCcAG\|C\|CCcCCCActCCAGcCCTTCCCCAAGGAtGCAGCCTccgcattCtcCAcaCCacgtttCCCcacagaaaatttccccacatcctggactggtagctgcccagcccaaccccat |
| 1:1204:6769:23442#2#R | \| \| |
| MQ=60 isize=-212 C=33 | CGGCCACAGTCCCag\|C\|CcCcCCACTCCAGTCCTTCCCCAAGGATGCAGCCTCAGCCTTCTCCaCACCACGTTTCCCCACAGACAAGTTCCCCACATCCTGGACTGGTAGCTGCCCAGGCCAACCCCATanGACTCCAATGAATTCGTGAAG |
| 1:2202:11465:24271#2#R | \| \| |
| MQ=60 isize=-212 C=27 | CggCCACagtCCCaG\|C\|CCcCCCACTCCAGTCCTTCCCCAAGGATGCAgcCTCAGCCTTCTCCACACCaCGTTTCCCCACAGACAAGTTCCCCACATCCTGGACTGGTAGCTGCCCAGGCCAACCCCATanGACTCCAATGCCTACAAAAGG |
| 2:1101:12776:40773#2#R | \| \| |
| MQ=60 isize=-212 C=2 | cggccccaggcccag\|c\|ccccccaCtccaGTCCTTCCCCAAGGATgCAGCCtCAGCCTTCTCcacaCCaCGTTTCCCCACAGACAAGTTCCCCACATCCTGGACTGGTAGCTGCCCAGGCCAACCCCATAGGACTCCAATGAGTCAAACGGC |
| 2:1213:6844:22903#2#R | \| \| |
| MQ=60 isize=-212 C=37 | CGGCCACAGTCCCAG\|C\|CCCCCCACtCCAGTCCTTCCCCAAGGATGCAGCCTCAGCCTTCTCCACaCCACGTTTCCCCACAGACAAGTTCCCCACATCCTGGACTGGTAGCTGCCCAGGCCAACCCCATAGGACTCCAATGAGTCAAACGGC |
| 2:2101:2285:24717#2#R | \| \| |
| MQ=60 isize=-168 C=2 | tccccccccaaaccaaggcccctcccccaggcctgtccctccccccggccccagtcccag\|c\|ccccccccccccatccctCCCCaAGGATgCAGCCTCAGCCTTCTCCaCACCACGTTTCCCCACAGACAGGACTCCAATATAGATAATGAG |
| 2:2201:19944:88814#2#R | \| \| |
| MQ=60 isize=-212 C=37 | CGGCCACAGTCCCAG\|C\|CCCCCCACTCCAGTCCTTCCCCAAGGATGCAGCCTCAGCCTTCTCCACACCACGTTTCCCCACAGACAAGTTCCCCACATCCTGGACTGGTAGCTGCCCAGGCCAACCCCATanGACTCCAATCTGCAACCATAA |
| 1:1214:7515:42774#2#F | \| \| |
| MQ=60 isize=172 C=37 | CCCCTCTCAAATATTGGAGTncTGGCCACAGTCCCAG\|C\|CCCCCCACTCCAGTCCTTCCCCAAGGATGCAGCCTCAGCCTTCTCCACACCACGTTTCCCCACAGACAAGTTCCCCACATCCtGGACTGGTAGCTGCCCAGGCCAACCCCATG |
| 2:1109:6398:70319#2#F | \| \| |
| MQ=60 isize=172 C=33 | TGTAAATAGATAATTGGAGTCntcTGTGGCCGTgGAGAAGGGACAGGCCACAGtCCcAG\|C\|CCCCCCacTCCAGtcCTtCCCcaagaatgaagcctacgccttctccacaccacgtttccccacaaacaagttccccccaccctggactgga |
| 2:1115:17358:72478#2#F | \| \| |
| MQ=60 isize=172 C=37 | GCGGCTGAGTGGATTGGAGTnctGGCCACAGTCCCAG\|C\|CCCCCCACTCCAGTCCTTCCCCAAGGATGCaGcCTCAGCCTTCTCCACACCACGTTTCCCCACAGACAAGTTCCCCACATCCtgGACTGGTAgCTGCCCaggCCAAcCCCATG |
| 2:2108:2557:39076#2#F | \| \| |
| MQ=60 isize=172 C=37 | GCGGAGGGGGCAATTGGAGTCCTGGCCACAGTCCCAG\|C\|CCCCCCACTCCAGTCCTTCCCCAAGGAtgcAgCCTCAgCCTTCtCCccaCcacgTTtCCCCccagccaagttccccccatcctggccggggacctccccagccaacccccagg |
| 1:1104:20164:90904#2#F | \| \| |
| MQ=60 isize=276 C=27 | gacgCcCCCTTgCtcgcccccncgcCACAGtCCCAG\|C\|CCCCCcactccagcccttccccaaggatgcagcctcagccttctccacaccccgtttccccacacacaagctccccacaccctggactggtagctgcccagcccacccccacgg |
| 1:2103:7593:20913#2#F | \| \| |
| MQ=60 isize=171 C=33 | GCCGTGCGTCCTATTGGAGTCCTGCCACAGTCCCAG\|C\|CCCCCCACTCCAGTCCTTCCCCAAGGATGCAGCCTCAGCCTTCTCCACACCACGTTTCCCCaCAGACAaGTTCCCCACATCCTGGACtGgtAGCTGCCCaGGCCAACCCCAtGG |
| 1:2208:3417:30806#2#F | \| \| |
| MQ=60 isize=171 C=37 | GGGGCCTGGTACATTGGAGTCntGCCACAGTCCCAG\|C\|CCCCCCACTCCAGTCCTTCCCCAAGGATGCAGCCTCAGCCTTCTCCACACCACGtTTCCCCACAGACAAGTTCCCCACATCCTGGACTGGTAGCTGCCCAGGCCAACCCCATgg |
| 2:1113:16414:57176#2#F | \| \| |
| MQ=60 isize=171 C=40 | TACACACTGGTAATTGGAGTncTGCCACAGTCCCAG\|C\|CCCCCCACTCCAGTCCTTCCCCAAGGATGCAGCCTCAGCCTTCTCCACACCACGtTTCCCCACAGACAAGTTCCCCACATCCTGGACtGGTAGCTGCCCAGGCCAACCCCATGG |
| 2:1202:1839:65682#2#F | \| \| |
| MQ=60 isize=171 C=40 | TTTAGCCGAAATATTGGAGTCntGCCACAGTCCCAG\|C\|CCCCCCACTCCAGTCCTTCCCCAAGGATGCAGCCTCAGCCTTCTCCACACCACGTTTCCCCACAGaCAAGTTCCCCACATCCTGGcctGGtAgCTgCCCaGgCCAACCCcatgg |
| 2:1214:7134:29647#2#F | \| \| |
| MQ=60 isize=171 C=37 | GGTTGCGGCCGGATTGGAGTCCTGCCACAGTCCCAG\|C\|CCCCCCACTCCAGTCCTTCCCCAAGGATGCAGCCTCAGCCTTCTCCACACCACGTTTCCCCACAGACAAGTTCCCCACATCCTGGACTGGTAGCTGCCCAGGCCAACCCCATGG |
| 2:2212:10204:28954#2#F | \| \| |
| MQ=60 isize=171 C=40 | GGCCTGGTTGCTATTGGAGTCCTGCCACAGTCCCAG\|C\|CCCCCCACTCCAGTCCTTCCCCAAGGATGCAGCCTCAGCCTTCTCCACACCACGTTTCCCCACAGACAAGTTCCCCACATCCTGGACTGGTAGCTGCCCAGGCCAACCCCatgG |
| 1:2207:20124:88770#2#R | \| \| |
| MQ=60 isize=-215 C=22 | gCCACAGtCCCaG\|C\|CCCCCCaCTCCAGtCCTTCCCCAAGGAtGCAGCCTCAGCCTTctCCACACCACGTTTCCCCACAGACAAGTTCCCCACATCCTGGACTGGTAGCTGCCCAGGCCAACCCCATGGanGACTCCAATGACTGCCTCTAG |
| 1:1102:9601:41674#2#F | \| \| |
| MQ=60 isize=275 C=15 | gTtTatTGGGACATTGGAGTCntctcCACAGTCCCAg\|c\|CCCCCCACtCCAGTCcTTCCccaaggatgcagcctcagccttctccacaccacgtttccccacagacaagctccccacatcctggactggtagctgcccaggccaaccccacc |
| 1:1202:13850:63806#2#F | \| \| |
| MQ=60 isize=275 C=40 | AGCTCGCTGGGGATTGGAGtCCTCCACAGTCCCAG\|C\|CCCCCCACTCCAGTCCTTCCCCAAGGATGCAgCCTCAGCCTTCtCCACACCaCGTTTCCCCACAGaCAcGTTCCCCACATCCTGgactggtagctgcccaggccacccccatgga |
| 1:2115:4151:22000#2#F | \| \| |
| MQ=60 isize=170 C=33 | GGATTCTTACTTATTGGAGTCCTCCACAGTCCCAG\|C\|CCCCCCACTCCAGTCCTTCCCCAaGGATGCAGCCTCAGCCTTCTCCaCACCACgTttCCCcaCAGACAAGtTCcCCACAtCCTGGacTGGtagcTgcCCaggCCAaCCCCATgga |
| 2:2113:12162:7125#2#F | \| \| |
| MQ=60 isize=170 C=37 | ATTGCACGTAACATTGGAGTncTCCACAGTCCCAG\|C\|CCCCCCACTCCAGTCCTTCCCCAAGGATGCAGCCTCAGCCTTCTCCACACCACGTtTCCCCACAgACAAGTTCCCCACATCCTGAaCtggTAGCTGcCCAGgCCAACCCCatgga |
| 1:1211:6278:47146#2#R | \| \| |
| MQ=60 isize=-216 C=2 | ccccactcccag\|c\|ccccccactccagtccttccccaagcatgcagcctcagccttctccacaccacgtttccccacagacaagttccccacatcctggactgGTAGCTGCCCAGGcCAACCCCATGGaaGGacgccaagCCcCCCCCACCC |
| 2:2113:12676:59368#2#R | \| \| |
| MQ=60 isize=-216 C=33 | CCACaGtCccAG\|C\|CCCcCcACTCCAGTCCTTCCcCAAGGATGCAGCCTCAGCcTTCTCCACACCACGTTtCCCCACAGACAAGTTCCCCACATCCTGGACTGGTAGCTGCCCAGGCCAACCCCATGGAAgnACTCCAATCGTACCTGAGGT |
| 1:2105:16558:28341#2#F | \| \| |
| MQ=60 isize=169 C=37 | TACAGAACAGCCATTGGAGTCntCACAGTCCCAG\|C\|CCCCCCACTCCAGTCCTTCCCCAAGGATGCAGCCTCAGCCTTCTCCACACCACGTTTCCCCACAGACAAGTTCCCCACATCCTGGACTGGTAGCTGCCCAGGCCAacCCCATGGAA |
| 1:2203:14171:3814#2#F | \| \| |
| MQ=60 isize=169 C=37 | ATTGTTCTAGAAATTGGAGTCCTCACAGTCCCAG\|C\|CCCCCCACTCCAGTCCTTCCCCAAGGATGCAGCCTCAGCCTTCTCCACACCACGTTTCCCCACAGAcAAGTTCCCCacATCCTGGACtGGtagCtgCCCagcccAaCCCCAgGGAa |
| 2:1111:16575:100240#2#F | \| \| |
| MQ=60 isize=169 C=37 | TTGCGGCACGGTATTGGAGTCCTCACAGTCCCAG\|C\|CCCCCCACTCCAGTCCTTCCCCAAGGATGCAGCCTCAGCCTTCTCCACACCACGTTTCCCCACaGacAaGTTCCCCACATCCTGGACTGGTAGCTGCCCAGGCCaaccccatggaa |
| 2:2214:18436:7169#2#F | \| \| |
| MQ=60 isize=169 C=37 | GGAGAGAGGCGAATTGgAGTCCTCACAGTCCCAG\|C\|CCCCCCACTCCAGTCCTTCCCCAAGGATGCAGCCTCAGCCTTCTCCACACCACGTTTCCCCACAGACAAGtTCCCCACATCCTGGACTGGTAGCtGCCCAGGCCAACCCCATGGAA |
| 2:1106:8512:10165#2#F | \| \| |
| MQ=60 isize=168 C=37 | GTTACTTTTATTATTGGAGTCntACAGTCCCAG\|C\|CCCCCCACTCCAGTCCTTCCCCAAGGATGCAGCCTCAGCCTTCTCCACaCCACGTTTCCCCACAGaCaaGTCCCCCccAtCCtggccTGGtaGCTgcCcaggCCAacCCcagggaaa |
| 2:1213:19743:82939#2#F | \| \| |
| MQ=60 isize=168 C=37 | TAGGGAGCCGGTATTGGAGTCCTACAGTCCCAG\|C\|CCCCCCACTCCAGTCCTTCCCCAAGGATGCAGCCTCAGCCTTCTCCACACCACGtTTCCCCACAGACAAGTTCCCCACATCCTGGACTGGTAGCTgCCCAGGCCAACCCCATGGAAC |
| 2:2116:1562:97991#2#F | \| \| |
| MQ=60 isize=168 C=40 | TATCACATTAGTATTGGAGTncTACAGTCCCAG\|C\|CCCCCCACTCCAGTCCTTCCCCAAGGAtGCAGCCTCAGCCTTCTCCACaCCACGTTTCCCCccagacaagTTCCCCacctccgggccgggagccgccccagccaaccccctgggaaa |
| 1:1109:2812:53631#2#R | \| \| |
| MQ=60 isize=-218 C=37 | aCAGTCCCAG\|C\|CCCCCCACTCCAGTCCTTCCCCAAGGATGCAGCCTCAGCCTTCTCCACACCACGTTTCCCCACAGACAAGTTCCCCACATCCTGGACTGGTAGCTGCCCTGGCCAACCCCATGGAACAGGACTCCAATCCCCAGATGAAT |
| 1:2202:15553:84023#2#R | \| \| |
| MQ=60 isize=-218 C=33 | CcaGTCCCAG\|C\|CCCCCCaCTCCAGTCCTTCCCCAAGGATGCAGCCTCAGCCTTCtCCaCACCaCGTTtCCCCACAGACAAGTTCCCCACATCCTGGACTGGTAGCTGCCCAGGCCAACCCCATGGAACanGACTCCAATTTGGTACGGGCG |
| 2:1105:2560:56998#2#F | \| \| |
| MQ=60 isize=167 C=37 | CTGGGAAGTGCGATTGGAGTCntCAGTCCCAG\|C\|CCCCCCACTCCAGtCCTTCCCCAaggatgcagcctcacccttctccacaccacgtttccccacagccaagtcccccccaccctgaccgggaagttccccggcccaccccctgggacaa |
| 1:1115:12237:7231#2#F | \| \| |
| MQ=60 isize=166 C=33 | GTATATTTCGTCATTGGAGTCCTAG\|C\|cCCCCCACTCcAGTCCTTCCCCAAgGATGCAGCCTCAGCCCTCTCCACACCaCGTTTcCCCACagaCAAGTTCCCCACATCCTGGaCtGGcagCTgCCCAGGCCAACCcCatGGAACAagGGCAt |
| 1:1114:9747:33082#2#F | \| \| |
| MQ=60 isize=299 C=40 | ACCTTCTCGTTTATTGGAGTCCTGTCCCAG\|C\|CCCCCCACTCCAGTCCTTCCCCAAGGATGCAGCCTCAGCCTTCTCCACACCACGTTTCCCCACAGACAAGTTCCCCACATCCTGGACTGGtAgCTGCCCAGgcCAacCCCATGGAacaag |
| 1:2109:9410:87206#2#F | \| \| |
| MQ=60 isize=165 C=37 | CTCCTCCCATATATTGGAGTCntGTCCCAG\|C\|CCCCCCACTCCAGTCCTTCCCCAAGGATGCAGCCTCAGCCTTCTCCACACCACGTTTCCCCACAGACAaGTtCCCCACATCCTGGACTGgTAGCTGCCCAGGCCAACCCCATGGAACAAG |
| 2:1106:3651:20079#2#F | \| \| |
| MQ=60 isize=165 C=37 | CTGTGTTGGGAATTGGAGTCCngtCCCAG\|C\|CCCCCCACTCCAGTCCTcCCCCAAGGAtgcagcctcAGCCTTCtCCacaCCAcgtTtcccCaCAgACaagttCCCCacaccctggccggctagctccccaggccaacccccgggaacaagg |
| 2:1114:18225:42372#2#F | \| \| |
| MQ=60 isize=165 C=37 | GTGTGCCAGCTGATTGGAGTncTGTCCCAG\|C\|CCCCCCACTCCAGTCCTTCCCCAAGGATGCAGCCTCAGCCTtctCCAcaCcaCGtttCCCCACAGaCAAgttcCcCACATCCTGGACTGGTAGCTGCCCAGGcCaACCCCATGGAActag |
| 2:2110:12653:59681#2#F | \| \| |
| MQ=60 isize=299 C=33 | CCGCGAAGCTCGATTGGAGTCCTGTCCCAG\|C\|CCCCCCACTCCAGTCCTTCCCCAAGGATGCAGCCTCAGCCTTCTCCACACCACGTTTCCCCACAGACAAGTTCCCCACATCCtgGACTGGtagCTgCCCagGcCAACcccttggaacaag |
| 2:2201:18215:17240#2#F | \| \| |
| MQ=60 isize=299 C=40 | CGTGGCTGAGTGATTGGAGTCntGTCCCAG\|C\|CCCCCCACTCCAGTCCTTCCCCAAGGATGCAGTCTCAGCCTTCTCCACACCACGTTTCCCCACAGACAAGTTCCCCACATCCTGGACTGGTAGCTGCCCAGgCCAACCCCATGGAACAAG |
| 1:1108:3107:32570#2#R | \| \| |
| MQ=60 isize=-220 C=33 | gaCCCAG\|C\|CCaCCccCTCCAGtCCttCCCCAAGGATGCAGCCTCAGCCTtCTCCACaCCaCgTTtCCCCACAGACAAGTTCCCCACAtCCTGGACTGGTAGCTGCCCAGGcCAACCCCATGGAACAAGanGACTCCAATAGGATAACGCCG |
| 1:2203:1844:33397#2#R | \| \| |
| MQ=60 isize=-220 C=15 | gTCccAG\|c\|ccCCCCACTCCAGtCCtTCCCCAAGGATGCAGCCTCAGCcTTCtCCACaCCACGTttcCCCACAGACAAGTTCCCCACaTcCTGGACTGGTAGCTGCCCAGGCCAACCCCATGGAACAAGAGGACTCCAATAGGATAACGCCG |
| 2:2108:16991:97946#2#R | \| \| |
| MQ=60 isize=-218 C=33 | ccgGTCCCag\|C\|CCCCCCactcCAGTCCttCCCcAAGGATGCAGCCTCAGCCTTCTCCACACCACGTttCcCCACAGACAAGTTCCCCACATCCTGGACTGGTAGCTGCCCAGGCCAACCCCATGGAACAGGACTCCAATCCGTCATTCCCG |
| 1:1109:20301:57215#2#F | \| \| |
| MQ=60 isize=164 C=37 | AAAGAACAACGGATTGGAGTCCTCCCAG\|C\|CCCCCCACTCCAGTCCTTCCCCAAGGATGCAGCCTCAGCCTTCTCCACACCACgTTTCCCCACAGACAAGTTCCCCACATCCTGGACTGGtAgCTGCCCAGgCCAACCCCAtGGAACAAggg |
| 1:1210:6528:76262#2#F | \| \| |
| MQ=60 isize=164 C=37 | CAGCGTGTGACGATTGGAGTCCTCCCAG\|C\|CCCCCCACTCCAGTCCTTCCCCAAGGATGCAGCCTCAGCCTTCTCCACACCACGTTTCCCCACAGACAAGTTCCCCACATCCTGGACTGGTAGCTGCCCAGGCCaACCCCATGGAACAAGGG |
| 1:2112:12044:44873#2#F | \| \| |
| MQ=60 isize=164 C=37 | CTGTGTTGGGAATTGGAGTCCTTTCCCAG\|C\|CCCCCCACTCCAGTCCTTCCCcAAGGAtgCaGCCTCAGCCTTCTCCACACCaCgtTTCCCCACAGaCAAGTTCCCCACatCCTGGACTGgtAGCTgCCCAGGCCAaCCCCATGgAACAAgg |
| 1:2216:16642:71979#2#F | \| \| |
| MQ=60 isize=164 C=37 | GCTCGAGTTCCCATTGGAGTncTCCCAG\|C\|CCCCCCACTCCAGTCCTTCCCCAAGGATGCAGCCTCAGCCTTCTCCACaCCACGTTTCCCCACAGaCAAGTTCCCCacATCCtGGaCTGGTAGCTgCCCAGGCCAACCCCATGgAacaaggg |
| 2:1114:7443:21640#2#F | \| \| |
| MQ=60 isize=298 C=37 | GTATGCGGACGAATTGGAGTncTCCCAG\|C\|CCCCCCACTCCAGTCCTTCCCCAAGGAtGCaGCctCAGCCTTCTCCACAcCAcGTTTCCccaCAGAcaAttTccCCACATCCtgGACaggtagctgCCCAggccaaccccatggaacaaggc |
| 2:1206:10137:75736#2#F | \| \| |
| MQ=60 isize=164 C=40 | ACTGTTCGCTTGATTGGAGTCntCCCAG\|C\|CCCCCCACTCCAGTCCTTCCCCAAGGATGCAGCCTCAGCCTTCTCCACACCACGTTTCCCCACAGACAAGTTCCCCACATCCTGGACTgGtAGCTGCCCAGgCCAACCCCATGGAACAAGGG |
| 2:1214:14793:73229#2#F | \| \| |
| MQ=60 isize=164 C=27 | AGgGGTGGGCGCATTGGAGTCCTCCCAG\|C\|CCCCCCACTCCAGTCCTTCCCCAAGGAtGCAGCCTCAgCcTTCTCCACAcCACGtttccccACagaaAAgatCcCCACATCCTtGACTggtAgctGCCcAGGccAACCCCctgGAACAAggg |
| 2:2107:7679:86244#2#F | \| \| |
| MQ=60 isize=164 C=33 | CGAGGAGGGGcGATTGGAGTCCTCCCAG\|C\|CCCCCCACTCCAGTCCTTCCCCAAGGATGCAGCCTCAGCCTTctccacaccccgtttccccacagacaagttcccaacatcctggactggtacctgcccgggccaaccccat\_gaacaaaggc |
| 2:2209:7383:32815#2#F | \| \| |
| MQ=60 isize=164 C=40 | GGCCCTGTGCACATTGGAGTCntCCCAG\|C\|CCCCCCACTCCAGTCCTTCCCCAAGGATGCAGCCTCAGCCTTCTCCACACCACGTTTCCCCACAGACAAGTTCCCCACATCCTGGACTGGTAGCTGCCCAGGCCAACCCCATGGAACAAGGg |
| 1:2102:9041:33643#2#R | \| \| |
| MQ=60 isize=-221 C=37 | tCCCAG\|C\|CCCCCCaCTCCAGTCCTTCCCCAAGGATGCAGCCTCAGCCTTCTCCACACCACGTTTCCCCACAGACAAGTTCCCCACATCCTGGACTGGTAGCTGCCCAGGCCAACCCCATGGAACAAGGAGGACTCCAATGCAGGGTGATAC |
| 1:2206:11983:61400#2#R | \| \| |
| MQ=60 isize=-221 C=2 | tcccag\|c\|cCCCCcaCtCCAGtCCTTCCCcaaggatgaAGCCTcaGcCTtctCCaCaCCaCGtTTCCCCACAGACAAGTTCCcCACATCCTGGACTGGTAGCTGcctAGGCCAtCcCCaTGGAACAAGganGACTCCAATGAcTGAGtcGCG |
| 1:1107:18847:38176#2#R | \| \| |
| MQ=60 isize=-222 C=37 | CCCAG\|C\|CCCCCCaCTCCAGTCCTTCCCCAAGGATGCAGCCTCAGCCTTCTCCACACCACGTTTCCCCACAGACAAGTTCCCCACATCCTGGACTGGTAGCTGCCCAGGCCAACCCCATGGAACAAGGGAGGACTCCAATCTGTACCATTAA |
| 2:1213:1785:62075#2#R | \| \| |
| MQ=60 isize=-222 C=37 | CCCaG\|C\|CCCCCCaCtCCAGTCCtTCCCCAAGGATgCagCCtCAGCCTCCTCcaCaCCACGTTTCCCCACAGACAAGTTCCCCaCATCCTGGACTGGTAGCTGCCCAGGCCAACCCCATGGAACAAGGGAGGACTCCAATCTGTACCATTAA |
| 2:2203:2269:39299#2#R | \| \| |
| MQ=60 isize=-212 C=2 | ccgcccaagacccag\|c\|ccccccccgcctgtccttccccaaggatgcacccctcgccttctccaccccacgtttccccacagaccagttccccacatcctggactggtagctgcccaggccaaccccatangactccaatgccgcccccccc |
| 1:1115:9615:7043#2#F | \| \| |
| MQ=60 isize=162 C=33 | GGCCGTGGGGGGATTGGAGTCCTCCAG\|C\|CCCCCCACTCCAGTCCTTCCCCAAGGATGCAGCCTCAGCCTTCTCCACACCACGTTTCCCCACAGACAAGTTCCCCACATCCTGGACTGGTAGCTGCCCAGGCCAACCCCATGGAACAAGGGC |
| 1:2209:5841:33720#2#F | \| \| |
| MQ=60 isize=162 C=37 | TTTAATCGTATTAtTGGAGTCntCCAG\|C\|CCCCCCACTCCAGTCCTTCCCCAAGGATGCAGCCTCAGCCTTCTCCACACCACGtTTCCCCACAGACAAGTTCCCCACATCCTGGACTGGTAGCTGCCCAGGCCAACcCCATGGAACAAGgGC |
| 2:1111:9784:65281#2#F | \| \| |
| MQ=60 isize=162 C=33 | GGCCGTGGGGGGATTGGAGTCCTCCAG\|C\|CCCCCCACTCCAGtCCTTCCCCAAGGATGCAGCCTCAGCCTTCTCCACACCACGtTTCCCCACAGACAAgtTCcCCACATCCTGGACtgGTagcTgCcCaGGcCAaCCCCAtgGaacAAGggc |
| 2:2111:14854:77175#2#F | \| \| |
| MQ=60 isize=162 C=37 | GGCGGATGCTGGATTGGAGTCCTCCAG\|C\|CCCCCCACTCCAGTCCTTCCCCAAGGATGCAGCCTCAGCCTTCTCCACACCACGtTTCCCCACAGACAAGtTCCCCaCATCCTGgACtGGTAgCTGCCCAGGCCaACCCCATGGAACAAGGGC |
| 1:2205:15776:22028#2#R | \| \| |
| MQ=60 isize=-224 C=2 | ccag\|c\|ccccccactccagtccttccccaaggatgcagcctcagccttctccacaccacgtttccccacagaaaagttcccCACAtCCTGGACTGGTAGCTGCCCAGGCCAACCCCATGGAACAAGGGCanGACTCcAatTCAAGGAAAATA |
| 1:1103:14617:92343#2#F | \| \| |
| MQ=60 isize=161 C=37 | ACCCCCCGTTTAATTGGAGTCCTCACAG\|C\|CTCCCCACTCCAGTCCTTCCCCAAGGATGCAGCCTCAGCCTTCTCCACACCACGTTTCCCCACAGACAAGtTCcCCACATCCTGGACTGGtAGCtGCCCAGGCCAACCCCATGGAACAAGGG |
| 1:2106:8826:22398#2#F | \| \| |
| MQ=60 isize=161 C=37 | CTGGCCGTTTCAATTGGAGTCntCAG\|C\|CCCCCCACTCCAGTCCTTCCCCAAGGATGCAGCCTCAGCCTTCTCCACACCACGTTTCCCCACAGACAAGTTCCCCACATCCTGGACTGGTAGCTGCCCAGGCCAACCCCATGGAACAAGGGCc |
| 2:1109:5484:54931#2#F | \| \| |
| MQ=60 isize=161 C=33 | CACGCCGAAGGGATTGGAGTCntCAG\|C\|CCCCCCACTCCAGTCCTTCCCCAAGGATGCAGCCTCAGCCTTCTCCaCACCaCgTTTCCCCACAGaCAagTtcCCCcCatCCTGGactGGtAGCtgcCCAGgCCaaccccctggaacaagggct |
| 2:2205:21308:36884#2#F | \| \| |
| MQ=60 isize=161 C=33 | TGTCTTTGGTAGATTGGAGTCntCAG\|C\|CCCCCCACTCCAGTCCTTgcccAAGGATGCAGCCTCAGCCTTCTCCACACCACGtTTCCCCACAGACAAGTTCCCCACATCCTGGACTGGTAGCTGCCCAGGCCAACCCCATGGAACAAGGGCT |
| 2:1207:2298:88640#2#R | \| \| |
| MQ=60 isize=-224 C=22 | CAG\|C\|CCCCcCaCtCCAGTCCTtCCCCAAGGATGCaGCCTCAGCcTtCtCcAcccCcCGTTTCCCCACaGACAAGTTCCCCACaTCCTGGACTGGTAGCTGCCCAGGCCAACCCCATGGAACAAGGGCAanGACTCCAATGCCACTGGAGCC |
| 1:1105:7987:6838#2#F | \| \| |
| MQ=60 isize=396 C=15 | TCGTCGGTGATGaTTGGAGTCntAG\|c\|CCCCCCAcTCCAGTCCTTCCCCAAGGAtGCAGCCTCAGCCTTCTCCACACCACgtttCcCCACAGACAAGTTCCCCACATCCTGGACtGGTAGCtgCCCAGgCCaacCCCatGGAACAagggcat |
| 1:2116:7579:88494#2#F | \| \| |
| MQ=60 isize=160 C=37 | GCGAGGCACTTGATTGGAGTCntaG\|C\|CCCCCCACTCCAGTCCTTCCCCAAGGATGCAGCCTCAGCCTTCTCCACACCACGTTTCCCCACAGACAAGTTCCCCACATCCTGGACTGGTAGCTGCCCAGGCCAACCCCATGGAACAAGGGCAT |
| 2:1204:8455:87434#2#F | \| \| |
| MQ=60 isize=160 C=37 | CGGGTGTTGGGCATTGGAGTCntAG\|C\|CCCCCCACTCCAGTCCTTCCCCAAGGATGCAGCCTCAGCCTTCTCCACACCACGTTTCCCCACAGACAAGTTCCCCACATCCTGGACTGGTAGCTGCCCAGGCCAACCCCATGGAACAAGGGCAT |
| 1:2114:9532:97361#2#F | \| \| |
| MQ=60 isize=159 C=37 | CCCCTATGTCCTATTGGAGTCCTG\|C\|CCCCCCACTCCAGTCCTTCCCCAAGGATGCAGCCTCAGCCTTCTCCACACCACGTTTCCCCACAGACAAGTTCCCCACATCCTGGACTGGTAGCTGCCCAGGCCAACCCCATGGAACAAGGGCATT |
| 2:2203:10789:20795#2#F | \| \| |
| MQ=60 isize=159 C=22 | GTCACCGGGGGCATTGGAGTCntg\|C\|CCCCCCaCTCCAGTCCTTCCCCAAGGATGCAGCCTCAGCCTTCTCCACACCacgtTTCCCCACAgAcAAgTTCCCCAcATCCTGGACTGGTAGCcgCCCAGgCCAACCCCatggaaCAAGGgcatt |
| 1:2110:4609:63254#2#R | \| \| |
| MQ=18 isize=-210 C=2 | tccccg\|c\|ccccccaccccagcccttccccaaggatgcggcctcgcccttctcccttccccgtttccccccagacaagttccccaacccctggaatggtagctgcctaggcgaacccccgggactaagtaggactccaatttcgcccccctc |
| 2:2107:2282:73369#2#R | \| \| |
| MQ=60 isize=-205 C=2 | cctccccccgcgcccaggccccg\|c\|cccccccccccagtccctccccaagggtgccgccccagccctccccacaccccgtttcCCCaCAGACAAGTTCCCCaCAtcCTGGACTGGTAGCTGCCCAGGCCAGGACTCCAATAATGATGAAAAA |
| 2:1206:7621:35403#2#F | \| \| |
| MQ=60 isize=158 C=27 | TGTCCGGCACGCATTGGAGTCnt\|C\|CCCCCCACTCCAGTCCTTCCCCAAGGATGCAGCCTCAGCCTTCTCCACACCACGTTTCCCCACAGACAAGTTCCCCACATCCTGGACTGgtAGCTGCCCAGGCCAACCCCAtGGAACAAGGGCatTT |
| 1:2112:20171:16565#2#R | \| \| |
| MQ=60 isize=-199 C=2 | \|c\|ccccccactccaggcctaccccaacgatgcagcctcagccttctccacacctcgagtccccacagacaagttccccacagccgggactggtagctgcccttgttccATGGGgttGGCCTGGgCAGcTAGGACTCCAATCaaCcgcacca |
